# A revised genome annotation of the model cyanobacterium *Synechocystis* based on start and stop codon-enriched ribosome profiling and proteogenomics

**DOI:** 10.64898/2025.12.20.695498

**Authors:** Lydia Hadjeras, Vanessa Krauspe, Rick Gelhausen, Benjamin Heiniger, Philipp Spät, Viktoria Reimann, Garance Jaques, Paul Minges, Raphael Bilger, Maren Gerstner, Boris Maček, Christian H. Ahrens, Rolf Backofen, Cynthia M. Sharma, Wolfgang R. Hess

## Abstract

Cyanobacteria are important primary producers and are used as microbial cell factories due to their ability to use solar light for oxygenic photosynthesis. *Synechocystis* sp. PCC 6803 is a popular model cyanobacterium, yet there are ambiguities in the precise coding regions of many genes, and numerous genes encoding small proteins have remained undetected. Here we present the results of a riboproteogenomic analysis, combining ribosome profiling (Ribo-seq) analysis involving inhibitors that stall ribosomes at translation initiation and termination sites (TIS– and TTS-Ribo-seq), with a proteogenomic reevaluation and reannotation of its entire genome. We report evidence for the translation of 3,055 annotated genes based on proteogenomics (83%), of 3,492 based on Ribo-seq (95.2%), and of 3,018 supported by both methods (82%). The data suggested unannotated protein-coding genes and corrections for annotated ones. We validated 15 small proteins translated from antisense RNAs, from intergenic and intragenic regions and provide proteogenomic support for up to 69 further, mostly small proteins. With *slr0489, slr1079* and *slr1082* we identified three genes with intragenic out-of-frame translons and show that both the internal and host reading frames are translated and that the resulting proteins interact with each other. Our data can be accessed via an intuitive and interactive genome browser platform at https://www.bioinf.uni-freiburg.de/∼ribobase/. They illustrate the enormous value of consolidating genome annotations in the context of integrated experimental data.

## INTRODUCTION

Oxygenic photosynthesis powers photoautotrophic organisms such as cyanobacteria enabling them to build complex organic molecules from inorganic carbon. As primary producers, cyanobacteria have a pivotal ecological significance^1^ and considerable value in biotechnological applications^2–4^. Among cyanobacteria, *Synechocystis* sp. PCC 6803 (from here: *Synechocystis* 6803), the third organism to have its complete genome sequenced^5^, is considered one of the most advanced models. However, despite careful annotation attempts, severe annotation ambiguities still exist. Cyanobacteria, like other prokaryotes, harbor a higher share of genes encoding small proteins compared to eukaryotes due to their more compact genome architecture^6^. Importantly, small protein-coding genes are frequently missed during standard genome annotation which aims to minimize the over-prediction of spurious short open reading frames (ORFs). Moreover, standard mass spectrometry-based approaches fail to comprehensively detect this class of proteins^7^, making small proteins a systematically underestimated component of the total proteome. Small proteins (here defined as ≤70 amino acids (aa)) are often considered too small to exert enzymatic activity. Nevertheless, many small proteins were demonstrated to interact with larger proteins or multi-protein complexes and have important functions^8^. Examples are the One-Helix Proteins in the assembly or stabilization of photosynthetic pigment-protein complexes in plant chloroplasts and cyanobacteria^9,10^, small structural proteins in photosystem (PS) I^11^, from which PsaJ (40 aa) is essential for the stable assembly and structural integrity of the entire complex^12^. Due to the extensive previous work on the photosynthetic apparatus, cyanobacteria provide a paradigm for small protein functions: Nineteen well-characterized proteins smaller than 50 aa are associated with the photosynthetic machinery alone^13^. Globally, small proteins operate as modulators of larger proteins’ enzymatic activities^8,14–16^, such as the 48 aa protein AtpΘ, which inhibits the futile reverse reaction of ATP synthase under unfavorable conditions^17^. Lastly, many small proteins are toxic, including those that are part of type I toxin–antitoxin systems^18^, or antimicrobial peptides (AMPs)^19^.

Ribosome profiling (Ribo-seq) detects translated mRNA segments on a genome-wide level^20^. Following cellular lysis and nuclease treatment, intact RNA associated with proteins – including the mRNA engaged with ribosomes during translation, is separated from cell debris by sucrose gradient ultracentrifugation followed by recovery of the associated RNA, cDNA synthesis and sequencing^21^. The analysis of RNA associated with the bacterial 70S ribosomes yields ribosome footprints with a length of 15–40 nt, peaking around 24 nt for *E. coli*^22^. Through comparison with an RNA-se^q^ library generated in parallel from total RNA harvested from the same culture and fragmented into similar size, the coverage by sequenced ribosome footprints then allows to infer ribosome occupancy, ORF boundaries, and translational efficiency. While Ribo-seq was initially developed to study translational regulation^23–25^, the discovery of extensive, previously unrecognized, coding potential in genomes of diverse organisms, such as yeast^26^, vertebrates including humans^27,28^, plants^29^, bacteria^30–32^, and viruses^33^ was another major benefit. The identification of potential short ORF encoded proteins (SEPs) catalogued by Ribo-seq in diverse organisms is gaining momentum. After the adaptation of the Ribo-seq workflow, this has only recently been extended to a range of bacteria and archaea^32,34–36^.

The antibiotic retapamulin, an inhibitor of the initiation of translation in bacteria, has revolutionized the view on the small proteome through Ribo-seq. Retapamulin (a pleuromutilin antibiotic) stalls ribosomes in treated cells at the translation initiation site (TIS), while elongating ribosomes finish the translation and run off the mRNA releasing the nascent peptide. Thus, retapamulin-assisted TIS-Ribo-seq results in defined peaks for translated proteins at their respective translational start sites, which allows to detect both unknown internal proteins, alternative versions of known proteins and unannotatedsmall proteins^30,37^. Further improvements have been obtained by the introduction of apidaecin (Api137), a proline-rich antimicrobial peptide, which arrests translation close to the stop codon^38^ and leads to the accumulation of ribosome footprints close to the 3′ ends of coding regions^39^, and thus can be used for globa^l^ mapping of stop codons via translation termination site (TTS)-Ribo-seq^32,40^.

Combining TIS– and TTS-Ribo-seq data can thus yield the precise borders of a translated sequence from start to stop^32,40^. These recent methodological breakthroughs applied alone have already yielded unprecedented insight not only into the small proteome of bacteria, but also suggested an unexpectedly high number of proteins translated from regions within annotated larger genes and even in antisense orientation^15,30–32,35,36,40,41^. However, thus far these approaches have only been applied to a small number of model species, and even more rarely as powerful combination of TIS-Ribo-seq and TTS-Ribo-seq. Given the large diversity among prokaryotes, substantial insight can be expected from describing the small and alternative proteome of species from diverse taxonomic ranks.

Previously collected Ribo-seq data for wild type and carbon-starved cells of *Synechocystis* 6803 showed allocation of ribosomes in carbon depletion^42^ and helpe^d^ to characterize the protein turnover at different time points of the light-dark cycle^43^. To our knowledge, there are no TIS– or TTS-Ribo-seq datasets at all and no Ribo-seq datasets for any other cyanobacterium available.

Altogether, 165 genes encoding small proteins ≤70 aa are annotated (Kazusa (KZS) annotation) in the *Synechocystis* 6803 genome (4.5% of all genes). We have recently characterized five additional small proteins NblD, AtpΘ, AcnSP, SliP4 and NirP1 that are functionally relevant in the energy and nutrient metabolism of *Synechocystis* 6803^9,17,44–46^ demonstrating the immense value of fully unraveling the composition and functional relevance of the small and alternative proteome.

Here, we report the results of a riboproteogenomic analysis combining TIS-, TTS-Ribo-seq and proteogenomics in the model cyanobacterium *Synechocystis* 6803. We present evidence for the translation of 3,492 and 3,055 annotated genes based on Ribo-seq and proteogenomics, and of previously unknown or unvalidated translons. We use translon as a unifying term for all categories of ORFs^47^. We detected an instance of ribosome frameshifting and small proteins translated from intragenic out-of-frame (IOF) alternative reading frames and from antisense RNAs (asRNAs). We integrated our Ribo-seq data with the annotations from several reference databases, performed a reanalysis of previously generated proteomic datasets^48^, and *ab initio* gene predictions. This integrated view of several annotations overlaid with experimental evidence allowed to consolidate many of the genome annotation differences. The proteogenomic approach, based on a search database that captures these differences and novel predictions^49^ also validated several Ribo-Seq candidates, frequently with an N-terminal peptide, and identified up to 69 additional, mostly small proteins. The consolidated data provide an updated and extended annotation of the *Synechocystis* 6803 genome that is available through an interactive web-based genome-browser at https://www.bioinf.uni-freiburg.de/∼ribobase/. The here developed generic approach provides an optimal basis to better exploit the genome sequence information; we expect it to be highly beneficial for many other model organisms and research communities.

## RESULTS

### Ribo-seq and retapamulin-based TIS-Ribo-seq captures the translatome of *Synechocystis* 6803

To globally measure translation of annotated ORFs and to discover potential new short ORF encoded proteins (SEPs), we applied Ribo-seq detecting actively translated translons. In parallel to the standard Ribo-seq approach, we added retapamulin to stall ribosomes on the start codon positions (set-up in **Fig. 1A**) for TIS-Ribo-seq. Cells were rapidly harvested to halt ribosomes along ORFs and preserve polysomes, and lysed, followed by converting polysomes into monosomes and trimming of ribosome footprints by MNase. Polysome profiles generated by sucrose density gradient centrifugation showed successful capture of translating ribosomes as well as their controlled digestion into monosomes (**Fig. 1A**). Following the results of our pre-experiments (**Fig. S1**), we used 10 μg/mL retapamulin in 100 mL *Synechocystis* 6803 cultures for TIS-Ribo-seq. Retapamulin treatment collapsed polysomes to monosomes even without micrococcal nuclease (MNase) digestion (**Fig. 1A**), as reported for *E. coli*^30^, validating our conditions for TIS profiling. Metagene plots of classical Ribo-seq data separately from TIS-Ribo-seq data document the overall quality of the ribosome profiling datasets and visualize the periodicity or phasing of elongating ribosomes (**Fig. S2**). The 5’ and 3’ ends of ribosomal footprints were mapped 12 nt up– and 19 nt downstream the first nt of the start codons, respectively. These signals were clearly enhanced if footprints were filtered for 32 nt length and more prominent in the presence of retapamulin. A 3’ nt periodicity of elongating ribosomes was clearly visible in the 5’ and 3’ mapping visualization without retapamulin.

**Fig. 1.**
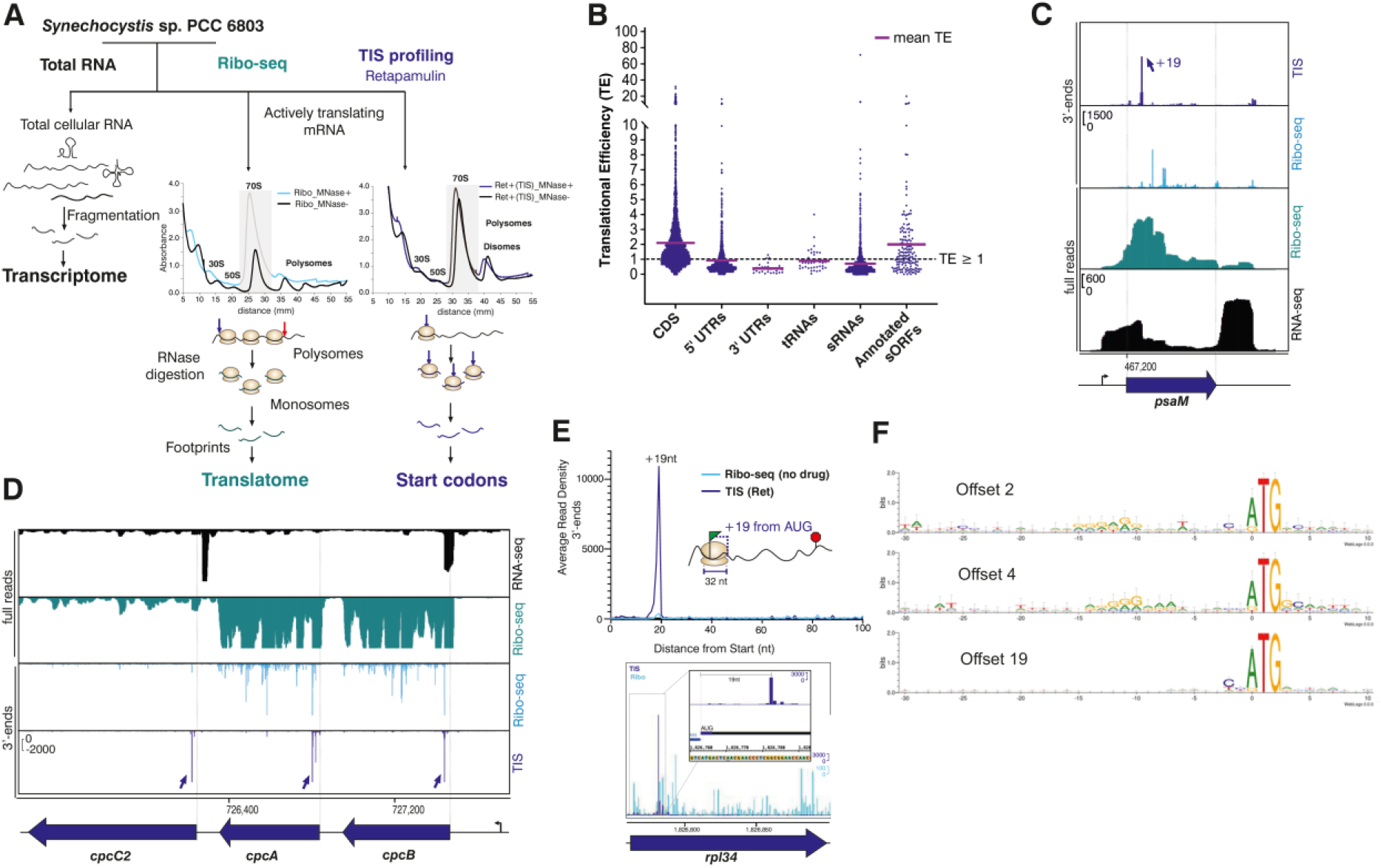
| Establishment of TIS-Ribo-seq for the cyanobacterium *Synechocystis* 6803. **A.** Experimental set-up. Ribosomes were captured in the presence or absence of retapamulin on the translated mRNAs. In parallel, total RNA was prepared from aliquots of the same cultures. After sucrose gradient ultracentrifugation, absorption profiles were taken at 260 nm comparing samples from cultures treated with retapamulin (10 μg/mL) for 15 min and digested with MNase (Ret+(TIS)_MNase+; dark blue) or not (Ret+(TIS)_MNase-; black). The middle panel shows samples without retapamulin treatment, either after MNase digestion (Ribo_MNase+, light blue), or without (Ribo_MNase-, black). The MNase treatment depleted polysomes between 40 and 60 mm, enriching the 70S monosome peak at 30 mm. Retapamulin treatment stalled ribosomes at translation initiation sites while elongating ribosomes finished translation, resulting in an enrichment of 70S monosomes. Approximately 30 nt long footprints protected by and co-purified with 70S ribosomes were then subjected to cDNA library preparation and deep sequencing to identify the translatome and start codons, respectively. **B.** Translational efficiency (TE) for different RNA categories. The scatter plots show global TEs (TE = Ribo-seq/RNA-seq) computed from Ribo-seq replicates for all annotated coding sequences (CDS), 5′– and 3′-UTRs, tRNAs, annotated small RNAs (sRNAs), and annotated sORFs encoding proteins of ≤70 aa. The purple lines indicate the mean TE of the respective transcript category. **C.** Upper 2 panels, read coverage from mapped 3′ ends from TIS-Ribo-seq (dark blue) and the Ribo-seq control (light blue), and from Ribo-seq and RNA-seq libraries (lower 2 panels) for the gene *psaM* encoding the photosystem I subunit XII. A sharp 3′ end was observed 19 nt downstream the annotated start codon. The Ribo-seq read coverage is enriched for the coding part, in contrast to the RNA-seq coverage, which starts at the position of the transcription start site (bent arrow). **D.** Read coverage for the *cpcBAC2* genes encoding the phycobilisome subunits CpcB, A and the rod linker polypeptide C2. **E.** Meta-analysis of footprints enriched in TIS-Ribo-seq. A sharp 3′ end was observed in 12,979 reads 19 nt downstream the annotated start codons of 582 genes (upper panel), exemplified in the lower panel for gene *rpl34* encoding the large ribosomal subunit protein L34. The 3′ ends of cDNA reads are depicted for ribosome-associated fractions after retapamulin treatment (dark blue) or without the drug (light blue lines). **F.** Enrichment of nucleotides associated with TIS-Ribo-seq 3′ end signals with an offset of 2 nt or 4 nt (51 genes), or 19 nt (407 genes) from the annotated start codons after filtering for ≥10 reads. Sequence logos of the 30 nt preceding and ten nt following the start codons are shown. The preference for ATG over GTG codons is visible, as well as a slight preference for preceding pyrimidine residues. An enrichment for upstream purine residues resembling a possible ribosome binding site was only seen for the TIS-Ribo-seq signals with an offset of 2 or 4 nt. For further details, see **Table S1**.

The mapped reads from the standard Ribo-seq analysis revealed the transcriptome-wide ribosome occupancy. Comparison to the RNA-seq coverage inferred in parallel allowed us to calculate the translation efficiency (TE) for a particular transcript (ratio Ribo-seq/RNA-seq coverage). The comparison between different annotated gene classes showed a much higher mean TE for protein-coding sequences than for sRNAs or 5′ and 3′ UTR sequences (**Fig. 1B**).

Typical examples for the obtained read profiles are shown for gene *psaM* encoding the small PSI subunit XII protein (**Fig. 1C**) and the tricistronic *cpcBAC2* genes encoding the phycobilisome subunits CpcB, CpcA and the rod linker polypeptide CpcC2 (**Fig. 1D**). The profiles show transcriptome coverage starting at the transcriptional initiation sites (TSS) and ending within the 3′UTR downstream the coding region. In contrast, the Ribo-seq read coverage is confined to the coding sections. As reported for MNase-generated Ribo-seq libraries in other bacteria^22,32^, the mRNA fragments protected by ribosomes showed a much cleaner, more defined cutoff at the 3′ ends, where the ribosomes’ footprints ended compared to the 5′ ends, where the footprints started (**Fig. S2**). As examples, we show the sharp 3′ ends observed 19 nt downstream the respective annotated start codons of *psaM* (**Fig. 1C**), the *cpcB, cpcA* and *cpcC2* genes (**Fig. 1D**), and *rpl34*, a house-keeping gene encoding the large ribosomal subunit protein L34 (**Fig. 1E**). For control, the 3′ ends of cDNA fragments generated from standard Ribo-Seq (in the absence of retapamulin), are plotted for all three examples (**Fig. 1C, D, E**). Meta-analysis of footprints enriched in TIS-Ribo-seq near annotated start codons revealed that the vast majority of their 3′ ends were located 19 nt downstream the annotated start codons (**Fig. 1E**). Disome footprints were excluded from TIS peak detection due to the size selection via PAA gels and the bioinformatic filtering for reads of length 32 nt. A parallel metagene analysis focusing on short ORFs (sORFs; ≤50 nt) was conducted to confirm that translation initiation patterns in these shorter sequences did not differ substantially from those observed in longer ORFs (**Fig. S3**).

Minor peaks for the footprint 3′ ends were observed 2 nt and 4 nt downstream of the annotated start codons (see **Table S1** for a list of these genes). The nature and conservation of possible ribosome binding sites for the initiation of translation in cyanobacteria is not known. Therefore, we searched for potentially conserved nucleotides 30 nt upstream and 10 nt downstream of the start codons, focusing on those genes contributing to the read density at the specific offsets (+2,+4,+19) and filtered them using a 10 read cutoff. This revealed a clear preference for ATG over GTG as start codon, and a slight preference for 2 preceding pyrimidine residues. A purine-rich motif that would indicate a likely ribosome binding site was found in only a subset of genes (**Fig. 1F)**. Most genes in this subset are highly expressed and often the first gene in an operon (**Table S1**), pointing at the relevance of these potential ribosome binding sites to achieve efficient translation.

Overall, metagene analysis and single-gene inspection suggested successful set-up of TIS-Ribo-seq profiling in *Synechocystis* 6803.

### Establishing TTS-Ribo-seq in a cyanobacterium

The comprehensive analysis and precise localization of stop codons is highly valuable for accurately calling previously overlooked translated reading frames. Translation termination site (TTS)-Ribo-seq is a method to infer stop codon positions based on the accumulation of ribosome footprints in the presence of inhibitors of termination^32,39,40^. This method has not been established yet for cyanobacteria. To facilitate TTS-Ribo-seq, we used the inhibitor Api137 that traps the release factors during translation termination, impeding the natural process of peptide liberation^38^. The experimental set-up is shown in **Fig. 2A**. In contrast to the polysome depletion in retapamulin-treated cultures (**Fig. 1A**), the gradient profile of Api137-treated cells did not change much in terms of polysome abundance (**Fig. 2A**). Remaining disome peaks were also observed quite frequently in samples digested with MNase, even without Api137. This observation is consistent with previous reports of ribosome queuing at termination sites^32,40^ indicating likely interference with MNase access, and was addressed here by inclusion of a dedicated disome library.

**Fig. 2.**
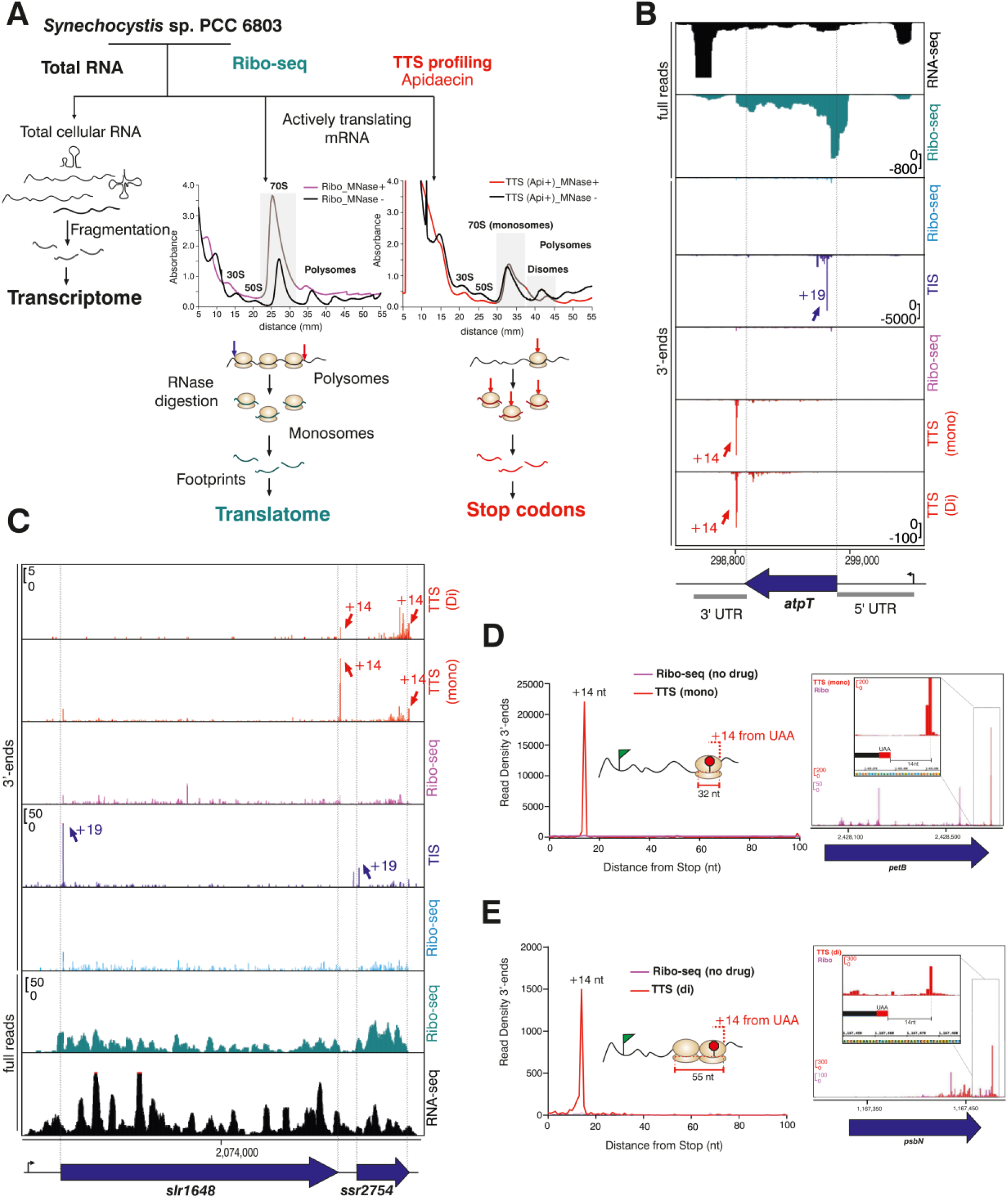
| Establishment of TTS-Ribo-seq for *Synechocystis* 6803. **A.** Experimental set-up. Gradient profiles comparing samples from cultures treated with 10 µM Api137 for 15 min (right panel) digested with MNase (TTS (Api+)_MNase +; red) or without (TTS (Api+)_MNase –; black) compared to samples without Api137 treatment (panel in the middle), after MNase digestion (Ribo_MNase +, pink), or without (Ribo_MNase –, black). After RNase treatment, approximately 30 nt long footprints protected by and co-purified with 70S ribosomes were obtained, subjected to cDNA library preparation and deep sequencing to identify the translatome and stop codons, respectively. In parallel, transcriptome samples were prepared and analyzed. **B.** TTS and TIS ribosome profiling for the gene *atpT* encoding the ATP synthase inhibitor AtpΘ^17^. Read coverage from RNA-seq and Ribo-seq libraries (upper 2 panels), mapped 3′ ends from Ribo-seq as control and TIS-Ribo-seq (middle panels) and from Ribo-seq control and TTS-Ribo-seq prepared from monosomes and disomes (lower 3 panels). The Ribo-seq read coverage is enriched for the coding part, in contrast to the RNA-seq coverage, which starts at the position of the transcription start site (bent arrow). The TIS-Ribo-seq 3′ ends map 19 nt downstream the start codon and the TTS-Riboseq 3′ ends 14 downstream the stop codons. **C.** TTS and TIS ribosome profiling for the genes *slr1648* and *ssr2754*. Ribosomes jam into blocked termination sites resulting in visible polysome fractions in Api137-treated samples prepared from disomes as well as monosomes (TTS-seq, upper 2 panels). The gene *slr1648* encodes the carotenoid oxygenase SynDiox2 involved in the biosynthesis of retinal^50^, *ssr2754* encodes a YefM-type antitoxin that forms an antitoxin-toxin gene pair with *ssr2755*^51^. **D.** Meta-analysis of footprints enriched in TTS-Ribo-seq from prepared monosomes (replicate 2, 32 nt reads, counts per million (CPM)-normalized). A sharp 3′ end was observed 14 nt downstream of the annotated stop codons across 1,055 genes (left panel), exemplified for gene *petB* (right panel). The 3′ ends of cDNA reads are depicted for ribosome-associated fractions after Api137 treatment (red) or without the drug (pink lines). **E.** Metagene analysis of footprints enriched in TTS-Ribo-seq from prepared disomes for 358 genes (replicate 2, 56 nt reads, CPM-normalized) (left panel), exemplified for the *psbN* gene (right panel).

The 3′ ends of sequenced footprints from treated samples compared to the Ribo-seq control were mapped for the TTS-Ribo-seq analysis. To document the overall quality, metagene plots of classical Ribo-seq data separately from TTS-Ribo-seq data are provided in **Fig. S4**. In both datasets, 5’ and 3’ ends of ribosomal footprints were mapped 14 nt up– and downstream the last nt of the stop codon. These signals were enhanced if footprints were filtered for 29 nt length and more prominent in the presence of Api137. A 3’ nt periodicity of elongating ribosomes was clearly visible in the 5’ mapping visualization without Api137.

As examples, we show the sharp 3′ ends observed 14 nt downstream of the respective stop codons of *norf1* (**Fig. 2B**), and the dicistronic *slr1648* and *ssr2754* genes encoding the carotenoid oxygenase SynDiox2^50^, and a YefM-type antitoxin^51^ (**Fig. 2C**). Note that the inclusion of TIS-Ribo-seq data allowed the precise definition of the *norf1* reading frame (**Fig. 2B**) that had not been annotated in any of the used reference genome annotations (KZS, GBK, RefSeq; see Methods). This gene has meanwhile been identified to encode the 48 aa ATP synthase inhibitor AtpΘ^17^.

Meta-analysis of footprints enriched in TTS-Ribo-seq from isolated monosomes revealed that the majority of their 3′ ends mapped 14 nt downstream of the respective stop codons (**Fig. 2D**). For illustration, we show the stop codon region of *petB* encoding the cytochrome b6 protein involved in photosynthetic electron transport (**Fig. 2D**). The analysis of cDNA 3′ ends enriched in TTS-Ribo-seq from prepared disomes revealed peaks mapping 14 nt downstream of the respective stop codons as well (**Fig. 2E**), exemplified for *psbN* encoding the 43 aa photosystem II protein PsbN.

To address the specificity of Api137 treatment, we performed metagene analyses around start codons. To compare our results to similar analyses in other bacteria, we also re-analyzed the respective datasets from *E. coli*^40^ and *Campylobacter*^32^ with identical settings. The results indicate that Api137 treatment leads to some enrichment of ribosome footprints also at initiation sites. This effect is not specific for *Synechocystis,* but can be generalized (**Fig. S5**).

To summarize, the 3′ ends of the footprints obtained from Api137-stalled ribosomes were observed on average 14 nt downstream of the respective stop codons. Metagene analysis and single-gene inspection suggested successful set-up of TTS-Ribo-seq profiling in *Synechocystis* 6803.

### ORFBounder identification of unannotated sORFs

Existing tools for sORF identification from ribosome profiling data have significant limitations for TIS-Ribo-seq and TTS-Ribo-seq analysis. While HRIBO^52^ incorporates DeepRibo for ORF detection, this tool is designed for conventional Ribo-seq data and not optimized for TIS or TTS datasets. At the time, the only available method for peak-based TIS detection from retapamulin-treated Ribo-seq data was RETscript^53^, which was developed for *Haloferax volcanii* and built on the methodological concepts of Meydan et al.^30^ However, RETscript was not meant for usage on other organisms and lacked support for TTS data. To address these points, we developed ORFBounder, a standalone, customizable tool specifically designed for calling TIS and TTS peaks from the respective Ribo-seq data types, with a particular focus on identifying small ORFs.

ORFBounder identifies translation sites based on ribosomal footprint enrichment following retapamulin treatment near potential start codons for TIS data. Optimal read lengths and P-site offsets for TIS based start codon prediction were determined by a metagene analysis (**Fig. 1E, Fig. S2**). All ORFBounder predictions that passed our filter criteria were corroborated by HRIBO and compared to an in-house curated genome annotation based on the KZS annotation that contained all previously mapped sRNAs, transcriptional units, 5′ and 3′ UTRs that were defined in comparative differential RNA-seq analyses over ten different growth conditions^54^.

### Validation by proteogenomic analysis and database visualization

An optimized proteogenomic workflow combining Ribo-seq and proteomic data can substantially support the identification of unannotated proteins or alternative protein forms^55^. To leverage the extensive proteomics data previously acquired for *Synechocystis* 6803^48^, we created integrated proteogenomics databases (iPtgxDBs) based on the chromosome and plasmid sequences and searched publicly available mass spectrometry datasets^48^ against them (**Fig. S6**). Besides the extended KZS annotation used for the Ribo-seq analysis, we integrated the annotations according to NCBI RefSeq 2022 (ID: GCF_000009725.1) (2022) and GenBank (2016) to distinguish differences in gene calling and annotation. Despite being based on the same genome sequence, these differed substantially (**Fig. 3B**). Furthermore, to capture the protein-coding potential comprehensively^49^, we also included two *ab initio* gene predictions (Prodigal 2.6.3^56^ and ChemGenome 2.1^57^), and an *in-silico* ORF prediction which corresponds to a six frame translation of the full genome sequence, considering all ORFs with a minimum length of 10 aa and starting with ATG or one of the three most common alternative start codons, namely TTG, GTG and CTG^58^. These annotations were hierarchically integrated with minimal redundancy into a standard iPtgxDB, ensuring that over 95% of the identifiable peptides unambiguously imply one protein sequence (**Fig. S6**). Due to the inclusion of *in-silico* ORFs, standard iPtgxDBs are large, leading to statistical limitations for protein identifications. Therefore, a much smaller custom iPtgxDB that replaces the *in-silico* ORFs with the top Ribo-seq candidates was created, as described previously^35^ (**Fig. 3**). For an overview of the peptide information content of different search databases, see **Fig. S7, Supplementary Dataset 4,** and **Methods**. These data have been integrated in our platform at https://www.bioinf.uni-freiburg.de/∼ribobase/.

**Fig. 3.**
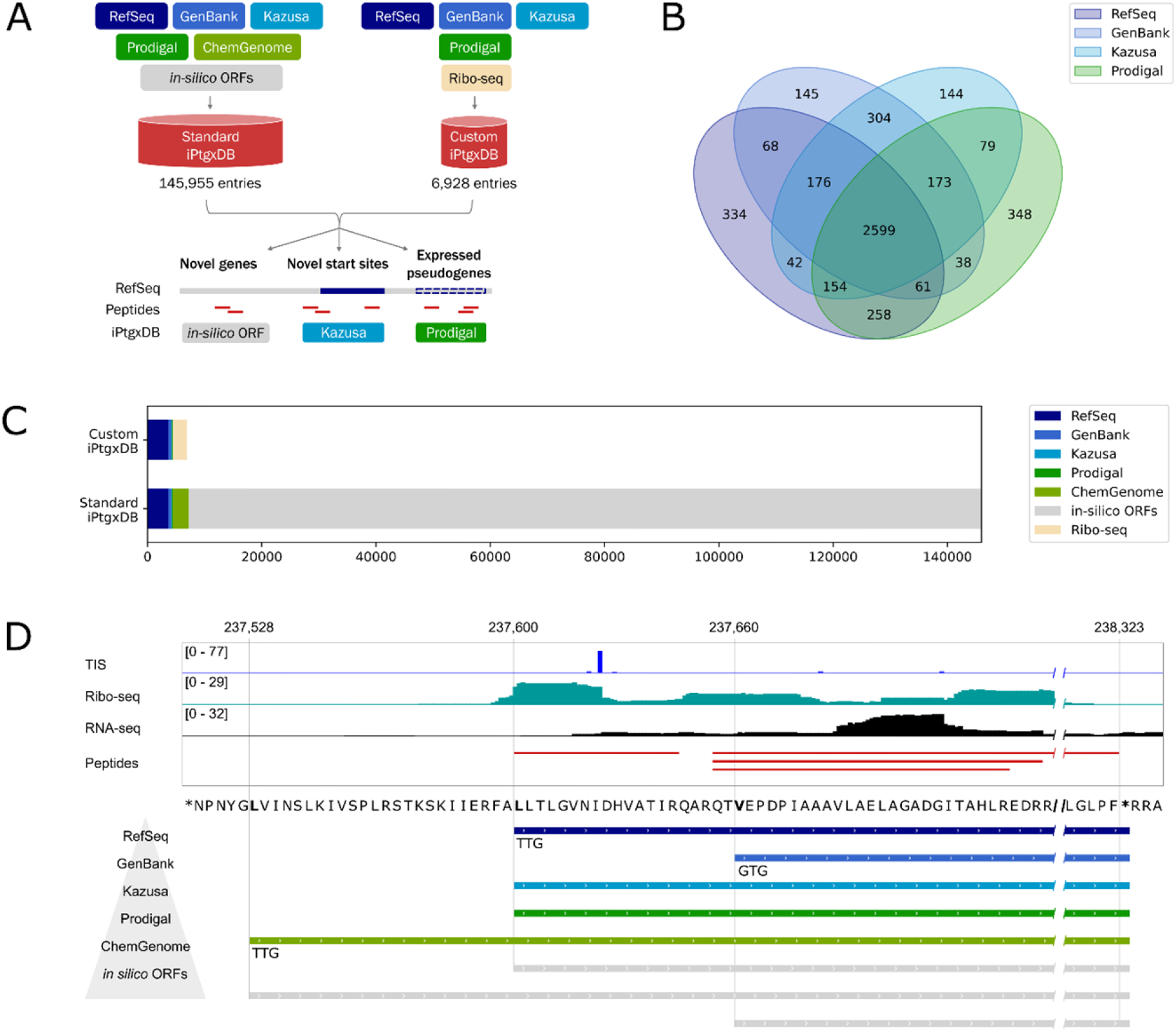
| Proteogenomics strategy and database construction. **A.** A standard integrated proteogenomics database (iPtgxDB) was created by integrating curated genome annotations (blue), *ab-initio* gene predictions (green) and a modified six-frame translation considering alternative start codons (*in-silico* ORFs, grey) in a minimally redundant way, resulting in a database containing close to 146,000 entries. A much smaller custom iPtgxDB was created by replacing the *in-silico* ORFs and ChemGenome predictions with the 2,708 top Ribo-seq candidates, resulting in a database containing only roughly 7,000 entries. Both of these iPtgxDBs could then be used in tandem with proteomics data to uncover novel genes, correct the start sites of already annotated genes and confirm the expression of pseudogenes. **B.** Overlap between different annotations of the *Synechocystis* 6803 genome. Shown are the precise matches in the number of CDS sharing identical start and stop codon assignments in the NCBI RefSeq 2022 (ID: GCF_000009725.1) (2022), the GenBank (2016) and the KZS annotations, plus an *ab initio* gene prediction using Prodigal 2.6.35. Only 2,599 CDS are precisely shared between the four datasets. **C.** Entries contributed to both iPtgxDBs by the included annotation sources. Entries are only considered if they are not predicted by a source higher in the annotation hierarchy (corresponding to the order in the legend). This shows that the *in-silico* ORFs are the main contributor to the large size of the standard iPtgxDB. **D.** Example of an annotation cluster (predictions from different annotations sharing the same stop coordinate). RefSeq, the highest entry in the annotation hierarchy (bottom left triangle), KZS and Prodigal predict the same start, while GenBank predicts a shorter version and ChemGenome a longer one. The respective start codons are shown below the entry. The TIS Ribo-seq data (blue) and Ribo-seq footprint (turquoise), as well as an n-terminal peptide detected with proteomics (red) confirm the RefSeq annotated start.

### Annotated proteins and SEPs confirmed by riboproteogenomics

Overall, 3,492 of the 3,669 KZS-annotated proteins were considered expressed according to the Ribo-seq results (95%, RIBO-WT-avg_TE ≥0.288) and 3,055 based on proteogenomics (83%), while 3,018 proteins were supported by both methods (82%) (**Table S2**). Of the 165 KZS-annotated SEPs ≤70aa (**Table S4**), 138 (83.6%) were considered expressed in our dataset based on the Ribo-seq data (RIBO-WT-avg_TE ≥0.288), and 97 had support from TIS-Ribo-seq. In addition, proteogenomics evidence was observed for 67 of these SEPs (39%) (**Table S4**), all of which had some form of Ribo-seq support. Previously, Baers et al. had validated 47 small proteins^59^. Our proteogenomic dataset missed 3 of those (*hliD, ssl5096* and *MYO_RS19000*), which, except MYO_RS19000, were covered by Ribo-seq data. Thus, based on our combined data, 138 annotated SEPs are translated, almost tripling the previously available experimental evidence for 47 annotated and validated SEPs. Among the remaining sORFs lacking evidence for translation, 12 are pseudogenes, mainly fragments of transposase genes.

### Previously unknown proteins identified by riboproteogenomics

An optimized proteogenomic workflow combining Ribo-seq and proteomic data can substantially support the identification of unannotated proteins or alternative protein forms^55^. The combination and filtering of all TIS-, TTS– and classical Ribo-seq data reduced the initial number of 146,175 predictions to 2,708 instances of suggested corrections of annotated genes, or potentially additional translons (1,069 unannotated, 1,191 Internal-Out of-Frame (IOF), 305 Internal-In-Frame (IIF), 116 longer and 27 shorter proteoforms) (**Table S2, S3**). The lengths of these additional translons were strongly biased towards smaller proteins (**Fig. S8**). After manual inspection of 808 randomly chosen instances from this list (30%), we considered 163 translons to be correct (20%) based on the Ribo-seq coverage. Among these 163 candidates, we found 90 unannotated translons, 51 IOF, 9 IIF, 4 N-terminal extensions and 9 possible N-terminal truncations. 124/163 (76%) are SEPs of 70 or less aa. A visualization of translon categories, length distributions, and validation rates per class is given in **Fig. S9**.

To identify previously not annotated proteins or those unique to one of the annotations with proteogenomics, we performed searches against the standard and custom iPtgxDB. A summary of the search results and the overlap for different classes is shown in **Table S5,** for detailed information see **Supplementary Dataset 1.** Overall, 69 proteins were identified that were not annotated by RefSeq, 11 of which were contained in the older KZS and/or GenBank annotations, i.e., examples of CDS that got wrongly removed from the genome annotation (**Fig. S10C**). The other 58 proteins identified through searches of the iPtgxDBs were entirely new (**Fig. S10**). The custom iPtgxDB uniquely uncovered 22 proteins with Ribo-seq evidence, while the standard iPtgxDB uniquely found 27 *in-silico* ORFs (not contained in the custom iPtgxDB), and 6 were predicted by both. Finally, two predictions from ChemGenome and one from Prodigal were only found using the much larger standard iPtgxDB, again illustrating the value of integrating several annotations^49^.

### Start codon usage by annotated and previously unknown genes

The power of combining the Ribo-seq and ORFBounder analysis with proteogenomic searches was well illustrated in the assignment of start codons. Integrated evidence from TIS and standard Ribo-seq data combined with N-terminal peptides identified by mass spectrometry–based proteomics provided direct confirmation of the annotated start site for 2,065 of 3,669 (56%) annotated genes (**Fig. S11**). The Ribo-seq data further suggested a corrected start site for 13 annotated proteins, while proteogenomics indicated 54 corrected starts.

As reported in other bacteria, start codon usage of the annotated genes strongly favors ATG (85%), followed by GTG (10%) and TTG (4%). While this was also found for the implied corrected start codons and riboproteogenomically detected proteins, especially the candidates detected by proteogenomics showed stronger preference for alternative start codons, with less than 50% starting with a canonical ATG and 23% starting with GTG and TTG, respectively. Interestingly, some of those candidates also start with CTG, confirming the validity of including this start codon when generating iPtgxDBs (**Fig. S12)**.

There was a single ATT start codon annotated (KZS), for gene *ureF/slr1899* encoding urease accessory protein F, while in the GBK and RefSeq annotations, *ureF* was annotated to start from a GTG. However, ORFBounder specified yet another, more distal TTG as start with a very good score, leading to a 243 aa protein, instead of the 217 aa (GBK and RefSeq) and 180 aa (KZS) forms. This N-terminally extended form was directly supported by a peptide MDDNLLPLLSGSAQLK identified in the standard iPtgxDB search with a spectral count of 5. It serves as direct proof for the N terminus, as the genomically encoded sequence would have been LDDNLLPLLSGSAQLK.

### TIS– and TTS-Ribo-seq permit improved translon identification and the detection of ribosome frameshifting

We observed that the annotation of reading frames and their respective start codons frequently differs between the considered annotation files, although these were based on identical genome sequences. This can cause substantial problems if proteins of interest are expressed only in a truncated version or contain unannotated N-terminal extensions. These problems can be resolved by considering the TIS– and TTS-Ribo-seq data and consolidating the different genome annotations on a case-by-case basis.

We illustrate this here for the gene *sll0553* encoding an alpha/beta fold hydrolase, that is widely conserved in cyanobacteria. The annotated longest gene model in GBK is 1,038 nt, encoding a 345 aa homolog with a GTG start codon (WP_010873707.1), while the corresponding gene model in the KZS annotation suggests a 306 aa protein. In contrast to both annotations, the Ribo-seq coverage and TIS-Ribo-seq mapping data indicated a reading frame encoding a 298 aa protein. DNA and protein sequence alignments show that the here reported start codon is conserved among different *Synechocystis* strains and matches the annotations of homologs in *Synechocystis* strains PCC 6714 (hereafter *Synechocystis* 6714) and 7339 (**Fig. 4A**). We further noticed that the protein length of 298 aa is supported by 19 different peptides from the proteogenomic analysis, while no peptides were found that support the longer N termini annotated in the GBK and KZS annotations. Hence, based on the combined evidence, the gene model for *sll0553* should be truncated by 47 codons.

**Fig. 4.**
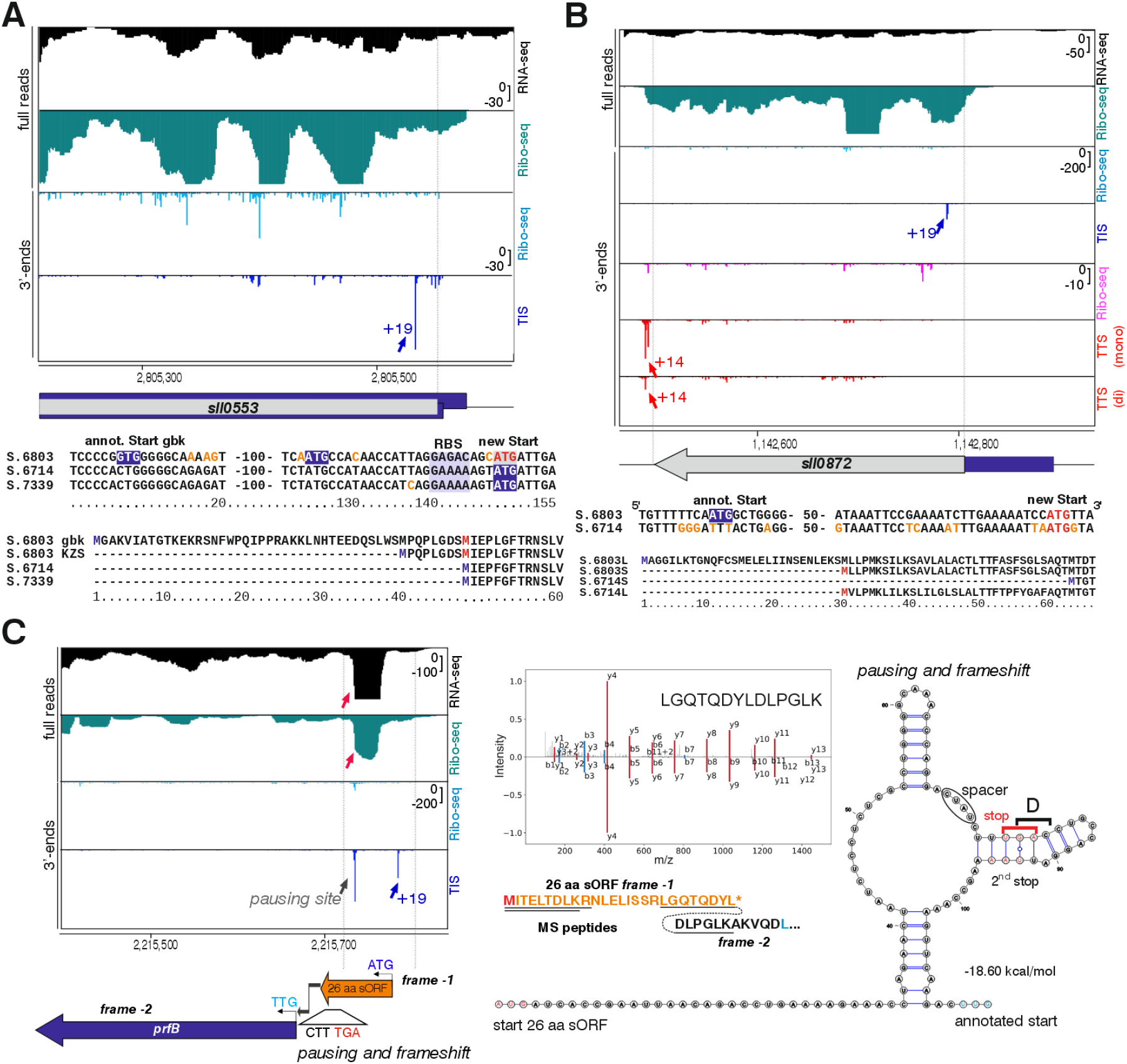
| Start codon re-annotation and detection of ribosomal frameshifting. **A.** Re-annotation of the start codon of *sll0553* based on TIS profiling data. The annotated longest gene model in GBK is 1038 nt, encoding a 345 aa homolog with a GTG start codon (WP_010873707.1), while the KZS annotation suggests a 306 aa protein. However, the Ribo-seq coverage and TIS-Ribo-seq mapping indicate a reading frame encoding a 298 aa protein. Blue: current annotations (345 and 306 aa). Grey: Novel annotation based on TIS profiling data (298 aa). The enriched TIS peak associated with the correct start codon is indicated with a black arrow in the TIS library and indicates an N-terminal truncation for the *sll0553*-encoded protein. The lower panel shows DNA and protein alignments of the translation initiation regions from different *Synechocystis* strains. Thus, the TIS data supports a shorter form of *sll0553* in *Synechocystis* 6803, consistent with the annotations in *Synechocystis* strains PCC 6714 and 7339. The annotated and new start codons are indicated. The corrected protein length of 298 aa is supported by 19 different peptides from the proteogenomics analysis, while no peptides were found for the elongated N termini based on the GBK and KZS annotations. **B.** Correction of Sll0872. It is annotated as a 393 nt ORF in *Synechocystis* 6803 (GBK and KZS annotations) yielding a 130 aa protein (S.6803L; BAL28778.1), while the reading frame of the homolog D082_22070 in *Synechocystis* 6714 with 69 codons is much shorter (S.6714S). The Ribo-seq data suggest that both variants are incorrect. The predicted new start is depicted in red in both strains. **C.** Ribosome frameshifting. The *prfB* gene (*sll1865*) encodes peptide chain release factor 2. RNA-seq coverage (top, black) indicates contiguous transcription from a TSS approximately 140 nt upstream the annotated reading frame beginning with a TTG start codon. Ribo-seq coverage in the 2^nd^ panel from top indicates translation in the upstream region, supported by a TIS-Ribo-seq signal 19 nt downstream of a suitable start codon (blue line +19 in the 4^th^ panel from top), pointing at a 26-codon reading frame finishing with a TGA stop codon. However, both datasets show an unusual, abundant read accumulation in the same subregion (red arrows) and another TIS-Ribo-seq signal, which is not associated with a start codon. This pattern is consistent with ribosomal pausing and +1 ribosomal frameshifting, which is strongly supported by mapped peptides (black horizontal lines under the aa sequence). Two (MITELTDLK and MITELTDLKR) map to the upstream 26 aa frame, including the start amino acid, while a third (LGQTQDYLDLPGLK) directly spans the frameshifting site (lines connected by a dashed line and mirror plot). The mirror plot shows the mass-to-charge ratio on the X-axis and the fragment ion intensities from the experimentally observed (upper part) and computationally predicted spectrum (lower part) on the Y-axis (b-ions retain the positive charge in the N-terminus, y-ions retain the positive charge in the C-terminal fragment). Elements involved in the frameshifting are annotated alongside the RNA secondary structure with a minimum free energy of –18.60 kcal/mol. The shift site is preceded by a short spacer sequence and followed by a second UAA stop site downstream, a conserved feature of these elements to suppress translation from re-initiation at the UUG. The RNA secondary structure was predicted by RNAfold^125^ with default parameters and visualized with VARNA version 3.93^126^. The technical data for the spectrum shown in the mirror plot are: charge, +2; collision energy, 27.0; precursor m/z, 780.91821289; mass difference, –0.00085449; percolator score, 1.00; hyper score, 42.78; spectral angle, 0.94; Pearson’s correlation, 1.00.

As a second example, we show the gene *sll0872,* annotated by both GBK and KZS to encode a 130 aa protein (BAL28778.1). However, the very closely related homolog D082_22070 in *Synechocystis* 6714 with its annotated 68 aa is much shorter. Here, our Ribo-seq data indicate that likely the annotations in both strains are incorrect. Ribo-seq coverage and a TIS signal 19 nt downstream of another suitable start codon suggest that Sll0782 is a 101 aa protein in *Synechocystis* 6803 (**Fig. 4B**), consistent with 9 detected peptides, while none was found matching the annotated 29 codons of the N-terminus before the here defined start. Sequence alignments show that a start codon at a matching position exists also in *Synechocystis* 6714, for which consequently the gene model for D082_22070 should be extended by 33 aa, yielding 101 aa proteins in both strains (**Fig. 4B**). Thus, the data suggest that the *sll0872* gene model should be truncated by 29 codons.

The predictive power of TIS– and TTS-Ribo-seq profiling is furthermore evident in the region around genes *slr1220* and *slr1222* encoding unknown proteins. TIS-Ribo-seq in combination with Ribo-seq coverage supported an N-terminal extension of Slr1222 by 116 aa and initiation from a rare TTG codon, yielding a 321 aa protein **(Fig. S13A**).

In many different bacteria, translation of *prfB* encoding release factor 2 (RF2 or PrfB) includes ribosomal frameshifting^60,61^. This *prfB* ribosomal frameshifting has not yet been demonstrated in cyanobacteria. However, the identification of two peptides by mass spectrometry provided proteogenomic support for the translation of an 26 aa SEP located upstream of *prfB*^48^, while the annotated *prfB* gene (*sll1865*) appears truncated and starts from a rare TTG codon (**Fig. 4C**). Here, we noticed abundant signals in both the Ribo-seq and RNA-seq coverage exactly upstream of the likely pausing site (**Fig. 4C**). The encoded release factor 2 mediates termination at UAA and UGA codons. Almost all described frameshifting instances in other bacteria involve CUU UGA as the shift site/stop codon. A cytosine base 3’ of the UGA stop codon was shown in *E. coli* to make UGA a relatively weak termination codon, contributing to the frameshifting^62^. These bases – CUU UGA C – are also found in the C-terminal region of the 26 aa sORF here. In the presence of high PrfB concentrations, most ribosomes terminate at the UGA stop codon and only the 26 aa peptide is formed. If the concentration of PrfB is low, the CUU codon can detach from the anticodon of peptidyl-tRNA_Leu_, mediating a shift from the +1 to the +2 frame^61^. Another described element involved in the frameshifting is a short spacer upstream of the shift site (often UAU^61^, here CUAU). All of these elements are also present here **(Fig. 4C**). However, direct proof for the ribosomal frameshifting comes from an identified peptide (LGQTQDYLDLPGLK) that directly spans the frameshifting site **(Fig. 4C**). This data corroborates the presence of this naturally occurring frameshift in cyanobacteria and provides a view of how such a site appears in ribosome profiling data. The abundant Ribo-seq coverage and TIS signals at the shift site provide direct evidence for ribosome stalling upstream of the termination codon and re-initiation at the internal SD element as evidenced from the TIS signal.

We conclude that the riboproteogenomics data are useful to spot ambiguities concerning functional start codons and support gene models that should be truncated or N-terminally extended. They furthermore illustrate an instance of ribosomal frameshifting in cyanobacteria.

### Ribosome profiling aids the discovery of toxin-antitoxin candidates

Our ribosome profiling data allowed us to identify additional protein-coding genes. Names were assigned conferring to their location in non-coding areas according to the location of the forward or reverse strand (*ncr* or *ncl*), their transcriptional unit (TU), or their location within an annotated gene. Two previously identified transcripts, Ncl1450 and Ncr1470, were thought to be two non-coding sRNAs complementary to each other^54^. Their origin from the same locus in forward and reverse orientation, their upregulation under low carbon^54^, as well as the arrangement of *ncl1450* resembling an excludon^63^ with regard to adjacent genes, prompted further investigation. Intriguingly, Ribo-seq coverage, TIS, and TTS signals indicated the translation of both Ncl1450 and Ncr1470 (**Fig. 5A**). Fusions of the reading frames with the P*_petE_* promoter led to their expression detected in western blots (**Fig. 5B**). Ncl1450 homologs are found in several other cyanobacteria (**Fig. 5C**), including *Pleurocapsa*, *Stanieria* (both 53% aa identity) and *Cyanothece* (50% aa identity), while *ncr1470* is only found in *Synechocystis* strains (**Fig. 5E**). Ncl1450 is a 36 aa protein that was predicted as containing a single transmembrane domain by DeepTMHMM^64^. Corresponding fractionation experiments into soluble and membrane fractions and comparison with the membrane protein PetC supported a membrane localization for Ncl1450 (**Fig. 5D**). The combination of a gene encoding a single transmembrane protein with an asRNA resembles the architecture of type I toxin-antitoxin systems^65^. We conclude that Ncl1450 and Ncr1470, previously considered as two non-coding sRNAs complementary to each other, encode SEPs of 36 and 40 aa and appear as a derivative of a type I toxin-antitoxin-like system.

**Fig. 5.**
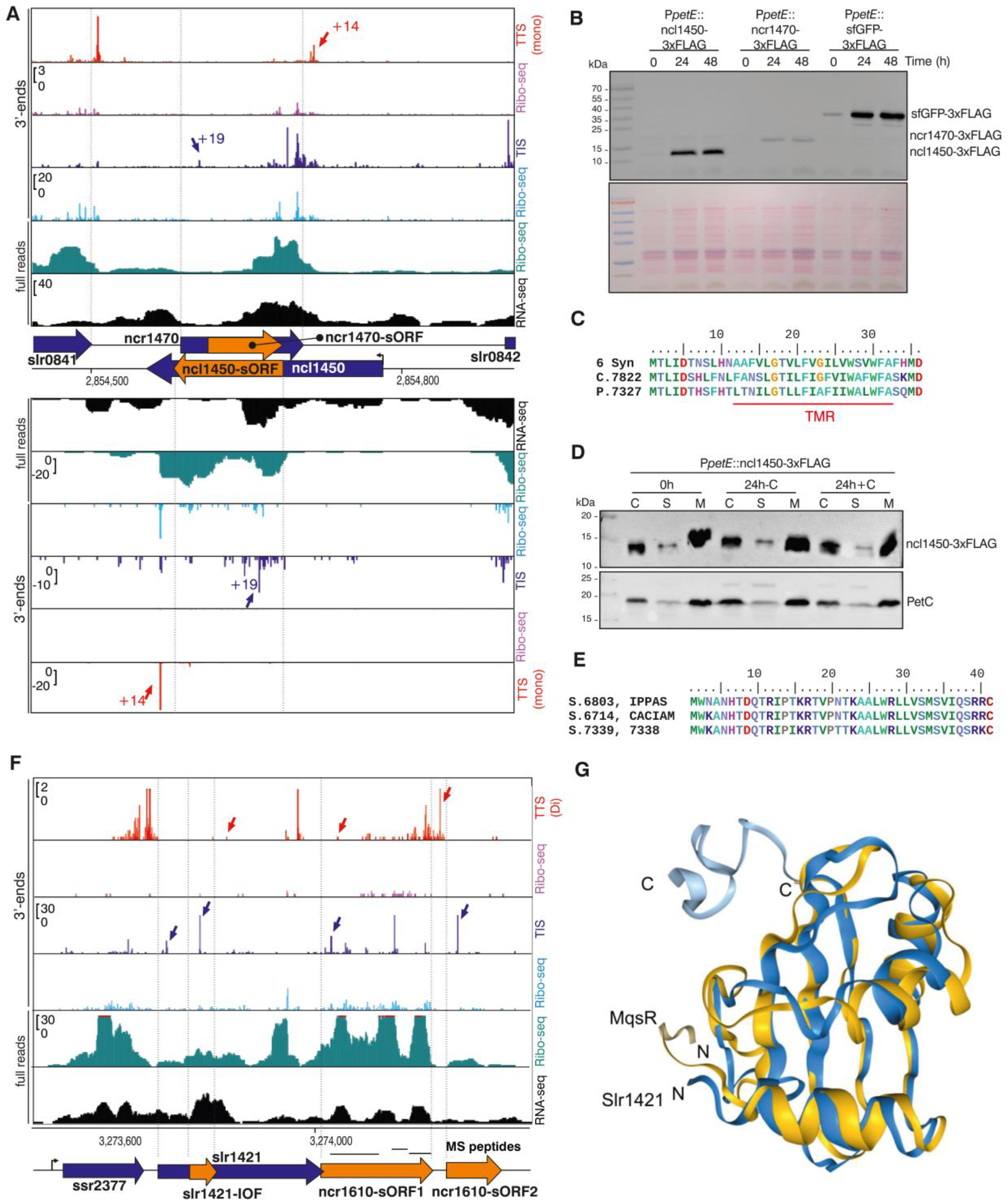
| Identification of sORFs encoding putative toxin-antitoxin candidate systems. **A.** Ncr1470 and Ncl1450 were annotated as a pair of overlapping sRNAs^54^, the genomic arrangement is shown in the center of panel A. Ribo-seq, and the comparison of TIS and TTS tracks with the respective “Ribo-seq” negative control panels supports the protein-coding potential of Ncr1470 and Ncl1450 (orange arrows). **B.** Both coding sequences were fused to the copper-regulated P*_petE_* promoter and a segment encoding a C-terminal 3xFLAG tag. A sequence encoding superfolder GFP (sfGFP) was engineered as control. The time course of protein induction after induction is shown. The lower panel shows the corresponding ponceau red-stained membrane. **C.** Alignment of 6 identical Ncl1450 homologs in *Synechocystis* strains 6803, 6714, 7338, 7339, IPPAS B-1465 and CACIAM 05 (6 Syn), with homologs from *Cyanothece* sp. PCC 7822 and *Pleurocapsa* sp. PCC 7327. A predicted transmembrane domain is indicated (red line, TMR). **D.** Fractionation of total cellular protein (C) into soluble (S) and membrane (M) fractions yielded a similar localization for Ncl1450 in the membrane fraction as the known membrane protein PetC. Mouse monoclonal anti-FLAG serum coupled to horseradish peroxidase (A8592, Sigma) was used at a titer of 1:5000, anti-PetC antiserum (AS08 330, Agrisera) was used at a titer of 1:2,000 and developed with horseradish peroxidase-conjugated anti rabbit secondary antiserum (A8275, Sigma-Aldrich) at a titer of 1:10,000. **E.** Ncr1470 is only found in *Synechocystis* strains. The alignment shows all putative homologs that could be identified. Three sequence variants were found, with identical sequences each in *Synechocystis* 6803 and IPPAS B-1465 (top), in strains CACIAM 05 and PCC 6714 (center), and in strains PCC 7339 and PCC 7338 (bottom). None of the respective genes is annotated. **F.** Coding potential around the *slr1421* locus. TIS– and TTS-Ribo-seq data suggest two sORFs encoding small proteins of 71 and 34 aa within the previously defined sRNA *ncr1610*^54^. The coding potential of sORF1 was verified immunologically (**Fig. 6A**) and by the detection of 3 peptides (horizontal tabs). Its start codon overlaps the stop codon of *slr1421*. Bent arrows indicate transcription start sites in panels A and F. **G.** Alphafold-predicted structure of Slr1421 (blue) and overlay with the best-matching structure identified by Foldseek (yellow), the protein MqsR (TM score: 0.6003).

An even more compelling case is provided by *ncr1610* previously considered as an sRNA^54^. Both TIS– and TTS-Ribo-seq data suggested that *ncr1610* contains two sORFs encoding small proteins of 71 and 34 aa (**Fig. 5F**). The coding potential of sORF1 was verified by the detection of 4 peptides from the search against the iPtgxDB and immunologically by FLAG tagging and western blot detection (**Fig. 6A**). Homologs to *ncr1610-sORF1* exist throughout the cyanobacterial phylum, in Pseudomonadota, Chloroflexota and other bacteria (**Fig. S13B**), while no conservation was found for *ncr1610-sORF2.* The *ncr1610-sORF1* start codon overlaps the stop codon of the preceding gene *slr1421* indicating a tight genetic association between the two genes. Searches using Foldseek^66^ suggested that Slr1421, a thus far uncharacterized DUF4258 protein, is a structural homolog of MqsR (**Fig. 5G**), an mRNA interferase toxin^67^. Therefore, the Ncr1610-sORF1 protein is suggested as the corresponding antitoxin.

**Fig. 6.**
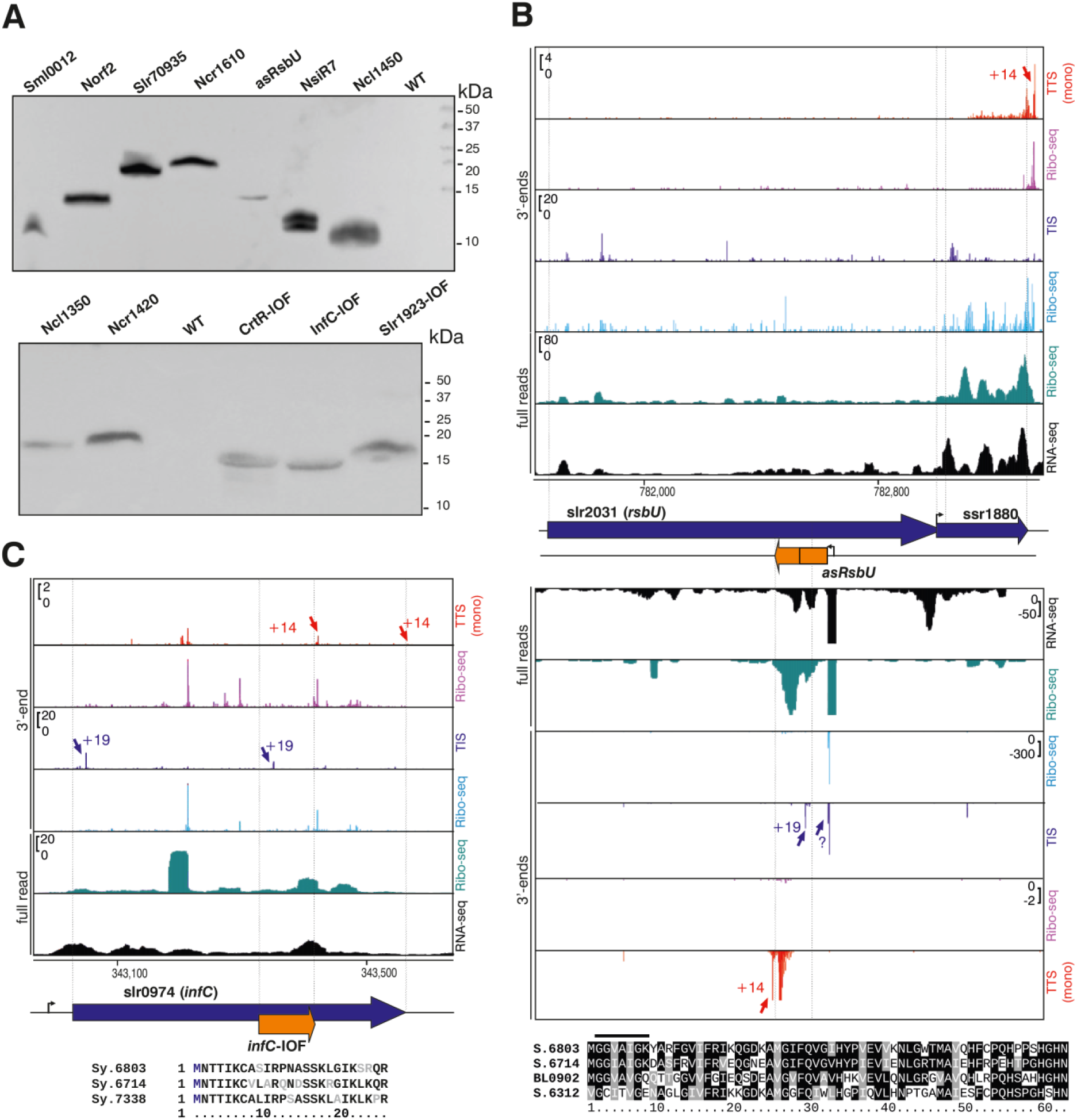
| Validation of sORF candidates. **A.** Western Blot verification using anti-FLAG serum showing signals for 7 3xFLAG-tagged sORFs (upper panel) and 5 dual SPA-tagged proteins (lower panel) expressed from plasmid pVZ322 under control of the P*_petE_* promoter induced by copper addition. *Synechocystis* wild type (WT) samples served as negative controls. **B.** Coding potential in an asRNA to *slr2031* (*rsbU*). Both DNA strands are expressed, supported by RNA-seq (black), Ribo-Seq (turquoise), TIS-Ribo-seq (dark blue) and TTS-Ribo-seq (red) datasets. The sequence alignment at the bottom shows possible homologs of the 62 aa asRsbU encoded in *Synechocystis* strains 6714 and 6312, and in *Leptolyngbya* sp. BL0902 identified by TBlastN in antisense orientation to their respective *rsbU* genes. However, it is unknown if corresponding asRNAs exist in these strains and the respective ORFs are longer than the shown sections. **C.** Evidence for an IOF encoding a 25 aa peptide within *slr0974* encoding the ribosome initiation factor IF-3. Bent arrows indicate transcription start sites.

Toxin-antitoxin proteins are often small, which makes the detection of the corresponding loci particularly difficult. Riboproteogenomics aids in the discovery of such systems, illustrated here by the antitoxin candidate Ncr1610-sORF1, and the *ncl1450/ncr1470* locus.

### Validation of SEPs

Based on the DeepRibo and ORFBounder predictions, we selected 15 sORFs for experimental validation in addition to the Ncl1450/Ncr1470 pair and Ncr1610-sORF1 discussed above. Six of these sORFs were intergenic, three were short IOFs, three were asRNA-located sORFs, and three were intragenic IIFs in the same reading frame as the main gene. Thus, these candidates represented all types of translons, with different levels of support. All sORFs were fused to C-terminal 3xFLAG tags and if these could not be detected, we repeated the experiment using an SPA tag. Using this strategy, 15 of the in total 18 candidates were validated by western blotting (**Fig. 6A**, **Table 1**).

**Table 1.**
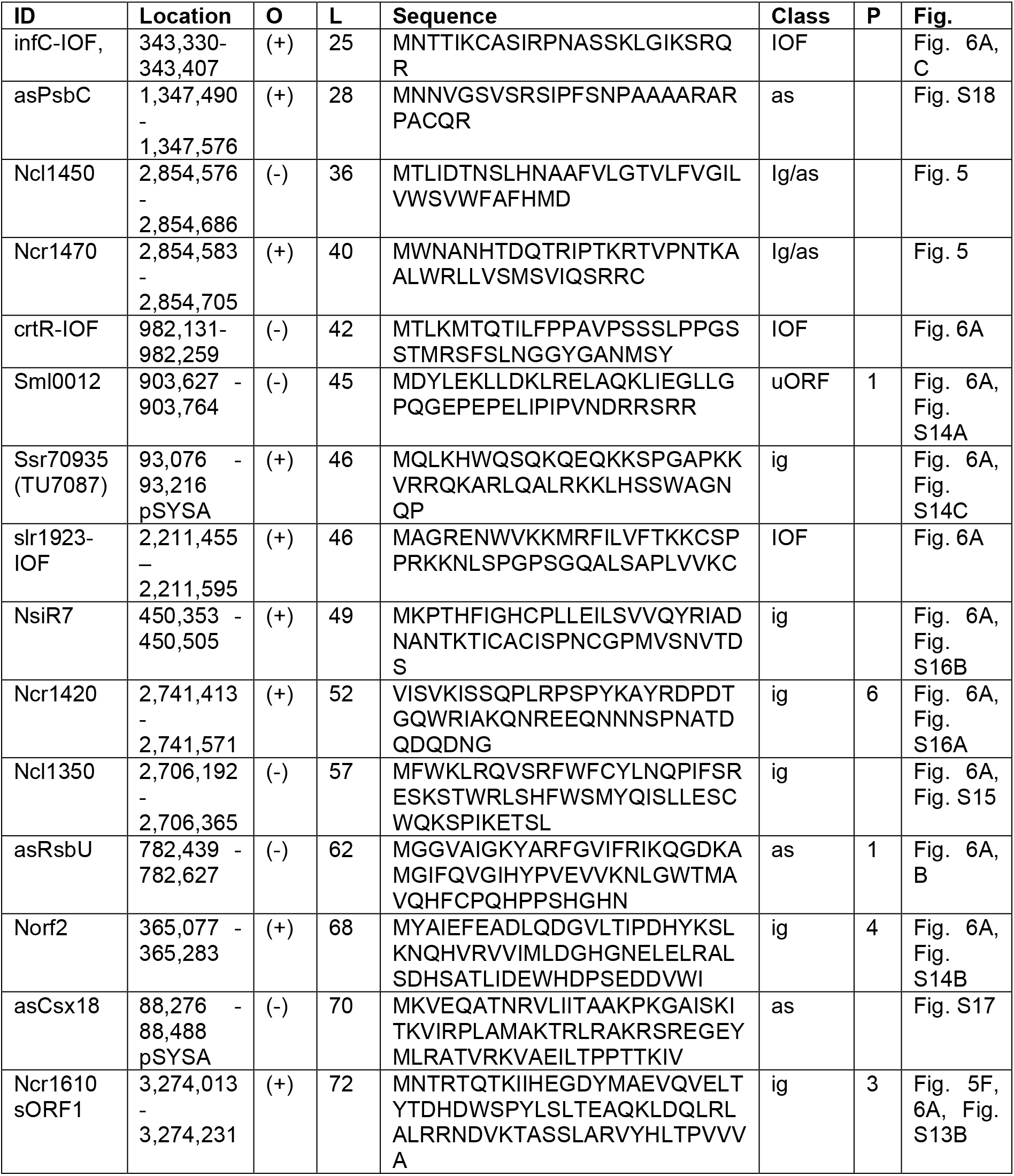
SEPs validated in this study by tagging and western blotting. The SEP ID is given (ID), followed by the coordinates and orientation (O). The SEPs are sorted by length (L), and the sequences are given. If proteogenomic support was found, the number of identified peptides is given in column P. For further details on the assigned peptides and spectra, see **Supplementary Dataset 2**. These translons are located on the chromosome (acc. NC_000911), and on plasmid pSYSA (AP004311).

#### Validated intergenically encoded SEPs

The genes *sml0012* and *norf2* were suggested before to encode hypothetical proteins^13,54,68^, but have never been validated. The gene *sml0012* encodes a 45 aa polypeptide and is located upstream of *aroQ*, the 3-dehydroquinate dehydratase involved in the shikimate/chorismate pathway, essential for the synthesis of aromatic amino acids. We validated it by western blotting (**Fig. 6A**) and detection of a single peptide in the iPtgxDB search (**Fig. S14A**). Homologs of Sml0012 are widespread in the cyanobacterial phylum, sharing 38 to 88% aa identity (**Fig. S14A**). Sml0012 might function as an attenuator peptide for *aroQ* translation, as the closely spaced arrangement of both genes in the same operon is conserved. Translation of the 67 aa protein Norf2 was well supported by western blotting, TIS– and TTS-Ribo-seq data, and the detection of 4 peptides in the iPtgxDB search (**Fig. S14B**). There is no homolog for *norf2* in other cyanobacteria, but some closely related proteins exist in gammaproteobacteria (**Fig. S14B**). The region surrounding the *norf2* gene in the *Synechocystis* 6803 genome has been characterized as a genomic island^69^, consistent with its paraphyly. The *norf2* gene overlaps by three codons with the downstream located gene *slr1062* encoding a PemK-type toxin^51^ suggesting Norf2 is possibly the cognate antitoxin.

We also found SEPs on the ∼100 kb plasmid pSYSA encoding the three distinct CRISPR-Cas systems of *Synechocystis* 6803^70^. There are substantial numbers of accessory proteins encoded with type III CRISPR-Cas gene cassettes^71^, and more such genes might have been overlooked. Indeed, there is an sORF close to the 3′ end of the CRISPR3 array encoding a 46 aa SEP, here called Ssr70935, supported by strong Ribo-seq, TIS– and TTS-Ribo-seq coverage (**Fig. S14C**) and by western blotting (**Fig. 6A**). Ssr70935 is a very basic protein due to the presence of 9 lysine and 8 glutamine residues (isoelectric point (pI) of 12.04), a hallmark of nucleic acid-binding proteins.

We validated the translation of three previously defined sRNAs, Ncl1350, Ncr1420 and NsiR7^54^. Ncl1350 and Ncr1420 encode a 57 and a 52 aa SEP, respectively. Homologs of similar size exist in many different cyanobacteria (**Fig. S15** and **S16**). The transcript level of *ncl1350* was previously found to be high and almost constitutive under ten different conditions. The transcription of *ncl1350* is linked with that of the downstream gene *sll0376* encoding a core protein of the cyanoglobule lipid droplet^72^, to which it may be functionally related. Translation of Ncl1420 was validated by six different peptides identified in the iPtgxDB. Transcription of *ncl1420* is highest during exponential growth and very low under adverse conditions suggesting involvement in vegetative growth. In contrast, NsiR7 is upregulated under nitrogen starvation^73^ and encodes a 51 aa SEP. Putative homologs of NsiR7 were identified only in a small number of other *Synechocystis* strains, which share the presence of cysteine residues at conserved positions (**Fig. S16B)** suggesting redox sensitivity.

#### SEPs translated from asRNAs

Three asRNA-located SEPs were chosen for validation. One of them, asRsbU, was found on an asRNA to *slr2031* encoding a homolog to the protein phosphatase RsbU^74^. The asRNA to *slr2031* was detected earlier and divergent transcript accumulation with the *slr2031* mRNA was observed under high light compared to standard conditions and cultivation in darkness^68^. In the Δ*slr2031* mutant, pigment contents (chlorophyll and phycocyanin) were significantly higher compared to wild type^74^. This finding was consistent with reports that spontaneous mutations can be observed in this gene^75^ and that deletion of a 154 bp DNA segment encoding part of the N-terminal domain of Slr2031 led to a glucose-resistant substrain^76^ that also displayed higher pigment contents and showed an increased high-light tolerance^77^. The RNA-seq and Ribo-seq signals are stronger for asRsbU on the asRNA than for the gene *slr2031* in sense orientation (**Fig. 6B**). Within this asRNA, we found two sORFs in-frame with each other, encoding a 62 or a 38 aa small protein. Support from TTS-Ribo-Seq is excellent for both, since they end with the same stop codon. TIS-Ribo-Seq support is very clear and at the optimal distance of 19 nt for the shorter sORF. However, a peptide was found in the custom iPtgxDB search, directly supporting the long variant and expression of the FLAG tagged protein was confirmed by western blotting (**Fig. 6A, B**). Therefore, it is also possible that the 38 aa variant is an IOF to the longer sORF. Corresponding contiguous ORFs exist in some other cyanobacteria including possible start codons at the corresponding positions for the 62 and 38 aa sORFs in *Synechocystis* 6803 (**Fig. 6B**). RsbU-type phosphatases such as Slr2031 commonly belong to a partner switching system that consists of four modules. These are RsbU, a phosphorylatable anti-sigma antagonist, a kinase anti-sigma factor and an alternative sigma factor^78^. This already complex system is made even more complex by the asRNA and the herein encoded SEP.

We also validated the coding potential in two more asRNAs, in as_csx18 to gene *slr7091,* encoding a CRISPR-associated Csx18 protein **(Fig. S17),** and in as_psbC to gene *sll0851,* encoding the photosystem II protein PsbC/CP43 **(Fig. S18)**. For as_csx18 and its cognate gene in sense orientation, *slr7091,* both RNA-seq, Ribo-seq, and TIS-Ribo-seq coverage supported the expression of both DNA strands. The coding potential within *as_csx18* was well supported by the characteristic TIS-Ribo-seq signals 3′ of the start codon at position +19 nt. Our TIS-Ribo-seq signals also indicated that the *slr7091* ORF encodes a 96 aa protein, which is shorter than the 122 aa protein in the KZS, RefSeq and GBK annotations, but consistent with the shorter Prodigal prediction (**Fig. S17A**). The translation of the *as_csx18* sORF was validated by western blot, detecting the SPA-tagged asCsx18 protein (**Fig. S17B**). Homologs of the asCsx18 protein could be encoded on asRNAs to the respective *csx18* gene in *Synechocystis* 6714, and possibly further cyanobacteria (**Fig. S17C**). The asRNA overlapping the 3′ end of the *psbC* coding region originates from a TSS at position 1,347,388 on the forward strand of the chromosome^54^. Thus, this TSS is located 86 nt downstream of the *psbC* stop codon, while the asRNA overlaps >300 nt of the *psbC* coding region, suggesting a regulatory function. Indeed, both transcripts are divergently regulated. While the *as_psbC* transcript is maximum under adverse conditions, *psbC* mRNA accumulation is maximum under optimal growth conditions and exponential growth^54^. Translation of the *as_psbC*-encoded 28 aa SEP was supported by Ribo-Seq, TIS-Ribo-seq, TTS-Ribo-seq data and western blotting **(Fig. S18A, B)**. The asPsbC protein in *Synechocystis* 6803 and several possible homologs in other cyanobacteria are very basic (pI of ∼12) and very rich in alanine **(Fig. S18C)**.

#### Translation of SEPs from alternative start sites within coding regions

In addition to SEPs encoded in intergenic regions or within asRNAs, we also found evidence for the translation of SEPs from alternative start sites within protein-coding genes Internal-Out of-Frame (IOF). Three examples were selected for validation, *infC*-IOF, *crtR*-IOF and *slr1923*-IOF. The *infC*-IOF encodes an only 25 aa SEP, located within the *infC* gene (*slr0974*) for translation initiation factor IF-3 (**Fig. 6C**). Although translation of the 42 aa SEP originating from the IOF within the *crtR* gene and of the SEP originating from the IOF within *cvrA* was well supported (**Fig. S19**), their reading frames appear not as widely conserved. Lastly, we picked three examples of IIF-encoded SEPs (within *sll0871, slr0601* and *slr1397*), but these could not be validated.

To summarize, in addition to the Ribo-seq support. we validated 15 of 18 selected sORFs by western blots, and five were supported by proteogenomic peptide evidence (**Table 1**). From these 15 validated instances, one was annotated as a hypothetical protein (*sml0012*), one was a previously predicted protein-coding gene based on transcriptomic evidence (*norf2*), while the others were known only as sRNAs or asRNAs, or are intragenic IOF-encoded SEPs. These results indicated the protein-coding potential of widely different types of loci.

### A family of genes encoding two proteins by IOF translation

Thus far we have reported on SEPs encoded in intergenic regions, antisense transcripts, or within coding regions. However, the Ribo-seq data also uncovered the existence of a hitherto uncharacterized family of genes that encoded a second, relatively long protein IOF. In **Fig. 7** we show that a 112 aa protein is encoded within the coding sequence for the 151 aa protein Slr0489. From here on, we call these proteins Slr0489L for the longer, annotated gene product, and Slr0489S for the shorter protein, encoded IOF. Reads from the standard Ribo-seq analysis cover the entire annotated *slr0489* gene, but the coverage is visibly higher for the internal segment encoding Slr0489S (**Fig. 7A)**. There are TIS-Ribo-seq signals for both reading frames at the characteristic position 19 nt 3′ of the respective start codons, and TTS-Ribo-seq signals mapped 14 nt 3′ of the respective stop codons. These signals are strikingly higher for the internal segment. Ultimate proof for the simultaneous expression of both proteins was obtained by the proteogenomic detection of 6 different peptides for Slr0489L and 5 for Slr0489S (**Fig. 7B, C**). An Alphafold prediction of the Slr0489L/Slr0489S heterodimer can be found in **Fig. S20.**

**Fig. 7.**
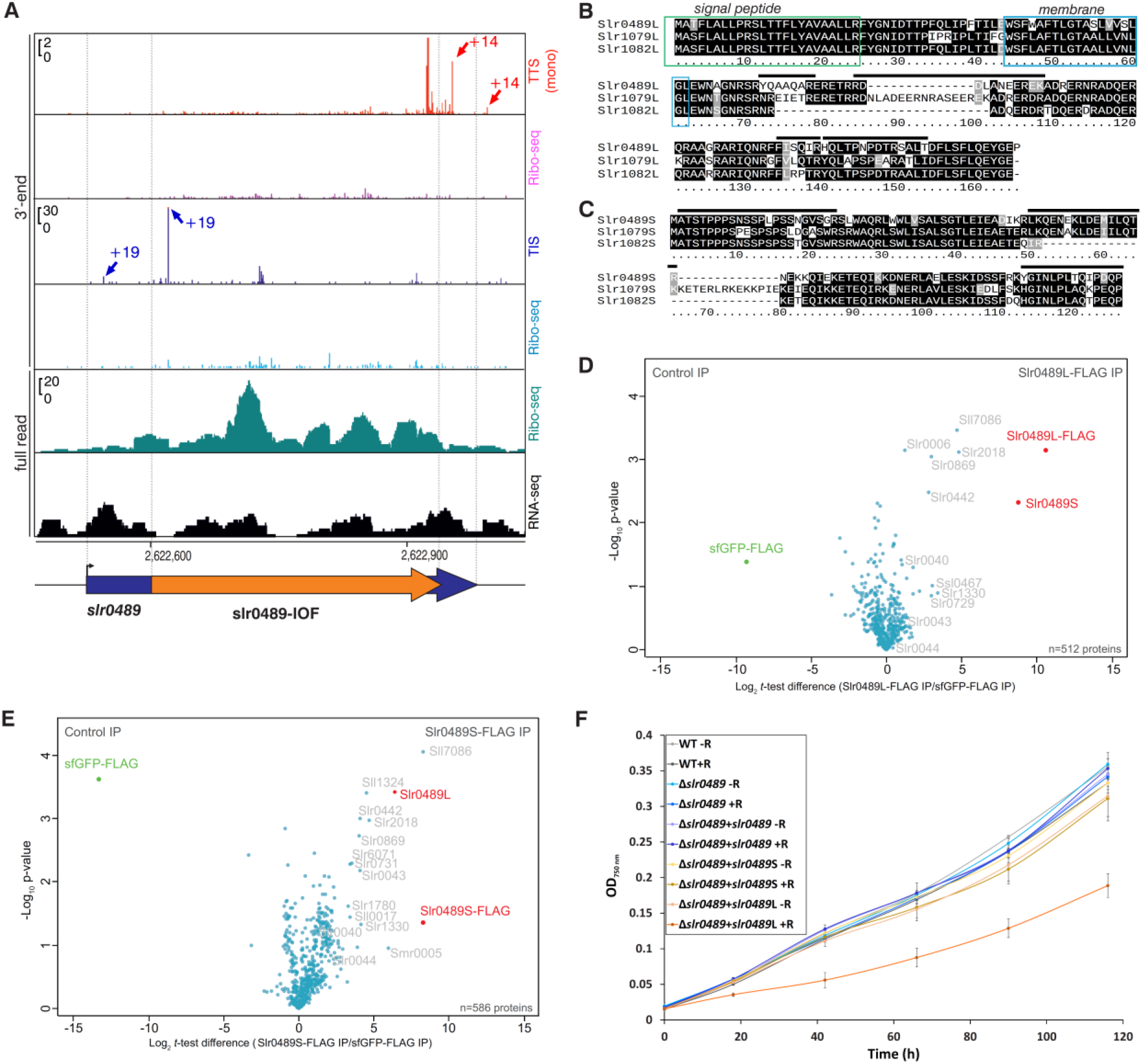
| A family of genes encoding proteins in long internal out-of-frame (IOF) reading frames. **A.** A protein Slr0489S is encoded within the *slr0489* reading frame (blue arrow) by a long IOF reading frame (orange arrow). Read coverage from Ribo-seq and RNA-seq libraries is shown in the lower 2 panels, the Ribo-seq read coverage is higher for the IOF segment. Mapped dual signals from TIS-Ribo-seq at the characteristic position 19 nt 3′ of the respective start codons (two central panels including control), and 14 nt 3′ of the respective stop codons from TTS-Ribo-seq (two upper panels including control) are shown. **B.** Sequence comparison of the proteins encoded by *slr0489* and two closely related loci, *slr1079* and *slr1082*. The proteins expressed from the annotated genes are labeled “L” for long. Signal peptides and transmembrane segments predicted by DeepTMHMM^64^ are boxed. Black horizontal lines indicate some of the proteogenomically detected peptides for Slr0489L. Altogether, 6 peptides were identified in the custom and 5 in the standard iPtgxDB. **C.** Sequence comparison of the proteins (labeled “S” for short) encoded IOF within *slr0489*, *slr1079* and *slr1082*. Black horizontal lines indicate some of the proteogenomically detected peptides for Slr0489S. **D.** Co-IP analysis of Slr0489L-3xFLAG in three replicates. The volcano plot shows the protein interaction partners of Slr0489L-3xFLAG identified by mass spectrometry compared to sfGFP-3xFLAG. The two most enriched proteins, Slr0489L and Slr0489S are colored red, the locus IDs are given for co-enriched proteins in grey. For the complete list of 512 detected proteins and further details, see **Table S8**. **E.** Co-IP analysis of Slr0489S-3xFLAG. Details as in panel D. For the complete list of 586 detected proteins, see **Table S9.** The results of an analogous co-IP experiment for *slr1079* encoding Slr1079L/S are shown in **Fig. S21**. **F.** Effects of overexpressing *slr0489* (without FLAG tag) in a *Synechocystis* 6803 Δ*slr0489* deletion mutant, either with both intact reading frames (Δ*slr0489*+*slr0489* +R), of Slr0489S separately (Δ*slr0489*+*slr0489*S +R), or of Slr0489L from a variant in which a single point mutation prevented the expression of Slr0489S (Δ*slr0489*+*slr0489*L +R). The overexpression was triggered by the addition of rhamnose (+R), which was omitted from the respective controls (-R). The wild type (WT) and Δ*slr0489* deletion mutant served as further controls. The lines show the averaged increase in OD_750_ over 120 h. Cultures were measured in technical duplicates of two independent biological replicates.

A search for related genes yielded two additional loci in *Synechocystis* 6803, *slr1079* and *slr1082*. These genes harbor IOF reading frames as well, and Ribo-seq data supported their expression (**Fig. S21**) and the detection of peptides in the iPtgxDB search directly supported the presence of Slr1079L, Slr1079S and Slr1082S (**Fig. S21D, E**). Notably, the internally encoded proteins Slr0489S, Slr1079S and Slr1082S are closely related to each other, as are Slr0489L, Slr1079L and Slr1082L (**Fig. 7B, C**). However, the respective longer and shorter proteins encoded within each of these loci are not related in sequence or structure. While Slr0489S has a slightly acidic pI of 5.56, Slr0489L has a pI of 9.38. Moreover, Slr0489L shares the presence of a signal peptide and a transmembrane domain with Slr1079L and Slr1082L (**Fig. 7B**). The presence of these three genes does not follow phylogeny — the closely related strain *Synechocystis* 6714 lacks a related gene, while in the only distantly related *Synechococcus* sp. PCC 7003, a close homolog exists (75% identical and 87% conserved aa with Slr0489, sequence ID WP_065714473.1).

To search for interacting proteins, we expressed a FLAG-tagged version of Slr0489L from a plasmid vector and performed coimmunoprecipitation (co-IP) analysis compared to a separate co-IP for sfGFP-3xFLAG as control. The most enriched protein was Slr0489S, suggesting that the internally encoded smaller and the longer 2TM proteins interact with each other (log_2_-fold enrichment of 10.6 for Slr0489L and 8.8 for Slr0489S, **Fig. 7D**). However, the overexpression of *slr0489* inherently leads to a higher level of both, Slr0489L and Slr0489S, as they are both encoded on the same mRNA. Therefore, we constructed a cell line separately overexpressing Slr0489S fused to a C-terminal 3xFLAG tag (Slr0489S-3xFLAG) and performed co-IP. In this case, a log_2_-fold enrichment of 8.3 was observed for Slr0489S and of 6.4 for Slr0489L (**Fig. 7E**). We noticed that in both cases two other highly enriched proteins were the CRISPR3 type III-Bv system Cmr5 protein Sll7086^79^, and the minor pilin Slr2018^80^ suggesting a connection to complexes close to the cell membrane. We conclude that the two different proteins produced from the *slr0489* locus interact with each other. In parallel, we performed the same type of co-IP experiments with Slr1079 (**Fig. S21**). The co-IP using Slr1079L as bait yielded Slr1079S as the most enriched protein (log_2_-fold enrichment of 12.1 for Slr1079L and 6.7 for Slr1079S, **Fig. S21B, Table S10**), while the co-IP using Slr1079S as bait yielded Slr1079L as the most enriched protein (log_2_-fold enrichment of 10.1 for Slr1079S and of 8.8 for Slr1079L, **Fig. S21C, Table S11**).

To measure possible physiological effects of the Slr0489L/S system, we constructed a *Synechocystis* 6803 Δ*slr0489* deletion mutant. In this mutant, we then overexpressed the native *slr0489* gene with intact reading frames for both Slr0489L and Slr0489S. Alternatively, we expressed only Slr0489S, or only Slr0489L. For the latter, we introduced a *slr0489* gene with a single point mutation substituting the ATG start codon by an ACG codon for threonine, thus preventing the translation of Slr0489S. For Slr0489L, this did not change the encoded amino acid sequence because codon 26 was changed from “TAT” to “TAC”, both encoding tyrosine. In a growth experiment lacking the inducer rhamnose, all gene variants had no or only negligible growth effects compared to wild type (**Fig. 7F**). In this line only, clear growth retardation was observed in the presence of the inducer rhamnose, while all other strains grew similarly over a 5-day time course (**Fig. 7F**). We conclude that the sole expression of Slr0489L had a toxic effect on cell growth, which was neutralized in the presence of Slr0489S.

## DISCUSSION

### Genome consolidation effort in *Synechocystis* 6803

*Synechocystis* 6803 is an important model organism, which is reflected by the fact that its genome was the overall third to be fully sequenced 30 years ago^5^. Nevertheless, similar to other model systems, precise determination of coding regions has remained challenging up to this date, visible in differences in the available annotations of the identical genome sequence, such as the ones used here, deriving from NCBI RefSeq (2022), GenBank (2016) and the extended KZS 2009 annotation. These annotations differ both in terms of the overall numbers of coding sequences predicted and their precise start sites (**Fig. 3**). Previously, among prokaryotes, TIS– and TTS-Ribo-seq were combined in *E. coli*^40^ and in *Campylobacter jejuni*^32^, while Ribo-seq was combined with mass spectrometry analyses in the work on the Archaea *Methanosarcina mazei*^34^ and *Haloferax volcanii*^36^ and on the plant growth promoting soil bacterium *Sinorhizobium meliloti*^35^. While the value of riboproteogenomic analyses is broadly acknowledged^81^, TIS-Ribo-seq, TTS-Ribo-seq and a proteogenomics analysis have not been combined in any prokaryote thus far.

As we show in this study, several reading frames should be extended or truncated (**Fig. 3**), which resolves ambiguities concerning functional start codons. Nevertheless, we are aware that ultimate proteomics proof requires in many instances specialized N-terminomics approaches^82^, beyond the scope of this work. We have integrated the different reference annotations, the *ab initio* gene predictions from Prodigal 2.6.3^56^ and ChemGenome 2.1^57^, plus the*in-silico* ORFs from a full six-frame translation prediction into a standard and a custom integrated proteogenomic search database (iPtgxDB). The custom iPtgxDBs integrated the top Ribo-Seq candidates, while the standard iPtgxDB captured almost the entire protein-coding potential of the genome. Consequently, the first is maximized for sensitivity and better search statistics and the second for completeness leading to differences, in particular for the identification of previously unknown proteins (see **Figure S10**). Both iPtgxDBs are accessible through RIBOBASE as a single platform. The generation of a database comprising different annotations is generic; it enables researchers to jointly consolidate the genome annotation of their favorite model organism in the context of overlaid experimental data. The addition of data from multiple conditions and sources forms the basis to more comprehensively discover the full coding potential of genome sequences. To facilitate access to the entire set of data, we developed a JBrowse 2 instance^83^, a web-accessible, fully interactive genome browser provided at https://www.bioinf.uni-freiburg.de/∼ribobase/.

While SEPs mediate multiple important functions in all domains of life, they frequently have escaped annotation. In line with these observations, our riboproteogenomic analysis identified previously unknown candidate genes, of which several were experimentally validated to be present (**Table 1**, **Figs. 5, 6, 7, S17–S18**). From the total of 808 manually checked instances of extended, truncated or previously unknown candidates supported by Ribo-seq, 163 were considered as likely correct. This allows to extrapolate a number of likely 571 true candidates, among them a number of 293 expected unannotated translons (**Table S12**). Our discovery of many new SEPs is in line with the recent prediction of extensive microprotein families in the Enterobacteriaceae^16^. More than 140 previously unknown small proteins (in this case even ≤50 aa) have been manually validated over the span of roughly ten years in *E. coli*^84^ with evidence for several hundred additional ones below 100 aa, which is consistent with our findings. We have previously characterized the *Synechocystis* 6803 SEPs AtpΘ as a regulator of ATP synthase^17^, NblD as a factor involved in the nitrogen starvation-induced degradation of the phycobilisomes^45^, AcnSP as a regulator of aconitase activity^44^ and NirP1 as regulating nitrite reductase^46^ demonstrating the physiological relevance of SEPs. In extension, many of the newly identified candidates in this study warrant further consideration. These include the *slr1421*-*ncr1610-sORF1* locus (**Fig. 5F and G**), or the *ncl1450/ncr1470* locus, which are related to toxin-antitoxin-like systems. Our data indicate that a SEP evolved in the asRNA Ncr1470, which can be detected in some but not the majority of related loci in other cyanobacteria possessing homologs of *ncl1450* (**Fig. 5C, E**). The *ncl1450* gene and its overlapping *ncr1470* are located in an arrangement between *slr0841* and *slr0842* resembling an excludon^63^. The Ncl1450/Ncr1470 pair might constitute an associated type I toxin-antitoxin candidate system, possibly with a stabilizing role^85^. Secondarily, the Ncr1470 sORF may have evolved in an originally non-coding asRNA.

### CRISPR-associated hidden genes

We present two instances of previously unknown CRISPR-associated sORFs and corrected the ORF for gene *slr7091/csx18* **(Fig. S14C, S17A)**. The here defined gene *ssr70935* encodes a 46 aa protein. It is located directly between the CRISPR array of a type III-Bv system^79^, and the genes *slr7094* and *slr7095* with which it is cotranscribed as part of a tricistronic operon. Homologs of *ssr70935* in other cyanobacteria are also located adjacent to CRISPR-Cas loci, indicating a genetic association. There are several examples for the relevance of small proteins encoded in atypical sORFs associated with CRISPR-Cas systems. In the archaeal CRISPR-Cas type I-B system of *Haloferax volcanii*, the Cas11b is encoded by an internal in-frame translation within the *cas11* gene^86^. In a study on the type I-D system of the here investigated *Synechocystis* 6803, cryo-EM structures of the purified complex revealed an additional small subunit Cas11d^87^. Initially, no gene could be assigned to this protein until it was discovered that it originated from an IIF translation within the gene of the large subunit *cas10d*^87^. Therefore, the here identified asCsx18 protein translated from an asRNA further illustrates the coding capacity of CRISPR-Cas systems to be extended by translation of ORFs from unorthodox locations. While currently neither the precise function of Csx18 nor of the asCsx18 protein is known, we noticed that Csx18 proteins are generally associated with type III-D and III-B CRISPR systems and likely adaptation-associated^71^. This would be consistent with the location of *slr7091* and its asRNA just upstream of the *cas1* and *cas2* adaptation module and next to the CRISPR3 effector genes on plasmid pSYSA in *Synechocystis* 6803^79^.

### Genes-in-genes

Our work not only provided evidence for additional sORFs, but also uncovered the existence of a hitherto uncharacterized family of 3 genes encoding a second, relatively long protein out-of-frame, which may be considered as genes-in-genes (**Fig. 7, Fig. S21**). This genetic arrangement resembles some bacteriophage loci encoding two-component spanins. Spanins are lysis proteins produced by phages that are required to disrupt the outer membrane during lysis of gram-negative host cells. Two-component spanins consist of an integral inner membrane i-spanin, and an outer membrane o-spanin, which is a lipoprotein^88^. In bacteriophage lambda, the reading frame Rz1 encoding the o-spanin is embedded within the reading frame Rz encoding the i-spanin^88^. The lambda phage i-spanin possesses a transmembrane domain and is located in the inner membrane. The C-terminal domain of the i-spanin, which extends into the periplasm, consists of alpha helical segments. In contrast, the o-spanin possesses a proline-rich region, is connected to the outer membrane, lacks a transmembrane domain and any other detectable structure^89^. Therefore, there are striking parallels to the here described *slr0489, slr1079* and *slr1082* loci. In addition to the genetic arrangement, the three proteins Slr0489L, Slr1079L and Slr1082L share the presence of a signal peptide and a transmembrane domain (**Fig. 7B**), while Slr0489S, Slr1079S and Slr1082S encoded by the embedded ORFs possess a conserved proline-rich motif at positions 6–8 (**Fig. 7C**), recognized as a membrane fusion motif in o-spanins^88^.

However, there are also important differences. These three loci do not belong to any bacteriophage or prophage genome and their overexpression did not cause the cells to lyse. However, we did observe that the expression of Slr0489L without Slr0489S led to growth retardation (**Fig. 7F**). Thus, the tight genetic connectivity between Slr0489L and Slr0489S first had to be disentangled to observe this effect, resembling a toxin-antitoxin effect in which binding of Slr0489S negates the deleterious effect of Slr0489L on our cultures. This resembles rather a defense-associated toxin–antitoxin relationship, as in the Panoptes defense system^90,91^, providing an exciting topic for further research.

### Outlook and limitations of this work

Metagene analyses were performed to compare our TIS– and TTS-Ribo-seq data to previous results from *E. coli* and *Campylobacter* (**Fig. S22**). While the footprint lengths were identical or very similar among the 3 bacteria (ribosomes stalled on start codons: 32 nt for both *Synechocystis* and *Campylobacter,* and 31 nt for *E. coli;* ribosomes stalled on stop codons: 29 nt), the cyanobacterial ribosomes were offset by 4 nt in the 3’ direction. This indicates a potentially fundamental difference in the ribosome structure of cyanobacteria compared to *E. coli* and *Campylobacter*, worth studying more deeply in the future.

There are two previously published instances of alternative proteins produced from single genes in *Synechocystis* 6803. These are the IIF-encoded CRISPR-Cas subunit Cas11d^87^ and the two alternative forms of ferredoxin:NADP oxidoreductase^92^. Translation of the corresponding genes *slr7011* and *slr1643/petH* can be seen in the Ribo-seq data, but not the initiation of translation at the internal sites. This points at a sensitivity limit and that we rather under– than overestimate the number of alternative ORFs. Part of this may relate to the fact that we have established TISv and TTS-Ribo-seq and performed that analysis under one condition only. The shorter ferredoxin:NADP oxidoreductase proteoform is mainly made under heterotrophic conditions^92^, which were not included here. On the other hand, we are aware of the fact that the induced overexpression of SEP candidates may lead to artefacts regarding the protein stability, localization and aggregation. Therefore, endogenous tagging and the use of native promoters should be preferred for physiological studies.

Nevertheless, we provide information on instances of previously unknown translons and suggest corrections to annotated genes. This list of translons includes candidates IOF within annotated genes and loci that were previously considered non-coding. We validated several instances related to the CRISPR-Cas systems, to photosystem II, and regulatory factors. The functional analysis of these translons is an exciting topic for further research.

## Methods

### Strains and growth conditions

*Synechocystis* 6803 cultures were grown in TES-buffered BG11^93^. One hundred mL cultures were grown in 250 mL Erlenmeyer flasks with shaking under an illumination of 40 µmol photons m^-2^ s^-1^ and a temperature of 30°C until an OD_750_ of 0.6-0.8 was reached. For TIS profiling, 10 µg/mL retapamulin final concentration (∼6× MIC) solved in DMSO was added to the culture and incubated for 15 min with continued cultivation. For control, an equal volume of DMSO was added (mock-treated cultures). For TTS profiling, the cultures were treated with 10 µM final concentration (∼12.5× MIC) of the proline-rich antimicrobial peptide Api137 (solved in ddH_2_O, NovoPro BioSciences, Shanghai) for 15 min. Mock-treated controls without the antibiotic were grown in parallel. Two independent biological replicates were prepared and processed for Ribo-seq, TIS and TTS profiling experiments. Immediately before cell harvest, aliquots were taken from the untreated cultures for total RNA analysis, mixed with 0.2 vol stop mix (5% buffer-saturated phenol (Roth) in 95% ethanol), and snap-frozen in liquid N_2_.

### MIC assays for TIS and TTS profiling

In *E. coli* TIS profiling, mutants lacking the gene for efflux transporter *tolC* were used because translation stalling effects were limited due to retapamulin secretion via the transporter in the wild type strain^30^. A homologous transporter is also present in *Synechocystis* 6803^94^ and might be responsible for the efficient efflux of retapamulin. This is unfavorable for TIS profiling, since the appropriate antibiotic levels need to be maintained in the cell to stall initiating ribosomes. Minimum inhibitory concentrations (MIC) of retapamulin (Sigma CDS023386) were determined in liquid cultures for the wild type and the Δ*tolC*, Δ*acrA* and Δ*acrB* transporter mutants. Added retapamulin inhibited all three mutants, while the wild type resisted retapamulin concentrations up to 1 μg/mL (**Fig. S1A, B**). Without antibiotics, all strains grew steadily from an OD_750_ of 0.2-0.4 to > 2 within 3 days. To avoid possible unphysiological effects of working with mutant strains, we chose the wild type for retapamulin-assisted TIS profiling, applying a ∼6× MIC of 10 μg/mL retapamulin in 100 mL cultures. To establish proper conditions for TTS-Ribo-seq, MIC were determined using the proline-rich antimicrobial peptide (prAMP) Oncocin 112 (Onc112, Novopro) in small scale liquid cultures. Onc112 is similar in chemical nature (sequence VDKPPYLPRPRPPRrIYNr-NH_2_) compared to Api137 (Gu-ONNRPVYIPRPRPPHPRL-OH (Gu=N,N,N’,N’-tetramethylguanidine and O=L-ornithine))^37^, but is less costly. Both prAMPs are supposed to be taken up by a SbmA-like transporter^95^. A MIC of 0.8 μM Onc112 was determined for *Synechocystis* 6803 wild type (**Fig. S1C**). Consequently, a 12.5-fold MIC of 10 μM Api137 was applied in the TTS approach.

### Ribosome Profiling

Cells were collected by fast filtration through hydrophilic polyethersulfone filters (Pall Supor®-800, 0.8 μm). The cells were scratched off the filters, quickly frozen in liquid nitrogen and stored at –80°C. Cells were then disrupted by cryogenic grinding (Retsch M-400) using prechilled grinding ball and chamber in the presence of 1 mL frozen lysis buffer (50 mM HEPES pH 8, 100 mM NaCl, 10 mM MgCl_2,_ 5 mM CaCl_2,_ 0.4 % Triton X-100, 2 mM DTT, 500 U RiboLock). In contrast to other approaches, chloramphenicol has been omitted in all buffers to avoid bias^22^. The grinding (15 Hz, 3 min) was re-peated four more times. The grinding jars were cooled with liquid nitrogen between cycles. Frozen powder was collected and then centrifuged (4°C, 8,000 × g, 10 min) to separate cell debris and lysate. The lysate was kept on ice for further processing, or stored at – 80°C. For the digestion, an aliquot containing ∼100 µg nucleic acid (quan-tified with nanodrop) was filled up to a volume of 500 µL with digestion buffer (50 mM HEPES pH 8, 100 mM NaCl, 10 mM MgCl_2,_ 10 mM CaCl_2,_ 0.4% Triton X-100, 2 U/mL TURBO DNase, 100 U/mL RiboLock) and digested with 300 U MNase (NEB) for 45-60 min at 25°C, 450 rpm. A negative control was treated the same way, but MNase was omitted. To stop the digestion, 5 mM EGTA (final concentration) was added. The lysate was immediately layered on a sucrose gradient with 10%-55% sucrose solved in gradient buffer (100 mM NH_4_Cl, 20 mM Tris-HCl pH 8, 5 mM CaCl_2,_ 10 mM MgCl_2_, 2 mM DTT). Gradients were formed by Gradient Master (Biocomp) in 1:50 min, with 80° angle and 21 rpm, at 4°C, and then centrifuged for 2.5 h, 217k × g at 4°C in a Beckman Coulter Optima L-80 XP ultracentrifuge and SW40 Ti rotor), the gradients were fractionated and the absorbance at 260 nm was measured with Gradient Frac-tionator (Biocomp) integrated TRIAX flow cell. For TIS profiling, the monosomes (70S) fractions were collected for both the untreated (Ribo-seq) and Retapamulin-treated samples (TIS). For TTS profiling, fractions corresponding to monosomes, disomes as well as trisomes were collected for both untreated (Ribo-seq) and Api137-treated sam-ples (TTS). The collected fractions were immediately frozen in liquid N_2_ until further analysis.

### Isolation of ribosome footprints

Ribosome footprints were prepared from monosomes, disomes, and trisomes. Total RNA was extracted from fractions or cell pellets using hot phenol-chloroform-isoamyl alcohol (25:24:1, Roth) or hot phenol (Roth), respectively^96^. Ribosomal RNA (rRNA) was depleted from 5 µg of TURBO DNase-digested total RNA by subtractive hybridization with the Pan-Prokaryotes riboPOOLs (siTOOLs, Germany) according to the manufacturer’s protocol with Dynabeads MyOne Streptavidin T1 beads (Invitrogen). Total RNA was fragmented with an RNA fragmentation reagent (Ambion) following the manufacturer’s instructions. Monosomes RNA and fragmented total RNA were size selected (26–34 nt) on 15% polyacrylamide (PAA), 7 M urea gels, as previously described^97^ using RNA oligonucleotides NI-19 and NI-20 as ladder. In parallel, RNA from disomes (50-80 nt) and trisomes (80-120 nt) from both untreated and Api137-treated samples were size selected on the same gel electrophoresis as indicated above using Ultra-low RNA ladder (RNA Decade Markers, #AM7778, ThermoFischer). RNA was extracted from the gel, cleaned and concentrated by isopropanol precipitation with 15 μg of GlycoBlue (Ambion) and dissolved in H_2_O. cDNA libraries were prepared by Vertis Biotechnologie AG (Freising, Germany) using the adapter ligation protocol without fragmentation. First, an oligonucleotide adapter was ligated to the 3′ end of the RNA molecules. First-strand cDNA synthesis was performed using M-MLV reverse transcriptase and the 3′ adapter as the primer. The first strand of cDNA was purified, and the 5′ Illumina TruSeq sequencing adapter was ligated to the 3′ end of the antisense cDNA. The resulting cDNA was PCR-amplified to approximately 10–20 ng/μL using a high-fidelity DNA polymerase. The DNA was purified using the Agencourt AMPure XP kit (Beckman Coulter Genomics) and analyzed by capillary electrophoresis. The primers used for PCR amplification were designed for TruSeq sequencing according to the instructions of Illumina. The following adapter sequences flank the cDNA inserts: TruSeq_Sense_primer: (NNNNNNNN = i5 Barcode for multiplexing) 5′-AATGATACGGCGACCACCGAGATCTACAC-NNNNNNNN-ACACTCTTTCCCTACA CGACGCTCTTCCGATCT-3′; TruSeq_Antisense_primer: (NNNNNNNN = i7 Barcode for multiplexing) 5′-CAAGCAGAAGACGGCATACGAGAT-NNNNNNNN-GTGACTGGAGTTCAGACGTGT GCTCTTCCGATCT-3′. cDNA libraries were pooled on an Illumina NextSeq 500 high-output flow cell and sequenced in single-end mode (75 cycles, with 20 million reads per library) at the Core Unit SysMed at the University of Würzburg.

### Ribo-seq, TIS and TTS profiling data analysis

Ribo-seq and TIS/TTS datasets were processed using the HRIBO workflow (version 1.7.0)^52^, which has previously been used for the analysis of other prokaryotic Ribo-seq data^35,36,98^. Sequencing read files were processed with a Snakemake^99^ workflow that downloads all required tools from bioconda^100^ and automatically determines the necessary processing steps. Adapters were trimmed from the reads with cutadapt (version 2.1)^101^ and then mapped against the *Synechocystis* genome with segemehl (version 0.3.4)^102^. Reads corresponding to rRNA and other multiple mapping reads were removed using SAMtools (version 1.9)^103^. Quality control was performed by creating read count statistics for each processing step and RNA class with Subread featureCounts (1.6.3)^104^. All processing steps were analyzed with FastQC (version 0.11.8)^105^, and the results were aggregated with MultiQC (version 1.7)^106^.

Read coverage files were generated with HRIBO using different full-read mapping approaches (global or centered) and single-nucleotide mapping strategies (5′ or 3′ end). Read coverage files using two different normalization methods were created (mil and min). For the mil normalization, read counts were normalized by the total number of mapped reads within the sample and scaled by a per-million factor. For the min normalization, the read counts were normalized by the total number of mapped reads within the sample and scaled by the minimum number of mapped reads among all analyzed samples. The coverage files generated using the min normalization and the global mapping (full read) approach were used for genome browser visualization.

### Metagene analysis of ribosome occupancy

A genome wide analysis of ribosome occupancy was conducted using TIS-Ribo-seq and TTS-Ribo-seq. To this end, all annotated coding sequences (CDS) were collected, and windows consisting of upstream and downstream regions around each annotated start codon (or stop codon, respectively) were defined. For each position in the analyzed window, reads occupying that position were counted for each annotated CDS. The total number of reads per position was subsequently reported and plotted in a metagene graph. To generate a high-confidence set of genes, multiple filters were applied. These filters removed genes in close-proximity (≤50 nt apart), genes below a coverage threshold of 10 RPKM and genes that fell below a minimum length cutoff (i.e., genes shorter than the downstream window for start codons or shorter than the upstream windows for stop codons). However, we performed a parallel metagene analysis focusing on 39 short ORFs (≤50 aa) that confirmed that translation initiation patterns in these shorter sequences did not differ substantially from those observed in longer ORFs (compare **Fig. 1E** and **Fig. S2**).

To assess sequence motifs in the 5’-UTR regions of annotated start codons, all annotated CDS contributing to positions with the highest ribosome occupancy (at offset of +2, +4, and +19) were collected, and sequence windows were extracted. Motif plots were then generated using weblogo (3.7.9)^107^.

All scripts used for metagene analysis are available in our Github repository (https://github.com/RickGelhausen/Synechocystis_pub_scripts_2025/).

### Ribo-seq ORF predictions using HRIBO modules

ORFs were called with an adapted variant of REPARATION^108^ using blast instead of usearch (see https://github.com/RickGelhausen/REPARATION_blast) and DeepRibo^109^. Generic feature format (GFF) track files with this information, plus potential start and stop codons and ribosome binding site information were created for in-depth manual genome browser inspection. Summary statistics for KZS and GenBank annotated and merged translons detected by REPARATION and DeepRibo were computed in a tabularized form, including, among other values, translation efficiency (TE), RPKM (reads per kilobase of transcript per million mapped reads) normalized read counts, codon counts, and nucleotide and aa sequences (see **Table S2** for annotated genes and **Table S3** for newly predicted candidates or known genes with suggested corrections). Annotated sORFs were classified as translated if they fulfilled an empirical mean TE cut-off of ≥ 0.288 and RNA-seq and Ribo-seq RPKM of ≥ 10 (cut-offs chosen based on the lowest TE and RPKM values associated with housekeeping genes [i.e., ribosomal protein genes] and the genes detected by proteomics). To identify sORF candidates based on Ribo-seq coverage, we inspected HRIBO ORF predictions from DeepRibo and REPARATION. As DeepRibo is prone to a high rate of false positives^110^, we first generated a reasonable set of potential sORFs by applying the following expression cut-off filters: mean TE of ≥0.288 and RNA-seq and Ribo-seq RPKM of ≥40 (in one of the replicates). In addition, the sORF candidates were required to have a DeepRibo prediction score^109^ of ≥-3.009 that allows for ORF candidate ranking. The filtered sORFs were then subjected to manual curation as described^110^. This parallel inspection of Ribo-seq and RNA-seq read coverage files in a genome browser allowed for asserting the translation status for the predicted sORFs. Briefly, the Ribo-seq and RNA-seq read coverage files were loaded in the Integrated Genome Browser (IGB) along the sequence of the reference genome and the KZS 2009 annotation (with annotated 5’– and 3’-UTRs and further manual curation from the Hess laboratory). The Ribo-seq and RNA-seq read coverage files (normalized to the lowest number of reads between the two) were visually inspected with similar scales. To assess translation of the predicted sORFs, we used the following criteria: 1) Ribo-seq read coverage within ORF boundaries with the detection of ribosome footprints in the UTRs (15-16 nt) near the start and stop codons resulting from initiating and terminating ribosomes; 2) exclusion of Ribo-seq read coverage from the residual 5’-and 3’-UTRs; 3) the shape of the Ribo-seq read coverage; here the evenness of the read coverage was considered and predicted sORFs with uneven read coverage (exhibiting peaks with plateau, which resulted from either RNA structures or cDNA library preparation artifacts) were not taken into account; 4) the Ribo-seq read signal was generally required to be comparable to or higher than the transcriptome signal from the RNA-seq library. For visualization, we developed an interactive web-based genome browser using JBrowse 2 (https://www.bioinf.uni-freiburg.de/∼ribobase/)^111^, where the coverage files for the Ribo-seq replicates, the different genome annotations, results of proteogenomic mappings and the predicted sORF can be accessed interactively.

### ORFBounder: TIS and TTS peak calling

ORFBounder is a standalone, command-line tool implemented in Python 3 for detecting translation initiation sites (TIS) and translation termination sites (TTS) from ribosome profiling data. It builds conceptually on peak-based TIS detection (RETscript^53^) and is a substantial reimplementation of our earlier StartStopFinder scripts^32^. The tool, documentation, a toy example, and unit tests are available at https://github.com/RickGelhausen/ORFBounder.

ORFBounder takes standard sorted and indexed BAM files together with a GFF3 annotation as input; no pre-computed density files are required. Run parameters — read lengths, normalization method, mapping strategy, peak-height thresholds, and the assignment of files to samples, replicates, and conditions — are set in a configuration spreadsheet, with read-length– and sample-specific P-site offsets supplied in JSON format. Internally, reads within the configured length range are assigned to single-nucleotide P-site positions using these offsets, and per-nucleotide densities are computed and normalized on each strand. Candidate TIS and TTS peaks are then called as local density maxima exceeding the peak-height threshold, scored by footprint enrichment, assigned to the nearest in-frame start or stop codon, and reconciled with the existing annotation. When matched RNA-seq libraries are provided, ORFBounder additionally computes RPKM, translational efficiency, and log₂ fold-changes, and merges results across all samples automatically. Outputs comprise GFF files for genome-browser visualization and a consolidated Excel report containing peak heights, scores, fold-changes, expression values, and codon and sequence information across samples.

For this study, TIS-Ribo-seq and TTS-Ribo-seq data were processed with ORFBounder (v2.0.0) using read lengths and P-site offsets determined by metagene analysis (**Fig. 1E, 2E; S2, S3 and S4**).

### Candidate selection for manual curation

To define a set of previously unknown translons for manual curation, the three different datasets resulting from (1) classical Ribo-seq, (2) TIS-Ribo-seq and (3) TTS-Ribo-seq were filtered separately. Then, the filtered results were combined; for a translon to be retained as a candidate, it had to be detected by Ribo-seq and at least one of the other two methods (TIS or TTS-Ribo-seq). In other words, candidates had to correspond to the union of the Ribo-seq ∩ TIS-Ribo-seq, Ribo-seq ∩ TTS-Ribo-seq, or Ribo-seq ∩ TIS-Ribo-seq ∩ TTS-Ribo-seq overlaps. Applying this scheme yielded 2,708 candidate translons that were carried forward for manual curation.

The three datasets were filtered as follows:

1. Ribo-seq data were analyzed using the workflow HRIBO (version 1.4.3)^52^. ORFs were initially called with two algorithms, REPARATION^108^, but using blast instead of usearch^35,36^, and with DeepRibo^109^. Summary statistics included TE, RPKM normalized read-counts and codon counts. The nt and aa sequences for all available annotated and merged previously unknown translons detected by REPARATION and DeepRibo were computed in a tabular format (**Tables S2** and **S3**). For benchmarking, we used a dataset of 15 independently validated small and alternative translons (Ncl1450, Ncr1470, NblD, NirP1, NsiR7, Norf4, SliP4, as_rsbU, AcnSP, AtpΘ, IOF slr0978, IOF sll1954, IOF slr1082, IOF slr0489 and IOF slr1079). The worst averaged translational efficiency (aTE) for these translons was detected for NsiR7 with 0.32. However, one of the annotated house-keeping genes, *rpl36*, encoding ribosomal protein L36 had an even lower TE of 0.288 in one of the replicates. Therefore, we set the minimum threshold for aTE to ≥0.288. Similarly, we set the DeepRibo prediction score at ≥-3.009 according to the score of –3.0091 for gene *psbF* (*smr0006*) encoding the cytochrome b_559_ beta chain (photosystem II protein VI). The other filter criteria were that the RPKM was ≥40 in either of the 2 replicates, the number of codons ≥12 including the stop codon and that none of these translons was annotated at the time of analysis.
2. TIS-Ribo-seq were analyzed using ORFBounder^110^. According to the results of our metagene analysis, we required the three different distances from the mapped 3’ end of the peak to be located at 19, 4 and 2 nt from the respective start codon. The other filter criteria applied to the potential candidates called by ORFBounder, were that the RPKM was ≥40 in either of the 2 replicates for the TIS as well as the associated Ribo-seq read mapping, a TIS/Ribo-seq log_2_FC of ≥1 in either of the 2 replicates, a number of codons ≥12, an aTE of ≥0.282 and that none of these ORFs was annotated.
3. The TTS-Ribo-seq filter criteria applied to the potential candidates called by ORFBounder, were an RPKM value ≥10 in either of the 2 replicates for the TTS as well as the associated Ribo-seq read mapping, a TTS/Ribo-seq log_2_FC of ≥1 in either of the 2 replicates, a codon count ≥12, an aTE of ≥0.15 and that none of these ORFs was annotated. These criteria were applied separately to the “TTS-next” and the “TTS-furthest” dataset. Further manual curation was performed on 808 randomly chosen instances from this list of 2,708 candidate translons (results in **Table S3**, extrapolation in **Table S12**). Criteria were if Ribo-seq coverage was contiguous or interrupted by segments of lacking coverage in the two replicates and if the TIS or TTS signals picked by the algorithm appeared to represent cDNA synthesis artefacts.

### Conservation and protein domain analysis

The identification of small protein homologues was performed using Blastp and TBlastn searches in bacteria at the National Center for Biotechnology Information website (https://blast.ncbi.nlm.nih.gov/Blast.cgi). The amino acid sequences of candidates identified in this study were used as query sequences. For TBlastn, the following parameters were used: the filter for low complexity regions off, a word size of 6, 60% coverage of the query sequence with at least 40% identity, an E value ≤100 to capture all potential orthologs, and an E value <0.1 for high-confidence hits^35^. Moreover, novel small protein candidates were further analyzed for signal peptides and transmembrane domains using deepTMHMM^64^.

### Integrated proteogenomics search databases

Integrated proteogenomics databases (iPtgxDBs) were generated based on the *Synechocystis* 6803 genome sequence (RefSeq ID: GCF_000009725.1). Annotations from reference genome annotation centers (NCBI RefSeq (2022), GenBank (2016; from here GBK) and the KZS annotation, two *ab initio* gene predictions (Prodigal 2.6.3^56^ and ChemGenome 2.1^57^), and an *in-silico* ORF prediction considering the in bacteria most frequent alternative start codons GTG, TTG and CTG^58^ were hierarchically integrated into a standard iPtgxDB (**Table S7A**) as previously described^49^. A minimum length cutoff of 10 aa was chosen for the *in-silico* ORFs (closely matching the minimal length of Ribo-seq identified novel CDS candidates of 11 aa) and the iPtgxDBs were generated using the strict trypsin cleavage pattern (cleavage also before P). Since ChemGenome can only process fastA files containing a single contig, ChemGenome and Prodigal predictions were generated separately for the chromosome and each plasmid and then combined. To improve search statistics, we also created a much smaller custom iPtgxDB (**Table S7B**) as described^35^. For this, we included the 2,708 top candidates implicated by TIS, TTS and Ribo-seq data instead of the 5,071 ChemGenome entries and 107,626 *in-silico* ORF predictions included in the much larger standard iPtgxDB (**Figure 3**, **Tables S7A and B**).

### Analysis of available mass spectrometry data and post-processing

Public proteomics data for *Synechocystis* 6803^48^ were downloaded from 2 large datasets deposited in PRIDE (PXD033243, 628 files; PXD033290, 367 files) using pridepy 0.0.2^112^ and converted to mzML format with pyrawr 0.1.0. The files contained six different data types (unlabeled, dimethyl labeled with and without phosphoproteomics, TMT-6 labeled, and TMT-10 labeled with quantification at the MS2 or MS3 level). For each data type, three searches were performed (**Fig. S6**): A regular search against an NCBI RefSeq database (November 2022) was carried out to establish the coverage of the annotated proteome (83%). Searches against the larger standard iPtgxDB and a smaller custom iPtgxDB enabled the discovery of proteomic evidence for the expression of up to 69 previously unknown SEPs (**see Supplementary Dataset 4**). FragPipe 21.1 was used to search the data with MSFragger 4.0^113^ and validate peptides with PeptideProphet (Philosopher 5.1.0^114^). Apart from specifying the labels for the TMT and dimethyl searches, standard parameters were used for the corresponding workflows. The search results in PEPxml format were processed using custom Java tools as described previously^49^: The results of all 6 searches were combined for each of the three databases and filtered based on the PeptideProphet probability using a false discovery rate (FDR) of 0.0065% at the peptide-spectrum match (PSM) level, leading to protein level FDRs of 0.83% for the searches against the standard iPtgxDB and 0.98% for the searches against the custom iPtgxDB. To obtain a list of proteins identified with higher confidence, they were further filtered for a minimum number of PSMs depending on the origin of the annotation (≥2 PSMs if annotation came from RefSeq, GBK, KZS, or Ribo-Seq; ≥3 PSMs if annotation was based on *ab initio* prediction by Prodigal or ChemGenome; ≥4 PSMs if based on a prediction of *in-silico* ORFs). Based on the iPtgxDB identifier, proteins were then categorized into annotated proteins, previously unknown proteins, longer and shorter proteoforms of annotated proteins, and expressed pseudogenes according to the RefSeq annotation. Importantly, all peptides were also classified with the PeptideClassifier tool^115^, adapted for prokaryotic proteogenomics^49^. This allowed to filter the most information-rich peptides that either unambiguously identify one unique novel proteoform (class 1a), a subset or all proteoforms encoded by a gene model (class 2a, class 2b) and de-select those peptides shared by proteoforms encoded by different gene models (class 3b) (**Fig. S7**). For more information, see (https://iptgxdb.expasy.org/creating_iptgxdbs/) and **Supplementary Dataset 4**. The results of the proteogenomics searches are provided in **Supplementary Dataset 1**.

### Consolidation of genome annotations

A consolidated version of major reference annotation sources included in the standard iPtgxDB (see above) was generated by post-processing the GFF file produced when creating the iPtgxDB using a custom Python script. A comparison of RefSeq, GBK, KZS and Prodigal illustrates that substantial differences exist (**Fig. 3**). Notably, the Hess laboratory has been using their manually curated and extended KZS genome annotation as basis, while we used the RefSeq2022 annotation as main reference for the proteogenomics approach. This can nevertheless be resolved; it in fact allowed us to identify evidence for ten annotations from Genbank 2016 and one KZS annotation that were removed in RefSeq, but still represent valid protein-coding genes as judged by the protein expression evidence we detected. We in fact assume that many research communities could benefit from using such an integrative approach. The consolidated GFF file was constructed such that for each annotation cluster (annotations from different sources that share the same stop codon position), the longest annotation was used to represent the cluster. A respective cluster was then colored according to the conservation among annotations: clusters represented in all reference annotation sources were colored grey if all annotations were identical and yellow if any of the annotations differed in length, while clusters only appearing in a subset of annotations were colored red. Coloring entries in JBrowse2 by the color attribute in the GFF file was achieved by writing a simple no-build plugin.

### Translon terminology

In RIBOBASE and in **Tables S1–S4** we used systematically the “NC_000911.1:3458023-3458145:+” type of identifiers combining the replicon ID (here NC_000911.1 for the chromosome) with the precise translon’s start and end sites and the respective orientation (+ and – for forward and reverse strand). In addition, we provide in Ribobase other relevant identifiers, such as, for example, “WP_014407100.1” (non-redundant RefSeq protein ID) and “SGL_RS08200” (RefSeq locus tag ID), “BAA17799.2” (GenBank ID), “KHHDGJBG_01286” (Prodigal *ab initio* prediction) for the gene *psbC* (*sll0851*).

Throughout the manuscript we use terms in the following hierarchy: **(i)** Widely accepted functional gene designations if available (e.g., *psbC, psbN*); **(ii)** *sll/slr* and *ssl/ssr* IDs for the reverse (“left) and forward (“right”) orientation, as used in the community since publication of the genome sequence^5^; **(iii)** IDs of ncRNAs (*ncl/ncr*) following the logic as described under (ii), as defined by extensive transcriptome analyses^54^ if a translon was identified within one of these previously defined ncRNAs (e.g., *ncr1470*). We added “sORF” and a number if more than one translon was found in an ncRNA (e.g., *ncr1610-sORF1* and *ncr1610-sORF2)*; **(iv)** Instances related to annotated features, e.g. translons within asRNAs or translons IOF and IIF were connected to these annotations, for example infC-IOF, or asPsbC; **(v)** if any of the instances (ii) to (iv), as well as a few additional existing IDs were renamed due to a functional analysis, we follow the rule as described under (i) (e.g., *ncr1071* became *nirP1*^46^).

### Genetic Engineering and protein verification

Genes of interest were cloned into the self-replicating expression plasmid pVZ322 flanked by the copper-inducible P*_petE_* promoter at the 5’ end and a coding segment for either a 3xFLAG tag (DYKDHDGDYKDHDIDYKDDDDK) or a SPA tag^116^. This was started by subcloning the coding sequences into plasmid pUC19. Therefore, a part of pUC19 (primers P11-45 + P11-46; **Table S6**) was amplified to generate the backbone. For asPsbC inserts, the P*_petE_* promoter and the *asPsbC* coding sequence were amplified from *Synechocystis* 6803 genomic DNA using primers G3-64 + VR278 and VR279 + VR280. The SPA tag, STOP codon, *ncl1450* 3’UTR and the *oop* terminator sequences were PCR-amplified from plasmid pVZ322::P*_petE_*_*ncr1470*_SPA using primers VR281 + G3-65. For asCsx18 inserts, the P*_petE_* promoter and *asCsx18* coding sequence were amplified using primers G3-64 + VR285 and VR286 + VR287, while the SPA tag, STOP codon, *ncl1450* 3’UTR and *oop* terminator sequences were amplified from plasmid pVZ322_P*_petE_*::*ncr1470*-SPA using primers VR288 + G3-65. The PCR-amplified pUC19 backbone and the respective insert fragments each were assembled using AQUA cloning^117^ and subcloned into *E. coli* Top10F’ (Thermo Fisher Scientific). The isolated plasmids were used as a PCR template with primers VR277 + VR282 to amplify the fragment P*_petE_*_*asPsbC* or *asCsx18*_SPA_STOP_*ncl1450* 3’UTR_*oop* with overhangs to the self-replicating expression plasmid pVZ322. The vector was therefore cut with *Sal*I and *Pst*I (NEB) and treated with FAST AP (Thermo Fisher). The linearized plasmid and the insert fragments were assembled and subcloned as before. All other genes of interest were cloned following the same strategy but using the respective primers listed in **Table S6.** All PCR reactions were performed with PCRBIO HiFi Polymerase (PCR Biosystems). All PCRs using a plasmid as template were treated with FastDigest *Dpn*I (Thermo Fisher Scientific). All PCR products were gel-purified (NucleoSpin Gel and PCR Clean-up, Macherey-Nagel). Sanger Sequencing Economy Run (Microsynth) was used to verify the correct sequence of all plasmids. The constructs were then conjugated into the *Synechocystis* 6803 wild type^70^. Conjugated clones were selected with 50 µg/mL kanamycin and 5 µg/mL gentamicin in BG11 medium^93^, grown to exponential phase (OD_750_ ∼ 0.6-0.8) and protein expression was stimulated by the addition of 1.25 or 2 µM CuSO_4_. For candidates that lacked Western blot signals, we added Bortezomib (20 µM) during the expression phase to inhibit the Clp protease system as this was previously introduced to improve SEP validation rates^40^. After 2 h of incubation, induced and control wild type cells were harvested by centrifugation (4000 × g, 4°C, 10 min) and lysed in phosphate buffered saline (137 mM sodium chloride, 2.7 mM potassium chloride, 10 mM disodium phosphate, 1.8 mM potassium dihydrogen phosphate, pH 7.4) supplemented with cOmplete protease inhibitor cocktail (Roche). Cell disruption was performed mechanically in a prechilled Precellys 24 (Bertin Instruments) with three rounds of shaking at 6,000 rpm (10 s) followed by a pause (5 s) per cycle, for a total of 5 cycles and using glass beads with a 0.1–0.25 mm diameter. Glass beads were removed by centrifugation (1,000 x g, 1 min, 4°C). Obtained cell lysates were mixed with denaturing and reducing loading dye (5× concentrated: 250 mM Tris-HCl pH 6.8, 25% glycerol, 10% SDS, 500 mM DTT, and 0.05% bromophenol blue G-250). The samples were subsequently separated by SDS-Tricine-PAGE with Precision Plus Protein Dual Xtra molecular weight marker (Bio-Rad) as standard followed by western blotting^13^.

Several vectors were constructed for the expression of the different versions of Slr0489 and Slr1079, including the presence or absence of a sequence encoding a C-terminal 3xFLAG tag. The genes *slr0489* and *slr1079* were amplified from *Synechocystis* 6803 genomic DNA using primer pairs RB1 + RB2 and RB3 + RB4, respectively. For the nested genes, *slr0489S* and *slr1079S*, we also amplified the 5’ UTR with primer pairs RB5 + RB6 and RB7 +RB8, respectively. The amplicons were cloned into the reverse-amplified pUC19P_rha_YFP3×FLAG vector with or without the FLAG tag^118^. For the wild-type version, pUC19P_rha_YFP3×FLAG was reverse amplified with RB9 + RB10 for the tagged version, or RB10 + RB11 for the untagged version. Similarly, *slr0489S* and *slr1079S* were amplified with RB12 and RB9 or RB11 for the untagged version. The inserts of interest were amplified and cloned into *Xmn*I-digested pVZ322s^119^. For the co-immunoprecipitation assay, the 3xFLAG constructs were transformed into wild-type *Synechocystis* 6803 by electroporation as described^46^.

To assess possible physiological effects of expressing either the short or the long Slr0489 version on *Synechocystis*, first the native version, encoding both the large and the small protein, was amplified with primer pairs RB13 + RB14 from a previous construct. The fragment was cloned into a shorter version of a modified pACYC-Duet-1 vector (with Km^R^ instead of Cm^R^), after amplification with primers RB15 + RB16. From this new construct (pACYC-*slr0489*WT-6×His), a single point mutation was inserted into *slr0489*WT to change the *slr0489S* start codon into a threonine codon, creating a construct expressing only the long protein (Slr0489L). The *slr0489* gene construct was reverse amplified with primers RB17 + RB18 and RB18, carrying 5’overhangs. For expression in *Synechocystis*, *slr0489*L was amplified with primers RB1 + RB19 from pACYC-slr0489L to generate overhangs to the linearized pUC19-Rha-YFP-3×FLAG vector and later cloned into pVZ322s^118^ without any tag sequence. The final pVZ322s-P*rha*-*slr0489L* was then conjugated into *Synechocystis* Δ*slr0489* via tri-parental mating. The constructs pVZ322s-P*rha*-*slr0489* and pVZ322s-P*rha*-*slr0489S* were also conjugated into Δ*slr0489*. The strains were then cultivated in BG11 under standard conditions. Expression was induced with 0.6 mg/mL L-rhamnose; uninduced cultures served as control. Growth was monitored by following the OD_750_ of 150 µL samples in a 96-well plate for 5 d. Each strain was cultivated in biological duplicates for each condition.

The Δ*slr0489* deletion mutant was generated by replacing *slr0489* by homologous recombination with a streptomycin resistance cassette (*aadA*) amplified from plasmid pRL692, and using pUC19 as a vector for subcloning. For this, flanking regions present up– and downstream of *slr0489* were amplified from *Synechocystis* 6803 genomic DNA using primer pairs 117/118 and 123/124. Transcription of *slr0489* occurs from a transcriptional start site at position 2,622,526 on the fw strand matching the first nt of the start codon (leaderless expression), but leading to a joint transcriptional unit TU2770 with the downstream genes *slr0491–slr0492*^54^. To allow expression of the downstream-located genes, the promoter region was amplified separately using primers121/122 and inserted at the end of the *aadA* resistance cassette. Note, that the *slr0489* 3′ end and the *slr0491* 5′ end overlap by 15 nt, which were kept intact behind the inserted *aadA-*promoter construct. The corresponding plasmid was used to transform *Synechocystis* 6803 cells by electroporation according to Kraus et al.^46^. Transformants were selected on 20 µg/mL streptomycin. The segregation of Δ*slr0489*/*aadA* from wild type alleles was followed by colony PCR using the primers 133/134.

### Separation of soluble and insoluble fractions

For localization of the Ncl1450 protein, wild type *Synechocystis* 6803, P*_petE_*::*ncl1450*-3×FLAG and P*_ncl1450_*::*ncl1450*-3×FLAG ectopic overexpression mutants were grown until exponential phase, then the first sample was taken. The P*_petE_* promoter was induced by 2 μM CuSO_4_. Carbon-depletion for the native promoter was induced by washing the pellet three times in carbon-depleted BG11 medium. 24 hours after induction, samples were taken and centrifuged (4,000 × g, 10 min, 4°C) to collect cells. The pellet was resuspended in membrane buffer (25 mM HEPES-NaOH, pH 7.0, 15 mM CaCl_2_, 5 mM MgCl_2_, 15% v/v glycerol) with protease inhibitor and then frozen in liquid nitrogen and stored at –80°C. Cells were disrupted in a Precellys homogenizer as described above. Whole cell lysate was centrifuged (21,000 × g, 20 min, 4°C) for separation of soluble and insoluble fractions, then supernatant (soluble fraction) and pellet were separated. The pellet was resuspended in membrane buffer. SDS-PAGE and anti-FLAG western blot were performed as described^13^.

### Co-IP, LC-MS/MS analyses and data processing of nested gene products

Tagged proteins (Slr0489L-3xFLAG; Slr0489S-3xFLAG; Slr1079L-3xFLAG; Slr1079S-3xFLAG; sfGFP-3xFLAG) were expressed in *Synechocystis* 6803 from plasmid pVZ322s^119^ under control of the P_rha_ promoter^120^ after induction by adding 0.6 mg/mL L-rhamnose at an OD_750_ of 0.5. After 24 h, cells from 50 mL batch cultures were harvested by centrifugation (3,750 × *g*, 4°C, 20 min), resuspended in 700 µL FLAG buffer (50 mM HEPES-NaOH pH 7; 5 mM MgCl_2_; 25 mM CaCl_2_, 150 mM NaCl; 10% glycerol; 0.1% Tween-20) supplemented with Protease Inhibitor (cOmplete, Roche) and mixed with 100 μL of glass beads (Ø 0.1–0.25 mm, Retsch). Cell disruption was performed in a prechilled Precellys 24. Crude cell debris and glass beads were removed by centrifugation (1,000 × *g*, 1 min, 4°C) and supernatants were subsequently incubated at gentle shaking in presence of 2% w/v n-dodecyl-ß –D-maltoside for 60 min at 4°C in the dark. Remaining cell debris was removed by centrifugation (21,300 × *g*, 4°C, 30 min). The cleared total cell lysate was subjected to co-IP using anti-FLAG M2 affinity magnetic beads (Sigma–Aldrich) rotating for 2 h at 4°C. The beads were washed 5 times with 600 µL FLAG buffer supplemented with protease inhibitors for 2 min, 4°C, rotating. Bound proteins were eluted by incubation with 150 µL FLAG peptide (Sigma–Aldrich, 100 ng/µL) in TBS buffer (150 mM NaCl in 20 mM Tris/HCl, pH 7.6) for 30 min at 4°C in two subsequent steps and subsequently combined. Three independent replicates were prepared for each Co-IP. Fractions (20 µL) of each eluate protein were analyzed by SDS-PAGE and anti-FLAG immunoblotting for quality control. Remaining eluates were lyophilized and resolubilized in denaturation buffer (6 M urea, 2 M thiourea in 100 mM Tris/HCl; pH 8.0) for LC-MS/MS analysis. Protein disulfide bonds were reduced with dithiothreitol at a final concentration of 1 mM for 45 min and the resulting thiol groups were alkylated with iodoacetamide at a final concentration of 5.5 mM for 45 min in the dark. Proteins were pre-digested with Lys-C (Santa Cruz Biotechnology) for 3 h, diluted with 4 volumes of 20 mM ammonium bicarbonate buffer, pH 8, and overnight digested with trypsin (sequencing grade, Promega) at room temperature with a protease/protein ratio of 1/100 for both proteases. Resulting peptide solutions were acidified with trifluoroacetic acid to pH 2.5, and purified by Stage Tips^121^. For LC-MS/MS-based protein analysis, 1/20 of each sample was loaded onto an in-house-packed reverse-phase (RP) column (20 cm; 75 µm ID, packed with ReproSil-Pur 1.9 µm C18 from Dr. Maisch, Germany) and separated by RP chromatography on an EASY-nLC 1200 (Thermo Fisher Scientific, USA) using linear 36 min gradients. Eluting peptides were directly ionized by electro spray ionization and analyzed on a Q-Exactive HF mass spectrometer (Thermo Fisher Scientific, USA) in the data dependent acquisition mode. MS1 resolution was set to 60,000 at a scan range of 300 – 1650 m/z. The 7 most abundant ions with charge states 2-5 were subjected to fragmentation by higher-energy collision induced fragmentation. Fragment ions were analyzed at an MS2 resolution of 60,000 and a scan range of 200-2000 m/z. Dynamic exclusion of previously sequenced ions was set to 30 s.

Raw spectra of each Co-IP experiment (Slr0489L; Slr0489S; Slr1079L and Slr1079S) were analyzed separately including all replicates and processed with the common control (sfGFP-FLAG) with the MaxQuant software (version 1.6.8.0)^122^ at default settings. Label-free quantification (LFQ), matching between runs and iBAQ options were enabled. Peak lists were searched against Co-IP specific modified target-decoy database of *Synechocystis* 6803^48^ with 3,959 protein entries, including the tagged bait protein sequences truncated for the indifferentiable 3xFLAG sequence. False discovery rates were limited to 1% at peptide and protein levels. Missing LFQ protein intensities were imputed to increase number of quantifiable proteins. Therefore, random values the low intensity range were given in case of missing values for each replicate separately within a width of 10% of the standard deviation of the data and a downshift corresponding to the 2-fold standard deviation using the Perseus software suite (version 1.6.5.0)^123^ (**Figs. S23–S26**). Co-IP enriched proteins were analyzed by Student’s *t*-test relative to the control (sfGFP-FLAG IP) with a permutation-based FDR of 0.05 (**Tables S8–S11**).

## DATA AVAILABILITY

Ribo-seq, TIS and TTS profiling datasets were deposited in the Gene Expression Omnibus (GEO) Database, with accession number GSE246336 (direct access: https://www.ncbi.nlm.nih.gov/geo/query/acc.cgi?acc=GSE246336). The mass spectrometry proteomics data generated in this study have been deposited at the ProteomeXchange Consortium via the PRIDE^124^ partner repository with the dataset identifier PXD060427. The datasets described in this study can be viewed with an interactive online JBrowse 2 instance at https://www.bioinf.uni-freiburg.de/∼ribobase/.

## CODE AVAILABILITY

Repository for Metagene Profiling Scripts: https://github.com/RickGelhausen/Synechocystis_pub_scripts_2025.

Repository for ORFBounder: https://github.com/RickGelhausen/ORFBounder.

## Supporting information

Table S12. Extrapolation of the number of expected true-positive translons based on the manual inspection of 808 candidates

## ACKNOWLEDGEMENTS

This work was supported by the German Research Foundation (DFG) priority program SPP2002 “Small Proteins in Prokaryotes, an Unexplored World” (grants HE 2544/12-1/2 to WRH, SH 580/7-1/2 to CMS, BA 2168/21-1/2 to ROB), by grants HE 2544/22-1 to WRH and MA 4918/4-1/2 to BM and by Germany’s DFG-funded Excellence Strategy (CIBSS – EXC-2189 – Project ID 390939984) to ROB. CHA acknowledges funding from the Swiss National Science Foundation for BH and GJ (grant 197391). The computational resources were supported by the de.NBI Cloud within the German Network for Bioinformatics Infrastructure (de.NBI) and ELIXIR-DE (Forschungszentrum Jülich and W-de.NBI-001, W-de.NBI-004, W-de.NBI-008, W-de.NBI-010, W-de.NBI-013, W-de.NBI-014, W-de.NBI-016, W-de.NBI-022). We thank Ann-Janine Imsiecke, Stephanie Färber and Philipp Kible (Würzburg, Germany) for the technical assistance in establishing the Ribo-seq workflow for *Synechocystis* 6803, Paulo Oliveira (Porto, Portugal) for the *tolC* transporter mutants and Joëlle Sasse Schläpfer (Zurich, Switzerland) for critical reading.

## AUTHOR CONTRIBUTIONS

WRH, CMS and ROB designed and supervised the Ribo-seq part of the project and secured funding. CHA supervised the proteogenomics part of the project. VK and PM constructed and cultivated strains. LH and VK designed and established Ribo-seq and TIS/TTS methods for *Synechocystis* 6803. RG performed bioinformatic processing of the Ribo-seq, TIS and TTS data, developed custom analysis tools and scripts, and created a JBrowse 2 instance for the project. VK, LH and WRH performed the manual analysis of Ribo-seq, TIS and TTS data including the curation of gene annotations. PS and BM performed mass spectrometry, BH, GJ and CHA performed the proteogenomic analysis to identify additional translons and proteoforms including iPtgxDB-based protein searches informed by Ribo-seq and TIS/TTS data, and contributed the generic approach for genome annotation consolidation. VK, VR and MG constructed strains and performed western blot analyses. RAB performed cultivation experiments with *slr0489* and *slr1079*. VK, LH, CHA and WRH wrote the manuscript with input from all authors. All authors read and agreed on the manuscript.

## REFERENCES

1. Flombaum, P. et al. Present and future global distributions of the marine Cyanobacteria *Prochlorococcus* and *Synechococcus*. Proc. Natl. Acad. Sci. USA 110, 9824–9829 (2013).

2. Hagemann, M. & Hess, W. R. Systems and synthetic biology for the biotechnological application of cyanobacteria. Curr Opin Biotechnol 49, 94–99 (2018).

3. Vijay, D., Akhtar, M. K. & Hess, W. R. Genetic and metabolic advances in the engineering of cyanobacteria. Curr. Opin. Biotechnol. 59, 150–156 (2019).

4. Klähn, S., Opel, F. & Hess, W. R. Customized molecular tools to strengthen metabolic engineering of cyanobacteria. Green Carbon 2, 149–163 (2024).

5. Kaneko, T. et al. Sequence analysis of the genome of the unicellular cyanobacterium Synechocystis sp. strain PCC6803. II. Sequence determination of the entire genome and assignment of potential protein-coding regions. DNA Res. 3, 109–136 (1996).

6. Zhang, J. Protein-length distributions for the three domains of life. Trends Genet. TIG 16, 107–109 (2000).

7. Ahrens, C. H., Wade, J. T., Champion, M. M. & Langer, J. D. A practical guide to small protein discovery and characterization using mass spectrometry. J. Bacteriol. 204, e0035321 (2022).

8. Burton, A. T., Zeinert, R. & Storz, G. Large roles of small proteins. Annu. Rev. Microbiol. 78, 1–22 (2024).

9. Alvarenga-Lucius, L. V. et al. The high-light-induced protein SliP4 binds to NDH1 and photosystems facilitating cyclic electron transport and state transition in Synechocystis sp. PCC 6803. New Phytol. 239, 1083–1097 (2023).

10. Komenda, J. & Sobotka, R. Cyanobacterial high-light-inducible proteins--Protectors of chlorophyll-protein synthesis and assembly. Biochim. Biophys. Acta 1857, 288–295 (2016).

11. Fromme, P., Melkozernov, A., Jordan, P. & Krauss, N. Structure and function of photosystem I: interaction with its soluble electron carriers and external antenna systems. FEBS Lett. 555, 40–44 (2003).

12. Xu, W., Tang, H., Wang, Y. & Chitnis, P. R. Proteins of the cyanobacterial photosystem I. Biochim. Biophys. Acta BBA – Bioenerg. 1507, 32–40 (2001).

13. Baumgartner, D., Kopf, M., Klähn, S., Steglich, C. & Hess, W. R. Small proteins in cyanobacteria provide a paradigm for the functional analysis of the bacterial micro-proteome. BMC Microbiol. 16, 285 (2016).

14. Brandenburg, F. & Klähn, S. Small but smart: On the diverse role of small proteins in the regulation of cyanobacterial metabolism. Life 10, 322 (2020).

15. Gray, T., Storz, G. & Papenfort, K. Small proteins; Big questions. J. Bacteriol. 204, e0034121 (2022).

16. Fesenko, I. et al. The hidden bacterial microproteome. Mol. Cell 85, 1024–1041 (2025).

17. Song, K. et al. AtpΘ is an inhibitor of F0F1 ATP synthase to arrest ATP hydrolysis during low-energy conditions in cyanobacteria. Curr. Biol. 32, 136–148.e5 (2022).

18. Jurënas, D., Fraikin, N., Goormaghtigh, F. & Van Melderen, L. Biology and evolution of bacterial toxin–antitoxin systems. Nat. Rev. Microbiol. 20, 335–350 (2022).

19. Santos-Júnior, C. D. et al. Discovery of antimicrobial peptides in the global microbiome with machine learning. Cell 187, 3761–3778.e16 (2024).

20. Ingolia, N. T., Ghaemmaghami, S., Newman, J. R. S. & Weissman, J. S. Genome-wide analysis in vivo of translation with nucleotide resolution using ribosome profiling. Science 324, 218–223 (2009).

21. Ingolia, N. T., Hussmann, J. A. & Weissman, J. S. Ribosome profiling: global views of translation. Cold Spring Harb. Perspect. Biol. 11, a032698 (2019).

22. Mohammad, F., Green, R. & Buskirk, A. R. A systematically-revised ribosome profiling method for bacteria reveals pauses at single-codon resolution. eLife 8, e42591 (2019).

23. Gerashchenko, M. V., Lobanov, A. V. & Gladyshev, V. N. Genome-wide ribosome profiling reveals complex translational regulation in response to oxidative stress. Proc. Natl. Acad. Sci. USA 109, 17394–17399 (2012).

24. Zupanic, A. et al. Detecting translational regulation by change point analysis of ribosome profiling data sets. RNA 20, 1507–1518 (2014).

25. Chotewutmontri, P. & Barkan, A. Ribosome profiling elucidates differential gene expression in bundle sheath and mesophyll cells in maize. Plant Physiol. 187, 59–72 (2021).

26. Xiao, Z. et al. De novo annotation and characterization of the translatome with ribosome profiling data. Nucleic Acids Res. 46, e61 (2018).

27. Ingolia, N. T., Lareau, L. F. & Weissman, J. S. Ribosome profiling of mouse embryonic stem cells reveals the complexity and dynamics of mammalian proteomes. Cell 147, 789–802 (2011).

28. Bazzini, A. A. et al. Identification of small ORFs in vertebrates using ribosome footprinting and evolutionary conservation. EMBO J. 33, 981–993 (2014).

29. Hsu, P. Y. et al. Super-resolution ribosome profiling reveals unannotated translation events in *Arabidopsis*. Proc. Natl. Acad. Sci. USA 113, E7126–E7135 (2016).

30. Meydan, S. et al. Retapamulin-assisted ribosome profiling reveals the alternative bacterial proteome. Mol. Cell 74, 481–493.e6 (2019).

31. Orr, M. W., Mao, Y., Storz, G. & Qian, S.-B. Alternative ORFs and small ORFs: shedding light on the dark proteome. Nucleic Acids Res. 48, 1029–1042 (2020).

32. Froschauer, K. et al. Complementary Ribo-seq approaches map the translatome and provide a small protein census in the foodborne pathogen *Campylobacter jejuni*. Nat. Commun. 16, 3078 (2025).

33. Weingarten-Gabbay, S. et al. Pan-viral ORFs discovery using massively parallel ribosome profiling. Science 388, 1218–1224 (2025).

34. Tufail, M. A. et al. Uncovering the small proteome of *Methanosarcina mazei* using Ribo-seq and peptidomics under different nitrogen conditions. Nat. Commun. 15, 8659 (2024).

35. Hadjeras, L. et al. Unraveling the small proteome of the plant symbiont *Sinorhizobium meliloti* by ribosome profiling and proteogenomics. microLife 4, uqad012 (2023).

36. Hadjeras, L. et al. Revealing the small proteome of *Haloferax volcanii* by combining ribosome profiling and small-protein optimized mass spectrometry. microLife 4, uqad001 (2023).

37. Weaver, J., Mohammad, F., Buskirk, A. R. & Storz, G. Identifying small proteins by ribosome profiling with stalled initiation complexes. mBio 10, (2019).

38. Florin, T. et al. An antimicrobial peptide that inhibits translation by trapping release factors on the ribosome. Nat. Struct. Mol. Biol. 24, 752–757 (2017).

39. Mangano, K. et al. Genome-wide effects of the antimicrobial peptide apidaecin on translation termination in bacteria. eLife 9, e62655 (2020).

40. Stringer, A., Smith, C., Mangano, K. & Wade, J. T. Identification of novel translated small open reading frames in *Escherichia coli* using complementary ribosome profiling approaches. J. Bacteriol. 204, e00352–21 (2022).

41. Vazquez-Laslop, N., Sharma, C. M., Mankin, A. & Buskirk, A. R. Identifying small open reading frames in prokaryotes with ribosome profiling. J. Bacteriol. 204, e00294–21 (2020).

42. Karlsen, J., Asplund-Samuelsson, J., Thomas, Q., Jahn, M. & Hudson, E. P. Ribosome profiling of *Synechocystis* reveals altered ribosome allocation at carbon starvation. Msystems 3, e00126–18 (2018).

43. Karlsen, J., Asplund-Samuelsson, J., Jahn, M., Vitay, D. & Hudson, E. P. Slow protein turnover explains limited protein-level response to diurnal transcriptional oscillations in cyanobacteria. Front. Microbiol. 12, 657379 (2021).

44. de Alvarenga, L. V., Hess, W. R., Hagemann, M. & Hagemann, M. AcnSP – a novel small protein regulator of aconitase activity in the cyanobacterium Synechocystis sp. PCC 6803. Front. Microbiol. 11, 1445 (2020).

45. Krauspe, V. et al. Discovery of a novel small protein factor involved in the coordinated degradation of phycobilisomes in cyanobacteria. Proc. Natl. Acad. Sci. USA 118, e2012277118 (2021).

46. Kraus, A. et al. Protein NirP1 regulates nitrite reductase and nitrite excretion in cyanobacteria. Nat. Commun. 15, 1911 (2024).

47. Œwirski, M. I., et al. Translon: a single term for translated regions. Nat. Methods 22, 2002–2006 (2025).

48. Spät, P., Krauspe, V., Hess, W. R., Maček, B. & Nalpas, N. Deep proteogenomics of a photosynthetic cyanobacterium. J. Proteome Res. 22, 1969–19 (2023).

49. Omasits, U. et al. An integrative strategy to identify the entire protein coding potential of prokaryotic genomes by proteogenomics. Genome Res. 27, 2083–2095 (2017).

50. Chen, Q. et al. Deletion of *sll1541* in *Synechocystis* sp. strain PCC 6803 allows formation of a far-red-shifted holo-proteorhodopsin in vivo. Appl. Environ. Microbiol. 84, e02435–17 (2018).

51. Kopfmann, S., Roesch, S. K. & Hess, W. R. Type II toxin-antitoxin systems in the unicellular cyanobacterium Synechocystis sp. PCC 6803. Toxins 8, 228.1–228.23 (2016).

52. Gelhausen, R. et al. HRIBO: high-throughput analysis of bacterial ribosome profiling data. Bioinformatics 37, 2061–2063 (2021).

53. Gelsinger, D. R. et al. Ribosome profiling in archaea reveals leaderless translation, novel translational initiation sites, and ribosome pausing at single codon resolution. Nucleic Acids Res. 48, 5201–5216 (2020).

54. Kopf, M. et al. Comparative analysis of the primary transcriptome of Synechocystis sp. PCC 6803. DNA Res. 21, 527–539 (2014).

55. Willems, P., Fijalkowski, I. & Van Damme, P. Lost and found: re-searching and re-scoring proteomics data aids genome annotation and improves proteome coverage. mSystems 5, e00833–20 (2020).

56. Hyatt, D. et al. Prodigal: prokaryotic gene recognition and translation initiation site identification. BMC Bioinformatics 11, 119 (2010).

57. Mishra, A., Siwach, P., Singhal, P. & Jayaram, B. ChemGenome2.1: An ab initio gene prediction software. in Gene Prediction: Methods and Protocols (ed. Kollmar, M.) 121–138 (Springer, New York, NY, 2019). doi:10.1007/978-1-4939-9173-0_7.

58. Hecht, A. et al. Measurements of translation initiation from all 64 codons in *E. coli*. Nucleic Acids Res. 45, 3615–3626 (2017).

59. Baers, L. L. et al. Proteome mapping of a cyanobacterium reveals distinct compartment organization and cell-dispersed metabolism. Plant Physiol. 181, 1721–1738 (2019).

60. Craigen, W. J., Cook, R. G., Tate, W. P. & Caskey, C. T. Bacterial peptide chain release factors: conserved primary structure and possible frameshift regulation of release factor 2. Proc. Natl. Acad. Sci. USA 82, 3616–3620 (1985).

61. Baranov, P. V., Gesteland, R. F. & Atkins, J. F. Release factor 2 frameshifting sites in different bacteria. EMBO Rep. 3, 373–377 (2002).

62. Tate, W. P. et al. Translational termination efficiency in both bacteria and mammals is regulated by the base following the stop codon. Biochem. Cell Biol. 73, 1095–1103 (1995).

63. Sesto, N., Wurtzel, O., Archambaud, C., Sorek, R. & Cossart, P. The excludon: a new concept in bacterial antisense RNA-mediated gene regulation. Nat. Rev. Microbiol. 11, 75–82 (2013).

64. Hallgren, J. et al. DeepTMHMM predicts alpha and beta transmembrane proteins using deep neural networks. 2022.04.08.487609 Preprint at 10.1101/2022.04.08.487609 (2022).

65. Shore, S. F. H., Leinberger, F. H., Fozo, E. M. & Berghoff, B. A. Type I toxin-antitoxin systems in bacteria: from regulation to biological functions. EcoSal Plus 12, eesp-0025-2022 (2024).

66. van Kempen, M. et al. Fast and accurate protein structure search with Foldseek. Nat. Biotechnol. 42, 243–246 (2024).

67. Yu, V., Ronzone, E., Lord, D., Peti, W. & Page, R. MqsR is a noncanonical microbial RNase toxin that is inhibited by antitoxin MqsA via steric blockage of substrate binding. J. Biol. Chem. 298, 102535 (2022).

68. Mitschke, J. et al. An experimentally anchored map of transcriptional start sites in the model cyanobacterium Synechocystis sp. PCC6803. Proc. Natl. Acad. Sci. USA 108, 2124–2129 (2011).

69. Kopf, M. et al. Comparative genome analysis of the closely related *Synechocystis* strains PCC 6714 and PCC 6803. DNA Res. 21, 255–266 (2014).

70. Scholz, I., Lange, S. J., Hein, S., Hess, W. R. & Backofen, R. CRISPR-Cas systems in the cyanobacterium Synechocystis sp. PCC6803 exhibit distinct processing pathways involving at least two Cas6 and a Cmr2 protein. PLoS ONE 8, e56470 (2013).

71. Shah, S. A. et al. Comprehensive search for accessory proteins encoded with archaeal and bacterial type III CRISPR-cas gene cassettes reveals 39 new cas gene families. RNA Biol. 16, 530–542 (2019).

72. Susanto, F. A., Jones, A. D., Donnelly, S. L. & Lundquist, P. K. Cyanoglobule lipid droplets are a stress-responsive metabolic compartment of cyanobacteria and the progenitor of plant plastoglobules. Plant Cell 37, koaf127 (2025).

73. Kopf, M. & Hess, W. R. Regulatory RNAs in photosynthetic cyanobacteria. FEMS Microbiol. Rev. 39, 301–315 (2015).

74. Huckauf, J., Nomura, C., Forchhammer, K. & Hagemann, M. Stress responses of *Synechocystis* sp. strain PCC 6803 mutants impaired in genes encoding putative alternative sigma factors. Microbiology 146, 2877–2889 (2000).

75. Trautmann, D., Voß, B., Wilde, A., Al-Babili, S. & Hess, W. R. Microevolution in cyanobacteria: re-sequencing a motile substrain of Synechocystis sp. PCC 6803. DNA Res. 19, 435–448 (2012).

76. Katoh, A., Sonoda, M. & Ogawa, T. A possible role of 154-base pair nucleotides located upstream of ORF440 on CO2 transport of Synechocystis sp. PCC 6803. in Photosynthesis: From Light to Biosphere vol. 3 481–484 (Kluwer Academic Publishers, Dordrecht, The Netherlands, 1995).

77. Kamei, A., Ogawa, T. & Ikeuchi, M. Identification of a novel gene (slr2031) involved in high-light resistance in the cyanobacterium Synechocystis sp. PCC 6803. in Photosynthesis: Mechanisms and Effects: Volume I–V: Proceedings of the XIth International Congress on Photosynthesis, Budapest, Hungary, August 17–22, 1998 (ed. Garab, G.) 2901–2904 (Springer Netherlands, Dordrecht, 1998). doi:10.1007/978-94-011-3953-3_680.

78. Bouillet, S., Arabet, D., Jourlin-Castelli, C., Méjean, V. & Iobbi-Nivol, C. Regulation of σ factors by conserved partner switches controlled by divergent signalling systems. Environ. Microbiol. Rep. 10, 127–139 (2018).

79. Behler, J. et al. The host-encoded RNase E endonuclease as the crRNA maturation enzyme in a CRISPR–Cas subtype III-Bv system. Nat. Microbiol. 3, 367–377 (2018).

80. Oeser, S. et al. Minor pilins are involved in motility and natural competence in the cyanobacterium Synechocystis sp. PCC 6803. Mol. Microbiol. 116, 743–765 (2021).

81. Fijalkowski, I., Snauwaert, V. & Van Damme, P. Proteins à la carte: riboproteogenomic exploration of bacterial N-terminal proteoform expression. mBio 15, e0033324 (2024).

82. Kaushal, P. & Lee, C. N-terminomics – its past and recent advancements. J. Proteomics 233, 104089 (2021).

83. Diesh, C. et al. JBrowse 2: a modular genome browser with views of synteny and structural variation. Genome Biol. 24, 74 (2023).

84. Hemm, M. R., Weaver, J. & Storz, G. *Escherichia coli* small proteome. EcoSal Plus 9, (2020).

85. Jurėnas, D., Fraikin, N., Goormaghtigh, F. & Van Melderen, L. Biology and evolution of bacterial toxin–antitoxin systems. Nat. Rev. Microbiol. 20, 335–350 (2022).

86. Sailer, A.-L. et al. Internal in-frame translation generates Cas11b, which is important for effective interference in an archaeal CRISPR-Cas system. Front. Microbiol. 16, 1543464 (2025).

87. McBride, T. M. et al. Diverse CRISPR-Cas complexes require independent translation of small and large subunits from a single gene. Mol. Cell 80, 971–979 (2020).

88. Kongari, R. et al. Phage spanins: diversity, topological dynamics and gene convergence. BMC Bioinformatics 19, 326 (2018).

89. Cahill, J. et al. Genetic analysis of the lambda spanins Rz and Rz1: identification of functional domains. G3 GenesGenomesGenetics 7, 741–753 (2017).

90. Doherty, E. E. et al. A miniature CRISPR–Cas10 enzyme confers immunity by inhibitory signalling. Nature 647, 997–1004 (2025).

91. Sullivan, A. E. et al. The Panoptes system uses decoy cyclic nucleotides to defend against phage. Nature 647, 988–996 (2025).

92. Thomas, J.-C., Ughy, B., Lagoutte, B. & Ajlani, G. A second isoform of the ferredoxin:NADP oxidoreductase generated by an in-frame initiation of translation. Proc. Natl. Acad. Sci. USA 103, 18368–18373 (2006).

93. Rippka, R., Deruelles, J., Waterbury, J. B., Herdman, M. & Stanier, R. Y. Generic assignments, strain histories and properties of pure cultures of cyanobacteria. Microbiology 111, 1–61 (1979).

94. Oliveira, P. et al. The versatile TolC-like Slr1270 in the cyanobacterium Synechocystis sp. PCC 6803. Environ. Microbiol. 18, 486–502 (2016).

95. Mattiuzzo, M. et al. Role of the *Escherichia coli* SbmA in the antimicrobial activity of proline-rich peptides. Mol. Microbiol. 66, 151–163 (2007).

96. Sharma, C. M., Darfeuille, F., Plantinga, T. H. & Vogel, J. A small RNA regulates multiple ABC transporter mRNAs by targeting C/A-rich elements inside and upstream of ribosome-binding sites. Genes Dev. 21, 2804–2817 (2007).

97. Ingolia, N. T., Brar, G. A., Rouskin, S., McGeachy, A. M. & Weissman, J. S. The ribosome profiling strategy for monitoring translation in vivo by deep sequencing of ribosome-protected mRNA fragments. Nat. Protoc. 7, 1534–1550 (2012).

98. Venturini, E. et al. A global data-driven census of Salmonella small proteins and their potential functions in bacterial virulence. microLife 1, uqaa002 (2020).

99. Köster, J. & Rahmann, S. Snakemake-a scalable bioinformatics workflow engine. Bioinforma. Oxf. Engl. 34, 3600 (2018).

100. Grüning, B. et al. Bioconda: sustainable and comprehensive software distribution for the life sciences. Nat. Methods 15, 475–476 (2018).

101. Martin, M. Cutadapt removes adapter sequences from high-throughput sequencing reads. EMBnet.journal 17, 10 (2011).

102. Otto, C., Stadler, P. F. & Hoffmann, S. Lacking alignments? The next-generation sequencing mapper segemehl revisited. Bioinforma. Oxf. Engl. 30, 1837–1843 (2014).

103. Danecek, P. et al. Twelve years of SAMtools and BCFtools. GigaScience 10, giab008 (2021).

104. Liao, Y., Smyth, G. K. & Shi, W. featureCounts: an efficient general purpose program for assigning sequence reads to genomic features. Bioinformatics 30, 923–930 (2014).

105. Wingett, S. W. & Andrews, S. FastQ Screen: A tool for multi-genome mapping and quality control. F1000Research 7, 1338 (2018).

106. Ewels, P., Magnusson, M., Lundin, S. & Käller, M. MultiQC: summarize analysis results for multiple tools and samples in a single report. Bioinformatics 32, 3047–3048 (2016).

107. Crooks, G. E., Hon, G., Chandonia, J.-M. & Brenner, S. E. WebLogo: A Sequence Logo Generator. Genome Res. 14, 1188–1190 (2004).

108. Ndah, E. et al. REPARATION: ribosome profiling assisted (re-) annotation of bacterial genomes. Nucleic Acids Res. 45, e168–e168 (2017).

109. Clauwaert, J., Menschaert, G. & Waegeman, W. DeepRibo: a neural network for precise gene annotation of prokaryotes by combining ribosome profiling signal and binding site patterns. Nucleic Acids Res. 47, e36–e36 (2019).

110. Gelhausen, R., et al. RiboReport-benchmarking tools for ribosome profiling-based identification of open reading frames in bacteria. Brief. Bioinform. 23, bbab549 (2022).

111. Buels, R. et al. JBrowse: a dynamic web platform for genome visualization and analysis. Genome Biol. 17, 66 (2016).

112. Kamatchinathan, S. et al. pridepy: A Python package to download and search data from PRIDE database. J. Open Source Softw. 10, 7563 (2025).

113. Kong, A. T., Leprevost, F. V., Avtonomov, D. M., Mellacheruvu, D. & Nesvizhskii, A. I. MSFragger: ultrafast and comprehensive peptide identification in mass spectrometry–based proteomics. Nat. Methods 14, 513–520 (2017).

114. da Veiga Leprevost, F., et al. Philosopher: a versatile toolkit for shotgun proteomics data analysis. Nat. Methods 17, 869–870 (2020).

115. Qeli, E. & Ahrens, C. H. PeptideClassifier for protein inference and targeted quantitative proteomics. Nat. Biotechnol. 28, 647–650 (2010).

116. Zeghouf, M. et al. Sequential Peptide Affinity (SPA) system for the identification of mammalian and bacterial protein complexes. J. Proteome Res. 3, 463–468 (2004).

117. Beyer, H. M. et al. AQUA cloning: a versatile and simple enzyme-free cloning approach. PLOS ONE 10, e0137652 (2015).

118. Hemm, L. et al. Interactors and effects of overexpressing YlxR/RnpM, a conserved RNA binding protein in cyanobacteria. RNA Biol. 21, 1–19 (2024).

119. Kaltenbrunner, A. et al. Regulation of pSYSA defense plasmid copy number in *Synechocystis* through RNase E and a highly transcribed asRNA. Front. Microbiol. 14, 1112307 (2023).

120. Kelly, C. L., Taylor, G. M., Hitchcock, A., Torres-Méndez, A. & Heap, J. T. A rhamnose-inducible system for precise and temporal control of gene expression in cyanobacteria. ACS Synth. Biol. 7, 1056–1066 (2018).

121. Ishihama, Y., Rappsilber, J. & Mann, M. Modular stop and go extraction tips with stacked disks for parallel and multidimensional peptide fractionation in proteomics. J. Proteome Res. 5, 988–994 (2006).

122. Cox, J. & Mann, M. MaxQuant enables high peptide identification rates, individualized p.p.b.-range mass accuracies and proteome-wide protein quantification. Nat. Biotechnol. 26, 1367–1372 (2008).

123. Tyanova, S. et al. The Perseus computational platform for comprehensive analysis of (prote)omics data. Nat. Methods 13, 731–740 (2016).

124. Perez-Riverol, Y. et al. The PRIDE database resources in 2022: a hub for mass spectrometry-based proteomics evidences. Nucleic Acids Res. 50, D543–D552 (2022).

125. Lorenz, R. et al. ViennaRNA Package 2.0. Algorithms Mol. Biol. 6, 26 (2011).

126. Darty, K., Denise, A. & Ponty, Y. VARNA: Interactive drawing and editing of the RNA secondary structure. Bioinformatics 25, 1974–1975 (2009).

