## Supplementary Information for "A revised genome annotation of the model cyanobacterium *Synechocystis* based on start and stop codon-enriched ribosome profiling and proteogenomics"

[#]Co-sharing first authors

### Supplementary Information

**Overview on Supplementary Datasets** **p. 2**

**Supplementary Tables:** **p. 4**

**Supplementary Figures:** **p. 10**

**Supplementary Code Description** **p. 47**

**Supplementary References:** **p. 50**

**Overview on Supplementary Datasets**

**Supplementary Dataset 1. Results of proteogenomics searches against the standard and custom iPtgxDBs.** The file contains five worksheets: **Worksheet 1** contains a legend explaining the contents of the four other worksheets, including the columns contained therein and an explanation on how to interpret iPtgxDB identifiers. **Worksheet 2** contains all identified proteins, **Worksheet 3** contains those that were already annotated by RefSeq, **Worksheet 4** contains cases where proteomics confirmed a new start codon for an already annotated protein and **Worksheet 5** contains information for up to 69 novel and predominantly small proteins identified in this study.

***See separate file.***

**Supplementary Dataset 2. Mirror plots of spectra assigned to peptides that support the validated SEPs in Table 1.** These mirror plots show the best spectrum for each peptide selected according to Percolator score and hyperscore in the top half of the plot. Where possible, the matching *in silico* predicted spectrum for the same peptide (based on the software Prosit) is shown in the bottom half. In cases where no prediction was possible, the reason is stated instead. Fragment ion intensities shown on the y-axis were normalized and their m/z values are shown on the x-axis. Details shown in the header consist of the peptide sequence including modifications and the protein name, followed by the peptide index for this protein. The second row shows the precursor charge, collision energy used, the m/z value of the precursor and the difference between theoretical and actual precursor mass in Dalton. The third row shows the Percolator score and hyperscore as calculated by Philosopher, followed by the spectral angle and Pearson's correlation coefficient between observed and theoretical spectrum as calculated in Gessulat et al. (2019)[1]. Modifications in the peptide sequence are indicated as follows:

| | |
|---|---|
| M[Ox] | Oxidation of Methionine |
| [Ac]- | N-terminal acetylation |
| [DmL] | Dimethyl label (light) |
| [DmM] | Dimethyl label (medium weight) |
| [DmH] | Dimethyl label (heavy) |
| [Ph] | Phosphorylation |
| [TMT] | TMT label |

***See separate file.***

**Supplementary Dataset 3. Mirror plots of spectra for the peptides supporting the set of up to 69 novel proteins detected with proteogenomics.** See description of **Supplementary Dataset 2** for more details. This table extends **Table S3**.

***See separate file.***

**Supplementary Dataset 4. Tryptic peptides from the standard and custom iPtgxDBs along with their peptide evidence class.** The Excel file contains two work sheets, one for the standard iPtgxDB and one for the custom iPtgxDB. Each work sheet lists in column 1 the peptide sequences of all theoretically possible tryptic peptides in the database with a length of six amino acids or above, the respective peptide length in column 2, the peptide class computed by PeptideClassifier in column 3, and the identifiers of the protein sequences that this peptide implies in column 4 (if multiple proteins are implied, they are separated by semicolons). **Figure S7B** summarizes these data graphically.

***See separate file.***

### Supplementary Tables

**Table S1. Genes with footprint 3′ ends located 2 nt and 4 nt downstream the annotated start codons.** This table extends **Figure 1F**.

***See separate Excel file.***

**Table S2. Ribo-seq and proteogenomic analyses of 3,669 KZS-annotated genes.** A separate sheet ("Legend") describes the content of the individual columns. Column A provides the identifiers beginning with the GenBank accession, followed by the nt coordinates and a "+" symbol to indicate location on the forward strand and a "-" symbol for the reverse strand (example: NC_000911.1:7229-8311:+ for *psbA2*, *slr1311* gene). Genes were considered translated if the RIBO-WT-avg_TE was ≥0.288 in column AU. Column M indicates how many peptides are in a size range theoretically detectable with mass spectrometry (7-40 aa) and column N marks genes for which a peptide that confirms the exact start site was detected. Columns O-AC and AD-AR summarize matching results from the proteogenomics searches against the custom and standard iPtgxDB if present. From left to right, the columns show the start site predicted by proteogenomics, the number of detected peptides and PSMs, the percentage of the protein sequence covered by peptides, the 6 PeptideClassifier classes (1a, 1b, 2a, 2b, 3a, 3b), the protein novelty classification, whether evidence disagrees with a KZS annotated start, and the sequence of the n-terminal peptide if detected. We also show the encoded amino acid preceding the start codon and whether this results in trypsin cleavage directly before the start codon (which would mean that this n-terminal peptide is not a direct confirmation of the start) and finally the sequences of all detected peptides.

***See separate Excel file.***

**Table S3**. **List of 2,708 genes classified as potentially novel or as ORF corrections.** A separate sheet ("Legend") describes the content of the individual columns. The type of novelty or correction is indicated in column A as "Unannotated", "Internal_OutofFrame" (IOF), "Internal_Inframe" (IIF), "N-terminal_extension" and "Truncated" (evidence that ORF should be truncated). The manual inspection of randomly selected entries (n=808) yielded 163 entries (20 %) that were considered real (column F), including 126 encoding proteins of ≤70 amino acids. The number of membrane helices predicted by the DeepTMHMM algorithm[2] is indicated in column C. Overlaps with the findings by Peng et al.[3] are indicated in column E (disclaimer: we just list their peptides as their data was not processed with our pipeline), details on the genomic context in columns G-I. The columns detailing our proteogenomics evidence (columns J-AO) match those listed above for **Table S2** (columns M-AR). In column AP, comments are listed for the manually inspected instances. Coordinates and genetic information for all entries can be found in columns AQ-AZ. Options to filter the data: Columns BA-BC contain the translational efficiencies calculated from Ribo-seq and RNA-seq coverage in two replicates, BD-BF provide the coverage from TIS-Ribo-seq.

The respective sequencing coverages given in Reads Per Kilobase of transcript per Million mapped reads (RPKM) are provided in columns BG-BN. Columns BO-CZ provide values for prepared disome and trisome fractions (if available) and for TTS-Ribo-seq. For further details on the assigned peptides and spectra, see **Supplementary Dataset 3.**

***See separate Excel file.***

**Table S4**. **List of 165 genes encoding proteins ≤70 aa (KZS annotation).** A separate sheet ("Legend") describes the content of the individual columns. The columns detailing our proteogenomics evidence (columns J-AO), again match those listed for **Table S2** (columns M-AR). Proteogenomically detected proteins are labeled "Yes" in column J. All entries were manually validated in the RIBOBASE database for the presence of 3' end peaks in TIS- and TTS-Ribo-seq (columns AP and AQ), and for contiguous Ribo-seq coverage (column AR). Proteins previously detected by mass spectrometry in the study by Baers et al.[4] are labeled "yes" in column AS. The average and respective translational efficiencies calculated from Ribo-seq and RNA-seq coverage in two replicates (columns AZ-BB), and from TIS-Ribo-seq (columns BC-BE) are given, followed by the respective coverages given in Reads Per Kilobase of transcript per Million mapped reads (RPKM); columns BF-BN).

***See separate Excel file.***

**Table S5. Overlap of proteins identified in searches against a standard and custom iPtgxDB.** The proteins are further grouped into different subcategories (all, RefSeq, novel start sites or novel CDS identifications).

| | **All** | **RefSeq** | **Starts** | **Novels** |
|---|---|---|---|---|
| **iPtgxDB standard** | 3111 | 3021 | 46 | 44 |
| **iPtgxDB custom** | 3128 | 3041 | 48 | 39 |
| **Both** | 3061 | 3007 | 40 | 14 |
| **Total** | **3178** | **3055** | **54** | **69** |

**Table S6. Deoxyribonucleotides and primers used in this manuscript.** All primers are given in 5'→3' orientation. Capital letters indicate segments binding to *Synechocystis* 6803 sequences.

| ID | Full Name | Sequence |
|---|---|---|
| VK1 | infC-in fwd | tacaatcgccaagaagtATGAACACGACTATC |
| VK2 | infC-in rev | tctcttttccgcggaCCGTTGCCTTGACTTT |
| VK3 | Slr1397 fwd v2 | tacaatcgccaagaagtATGGATAATCTAACCG |
| VK4 | Slr1397 rev | tctcttttccgcggaTCTTCCATCGATTGG |
| VK5 | Slr0601 fwd v2 | tacaatcgccaagaagtATGTCCACGGAAAAC |
| VK6 | Slr0601 rev | tctcttttccgcggagTGATTGCTGTGCTC |
| VK7 | Sll0871 fwd v2 | tacaatcgccaagaagtATGAACACTTTTTTGGC |
| VK8 | Sll0871 rev | tctcttttccgcggagCCCCGGACCAACAG |
| VK9 | crtR-in fwd | cgccaagaagtATGACCCTGAAAATG |
| VK10 | crtR-in rev | tctcttttccgcggagTAGCTCATATTTGC |
| VK11 | slr1923-in fwd v2 | tacaatcgccaagaagtATGGCCGGTCGAGAG |
| VK12 | slr1923-in rev | tctcttttccgcggagCATTTCACAACCAATG |
| VK13 | Toop_PstI_puc19_short fwd3 | ttcgctcggttgccgccgggcgtttttattactgcaggagacgaaagggcctcgtg |
| VK14 | puc19_short_SalI rev | gtcgacaaaggccaggaaccgtaaaaagg |
| VK15 | PpetE_puc19_short fwd 2 | cctttttacggttcctggcctttgtcgacCTGGGCCTACTGGGCTATTC |
| VK16 | ncl1450_3'UTR rev 2 | aggccctttcgtctcctgcagtaataaaaaacgcccggcggcaaccgagcgaaTAATTCCCAACGAAGGCAAGC |
| VK17 | PpetE rev | ACTTCTTGGCGATTGTATCTATAGG |
| VK18 | 3 x FLAG_ 3'UTR_Toop fwd | gattataaagatcatgatgg |
| VK19 | ncl1450 rev | tgagcgtcatACTTCTTGGCGATTGTATCTATAGG |
| VK20 | Ncl1450 fwd | gccaagaagtATGACGCTCATCGACACC |
| VK21 | TU1220#1 fwd | agatacaatcgccaagaagtATGAAACATTCACGGAGAAATTTTTTCG |
| VK22 | TU1220#1 rev | ccatcatgatctttataatcATATGGTGCGAGTCCCCC |
| VK23 | TU1220#2 fwd | agatacaatcgccaagaagtATGATGGGGAATATCAGTTATG |
| VK24 | TU1220#2 rev | ccatcatgatctttataatcTTCACATTTACCACGGCAAG |
| VK25 | TU1183 fwd | agatacaatcgccaagaagtATGAGCAAATCTGCCGTTC |
| VK26 | TU1183 rev | ccatcatgatctttataatcGTTGTCCTTATTGCCACAG |
| VK27 | Ncr1610 fwd | agatacaatcgccaagaagtATGAACACCAGAACCCAAAC |
| VK28 | Ncr1610 rev | ccatcatgatctttataatcAGCGACCACAACGGGGGT |
| VK29 | TU1447 fwd | agatacaatcgccaagaagtATGGTGGGGGCGGGCTGG |
| VK30 | TU1447 rev | ccatcatgatctttataatcACCCATCTGGCCGTCTCGG |
| VK31 | Ncr1470 fwd | agatacaatcgccaagaagtATGTGGAACGCGAACCAC |
| VK32 | Ncr1470 rev | ccatcatgatctttataatcACACCTCCTTGATTGTATGAC |
| VK33 | TU2407 fwd | agatacaatcgccaagaagtGTGGAGGTCAAAGGGCGATC |
| VK34 | TU2407 rev | ccatcatgatctttataatcGCAGTATCTAGAACAACTAGAGCC |
| VK35 | Slr2031_as fwd | agatacaatcgccaagaagtATGGGCGGCGTTGCTATAG |
| VK36 | Slr2031_as rev | ccatcatgatctttataatcATTATGACCATGACTAGGGGG |
| VK37 | Norf2 fwd | agatacaatcgccaagaagtATGTACGCAATAGAGTTTG |
| VK38 | Norf2 rev | ccatcatgatctttataatcGATCCATACATCATCCTC |
| VK39 | TU7087 fwd | agatacaatcgccaagaagtATGCAACTAAAACACTGGCAATCTC |
| VK40 | TU7087 rev | ccatcatgatctttataatcTGGCTGGTTTCCAGCCCA |
| VK41 | nsiR7_ORF fwd | agatacaatcgccaagaagtATGAAACCGACTCATTTC |
| VK42 | nsiR7_ORF rev | ccatcatgatctttataatcACTATCCGTAACATTTGAC |
| VK43 | SPA-tag backbone fwd | tgacaagtagCGCCTCCATTCCCCAACG |
| VK44 | SPA-tag fwd | ccgcggaaaagagaagatgg |
| VK45 | SPA-tag rev | AATGGAGGCGctacttgtcatcgtcatcc |
| VK46 | Ncr1610 backbone rev | ttttccgcggaAGCGACCACAACGGGGGTC |
| VK47 | TU7087 backbone rev | ctcttttccgcggaTGGCTGGTTTCCAGCCCA |
| VK48 | Slr2031-as backbone rev | ctcttttccgcggaATTATGACCATGACTAGGGGG |
| VK49 | TU1220#1 backbone rev | ctcttttccgcggaATATGGTGCGAGTCC |
| VK50 | TU1220#2 backbone rev | ctcttttccgcggaTTCACATTTACCACG |
| VK51 | TU1183 backbone rev | ctcttttccgcggaGTTGTCCTTATTGC |
| VK52 | TU1447 backbone rev | ctcttttccgcggaACCCATCTGGCCGT |
| VK52 | Ncr1470 backbone rev | ctcttttccgcggaACACCTCCTTGATT |

| | | |
|---|---|---|
| VK53 | TU2407 backbone rev | ctcttttccgcggaGCAGTATCTAGAACA |
| VK54 | Ncr1420 fwd | tacaatcgccaagaaGTGTGATATCTGTGAA |
| VK55 | Ncr1420 rev | tctcttttccgcggagCCATTGTCCTGGTC |
| VK56 | ssr0758_fwd | cgccaagaagtATGAAGAAAGAGTA |
| VK57 | Ssr0758 rev | tctcttttccgcggaAAATGAAAGTTCTCG |
| VK58 | Ncl1350 fwd | tacaatcgccaagaagtATGTTCTGGAAGC |
| VK59 | Ncl1350 rev | tctcttttccgcggagAGGGAAGTTTCCTTG |
| VK60 | pVZ322_seq fwd | tggttaattggttgtaacactggcag |
| VK61 | pVZ322_seq rev | gtaataccatgaaaaataccatgctcag |
| VK62 | pUC19_shorter_seq fwd | gaaatgttgaatactcatactcttcc |
| VK63 | pUC19_shorter_seq rev | atagtcctgtcgggtttcgcc |
| P11-45 | pUC19_fwd_shorterbackbone2 | agctcactcaaaggcggtaa |
| P11-46 | pUC19_rev_shorterbackbone2 | tcaccgtcatcaccgaaacg |
| G3-64 | pUC19s-PpetE_Fwd | cgtttcggtgatgacggtgaCTGGGCCTACTGGGCTATTC |
| G3-65 | oop-pUC19s_Rev | ttaccgcctttgagtgagctaataaaaaacgcccggcg |
| VR277 | PpetE_fwd2 | ggattacagatcctctagagCTGGGCCTACTGGGCTATTC |
| VR278 | PpetE_rev1 | cgttgttcatACTTCTTGGCGATTGTATCTATAGG |
| VR279 | as_psbC_fwd | gccaagaagtATGAACAACGTGGGTTCCG |
| VR280 | as_psbC_rev | tttccgcggaCCTCTGGCATGCTGGTCG |
| VR281 | Spa-3UTR-oop_fwd | atgccagaggtccgcggaaaagagaagatg |
| VR282 | Spa-3UTR-oop_rev | tatgctcttctgctcctgcaaataaaaaacgcccggcg |
| VR285 | PpetE_rev2 | cgactttcatACTTCTTGGCGATTGTATCTATAGG |
| VR286 | as_Csx18_fwd | gccaagaagtATGAAAGTCGAACAGGCC |
| VR287 | as_Csx18_rev | tttccgcggaTACTATTTTTGTGGTTGGGG |
| VR288 | SPA_UTR_oop_fwd | aaaaatagtatccgcggaaaagagaagatg |
| 101 | Slr1079-Leadless_fwd | ctttttagactggtcgtaatgaaATGGCAAGTTTTCTGGCTTTAC |
| 102 | Slr1079+Flag_rev_new | ccatcatgatcatgatctttataatccatTTCGCCGTATTCCTGGAG |
| 105 | NesSlr1079+SynRBS_fwd | aatatacaaaggaggtagaaATGGCAACATCGACACCACCCCCATCCCC |
| 107 | Nes1079-Flag_rev | gaatttggtaccgagctgcagTTAGGGTTGCTCTGGCTTCTGGGCTAGGG |
| 109 | Slr0489-Leadless_fwd | ctttttagactggtcgtaatgaaATGGCAACCTTCTTGGCCC |
| 113 | NesSlr0489+SynRBS_fwd | gtttataatatacaaaggaggtagaaATGGCAACATCGACACCACCCCCTTCCAACTC |
| 115 | Nes0489-Flag_rev | ggaatttggtaccgagctgcagTCAGGGCTGATCTGGTATCTGGGTTAGGGG |
| 117 | Slr0489_5'Fl_fwd | cacgaggccctttcgtctATGGGTAAATTGCGGCTGAG |
| 118 | Slr0489_5'Fl_rev | gcgttgacatcactctgtacGTAACAATATTGATCTGTGCTGGAAC |
| 119 | Slr0489_StrepR_fwd | GCACAGATCAATATTGTTACgtacagagtgatgtcaacgcc |
| 120 | Slr0489_StrepR_rev | CTAAAAAACCTAACTCTTTCATtattatcgtagttgctctcagagttg |
| 121 | Slr0489_5'UTR_fwd | ctgagagcaactacgataataATGAAAGAGTTAGGTTTTTTAGATGTTCC |
| 122 | Slr0489_5'UTR_rev | caataaattagggctcgccatGGCATCATTCTAGCTACTTCAAGC |
| 123 | Slr0489_3'Fl_fwd | gaagtagctagaatgatgccATGGCGAGCCCTAATTTATTGC |
| 124 | Slr0489_3'Fl_rev | ctttttacggttcctggcctttAACTGCTTTAAACCGTCCACTG |
| 133 | 133-SegSlr0489_fwd | cgtgttgcaatggttggctc |
| 134 | 134-SegSlr0489_rev | ggaagcagacaaaaactattaatttggcc |
| RB2/110 | Slr0489+Flag_rev_new | ccatcatgatctttataatccatGGGCTCGCCATACTCTTGG |
| RB3/104 | Slr1079+SynRBS_fwd | aatatacaaaggaggtagaaATGGCAAGTTTTCTGGCTTTAC |
| RB4 | Slr1079+Flag_rev_new | ccatcatgatctttataatccatTTCGCCGTATTCCTGGAG |
| RB5/116 | NesSlr0489+NatUTR_fwd | ctttttagactggtcgtaatgaaTTTCTGTATGCTGTAGCGGCAT |
| RB6/114 | NesSlr0489+Flag_rev_new | cgccatcatgatctttataatccatGGGCTGATCTGGTATCTGGGTTAGGGG |
| RB7/108 | NesSlr1079+NatUTR_fwd | ctttttagactggtcgtaatgaaTTCCTTTATGCCGTTGCGGCCC |
| RB8/106 | NesSlr1079+Flag_rev_new | catgatctttataatccatGGGTTGCTCTGGCTTCTGGGCTAGGG |
| RB9 | pUC19_Rha_3xFLAG_fwd | atggattataaagatcatgatgg |
| RB10 | pUC19_Rha_3xFLAG_rev | ttctacctcctttgtatattataaac |
| RB11 | pUC19_Rha_fwd | ctgcagctcggtaccaaattc |
| RB12 | pUC19_Rha_noRBS_rev | ttcattacgaccagtctaaaaag |
| RB13 | slr0489_fw | ttaataaggagatataccATGGCAACCTTCTTGGCC |

| | | |
|---|---|---|
| RB14 | pACYDuet_slr0489-6H-rev | gccccaaggggttatgctagttagtggtgatgatggtgatgGGGCTCGCCATACTCTTGG |
| RB15 | His_tag_pACYCDuet_fw | catcaccatcatcaccactaac |
| RB16 | pACYCDuet_rev | ggtatatctccttattaaagttaaac |
| RB17 | slr0489short_mut_fw | ATTGAGGTTTTAcGGCAACATCGACACC |
| RB18 | slr0489short_mut_rev | TAAAACCTCAATAATGCCGCTAC |
| RB19/ 111 | Slr0489-Flag_rev | ggaatttggtaccgagctgcagTTAGGGCTCGCCATACTCTTGG |

**Table S7A**. **A standard iPtgxDB search database with 145,955 protein entries (plus contaminants) was created.** Excluded were 3,030 possible N-terminal extensions shorter than 6 aa (not identifiable in a mass spectrometer), 70 entries annotated as pseudogenes by RefSeq and 372 entries whose internal start site would not be distinguishable from a shorter proteoform of the longer RefSeq annotation (if both start with a methionine).

| Name | Annotati-ons | Clus-ters | New clus-ters | New trun-cations | New exten-sions | Total clus-ters | Total ids |
|---|---|---|---|---|---|---|---|
| RefSeq | 3,692 | 3,692 | 3,692 | 0 | 0 | 3,692 | 3,692 |
| GenBank | 3,564 | 3,564 | 89 | 146 | 425 | 3,781 | 4,352 |
| Kazusa | 3,275 | 3,274 | 38 | 148 | 16 | 3,819 | 4,554 |
| Prodigal | 3,724 | 3,724 | 101 | 205 | 65 | 3,920 | 4,925 |
| ChemGenome | 5,071 | 5,071 | 1,802 | 63 | 997 | 5,722 | 7,787 |
| In-silico ORFs | 153,404 | 107,718 | 102,033 | 61 | 39,546 | 107,755 | 149,427 |

**Table S7B**. **A custom iPtgxDB search database with 6,928 protein entries (plus contaminants) was created.** Excluded were 109 possible N-terminal extensions shorter than 6 aa (not identifiable in a mass spectrometer), 70 entries annotated as pseudogenes by RefSeq, and 523 entries whose internal start site would not be distinguishable from a shorter proteoform of the longer RefSeq annotation (if both start with a methionine).

| Name | Annotati-ons | Clus-ters | New clus-ters | New trunca-tions | New extensi-ons | Total clus-ters | Total ids |
|---|---|---|---|---|---|---|---|
| RefSeq | 3,692 | 3,692 | 3,692 | 0 | 0 | 3,692 | 3,692 |
| GenBank | 3,564 | 3,564 | 89 | 146 | 425 | 3,781 | 4,352 |
| Kazusa | 3,275 | 3,274 | 38 | 148 | 16 | 3,819 | 4,554 |
| Prodigal | 3,724 | 3,724 | 101 | 205 | 65 | 3,920 | 4,925 |
| Ribo-Seq | 2,711 | 2,627 | 2,220 | 322 | 163 | 6,140 | 7,630 |

**Table S8. List of proteins enriched in co-IP experiments with Slr0489L-3xFLAG and control, ranked by statistical significance (Student‘s t-test).** The tagged proteins used as bait in the experiment (Slr0489L-3xFLAG) and in the control (sfGFP-3xFLAG) are highlighted in boldface letters. This table relates to results shown in **Figure 7D**. For the histograms of the label-free quantification (LFQ) value distribution, see **Figure S23**.

***See separate Excel file.***

**Table S9. List of proteins enriched in co-IP experiments with Slr0489S-3xFLAG and control, ranked by statistical significance (Student‘s t-test).** The tagged proteins used as bait in the experiment (Slr0489S-3xFLAG) and in the control (sfGFP-3xFLAG) are highlighted in boldface letters. This table relates to results shown in **Figure 7E**. For the histograms of the LFQ data distribution, see **Figure S24**.

***See separate Excel file.***

**Table S10. List of proteins enriched in co-IP experiments with Slr1079L-3xFLAG and control, ranked by statistical significance (Student‘s t-test).** The tagged proteins used as bait in the experiment (Slr1079L-3xFLAG) and in the control (sfGFP-3xFLAG) are highlighted in boldface letters. This table relates to results shown in **Figure S21B**. For the histograms of the LFQ data distribution, see **Figure S25**.

***See separate Excel file.***

**Table S11. List of proteins enriched in co-IP experiments with Slr1079S-3xFLAG and control, ranked by statistical significance (Student‘s t-test).** The tagged proteins used as bait in the experiment (Slr0489S-3xFLAG) and in the control (sfGFP-3xFLAG) are highlighted in boldface letters. This table relates to results shown in **Figure S21C**. For the histograms of the LFQ data distribution, see **Figure S26**.

***See separate Excel file.***

**Table S12. Extrapolation of the number of expected true-positive translons based on the manual inspection of 808 candidates**.

***See separate Excel file.***

**Supplementary Figures**

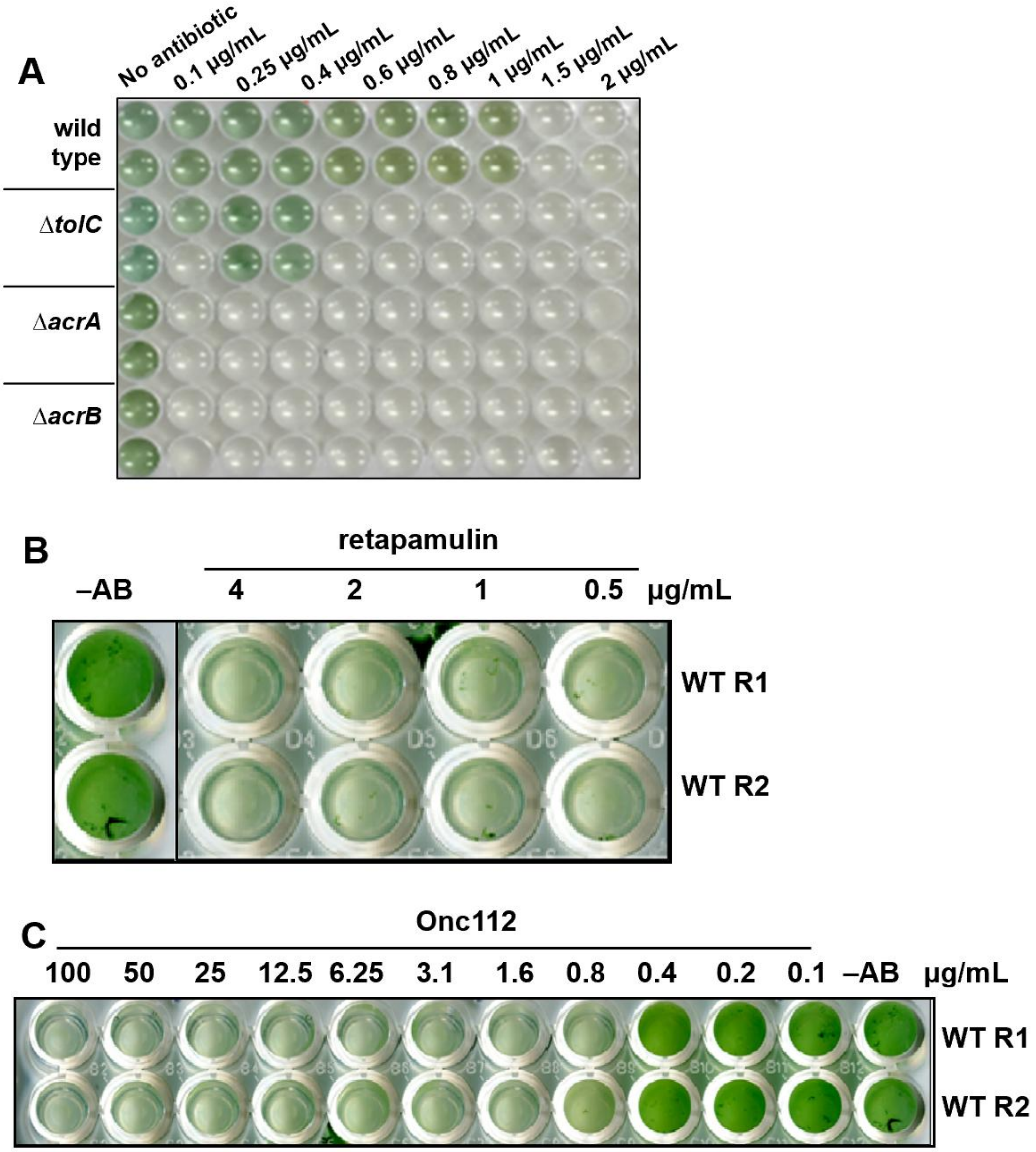


**Figure S1. Assays to define the minimal inhibitory concentration for retapamulin and Onc112 in *Synechocystis* 6803. A.** The wild type and different *tolC*-like transporter mutants[5] were grown in BG11 liquid culture volumes of 200 µL in the presence of retapamulin and documented after 7 days in two replicates. **B.** Wild type sensitivity test toward higher retapamulin concentrations in BG11 liquid cultures after 7 d in two replicates. **C.** Test of wild type sensitivity toward Onc112 after 7 d in two replicates. The respective concentrations are indicated in all panels (-AB, no antibiotic was added).

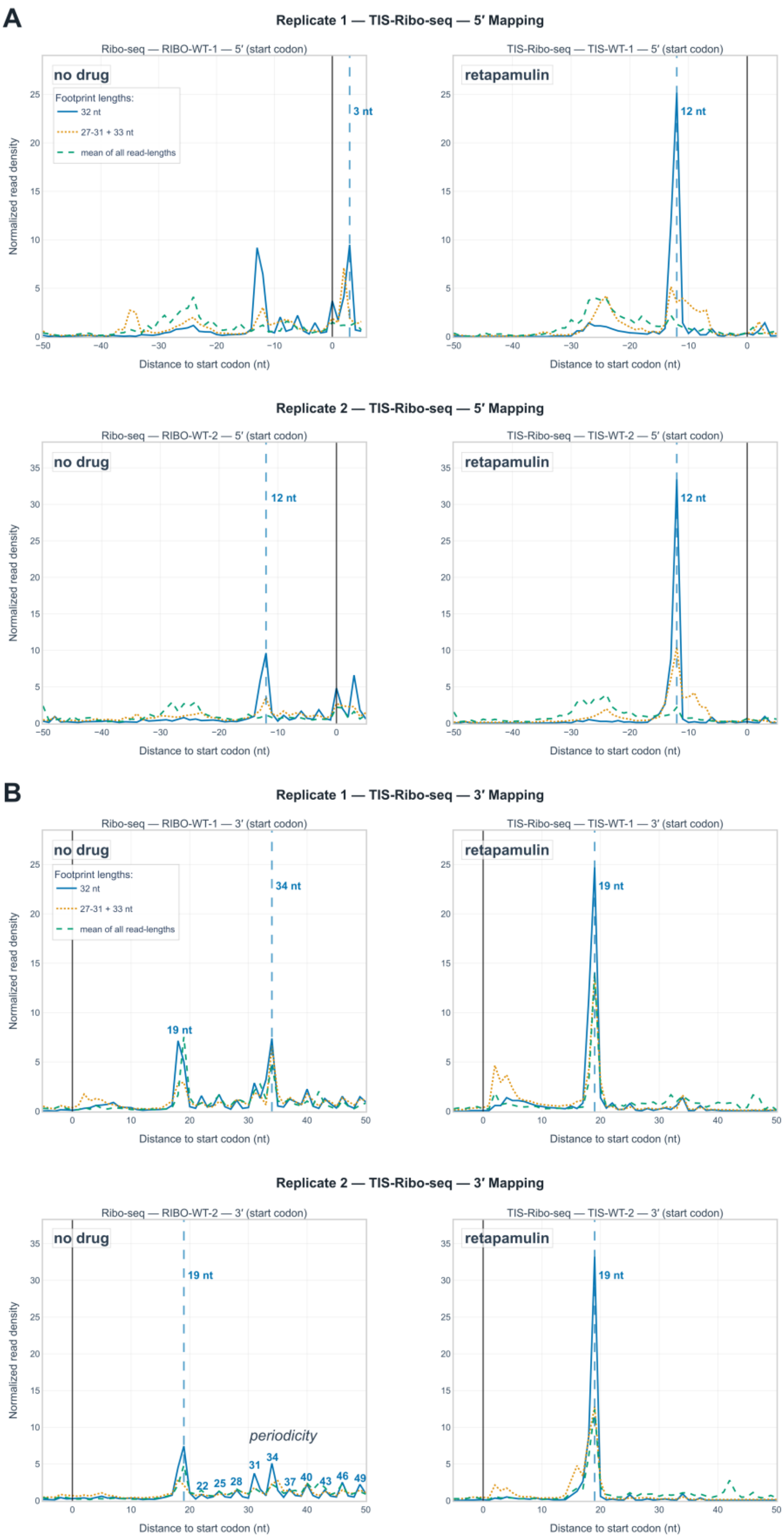


**Figure S2. Metagene analysis of ribosome occupancy (no drug vs. Ret) at start codons for indicated footprint lengths. A.** 5′ read end mapping. The vertical solid black lines indicate the position of the first nt of the start codon. The vertical dashed blue lines indicate the respective peaks. **B.** 3’ read end mapping. The vertical lines indicate start codon and peak positions as in panel A. Distances to the start codon are numbered for the lower peaks in the bottom-left panel to highlight the visible 3-nt periodicity.

In both A and B, the left panels show Ribo-seq (no drug), the right panels TIS-Ribo-seq (in the presence of retapamulin). In all panels, 32 nt-long reads are indicated by solid blue lines; reads of lengths 27-31 or 33 nt by the dotted brown lines and the means of all lengths by the dashed green lines. Footprint counts were normalized to the window length. For additional details, see https://www.bioinf.uni-freiburg.de/~ribobase/synechocystisbrowsepublic/data/metagene/tis_metagene_profiling.html. This figure extends **Figure 1**.

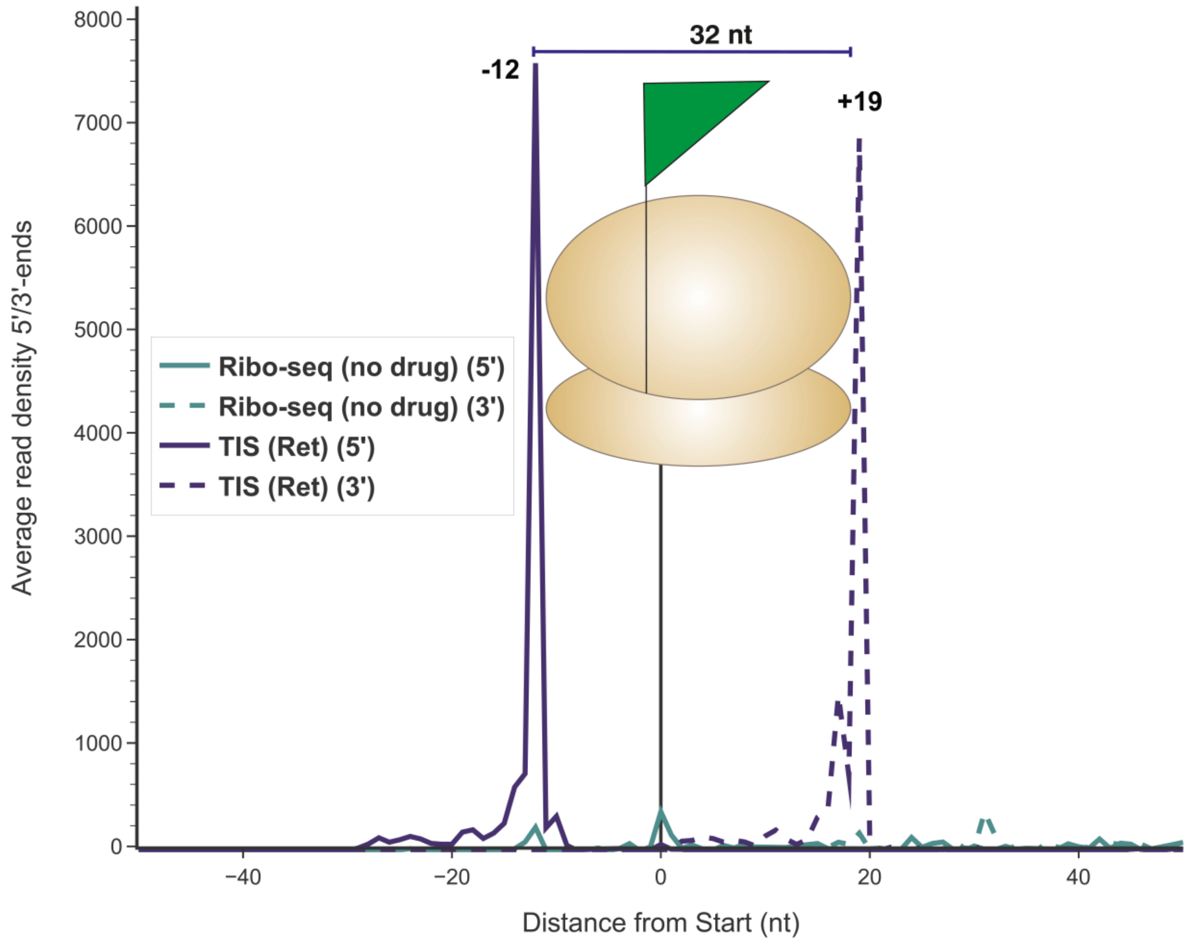


**Figure S3. Metagene analysis of footprints enriched in TIS-Ribo-seq for very short translons.** A sharp 5' end was observed for on average 6,567 reads 12 nt upstream and 19 nt downstream of the start codons of 39 small ORFs (≤50 aa). This demonstrates that sORFs exhibit translational initiation patterns comparable to those observed for other translons in **Figure 1E**.

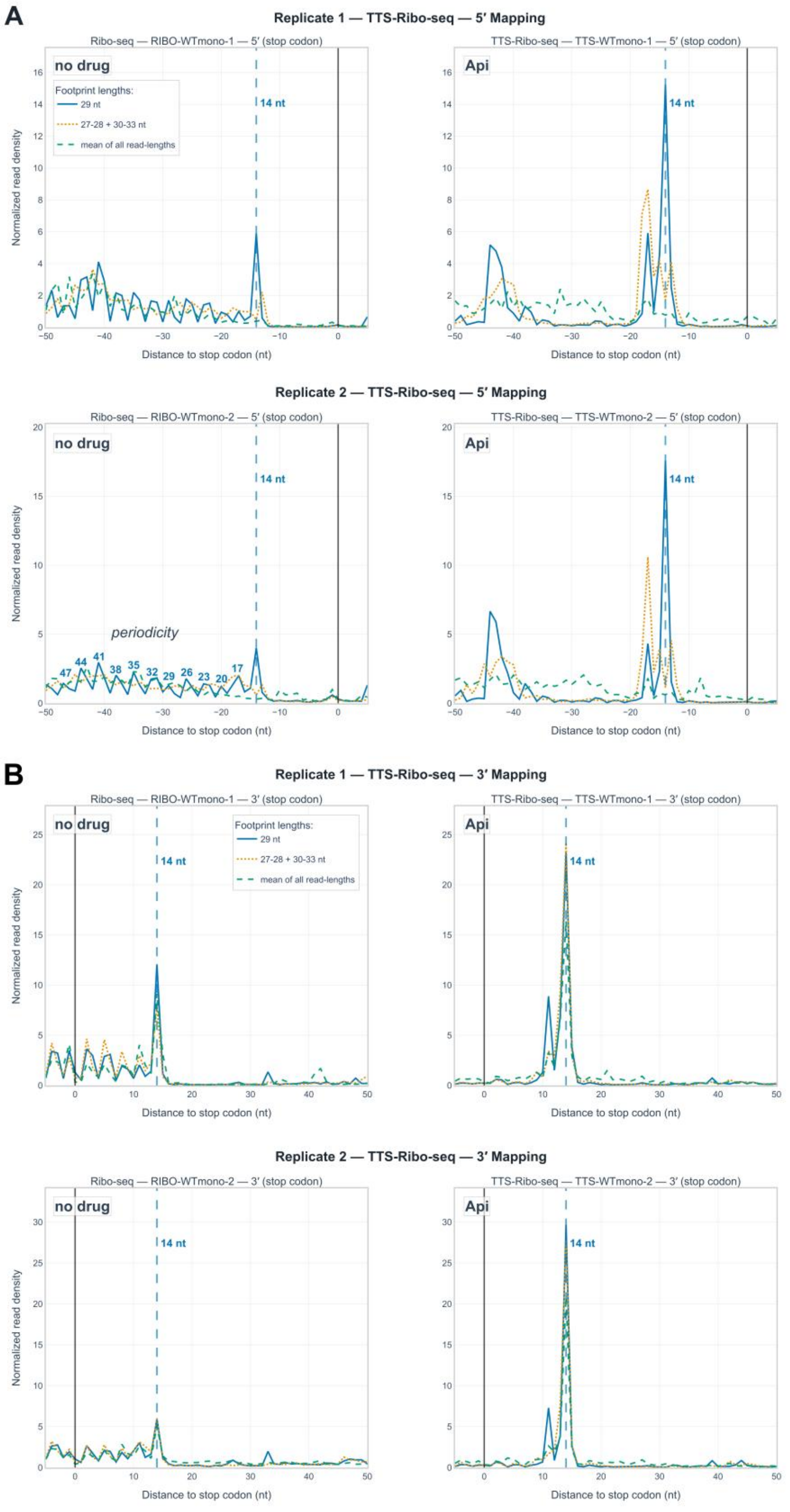


**Figure S4. Metagene analysis of ribosome occupancy (no drug vs. Api137) at stop codons for indicated footprint lengths. A.** 5′ read end mapping. The vertical solid black lines indicate the position of the last nt of the stop codon. The vertical dashed blue lines indicate the respective peaks and their distance to the last nt of the stop codon. The visible 3-nt periodicity is highlighted in the bottom-left panel by giving the distances of lower peaks to the stop codon in nt. **B.** 3' read end mapping. The vertical lines indicate stop codon and peak positions as in panel A.

In both A and B, the left panels show Ribo-seq (no drug), the right panels show TTS-Ribo-seq (in the presence of Api137). In all panels, 29 nt-long reads are indicated by solid blue lines; reads of lengths 27-28 or 30-33 nt by the dotted brown lines and the means of all lengths by the dashed green lines. Footprint counts were normalized to the window length. For additional details, see https://www.bioinf.uni-freiburg.de/~ribobase/synechocystisbrowsepublic/data/metagene/tts_metagene_profiling.html. This figure extends **Figure 2**.

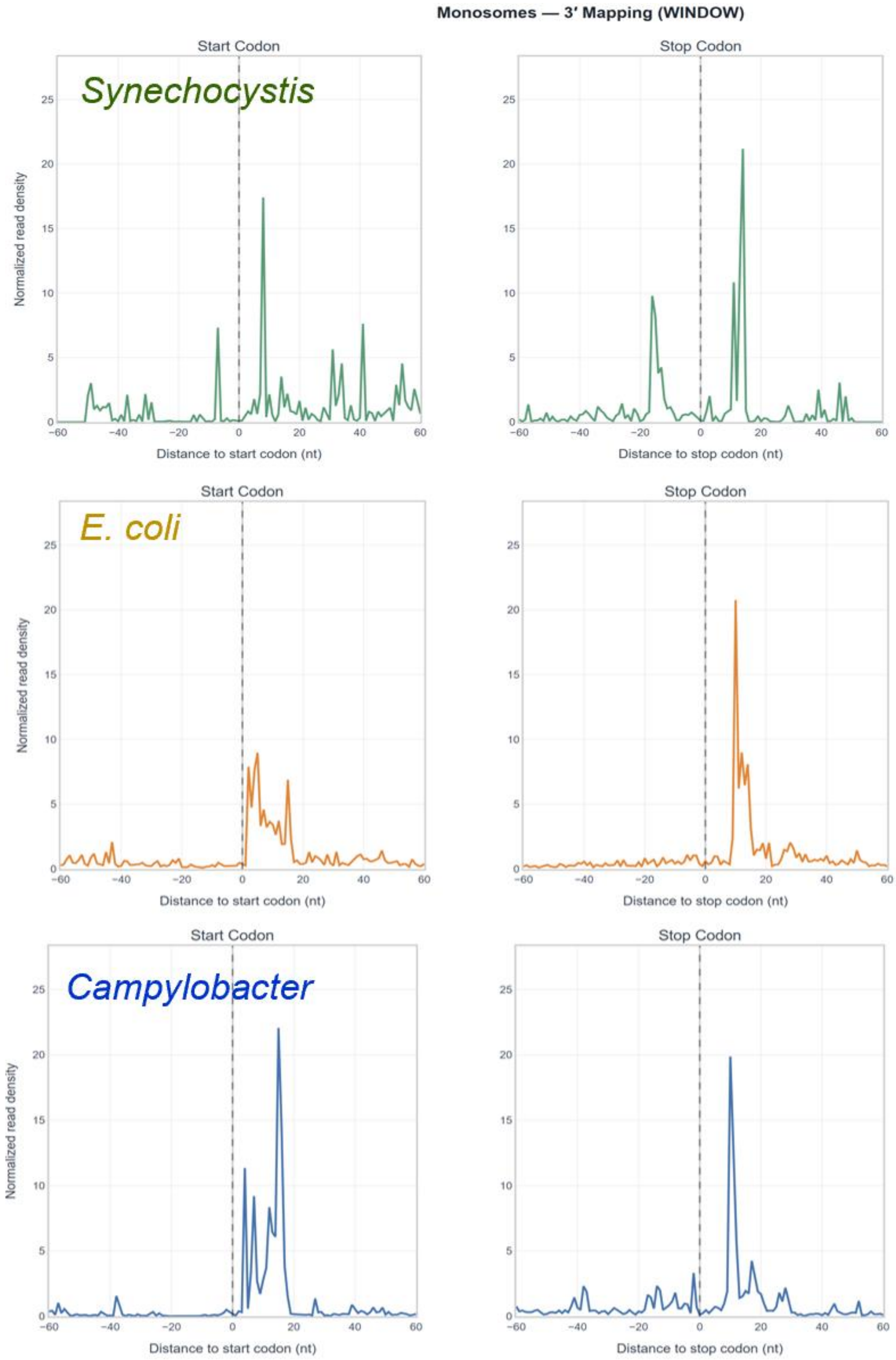


**Figure S5. Metagene analyses of ribosomal footprints at start and stop codons in the presence of Api137 in three different bacteria.** The 3’ ends of reads are shown mapped around start codons (left) and stop codons (right) in the Api137-treated datasets from *Synechocystis* (top, this study), *E. coli*[6] (middle) and *Campylobacter*[7] (bottom). The data from previous analyses were processed in an identical way alongside the *Synechocystis* results. For further replicates and additional details, see https://www.bioinf.uni-freiburg.de/~ribobase/synechocystisbrowsepublic/data/metagene/comparison_metagene_between_organisms.html.

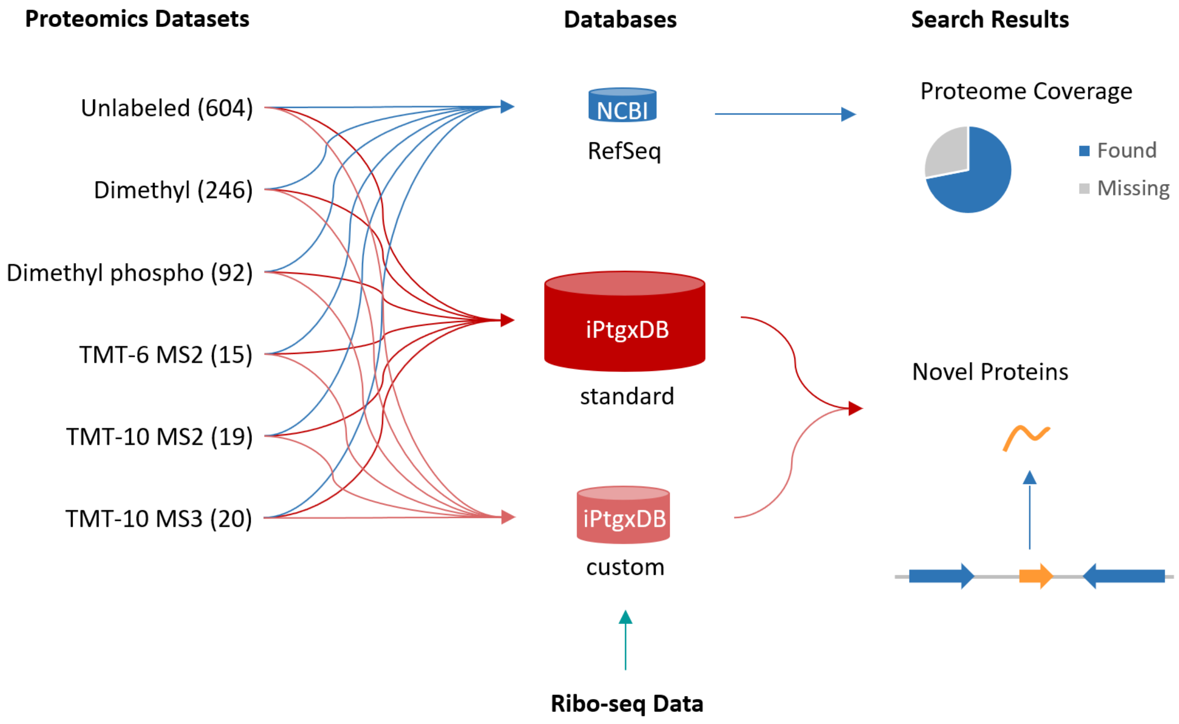


**Figure S6. Proteogenomic analyses.** Searches against three different databases (DBs) were carried out aiming to first establish a rough estimate of the overall proteome coverage (using NCBI RefSeq as search DB), which amounted to roughly 83% of the theoretical proteome, and to then provide protein expression evidence for so far unannotated protein-coding genes of *Synechocystis* 6803 (see **Table S5**). More information about the annotation resources considered for the standard and custom iPtgxDBs is provided in **Figure 3** of the main manuscript.

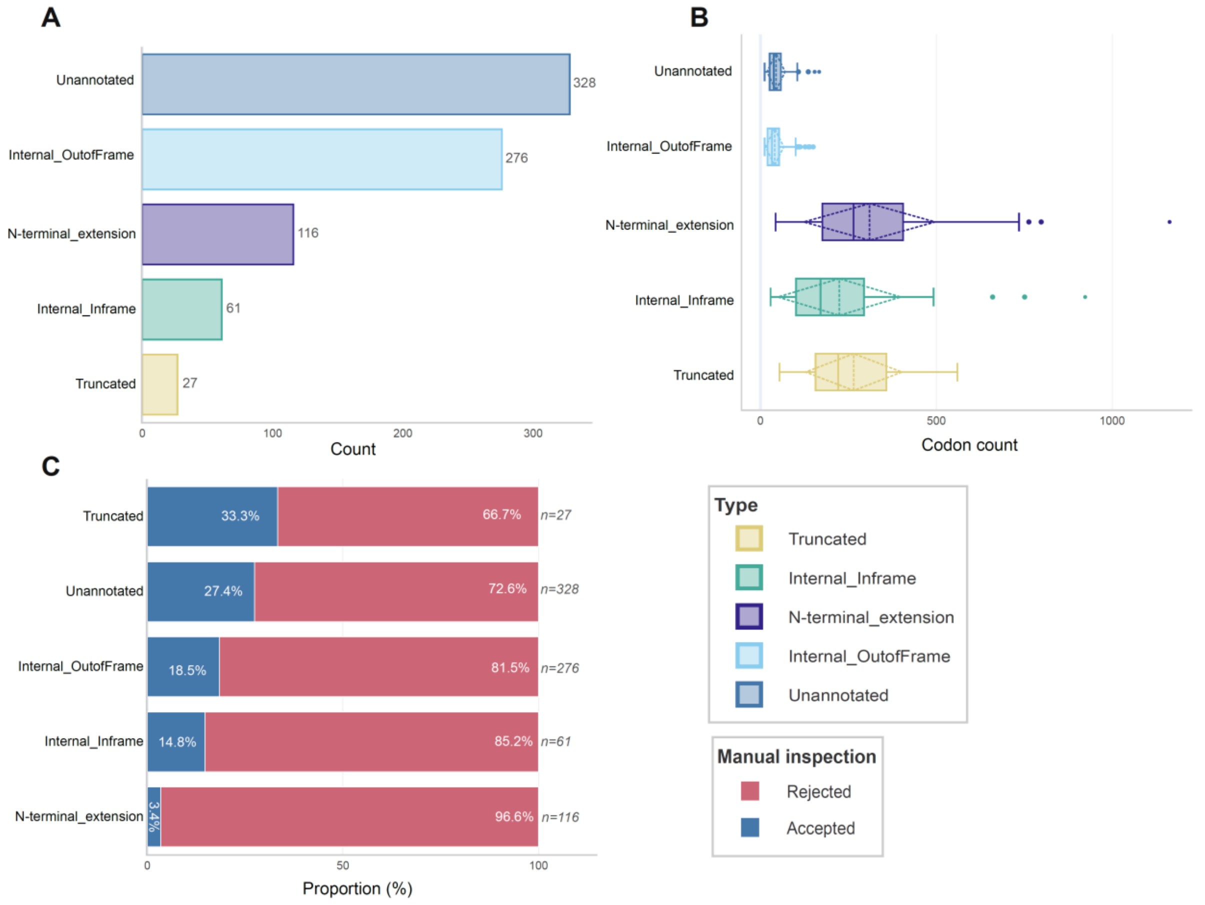


**Figure S7. Overview of predicted translon categories for candidates subjected to manual curation in *Synechocystis* 6803. A**. Number of translons per translon category. **B**. Distribution of codon counts per translon category; box plots show median (centre line), interquartile range (box), and 1.5× IQR (whiskers); diamond indicates the mean and dashed lines show ±1 s.d. **C**. Acceptance rate following manual inspection of Ribo-seq coverage. Of 808 manually inspected instances (30% of all predictions), 163 (20%) were accepted as genuine translons. Sample sizes per category are shown on the right. Extrapolated over the entire set of 2,708 potentially novel translons, an estimated number of 571 new, truncated or extended translons can be expected (**Table S12**).

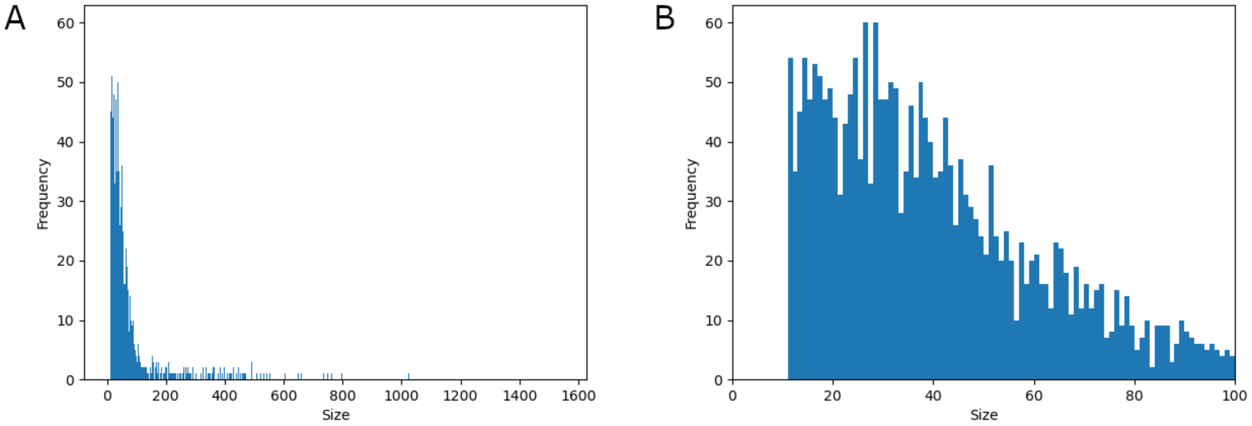


**Figure S8. Histogram of the lengths of 2,708 Ribo-seq candidates**. **A.** Size histogram (in amino acids (aa)) showing the lengths of all top candidates, which are enriched in candidates below 150 aa. **B.** Zoom in to the subset ≤100 aa, here with a bin size of 1.

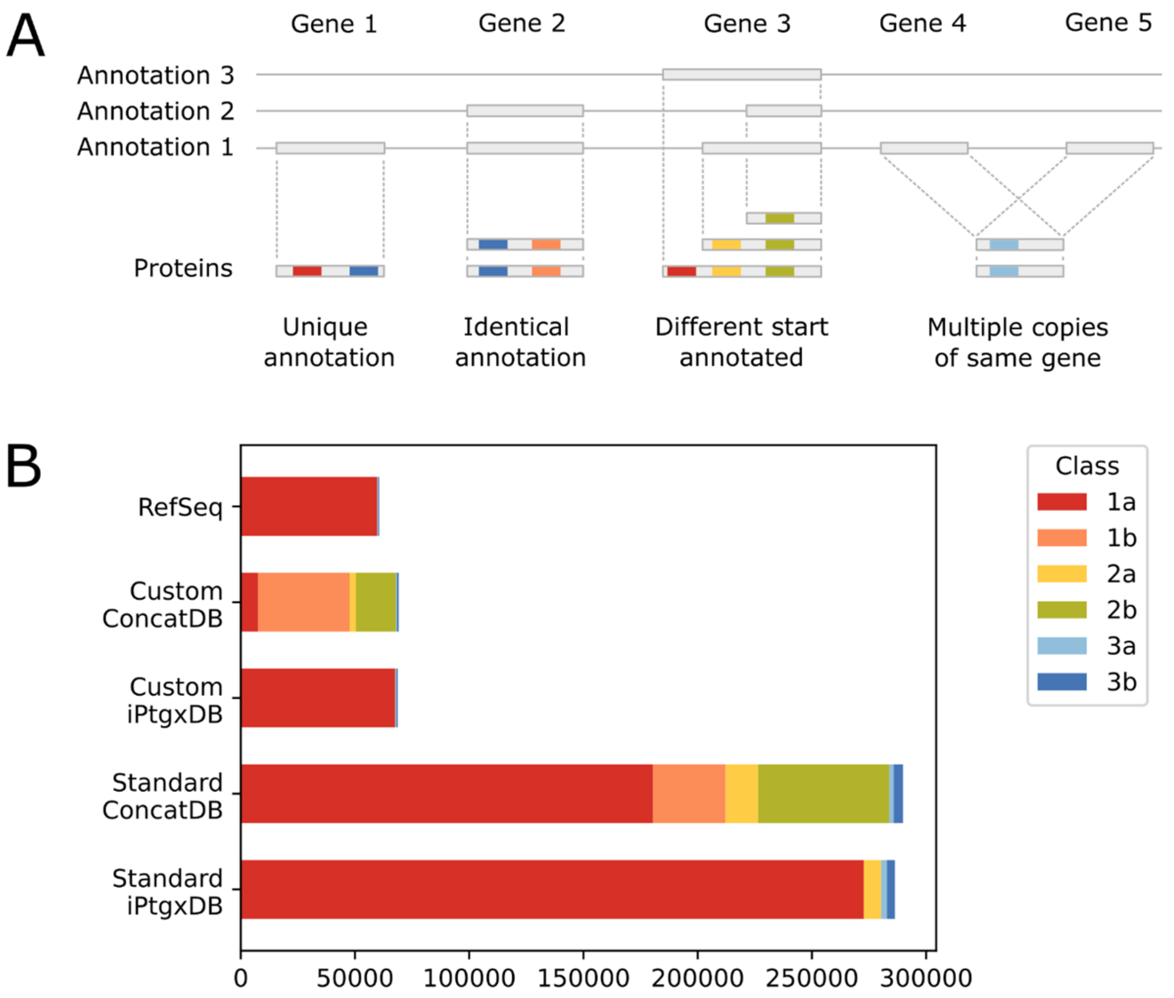


**Figure S9. Concept behind peptide evidence classes used for the proteomic data analysis. A**. Explanation of the six peptide classes distinguished by PeptideClassifier[8] and adapted for proteogenomics in prokaryotes[9]. 1a peptides uniquely identify a single gene annotation while 1b peptides match multiple predictions for the exact same gene by different annotation sources. Peptides of class 2 are ambiguous among all (2b) or multiple (2a) entries of an annotation cluster, that is gene predictions sharing the same stop coordinate. Finally, peptides of class 3 map to multiple genes in different locations, either coding for identical (3a) or different (3b) proteins. **B.** Overview of the peptide information content of different search databases. Concatenation of proteins from different annotation sources (ConcatDBs) leads to much higher percentages of ambiguous peptides. The pre-processing step for the creation of iPtgxDBs, a hierarchical integration that aims to create a minimally redundant, yet highly informative database that covers almost the entire protein coding potential of a prokaryotic genome, eliminates these ambiguities for the most part (see also https://iptgxdb.expasy.org). In the standard iPtgxDB for *Synechocystis* 6803, > 95% of the tryptic peptides unambiguously imply one of the roughly 146,000 proteins (class 1a) and this percentage is even higher in the custom iPtgxDB that contains fewer than 7,000 proteins. This figure extends **Figure 3**.

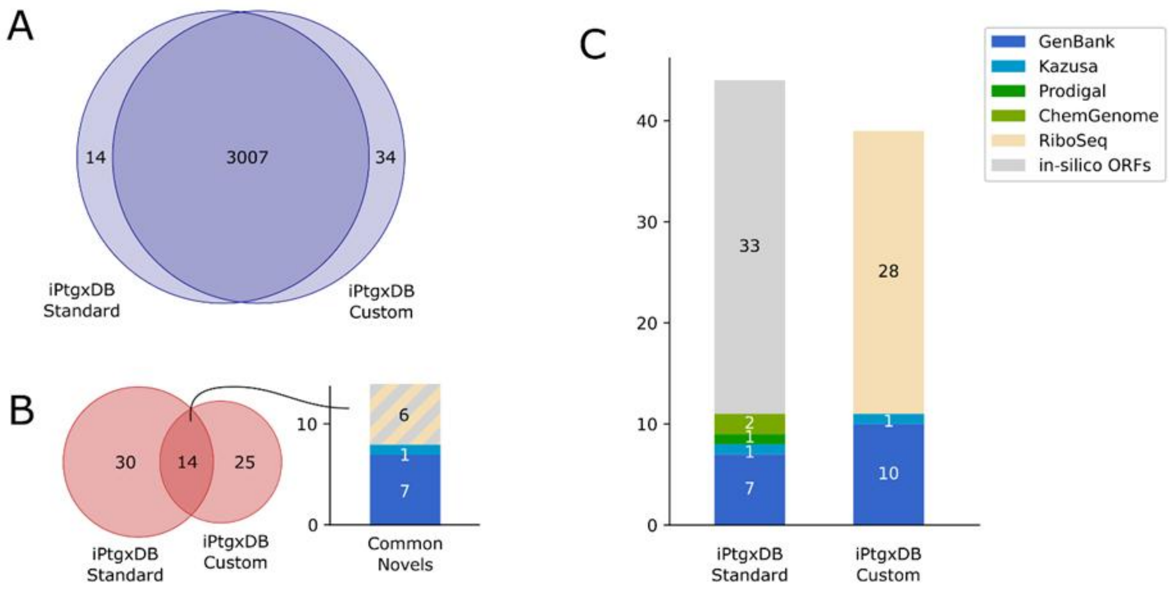


**Figure S10**. **Overlap of known and novel proteins identified by searches against standard and custom iPtgxDBs. A.** The search results obtained for RefSeq proteins are virtually identical (see also **Table S5**). As reported previously, slightly more RefSeq proteins were identified in searches against the smaller custom iPtgxDB[10]. **B**. Overlap of novel proteins identified in both iPtgxDB searches. Overall, up to 69 novel proteins were identified (11 of these were previously predicted by Genbank or Kazusa -see panel C- but were subsequently removed and are missing in the RefSeq 2022 annotation). The subset of 14 novel proteins found in both searches is further detailed: 6 novels were among the 28 novel proteins identified in the custom iPtgxDB that were solely annotated based on RiboSeq data, and among the 33 *in-silico* ORFs identified in the standard iPtgxDB (see also panel C). One Kazusa and 7 Genbank ORFs were also identified in both searches (they are novel compared to the RefSeq2022 annotation). **C**. Summary over the annotation sources for novel proteins identified with standard and custom iPtgxDBs.

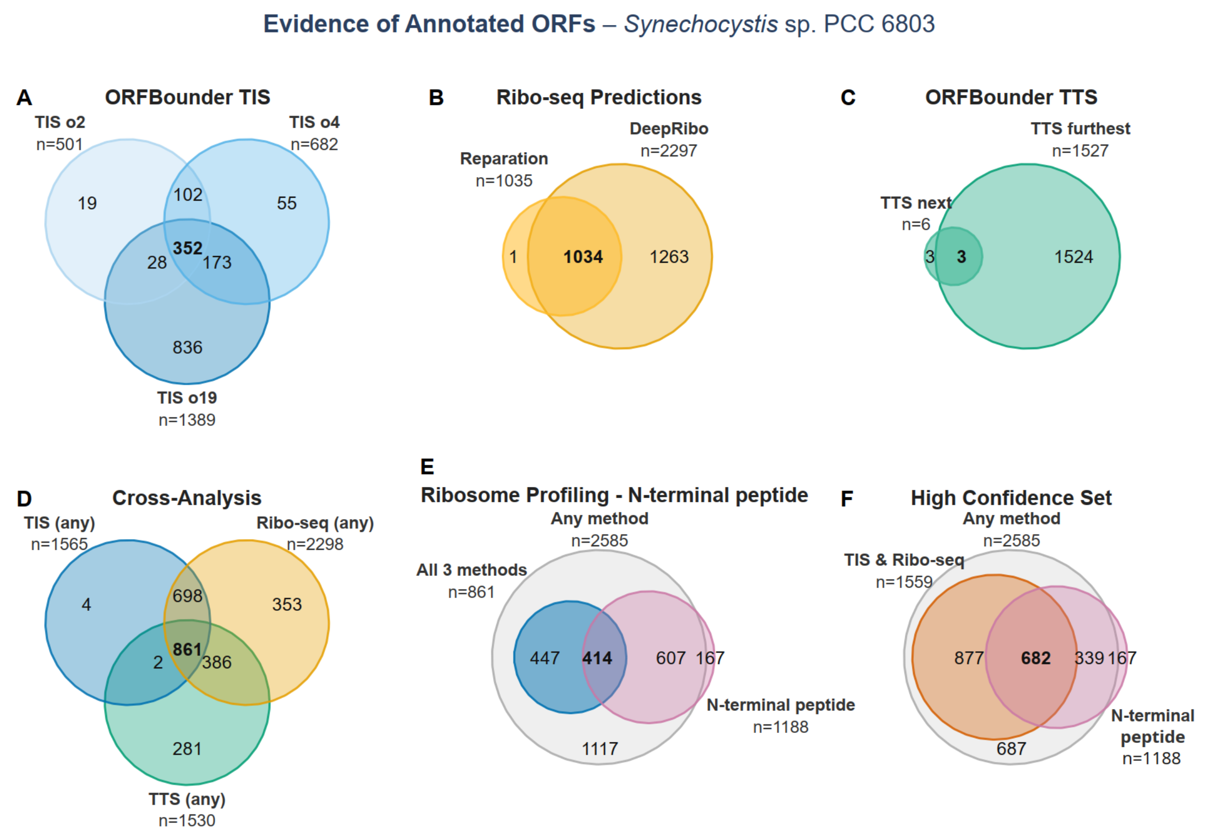


**Figure S11. Multi-evidence overview of annotated ORFs in *Synechocystis* 6803.** All comparisons are based on the Kazusa 2009 genome annotation. Filters match those applied during candidate selection for manual curation. Translational efficiency (TE) is calculated as the ratio of Ribo-seq RPKM to RNA-seq RPKM and reflects the translational output per mRNA molecule. The $\log_2$ fold-change ($\log_2$FC) is calculated as the ratio of site-specific ribosome footprint coverage (TIS-Ribo-seq or TTS-Ribo-seq) to total Ribo-seq coverage. Unless stated otherwise, all per-replicate thresholds (RPKM, $\log_2$FC) are met if at least one replicate passes.

**A.** Overlap among translons detected by ORFBounder from TIS-Ribo-seq data at three offset positions relative to the annotated start codon (+2, +4, +19). Each ORF was required to meet: RPKM ≥ 40 in both TIS-Ribo-seq and Ribo-seq libraries, $\log_2$FC ≥ 1, and average TE ≥ 0.288.

**B.** Overlap among translons predicted by the Ribo-seq–based tools DeepRibo and Reparation. Both sets share a common base filter (RPKM ≥ 40 in both TIS-Ribo-seq and Ribo-seq libraries, average TE ≥ 0.288). ORFs were additionally required to reach a DeepRibo score ≥ −3.009 or a Reparation probability ≥ 0.5, respectively.

**C.** Overlap among translons detected by ORFBounder from monosome TTS-Ribo-seq data using two start-codon assignment modes: "Next" (ORF defined using the nearest upstream in-frame TIS relative to the detected termination site) and "Furthest" (ORF defined using the most distant upstream in-frame TIS). Each ORF was required to meet: RPKM ≥ 10 in both TTS-Ribo-seq and Ribo-seq libraries, $\log_2$FC ≥ 1, and average TE ≥ 0.15. Only monosome libraries were used for TTS-Ribo-seq analysis.

**D.** Cross-method comparison of the three evidence classes. Each circle represents the union of ORFs from the respective evidence class: TIS-Ribo-seq (union of positions +2, +4, +19; panel A), Ribo-seq predictions (union of DeepRibo and Reparation; panel B), and TTS-Ribo-seq (union of Next and Furthest modes; panel C).

**E.** Relationship between ribosome profiling evidence and proteomics-based evidence that identifies a start. "Union" contains all ORFs recovered by any method shown in (D); "Intersect" contains ORFs jointly supported by all three evidence classes. "N-terminal peptide" indicates ORFs for which an N-terminal peptide was identified by mass spectrometry–based proteomics, providing direct confirmation of *in vivo* translation and correct start-site usage.

**F.** Translons classified into confidence levels based on the number of independent supporting evidence types. The outermost set (Any method) encompasses all ORFs recovered by any ribosome profiling approach. The intermediate tier (TIS-Ribo-seq + Ribo-seq predictions) reflects ORFs supported by both start-site evidence classes. The innermost tier (All three methods) requires concordance across TIS-Ribo-seq, Ribo-seq predictions, and TTS-Ribo-seq. Overlap with the N-terminal peptide circle (as defined in E) shows how mass spectrometry evidence distributes across confidence tiers. The 167 cases with an N-terminal peptide only indicate they were only found by proteomics (no Ribo-seq signal).

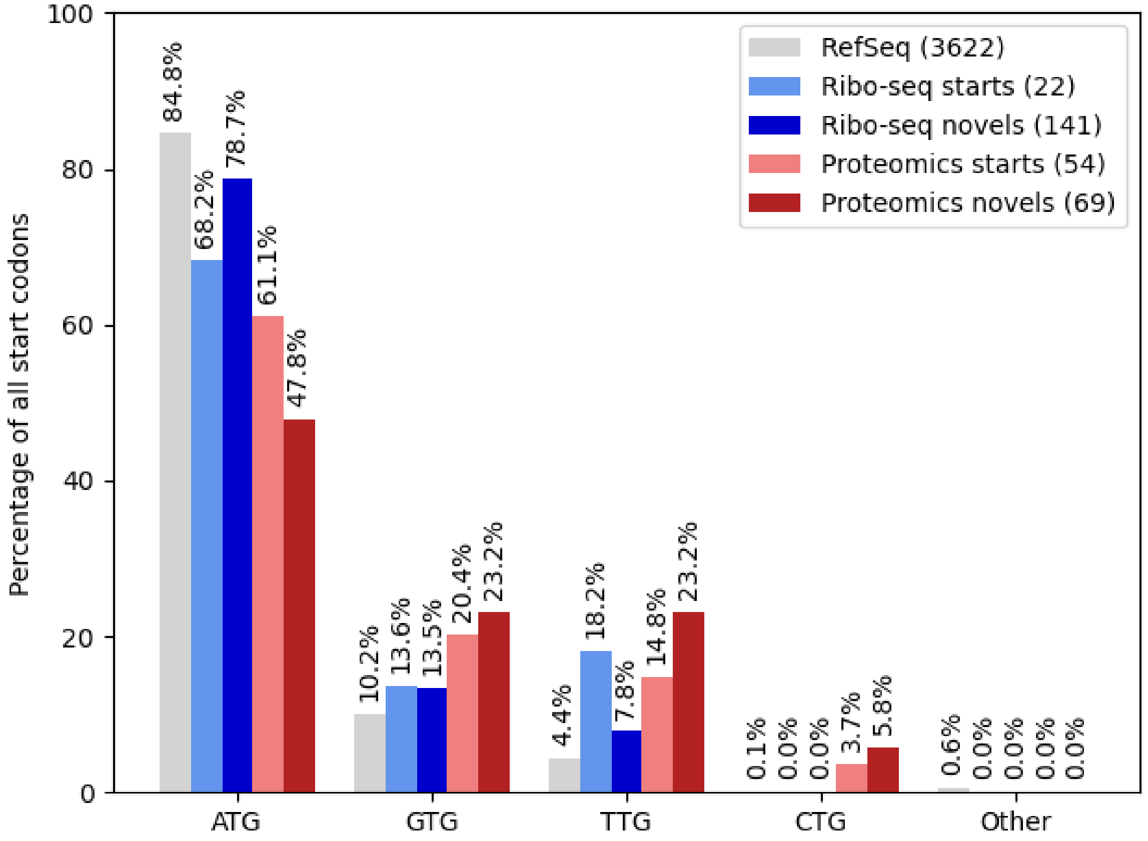


**Figure S12. Start codon distribution of annotated genes compared to new starts and novel gene candidates identified with riboproteogenomics.** Annotated genes (grey) preferentially start with the canonical ATG, while a minority starts with alternative start codons, mostly GTG, followed by TTG. This trend is similar among the Ribo-seq detected new starts (light blue) and novel genes (dark blue), whereas for proteogenomics based new starts (light red) and novel genes (dark red) a slight enrichment in TTG and CTG alternative start codons at the expense of canonical ATG starts can be observed.

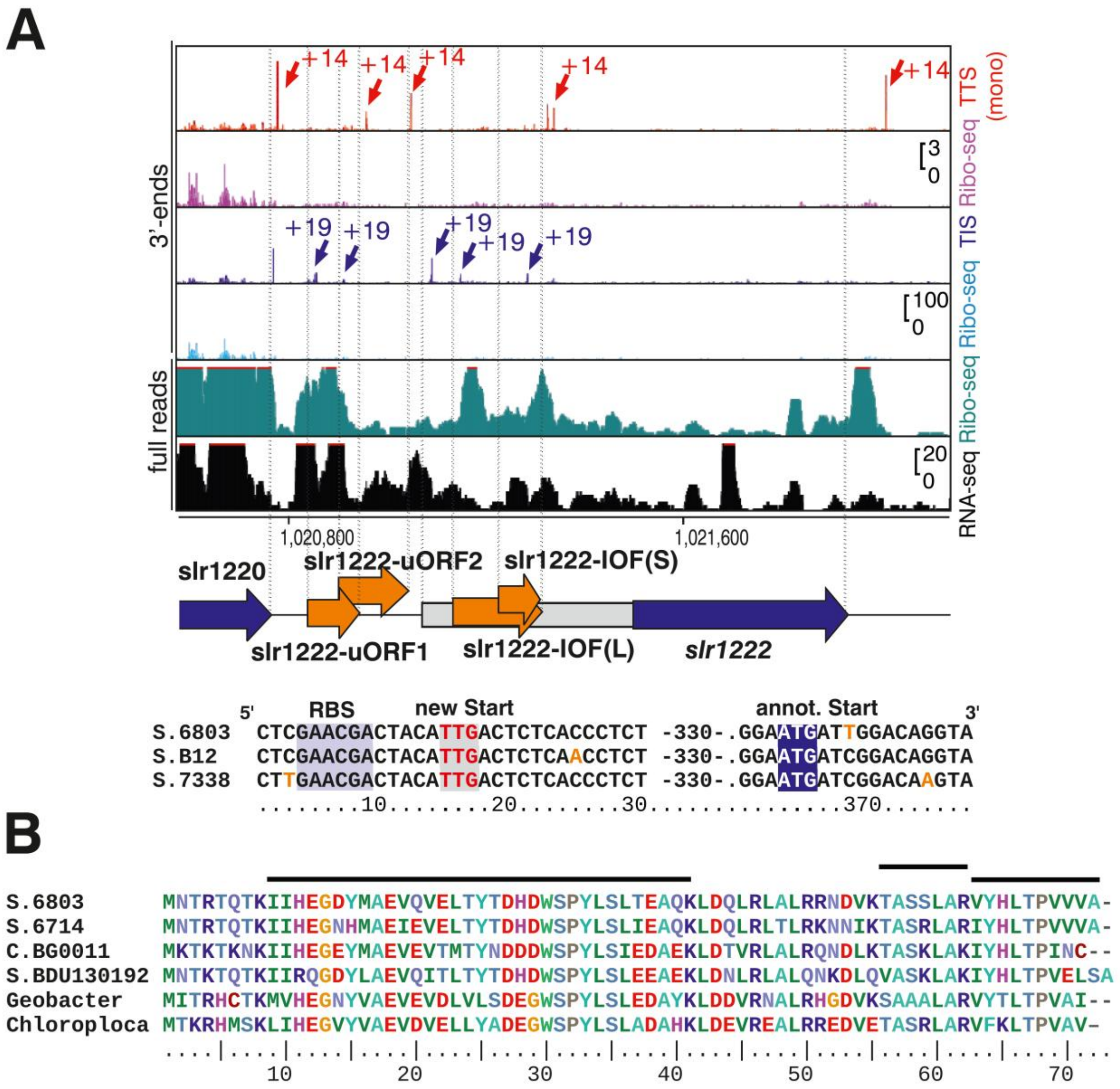


**Figure S13. Ribosome profiling aids the re-annotation of genes and the discovery of novel SEPs.**

**A.** TIS and TTS profiling data re-annotate the start codon of *slr1222* and suggest the existence of four potential novel sORFs in the associated region. TIS profiling (purple) in combination with Ribo-seq coverage (turquoise, full reads) supports an N-terminal extension by 116 aa for Slr1222, yielding 321 aa in total. DNA sequence alignments of the translation initiation regions from different *Synechocystis* strains indicated the annotated and new start codons (lower panel). In combination with Ribo-seq & TTS profiling coverage (red track on top), TIS profiling supports the detection of two internal out-of-frame, overlapping sORFs, IOF(L) and (S) (60 aa and 15 aa) within the N-terminal extension of Slr1222, and two un-annotated upstream sORFs (slr1222-uORF1 and 2, 35 aa and 44 aa, respectively) in the intergenic region between *slr1220* and *slr1222*. Arrows in purple indicate the respective TIS signals, arrows in red TTS signals. Note, that a transmembrane domain was predicted in Slr1222-IOF(L).

**B.** Homologs of Ncr1610-sORF1 exist in several cyanobacteria and many other species. The multiple sequence alignment shows in addition to Ncr1610-sORF1 putative homologs from the cyanobacteria *Synechocystis* 6714 (AIE73341), *Cyanothece* sp. BG0011 (WP_107667291), *Synechococcus* sp. BDU 130192 (WP_099239310), the Pseudomonadota *Geobacter* sp. DSM 9736 (WP_088535776) and the Chloroflexota ‘Candidatus *Chloroploca mongolica*‘ (WP_135477305). Horizontal lines above the alignment indicate peptides found for Ncr1610-sORF1. This figure extends **Figure 5F**.

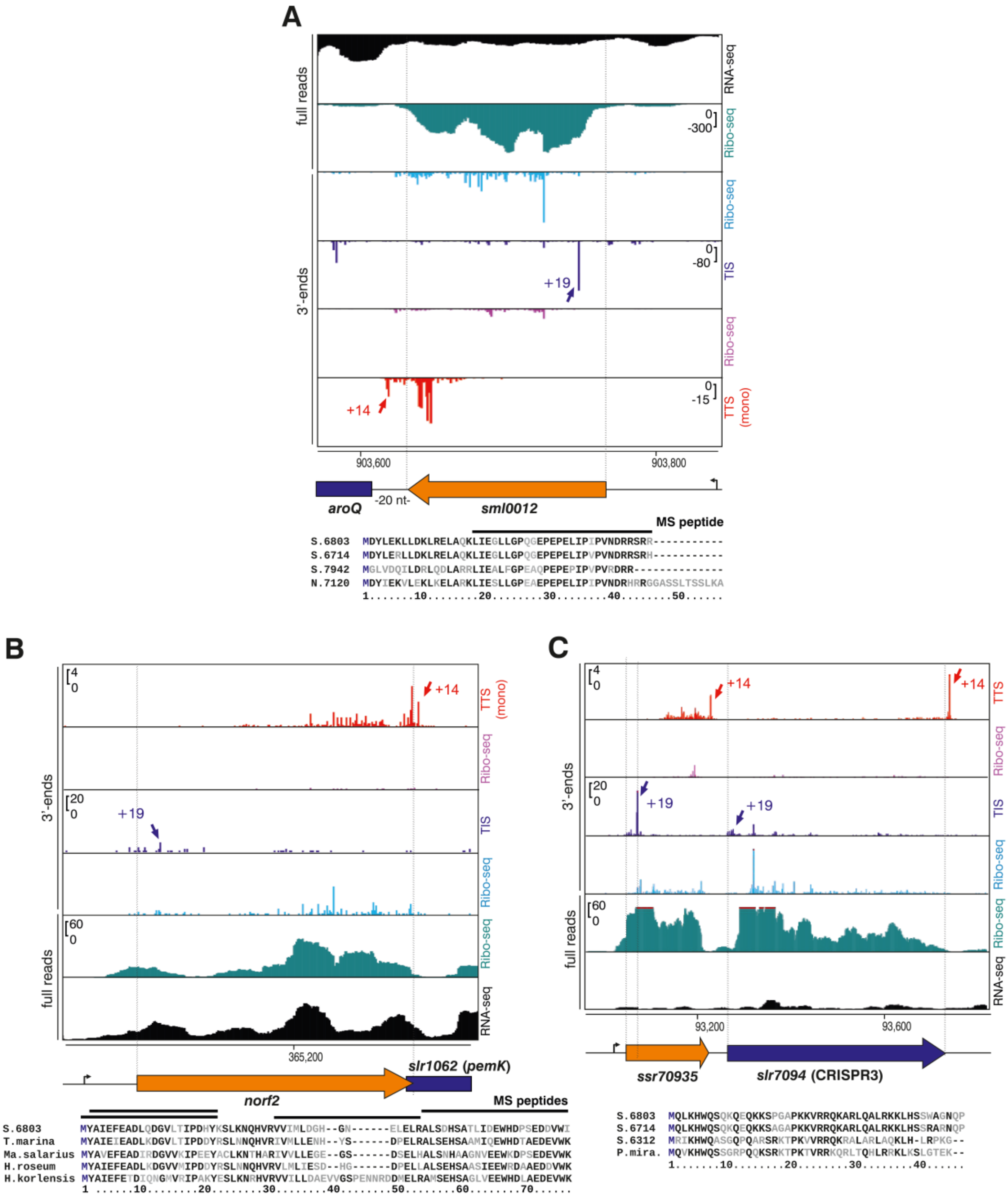


**Figure S14. Small proteins encoded in sRNAs and unannotated genome regions.** **A.** Sml0012 is a 45 aa protein with homologs in many other cyanobacteria. Read coverage from RNA-seq (top, black), Ribo-seq (second panel from top, turquoise), TIS- (purple) and TTS-Ribo-seq (red) indicates translation of the *sml0012* ORF. The TIS-Ribo-seq 3' ends map 19 nt downstream the start codon and the TTS-Ribo-seq 3' ends 14 downstream the stop codon (blue and red arrows). The Ribo-seq read coverage is enriched for the coding part, in contrast to the RNA-seq coverage, which starts at the transcription start site (bent arrow) 199 nt upstream the start codon. Transcription is contiguous with the downstream located gene *aroQ* encoding the enzyme 3-dehydroquinate dehydratase that leads via the shikimate pathway to the synthesis of aromatic amino acids Y, F and Q. The multiple sequence alignment shows the comparison to Sml0012 homologs from *Synechocystis* 6714 (gene D082_00160), *Synechococcus elongatus* sp. PCC 7942 (accession WP_011242938), and *Nostoc* sp. PCC 7120 (gene *asr2781*). The location 17 to 25 nt upstream of *aroQ* is conserved in these strains. The horizontal line above the alignment indicates a peptide found for Sml0012. The protein was moreover validated after FLAG tagging by Western blotting (**Figure 6A**).

**B.** Norf2 was previously hypothesized as protein-coding gene *norf2* ("novel ORF 2") based on its transcription and the conserved reading frame [11]. Here, its translation as a 68 aa protein is supported by the Ribo-seq, TIS- and TTS-Ribo-seq data, by the identification of 4 peptides in the standard iPtgxDB that cover 86% of it (horizontal lines), and by Western blotting after adding a C-terminal FLAG tag (**Figure 6A**). We found no homologs for Norf2 in any other cyanobacteria, but some closely related proteins exist in gamma-proteobacteria. The alignment shows the comparison to homologs from *Thiocapsa marina* (WP_007191346), *Marinobacter salarius* (WP_269400048), *Halochromatium roseum* (WP_201215208), and *Halomonas korlensis* (WP_089797407). These proteins are annotated as hypothetical proteins; however, the gene location in the *Synechocystis* 6803 genome has been characterized as a genomic island [12], and a role in a toxin-antitoxin system with the overlapping gene *slr1062* encoding a PemK-type toxin[13] as an antitoxin is likely.

**C.** The here defined gene *ssr70935* is located directly downstream the CRISPR3 array, between *slr7093* and *slr7094*, on plasmid pSYSA[14], upstream of *slr7094* and *slr7095* with which it is cotranscribed as part of TU7087[15], hence forming a tricistronic operon. Its translation is supported by strong Ribo-seq, TIS- and TTS-Ribo-seq coverage and by Western blotting after FLAG tagging (**Figure 6A**). In the lower part, Ssr70935 is compared to homologs from *Synechocystis* 6714 (gene D082_40650), *Synechocystis* sp. PCC 6312 (AFY60879), *Synechococcus* sp. PCC 6312 (gene Syn6312_1729), and *Petrachloros mirabilis* (WP_275072620).

.

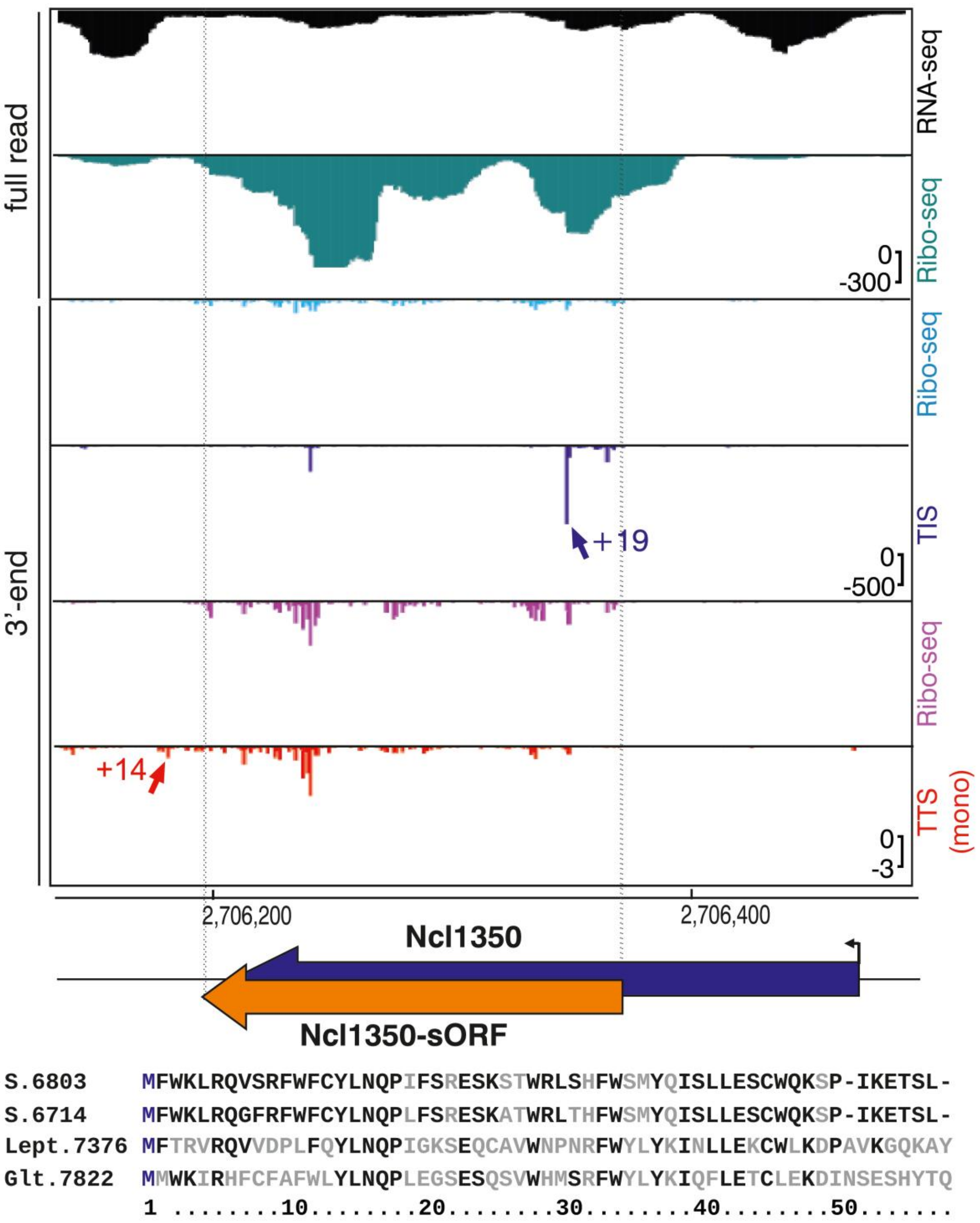


**Figure S15.** Ncl1350 was originally classified as an sRNA[15], but actually contains a translated sORF of 57 aa as indicated by the coverage from Ribo-seq, TIS- and TTS-Ribo-seq that was after SPA tagging validated by Western blotting (**Figure 6A**). The TSS was mapped to position 2,706,438 yielding a 5'UTR of 71 nt. Homologues of similar size exist in many different cyanobacterial species, here shown in the alignment with putative proteins from *Synechocystis* 6714, *Leptolyngbya* sp. PCC 7376 and *Gloeothece verrucosa* PCC 7822. These proteins are annotated (accession numbers AIE75908, AFY36466 and ADN14821.

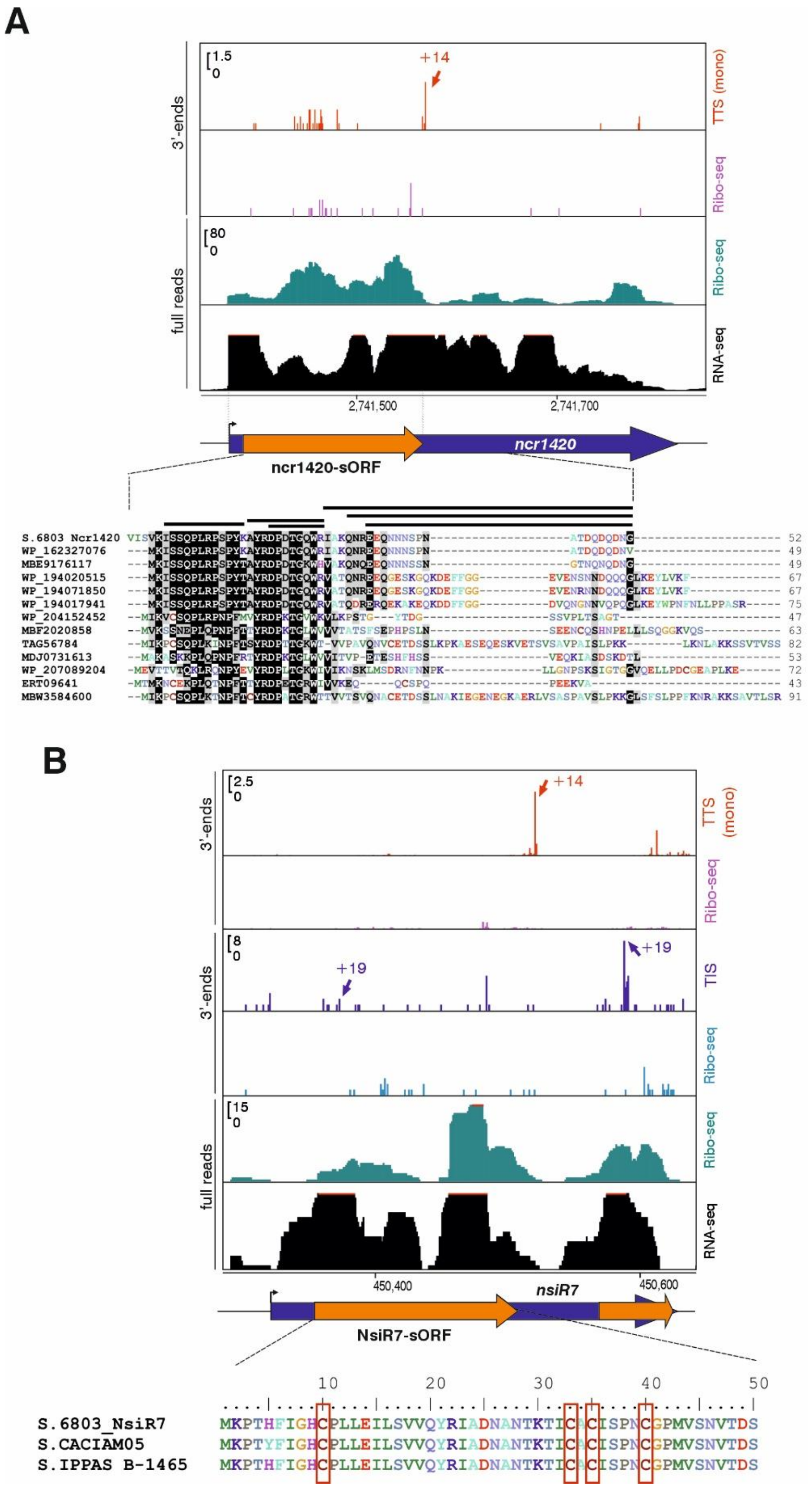


**Figure S16. Homologs of small proteins encoded on previously defined sRNAs in *Synechocystis* 6803, as predicted on the basis of TIS- and TTS-Ribo-seq data.** **A.** The sRNA Ncr1420 (synonym SyR21) was originally classified as a non-coding RNA within TU2899[15], but actually is translated into a 52 aa SEP, supported by TTS data with a 3’ end offset of 14 nt (red arrow) and confirmed after SPA tagging by Western blotting **(Figure 6A**). Moreover, six different peptides were identified in the iPtgxDB. The gene was annotated in RefSeq 2022 as SGL_RS19480. Homologues of similar size exist in several different cyanobacteria, the alignment shows the comparison to homologs from *Synechocystis* sp. CACIAM 05 (WP_162327076), *Synechocystis salina* LEGE 06155 (MBE9176117), *Synechocystis salina_1* (WP_194020515), *Synechocystis* sp. LEGE 06083 (WP_194071850), *Synechocystis salina_2* (WP_194017941), *Leptolyngbya* sp. CCY15150 (WP_204152452), *Hydrococcus* sp. (C42_A2020_068), *Hydrococcus* sp. C42_A2020_068 (MBF2020858), *Oscillatoriales cyanobacterium* (TAG56784), *Crocosphaera* sp. (MDJ0731613), *Phormidium pseudopriestleyi* (WP_207089204), *Lyngbya aestuarii* BL J (ERT09641), and *Cyanobacteria bacterium* 0813 (MBW3584600). Several more putative homologs can be identified by TBlastN.

**B.** The sRNA NsiR7 was originally classified as non-coding RNA Ncr0210 in TU3138[15]. It belongs to the 33 most abundant ncRNAs in *Synechocystis* 6803[16] and is strongly induced under nitrogen starvation and was therefore renamed to nitrogen starvation induced RNA 7 (NsiR7), controlled by the transcription factor NtcA[17]. It contains a translated sORF encoding a 50 aa protein, confirmed by Western blotting after 3xFLAG tagging **(Figure 6A**). Homologs of similar size exist in several different *Synechocystis*, an identical protein is encoded in *Synechocystis* sp. IPPAS B-1465. The alignment shows the comparison to homologs in *Synechocystis* sp. CACIAM 05 (CA05) and *Synechocystis* S. IPPAS B-1465. A downstream encoded very short sORF potentially encodes a protein of 17 aa (NsiR7-dORF), for which homologs can be predicted in *Synechocystis* sp. IPPAS B-1465 and *Synechocystis* sp. CACIAM 05, but not in *Synechocystis* 6714. Four conserved cysteine residues are boxed. Both sORFs are supported by TIS data with a 3’end offset of 19 nt and a clear TTS signal with a 3’ end offset of 14 nt in case of NsiR7 (red arrow).

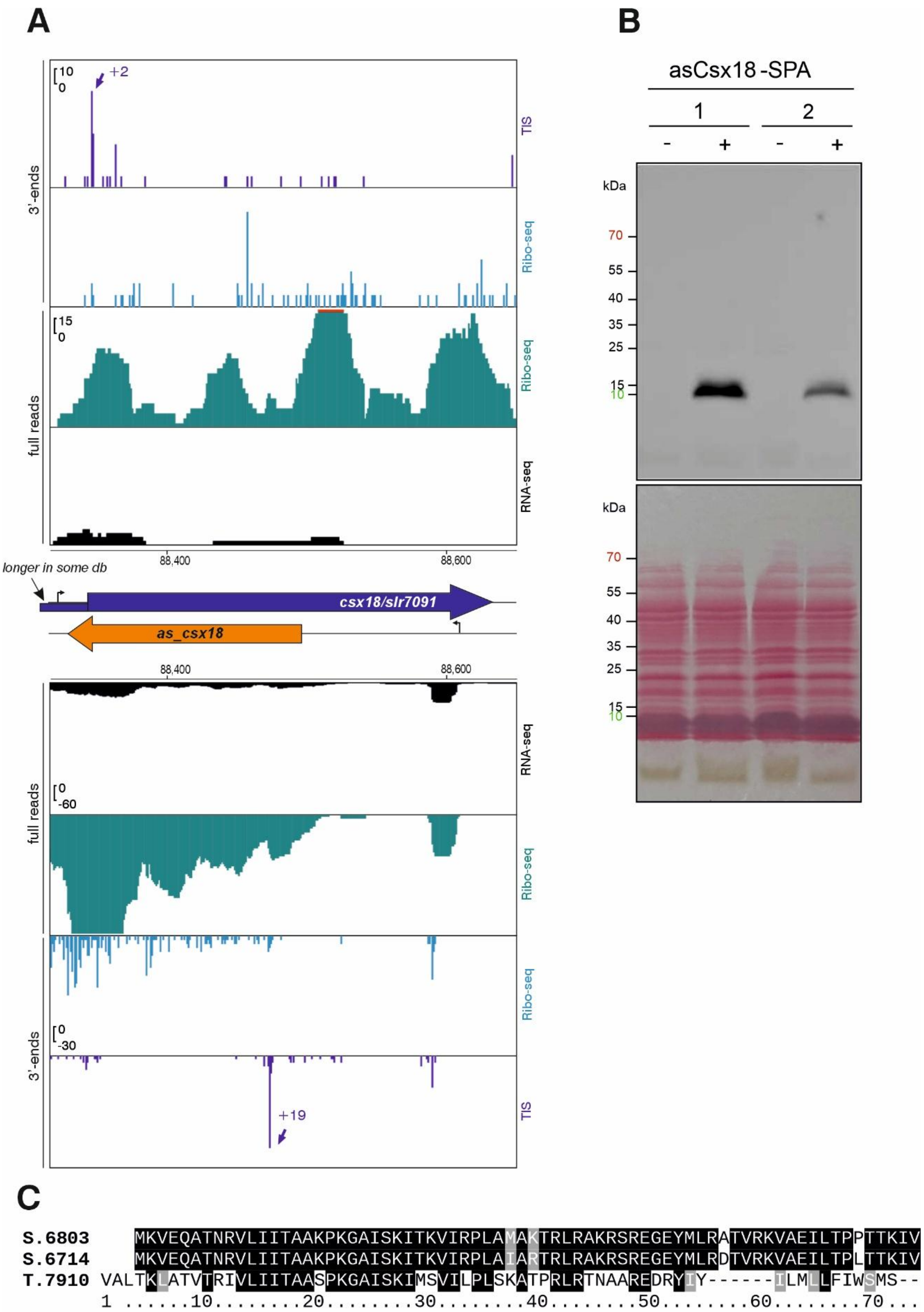


**Figure S17. Coding potential in the asRNA as_csx18 to gene *slr7091* encoding a CRISPR-associated Csx18 protein. A.** Both DNA strands are expressed, supported by RNA-seq (black), Ribo-Seq (turquoise), and TIS-Ribo-seq (purple) datasets. The characteristic TIS-Ribo-seq signals 3’ of the start codons were mapped at position +19 for the sORF in *as_csx18* and at position +2 (which was one of the offsets found for the TIS-Ribo-seq 3′ end signals, cf. **Figure 1F**) for *slr7091*. Note, that Slr7091 is annotated in the KZS, RefSeq and GBK databases as a 122 aa protein, but according to our data it is translated from a shorter ORF encoding a 96 aa protein, consistent with the Prodigal prediction. This example illustrates how experimental data overlaid on top of integrated annotations from different genome centers can be used to consolidate the annotation differences. **B.** Western Blot verification of the dual SPA-tagged asCsx18 protein expressed from plasmid pVZ322 under control of the $P_{petE}$ promoter, 22 h after induction by copper addition to 1.25 µM (+). Samples without induction served as negative controls (-). The labels ‘1’ and ‘2’ refer to biological replicates. In this gel, ten µg protein was loaded per lane, separated on a 12% SDS-PAA gel. The antiserum was M2 anti-FLAG HRP (Sigma-Aldrich A8592). The lower panel shows the ponceau red-stained membrane. **C.** The sequence alignment shows possible homologs of the 70 aa asCsx18 protein encoded by the *as_csx18* in *Synechocystis* 6714 and *Tolypothrix* sp. PCC 7910 identified by TBlastN in antisense orientation to their respective *csx18* genes. The respective ORFs are shown in full lengths. The gene in *Synechocystis* 6714 is located on plasmid pSYLA[18] and according to dRNA-seq data it is also associated with an asRNA[19]. All three *csx18* loci are located close to a CRISPR array (distance of three genes or less).

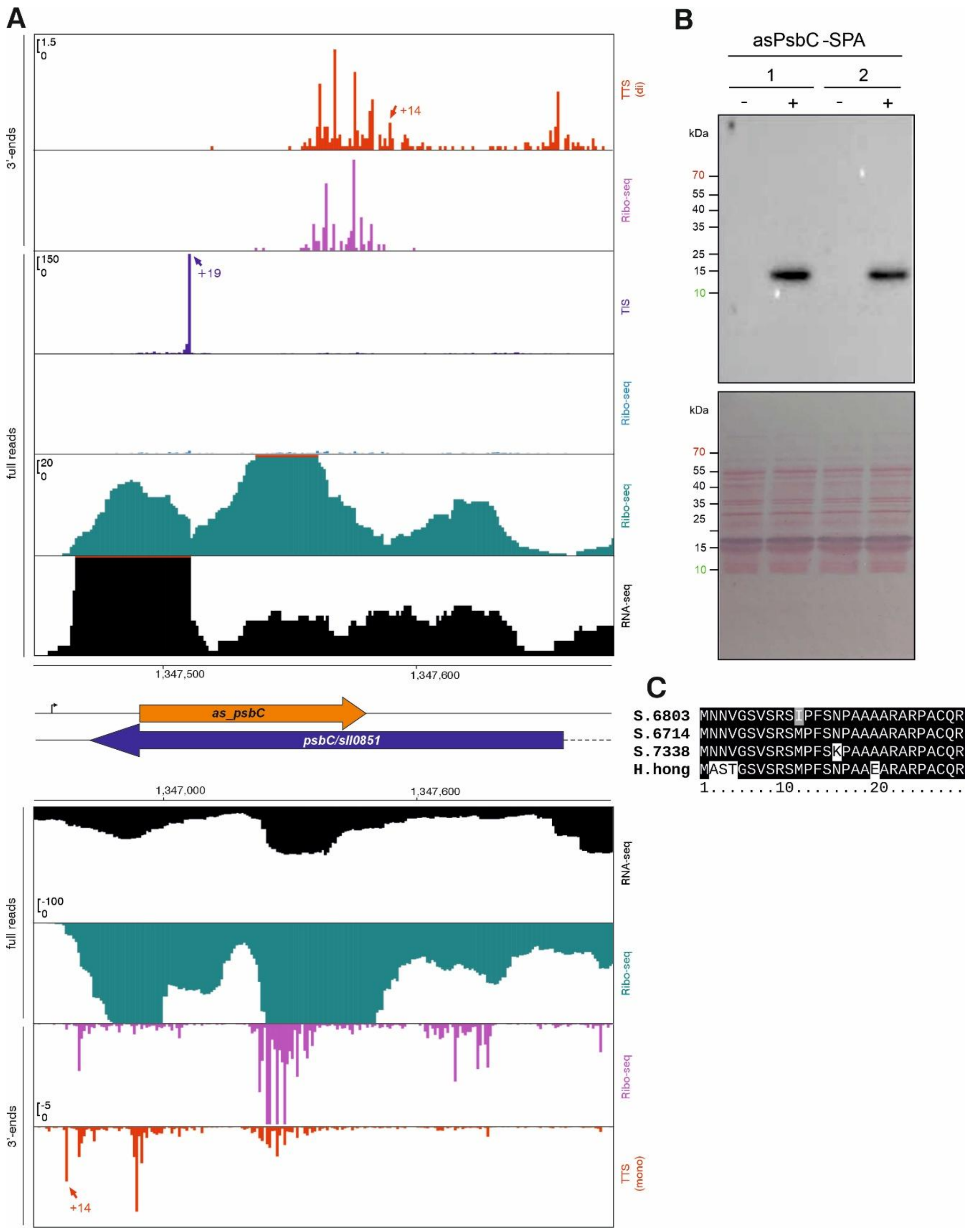


**Figure S18. Coding potential for a 28 aa protein in the asRNA *as_psbC* to gene *sll0851* encoding the photosystem II protein PsbC/CP43.** Both DNA strands are expressed, supported by RNA-seq (black), Ribo-Seq (turquoise), TIS-Ribo-seq (purple) and TTS-Ribo-seq (red) datasets. The characteristic TIS-Ribo-seq signal located 19 nt 3' of the start codon in the *as_psbC* transcript were mapped at position +19 (out of the shown region for *psbC*). TTS-Ribo-seq signals located 14 nt 3' of the stop codons in the as_psbC transcript and *psbC* mRNA are labeled by red arrows.

**B.** Western Blot verification of the SPA-tagged asPsbC protein expressed from plasmid pVZ322 under control of the $P_{petE}$ promoter, 22 h after induction by copper addition to a final concentration of 1.25 µM (+). Samples without induction served as negative controls (-). The labels ‘1’ and ‘2’ refer to biological replicates. In this gel, 3 µg protein was loaded per lane, separated on a 15% SDS-PAA gel. The antiserum was M2 anti-FLAG HRP (Sigma-Aldrich A8592). The lower panel shows the ponceau red-stained membrane.

**C.** The sequence alignment shows possible homologs of the asPsbC protein in *Synechocystis* strains 6714 and 7338, and in *Halomicronema hongdechloris* C2206 identified by TBlastN in antisense orientation to their respective *psbC* genes. However, it is unknown if corresponding asRNAs exist in these strains and the respective ORFs are also longer than the displayed segments.

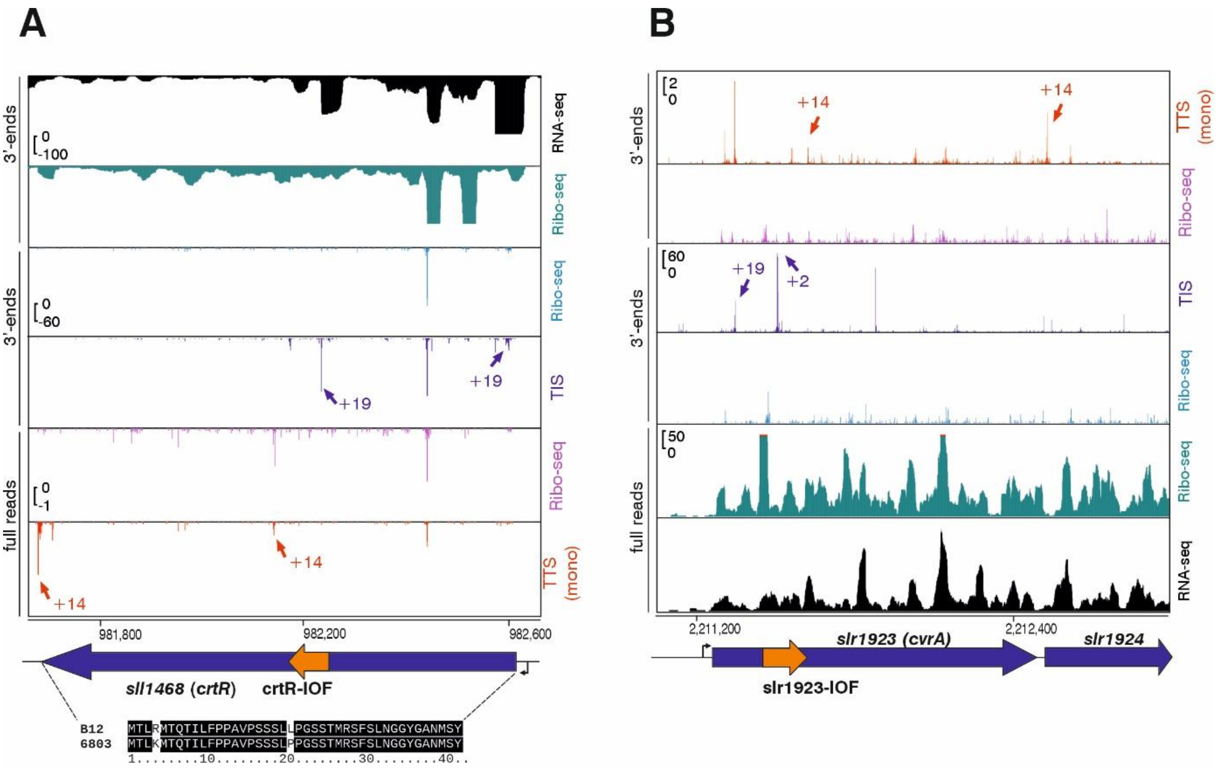


**Figure S19. sORFs identified in mRNAs out-of-frame**. Alternative, gene-internal out-of-frame (IOF) start sites were predicted for **A.** *crtR* and **B.** *cvrA* on the basis of TIS-RiboSeq (blue) and TTS-RiboSeq (red) data. TIS-RiboSeq reads 3'ends accumulated close to the potential internal start codon with a 3'end offset of 19 nt for the main genes and the IOF in *crtR/sll1468*, while a 3'end offset of 2 nt was detected for the IOF in *cvrA*/*slr1923*. Both main genes encode important enzymes of pigment biosynthesis. CrtR is the beta-carotene hydroxylase converting beta-carotene to zeaxanthin[20], CvrA is the vinyl reductase essential for the conversion from divinylchlorophyll(ide) to normal chlorophyll(ide)[21,22]. The sORF in *crtR* can also be identified in *Synechocystis* sp. B12, but generally these IOF sORFs are not conserved. In all panels, the Ribo-seq read coverage (turquoise) is enriched for the coding segments, in contrast to the RNA-seq coverage (black), which starts at the position of the transcription start site (bent arrow). The here identified sORFs are indicated by orange arrows.

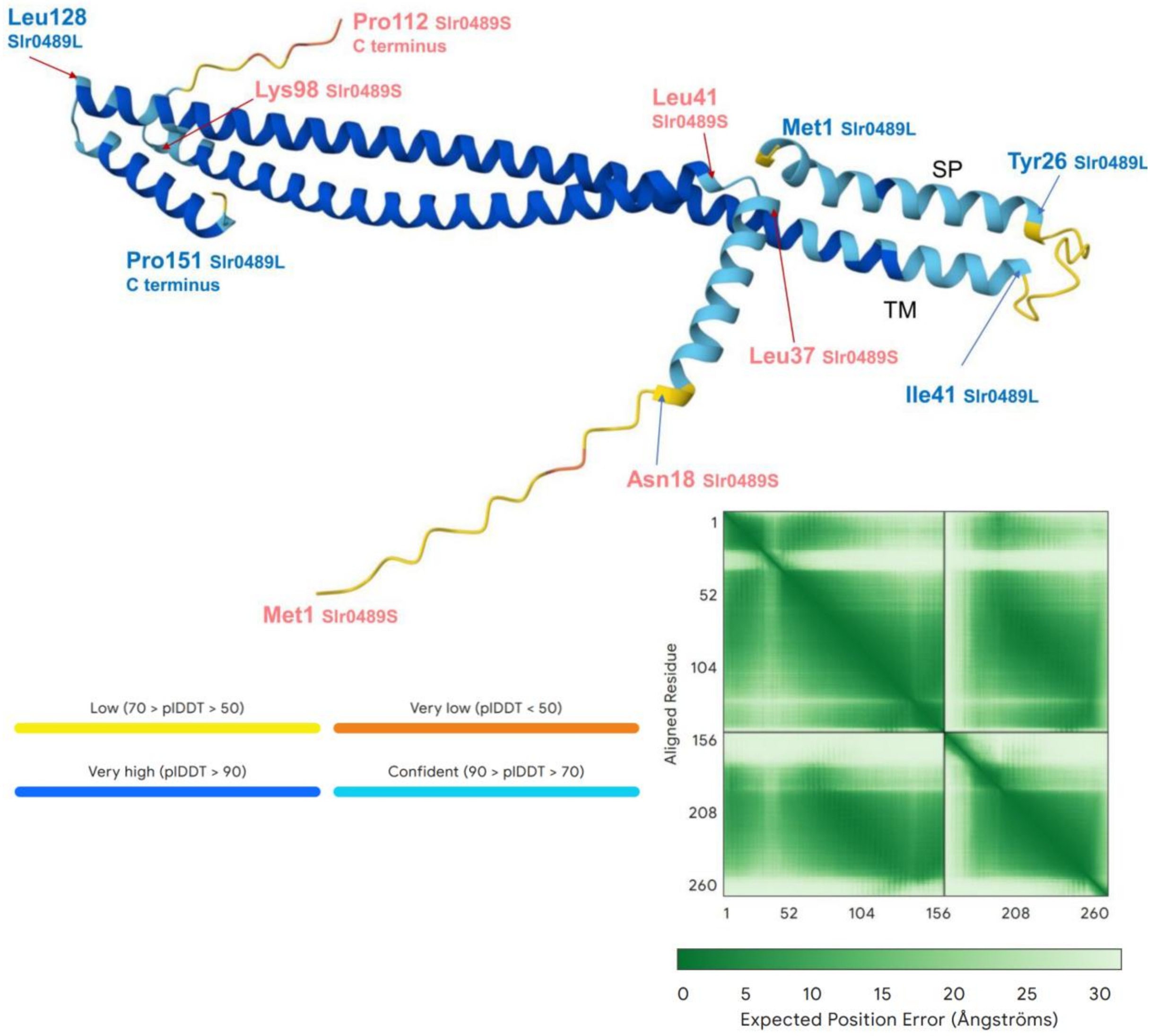


**Figure S20. Alphafold prediction of the Slr0489L/Slr0489S heterodimer**. Alphafold models long helical regions in both proteins which align over ~50% of their respective lengths. Several residues are given for orientation for Slr0489L (light blue) and Slr0489S (rose). The two predicted membrane-spanning regions in Slr0489L are indicated (TM1 and TM2). Support scores were ipTM = 0.58 and pTM = 0.59, note that support was with ipTM = 0.65 and pTM = 0.67 better for the heterooctamer (4 copies of Slr0489L and Slr0489S each). Here, the heterodimer is shown for better clarity.

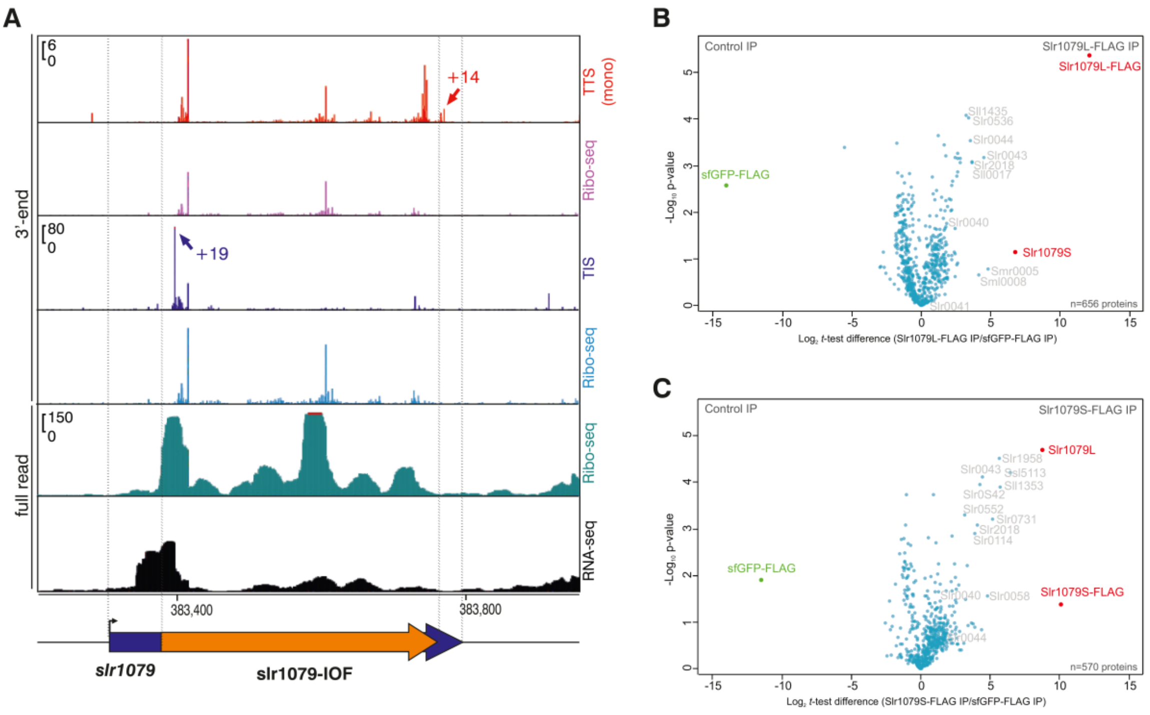


**D**

>Slr1079L
MASFLALLPRSLTTFLYAVAALLRFYGNIDTTPIPRIPLTIFGWSFLAFTLGTAALLVNLGLEWNTGNRSRNREIETRERETRRD
NLADEERNRASEEREKADRERDRADQERNRADQERKRAASRARIQNRGFVLQTRYQLAPSPEARATLIDFLSFLQEYGE

>Slr1079S
MATSTPPPSPESPSPSLDGASWRSRWAQRLSWLISALSGTLEIEAETERLKQENAKLDEIILQTKKETERLRKEKKPIEKEIE
QIKKETEQIRKENERLAVLESKIEDLFSKHGINLPLAQKPEQP

**E**

>Slr1082L
MASFLALLPRSLTTFLYAVAALLRFYGNIDTTPFQLIPLTILDWSFLAFTLGTAALLVNLGLEWNSGNRSRNRADQERDRTDQE
RDRADQERQRAARRARIQNRFFLRPTRYQLTPSPDTRAALIDFLSFLQEYGE

>Slr1082S
MATSTPPPSNSSPSPSSTGVSWRSRWAQRLSWLISALSGTLEIEAETEQIRKETEQIKKETEQIRKDNERLAVLESKIDSSFD
QHGINLPLAQTPEQP

**Figure S21. A protein Slr1079S is encoded by a long IOF (orange arrow) within the *slr1079* reading frame (blue arrow). A.** Read coverage from Ribo-seq and RNA-seq libraries is shown in the lower 2 panels. The Ribo-seq read coverage is clearly higher at the beginning of the IOF segment. Mapped signals from TIS-Ribo-seq at the characteristic position 19 nt 3' of the respective start codon (two central panels including control), and 14 nt 3' of the respective stop codon from TTS-Ribo-seq (two upper panels including control) are highlighted by the blue and red arrows.

**B.** Co-IP analysis of Slr1079L. The volcano plot shows the protein interaction partners of Slr1079L-3xFLAG identified by mass spectrometry compared to sfGFP-3xFLAG. Triplicate samples were analyzed (**Figure S25**). The two most enriched proteins,

Slr1079L and Slr1079S are colored red, the locus IDs are given for co-enriched proteins in grey. For the complete list of 656 detected proteins and further details, see **Table S10**.

**C.** Co-IP analysis of Slr1079S. Details as in panel B and in **Figure S26**. For the complete list of 570 detected proteins, see **Table S11.** The results of an analogous co-IP experiment for *slr0489* encoding Slr0489L/S are shown in **Figure 7D**. **D.** Sequences of Slr1079L and Slr1079S. The black horizontal lines indicate proteogenomically detected peptides for both proteins directly proving their simultaneous presence in the cells. **E.** Sequences of Slr1082L and Slr1082S. Proteogenomically detected peptides were only found for Slr1082S (horizontal lines).

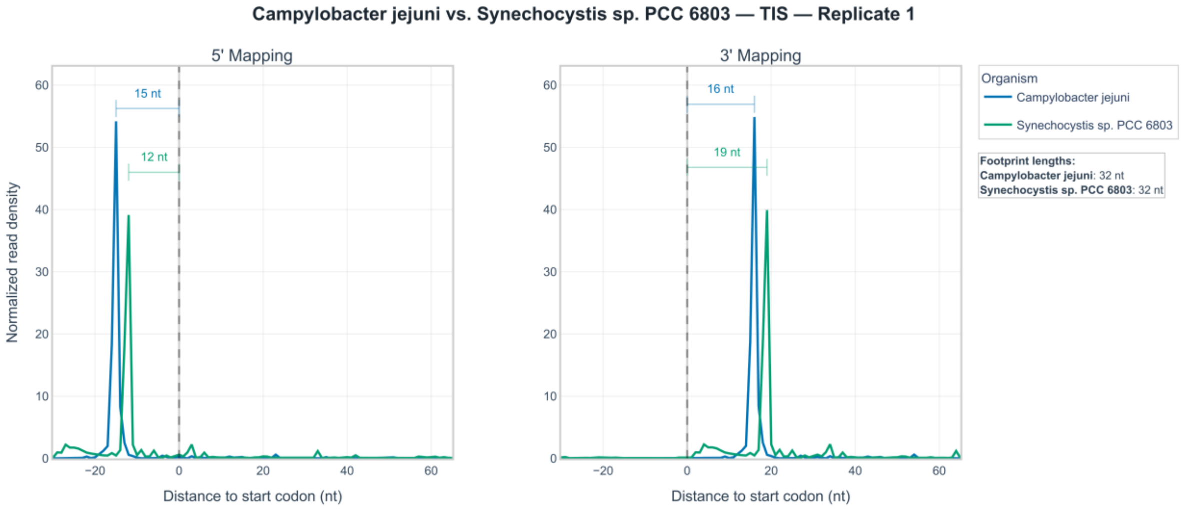


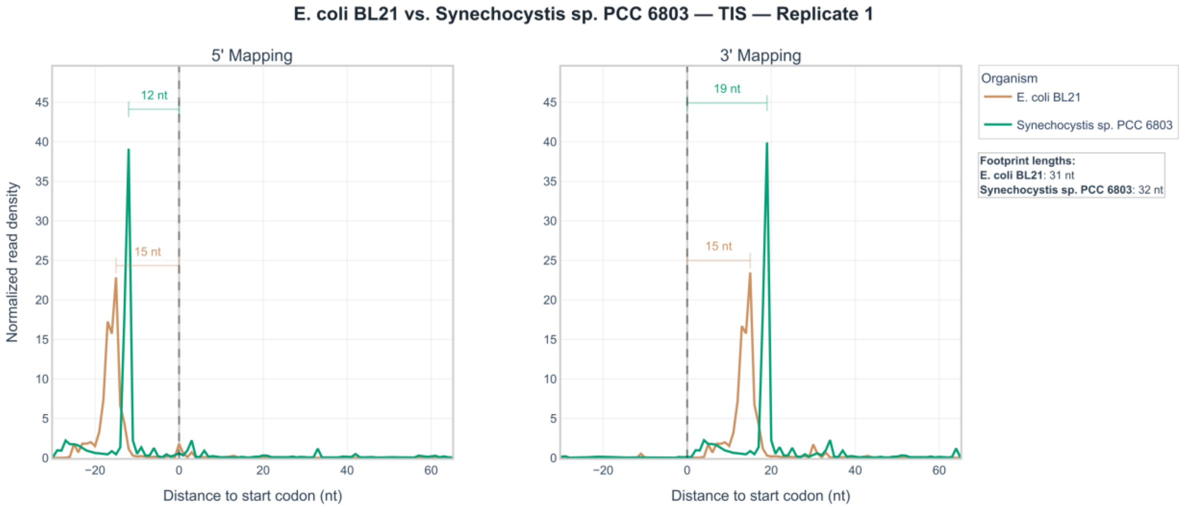


**Figure S22. Metagene analysis comparing 5' and 3' mapped ribosome footprints from *Synechocystis* 6803 with previously published results from *E. coli*[23] and *Campylobacter*[7].** All datasets were reprocessed through the same pipeline used in this study, and footprint lengths were selected to match those reported in the original publications. Read density is shown for each position from 30 nt upstream to 65 nt downstream of every annotated ORF start, with total read counts per position normalized to the window length. Position 0 represents the first nucleotide of each start codon. Additional comparisons including further replicates and filtering options can be accessed here: [https://www.bioinf.uni-freiburg.de/~ribobase/synechocystisbrowsepublic/data/metagene/comparison_metagene_between_organisms.html](https://www.bioinf.uni-freiburg.de/~ribobase/synechocystisbrowsepublic/data/metagene/comparison_metagene_between_organisms.html).

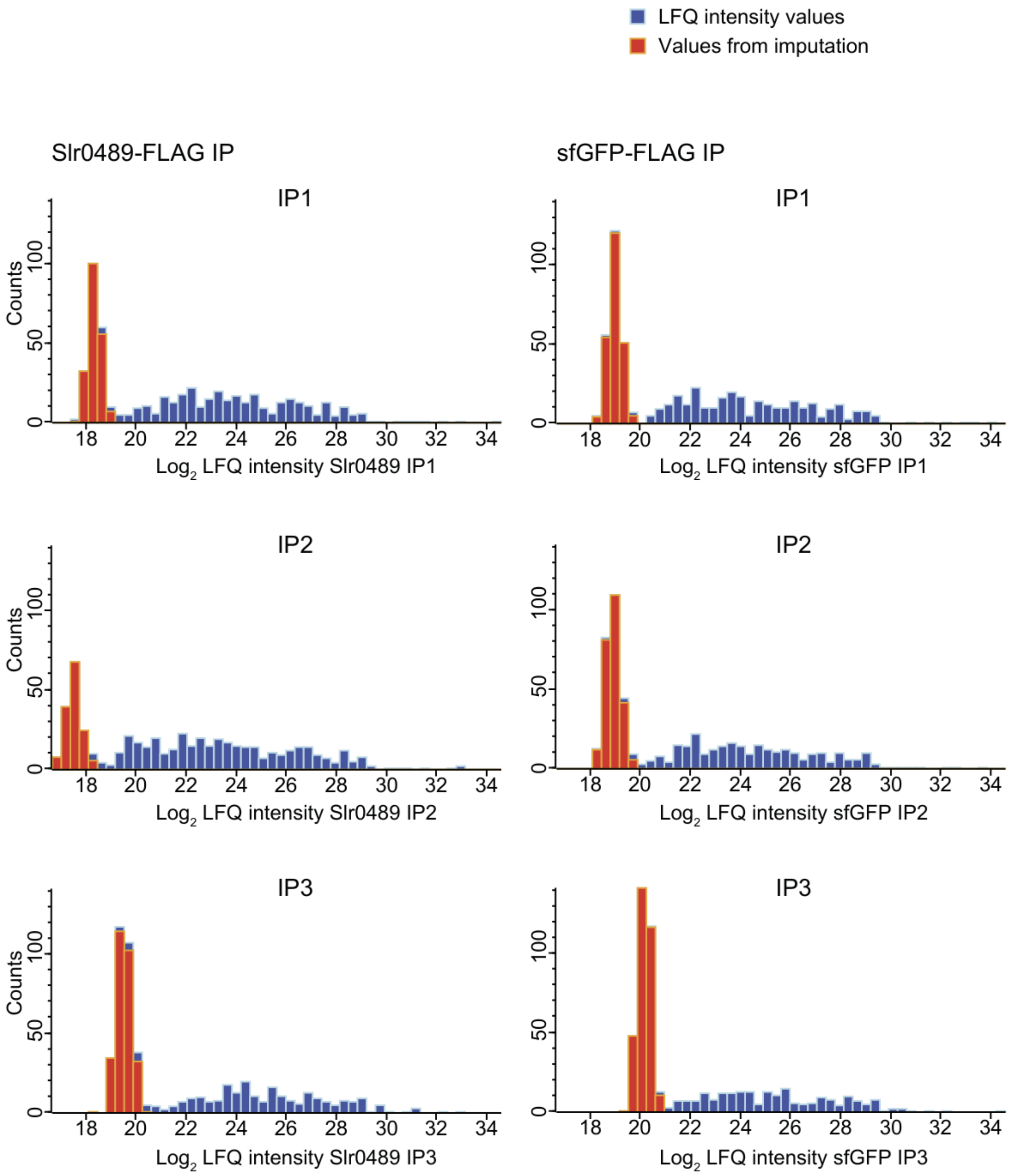


**Figure S23. Histograms of the LFQ data distribution for the co-IP with Slr0489L-3xFLAG compared to sfGFP-3xFLAG.** This figure extends **Figure 7D** and **Table S8**.

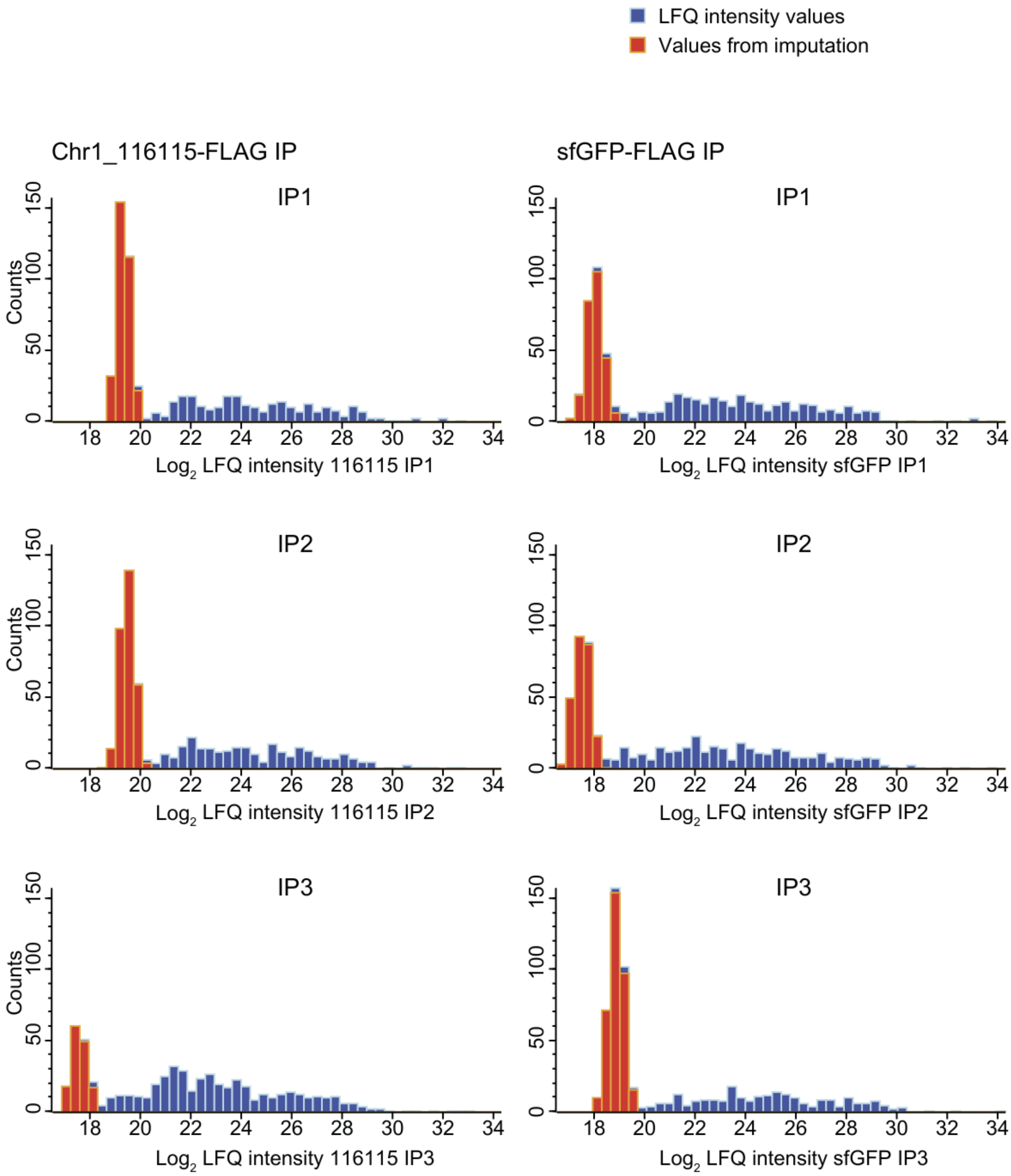


**Figure S24. Histograms of the LFQ data distribution for the co-IP with Slr0489S-3xFLAG compared to sfGFP-3xFLAG.** This figure extends **Figure 7E** and **Table S9**.

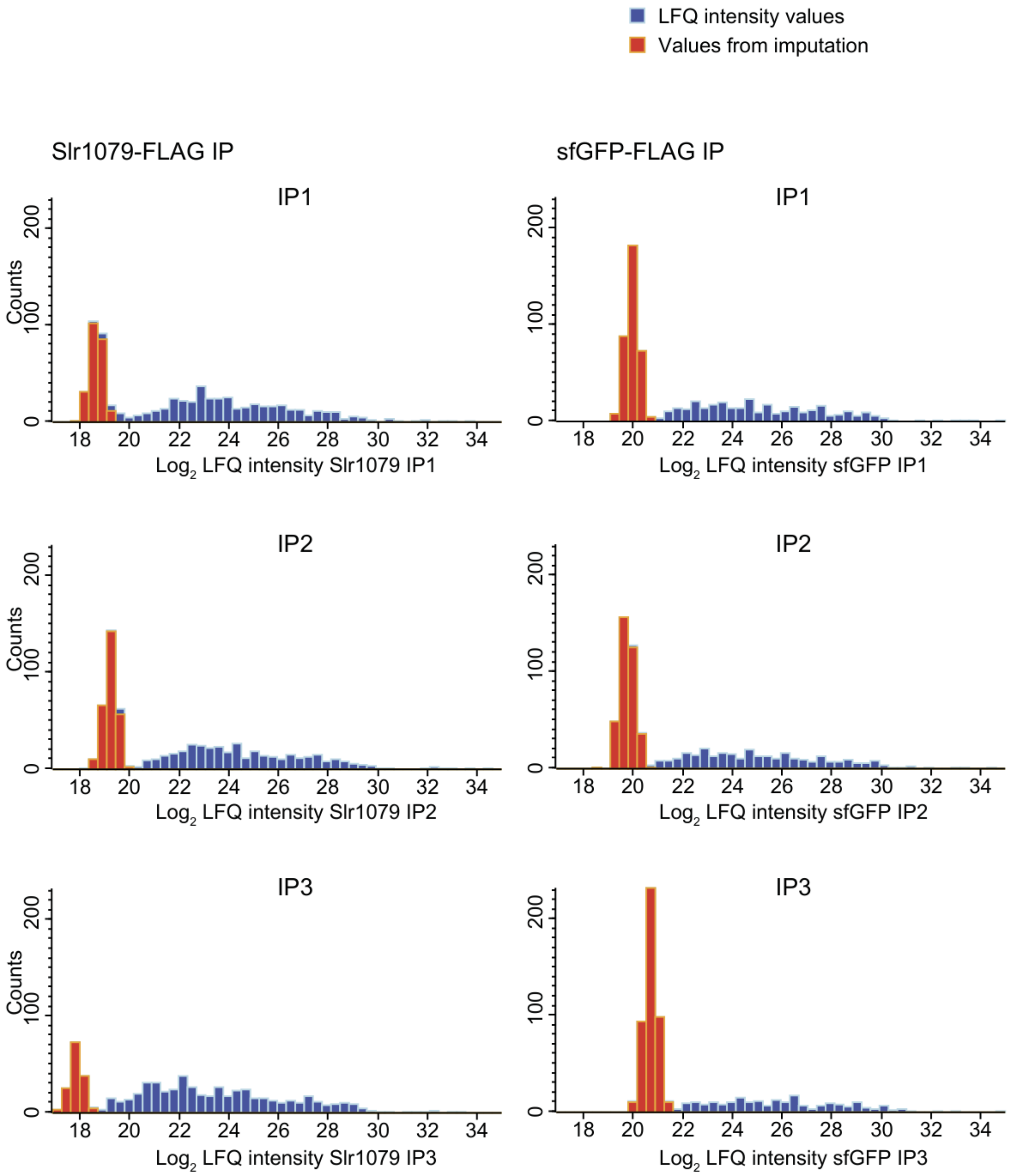


**Figure S25. Histograms of the LFQ data distribution for the co-IP with Slr1079L-3xFLAG compared to sfGFP-3xFLAG.** This figure extends **Figure S21B** and **Table S10**.

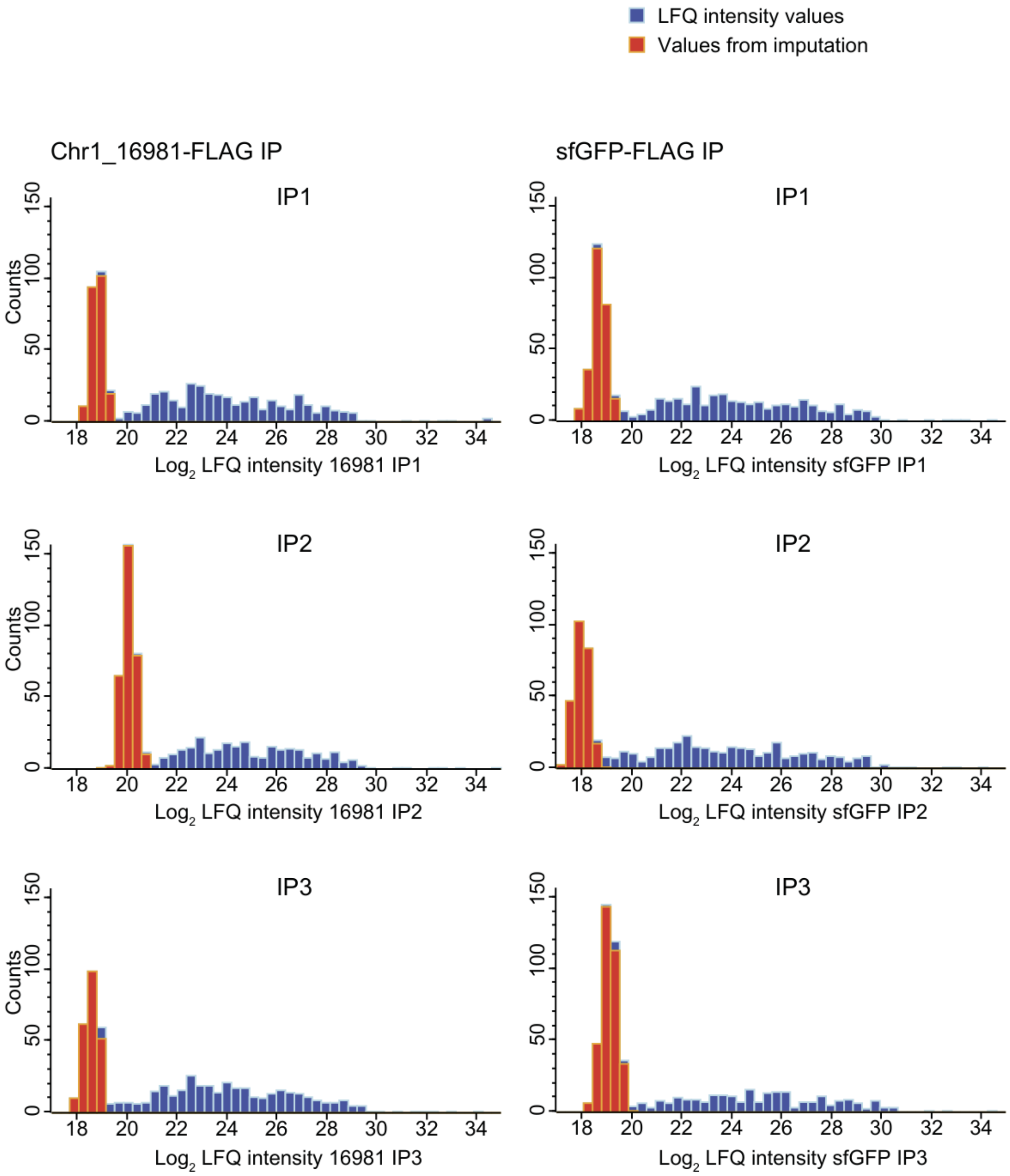


**Figure S26. Histograms of the LFQ data distribution for the co-IP with Slr1079S-3xFLAG compared to sfGFP-3xFLAG.** This figure extends **Figure S21C** and **Table S11**.

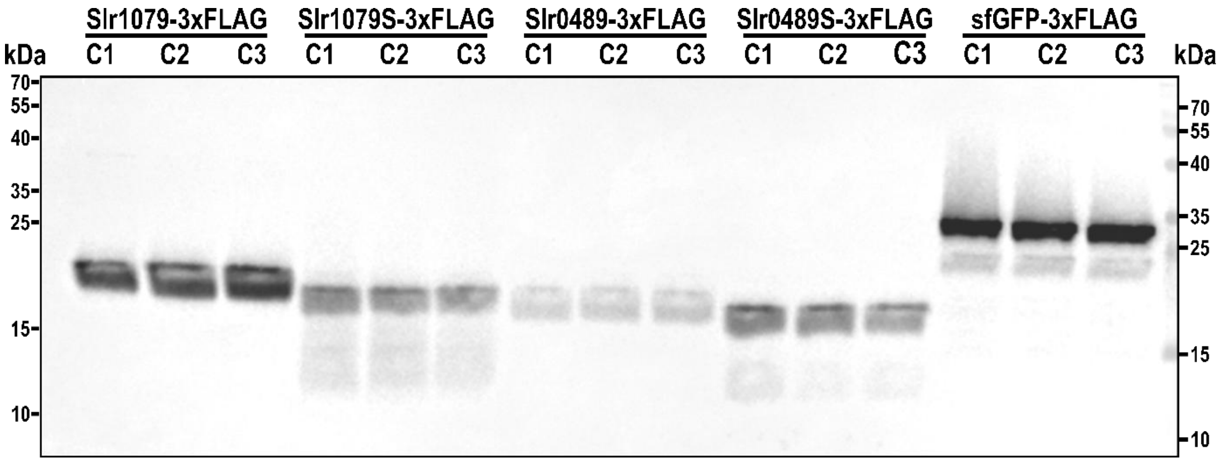


**Figure S27. Western blot of overexpressed Slr0489L/S and Slr1079L/S.** The genes *slr0489* and *slr1079* were amplified and cloned from the *Synechocystis* 6803 genome and inserted into expression vector pVZ322s under the control of the $P_{rha}$ promoter and with a sequence encoding a C-terminal 3xFLAG tag. In parallel, the IOF for each gene was cloned with their native 5´UTRs into the same expression vector. The constructs were transformed into wild-type *Synechocystis* 6803. A construct encoding tagged sfGFP was used as control. The expression was induced upon addition of 0.6 mg/mL L-rhamnose for 24 h before harvesting. Co-immunoprecipitation assays were performed and bound proteins were eluted with 100 ng/µL 3xFLAG-peptide. For each elution, 20 µL were loaded on a 12% SDS-PAA gel. The 3xFLAG tagged proteins were revealed by Western Blotting with M2 anti-FLAG antiserum. Each strain was tested in triplicates (C1-3). This figure extends **Figure 7D/E** and **Figure S21B, C**.

### Supplementary Code Description

#### Metagene Profiling Scripts

Repository: https://github.com/RickGelhausen/Synechocystis_pub_scripts_2025

The repository contains all code that was used to conduct the metagene analyses performed for the study. It also contains the scripts used for detailed analysis of specific peaks in the metagene profile and the creation of weblogos.

*Details on installation and usage are available in the provided repository.*

A genome-wide analysis of ribosome occupancy was conducted using TIS-Ribo-seq and TTS-Ribo-seq. To this end, all annotated coding sequences (CDS) were collected, and windows consisting of upstream and downstream regions around each annotated start codon (or stop codon, respectively) were defined. For each position in the analyzed window, reads occupying that position were counted for each annotated CDS.

The total number of reads per position was subsequently reported and plotted in a metagene graph. To generate a high-confidence set of genes, multiple filters were applied. These filters removed genes in close-proximity (≤50 nt apart), genes below a coverage threshold of 10 RPKM and genes that fell below a minimum length cutoff (i.e., genes shorter than the downstream window for start codons or shorter than the upstream windows for stop codons). However, we performed a parallel metagene analysis focusing on 39 short ORFs (≤50 aa) that confirmed that translation initiation patterns in these shorter sequences did not differ substantially from those observed in longer ORFs (compare **Figure 1E** and **Figure S2**).

Extreme outliers in the metagene graph were investigated by determining exact read numbers per gene involved in the offending position. These genes were subsequently inspected in the genome browser and filter out if they were clear sequencing artifacts.

To assess sequence motifs in the 5'-UTR regions of annotated start codons, all annotated CDS contributing to positions with the highest ribosome occupancy (at offset of +2, +4, and +19) were collected, and sequence windows were extracted.

Motif plots were then generated using weblogo (3.7.9).

### ORFBounder

Repository: https://github.com/RickGelhausen/ORFBounder

ORFBounder is a computational tool for calling peaks from ribosome profiling (Ribo-seq) data and subsequently determining open reading frame (ORF) boundaries. The tool identifies translation initiation sites (TIS) and translation termination sites (TTS) by integrating metagene profiling results with coverage information derived from ribosome-protected footprints. The tool calculates coverage directly from input alignment files, supporting three-prime, five-prime, centered, and global read mapping strategies, each available as raw or normalized (min, mil) coverage tracks.

*Details on installation and usage are available in the provided repository.*

***Analysis description***

The analysis proceeds through the following steps:

**Step 1 — Metagene-guided offset selection.** Prior to running the tool, optimal P-site offsets are determined from metagene profiling output for each mapping and normalization combination of interest. These offsets are provided as input parameters to the tool. To determine these offsets, the scripts described in the section „**Metagene Profiling Scripts**" can be used.

**Step 2 — Candidate codon identification.** All potential start codons (for TIS analysis) or stop codons (for TTS analysis) are extracted from the genome annotation. Each codon position is expanded into a ±2 nt window (5 nt total) around the first nucleotide of the codon.

**Step 3 — Coverage quantification.**

Alignment files are read and each read is adjusted by the appropriate P-site offset to align the previously determined candidate codon windows with the expected ribosome footprint signal. This coverage data is stored in a positional dictionary.

Overlaps between this coverage signal and each candidate codon interval are computed. At each overlapping position, if the coverage exceeds a user-defined read count threshold (default: 5), the signal is added to the cumulative peak height for the corresponding codon interval. Where multiple codon intervals overlap a single genomic position, the coverage is attributed to all overlapping intervals, as unambiguous assignment is not possible.

**Step 4 — ORF prediction.** For each candidate codon with a non-zero peak height, the pipeline searches for the corresponding in-frame boundary codon: for TIS predictions, the nearest in-frame stop codon downstream of the start codon is identified; for TTS predictions, the nearest in-frame start codon upstream of the stop codon is identified, alternatively the furthest in-frame start codon prior to the next in-frame stop codon can be used instead. The resulting predicted ORFs are reported in both GFF and XLSX format, with optional separate GFF files generated for each gene type to facilitate genome browser inspection.

**Step 5 — Cross-sample integration.** When multiple Ribo-seq, RNA-seq, TIS, or TTS datasets are analyzed, individual result tables are consolidated into a single spreadsheet. ORFs predicted across multiple samples are merged into a single entry, with sample-specific peak heights reported in separate columns. The output includes comprehensive annotation for each predicted ORF, including gene type, genomic coordinates, strand, locus tag (if annotated), codon count, the 15 nt sequence upstream of the start codon, full nucleotide and amino acid sequences, and peak height values. If both Ribo-Seq and TIS or TTS-Seq data are available, $\log_2$ fold-change columns are automatically added for matching condition-replicate pairs.

**Step 6 — Expression quantification.** Finally, expression metrics are computed for all predicted ORFs across all libraries. Read counts are obtained using featureCounts (Subread package) and used to calculate reads per kilobase per million mapped reads (RPKM) values. For samples where matched Ribo-seq and RNA-seq data are available, translational efficiency (TE) is calculated as the ratio of Ribo-seq to RNA-seq RPKM. These values are added to the consolidated results table.

### Supplementary References


1. Gessulat, S., Schmidt, T., Zolg, D.P., Samaras, P., Schnatbaum, K., Zerweck, J., Knaute, T., Rechenberger, J., Delanghe, B., Huhmer, A., et al. (2019). Prosit: proteome-wide prediction of peptide tandem mass spectra by deep learning. Nat. Methods *16*, 509–518. https://doi.org/10.1038/s41592-019-0426-7.

2. Hallgren, J., Tsirigos, K.D., Pedersen, M.D., Armenteros, J.J.A., Marcatili, P., Nielsen, H., Krogh, A., and Winther, O. (2022). DeepTMHMM predicts alpha and beta transmembrane proteins using deep neural networks. Preprint at bioRxiv, https://doi.org/10.1101/2022.04.08.487609.

3. Peng, Z., Xiao, Q., and Wan, C. (2025). Identification and validation of SmORF-encoded peptides by genomics and proteomics in five cyanobacteria. J. Proteome Res. *24*, 5818−5829. https://doi.org/10.1021/acs.jproteome.5c00685.

4. Baers, L.L., Breckels, L.M., Mills, L.A., Gatto, L., Deery, M.J., Stevens, T.J., Howe, C.J., Lilley, K.S., and Lea-Smith, D.J. (2019). Proteome mapping of a cyanobacterium reveals distinct compartment organization and cell-dispersed metabolism. Plant Physiol. *181*, 1721–1738. https://doi.org/10.1104/pp.19.00897.

5. Oliveira, P., Martins, N.M., Santos, M., Pinto, F., Büttel, Z., Couto, N.A., Wright, P.C., and Tamagnini, P. (2016). The versatile TolC-like Slr1270 in the cyanobacterium *Synechocystis* sp. PCC 6803. Environ. Microbiol. *18*, 486–502. https://doi.org/10.1111/1462-2920.13172.

6. Stringer, A., Smith, C., Mangano, K., and Wade, J.T. (2022). Identification of novel translated small open reading frames in *Escherichia coli* using complementary ribosome profiling approaches. J. Bacteriol. *204*, e00352-21. https://doi.org/10.1128/JB.00352-21.

7. Froschauer, K., Svensson, S.L., Gelhausen, R., Fiore, E., Kible, P., Klaude, A., Kucklick, M., Fuchs, S., Eggenhofer, F., Yang, C., et al. (2025). Complementary Ribo-seq approaches map the translatome and provide a small protein census in the foodborne pathogen *Campylobacter jejuni*. Nat. Commun. *16*, 3078. https://doi.org/10.1038/s41467-025-58329-w.

8. Qeli, E., and Ahrens, C.H. (2010). PeptideClassifier for protein inference and targeted quantitative proteomics. Nat. Biotechnol. *28*, 647–650. https://doi.org/10.1038/nbt0710-647.

9. Omasits, U., Varadarajan, A.R., Schmid, M., Goetze, S., Melidis, D., Bourqui, M., Nikolayeva, O., Québatte, M., Patrignani, A., Dehio, C., et al. (2017). An integrative strategy to identify the entire protein coding potential of prokaryotic genomes by proteogenomics. Genome Res. *27*, 2083–2095. https://doi.org/10.1101/gr.218255.116.

10. Hadjeras, L., Heiniger, B., Maaß, S., Scheuer, R., Gelhausen, R., Azarderakhsh, S., Barth-Weber, S., Backofen, R., Becher, D., Ahrens, C.H., et al. (2023). Unraveling the small proteome of the plant symbiont *Sinorhizobium meliloti* by ribosome profiling and proteogenomics. microLife *4*, uqad012. https://doi.org/10.1093/femsml/uqad012.

11. Mitschke, J., Georg, J., Scholz, I., Sharma, C.M., Dienst, D., Bantscheff, J., Voß, B., Steglich, C., Wilde, A., Vogel, J., et al. (2011). An experimentally anchored map of transcriptional start sites in the model cyanobacterium *Synechocystis* sp. PCC6803. Proc. Natl. Acad. Sci. USA *108*, 2124–2129. https://doi.org/10.1073/pnas.1015154108.

12. Kopf, M., Klähn, S., Pade, N., Weingärtner, C., Hagemann, M., Voß, B., and Hess, W.R. (2014). Comparative genome analysis of the closely related *Synechocystis* strains PCC 6714 and PCC 6803. DNA Res. *21*, 255–266. https://doi.org/10.1093/dnares/dst055.

13. Kopfmann, S., Roesch, S.K., and Hess, W.R. (2016). Type II toxin-antitoxin systems in the unicellular cyanobacterium *Synechocystis* sp. PCC 6803. Toxins *8*, 228.1-228.23. https://doi.org/10.3390/toxins8070228.

14. Scholz, I., Lange, S.J., Hein, S., Hess, W.R., and Backofen, R. (2013). CRISPR-Cas systems in the cyanobacterium *Synechocystis* sp. PCC6803 exhibit distinct processing pathways involving at least two Cas6 and a Cmr2 protein. PLoS ONE *8*, e56470. https://doi.org/10.1371/journal.pone.0056470.

15. Kopf, M., Klähn, S., Scholz, I., Matthiessen, J.K.F., Hess, W.R., and Voß, B. (2014). Comparative analysis of the primary transcriptome of *Synechocystis* sp. PCC 6803. DNA Res. *21*, 527–539. https://doi.org/10.1093/dnares/dsu018.

16. Kopf, M., and Hess, W.R. (2015). Regulatory RNAs in photosynthetic cyanobacteria. FEMS Microbiol. Rev. *39*, 301–315. https://doi.org/10.1093/femsre/fuv017.

17. Giner-Lamia, J., Robles-Rengel, R., Hernández-Prieto, M.A., Muro-Pastor, M.I., Florencio, F.J., and Futschik, M.E. (2017). Identification of the direct regulon of NtcA during early acclimation to nitrogen starvation in the cyanobacterium *Synechocystis* sp. PCC 6803. Nucleic Acids Res. *45*, 11800–11820. https://doi.org/10.1093/nar/gkx860.

18. Kopf, M., Klähn, S., Voss, B., Stüber, K., Huettel, B., Reinhardt, R., and Hess, W.R. (2014). Finished genome sequence of the unicellular cyanobacterium *Synechocystis* sp. strain PCC 6714. Genome Announc. *2*. https://doi.org/10.1128/genomeA.00757-14.

19. Kopf, M., Klähn, S., Scholz, I., Hess, W.R., and Voß, B. (2015). Variations in the non-coding transcriptome as a driver of inter-strain divergence and physiological adaptation in bacteria. Sci. Rep. *5*, 9560. https://doi.org/10.1038/srep09560.

20. Masamoto, K., Misawa, N., Kaneko, T., Kikuno, R., and Toh, H. (1998). Beta-carotene hydroxylase gene from the cyanobacterium *Synechocystis* sp. PCC6803. Plant Cell Physiol. *39*, 560–564. https://doi.org/10.1093/oxfordjournals.pcp.a029405.

21. Islam, M.R., Aikawa, S., Midorikawa, T., Kashino, Y., Satoh, K., and Koike, H. (2008). *slr1923* of *Synechocystis* sp. PCC6803 is essential for conversion of 3,8-divinyl(proto)chlorophyll(ide) to 3-monovinyl(proto)chlorophyll(ide). Plant Physiol. *148*, 1068–1081. https://doi.org/10.1104/pp.108.123117.

22. Ito, H., Yokono, M., Tanaka, R., and Tanaka, A. (2008). Identification of a novel vinyl reductase gene essential for the biosynthesis of monovinyl chlorophyll in *Synechocystis* sp. PCC6803. J. Biol. Chem. *283*, 9002–9011. https://doi.org/10.1074/jbc.M708369200.

23. Meydan, S., Marks, J., Klepacki, D., Sharma, V., Baranov, P.V., Firth, A.E., Margus, T., Kefi, A., Vázquez-Laslop, N., and Mankin, A.S. (2019). Retapamulin-assisted ribosome profiling reveals the alternative bacterial proteome. Mol. Cell *74*, 481-493.e6. https://doi.org/10.1016/j.molcel.2019.02.017.
