## Supplementary Dataset 3 for "A revised genome annotation of the model cyanobacterium *Synechocystis* based on start and stop codon-enriched ribosome profiling and proteogenomics"

**STLLNMVSGFNQPTQGTVK | NC_000911.1 : 1005181 : −. pep01**

Charge: +2 | Collision Energy: 27.0 | Percursor m/z: 1011.521484 | Mass Diff. 0.006104 Da

Percolator Score: 1.00 | Hyperscore: 57.92 | Spectral Angle: 0.93 | Pearson's Correlation: 0.99

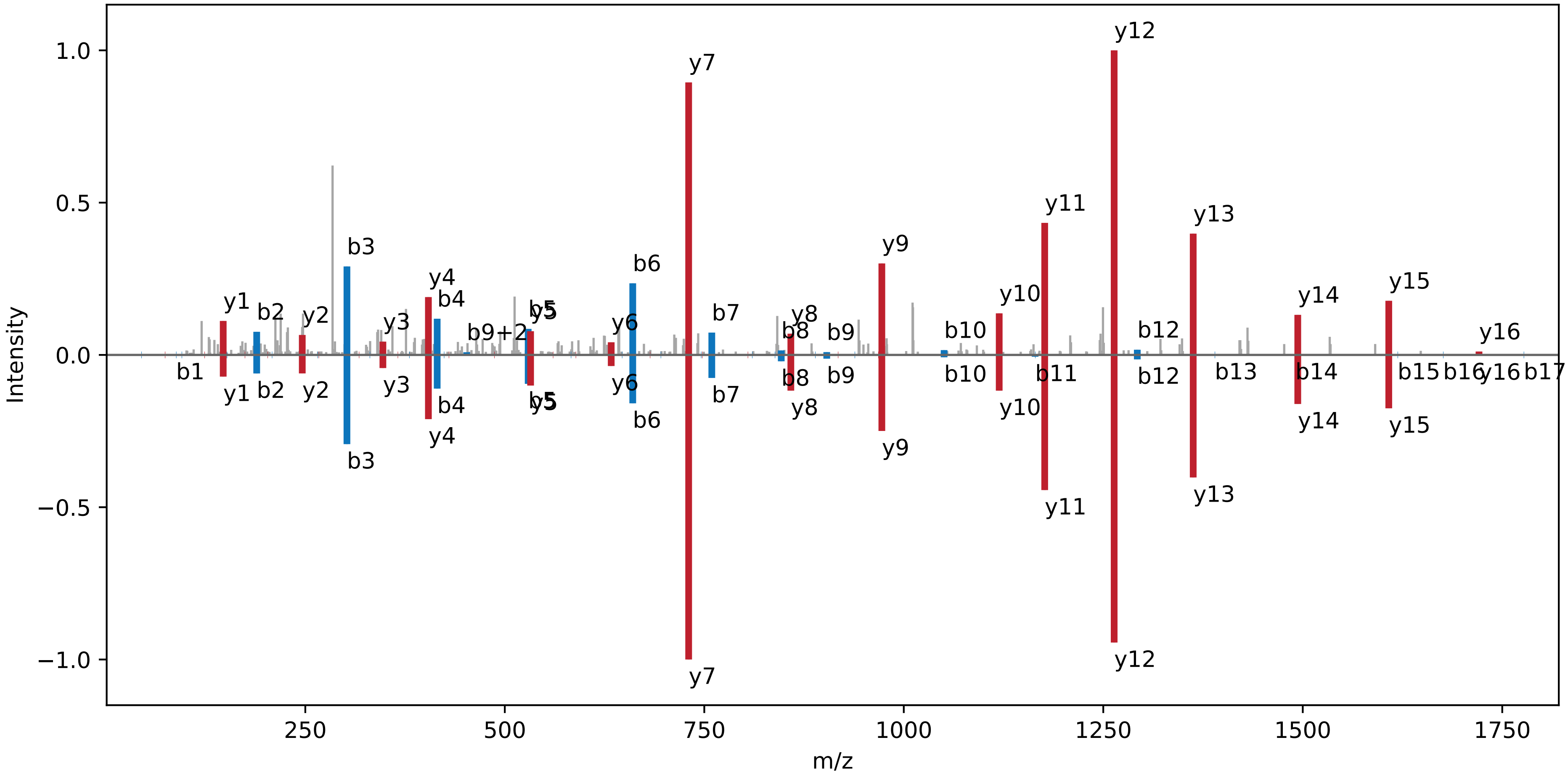

**M[Ox]QTM[Ox]NVNDPILTEEKTAFLAIENVSK | NC_000911.1 : 1005181 : −. pep02**

Charge: +3 | Collision Energy: 30.0 | Percursor m/z: 990.493652 | Mass Diff. 1.007568 Da

Percolator Score: 1.00 | Hyperscore: 18.55 | Spectral Angle: 0.53 | Pearson's Correlation: 0.73

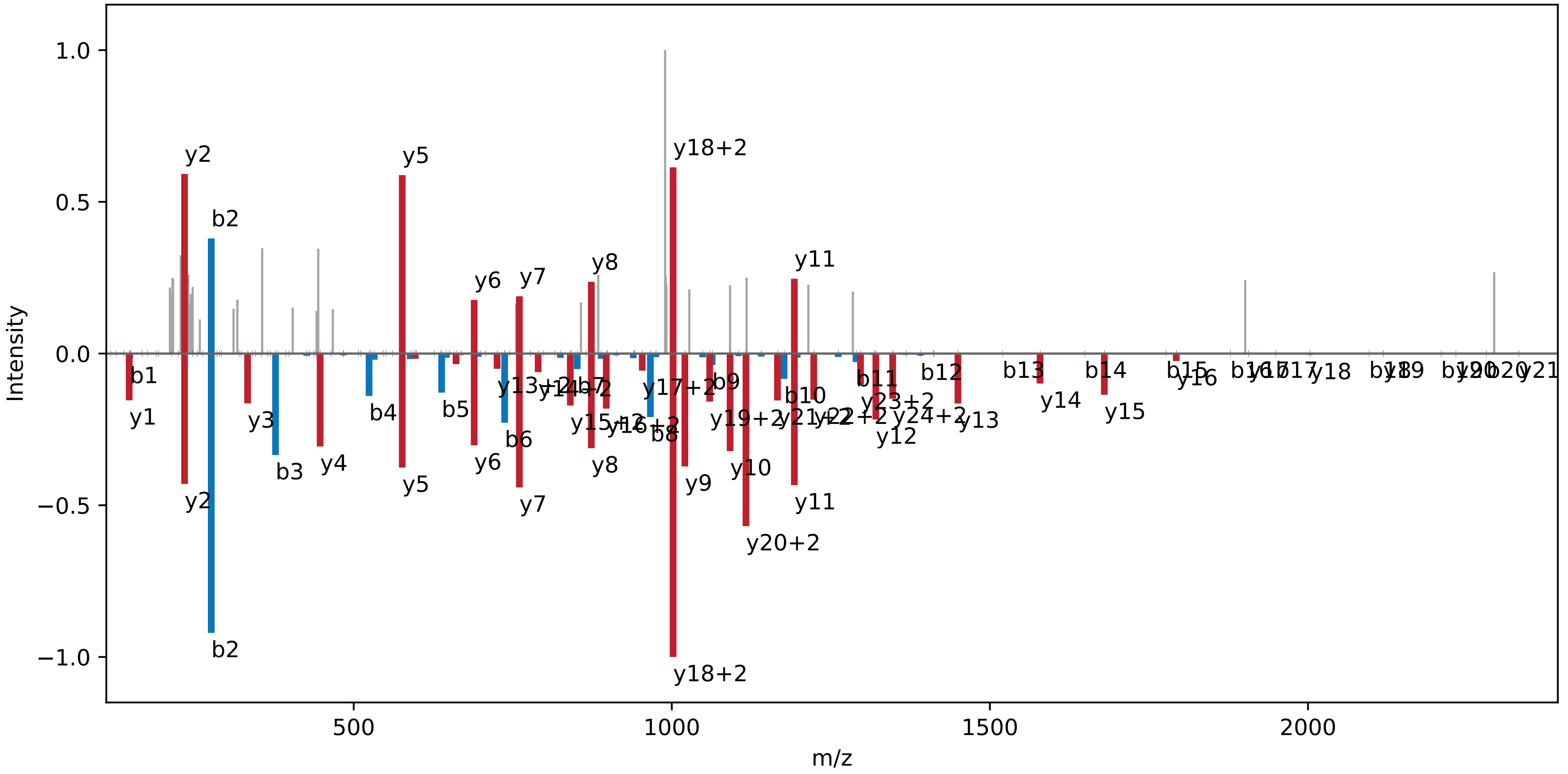

**YAHDDDDDLK | NC_000911.1 : 1005181 : −. pep03**

Charge: +3 | Collision Energy: 27.0 | Percursor m/z: 402.834503 | Mass Diff. -0.000122 Da

Percolator Score: 1.00 | Hyperscore: 27.87 | Spectral Angle: 0.91 | Pearson's Correlation: 0.99

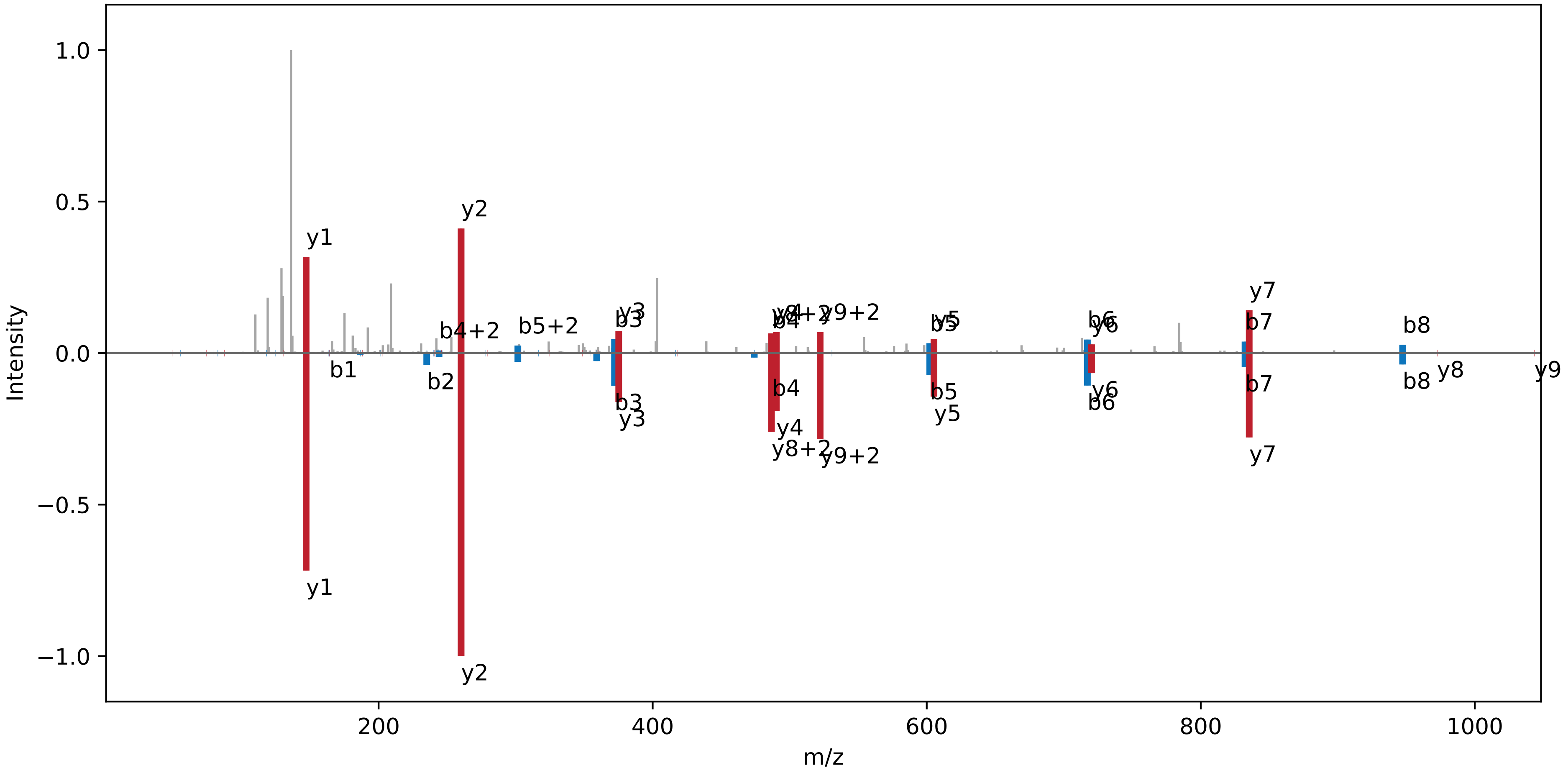

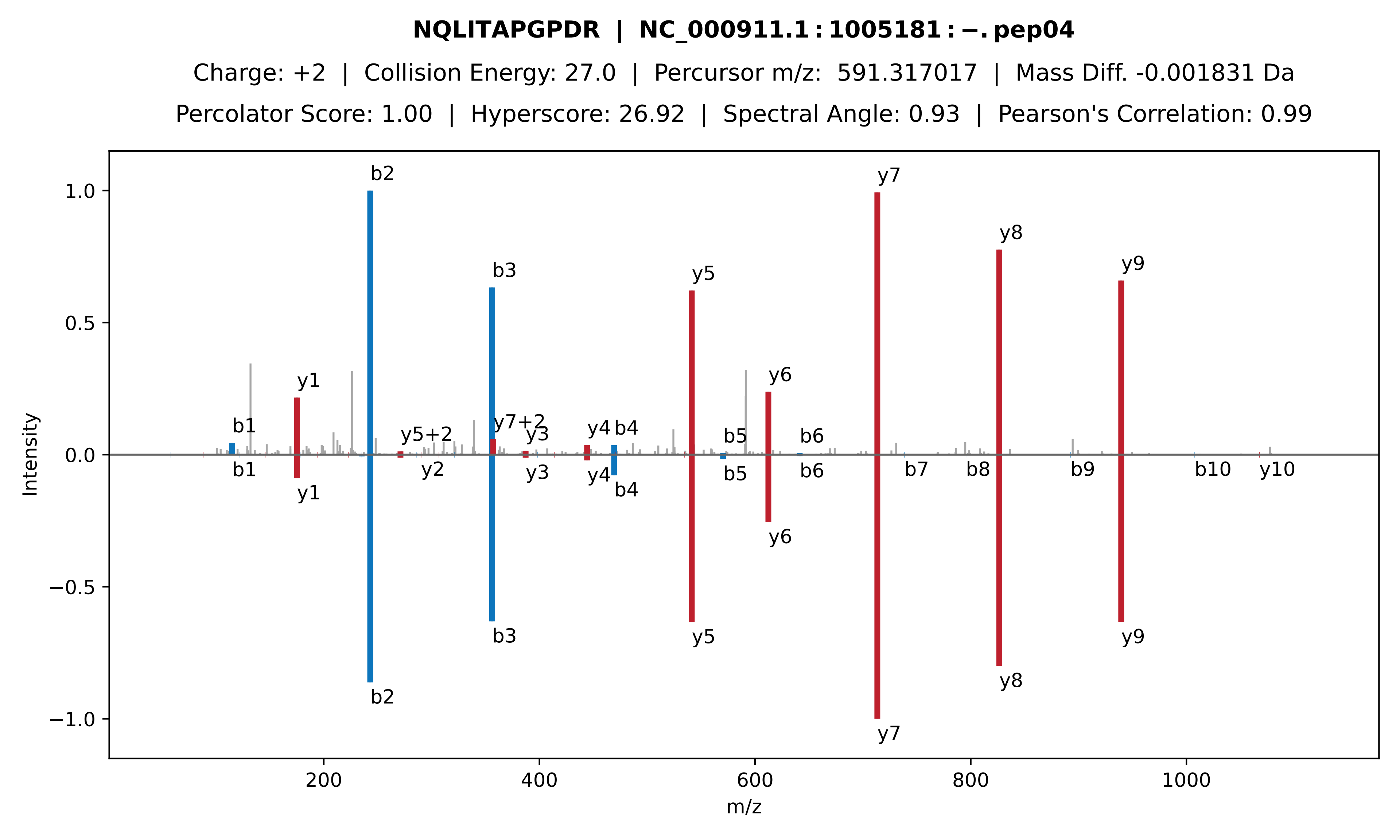
NQLITAPGPDR | NC_000911.1 : 1005181 : −. pep04
Charge: +2 | Collision Energy: 27.0 | Percursor m/z: 591.317017 | Mass Diff. -0.001831 Da
Percolator Score: 1.00 | Hyperscore: 26.92 | Spectral Angle: 0.93 | Pearson's Correlation: 0.99
Intensity
m/z
b1
y1
b2
y5+2
y7+2
b3
y3
y4
b4
y5
b5
y6
b6
y7
y8
y9
y2
b7
b8
b9
b10
y10

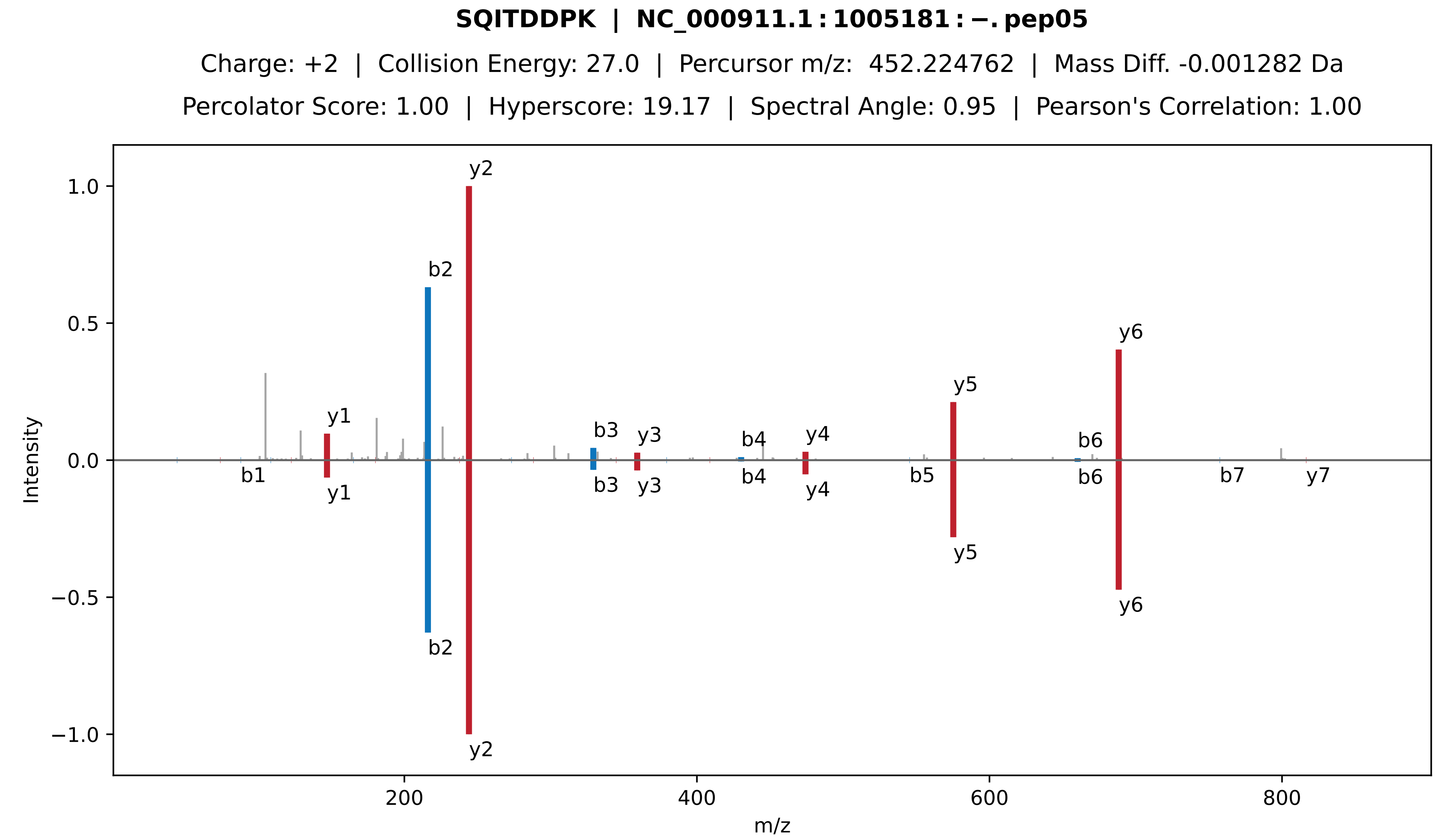
SQITDDPK | NC_000911.1 : 1005181 : −. pep05
Charge: +2 | Collision Energy: 27.0 | Percursor m/z: 452.224762 | Mass Diff. -0.001282 Da
Percolator Score: 1.00 | Hyperscore: 19.17 | Spectral Angle: 0.95 | Pearson's Correlation: 1.00
Intensity
m/z
y1
b1
b2
y2
b3
y3
b4
y4
b5
y5
b6
y6
b7
y7
1.0
0.5
0.0
−0.5
−1.0
200
400
600
800

**VYPTAQGPYTVLDGVNLEVK | NC_000911.1 : 1005181 : −. pep06**

Charge: +2 | Collision Energy: 27.0 | Percursor m/z: 1082.073400 | Mass Diff. 0.000488 Da

Percolator Score: 1.00 | Hyperscore: 59.24 | Spectral Angle: 0.91 | Pearson's Correlation: 0.99

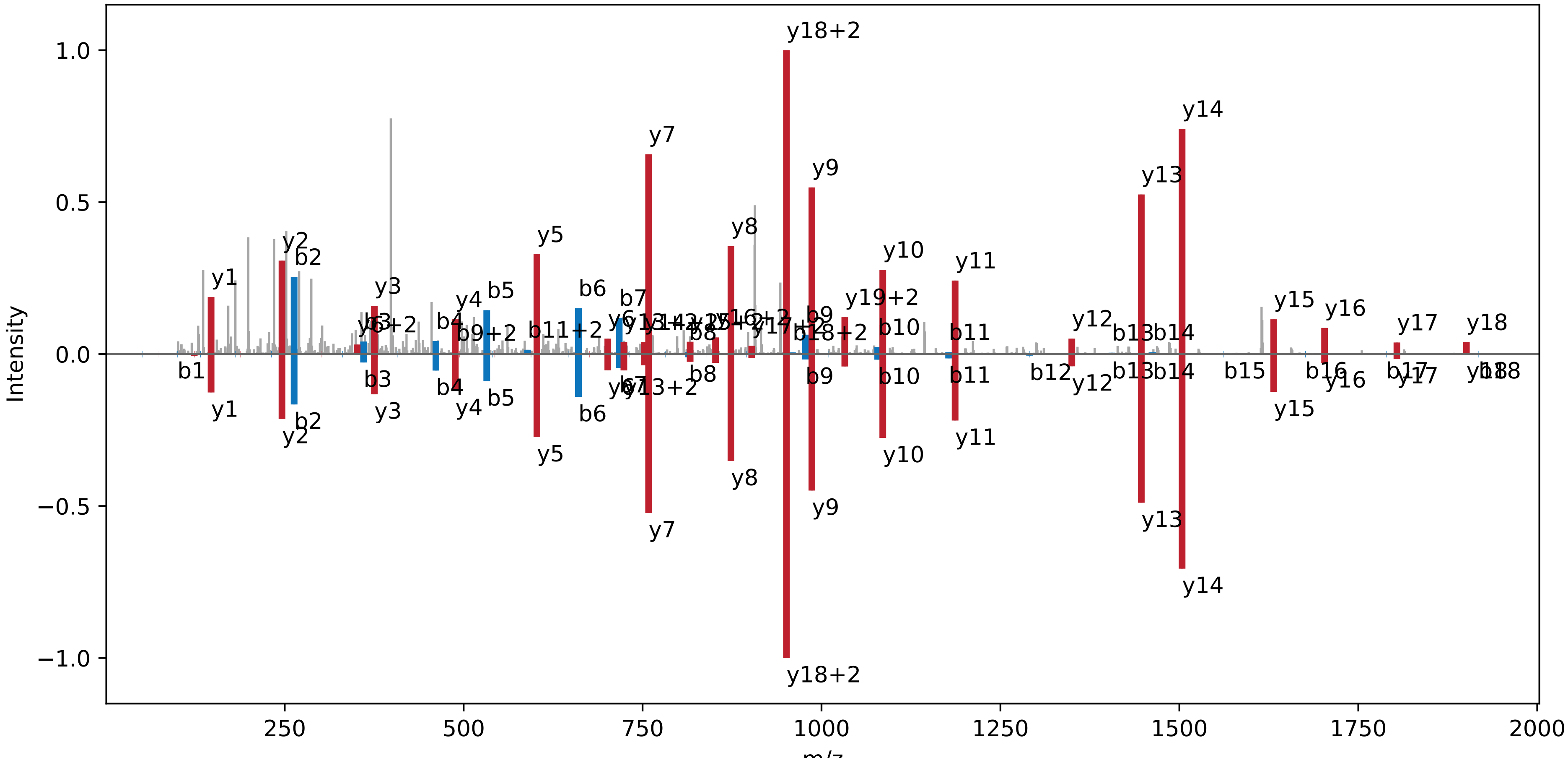

**VM[Ox]AGALGLVAIALVGVGLVQTFGSK | NC_000911.1 : 1005181 : −. pep07**

Charge: +3 | Collision Energy: 27.0 | Percursor m/z: 796.464233 | Mass Diff. 0.002686 Da

Percolator Score: 1.00 | Hyperscore: 34.87 | Spectral Angle: 0.66 | Pearson's Correlation: 0.85

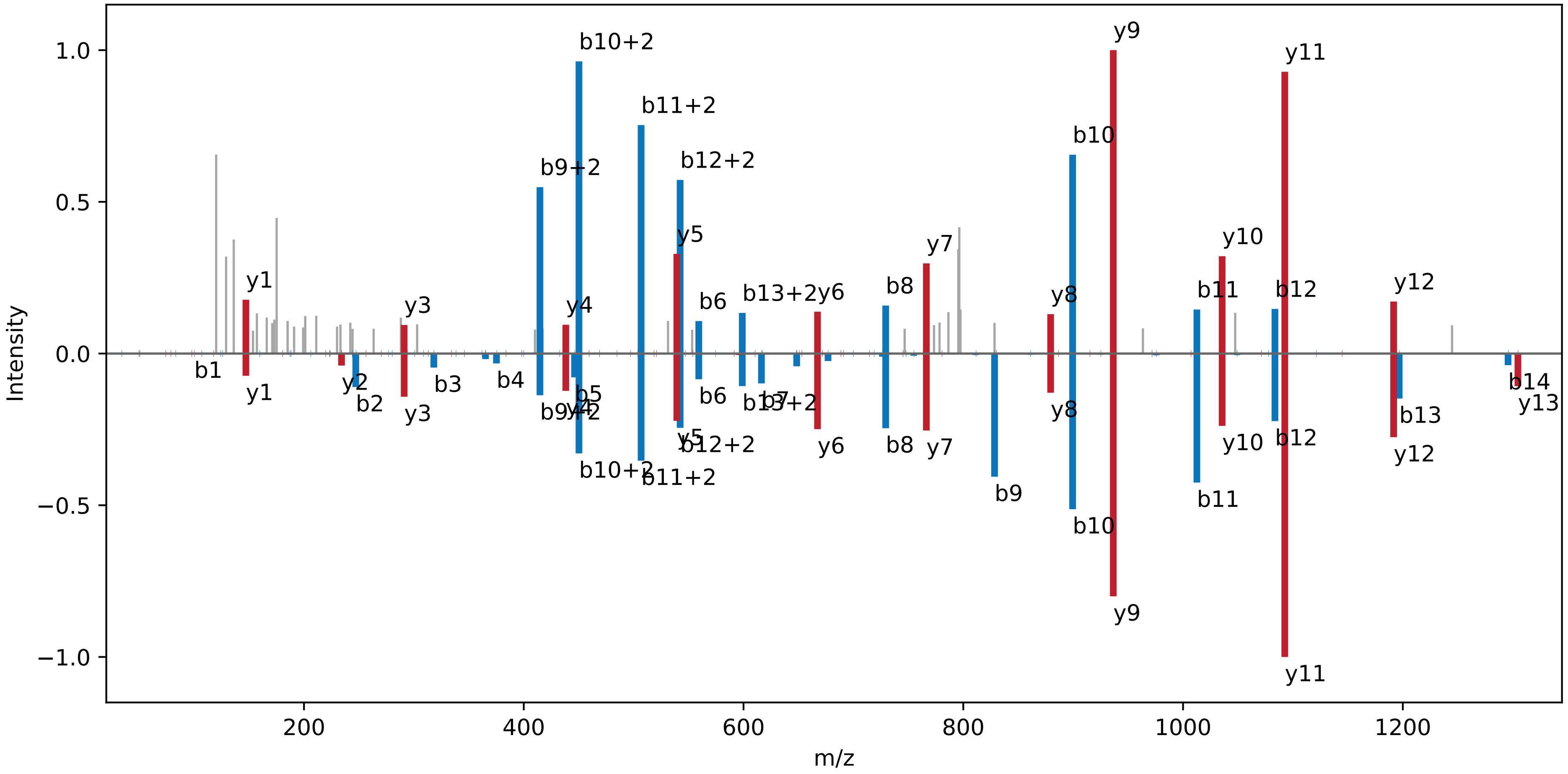

**M[Ox]QTM[Ox]NVNDPILTEEK | NC_000911.1 : 1005181 : −. pep08**

Charge: +2 | Collision Energy: 27.0 | Percursor m/z: 897.915649 | Mass Diff. -0.001465 Da

Percolator Score: 1.00 | Hyperscore: 41.34 | Spectral Angle: 0.94 | Pearson's Correlation: 0.99

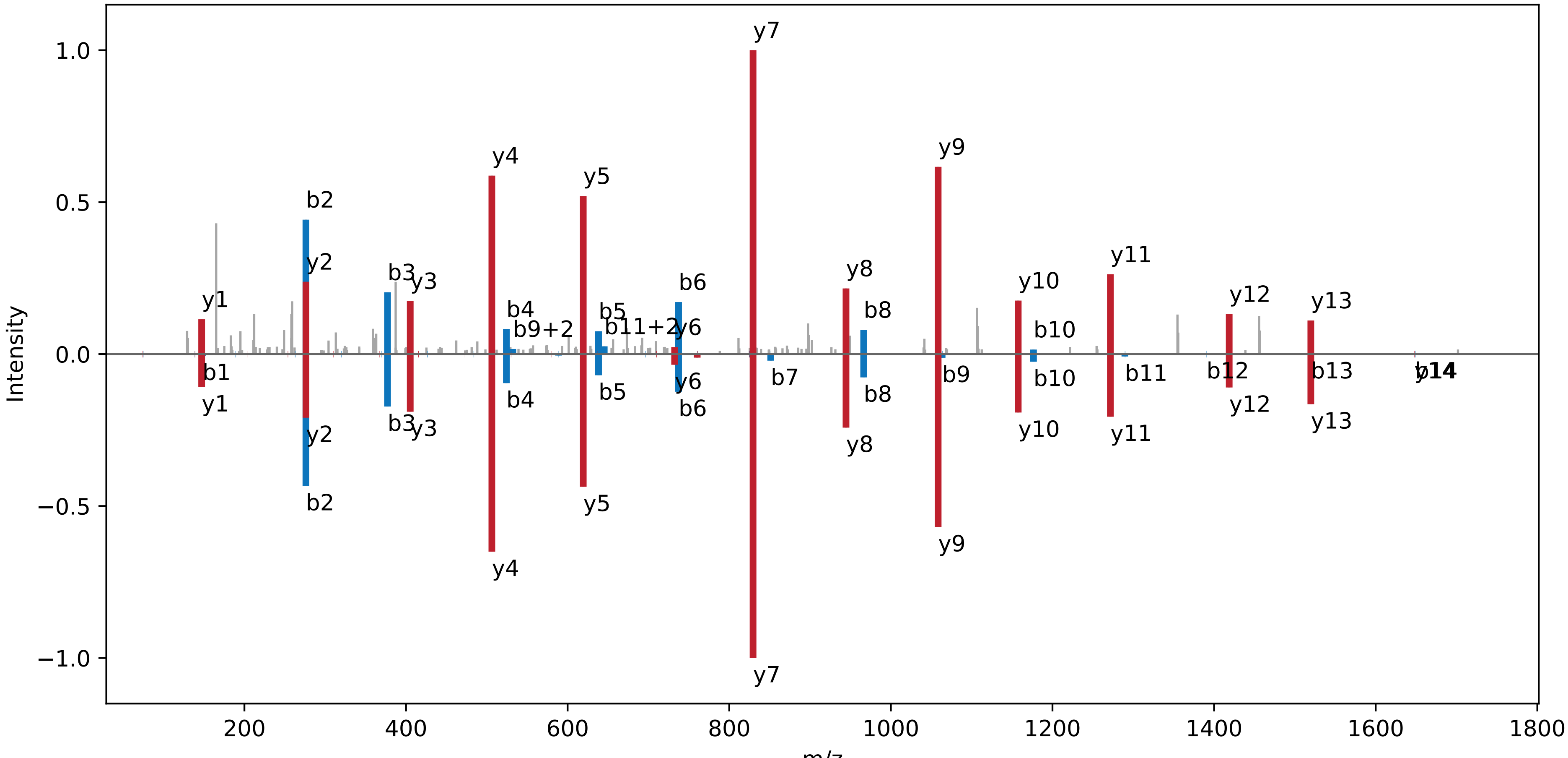

**IGEILDIPFDRPR | NC_000911.1 : 1005181 : −. pep09**

Charge: +3 | Collision Energy: 27.0 | Percursor m/z: 514.287781 | Mass Diff. -0.001709 Da

Percolator Score: 1.00 | Hyperscore: 38.00 | Spectral Angle: 0.45 | Pearson's Correlation: 0.60

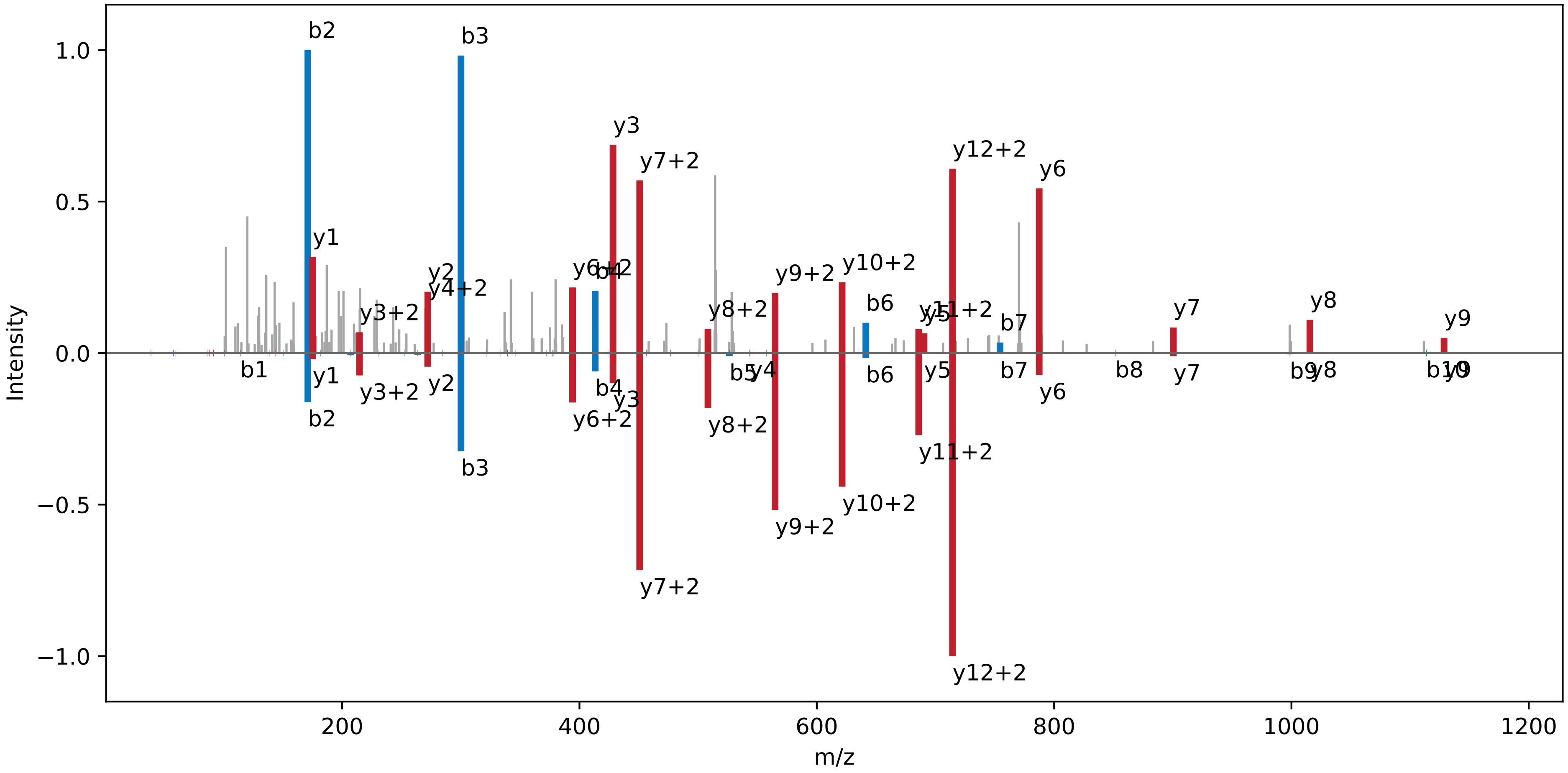

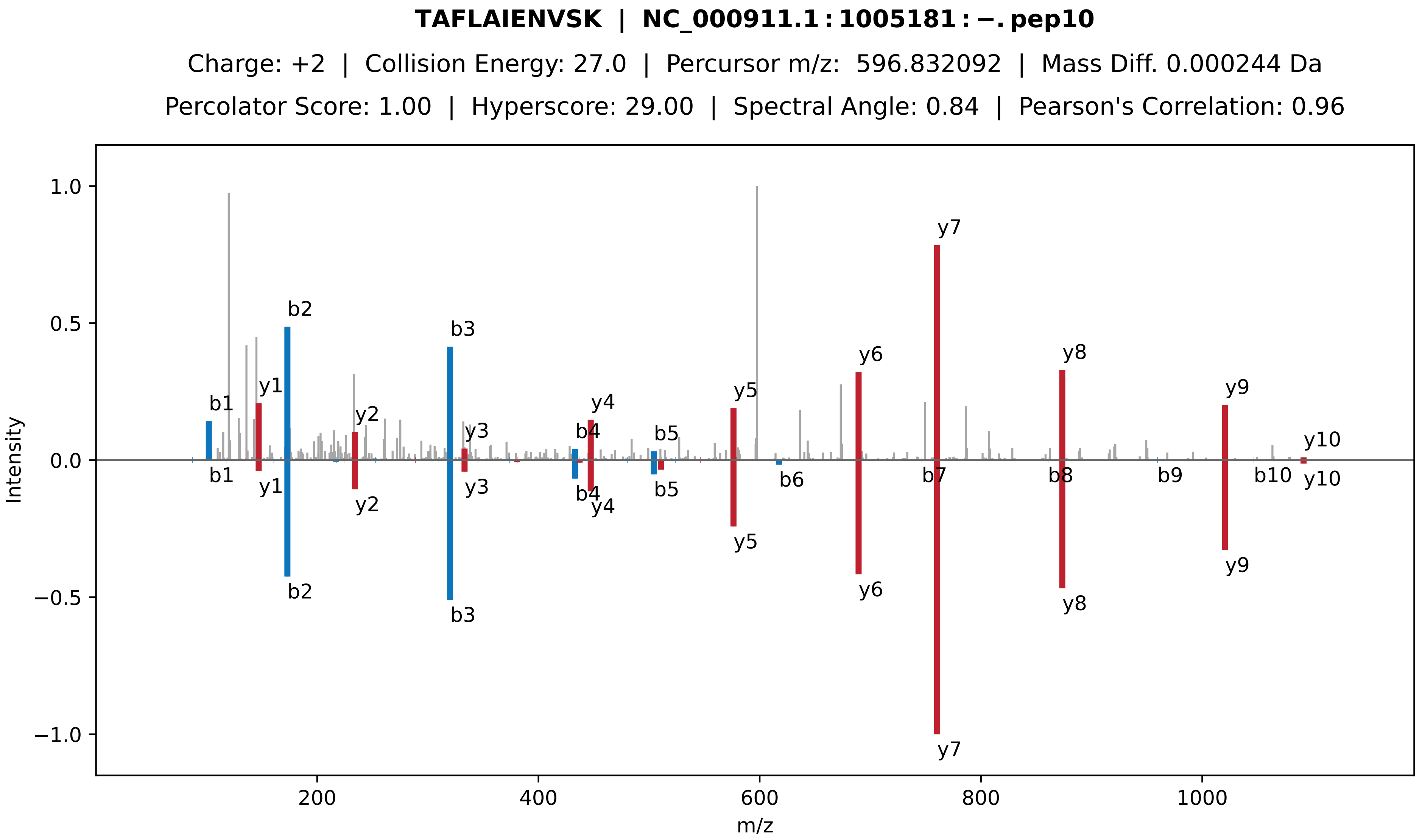

**TAFLAIENVSK | NC_000911.1 : 1005181 : −. pep10**
Charge: +2 | Collision Energy: 27.0 | Percursor m/z: 596.832092 | Mass Diff. 0.000244 Da
Percolator Score: 1.00 | Hyperscore: 29.00 | Spectral Angle: 0.84 | Pearson's Correlation: 0.96
Intensity
m/z
b1
b2
b3
b4
b5
b6
b7
b8
b9
b10
y1
y2
y3
y4
y5
y6
y7
y8
y9
y10

**TEKEELVAHYLEMVGLTEAAQK | NC_000911.1 : 1005181 : −. pep11**

Charge: +3 | Collision Energy: 27.0 | Percursor m/z: 830.761963 | Mass Diff. 1.007324 Da

Percolator Score: 1.00 | Hyperscore: 41.69 | Spectral Angle: 0.78 | Pearson's Correlation: 0.93

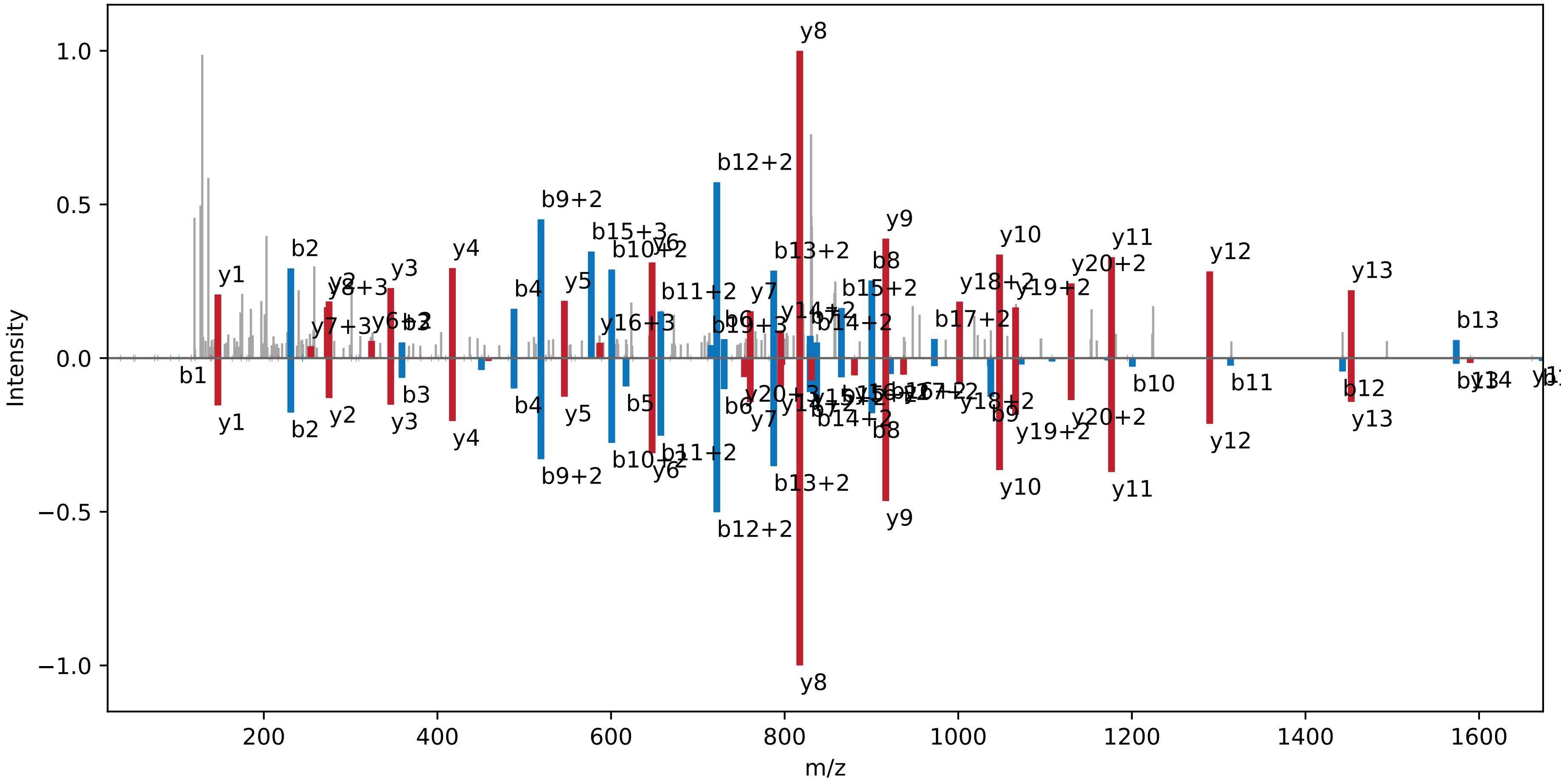

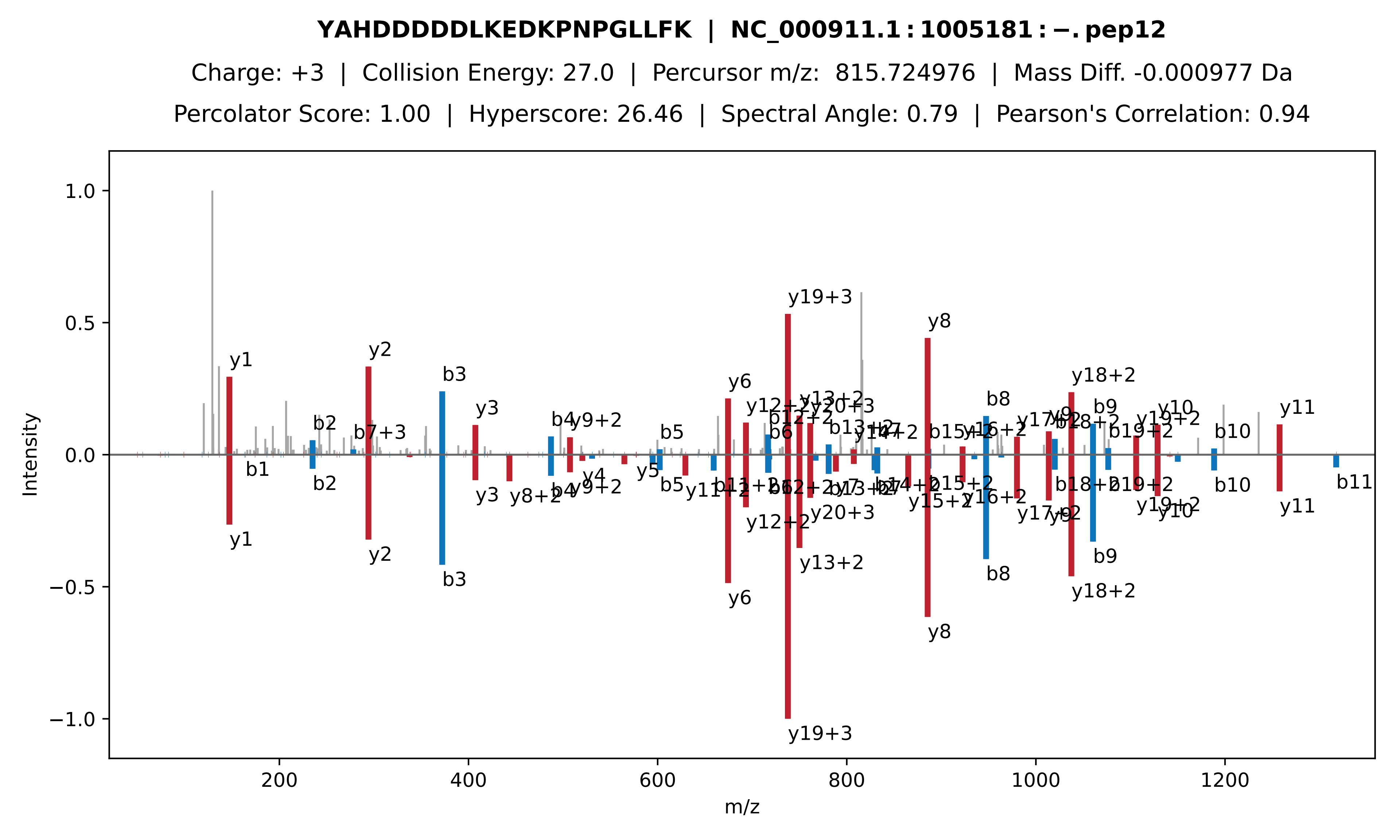
YAHDDDDDLKEDKPNPGLLFK | NC_000911.1 : 1005181 : −. pep12
Charge: +3 | Collision Energy: 27.0 | Percursor m/z: 815.724976 | Mass Diff. -0.000977 Da
Percolator Score: 1.00 | Hyperscore: 26.46 | Spectral Angle: 0.79 | Pearson's Correlation: 0.94
Intensity
m/z
y1
y2
b3
y3
y6
y19+3
y8
y18+2
b8
b9
y11
b10
b11
b1
b2
b4
b5
b6
y4
y5
y9+2
y12+2
y13+2
y20+3
y10
y19+2
b7+3
y8+2

**ENNLTVLMITHDIDEALFLADR | NC_000911.1 : 1005181 : −. pep13**

Charge: +3 | Collision Energy: 27.0 | Percursor m/z: 848.767883 | Mass Diff. 1.001709 Da

Percolator Score: 1.00 | Hyperscore: 45.10 | Spectral Angle: 0.37 | Pearson's Correlation: 0.50

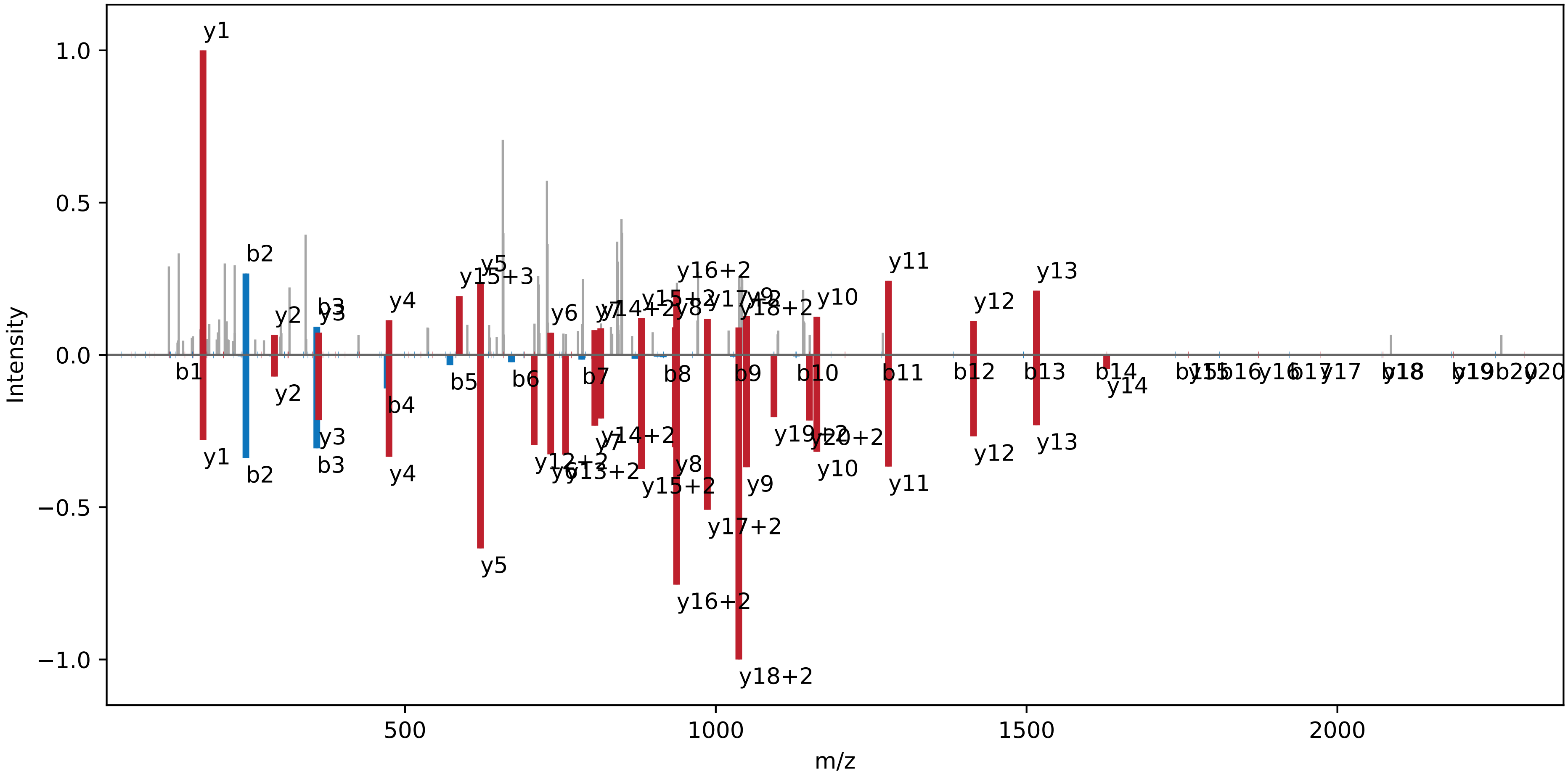

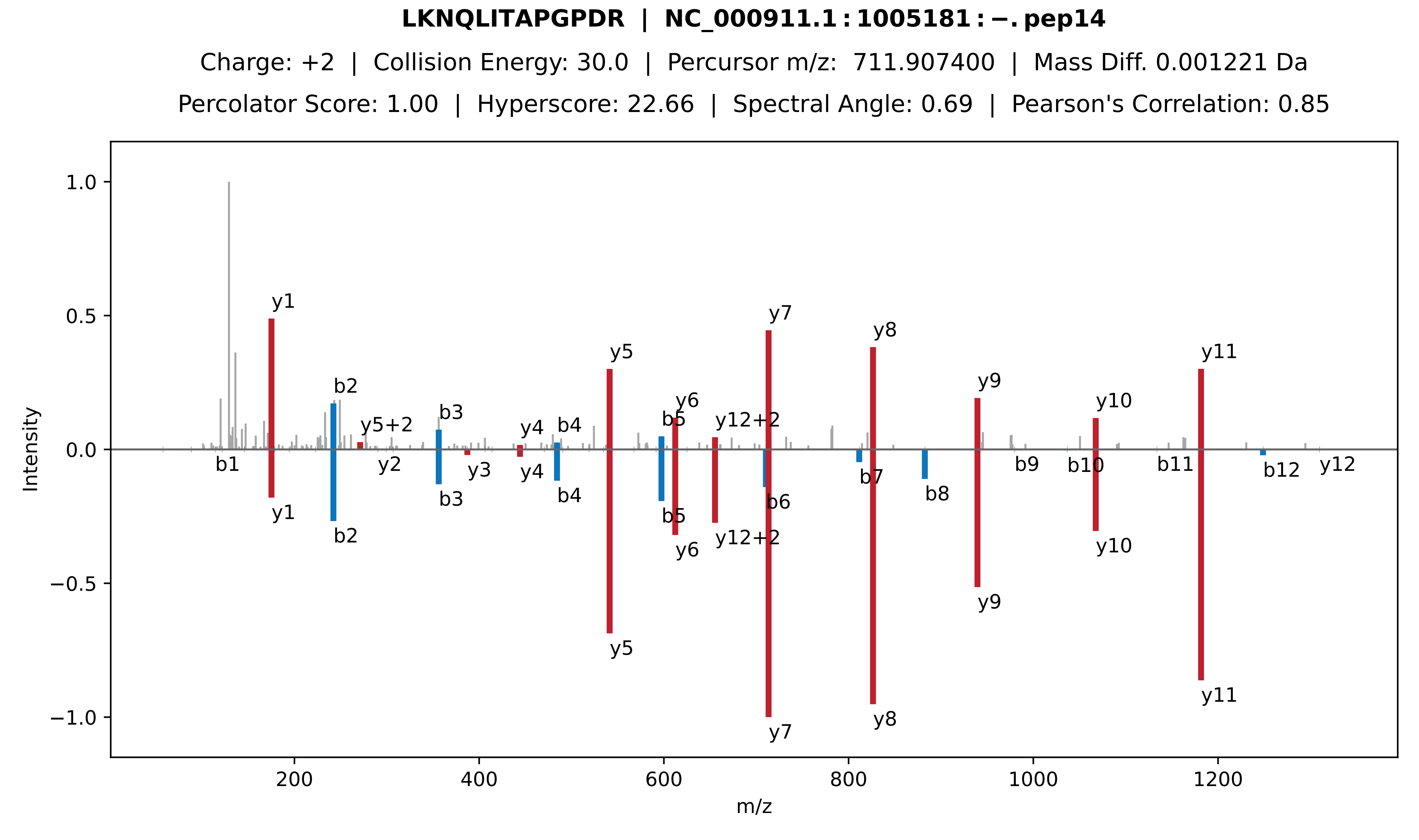
LKNQLITAPGPDR | NC_000911.1 : 1005181 : −. pep14
Charge: +2 | Collision Energy: 30.0 | Percursor m/z: 711.907400 | Mass Diff. 0.001221 Da
Percolator Score: 1.00 | Hyperscore: 22.66 | Spectral Angle: 0.69 | Pearson's Correlation: 0.85
Intensity
m/z
1.0
0.5
0.0
−0.5
−1.0
200
400
600
800
1000
1200
y1
b2
y5+2
b3
y4
b4
y5
b5
y6
y12+2
y7
y8
y9
y10
y11
b1
y2
y3
b6
b7
b8
b9
b10
b11
b12
y12

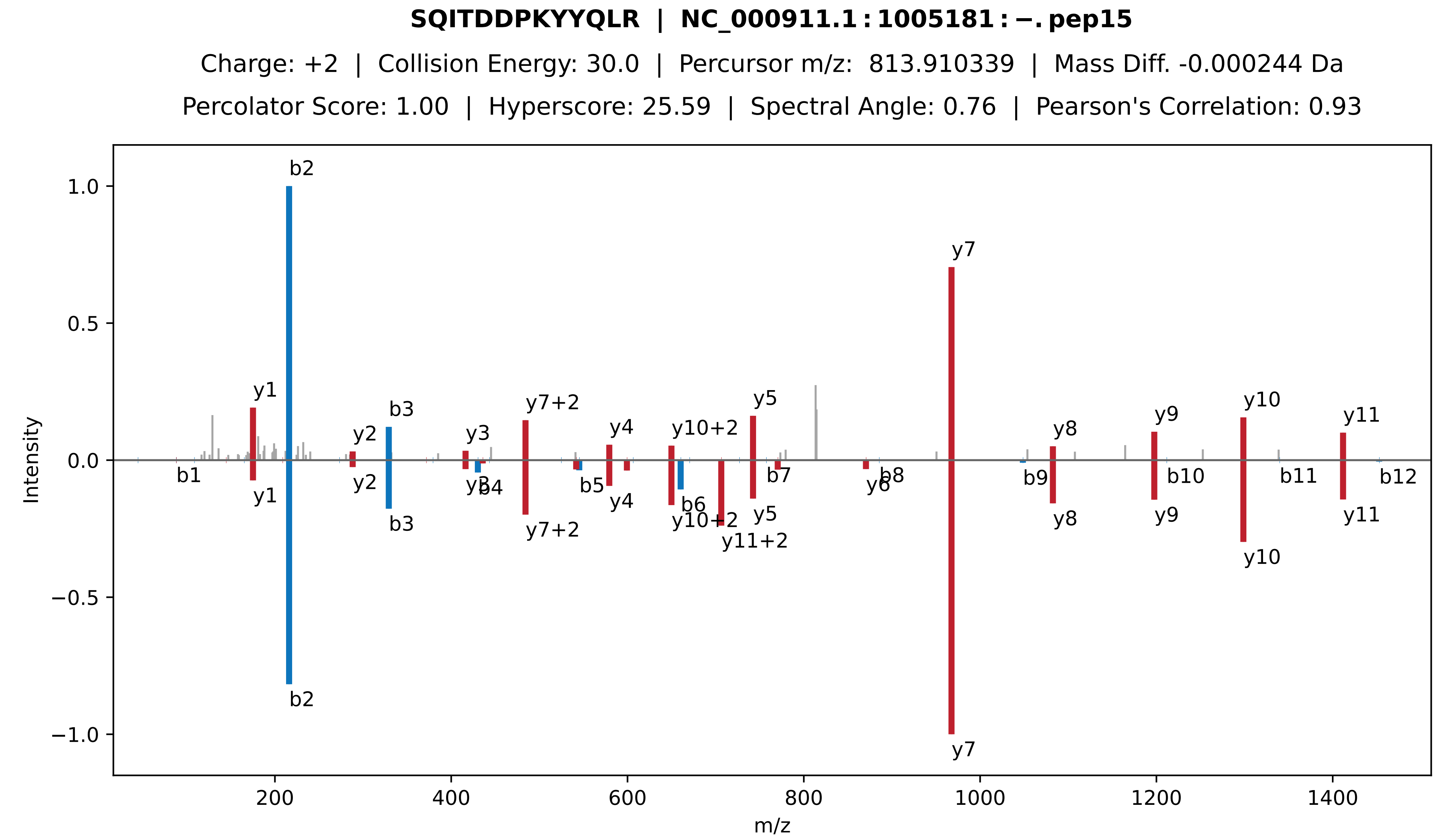

SQITDDPKYYQLR | NC_000911.1 : 1005181 : −. pep15
Charge: +2 | Collision Energy: 30.0 | Percursor m/z: 813.910339 | Mass Diff. -0.000244 Da
Percolator Score: 1.00 | Hyperscore: 25.59 | Spectral Angle: 0.76 | Pearson's Correlation: 0.93
Intensity
m/z
b1
b2
b3
b4
b5
b6
b7
b8
b9
b10
b11
b12
y1
y2
y3
y4
y5
y6
y7
y8
y9
y10
y11
y7+2
y10+2
y11+2

**SAYDNVSLAVESVYPDK | NC_000911.1 : 1005181 : −. pep16**

Charge: +2 | Collision Energy: 27.0 | Percursor m/z: 928.952087 | Mass Diff. -0.000854 Da

Percolator Score: 1.00 | Hyperscore: 49.10 | Spectral Angle: 0.93 | Pearson's Correlation: 0.99

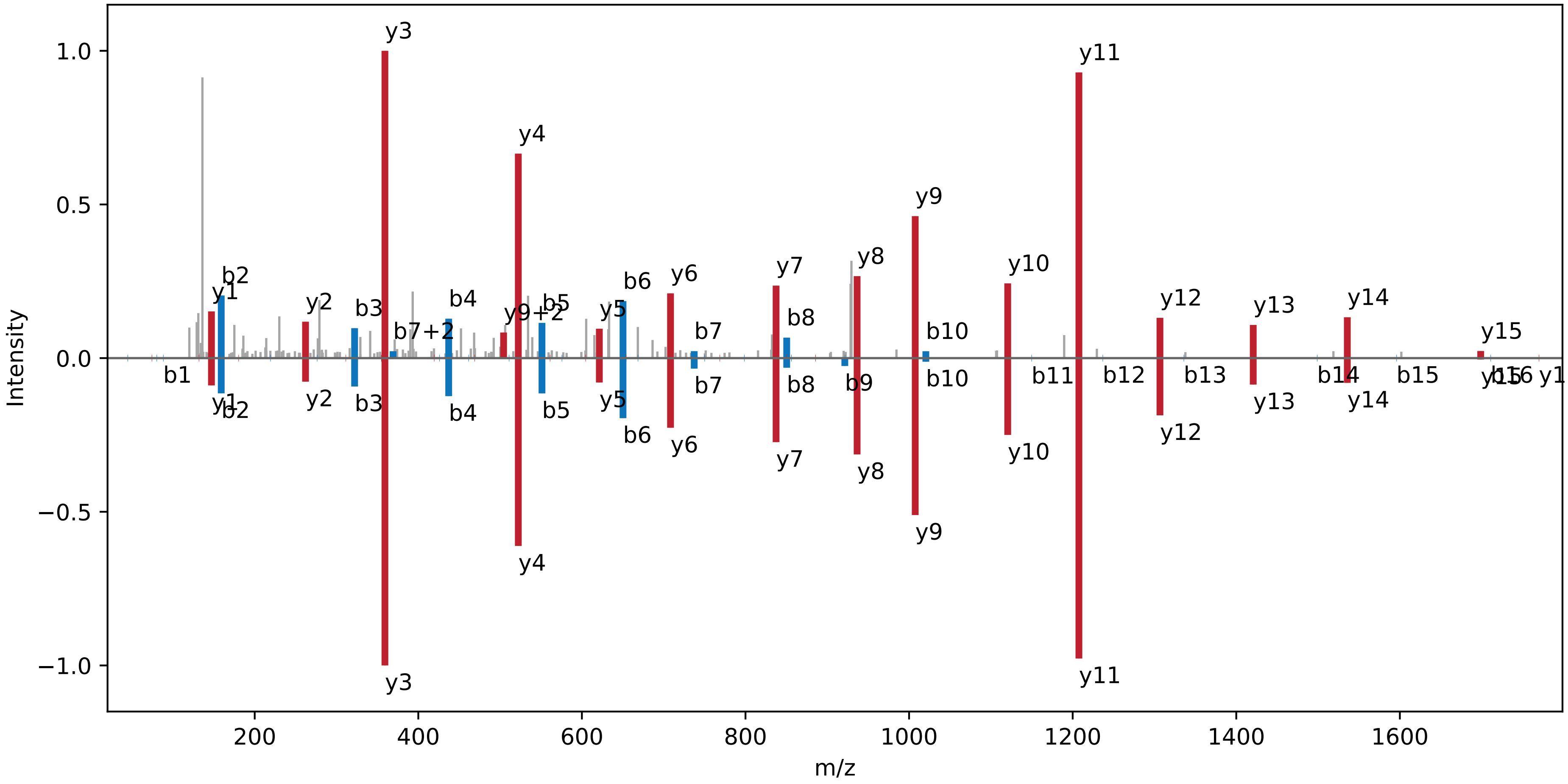

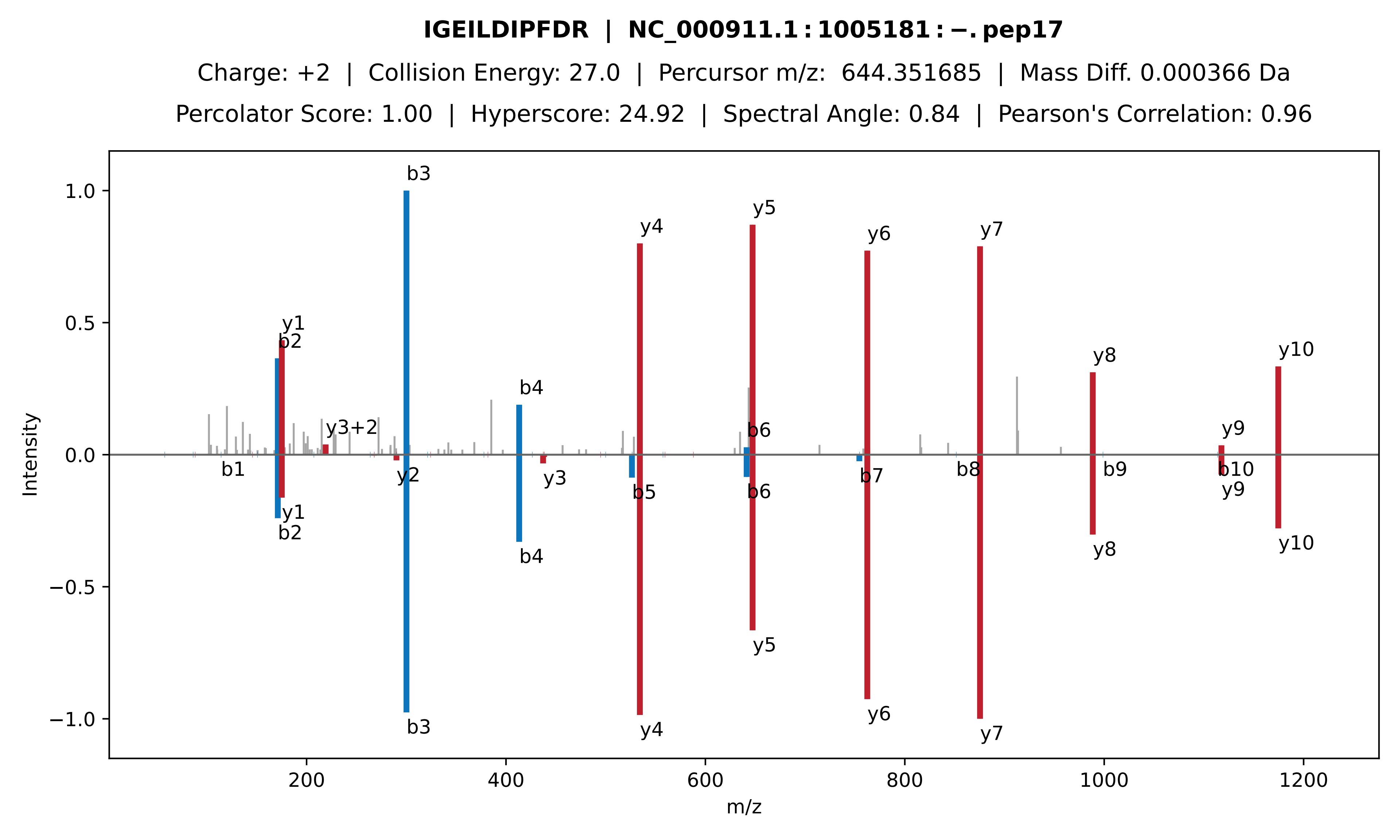
IGEILDIPFDR | NC_000911.1:1005181:-.pep17
Charge: +2 | Collision Energy: 27.0 | Percursor m/z: 644.351685 | Mass Diff. 0.000366 Da
Percolator Score: 1.00 | Hyperscore: 24.92 | Spectral Angle: 0.84 | Pearson's Correlation: 0.96
Intensity
m/z
b1
b2
b3
b4
b5
b6
b7
b8
b9
b10
y1
y2
y3
y3+2
y4
y5
y6
y7
y8
y9
y10

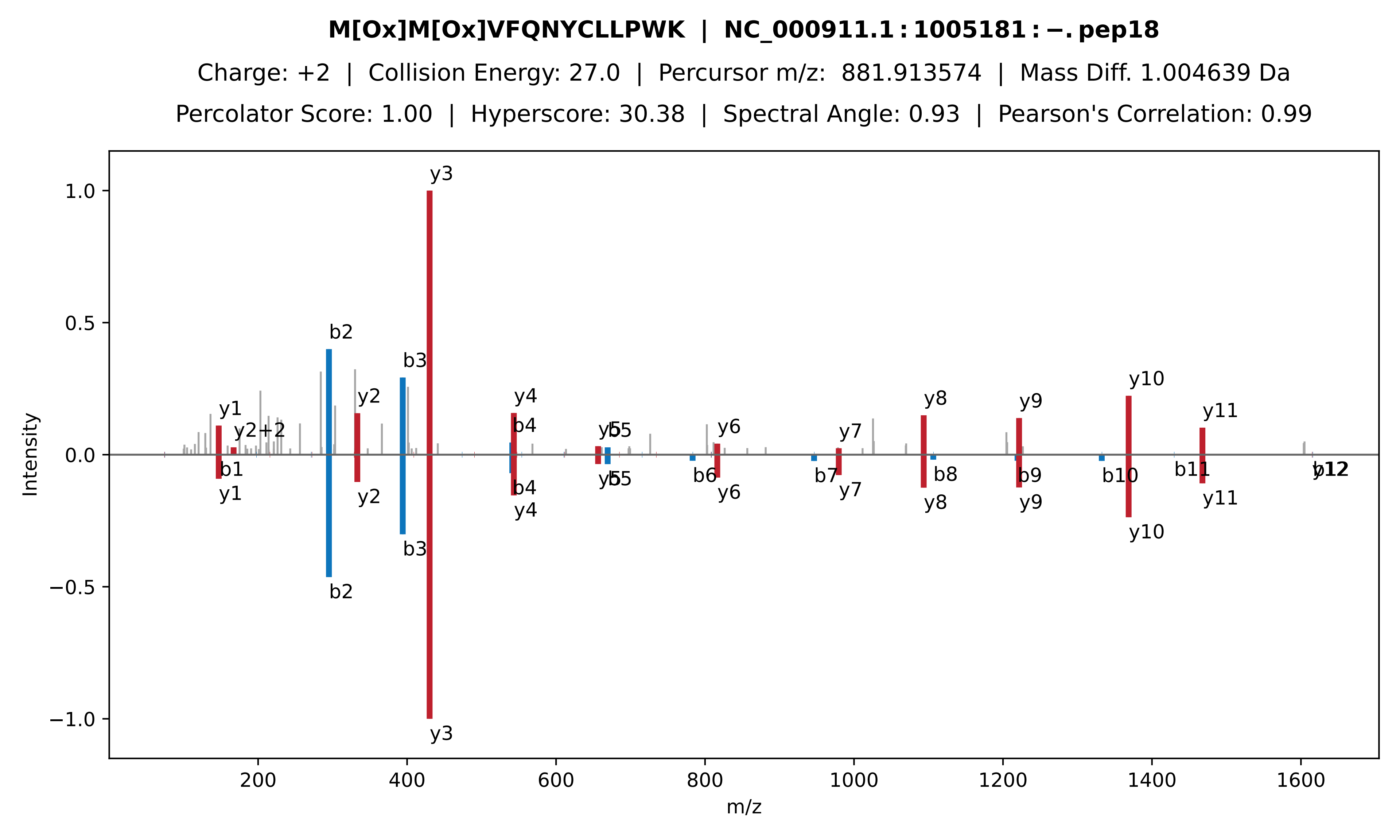
M[Ox]M[Ox]VFQNYCLLPWK | NC_000911.1 : 1005181 : −. pep18
Charge: +2 | Collision Energy: 27.0 | Percursor m/z: 881.913574 | Mass Diff. 1.004639 Da
Percolator Score: 1.00 | Hyperscore: 30.38 | Spectral Angle: 0.93 | Pearson's Correlation: 0.99
Intensity
m/z
1.0
0.5
0.0
−0.5
−1.0
200
400
600
800
1000
1200
1400
1600
y1
y2+2
b1
b2
b3
y2
y3
y4
b4
y5
b5
b6
y6
b7
y7
y8
b8
y9
b9
y10
b10
y11
b11
b12
y12

**EDKPNPGLLFK | NC_000911.1 : 1005181 : −. pep19**

Charge: +3 | Collision Energy: 27.0 | Percursor m/z: 419.899933 | Mass Diff. -0.001465 Da

Percolator Score: 1.00 | Hyperscore: 28.48 | Spectral Angle: 0.87 | Pearson's Correlation: 0.97

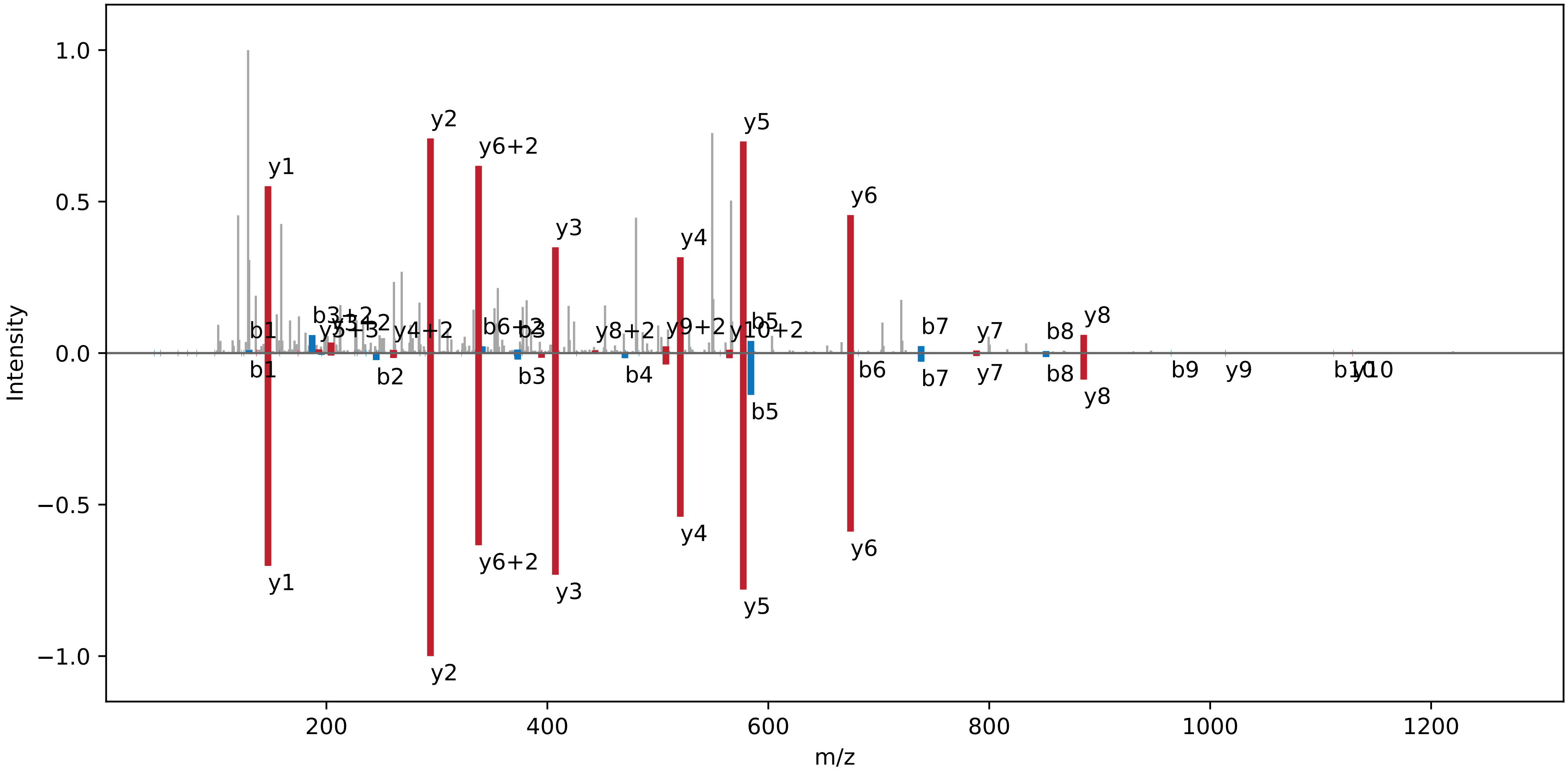

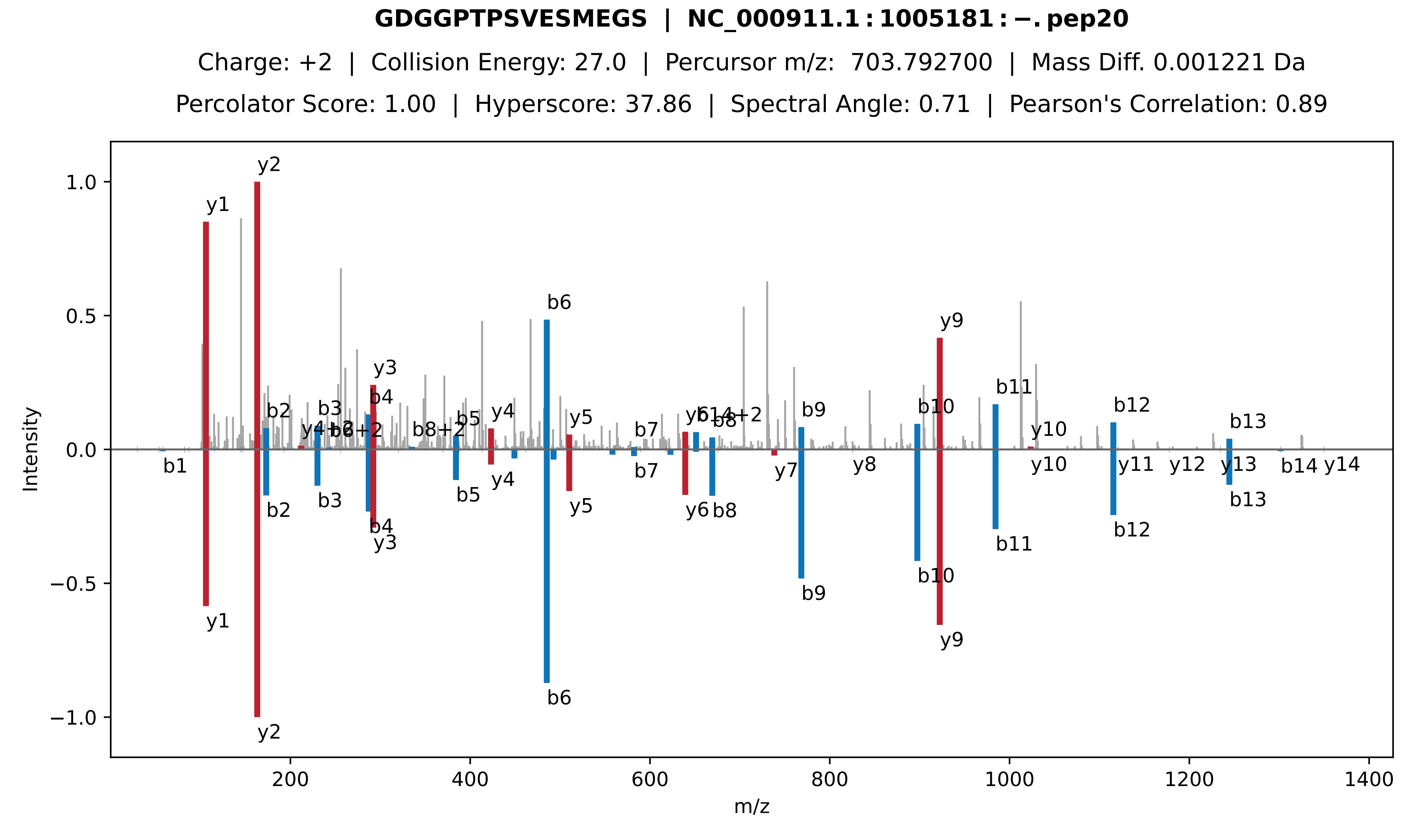
GDGGPTPSVESMEGS | NC_000911.1 : 1005181 : −. pep20
Charge: +2 | Collision Energy: 27.0 | Percursor m/z: 703.792700 | Mass Diff. 0.001221 Da
Percolator Score: 1.00 | Hyperscore: 37.86 | Spectral Angle: 0.71 | Pearson's Correlation: 0.89
Intensity
m/z
y1
y2
y3
y4
y5
y6
y7
y8
y9
y10
y11
y12
y13
y14
b1
b2
b3
b4
b5
b6
b7
b8
b9
b10
b11
b12
b13
b14
b8+2
b14+2

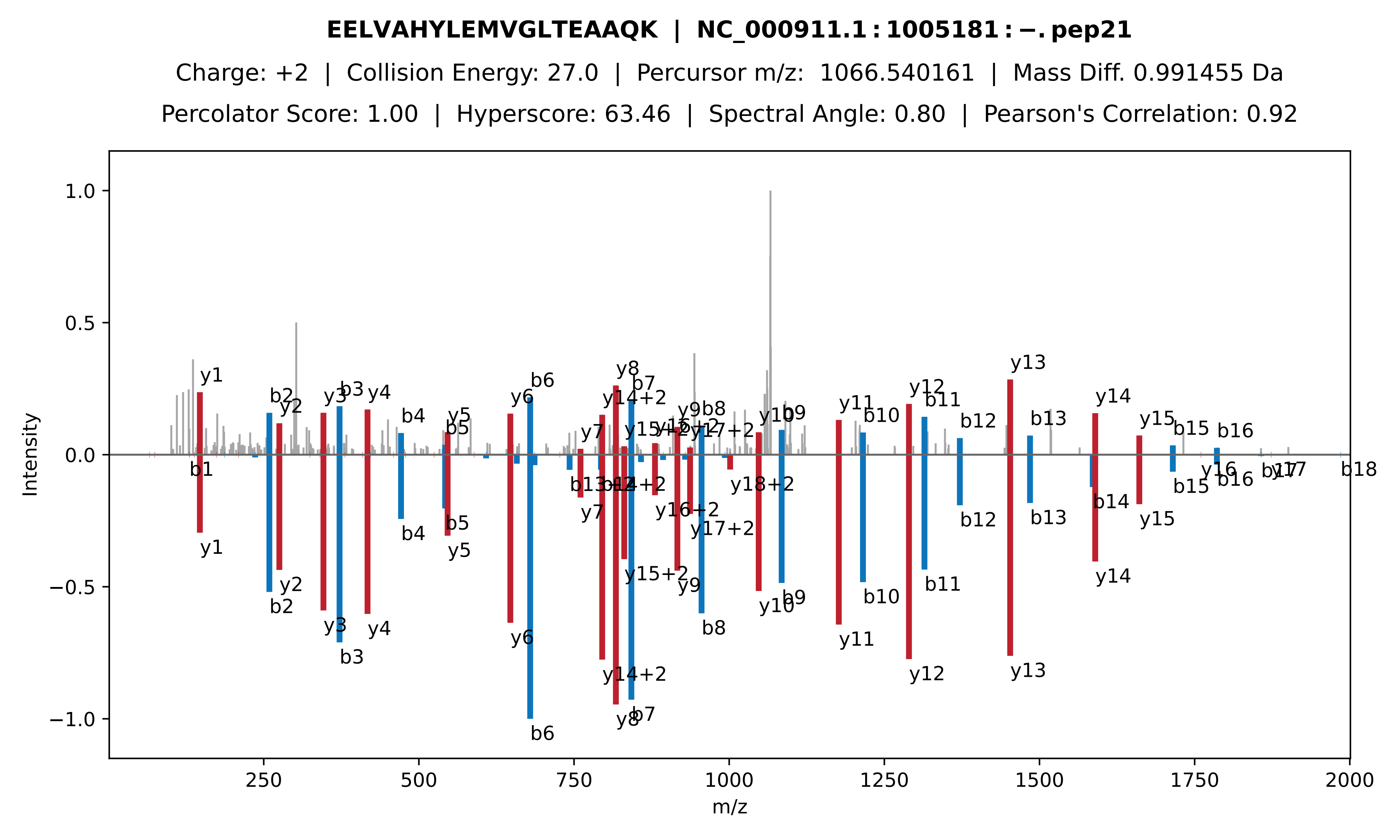
EELVAHYLEMVGLTEAAQK | NC_000911.1:1005181:−.pep21
Charge: +2 | Collision Energy: 27.0 | Percursor m/z: 1066.540161 | Mass Diff. 0.991455 Da
Percolator Score: 1.00 | Hyperscore: 63.46 | Spectral Angle: 0.80 | Pearson's Correlation: 0.92
Intensity
m/z

**LVMMTNGPSAK | NC_000911.1 : 1005181 : −. pep22**

Charge: +2 | Collision Energy: 27.0 | Percursor m/z: 574.793701 | Mass Diff. -0.001465 Da

Percolator Score: 1.00 | Hyperscore: 28.29 | Spectral Angle: 0.90 | Pearson's Correlation: 0.99

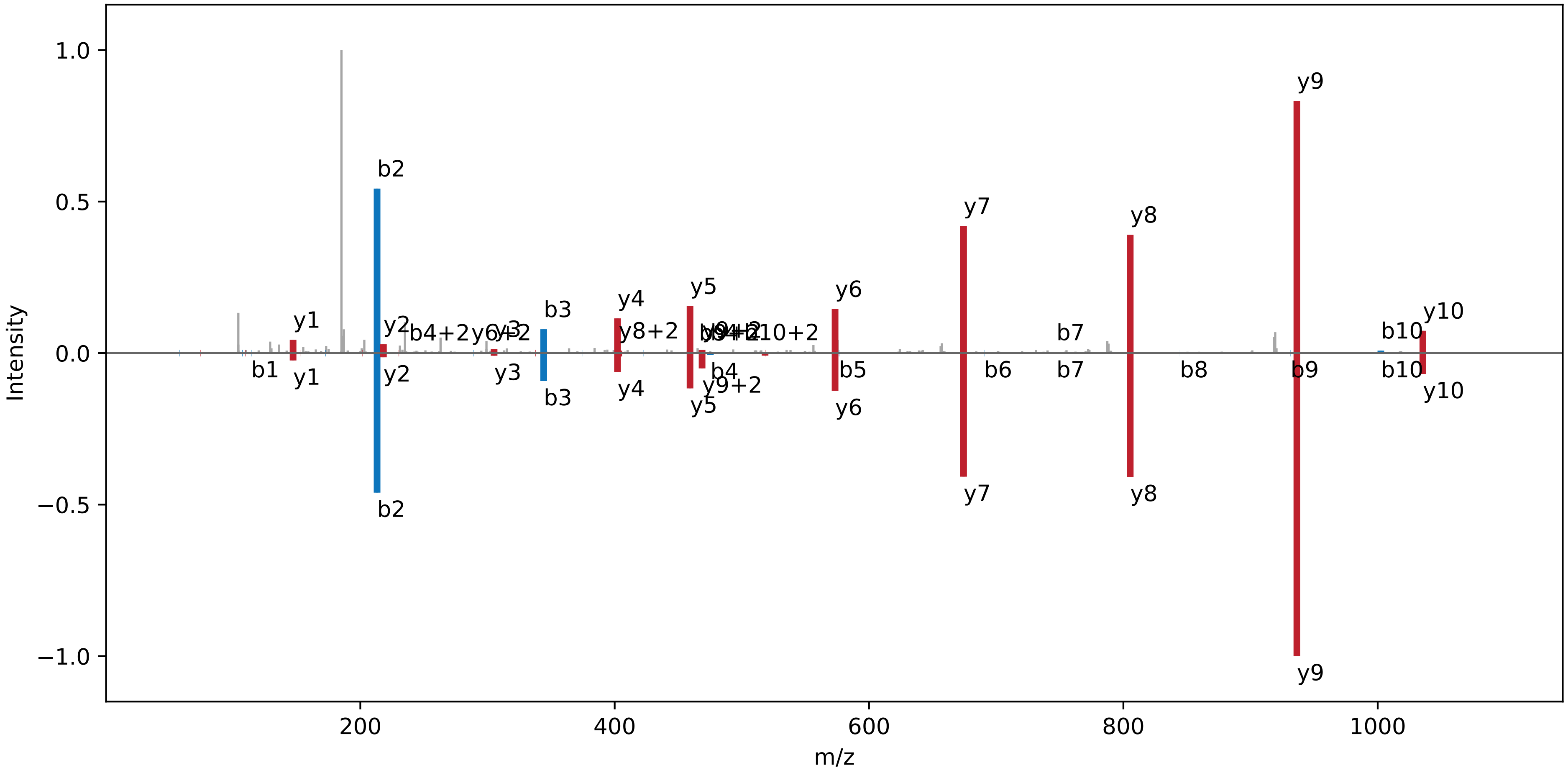

**NYALDFLFHR | NC_000911.1:1005181:−.pep23**

Charge: +2 | Collision Energy: 27.0 | Percursor m/z: 648.331238 | Mass Diff. 0.000610 Da

Percolator Score: 1.00 | Hyperscore: 29.08 | Spectral Angle: 0.89 | Pearson's Correlation: 0.98

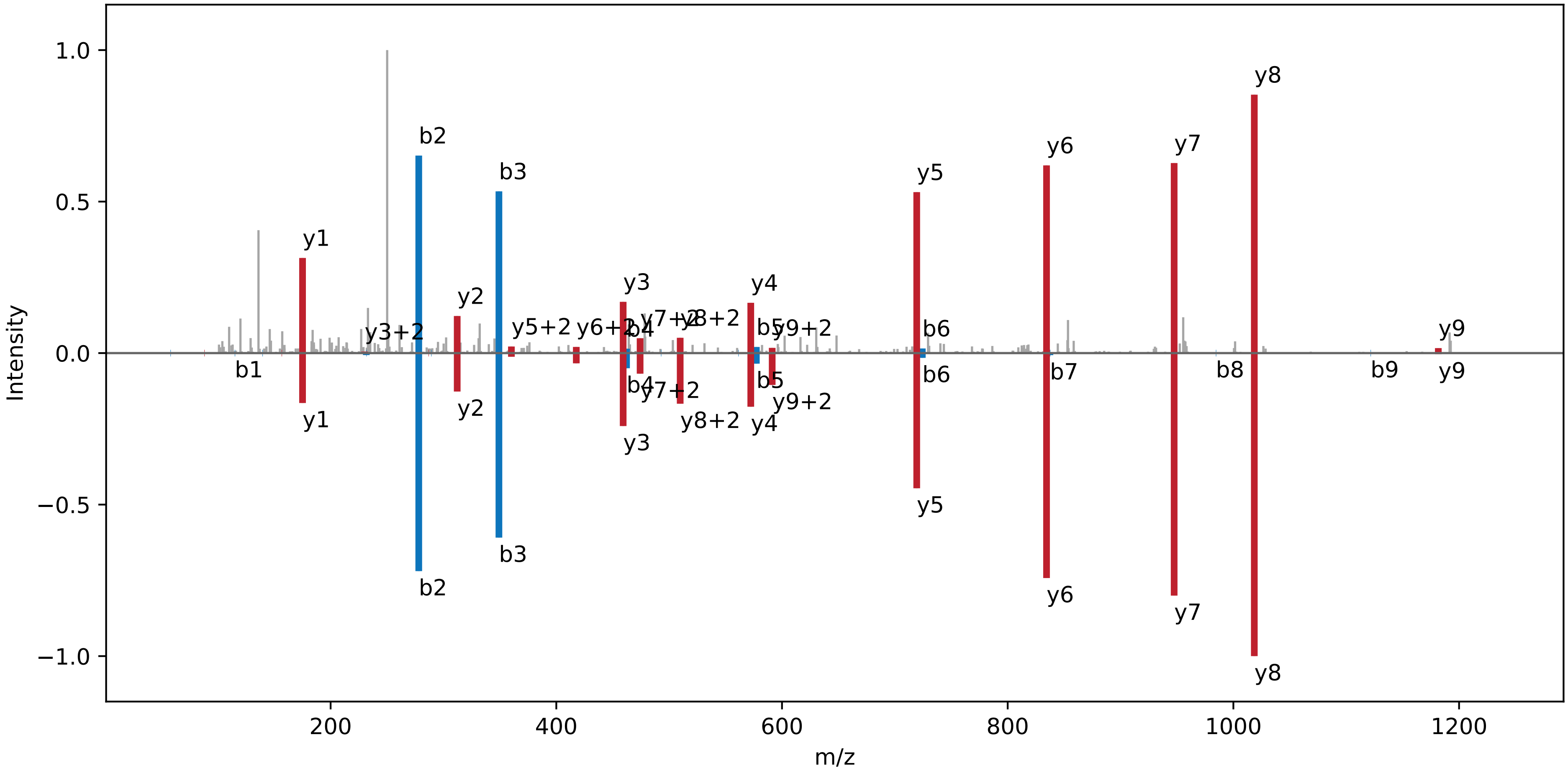

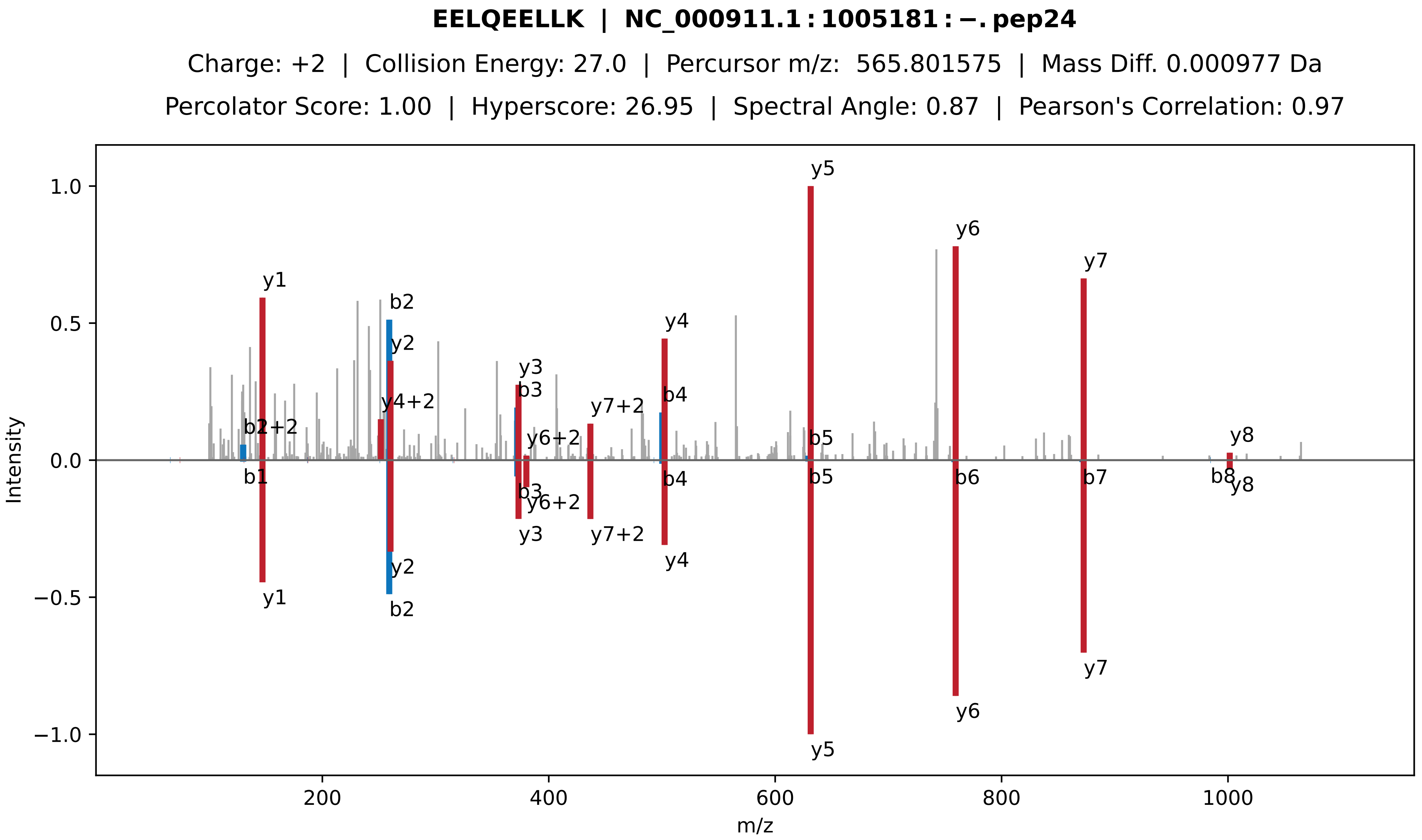
EELQEELLK | NC_000911.1 : 1005181 : −. pep24
Charge: +2 | Collision Energy: 27.0 | Percursor m/z: 565.801575 | Mass Diff. 0.000977 Da
Percolator Score: 1.00 | Hyperscore: 26.95 | Spectral Angle: 0.87 | Pearson's Correlation: 0.97
y1
b2
y2
y4+2
y3
b3
y6+2
y7+2
y4
b4
y5
b5
y6
y7
y8
b1
b6
b7
b8
1.0
0.5
0.0
−0.5
−1.0
Intensity
200
400
600
800
1000
m/z

**ALAISPEVLILDEPFGALDAITKEELQEELLK | NC_000911.1 : 1005181 : −. pep25**

Charge: +3 | Collision Energy: 27.0 | Percursor m/z: 1169.975100 | Mass Diff. 0.000977 Da

Percolator Score: 1.00 | Hyperscore: 74.46 | Spectral Angle: 0.00 | Pearson's Correlation: 0.00

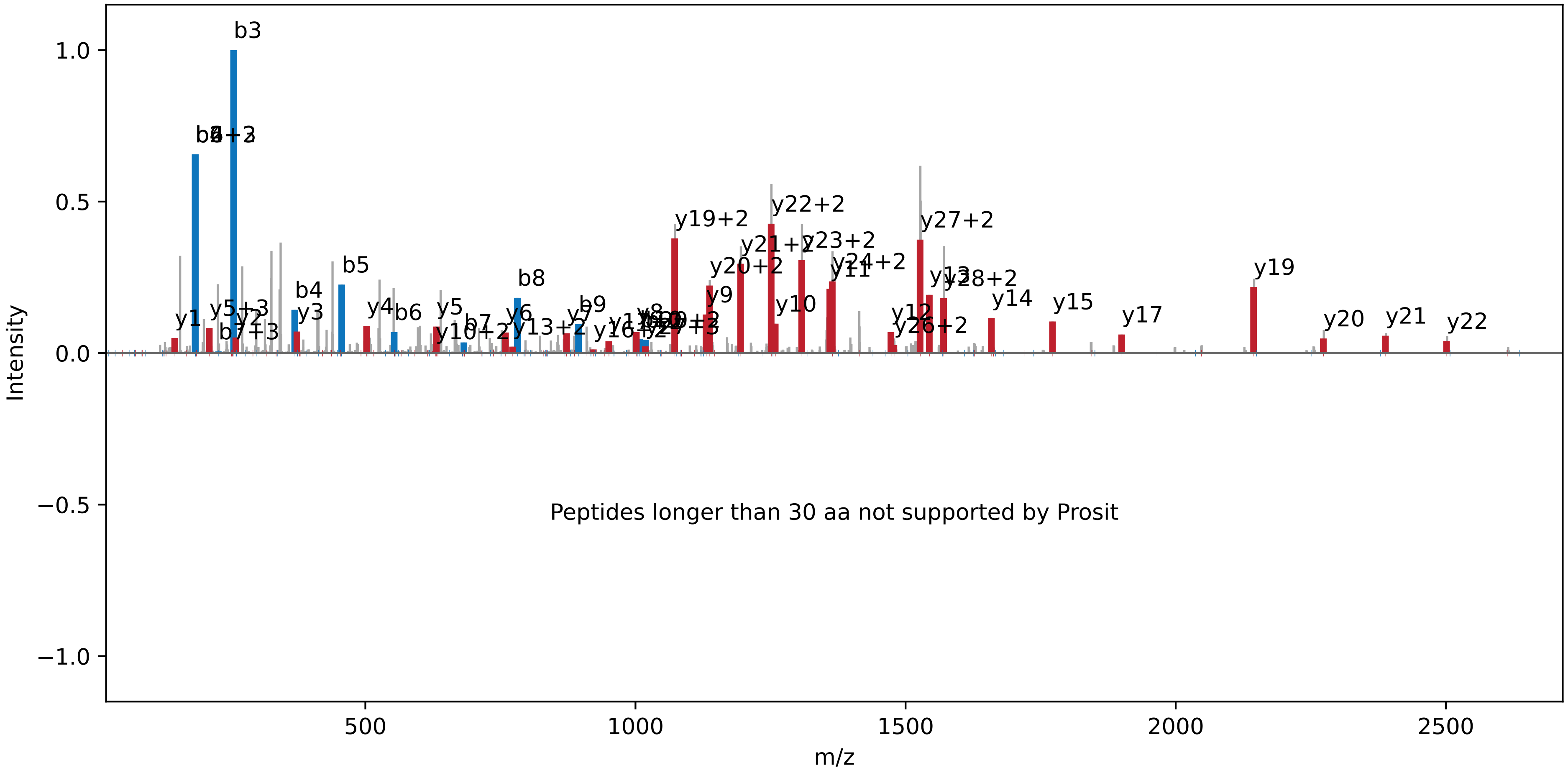

**ALAISPEVLILDEPFGALDAITK | NC_000911.1 : 1005181 : −. pep26**

Charge: +2 | Collision Energy: 27.0 | Percursor m/z: 1198.670776 | Mass Diff. -0.001709 Da

Percolator Score: 1.00 | Hyperscore: 64.13 | Spectral Angle: 0.82 | Pearson's Correlation: 0.95

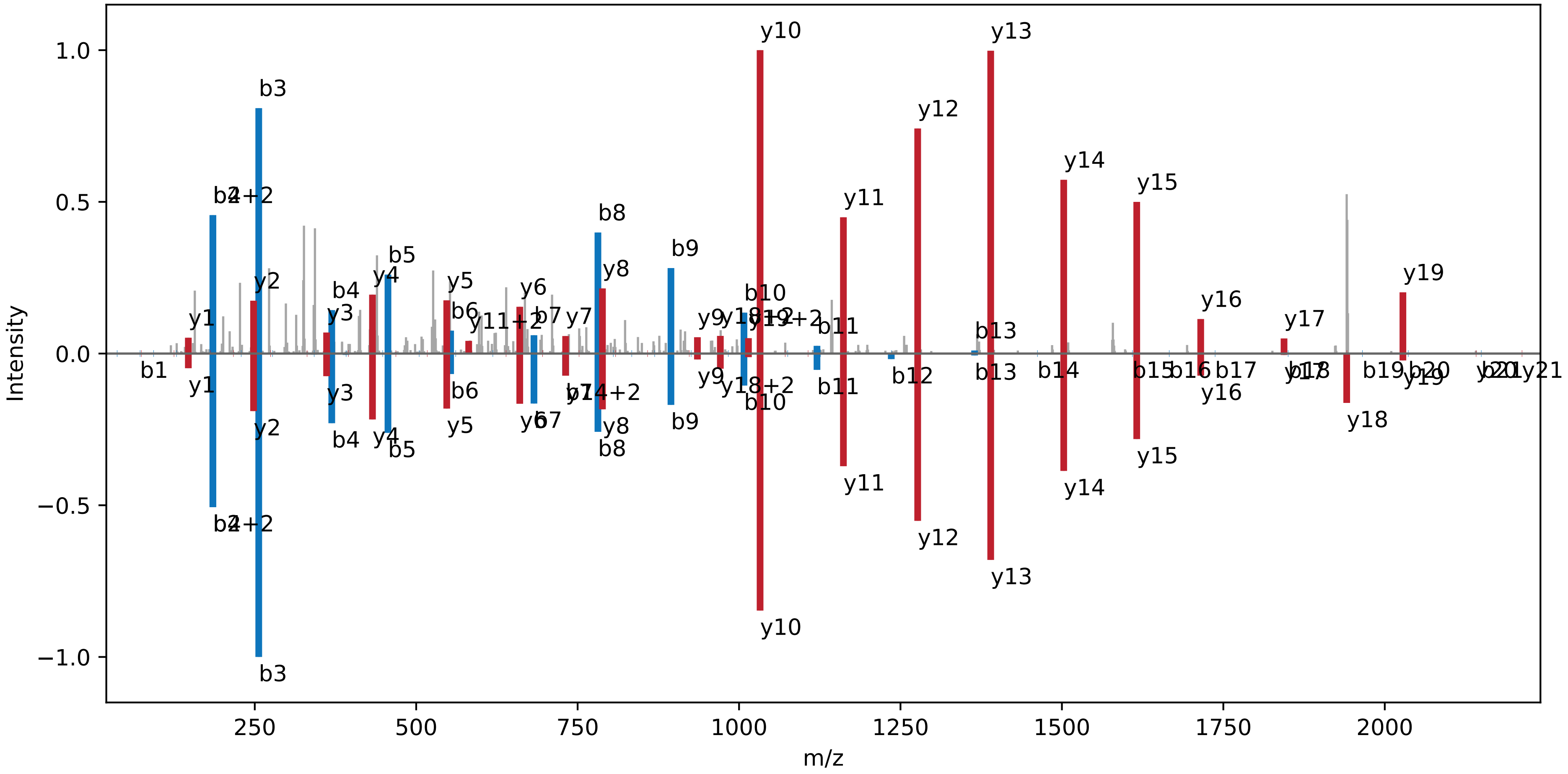

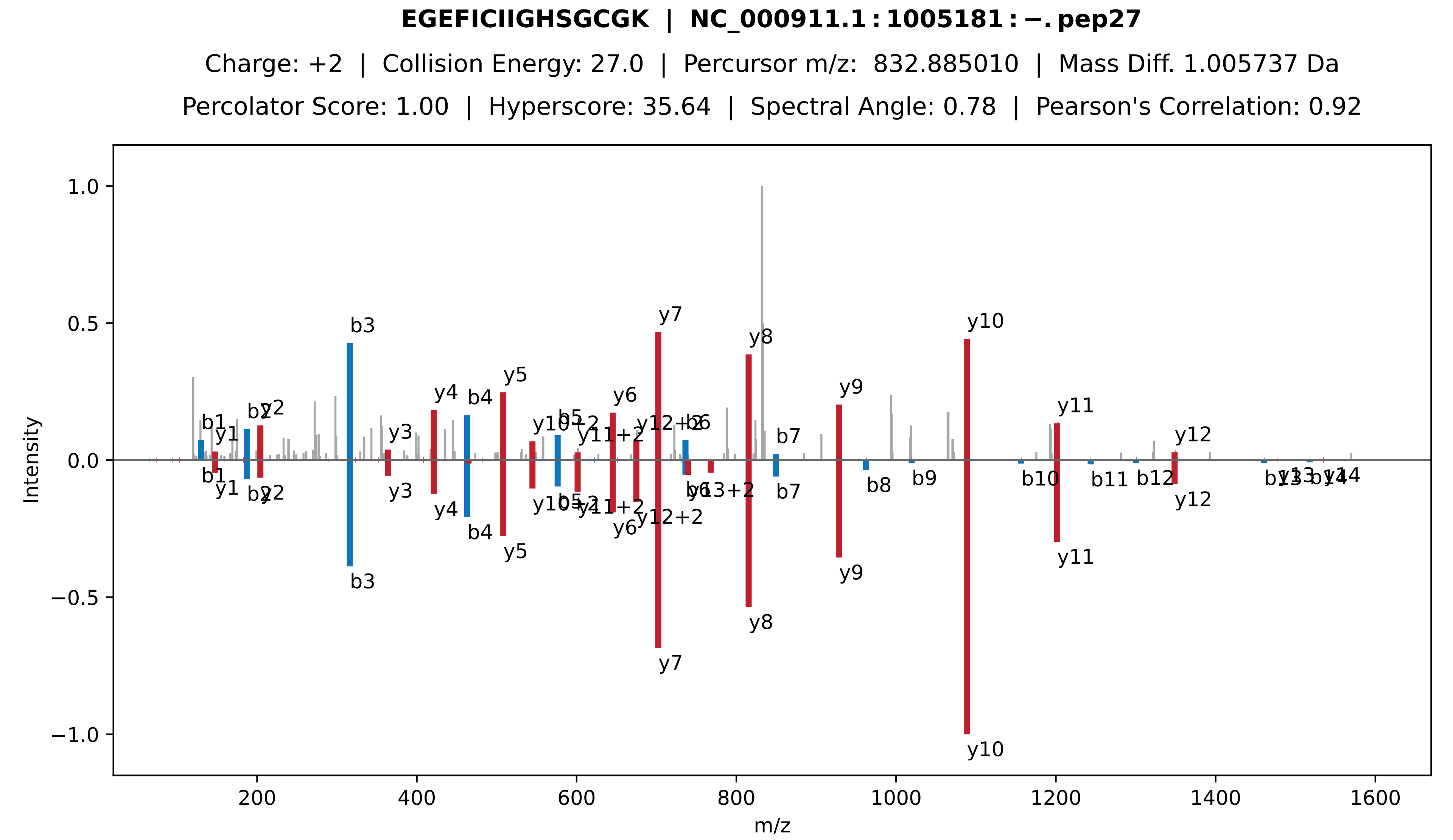

EGEFICIIGHSGCGK | NC_000911.1:1005181:−.pep27
Charge: +2 | Collision Energy: 27.0 | Percursor m/z: 832.885010 | Mass Diff. 1.005737 Da
Percolator Score: 1.00 | Hyperscore: 35.64 | Spectral Angle: 0.78 | Pearson's Correlation: 0.92
Intensity
m/z

**QQYAAIGPIK | NC_000911.1 : 2204654 : +. pep01**

Charge: +2 | Collision Energy: 27.0 | Percursor m/z: 544.808350 | Mass Diff. -0.000122 Da

Percolator Score: 1.00 | Hyperscore: 27.16 | Spectral Angle: 0.89 | Pearson's Correlation: 0.98

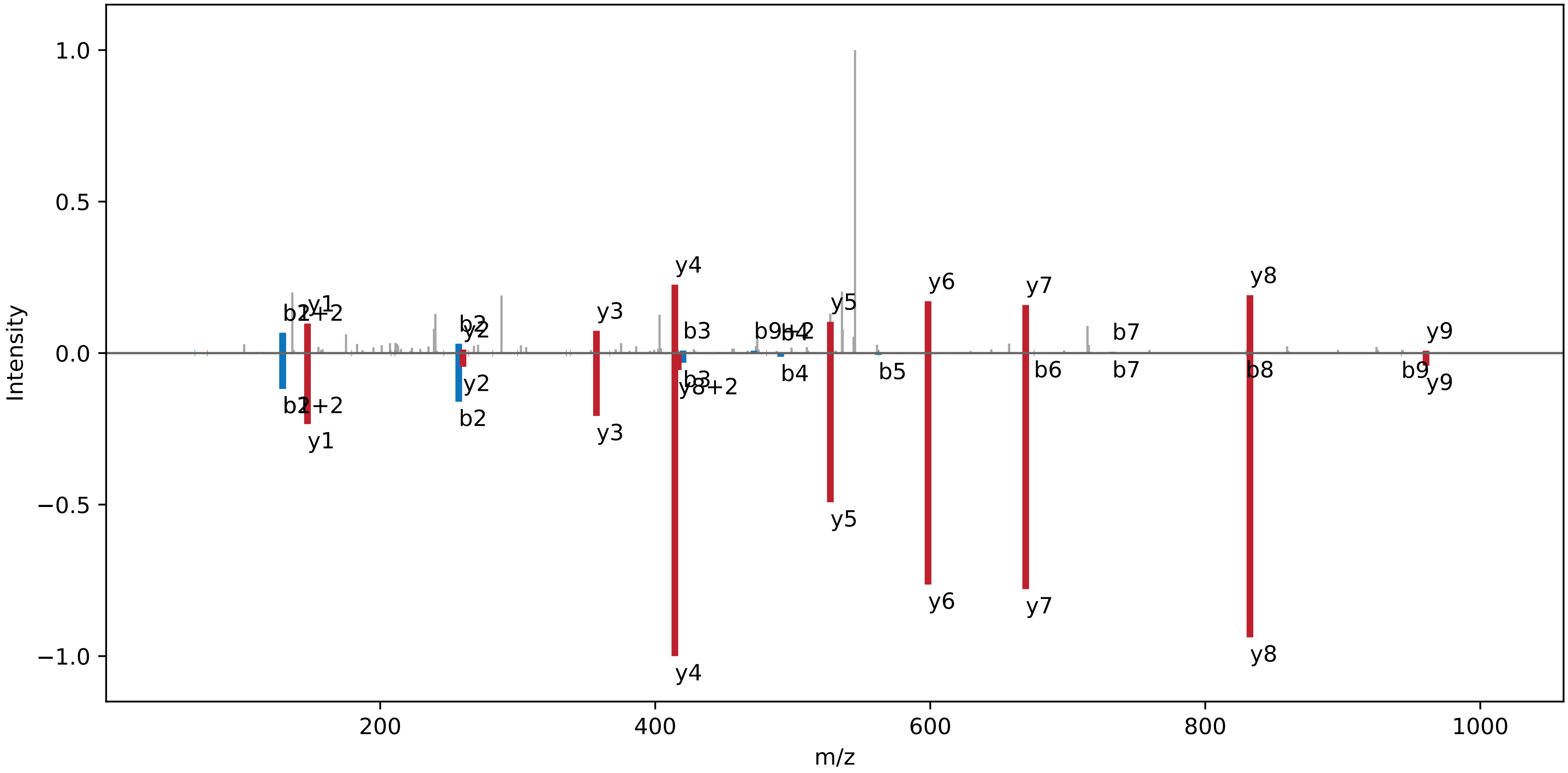

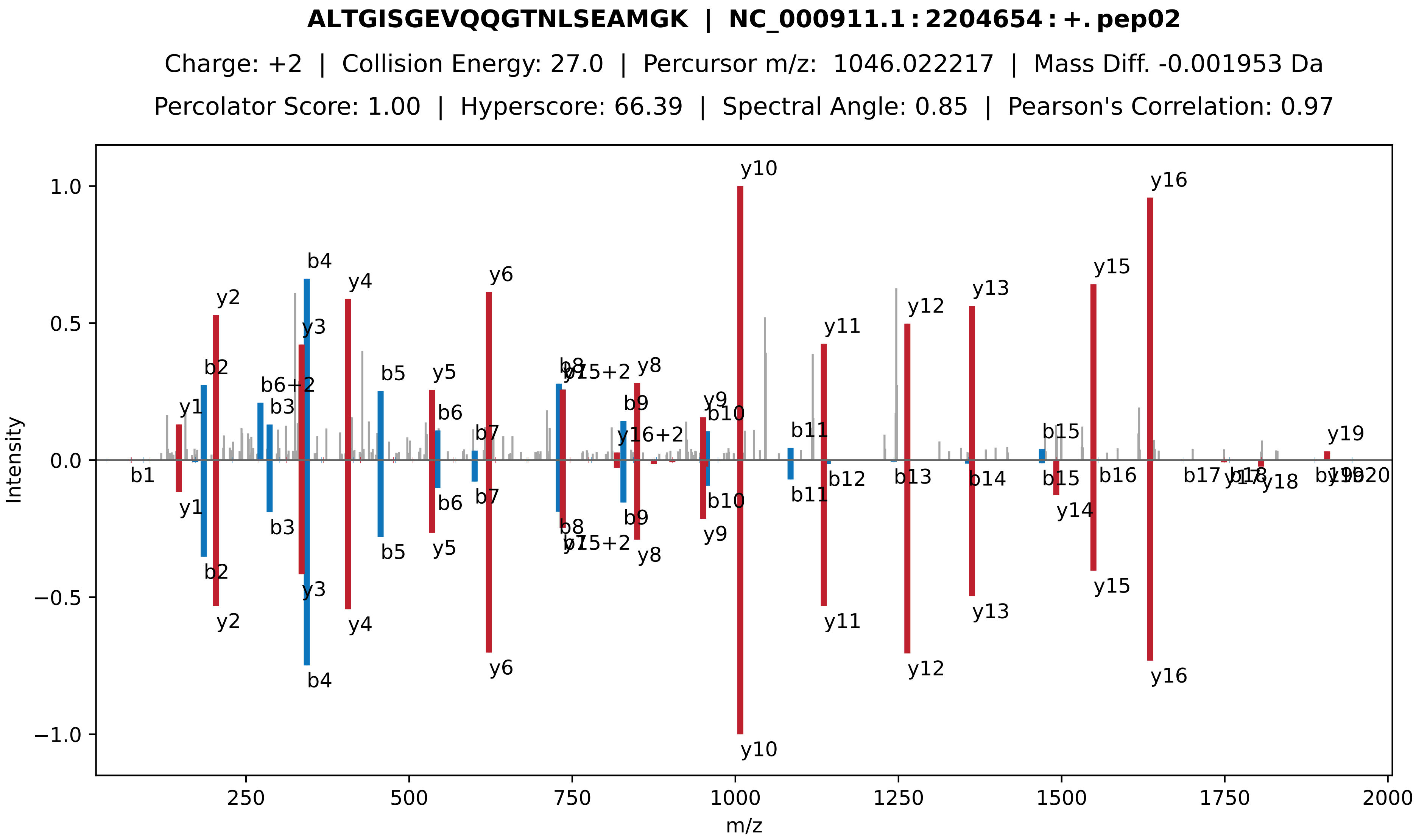
ALTGISGEVQQGTNLSEAMGK | NC_000911.1 : 2204654 : +. pep02
Charge: +2 | Collision Energy: 27.0 | Percursor m/z: 1046.022217 | Mass Diff. -0.001953 Da
Percolator Score: 1.00 | Hyperscore: 66.39 | Spectral Angle: 0.85 | Pearson's Correlation: 0.97
Intensity
m/z
y10
y16
y15
y13
y12
y11
y6
y4
y2
b4
y3
b2
b6+2
b3
b5
y5
b6
b7
y1
b8
y15+2
y8
b9
y16+2
y9
b10
b11
b12
b13
b14
b15
b16
b17
y17
y18
y19
y14
b1

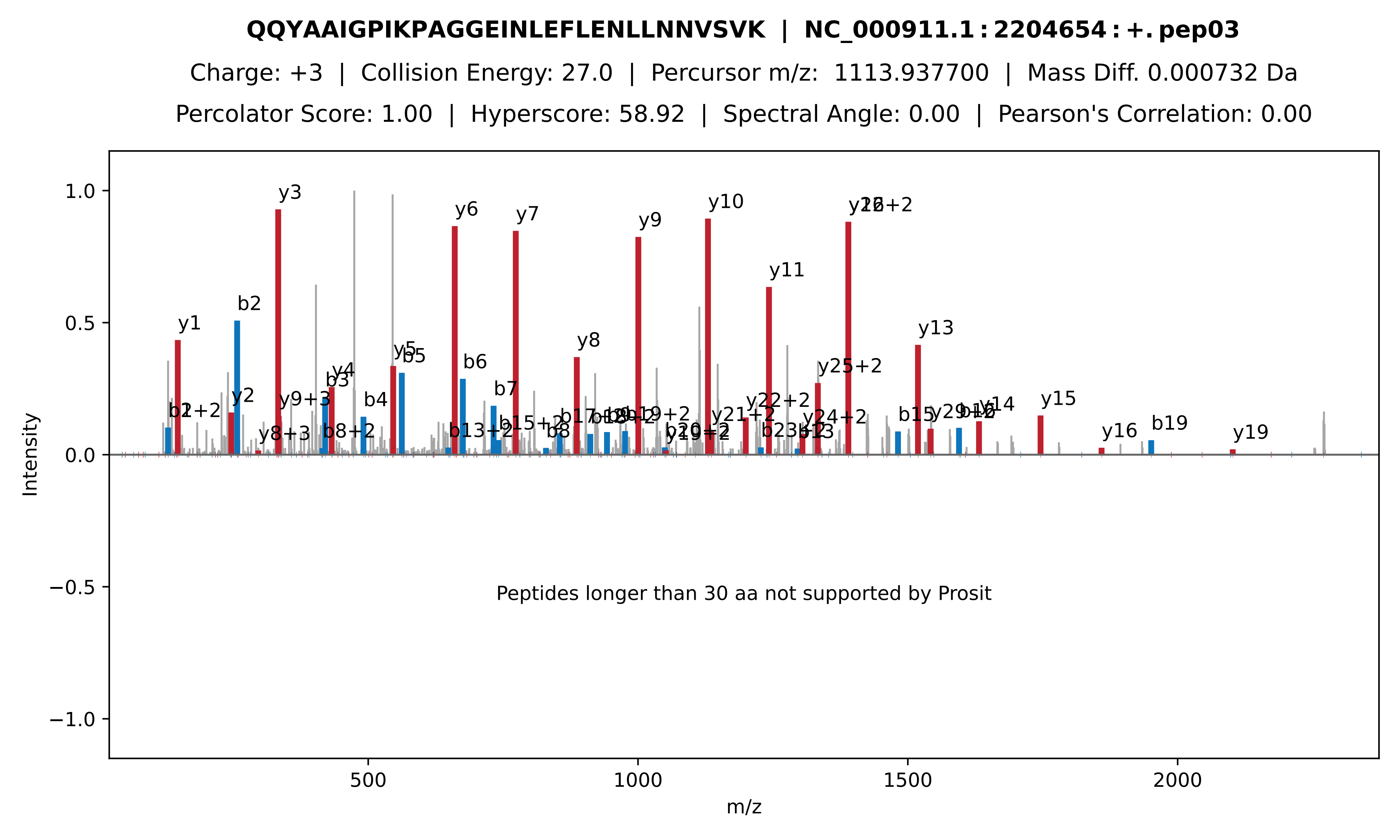
QQYAAIGPIKPAGGEINLEFLENLLNNVSVK | NC_000911.1:2204654:+.pep03
Charge: +3 | Collision Energy: 27.0 | Percursor m/z: 1113.937700 | Mass Diff. 0.000732 Da
Percolator Score: 1.00 | Hyperscore: 58.92 | Spectral Angle: 0.00 | Pearson's Correlation: 0.00
Intensity
m/z
Peptides longer than 30 aa not supported by Prosit
y1
b2
y2
y3
y9+3
y8+3
b3
y4
b8+2
b4
y5
b5
y6
b6
b13+2
b7
y7
b15+2
b8
y8
y9
y10
y11
y21+2
y22+2
y24+2
y25+2
y13
b15
y14
y15
y16
b19
y19

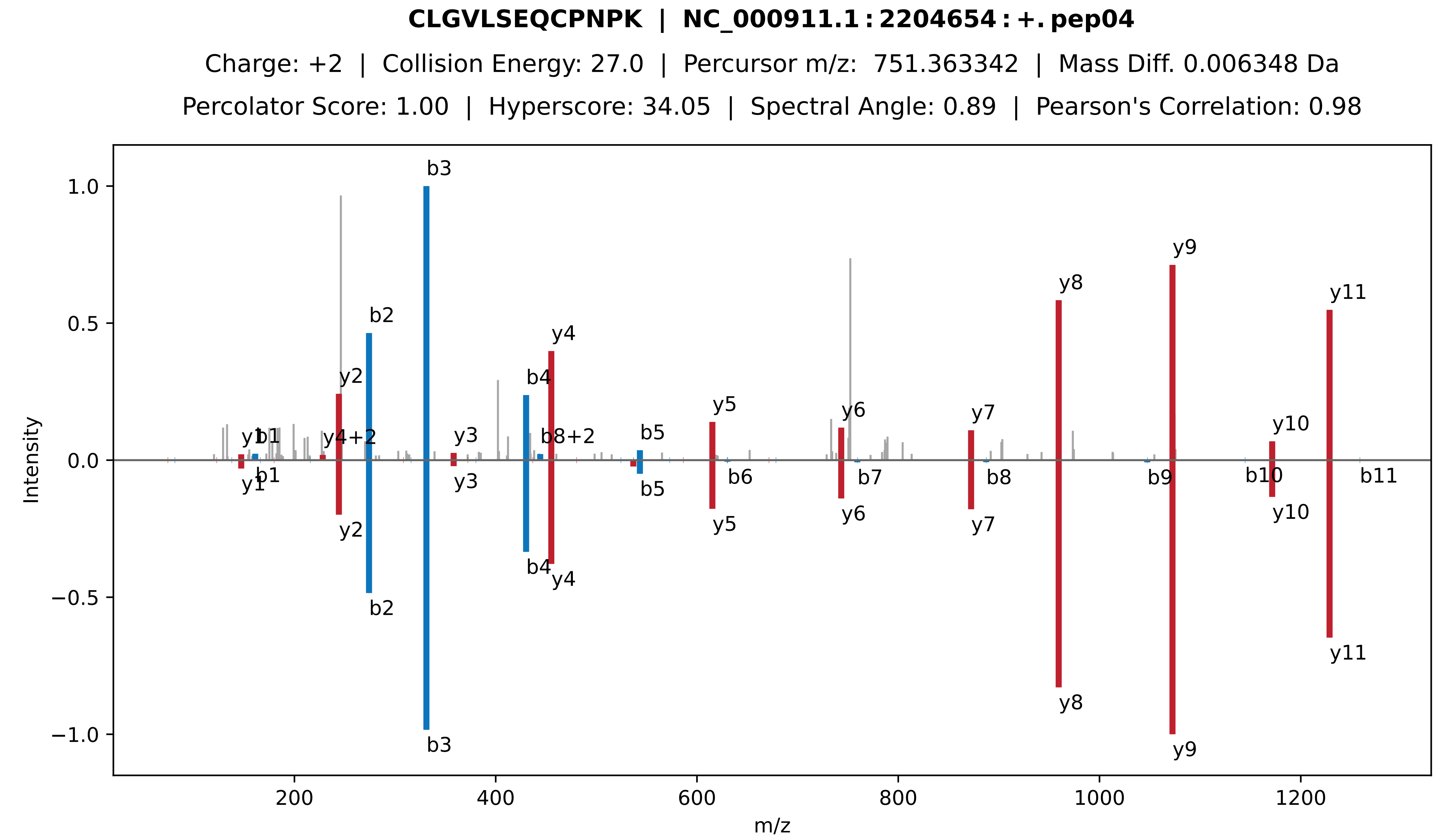

CLGVLSEQCPNPK | NC_000911.1:2204654:+.pep04
Charge: +2 | Collision Energy: 27.0 | Percursor m/z: 751.363342 | Mass Diff. 0.006348 Da
Percolator Score: 1.00 | Hyperscore: 34.05 | Spectral Angle: 0.89 | Pearson's Correlation: 0.98
Intensity
m/z

**ATFVAQVK | NC_000911.1 : 2204654 : +. pep05**

Charge: +2 | Collision Energy: 27.0 | Percursor m/z: 432.253479 | Mass Diff. -0.000488 Da

Percolator Score: 1.00 | Hyperscore: 20.82 | Spectral Angle: 0.96 | Pearson's Correlation: 1.00

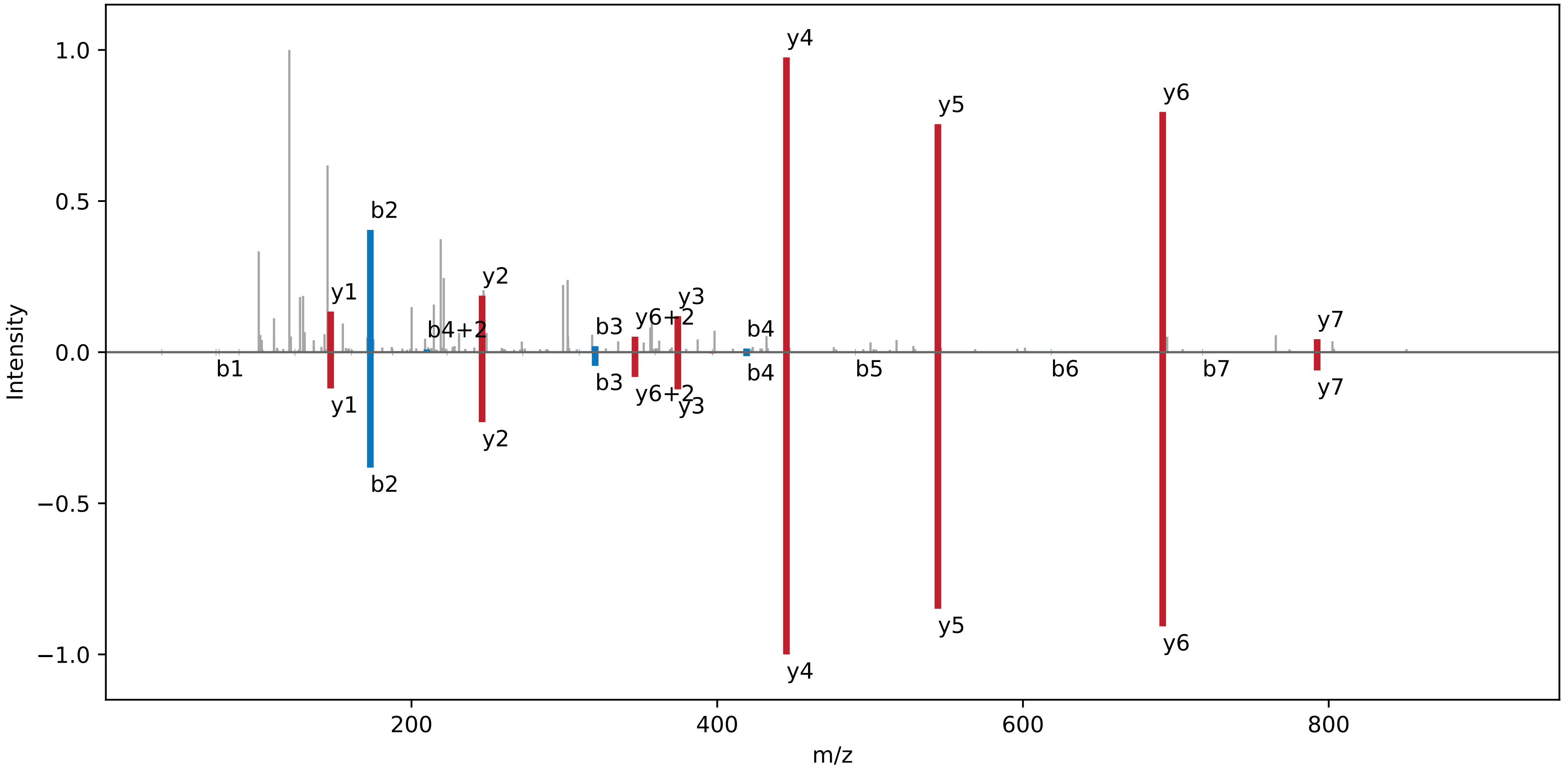

**RALTGISGEVQQGTNLSEAMGK | NC_000911.1 : 2204654 : +. pep06**

Charge: +3 | Collision Energy: 27.0 | Percursor m/z: 749.720600 | Mass Diff. 0.001953 Da

Percolator Score: 1.00 | Hyperscore: 57.49 | Spectral Angle: 0.87 | Pearson's Correlation: 0.98

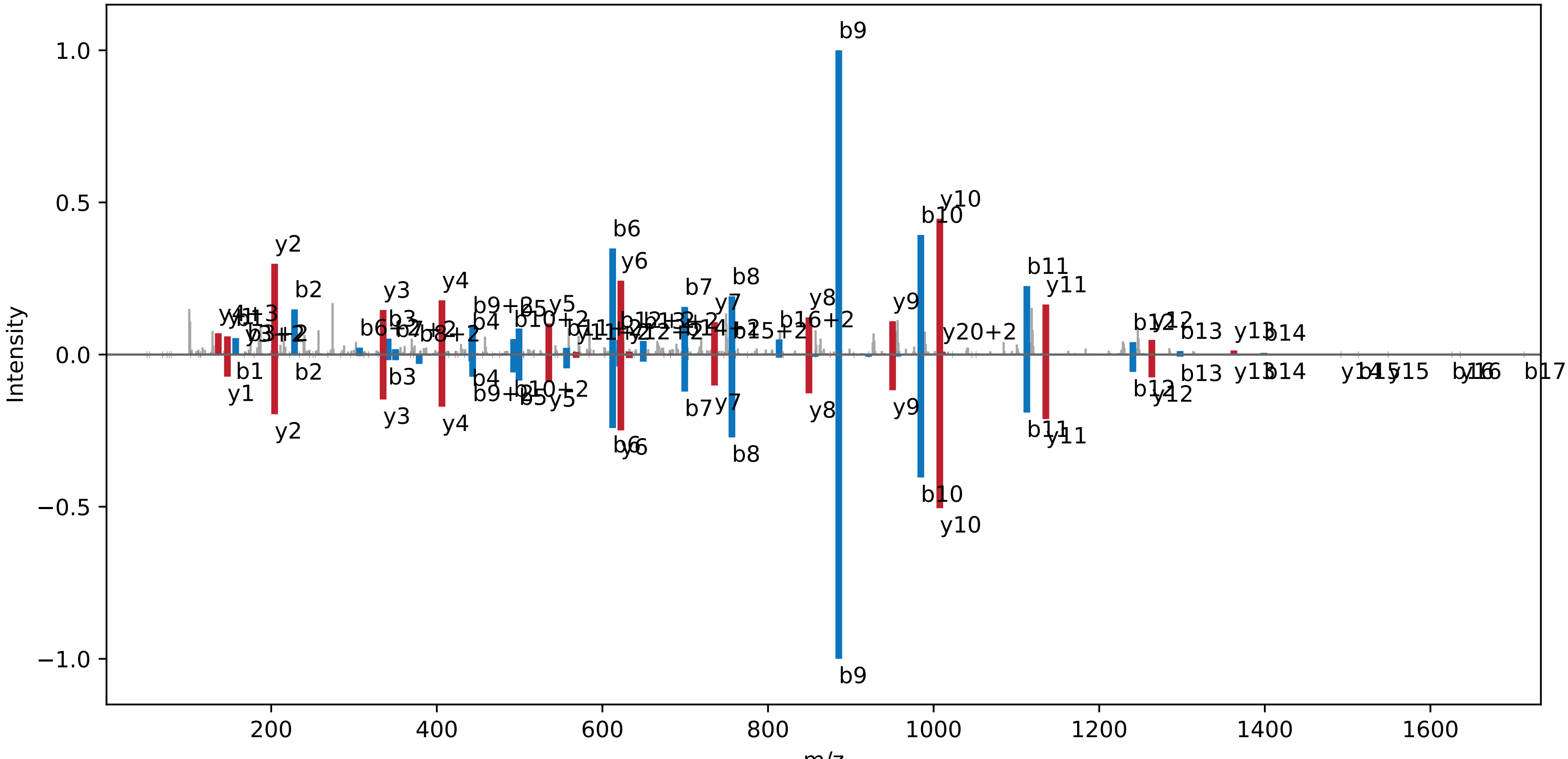

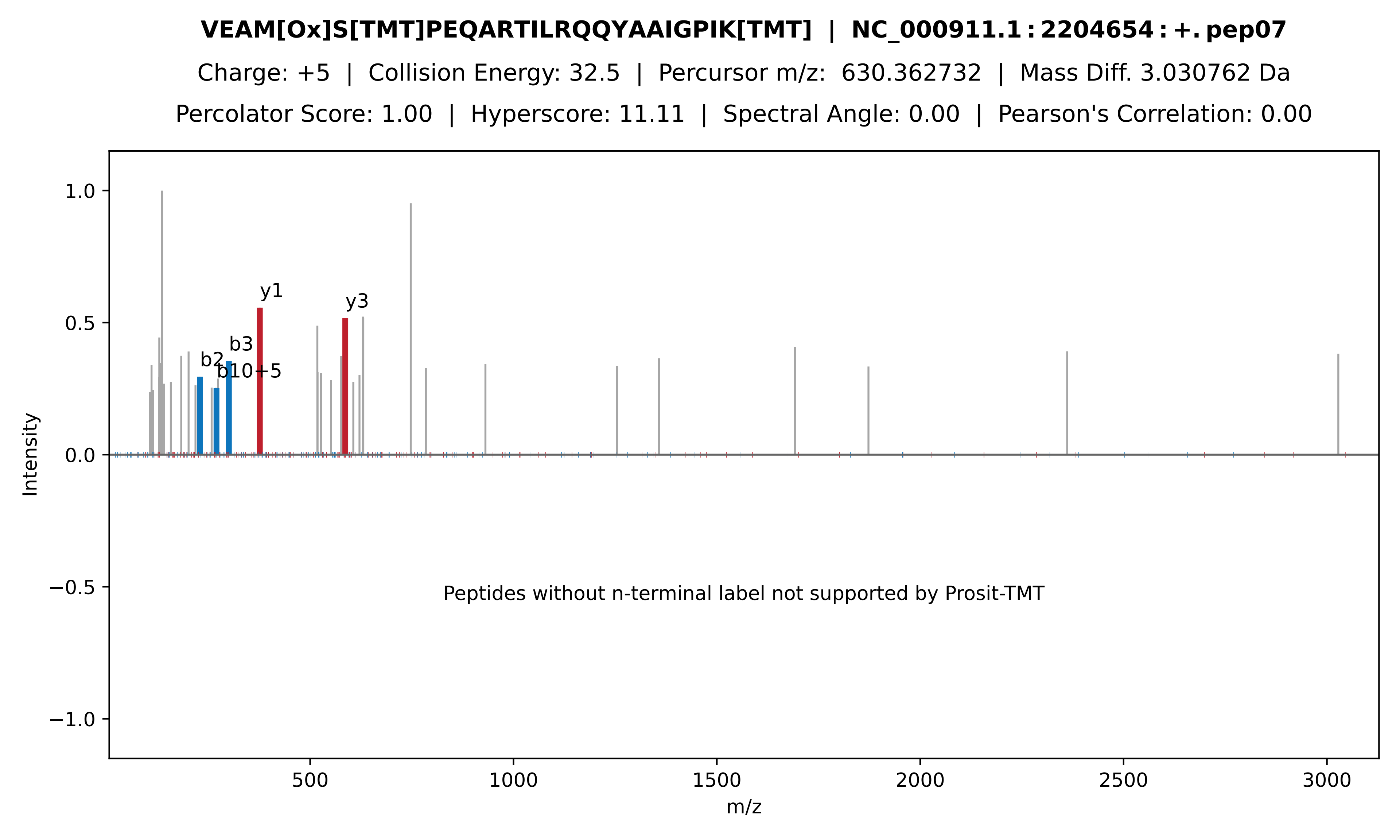

VEAM[Ox]S[TMT]PEQARTILRQQYAAIGPIK[TMT] | NC_000911.1 : 2204654 : +. pep07
Charge: +5 | Collision Energy: 32.5 | Percursor m/z: 630.362732 | Mass Diff. 3.030762 Da
Percolator Score: 1.00 | Hyperscore: 11.11 | Spectral Angle: 0.00 | Pearson's Correlation: 0.00
y1
y3
b3
b2
b10+5
1.0
0.5
0.0
−0.5
−1.0
Intensity
Peptides without n-terminal label not supported by Prosit-TMT
500
1000
1500
2000
2500
3000
m/z

**PAGGEINLEFLENLLNNVSVK | NC_000911.1 : 2204654 : +. pep08**

Charge: +3 | Collision Energy: 27.0 | Percursor m/z: 757.406500 | Mass Diff. -0.002686 Da

Percolator Score: 1.00 | Hyperscore: 57.85 | Spectral Angle: 0.82 | Pearson's Correlation: 0.95

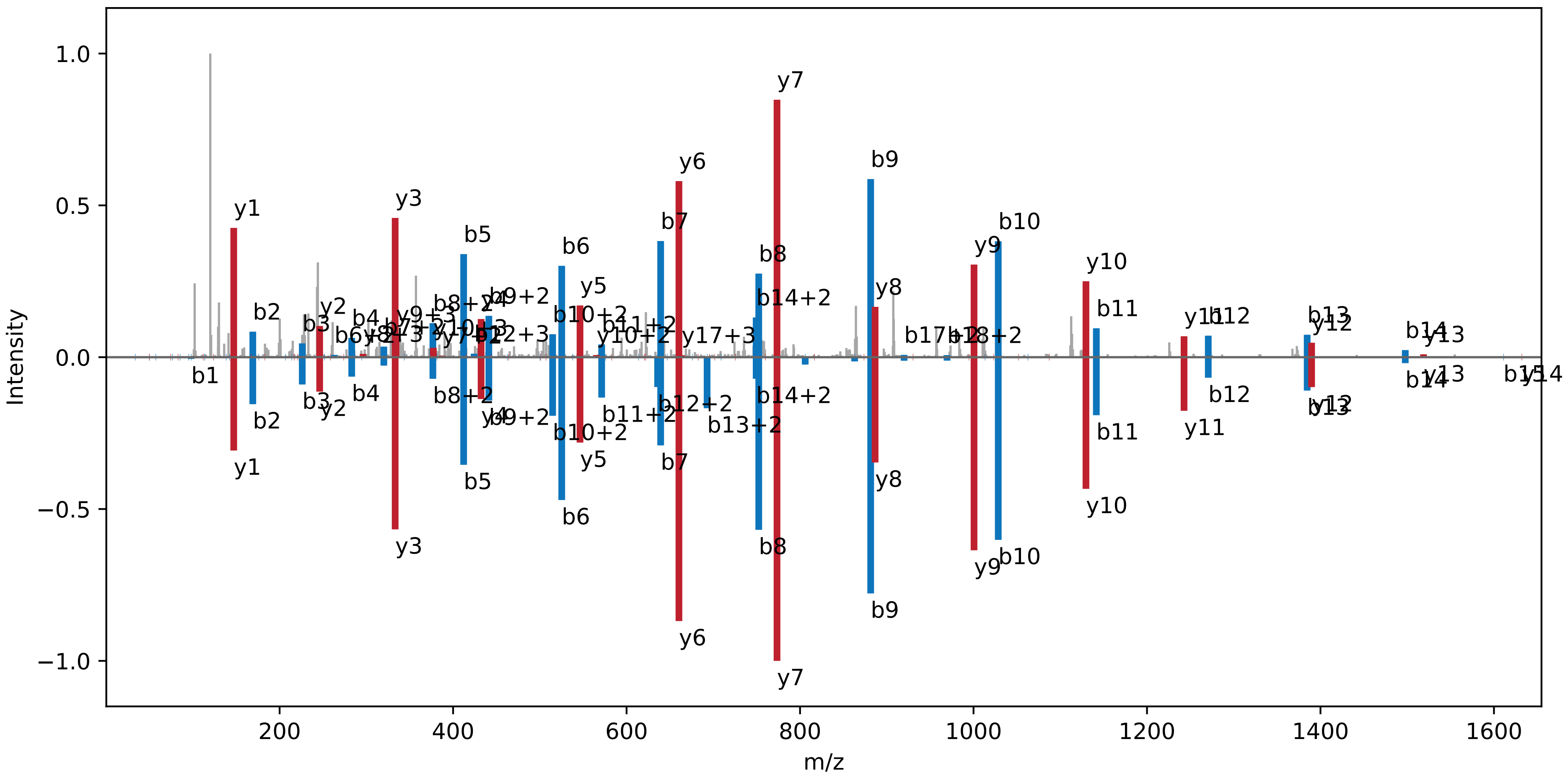

**AKVEAMSPEQAR | NC_000911.1:2204654:+.pep09**

Charge: +2 | Collision Energy: 27.0 | Percursor m/z: 658.835938 | Mass Diff. -0.000854 Da

Percolator Score: 1.00 | Hyperscore: 39.76 | Spectral Angle: 0.94 | Pearson's Correlation: 0.99

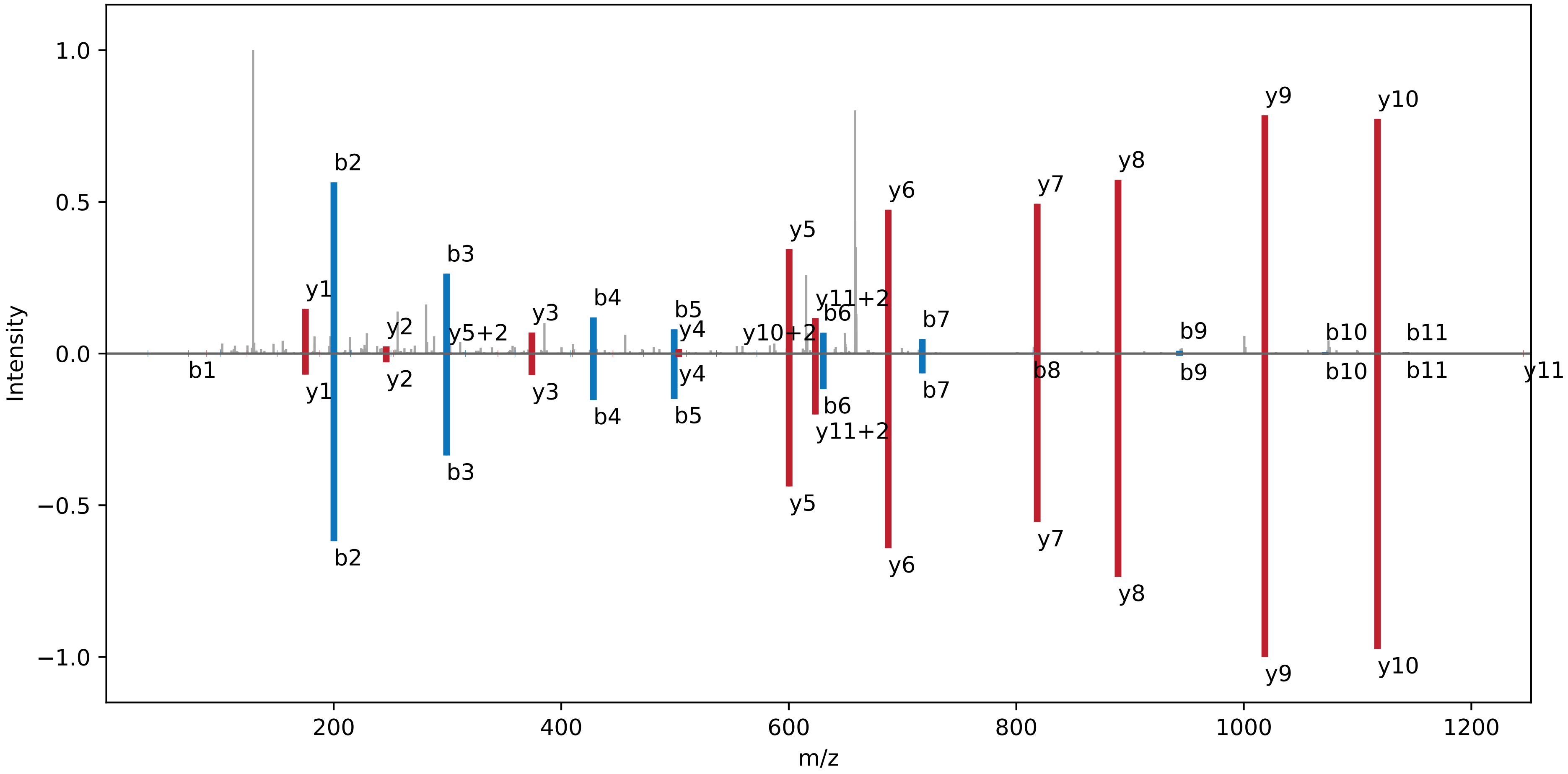

**VEAM[Ox]SPEQAR | NC_000911.1 : 2204654 : +. pep10**

Charge: +2 | Collision Energy: 27.0 | Percursor m/z: 567.265137 | Mass Diff. -0.001465 Da

Percolator Score: 1.00 | Hyperscore: 26.52 | Spectral Angle: 0.87 | Pearson's Correlation: 0.98

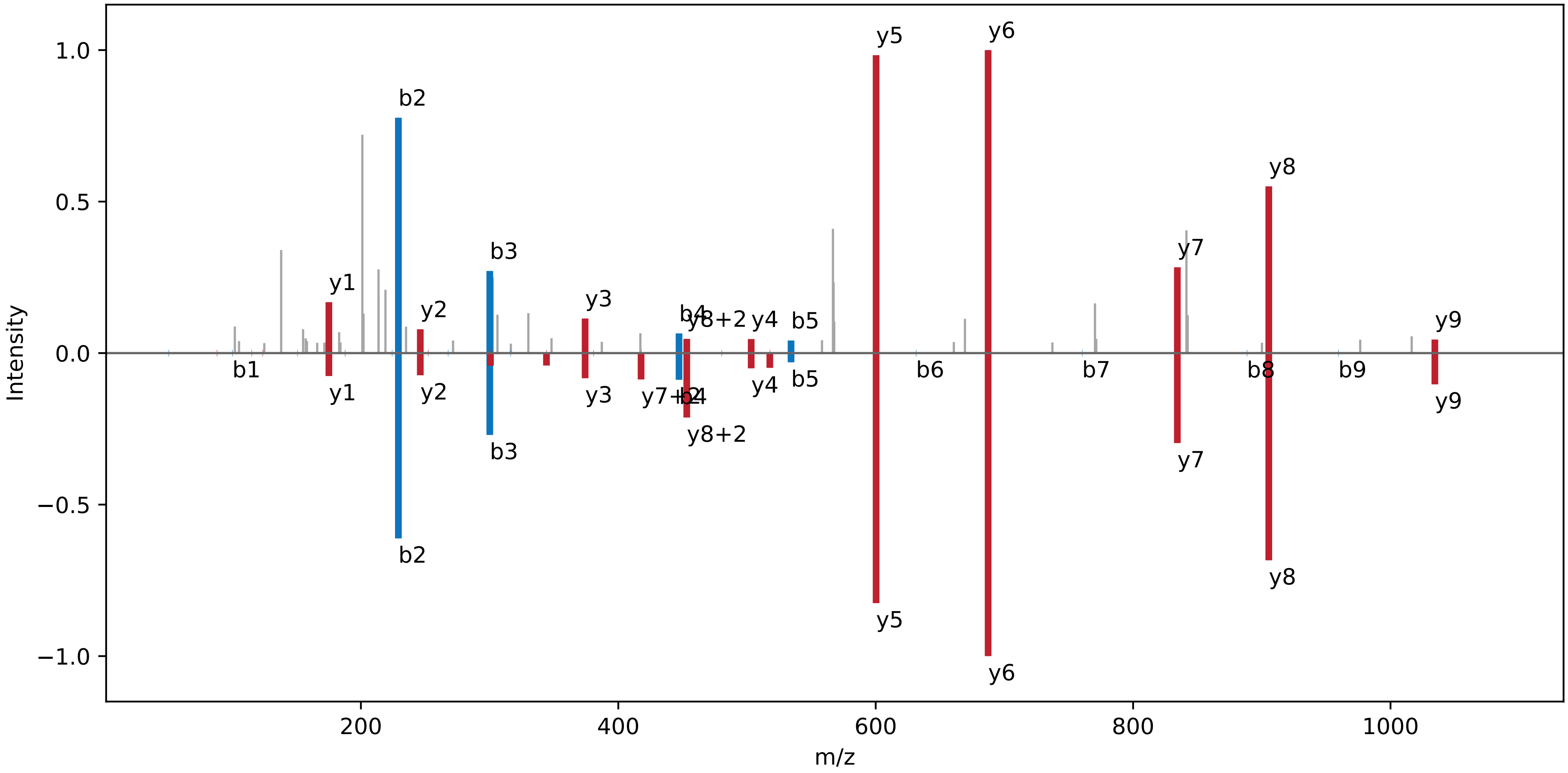

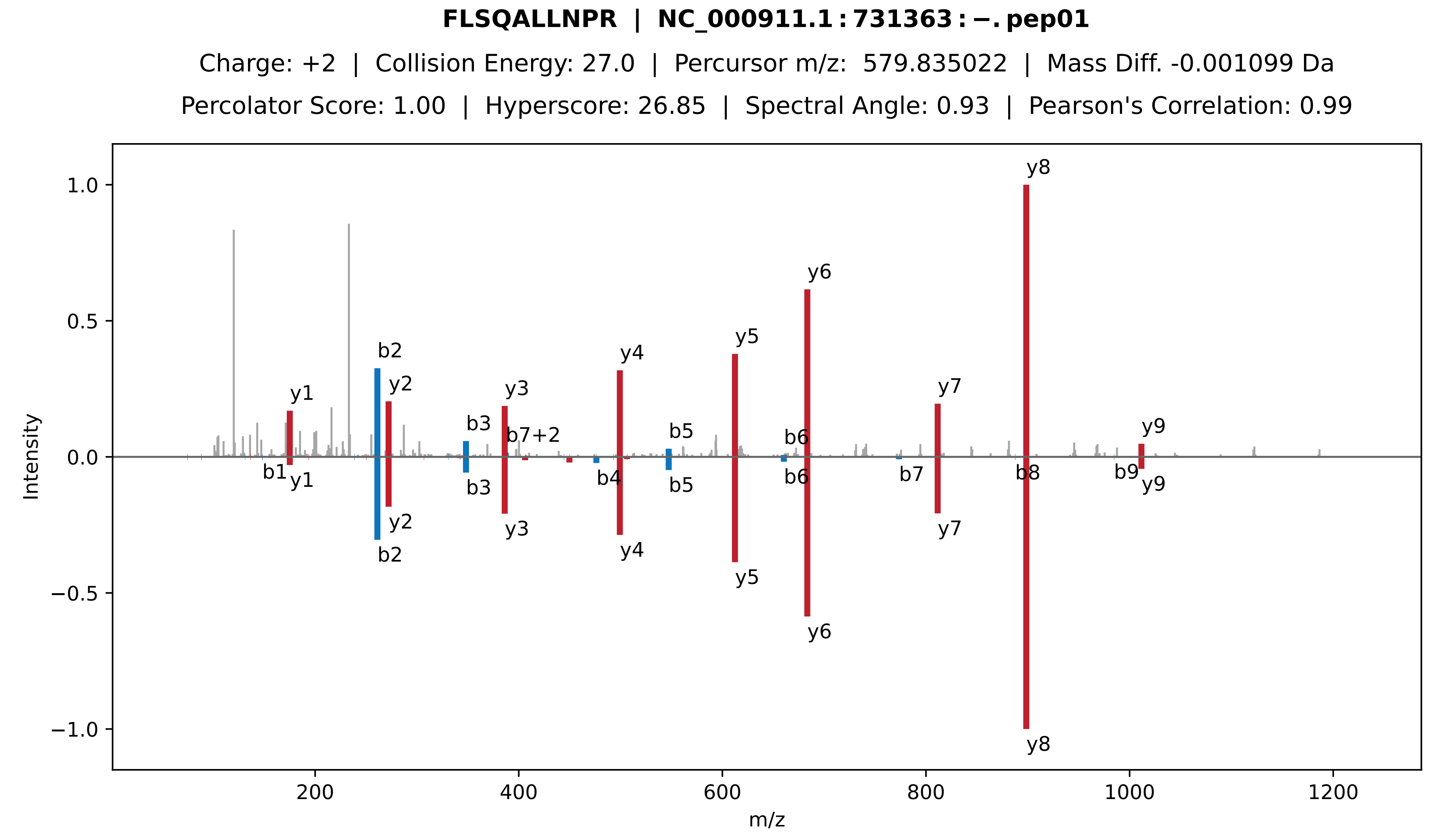
FLSQALLNPR | NC_000911.1 : 731363 : −. pep01
Charge: +2 | Collision Energy: 27.0 | Percursor m/z: 579.835022 | Mass Diff. -0.001099 Da
Percolator Score: 1.00 | Hyperscore: 26.85 | Spectral Angle: 0.93 | Pearson's Correlation: 0.99
Intensity
m/z
y1
b1
b2
y2
b3
y3
b7+2
b4
y4
b5
y5
b6
y6
b7
y7
y8
b8
b9
y9
1.0
0.5
0.0
−0.5
−1.0
200
400
600
800
1000
1200

**LLANRYQLVELVGSGAMGQVYRAEDK | NC_000911.1 : 731363 : −. pep02**

Charge: +4 | Collision Energy: 27.0 | Percursor m/z: 721.384583 | Mass Diff. 1.996826 Da

Percolator Score: 1.00 | Hyperscore: 31.81 | Spectral Angle: 0.65 | Pearson's Correlation: 0.84

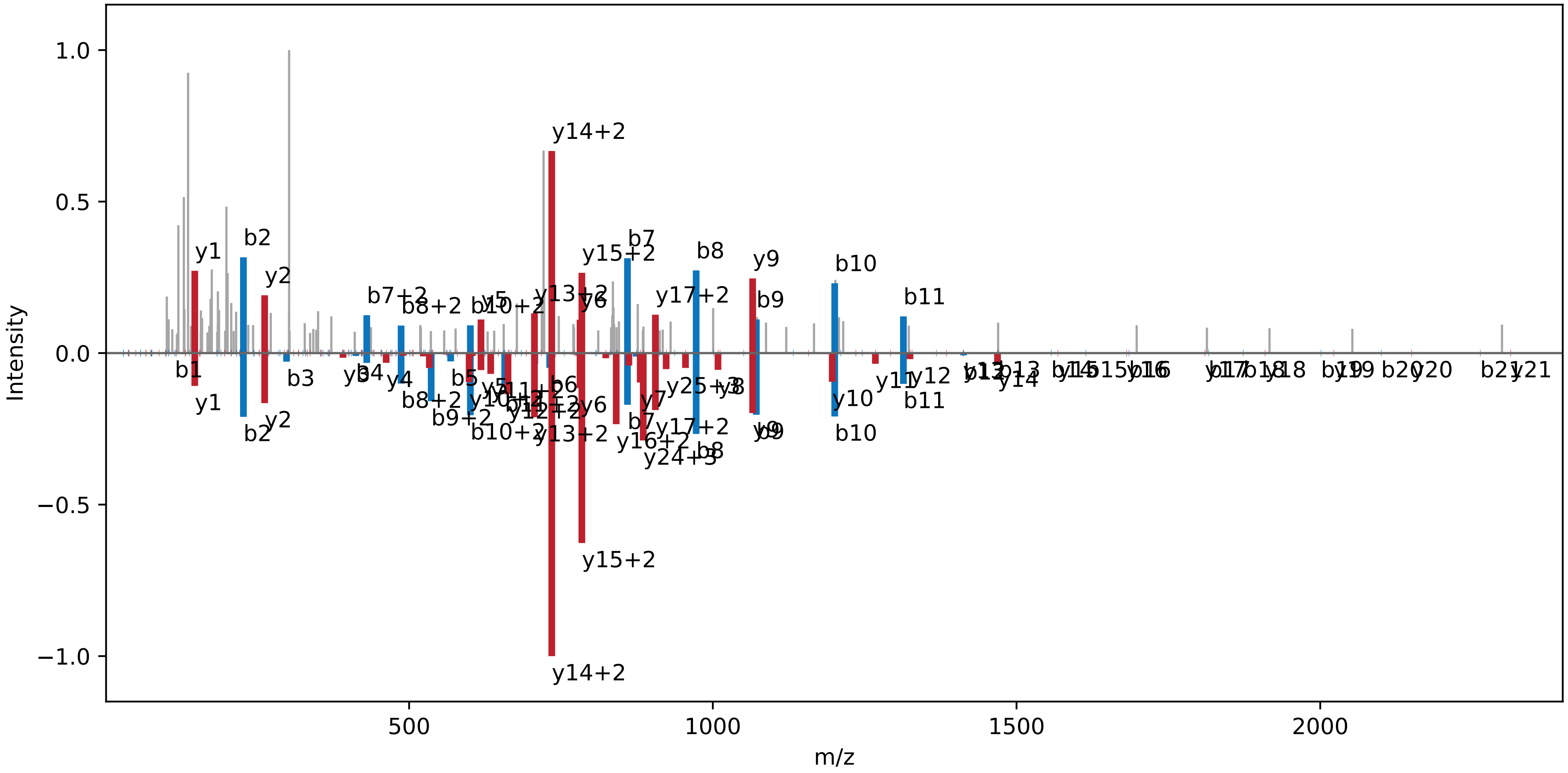

**LLGGVTVAVK | NC_000911.1 : 731363 : −. pep03**

Charge: +2 | Collision Energy: 27.0 | Percursor m/z: 478.810883 | Mass Diff. 0.000488 Da

Percolator Score: 1.00 | Hyperscore: 24.59 | Spectral Angle: 0.90 | Pearson's Correlation: 0.99

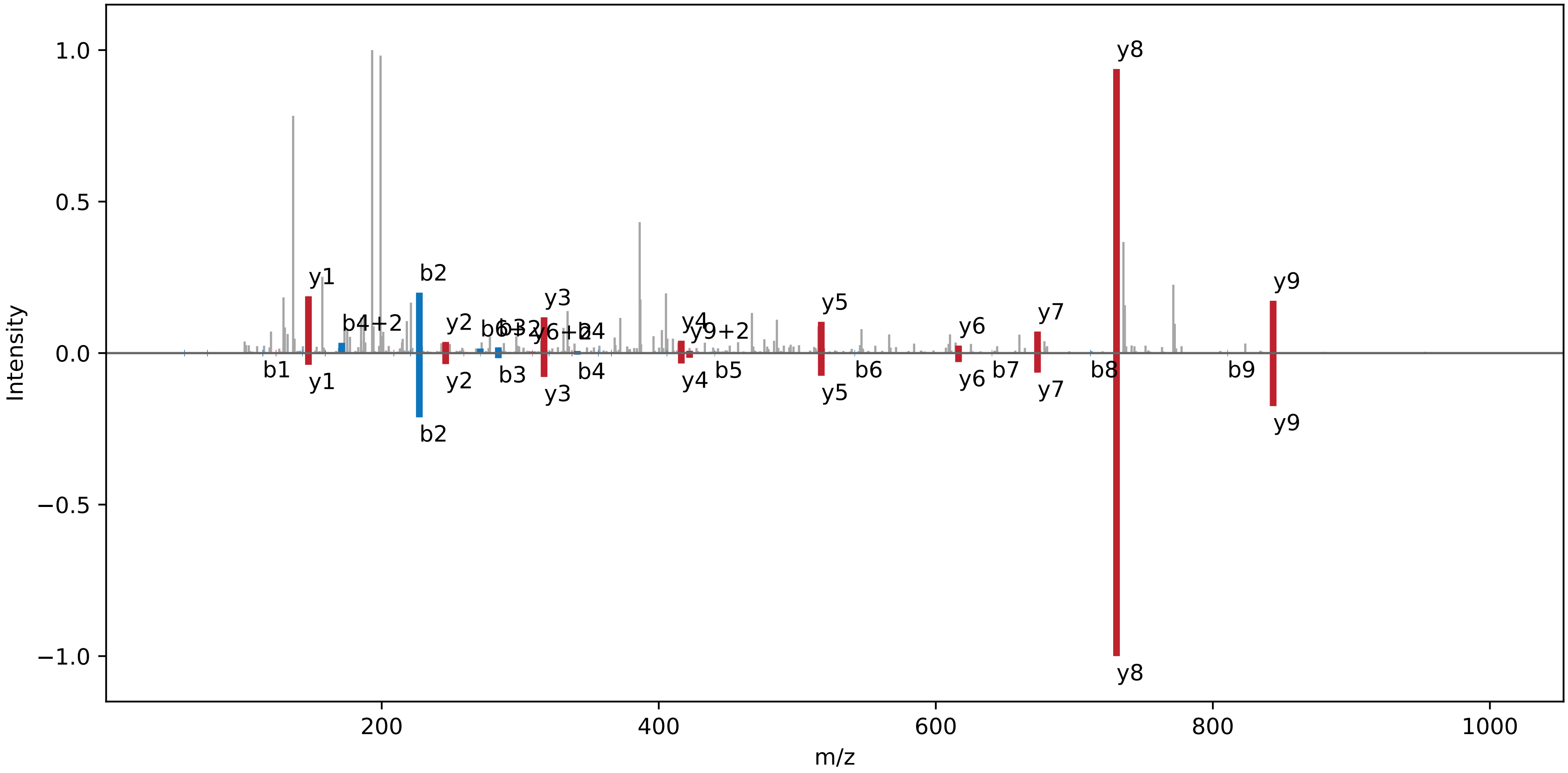

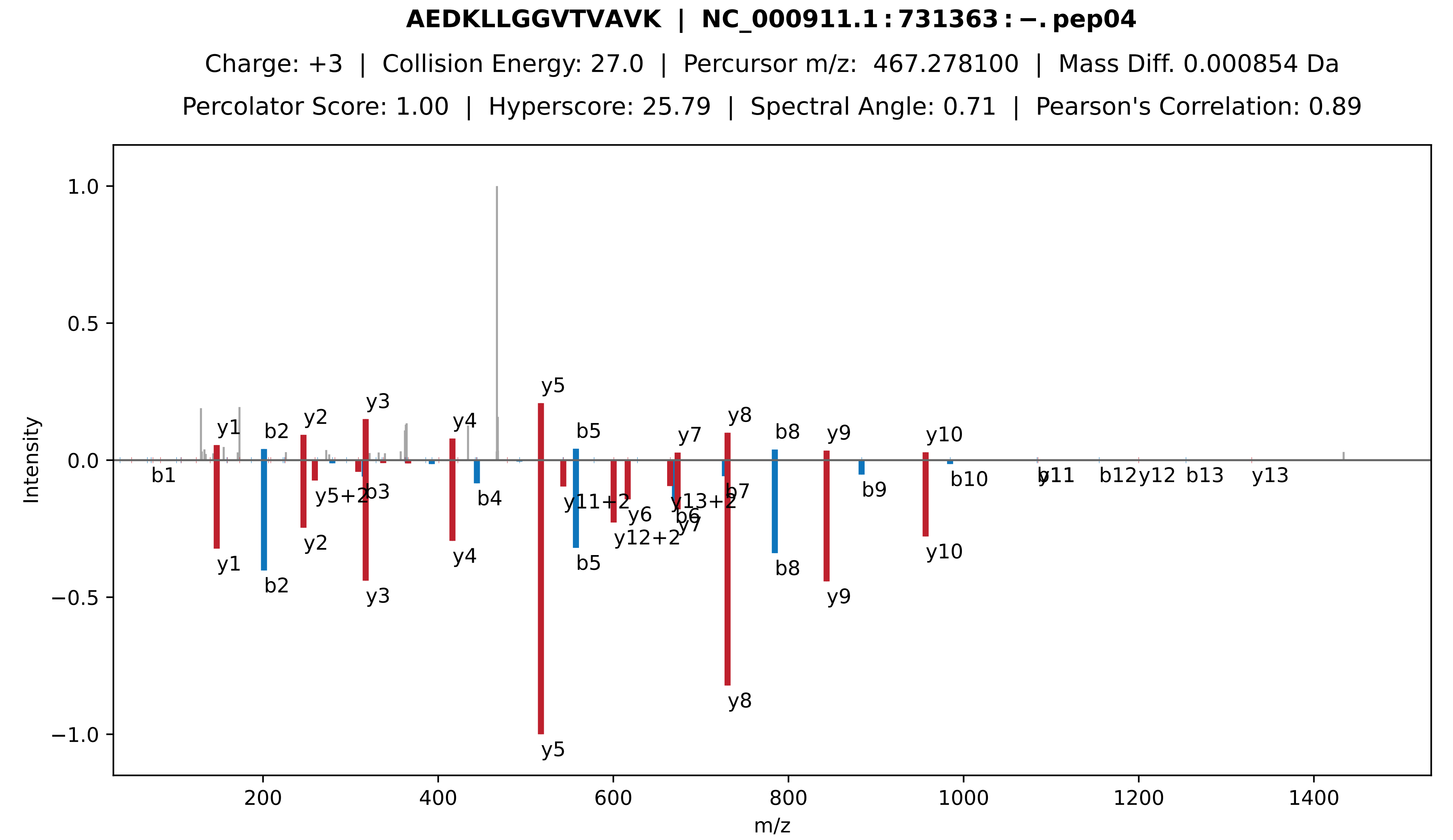
**AEDKLLGGVTVAVK | NC_000911.1 : 731363 : −. pep04**
Charge: +3 | Collision Energy: 27.0 | Percursor m/z: 467.278100 | Mass Diff. 0.000854 Da
Percolator Score: 1.00 | Hyperscore: 25.79 | Spectral Angle: 0.71 | Pearson's Correlation: 0.89
Intensity
m/z
1.0
0.5
0.0
−0.5
−1.0
200
400
600
800
1000
1200
1400
y1
b2
y2
y3
y4
y5
b5
y7
y8
b8
y9
y10
b1
y5+2
b3
b4
y11+2
y6
y12+2
y13+2
b6
y7
b7
b9
b10
b11
y11
b12
y12
b13
y13
y1
b2
y2
y3
y4
y5
b5
b8
y8
y9
y10

**YQLVELVGSGAMGQVYRAEDK | NC_000911.1 : 731363 : −. pep05**

Charge: +3 | Collision Energy: 27.0 | Percursor m/z: 772.064209 | Mass Diff. 1.035645 Da

Percolator Score: 1.00 | Hyperscore: 49.50 | Spectral Angle: 0.73 | Pearson's Correlation: 0.91

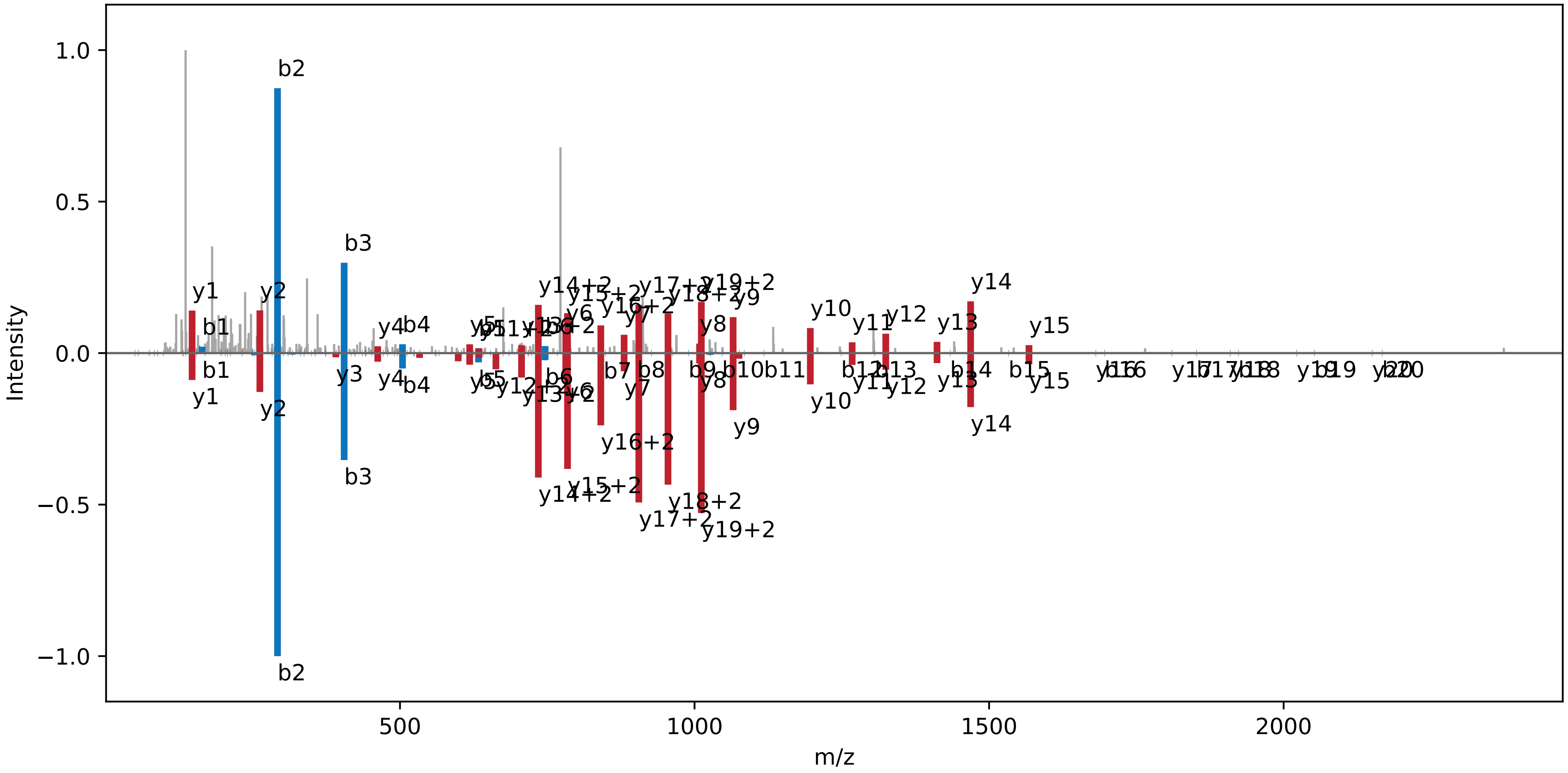

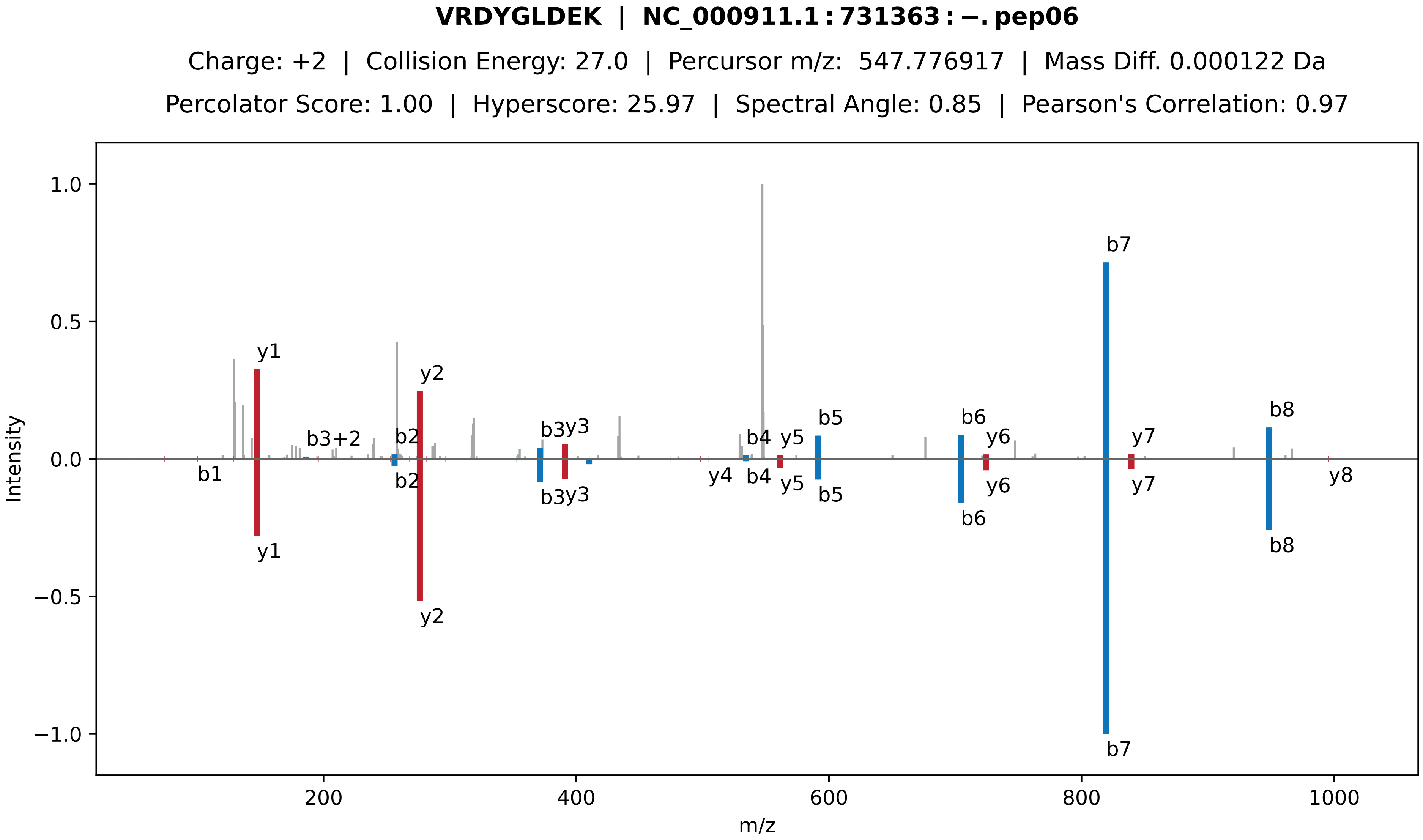

VRDYGLDEK | NC_000911.1 : 731363 : −. pep06
Charge: +2 | Collision Energy: 27.0 | Percursor m/z: 547.776917 | Mass Diff. 0.000122 Da
Percolator Score: 1.00 | Hyperscore: 25.97 | Spectral Angle: 0.85 | Pearson's Correlation: 0.97
Intensity
m/z
1.0
0.5
0.0
−0.5
−1.0
200
400
600
800
1000
y1
b3+2
b2
y2
b3y3
b4
y5
b5
b6
y6
b7
y7
b8
b1
y4
y8

**EATISALLGEK | NC_000911.1 : 731363 : −. pep07**

Charge: +2 | Collision Energy: 27.0 | Percursor m/z: 566.316101 | Mass Diff. -0.000366 Da

Percolator Score: 1.00 | Hyperscore: 27.25 | Spectral Angle: 0.94 | Pearson's Correlation: 0.99

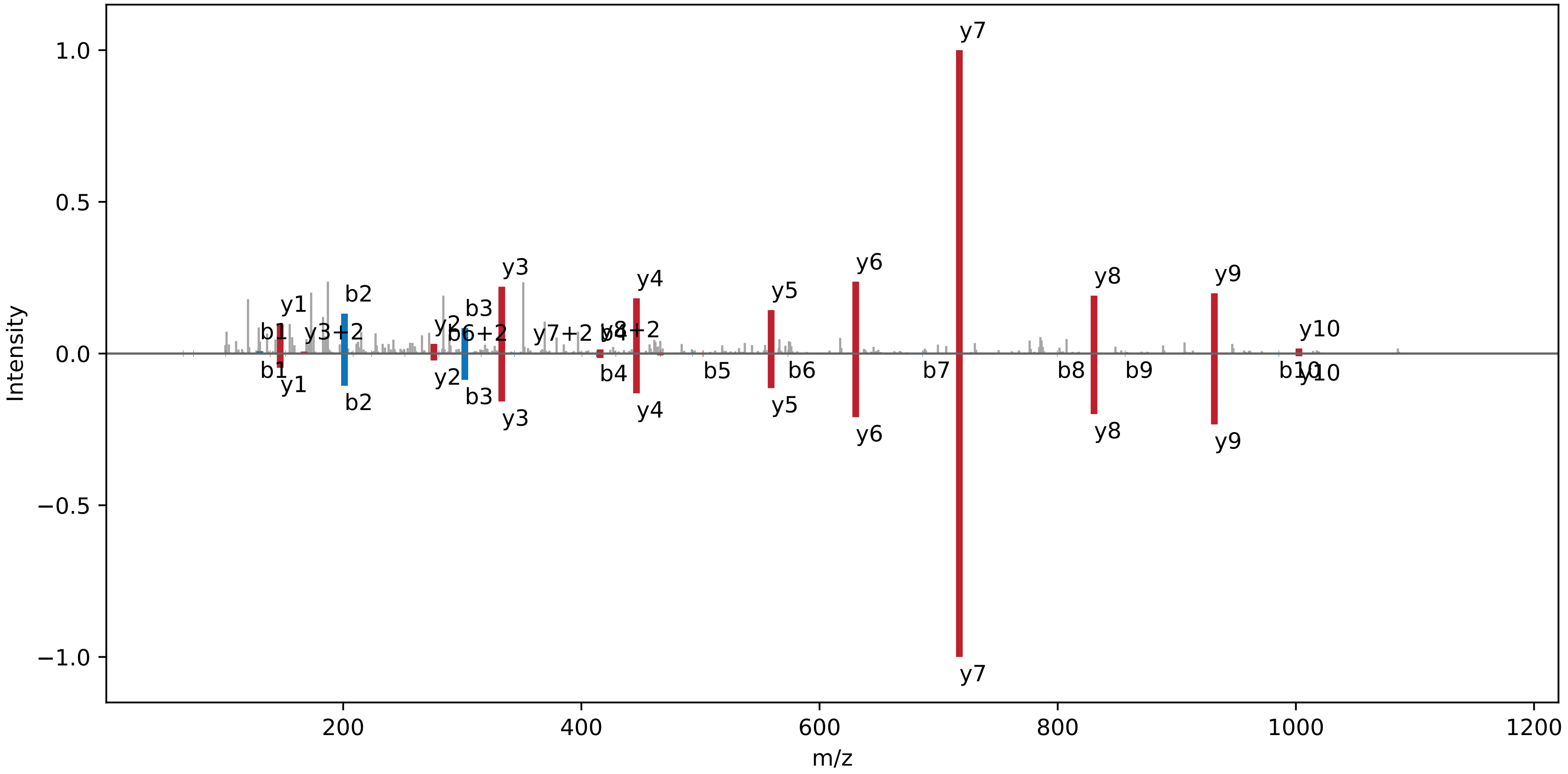

### YQLVELVGSGAMGQVYR | NC_000911.1 : 731363 : −. pep08

Charge: +2 | Collision Energy: 27.0 | Percursor m/z: 935.480000 | Mass Diff. 0.000244 Da

Percolator Score: 1.00 | Hyperscore: 45.32 | Spectral Angle: 0.86 | Pearson's Correlation: 0.97

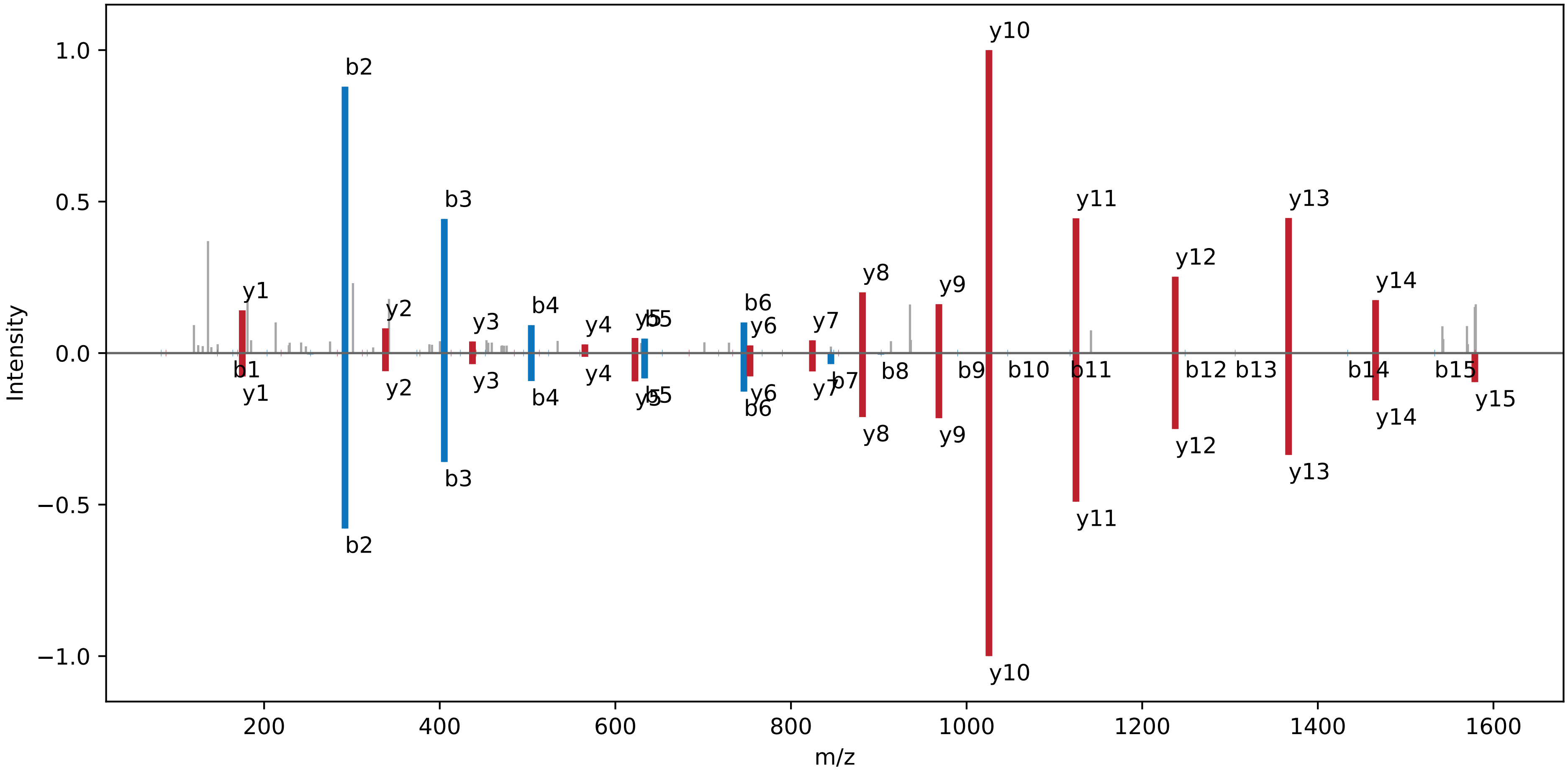

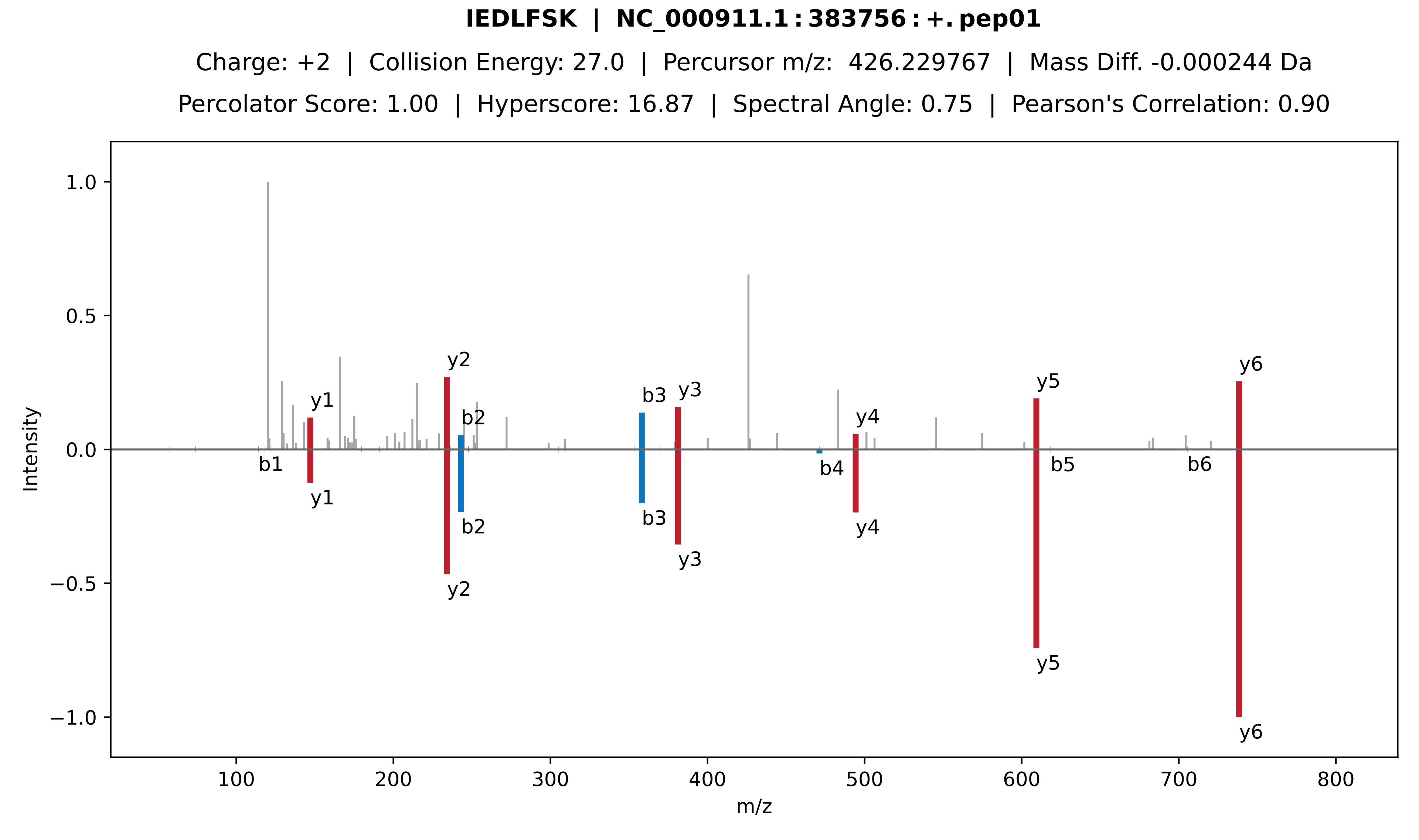
IEDLFSK | NC_000911.1:383756:+.pep01
Charge: +2 | Collision Energy: 27.0 | Percursor m/z: 426.229767 | Mass Diff. -0.000244 Da
Percolator Score: 1.00 | Hyperscore: 16.87 | Spectral Angle: 0.75 | Pearson's Correlation: 0.90
Intensity
m/z
1.0
0.5
0.0
−0.5
−1.0
100
200
300
400
500
600
700
800
y1
y2
b2
b3
y3
y4
y5
y6
b1
b4
b5
b6

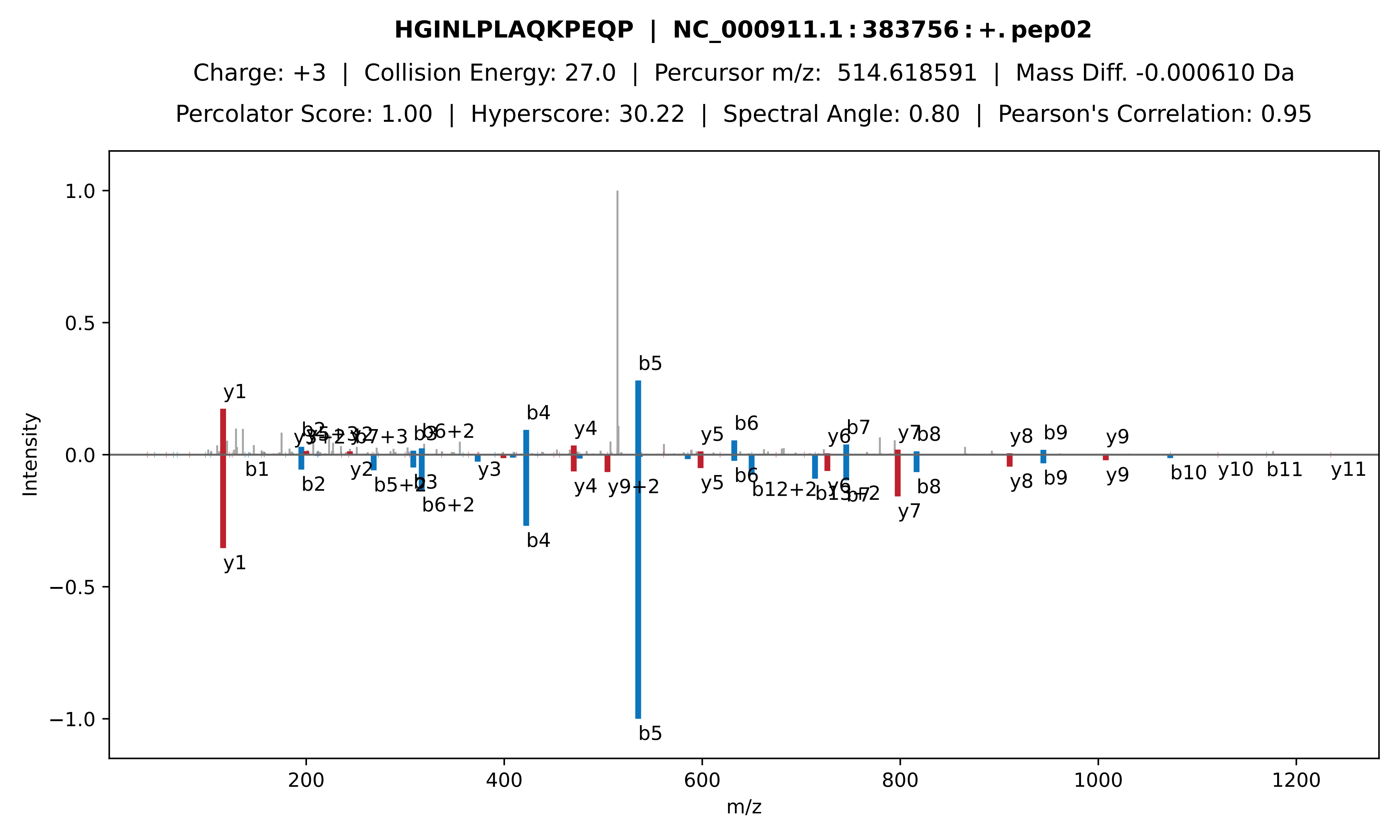
HGINLPLAQKPEQP | NC_000911.1:383756:+.pep02
Charge: +3 | Collision Energy: 27.0 | Percursor m/z: 514.618591 | Mass Diff. -0.000610 Da
Percolator Score: 1.00 | Hyperscore: 30.22 | Spectral Angle: 0.80 | Pearson's Correlation: 0.95
Intensity
m/z
1.0
0.5
0.0
−0.5
−1.0
200
400
600
800
1000
1200
y1
b5
b4
y4
b6
y5
b7
y6
y7
b8
y8
b9
y9
b1
b2
y2
b5+2
b3
b6+2
y3
y9+2
b12+2
b10
y10
b11
y11

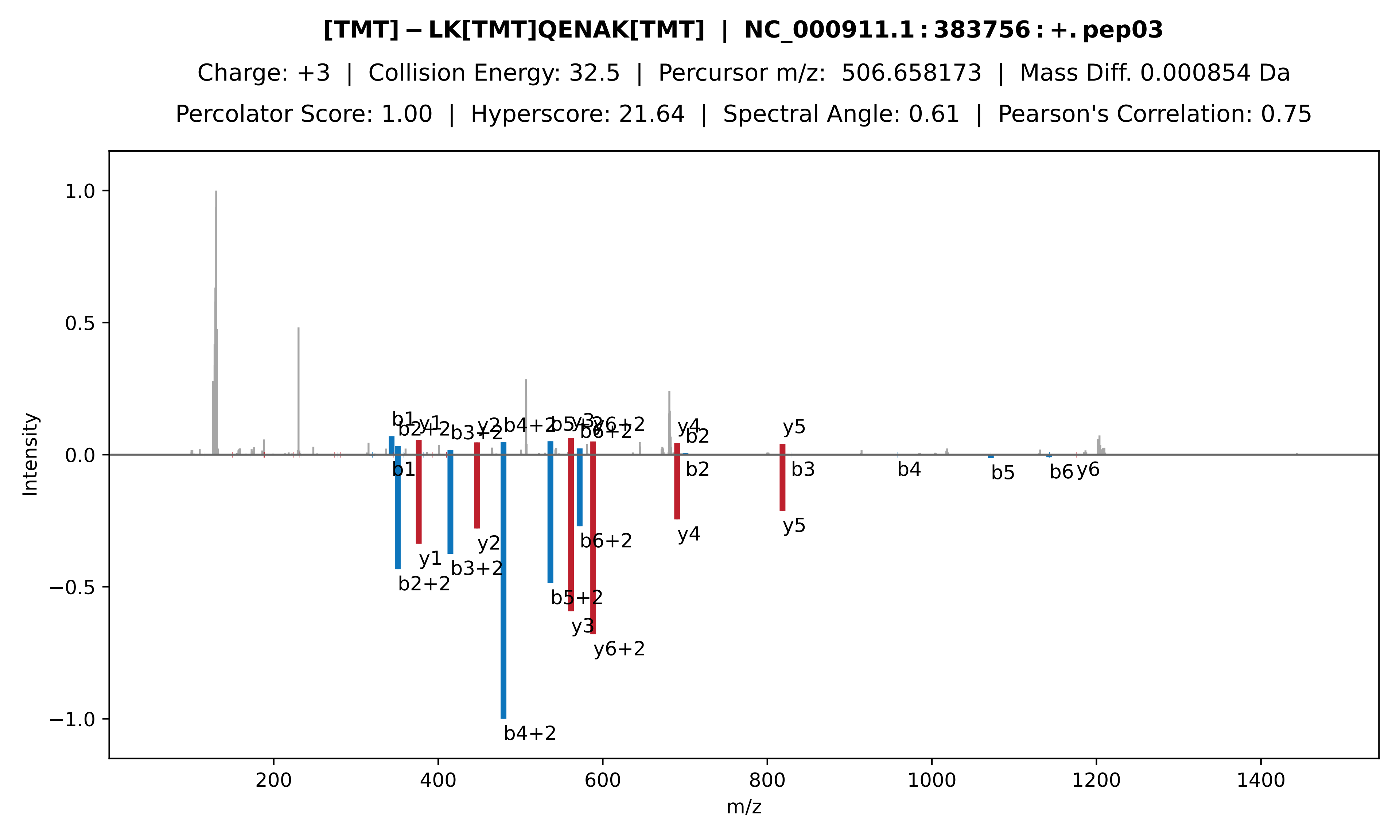

[TMT]-LK[TMT]QENAK[TMT] | NC_000911.1:383756:+.pep03
Charge: +3 | Collision Energy: 32.5 | Percursor m/z: 506.658173 | Mass Diff. 0.000854 Da
Percolator Score: 1.00 | Hyperscore: 21.64 | Spectral Angle: 0.61 | Pearson's Correlation: 0.75
Intensity
m/z
1.0
0.5
0.0
−0.5
−1.0
200
400
600
800
1000
1200
1400
b1
y1
b2+2
b3+2
y2
b4+2
b5+2
y3
y6+2
b6+2
y4
b2
y5
b3
b4
b5
b6
y6

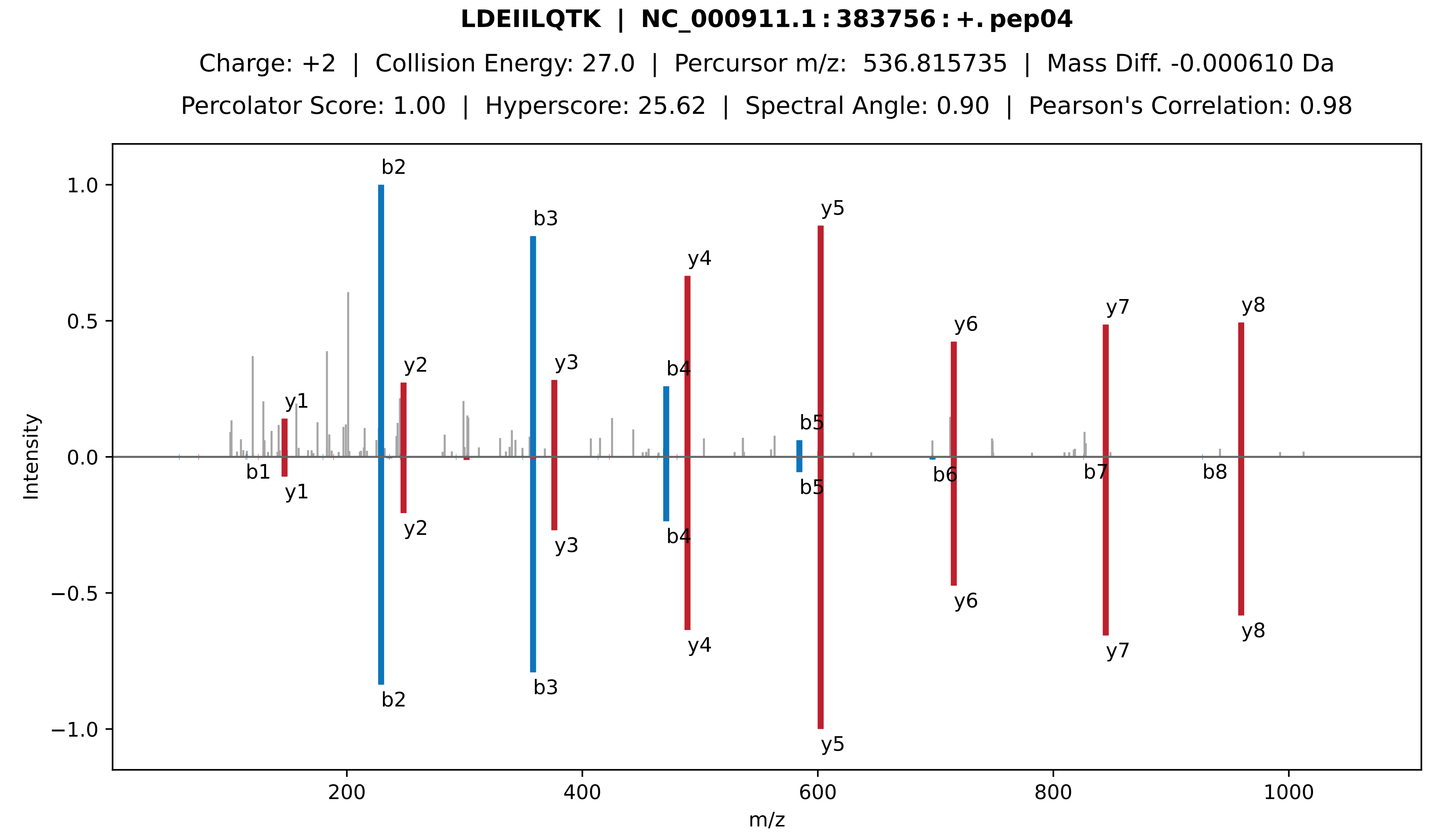

LDEIILQTK | NC_000911.1:383756:+.pep04
Charge: +2 | Collision Energy: 27.0 | Percursor m/z: 536.815735 | Mass Diff. -0.000610 Da
Percolator Score: 1.00 | Hyperscore: 25.62 | Spectral Angle: 0.90 | Pearson's Correlation: 0.98
Intensity
m/z
b1
b2
b3
b4
b5
b6
b7
b8
y1
y2
y3
y4
y5
y6
y7
y8

**LAVLESKIEDLFSK | NC_000911.1 : 383756 : +. pep05**

Charge: +2 | Collision Energy: 27.0 | Percursor m/z: 796.451000 | Mass Diff. 0.000000 Da

Percolator Score: 1.00 | Hyperscore: 42.93 | Spectral Angle: 0.75 | Pearson's Correlation: 0.90

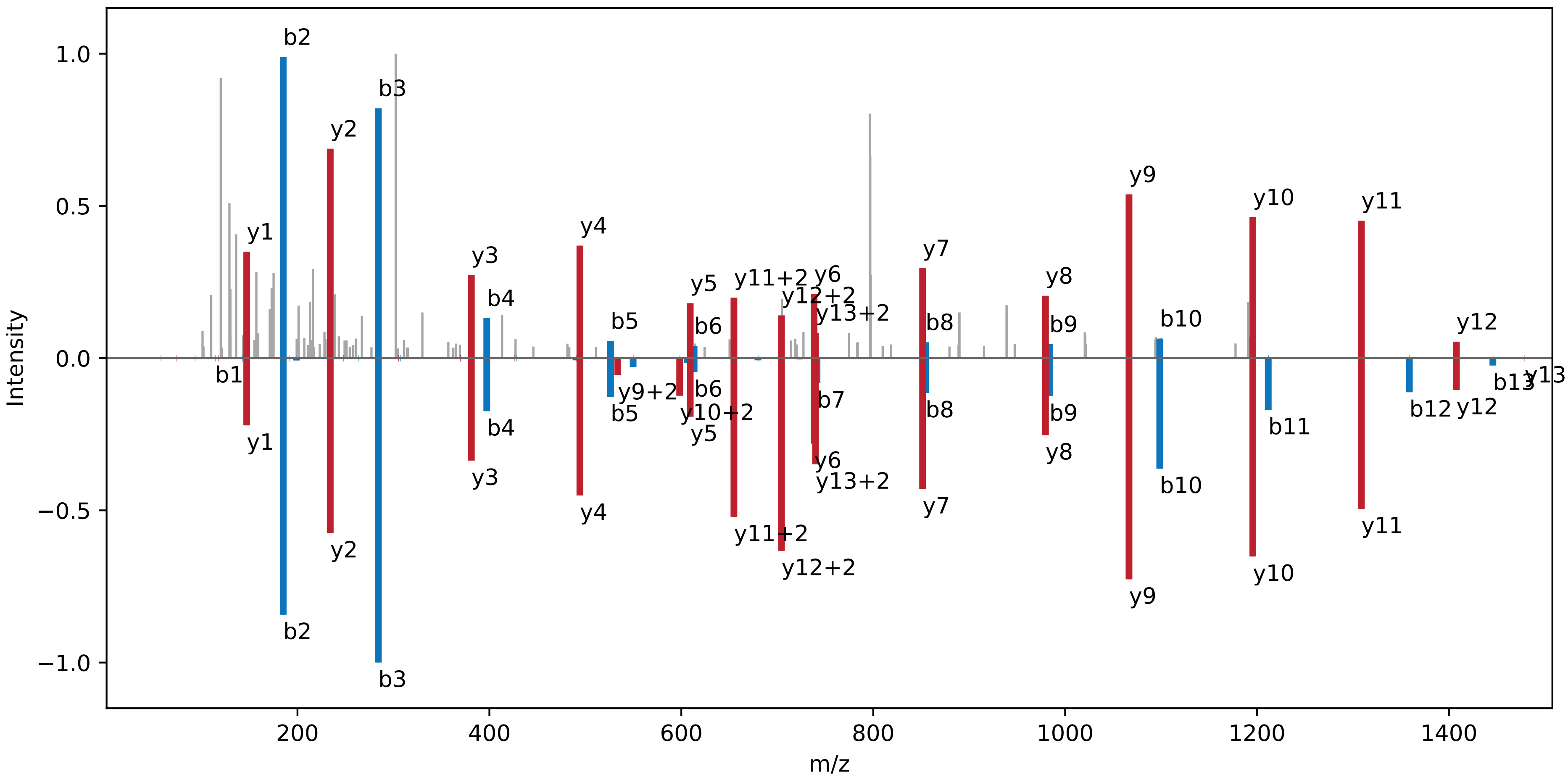

**ATSTPPPSPESPSPSLDGASWR | NC_000911.1 : 383756 : +. pep06**

Charge: +2 | Collision Energy: 35.0 | Percursor m/z: 1112.526400 | Mass Diff. -0.000732 Da

Percolator Score: 1.00 | Hyperscore: 43.51 | Spectral Angle: 0.13 | Pearson's Correlation: 0.17

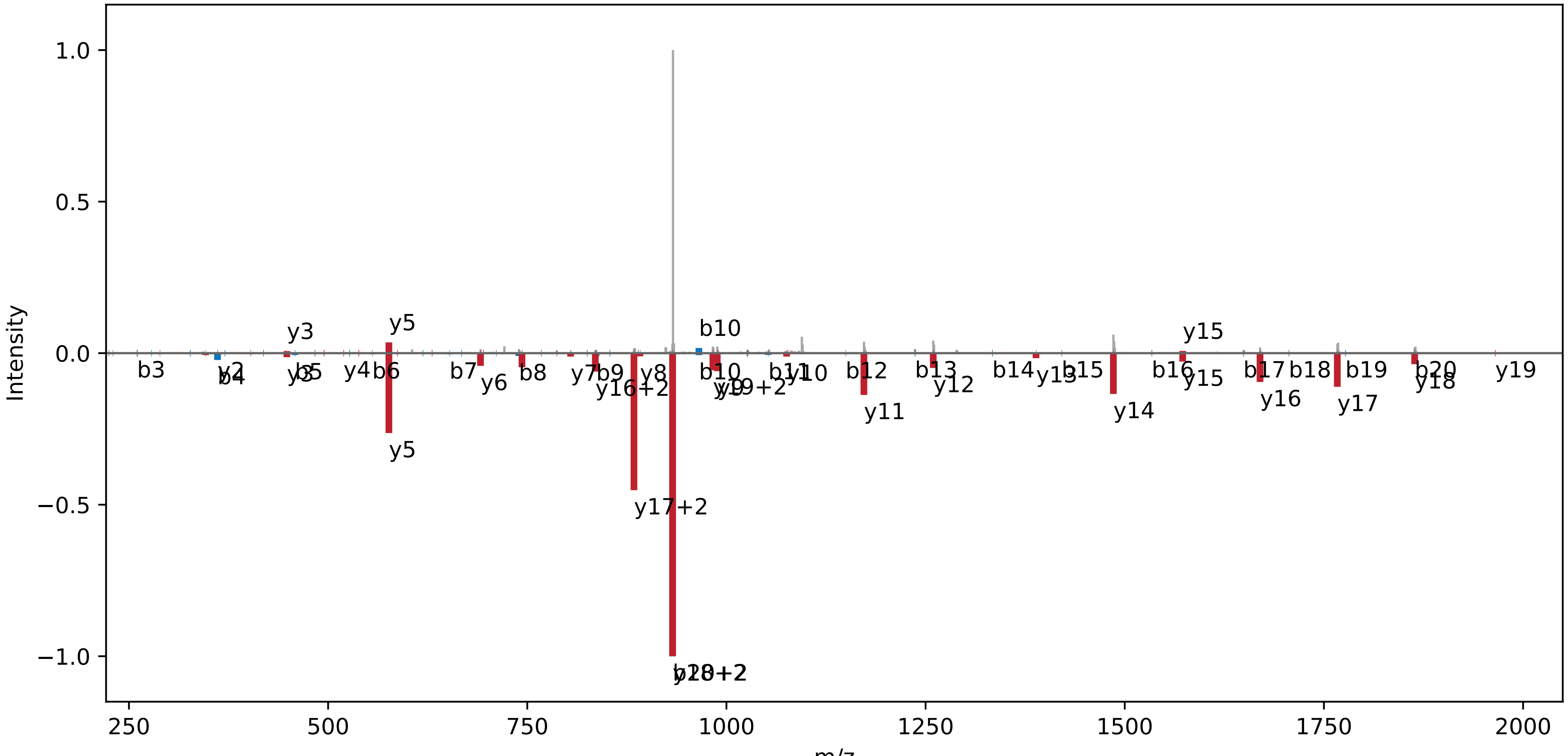

**[TMT]-EIEQIK[TMT]K[TMT] | NC_000911.1:383756:+.pep07**

Charge: +3 | Collision Energy: 32.5 | Percursor m/z: 525.674927 | Mass Diff. 0.002197 Da

Percolator Score: 1.00 | Hyperscore: 23.76 | Spectral Angle: 0.35 | Pearson's Correlation: 0.38

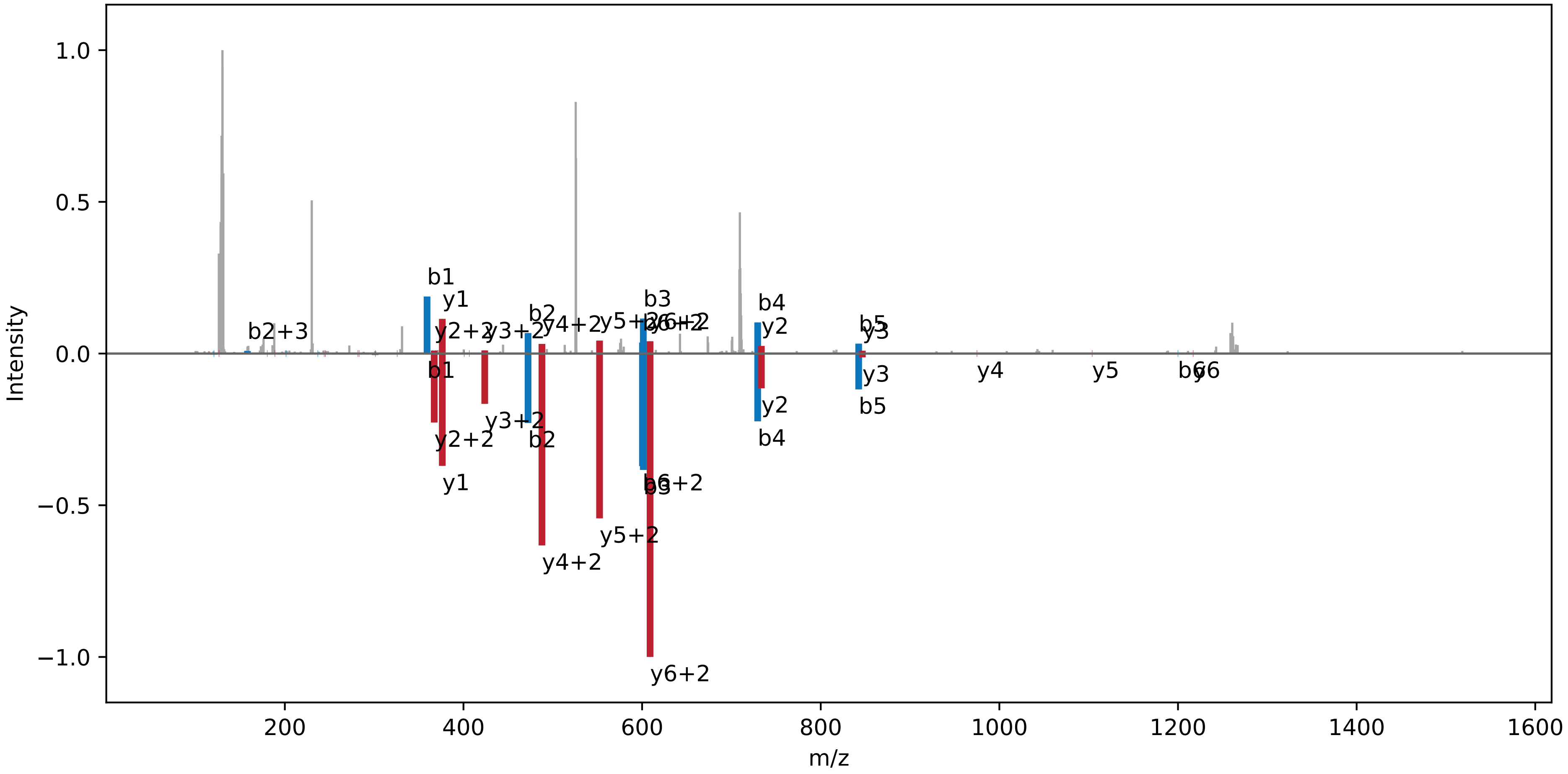

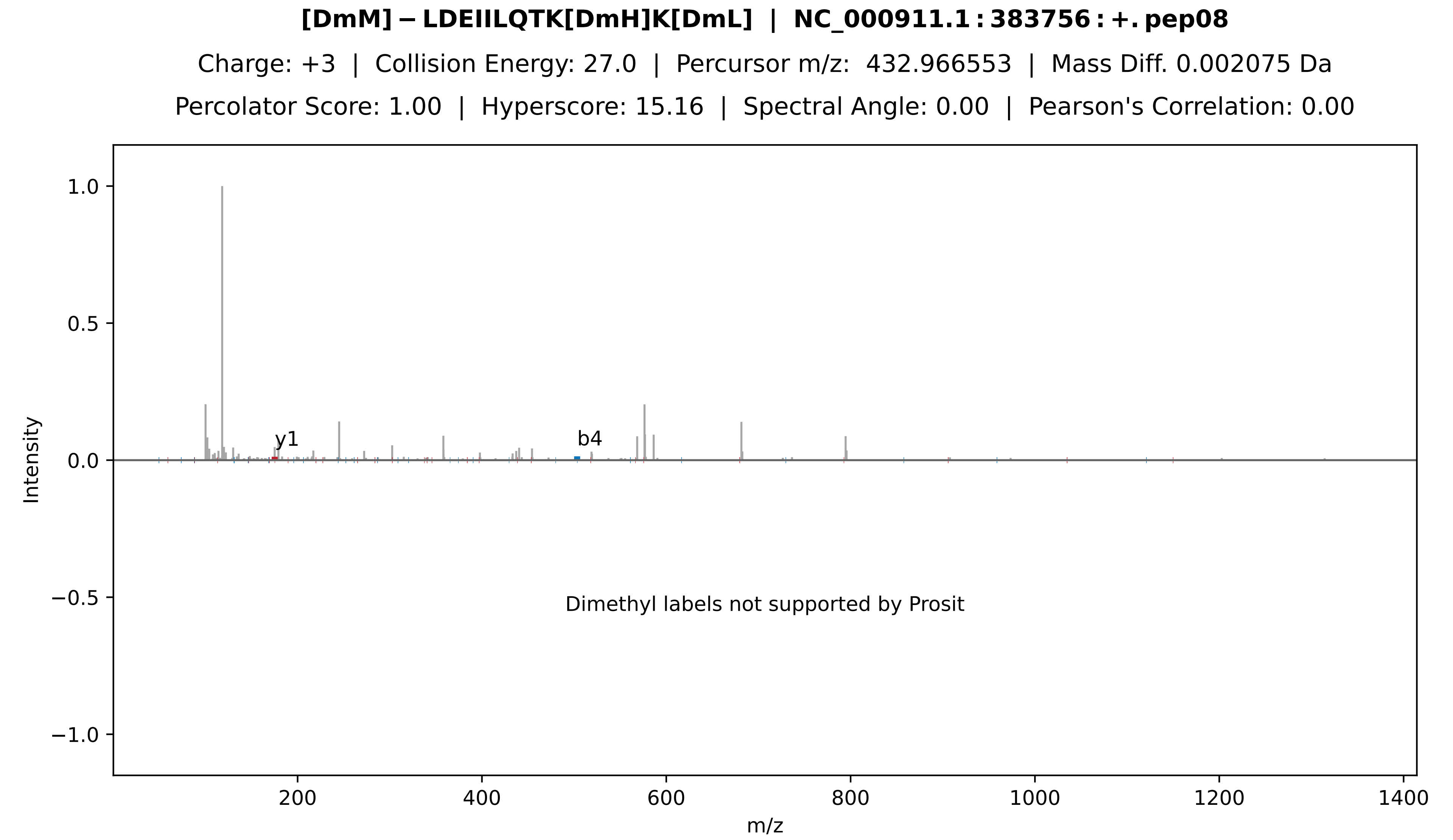
[DmM]−LDEIILQTK[DmH]K[DmL] | NC_000911.1:383756:+.pep08
Charge: +3 | Collision Energy: 27.0 | Percursor m/z: 432.966553 | Mass Diff. 0.002075 Da
Percolator Score: 1.00 | Hyperscore: 15.16 | Spectral Angle: 0.00 | Pearson's Correlation: 0.00
1.0
0.5
0.0
−0.5
−1.0
Intensity
y1
b4
Dimethyl labels not supported by Prosit
200
400
600
800
1000
1200
1400
m/z

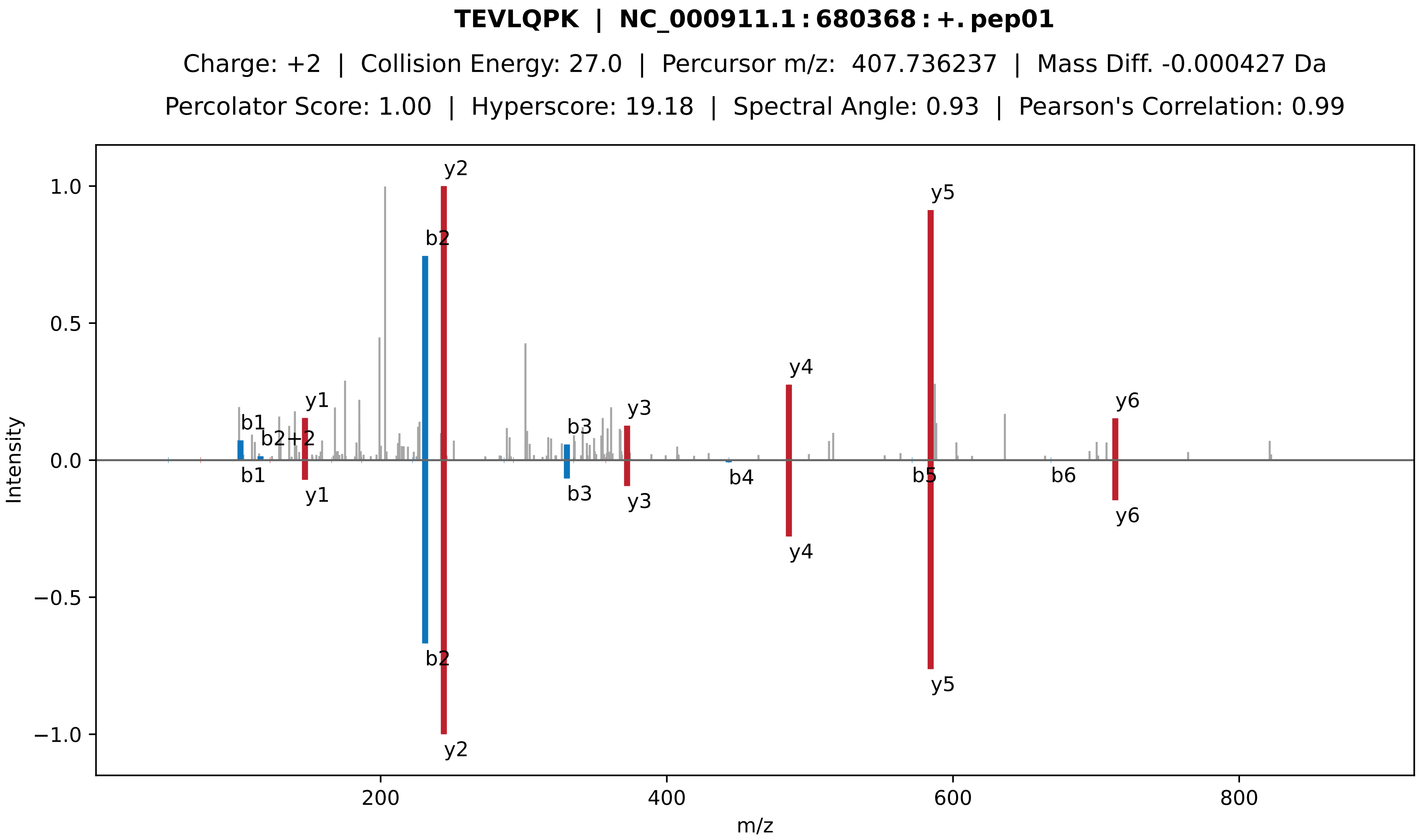

TEVLQPK | NC_000911.1 : 680368 : +. pep01
Charge: +2 | Collision Energy: 27.0 | Percursor m/z: 407.736237 | Mass Diff. -0.000427 Da
Percolator Score: 1.00 | Hyperscore: 19.18 | Spectral Angle: 0.93 | Pearson's Correlation: 0.99
Intensity
m/z
1.0
0.5
0.0
−0.5
−1.0
200
400
600
800
y2
b2
y5
y4
y1
b1
b2+2
b3
y3
y6
b1
y1
b3
y3
b4
b5
b6
y6
y4
b2
y5
y2

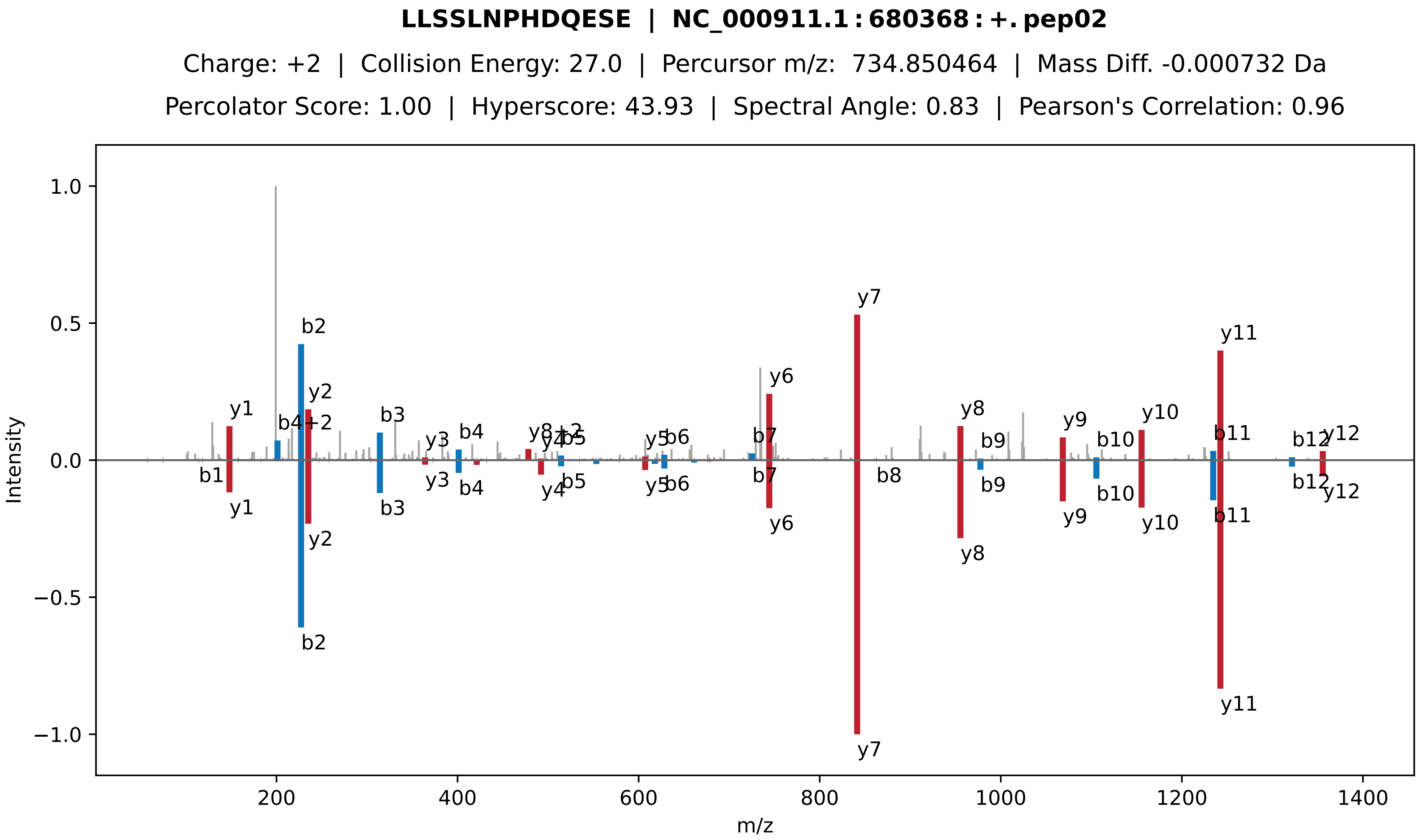
LLSSLNPHDQESE | NC_000911.1 : 680368 : +. pep02
Charge: +2 | Collision Energy: 27.0 | Percursor m/z: 734.850464 | Mass Diff. -0.000732 Da
Percolator Score: 1.00 | Hyperscore: 43.93 | Spectral Angle: 0.83 | Pearson's Correlation: 0.96
Intensity
m/z
y1
b2
y2
b4+2
b3
y3
b4
y8+2
y4
b5
y5
b6
b7
y6
y7
b8
y8
b9
y9
b10
y10
b11
y11
b12
y12
b1

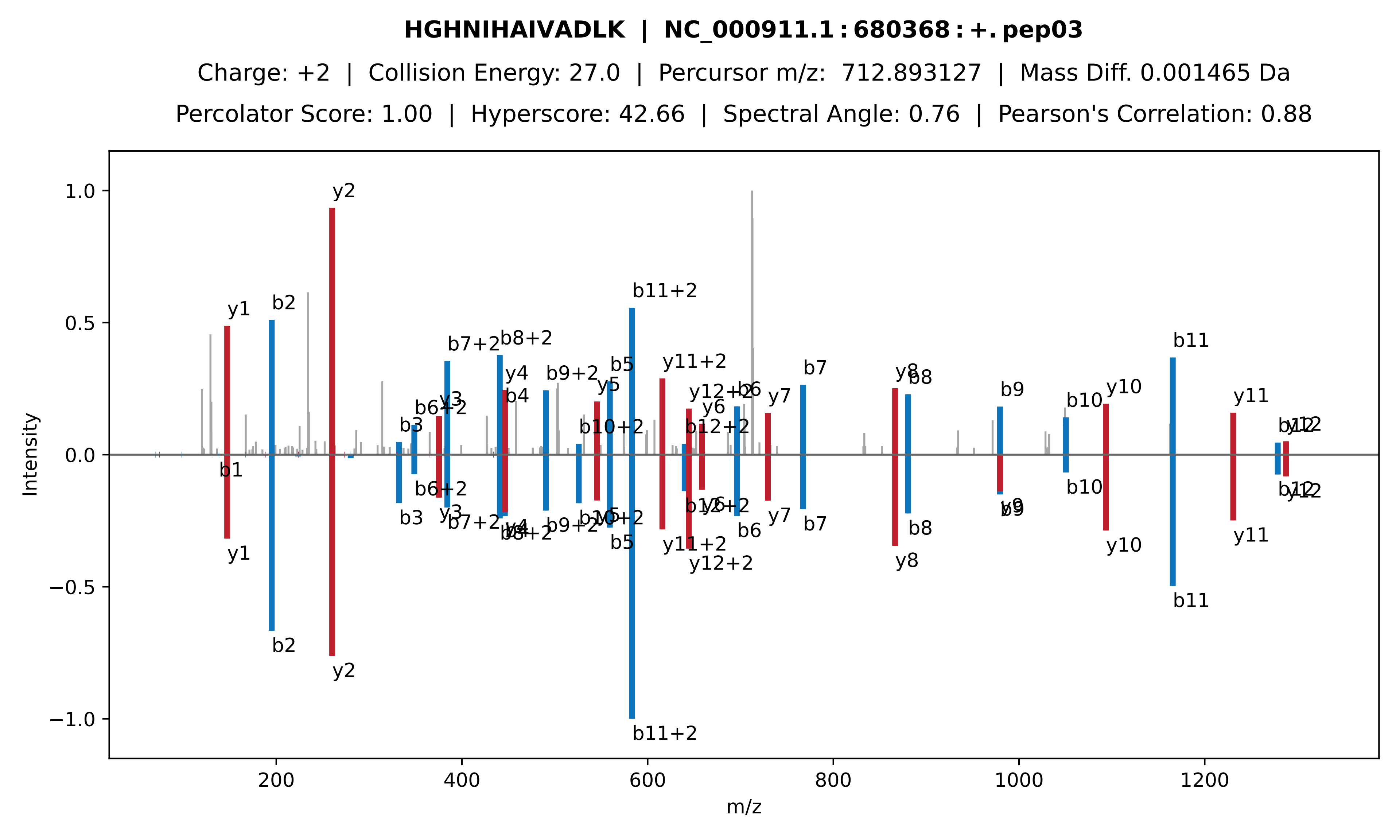
HGHNIHAIVADLK | NC_000911.1 : 680368 : +. pep03
Charge: +2 | Collision Energy: 27.0 | Percursor m/z: 712.893127 | Mass Diff. 0.001465 Da
Percolator Score: 1.00 | Hyperscore: 42.66 | Spectral Angle: 0.76 | Pearson's Correlation: 0.88
Intensity
m/z
1.0
0.5
0.0
−0.5
−1.0
200
400
600
800
1000
1200
y1
b2
y2
b1
b3
b6+2
y3
b7+2
b8+2
y4
b4
b9+2
b5
y5
b10+2
b11+2
y11+2
y12+2
b6
y6
b12+2
y7
b7
y8
b8
b9
y9
b10
y10
b11
y11
b12
y12

**[DmH] − RLRDEYAAK[DmH] | NC_000911.1 : 680368 : +. pep04**

Charge: +3 | Collision Energy: 27.0 | Percursor m/z: 398.589935 | Mass Diff. -0.000732 Da

Percolator Score: 1.00 | Hyperscore: 18.25 | Spectral Angle: 0.00 | Pearson's Correlation: 0.00

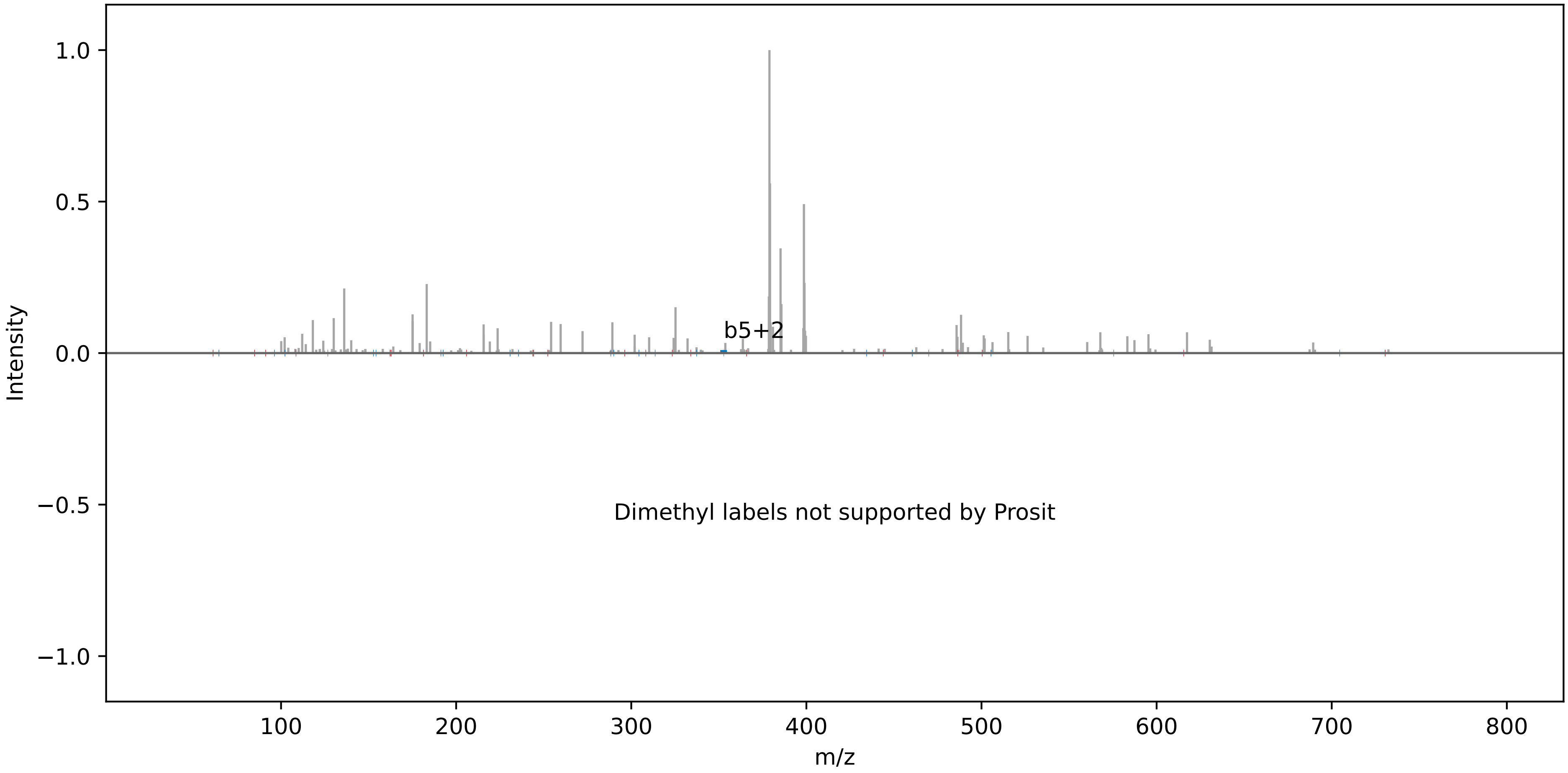

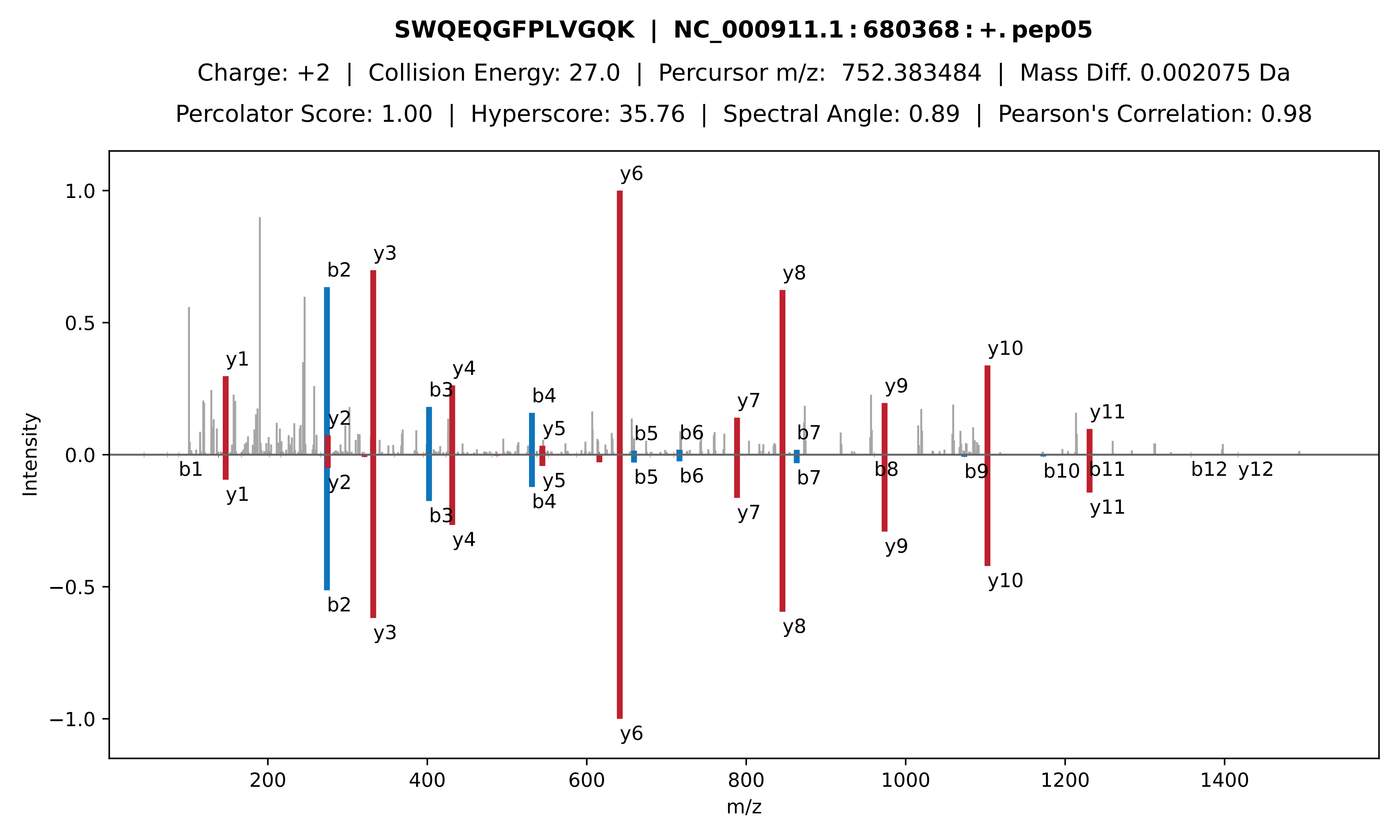

SWQEQGFPLVGQK | NC_000911.1:680368:+.pep05
Charge: +2 | Collision Energy: 27.0 | Percursor m/z: 752.383484 | Mass Diff. 0.002075 Da
Percolator Score: 1.00 | Hyperscore: 35.76 | Spectral Angle: 0.89 | Pearson's Correlation: 0.98
Intensity
m/z
y1 y2 y3 y4 y5 y6 y7 y8 y9 y10 y11 y12
b1 b2 b3 b4 b5 b6 b7 b8 b9 b10 b11 b12

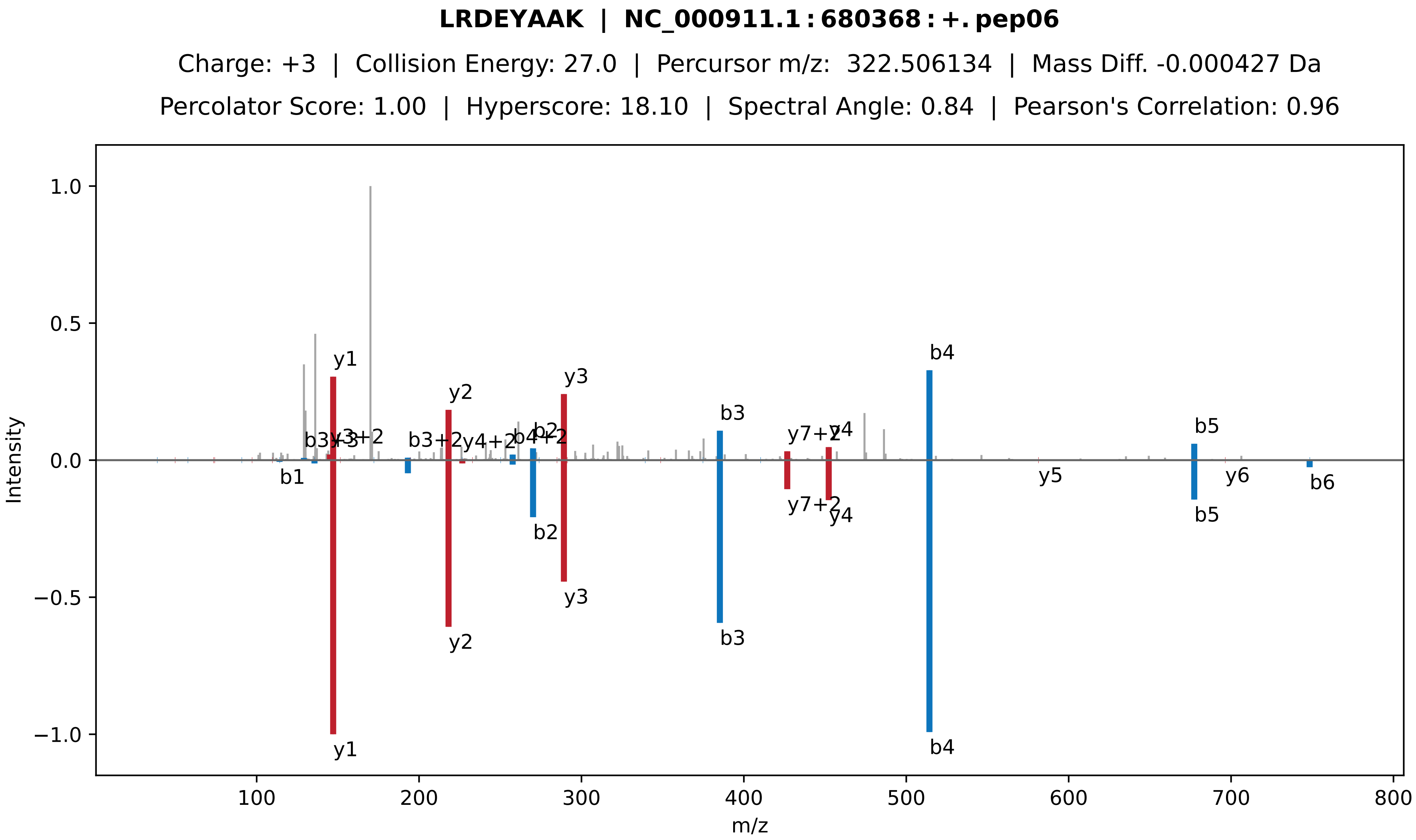

**LRDEYAAK | NC_000911.1 : 680368 : +. pep06**
Charge: +3 | Collision Energy: 27.0 | Percursor m/z: 322.506134 | Mass Diff. -0.000427 Da
Percolator Score: 1.00 | Hyperscore: 18.10 | Spectral Angle: 0.84 | Pearson's Correlation: 0.96
Intensity
m/z
y1
y2
y3
b3
b4
b5
b6
b1
b2
y5
y6
y4
y7+2
b3+2
y4+2
b4+2
y3+2
b3+3
y3+3

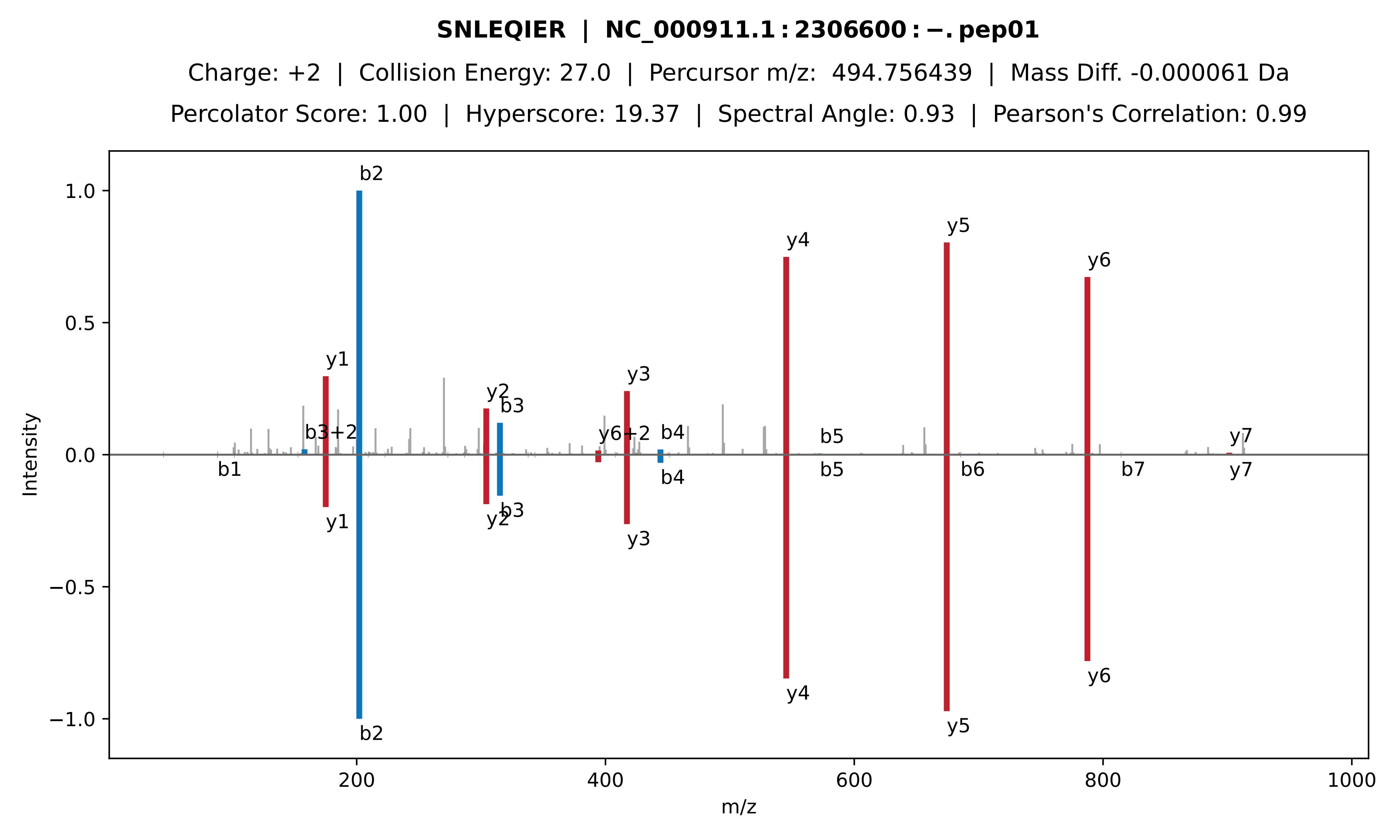

**SNLEQIER | NC_000911.1 : 2306600 : −. pep01**
Charge: +2 | Collision Energy: 27.0 | Percursor m/z: 494.756439 | Mass Diff. -0.000061 Da
Percolator Score: 1.00 | Hyperscore: 19.37 | Spectral Angle: 0.93 | Pearson's Correlation: 0.99
Intensity
m/z
b1
b2
b3
b3+2
b4
b5
b6
b7
y1
y2
y3
y4
y5
y6
y6+2
y7

**QLSPSDFAK | NC_000911.1 : 2306600 : −. pep02**

Charge: +2 | Collision Energy: 27.0 | Percursor m/z: 496.756073 | Mass Diff. 0.000854 Da

Percolator Score: 1.00 | Hyperscore: 24.74 | Spectral Angle: 0.88 | Pearson's Correlation: 0.98

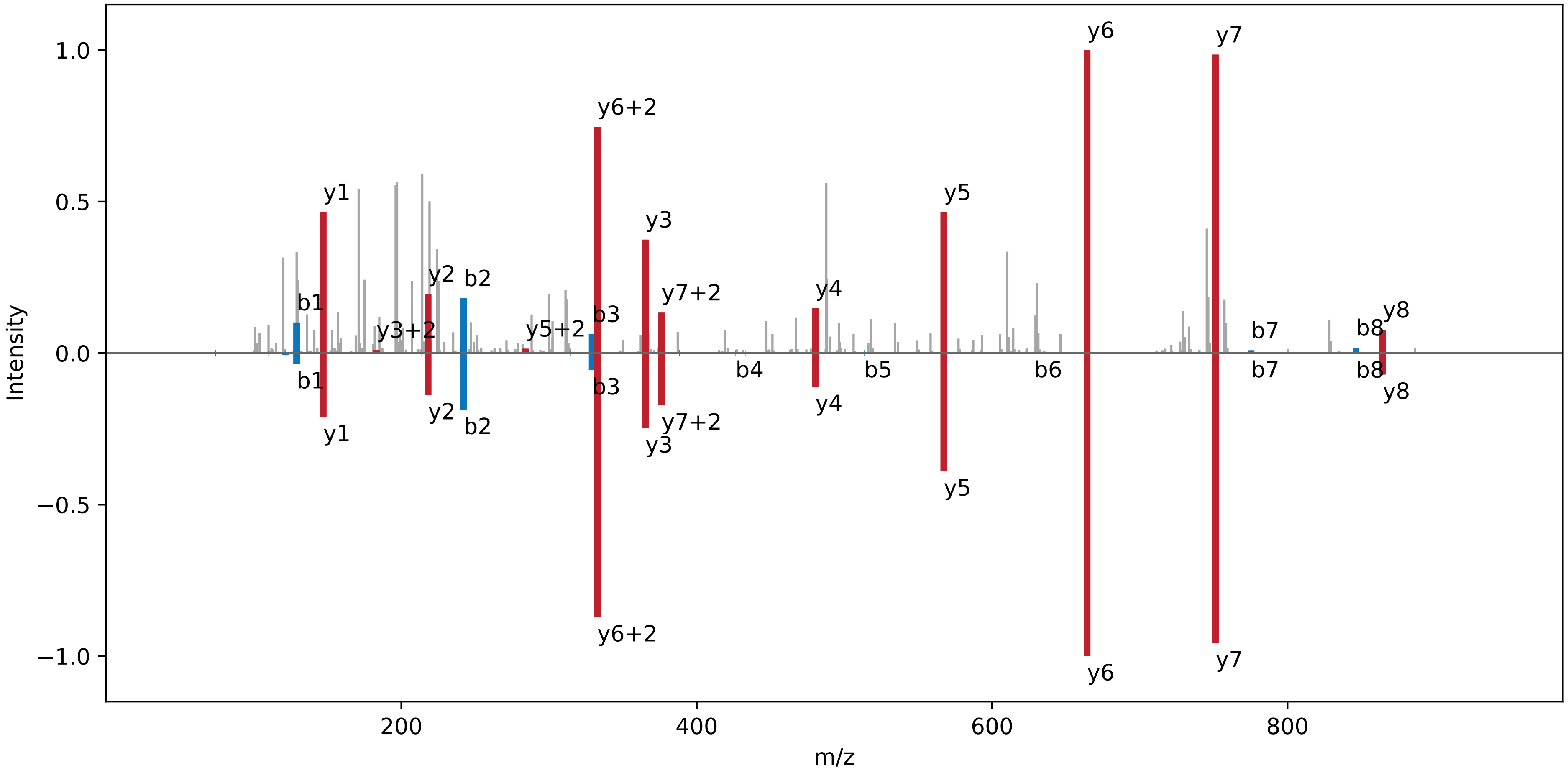

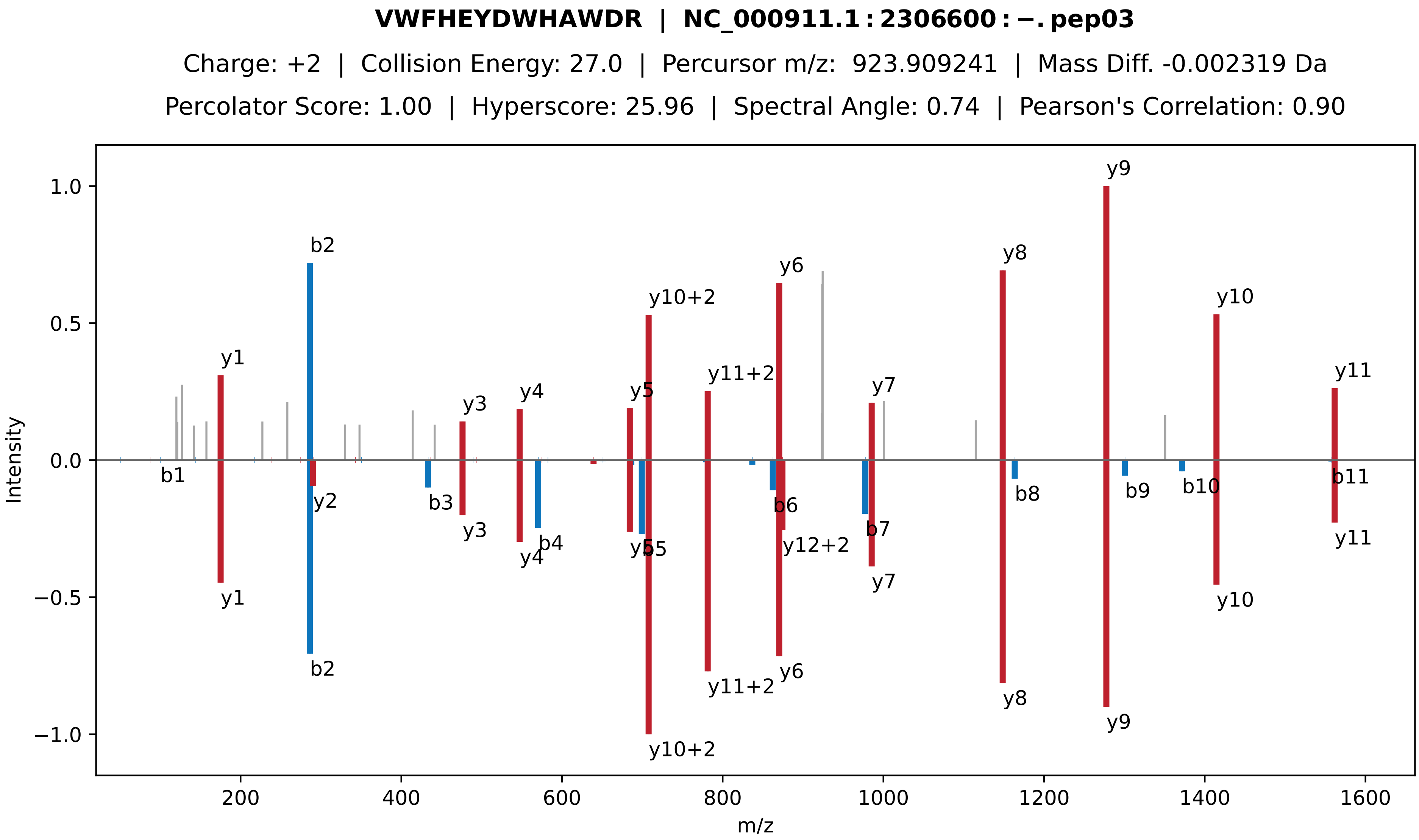

**VWFHEYDWHAWDR | NC_000911.1 : 2306600 : −. pep03**
Charge: +2 | Collision Energy: 27.0 | Percursor m/z: 923.909241 | Mass Diff. -0.002319 Da
Percolator Score: 1.00 | Hyperscore: 25.96 | Spectral Angle: 0.74 | Pearson's Correlation: 0.90
Intensity
m/z
y1
b2
y3
y4
y5
y10+2
y11+2
y6
y7
y8
y9
y10
y11
b1
y2
b3
b4
b5
b6
y12+2
b7
b8
b9
b10
b11

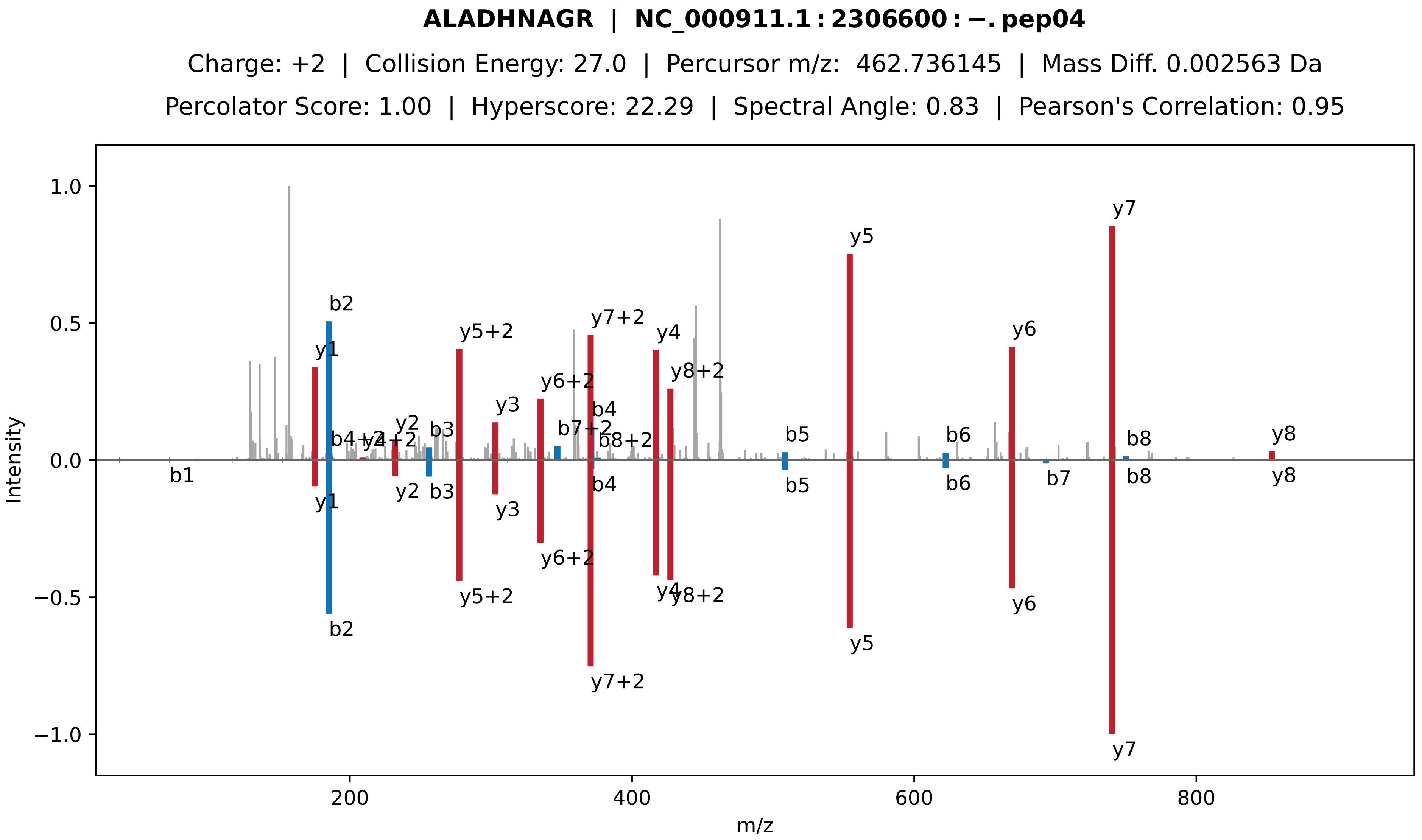
ALADHNAGR | NC_000911.1 : 2306600 : −. pep04
Charge: +2 | Collision Energy: 27.0 | Percursor m/z: 462.736145 | Mass Diff. 0.002563 Da
Percolator Score: 1.00 | Hyperscore: 22.29 | Spectral Angle: 0.83 | Pearson's Correlation: 0.95
Intensity
m/z
b1
b2
b3
b4
b5
b6
b7
b8
y1
y2
y3
y4
y5
y6
y7
y8
y5+2
y6+2
y7+2
y8+2
b7+2
b8+2
b4+2
y4+2

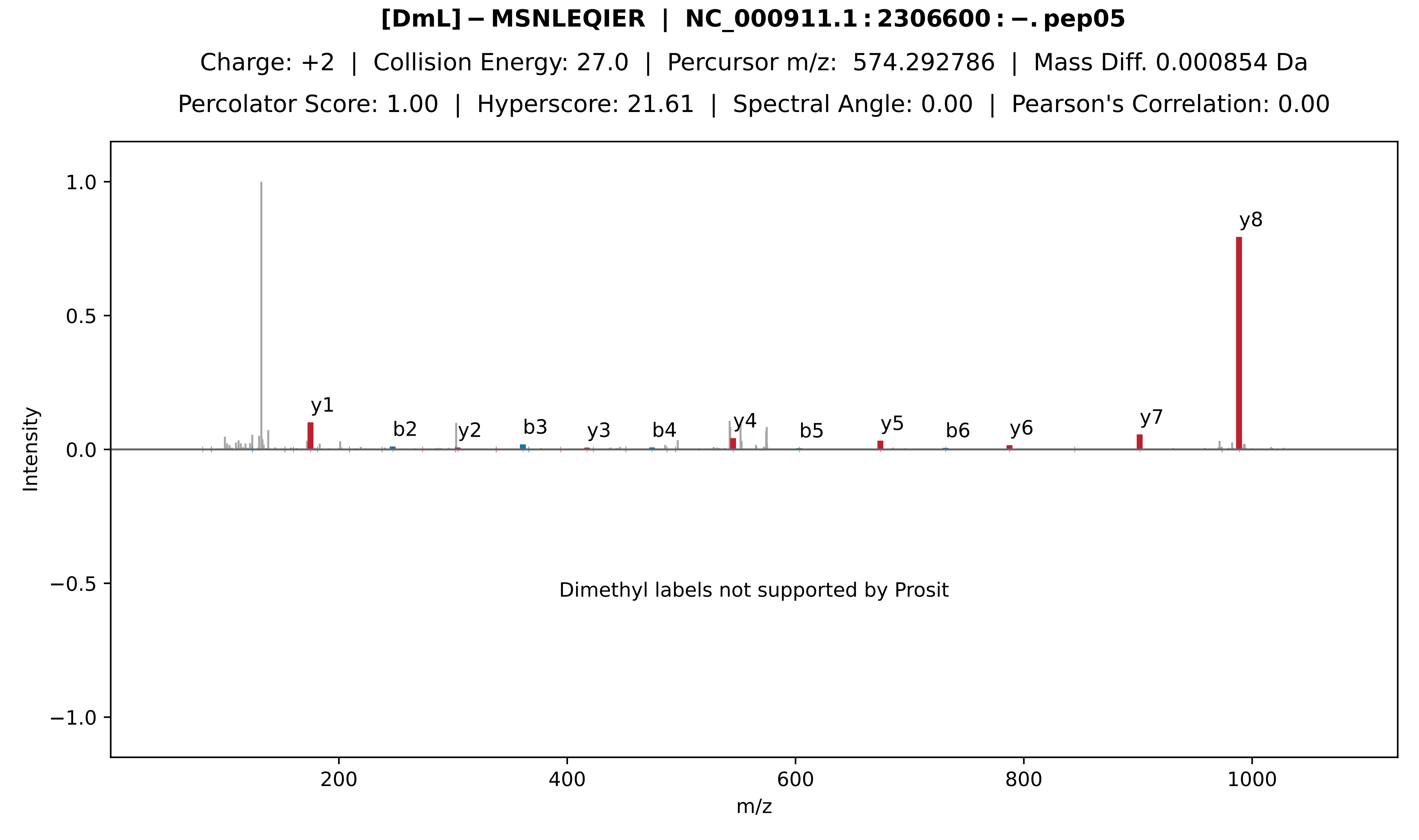
[DmL]−MSNLEQIER | NC_000911.1:2306600:−.pep05
Charge: +2 | Collision Energy: 27.0 | Percursor m/z: 574.292786 | Mass Diff. 0.000854 Da
Percolator Score: 1.00 | Hyperscore: 21.61 | Spectral Angle: 0.00 | Pearson's Correlation: 0.00
y8
y1
b2
y2
b3
y3
b4
y4
b5
y5
b6
y6
y7
Dimethyl labels not supported by Prosit
Intensity
m/z
1.0
0.5
0.0
−0.5
−1.0
200
400
600
800
1000

**QIEQDSK | NC_000911.1 : 2306600 : −. pep06**

Charge: +2 | Collision Energy: 27.0 | Percursor m/z: 424.212128 | Mass Diff. -0.000122 Da

Percolator Score: 0.54 | Hyperscore: 11.01 | Spectral Angle: 0.22 | Pearson's Correlation: 0.19

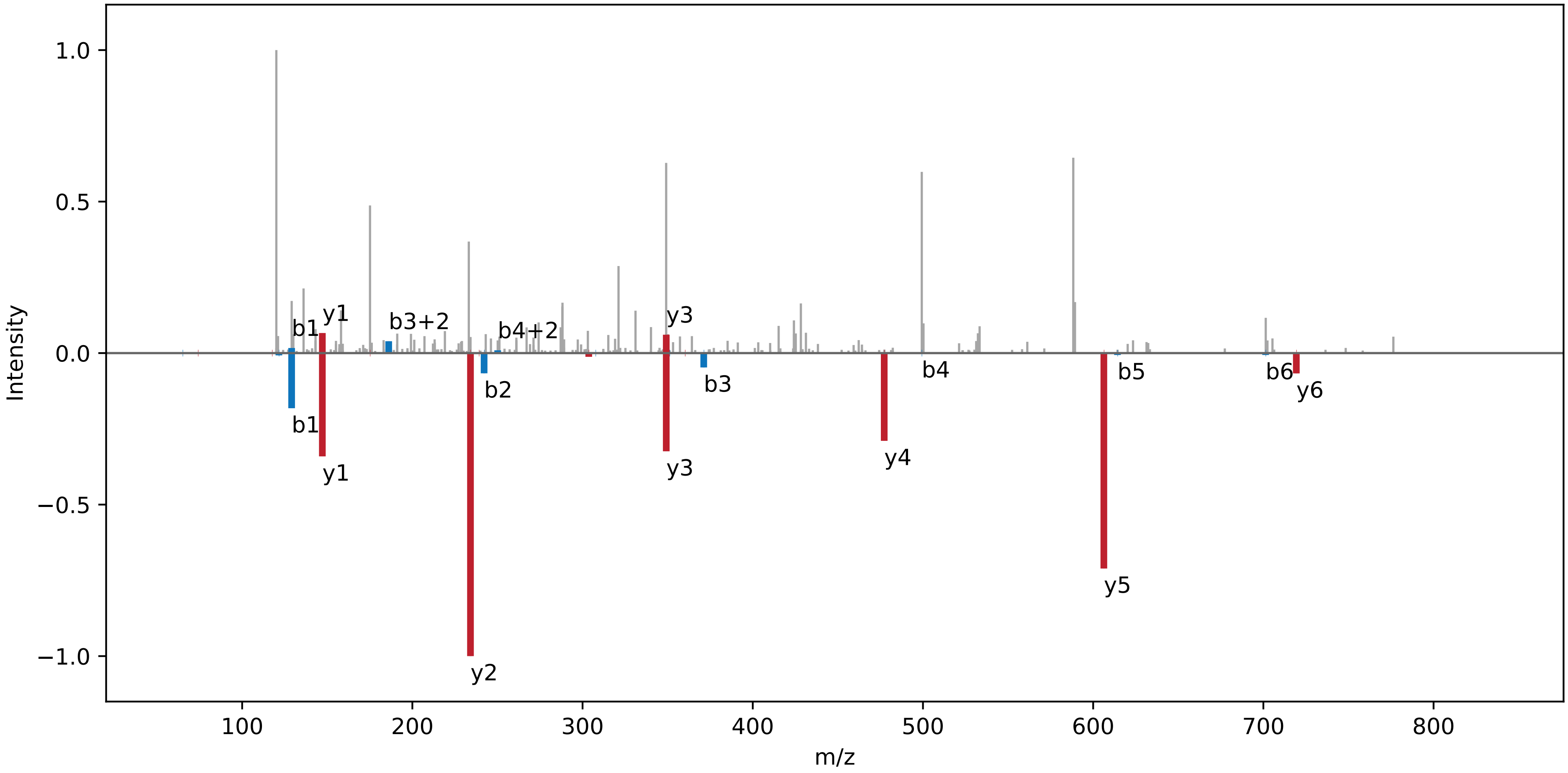

**[TMT]-LK[TMT]QENEK[TMT] | NC_000911.1:2622940:+.pep01**

Charge: +3 | Collision Energy: 32.5 | Percursor m/z: 525.993286 | Mass Diff. -0.000610 Da

Percolator Score: 1.00 | Hyperscore: 23.06 | Spectral Angle: 0.61 | Pearson's Correlation: 0.75

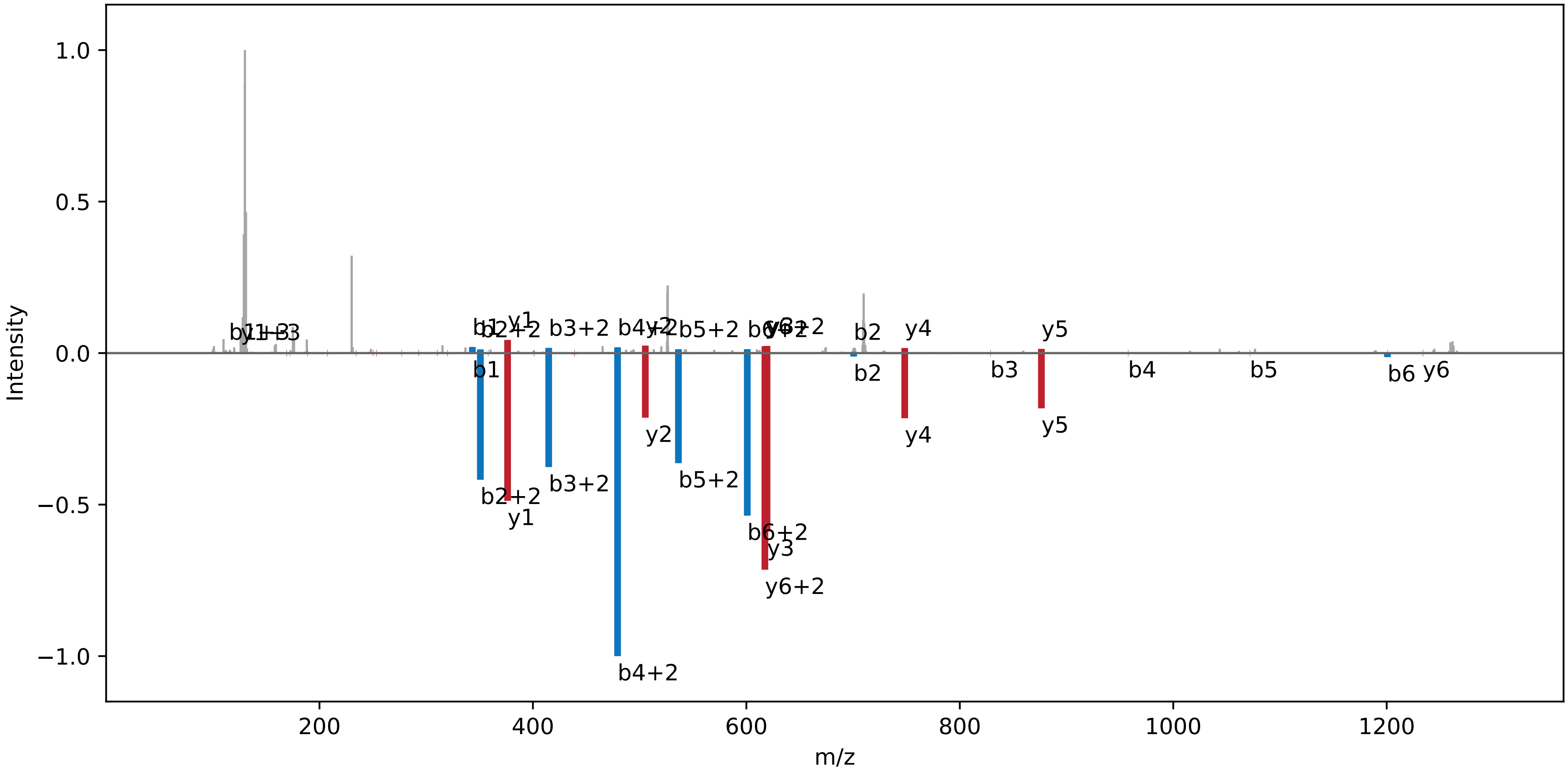

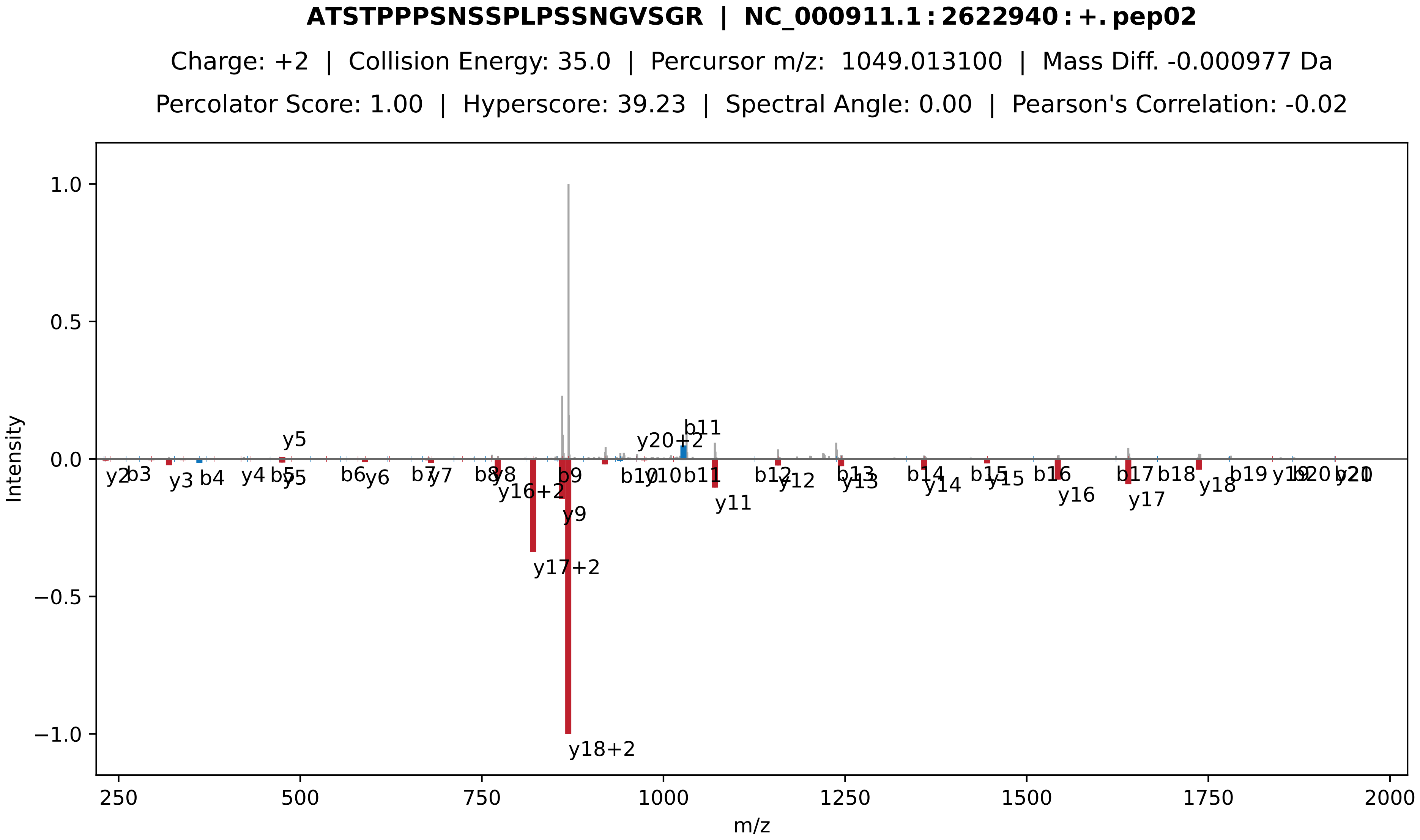
ATSTPPPSNSSPLPSSNGVSGR | NC_000911.1:2622940:+.pep02
Charge: +2 | Collision Energy: 35.0 | Percursor m/z: 1049.013100 | Mass Diff. -0.000977 Da
Percolator Score: 1.00 | Hyperscore: 39.23 | Spectral Angle: 0.00 | Pearson's Correlation: -0.02
Intensity
m/z
y5
y20+2
b11
y2
b3
y3
b4
y4
b5
y5
b6
y6
b7
y7
b8
y8
y16+2
b9
y9
y17+2
y18+2
b10
y10
b11
y11
b12
y12
b13
y13
b14
y14
b15
y15
b16
y16
b17
y17
b18
y18
b19
y19
b20
b21
y21

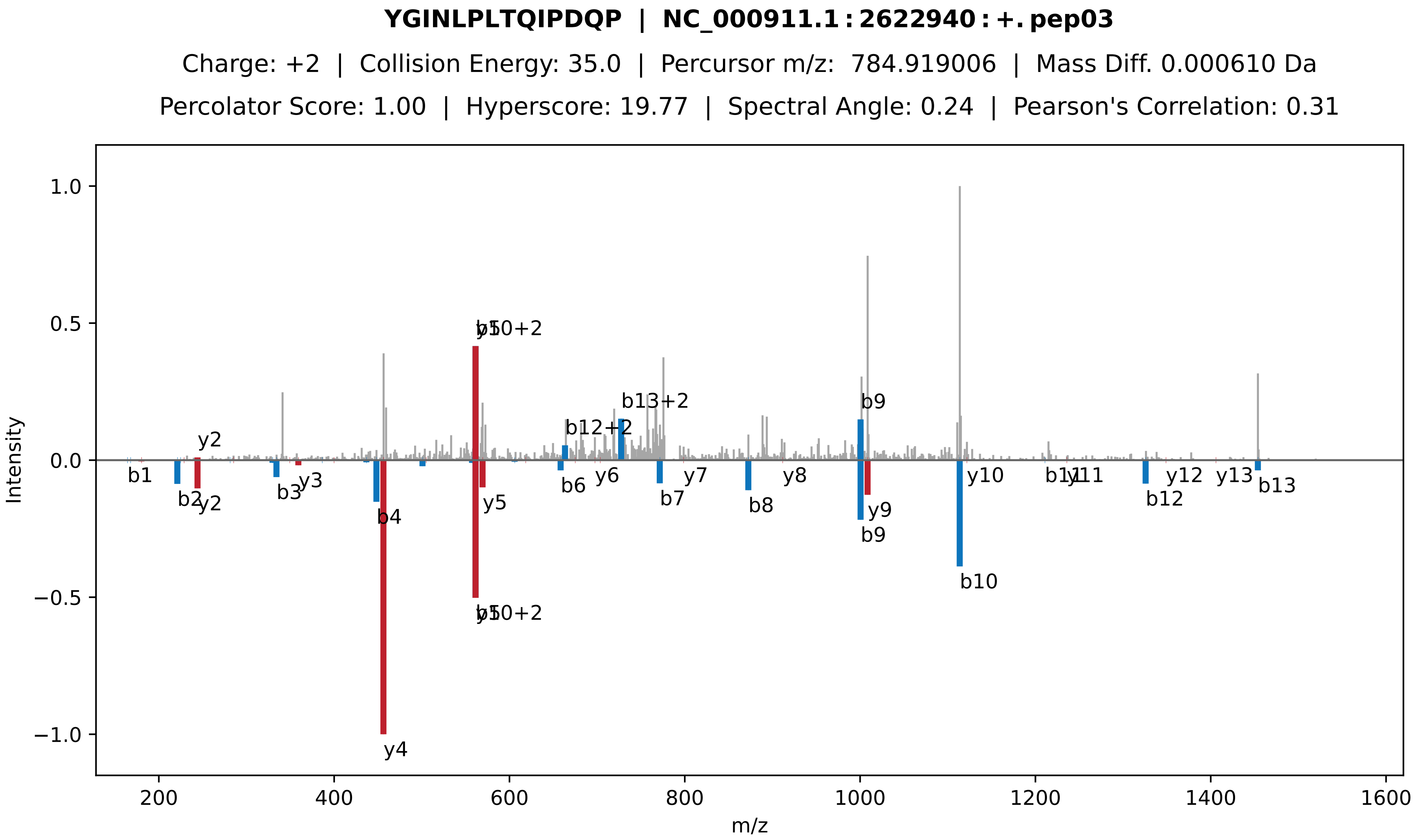
YGINLPLTQIPDQP | NC_000911.1:2622940:+.pep03
Charge: +2 | Collision Energy: 35.0 | Percursor m/z: 784.919006 | Mass Diff. 0.000610 Da
Percolator Score: 1.00 | Hyperscore: 19.77 | Spectral Angle: 0.24 | Pearson's Correlation: 0.31
Intensity
m/z
1.0
0.5
0.0
−0.5
−1.0
200
400
600
800
1000
1200
1400
1600
y2
b1
b2
b3
y3
b4
y4
y5
b10+2
b6
y6
b12+2
b13+2
b7
y7
b8
y8
b9
y9
y10
b10
b11
y11
b12
y12
y13
b13

**LDEM[Ox]ILQTR | NC_000911.1 : 2622940 : +. pep04**

Charge: +2 | Collision Energy: 28.0 | Percursor m/z: 567.797546 | Mass Diff. 0.001465 Da

Percolator Score: 1.00 | Hyperscore: 22.60 | Spectral Angle: 0.84 | Pearson's Correlation: 0.96

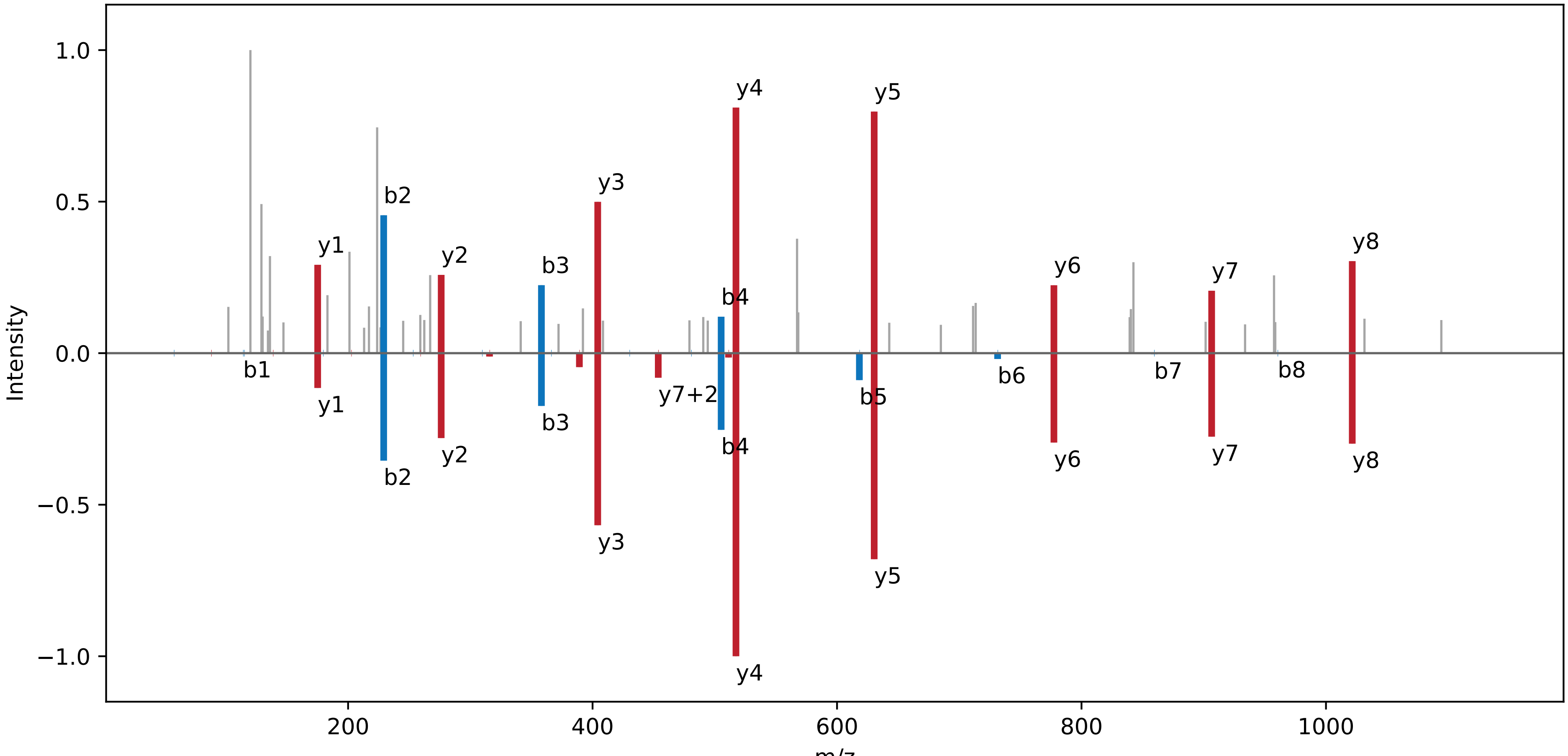

**LKQENEKLDEMILQTR | NC_000911.1 : 2622940 : +. pep05**

Charge: +3 | Collision Energy: 40.0 | Percursor m/z: 663.352800 | Mass Diff. 0.000488 Da

Percolator Score: 1.00 | Hyperscore: 23.67 | Spectral Angle: 0.51 | Pearson's Correlation: 0.67

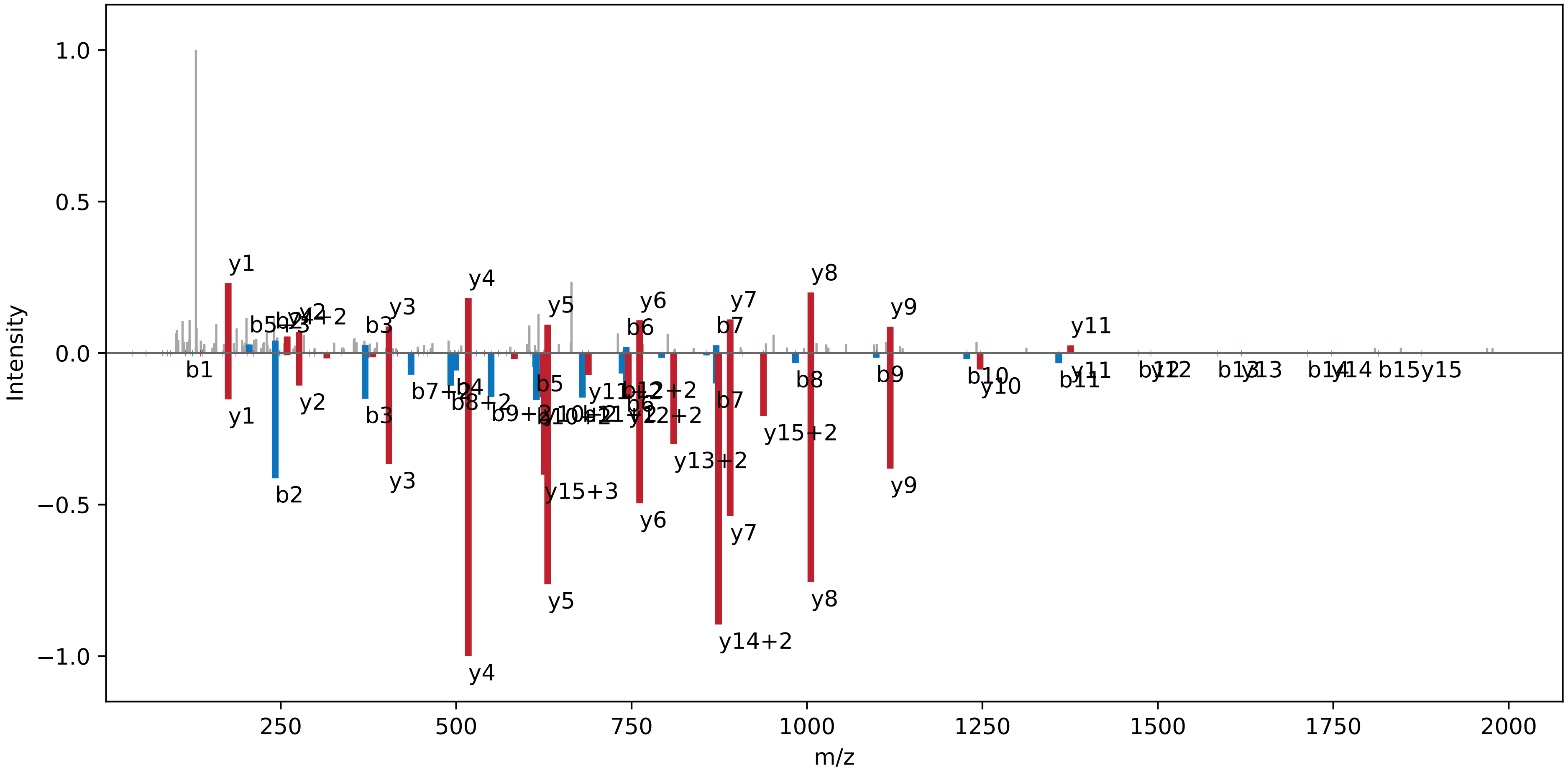

**QGSSPSSSDSEPILCPHCR | NC_000911.1 : 2905488 : +. pep01**

Charge: +3 | Collision Energy: 27.0 | Percursor m/z: 700.975830 | Mass Diff. 0.000000 Da

Percolator Score: 1.00 | Hyperscore: 21.06 | Spectral Angle: 0.75 | Pearson's Correlation: 0.92

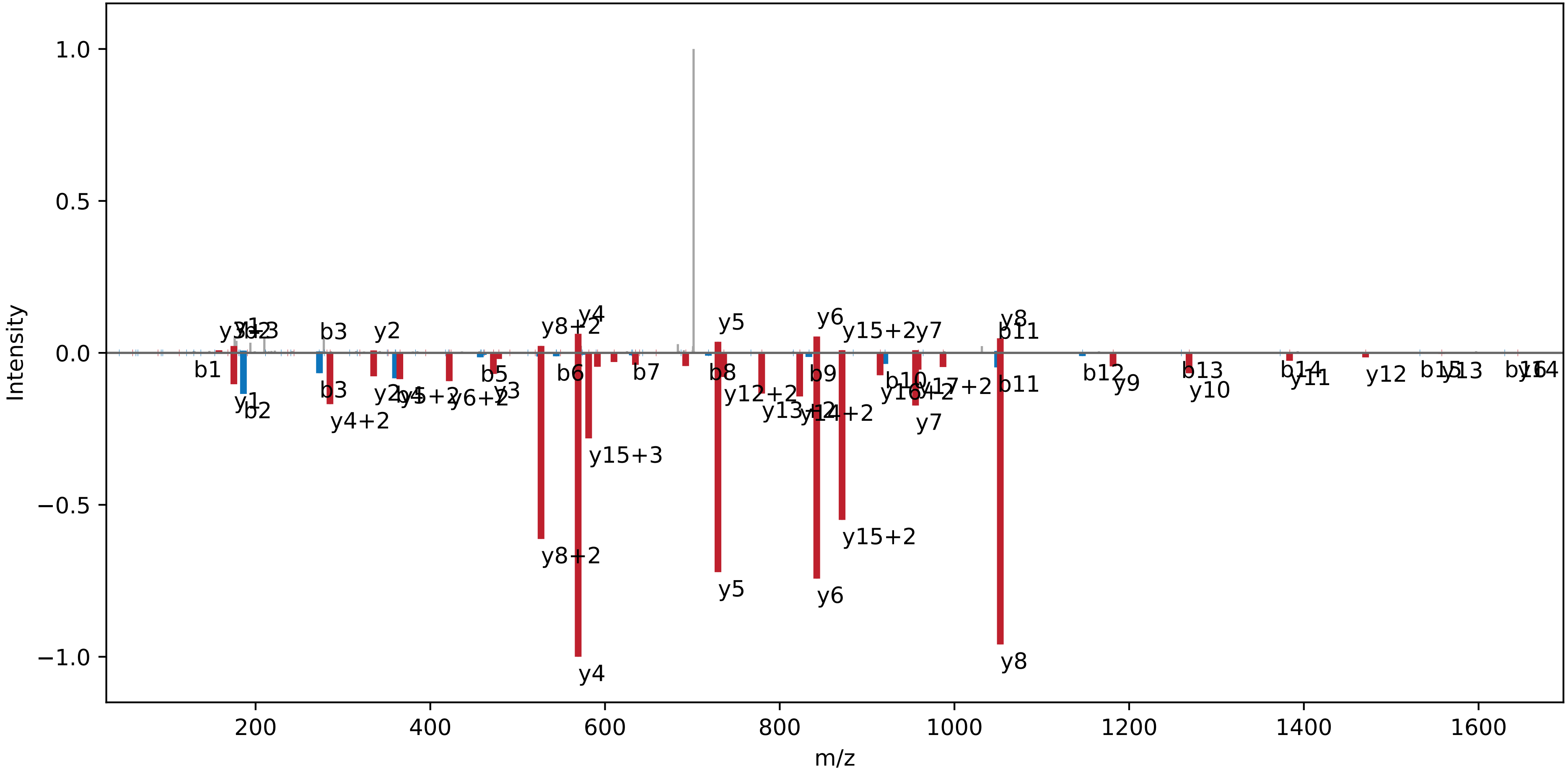

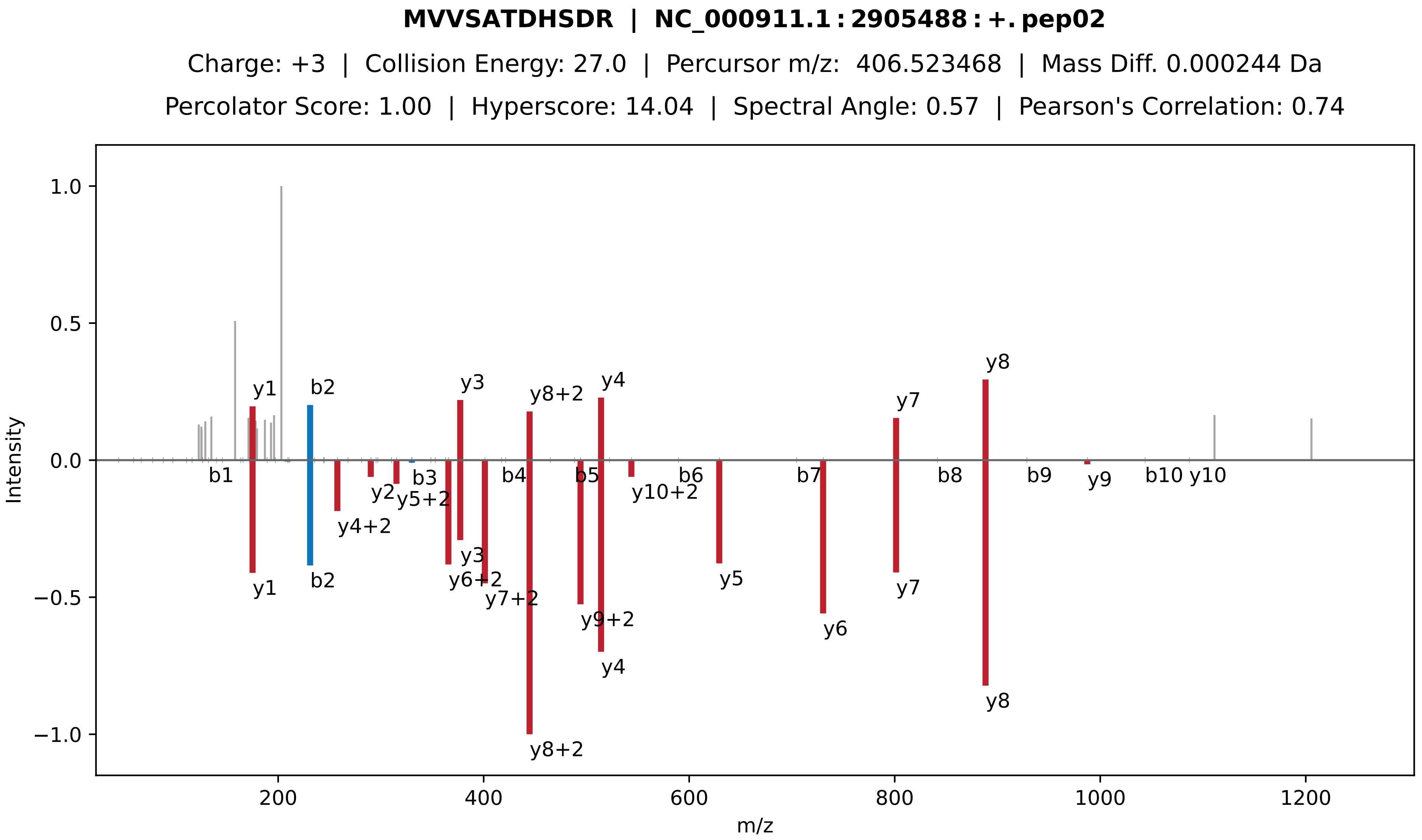

MVVSATDHSDR | NC_000911.1:2905488:+.pep02
Charge: +3 | Collision Energy: 27.0 | Percursor m/z: 406.523468 | Mass Diff. 0.000244 Da
Percolator Score: 1.00 | Hyperscore: 14.04 | Spectral Angle: 0.57 | Pearson's Correlation: 0.74
Intensity
m/z
y1
b2
y3
y8+2
y4
y7
y8
b1
y2
b3
y5+2
y4+2
b4
b5
b6
y10+2
y6+2
y7+2
y9+2
y5
b7
y6
b8
b9
y9
b10
y10

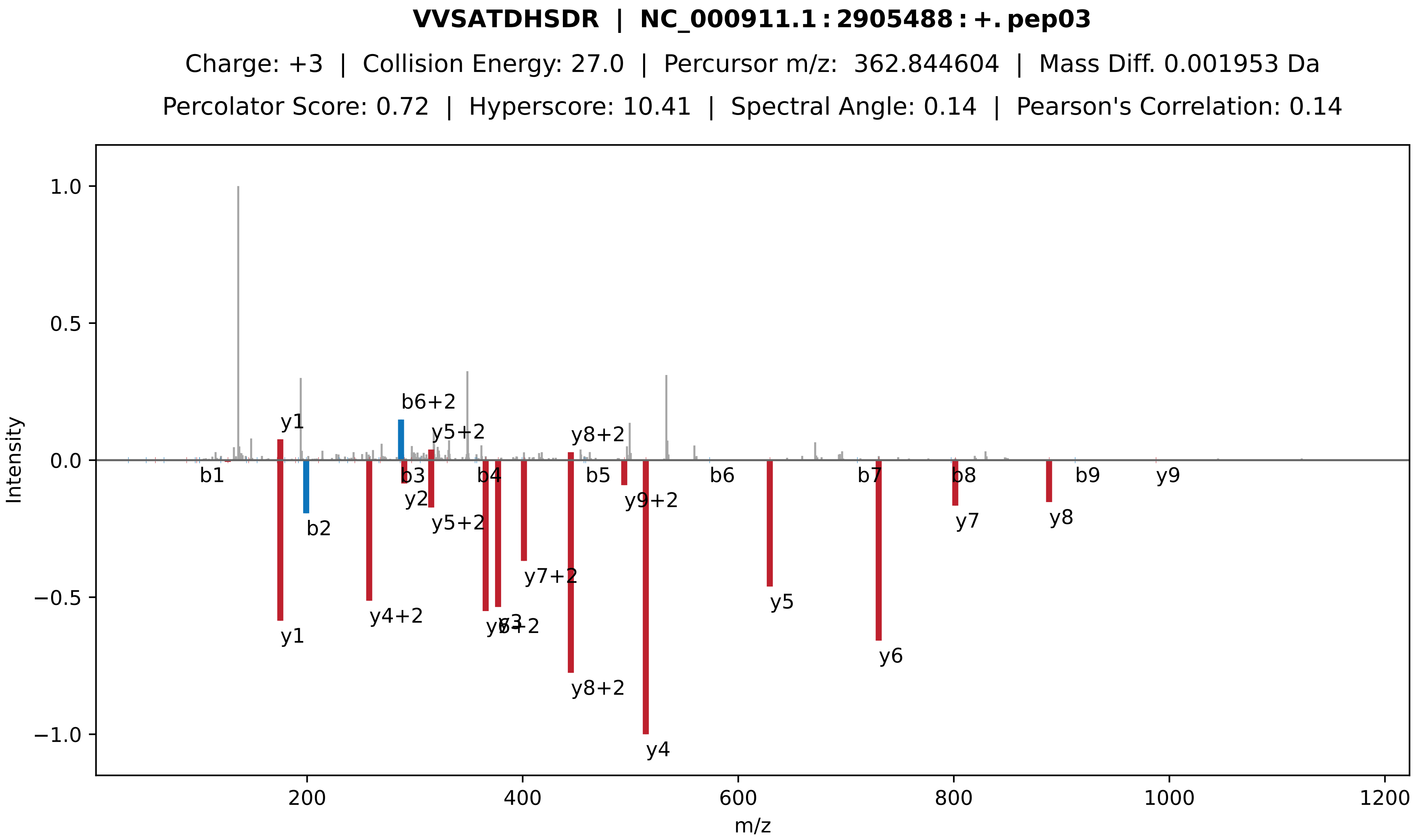
VVSATDHSDR | NC_000911.1:2905488:+.pep03
Charge: +3 | Collision Energy: 27.0 | Percursor m/z: 362.844604 | Mass Diff. 0.001953 Da
Percolator Score: 0.72 | Hyperscore: 10.41 | Spectral Angle: 0.14 | Pearson's Correlation: 0.14
Intensity
m/z
y1
b6+2
y5+2
y8+2
b1
b2
b3
y2
y5+2
b4
y4+2
y1
y3
y6+2
y7+2
y8+2
b5
y9+2
y4
b6
y5
b7
y6
b8
y7
y8
b9
y9

**VVSATDHSDRTTSQTGDR | NC_000911.1 : 2905488 : +. pep04**

Charge: +4 | Collision Energy: 27.0 | Percursor m/z: 483.980896 | Mass Diff. 0.003174 Da

Percolator Score: 1.00 | Hyperscore: 39.09 | Spectral Angle: 0.74 | Pearson's Correlation: 0.92

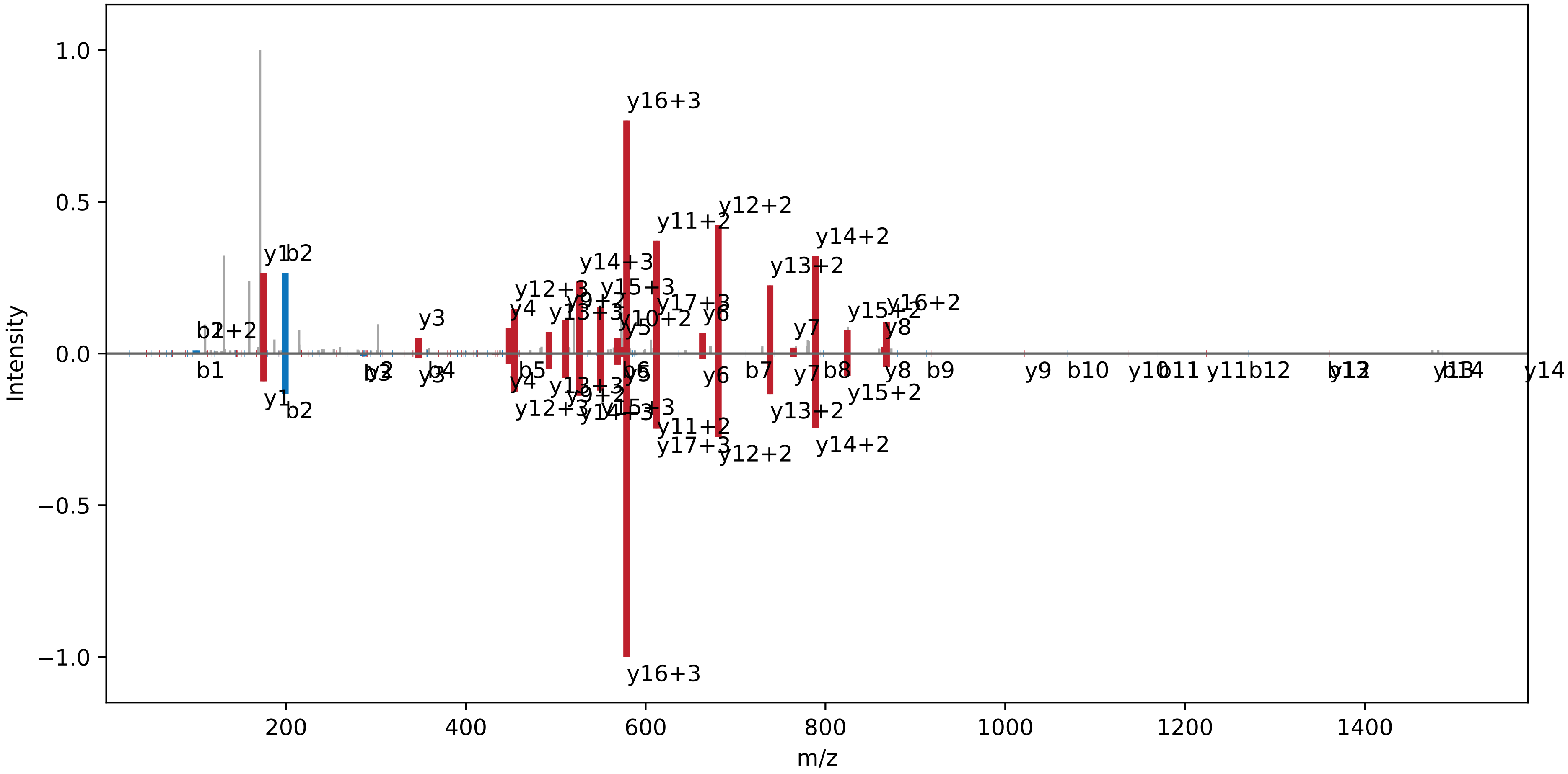

**ELEEWAQTDER | NC_000911.1 : 3042487 : −. pep01**

Charge: +2 | Collision Energy: 27.0 | Percursor m/z: 703.315735 | Mass Diff. 0.000610 Da

Percolator Score: 1.00 | Hyperscore: 19.51 | Spectral Angle: 0.83 | Pearson's Correlation: 0.96

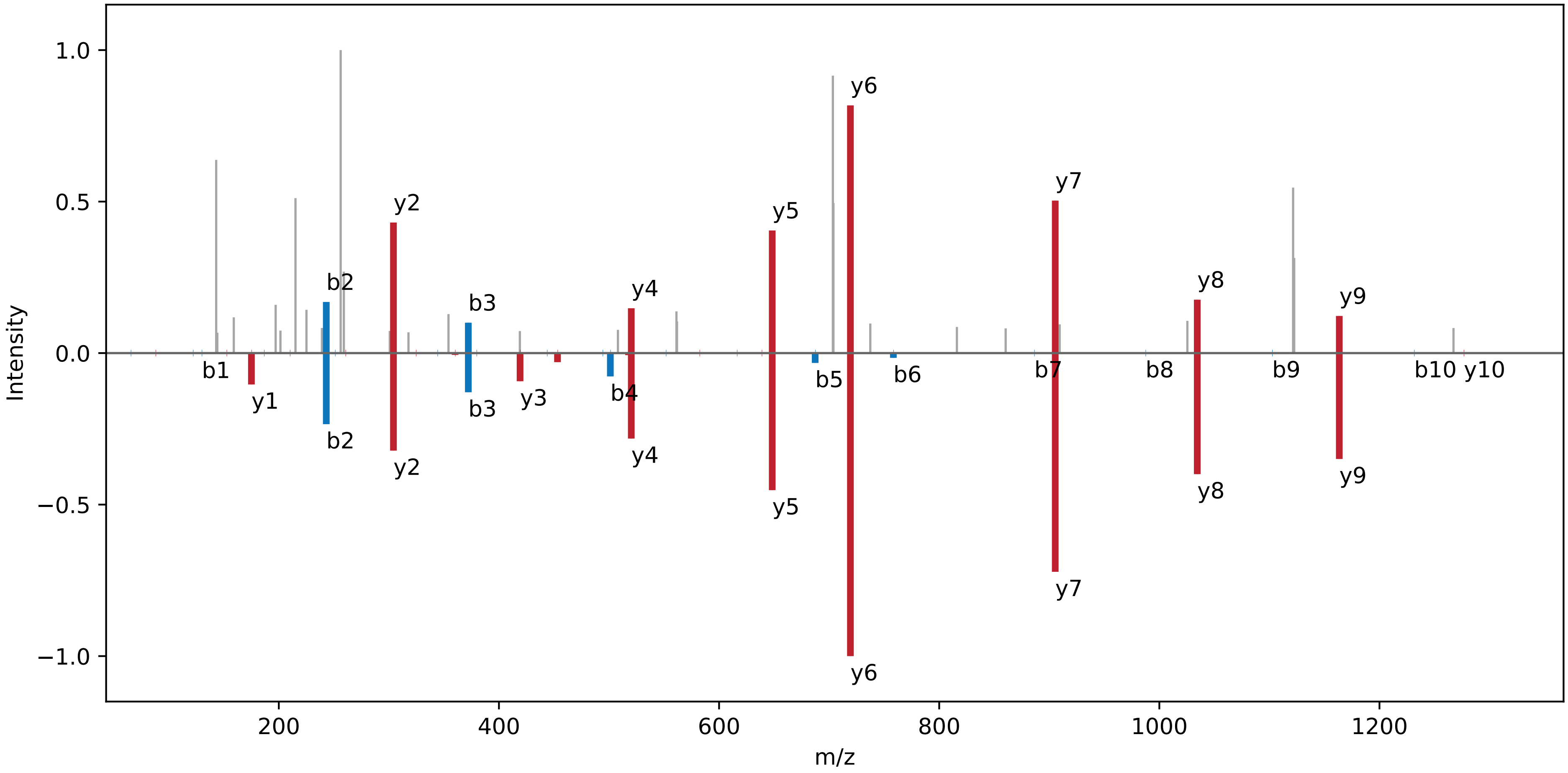

**[TMT] − SVSWLVAK[TMT] | NC_000911.1 : 3042487 : −. pep02**

Charge: +2 | Collision Energy: 32.5 | Percursor m/z: 674.422791 | Mass Diff. -0.000122 Da

Percolator Score: 1.00 | Hyperscore: 28.77 | Spectral Angle: 0.66 | Pearson's Correlation: 0.76

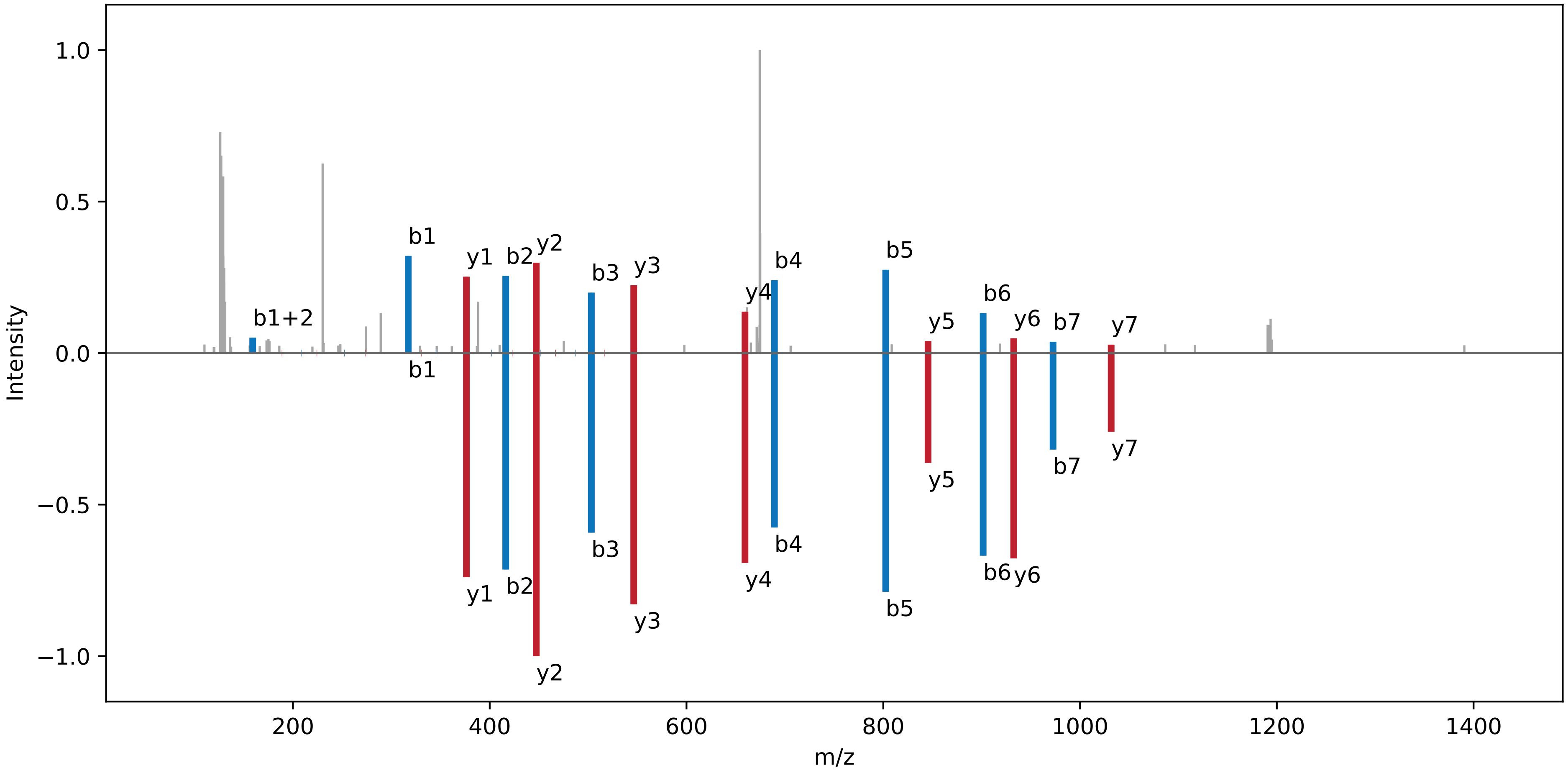

**VVAYIPAER | NC_000911.1 : 3042487 : −. pep03**

Charge: +2 | Collision Energy: 27.0 | Percursor m/z: 509.291473 | Mass Diff. 0.000977 Da

Percolator Score: 1.00 | Hyperscore: 24.40 | Spectral Angle: 0.94 | Pearson's Correlation: 0.99

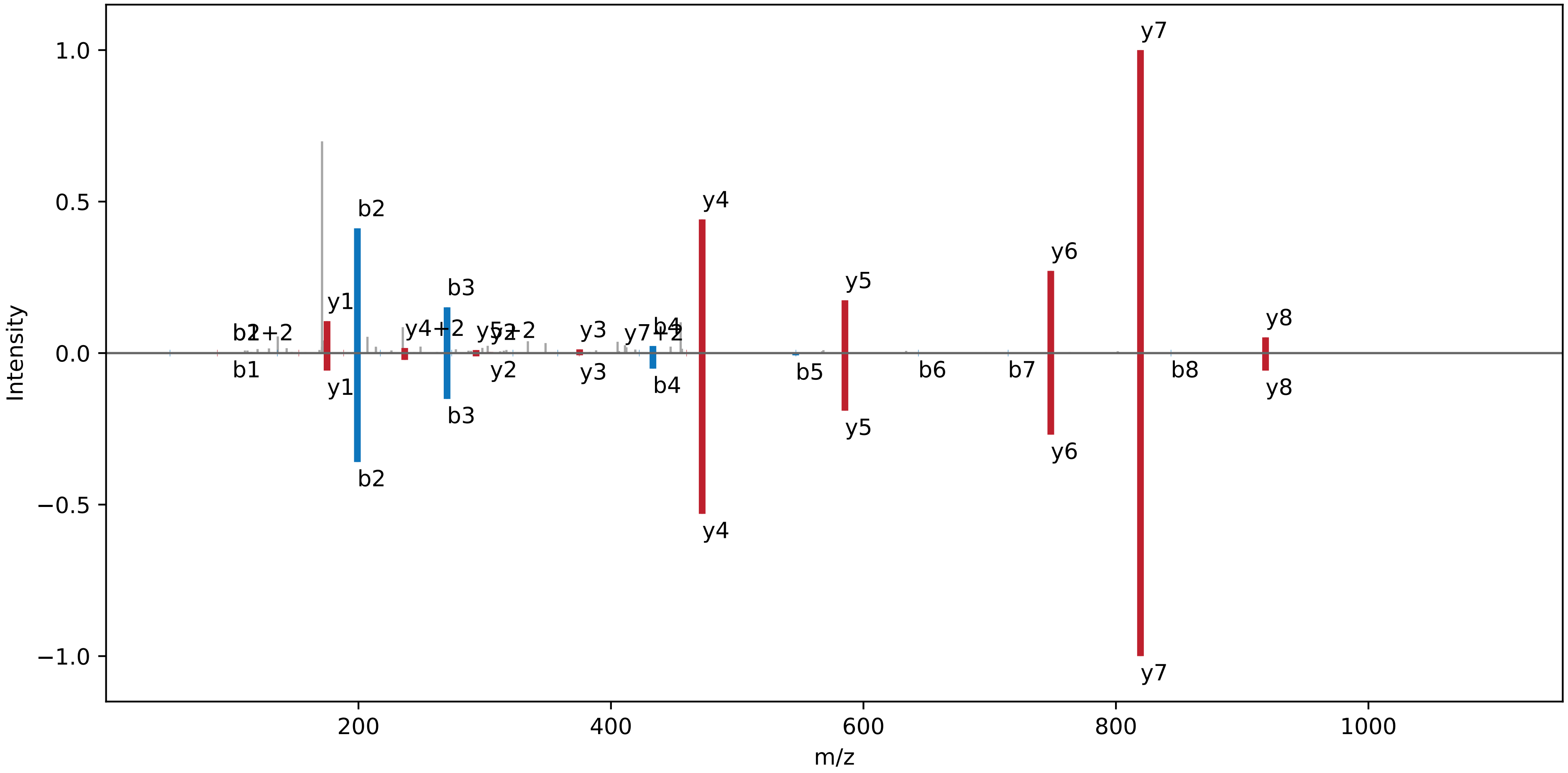

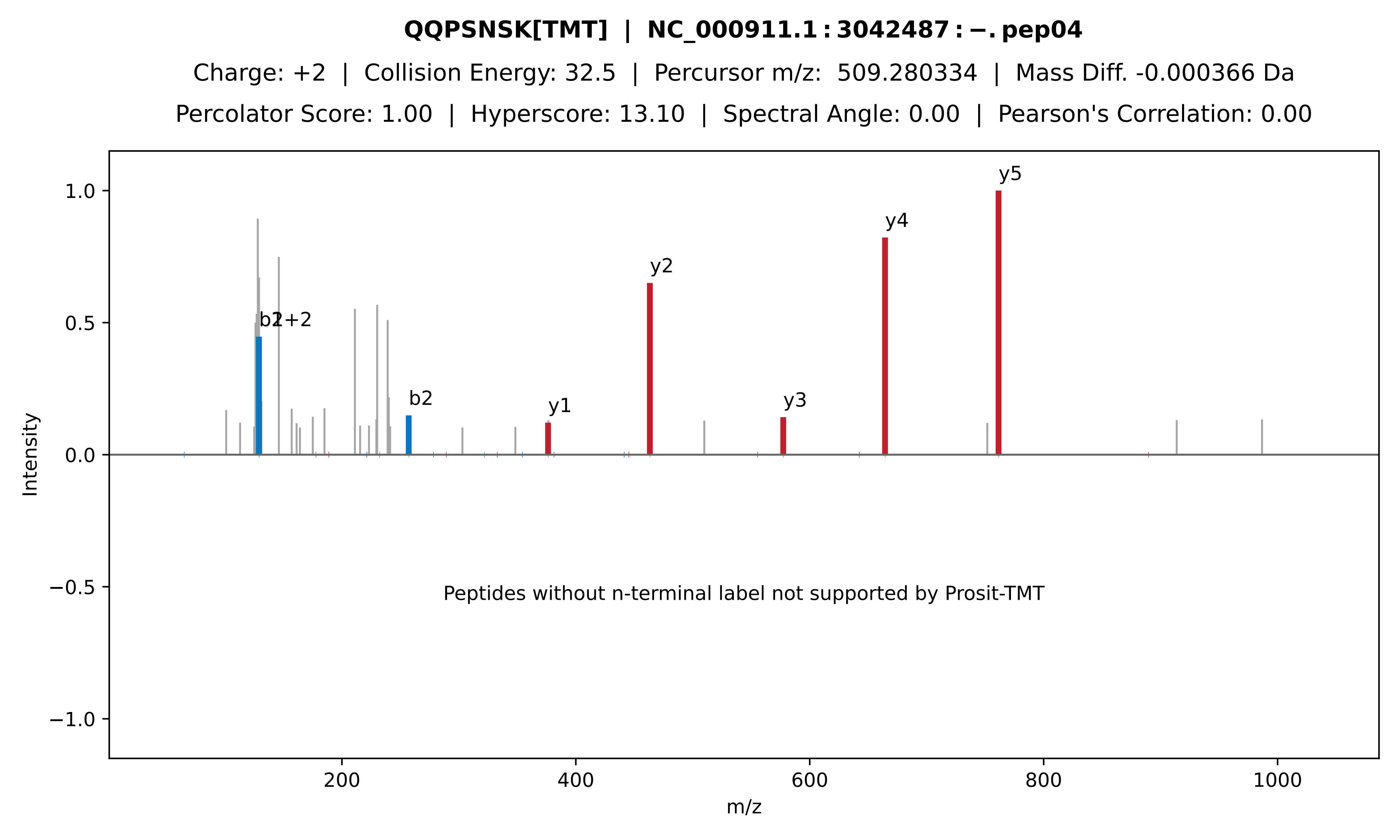
QQPSNSK[TMT] | NC_000911.1 : 3042487 : −. pep04
Charge: +2 | Collision Energy: 32.5 | Percursor m/z: 509.280334 | Mass Diff. -0.000366 Da
Percolator Score: 1.00 | Hyperscore: 13.10 | Spectral Angle: 0.00 | Pearson's Correlation: 0.00
Intensity
m/z
y5
y4
y2
y3
y1
b2
Peptides without n-terminal label not supported by Prosit-TMT

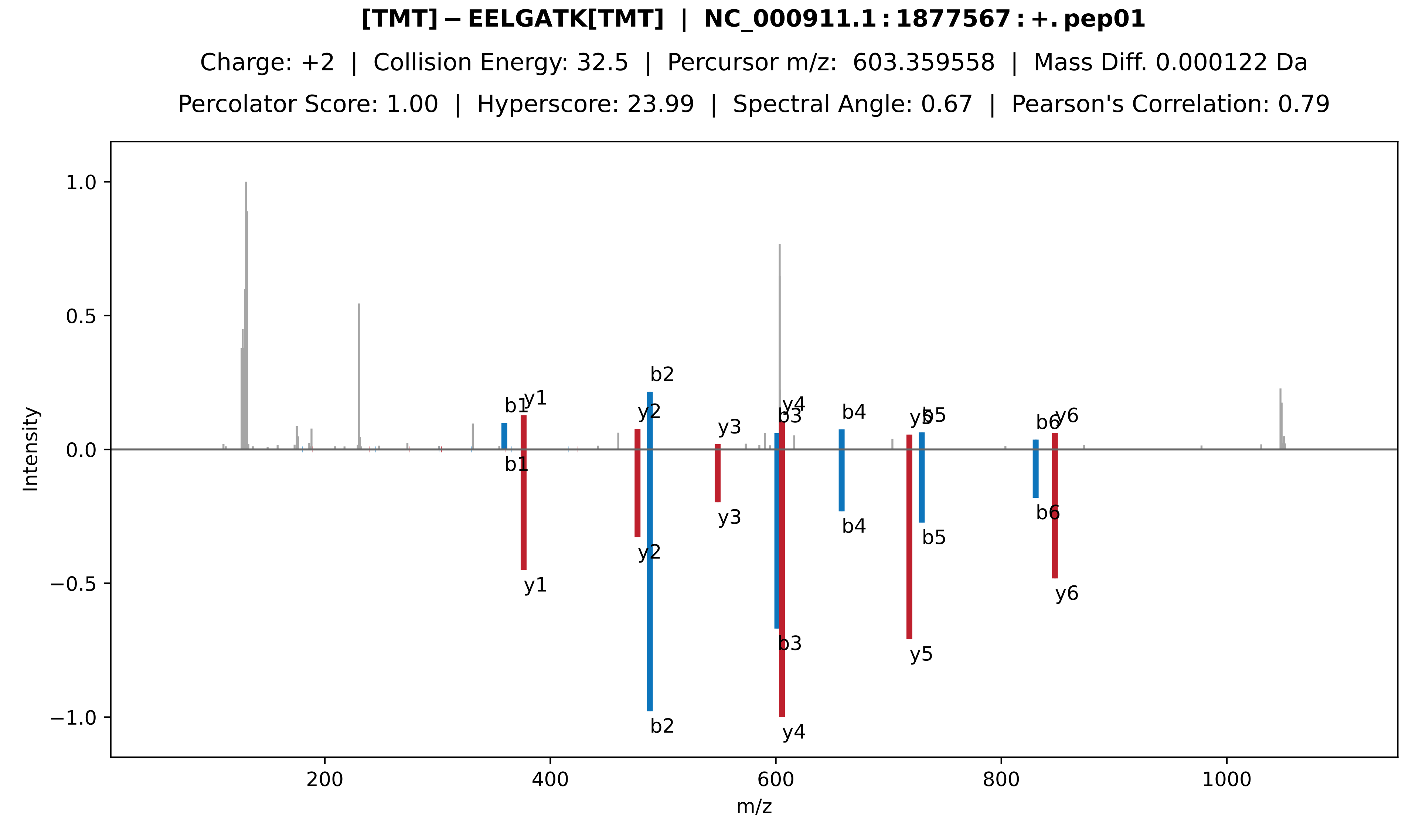
[TMT]−EELGATK[TMT] | NC_000911.1:1877567:+.pep01
Charge: +2 | Collision Energy: 32.5 | Percursor m/z: 603.359558 | Mass Diff. 0.000122 Da
Percolator Score: 1.00 | Hyperscore: 23.99 | Spectral Angle: 0.67 | Pearson's Correlation: 0.79
Intensity
m/z
b1
y1
b2
y2
y3
b3
y4
b4
y5
b5
b6
y6

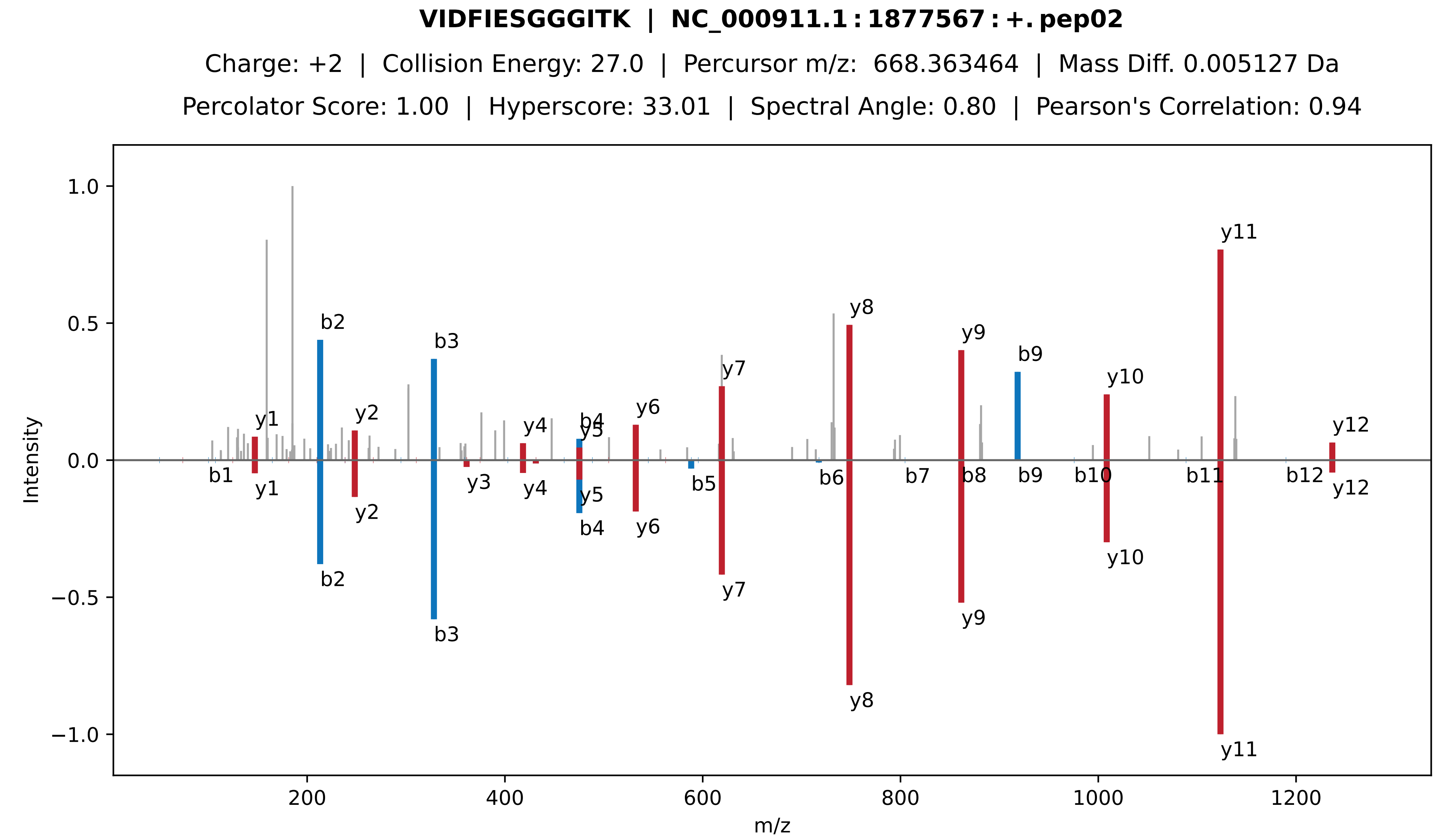
VIDFIESGGGITK | NC_000911.1 : 1877567 : +. pep02
Charge: +2 | Collision Energy: 27.0 | Percursor m/z: 668.363464 | Mass Diff. 0.005127 Da
Percolator Score: 1.00 | Hyperscore: 33.01 | Spectral Angle: 0.80 | Pearson's Correlation: 0.94
Intensity
1.0
0.5
0.0
−0.5
−1.0
m/z
200
400
600
800
1000
1200
y1
b2
y2
b3
y3
y4
b4
y5
y6
y7
y8
y9
b9
y10
y11
y12
b1
b5
b6
b7
b8
b10
b11
b12

**FGVTPASLCYQFK | NC_000911.1 : 1877567 : +. pep03**

Charge: +2 | Collision Energy: 30.0 | Percursor m/z: 759.380200 | Mass Diff. 0.002441 Da

Percolator Score: 1.00 | Hyperscore: 17.39 | Spectral Angle: 0.50 | Pearson's Correlation: 0.65

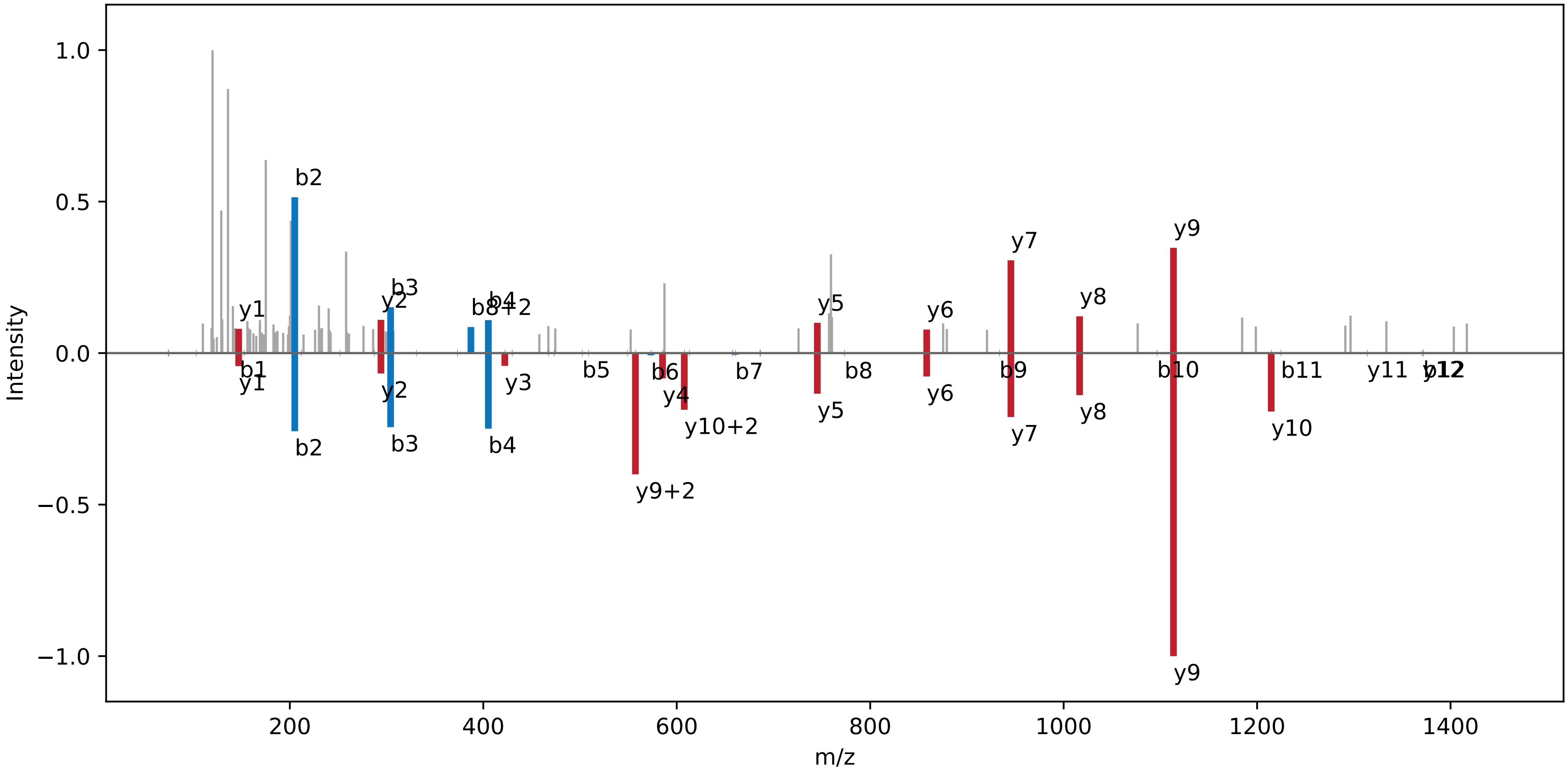

**TTNSPSPNGPASASPGLLSR | NC_000911.1 : 1604689 : +. pep01**

Charge: +2 | Collision Energy: 35.0 | Percursor m/z: 955.980300 | Mass Diff. 0.000366 Da

Percolator Score: 1.00 | Hyperscore: 40.43 | Spectral Angle: 0.16 | Pearson's Correlation: 0.16

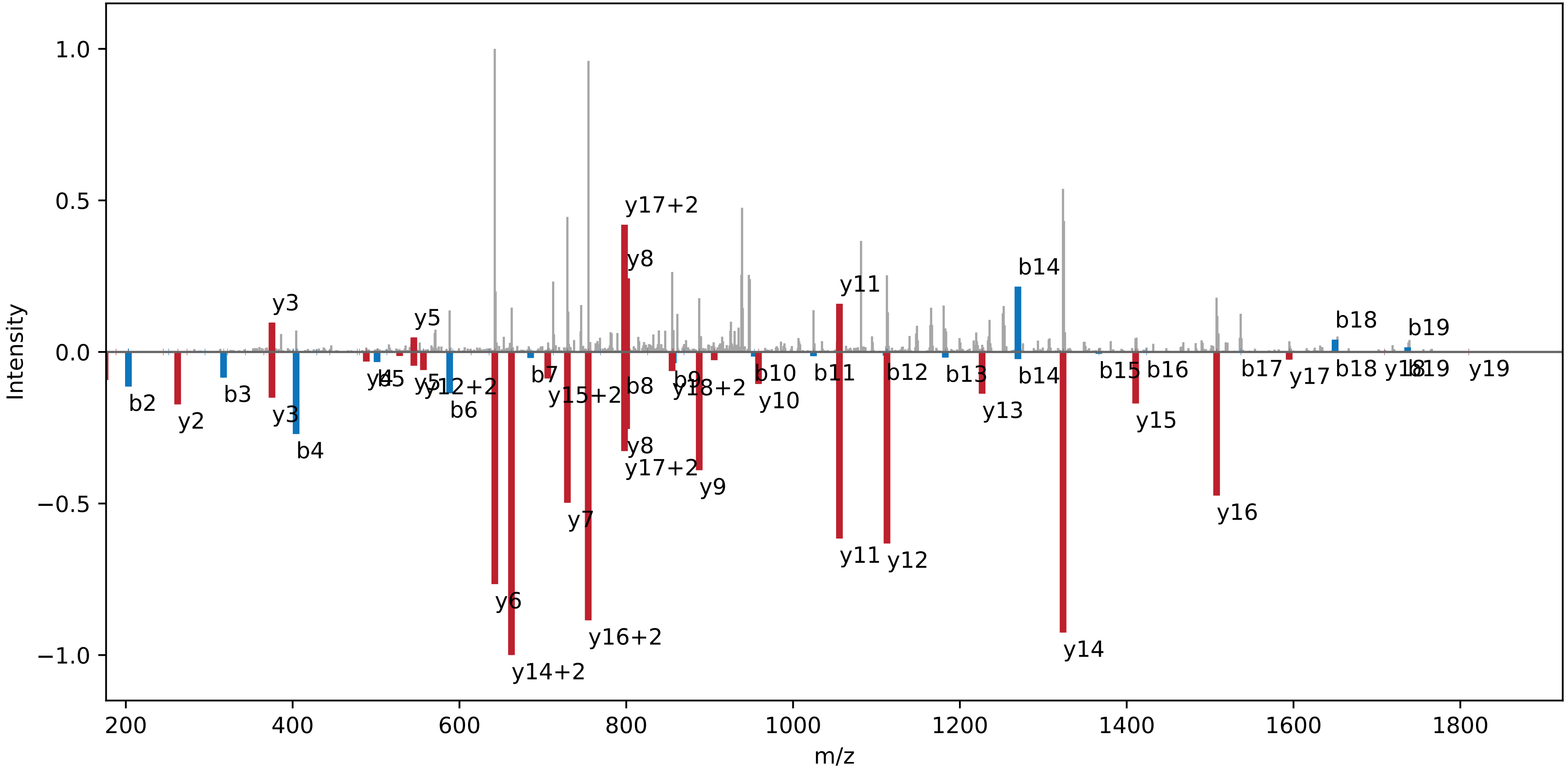

**KYNEEM[Ox]R | NC_000911.1 : 1604689 : +. pep02**

Charge: +2 | Collision Energy: 27.0 | Percursor m/z: 493.221344 | Mass Diff. -0.008972 Da

Percolator Score: 0.19 | Hyperscore: 8.11 | Spectral Angle: 0.47 | Pearson's Correlation: 0.63

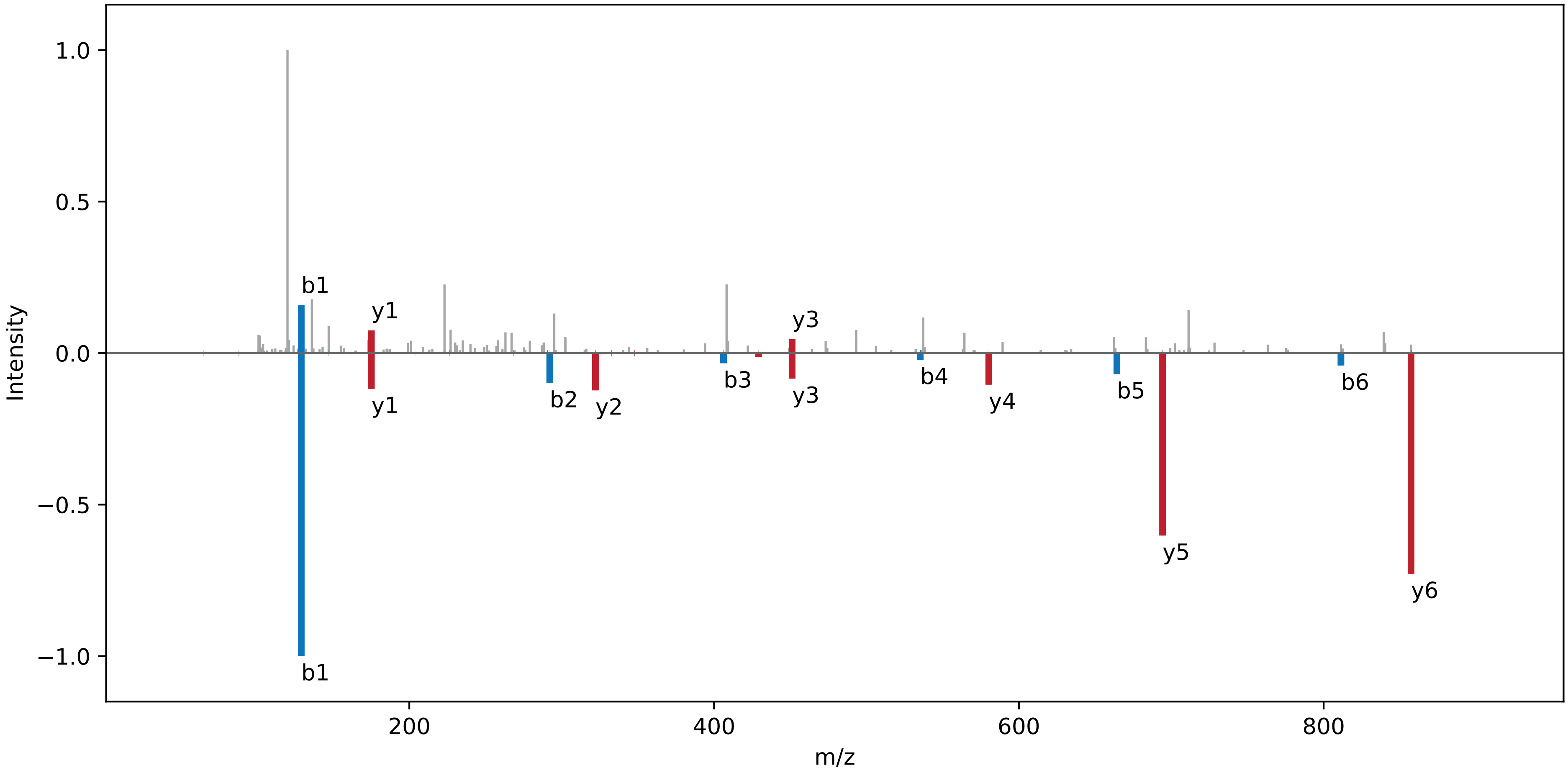

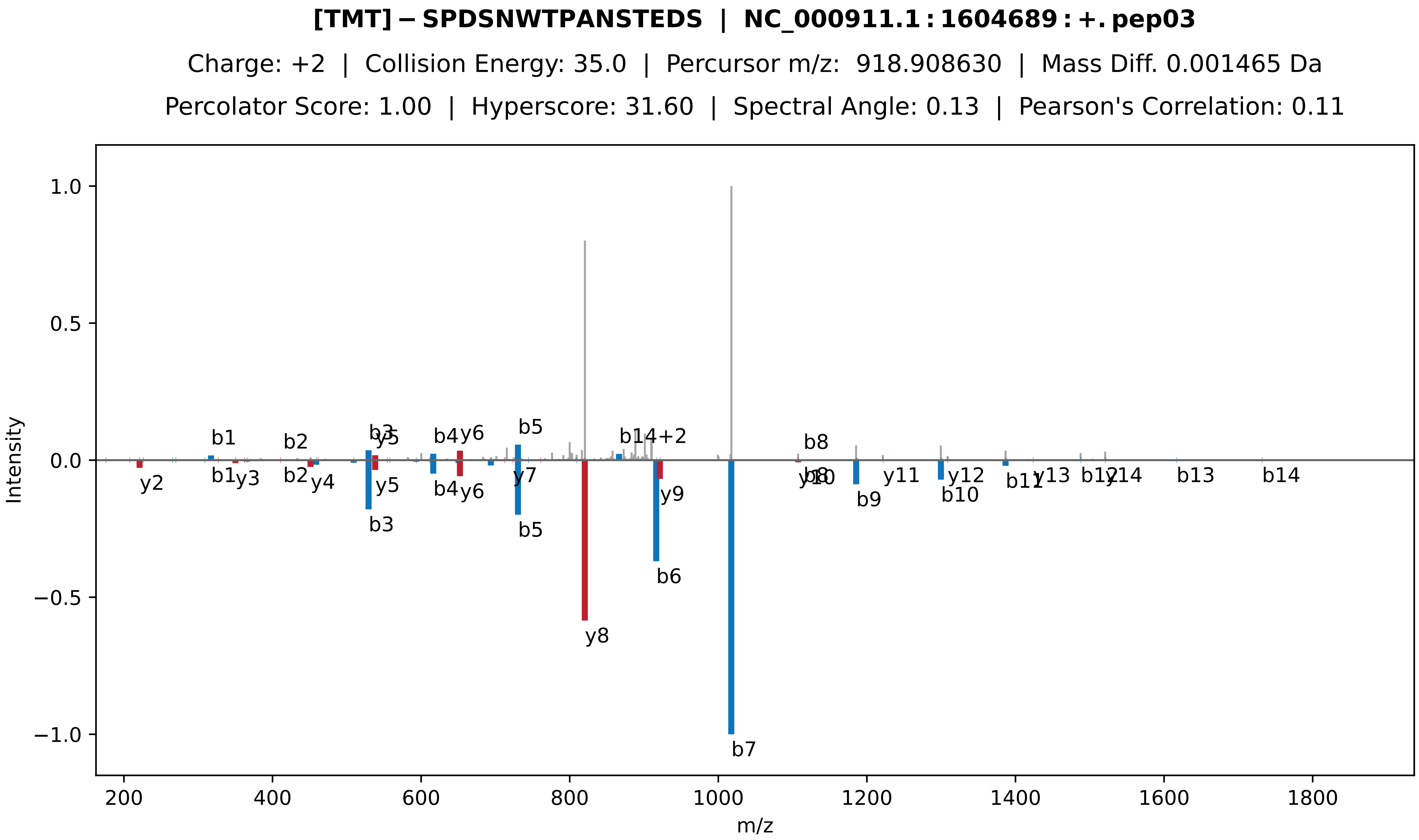

[TMT]−SPDSNWTPANSTEDS | NC_000911.1:1604689:+.pep03
Charge: +2 | Collision Energy: 35.0 | Percursor m/z: 918.908630 | Mass Diff. 0.001465 Da
Percolator Score: 1.00 | Hyperscore: 31.60 | Spectral Angle: 0.13 | Pearson's Correlation: 0.11
Intensity
m/z
1.0
0.5
0.0
−0.5
−1.0
200
400
600
800
1000
1200
1400
1600
1800
b1
b2
b3
y5
b4
y6
b5
b14+2
b8
y2
y3
y4
y7
y8
y9
b6
b7
b9
y11
y12
b10
b11
y13
y14
b13
b14

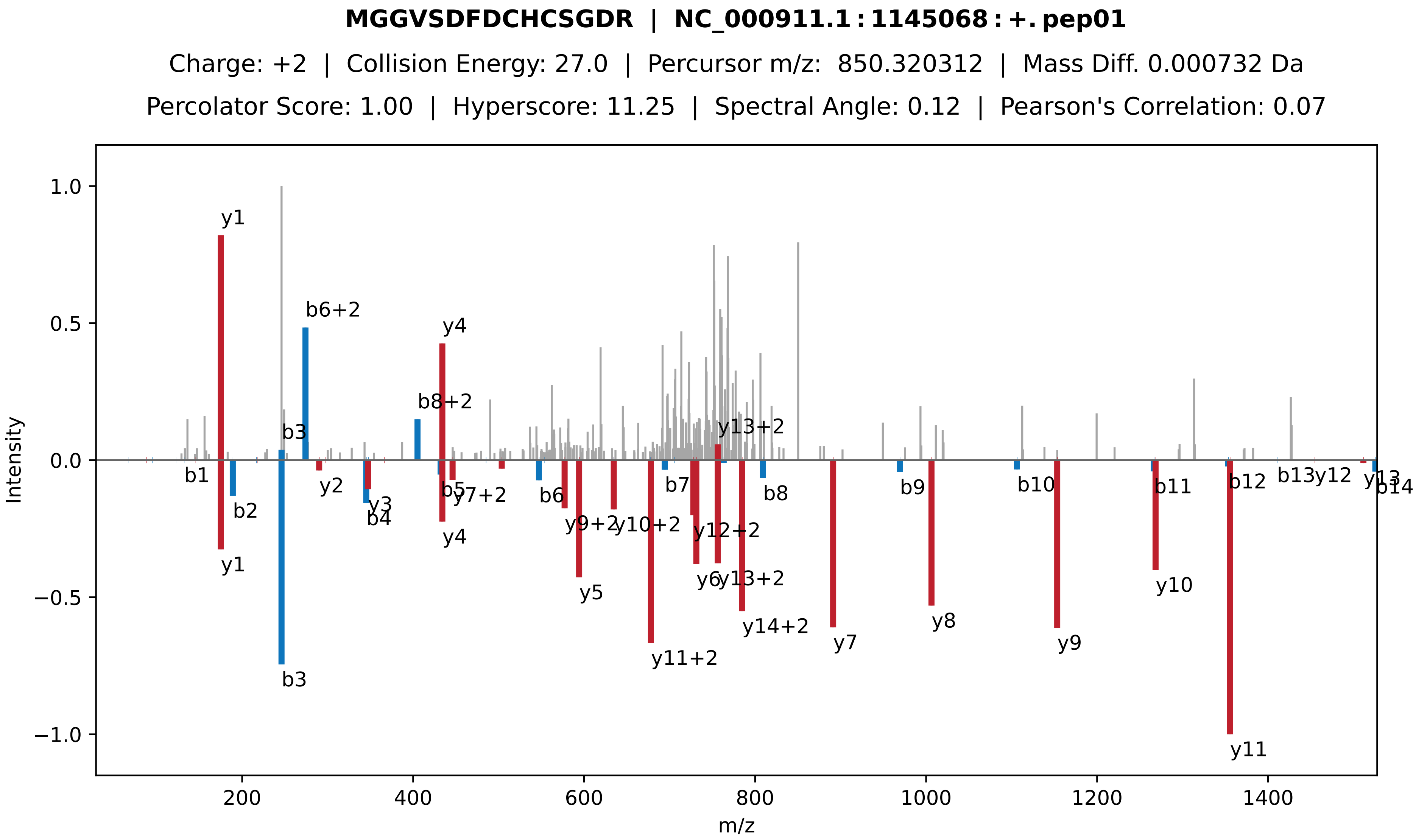
MGGVSDFDCHCSGDR | NC_000911.1 : 1145068 : +. pep01
Charge: +2 | Collision Energy: 27.0 | Percursor m/z: 850.320312 | Mass Diff. 0.000732 Da
Percolator Score: 1.00 | Hyperscore: 11.25 | Spectral Angle: 0.12 | Pearson's Correlation: 0.07
Intensity
m/z
1.0
0.5
0.0
−0.5
−1.0
200
400
600
800
1000
1200
1400
y1
b6+2
y4
b8+2
b3
y13+2
b1
b2
y2
y3
b4
b5
y7+2
b6
y9+2
y10+2
b7
y12+2
b8
b9
b10
b11
b12
b13
y12
y13
b14
y5
y6
y14+2
y11+2
y7
y8
y9
y10
y11

**GGVSDFDCHCSGDRRGGHR | NC_000911.1 : 1145068 : +. pep02**

Charge: +2 | Collision Energy: 27.0 | Percursor m/z: 1066.464400 | Mass Diff. 0.024414 Da

Percolator Score: 0.61 | Hyperscore: 9.04 | Spectral Angle: 0.04 | Pearson's Correlation: 0.01

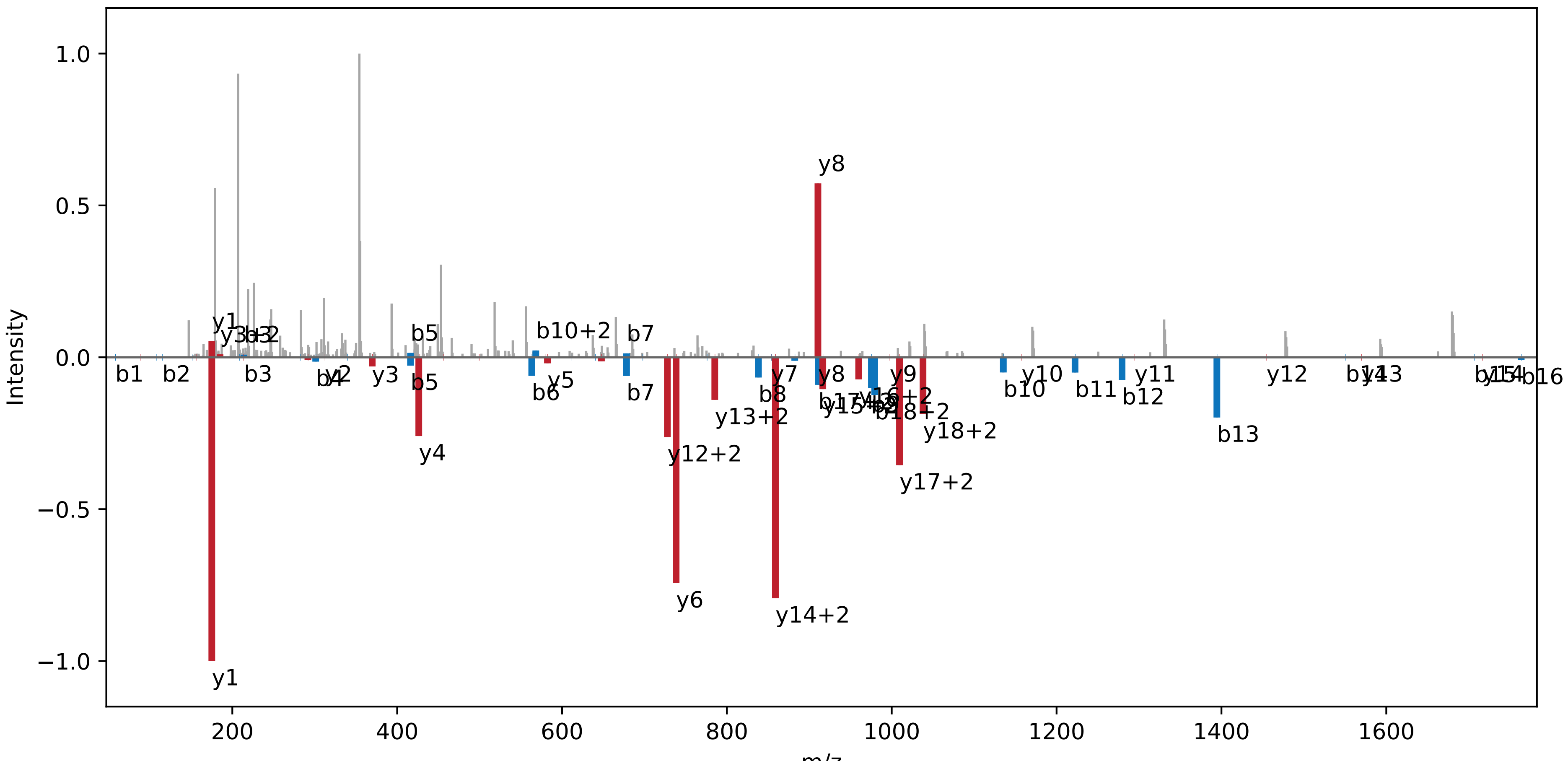

**IDSSFDQHGINLPLAQTPEQP | NC_000911.1 : 385352 : +. pep01**

Charge: +2 | Collision Energy: 27.0 | Percursor m/z: 1154.069100 | Mass Diff. 0.000244 Da

Percolator Score: 1.00 | Hyperscore: 35.64 | Spectral Angle: 0.80 | Pearson's Correlation: 0.95

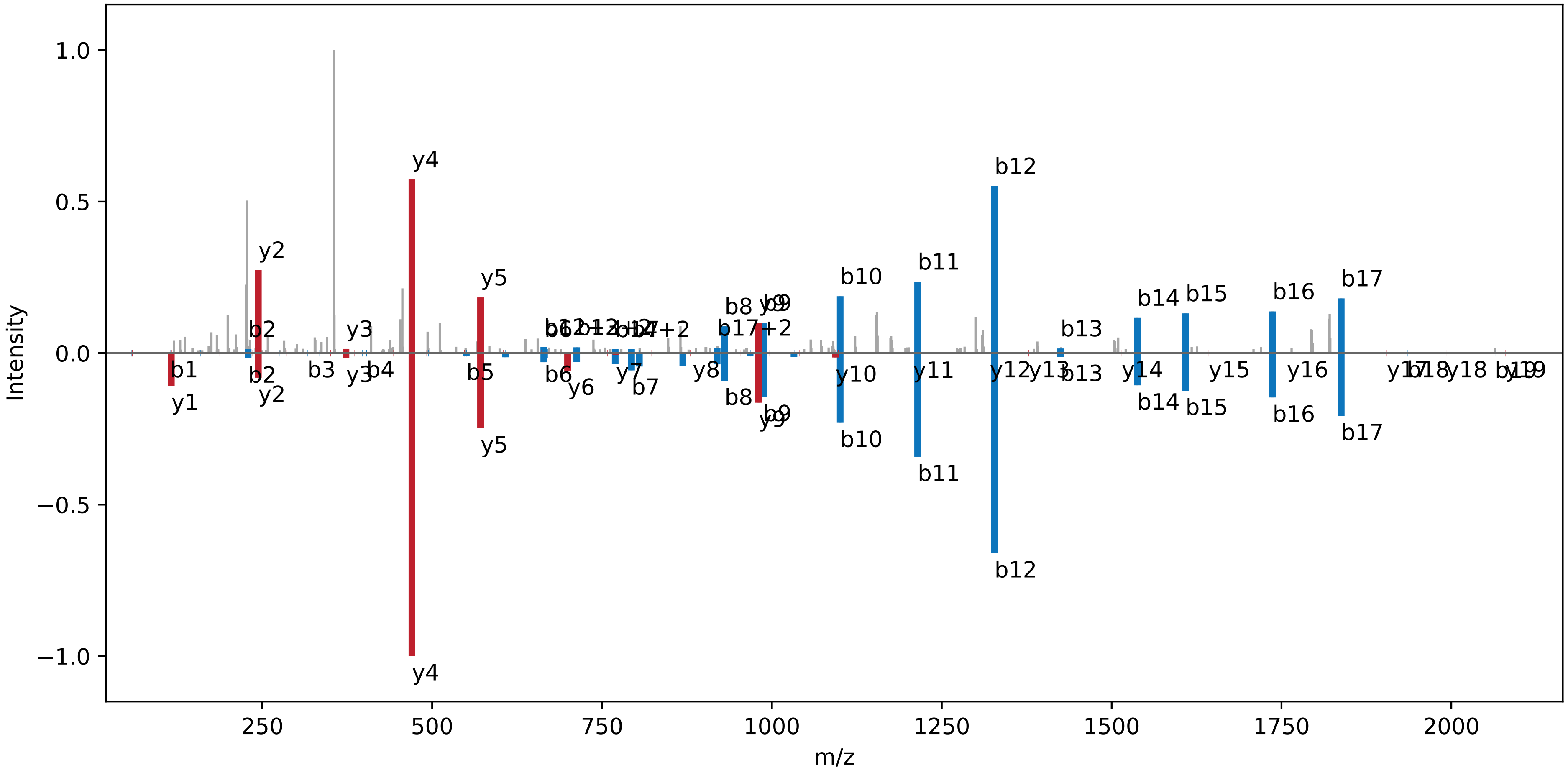

**ATSTPPPSNSSPSPSSTGVSWR | NC_000911.1:385352:+.pep02**

Charge: +2 | Collision Energy: 27.0 | Percursor m/z: 1094.017822 | Mass Diff. -0.001465 Da

Percolator Score: 1.00 | Hyperscore: 35.29 | Spectral Angle: 0.89 | Pearson's Correlation: 0.99

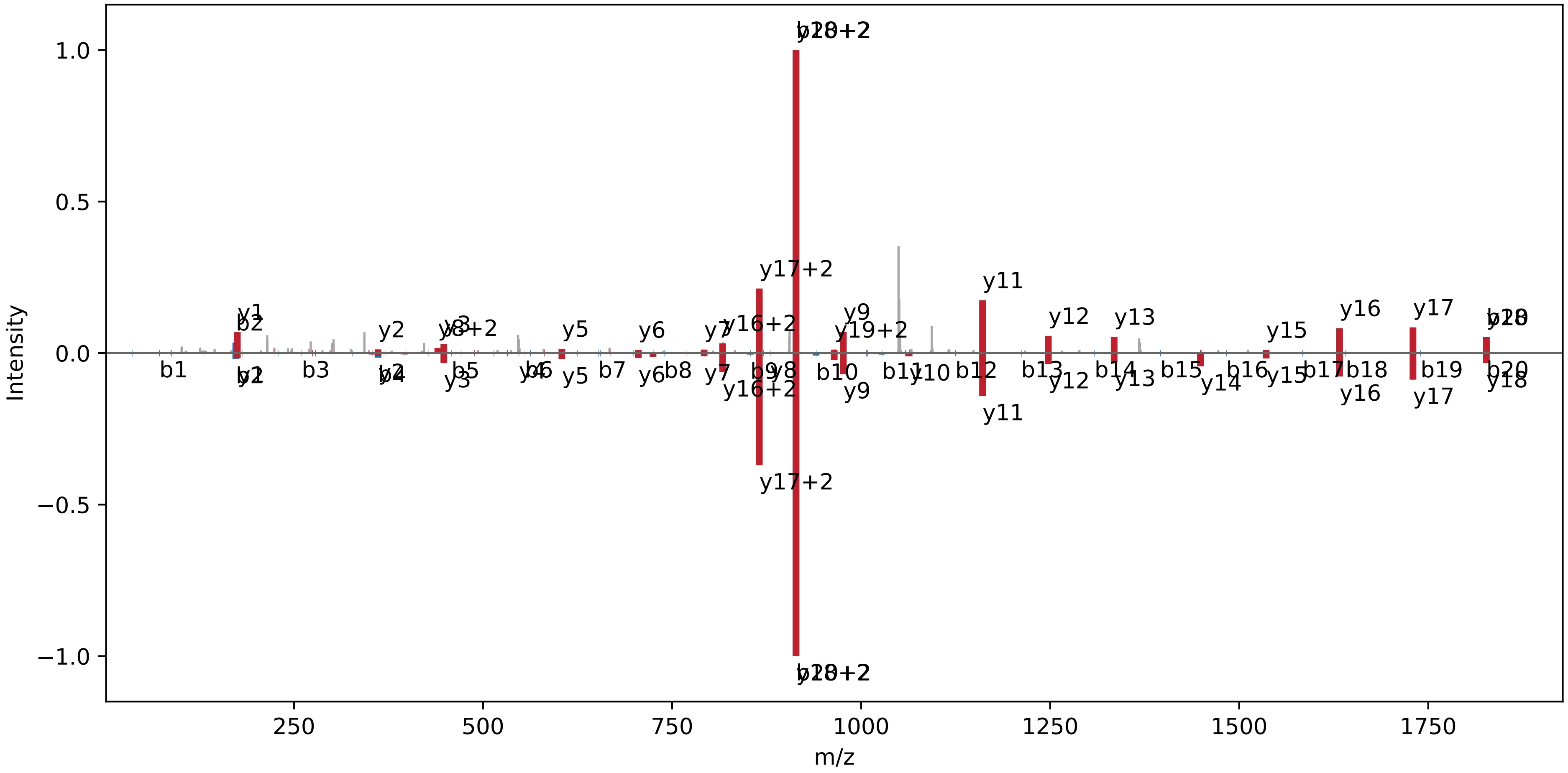

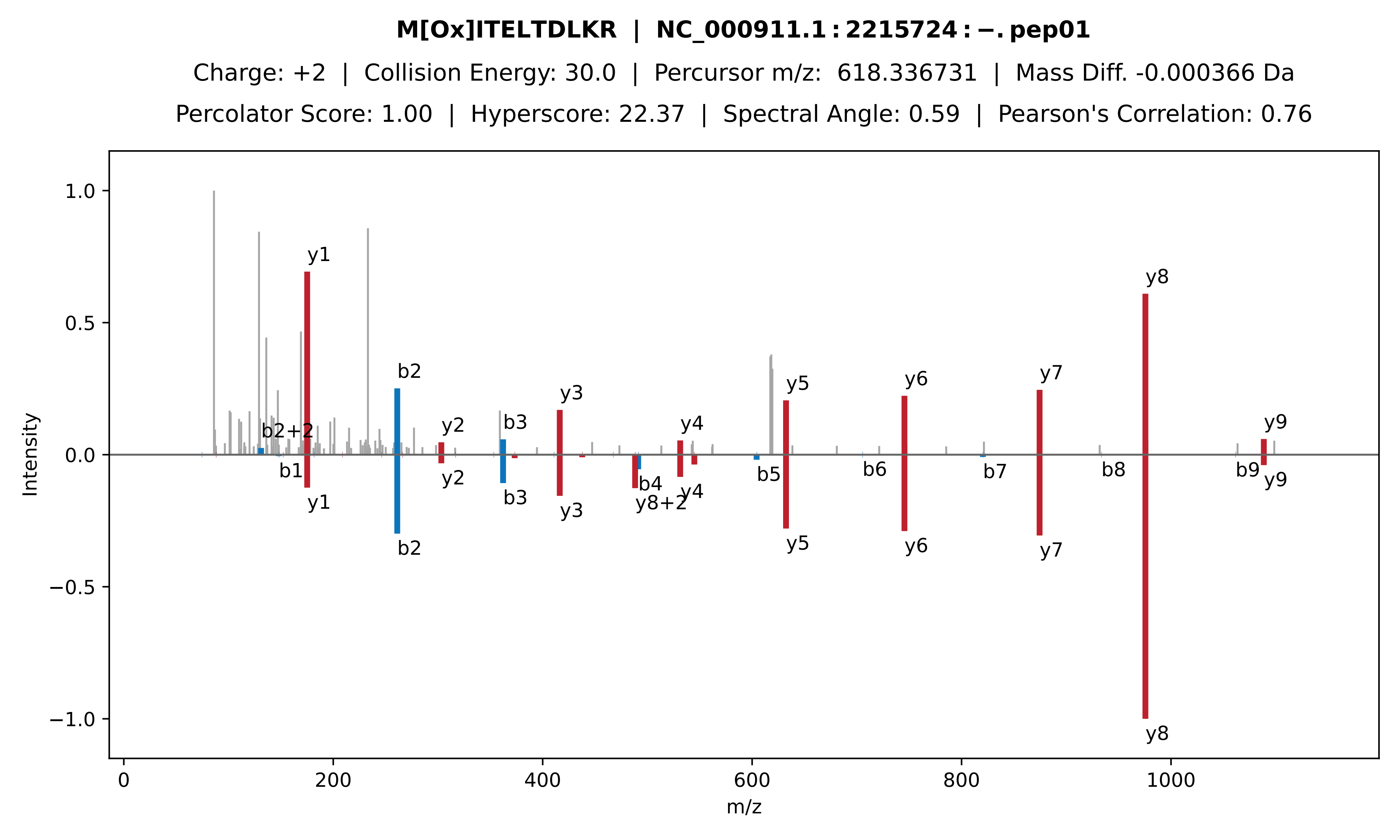

M[Ox]ITELTDLKR | NC_000911.1 : 2215724 : −. pep01
Charge: +2 | Collision Energy: 30.0 | Percursor m/z: 618.336731 | Mass Diff. -0.000366 Da
Percolator Score: 1.00 | Hyperscore: 22.37 | Spectral Angle: 0.59 | Pearson's Correlation: 0.76
Intensity
m/z
1.0
0.5
0.0
−0.5
−1.0
0
200
400
600
800
1000
y1
b2+2
b1
b2
y2
b3
y3
b4
y8+2
y4
b5
y5
b6
y6
b7
y7
b8
y8
b9
y9

**MITELTDLK | NC_000911.1 : 2215724 : −. pep02**

Charge: +2 | Collision Energy: 27.0 | Percursor m/z: 532.287659 | Mass Diff. -0.001709 Da

Percolator Score: 1.00 | Hyperscore: 25.39 | Spectral Angle: 0.90 | Pearson's Correlation: 0.99

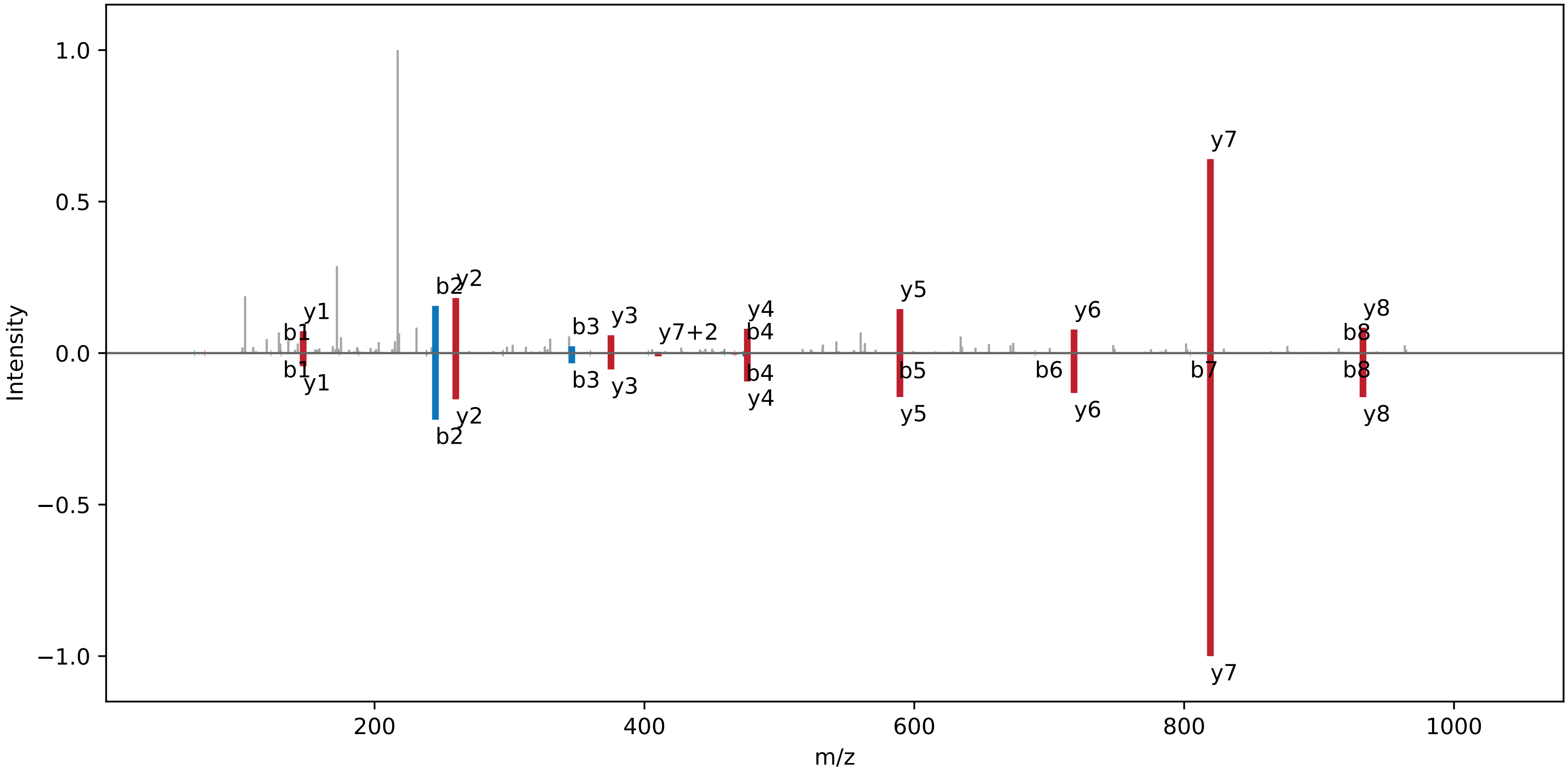

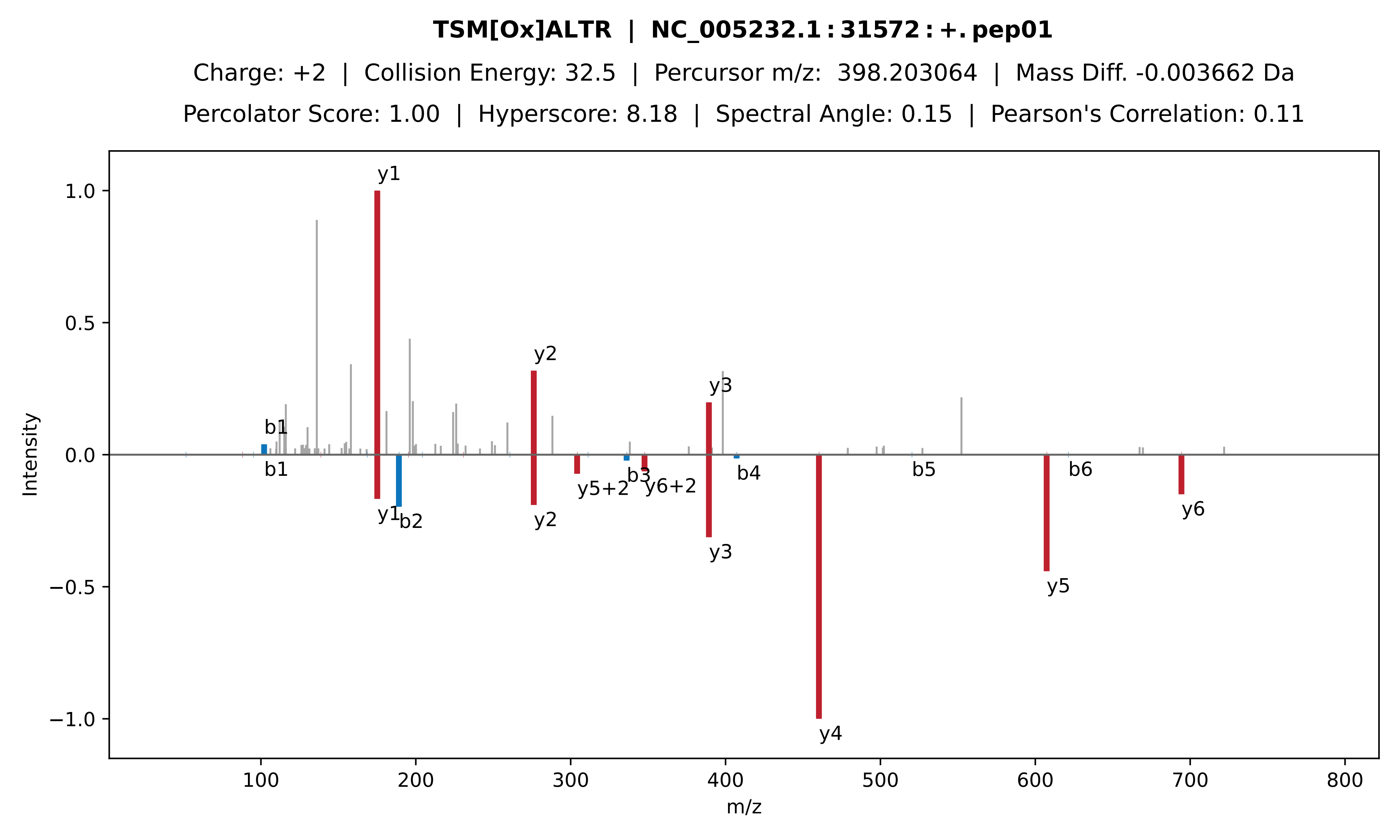

TSM[Ox]ALTR | NC_005232.1:31572:+.pep01
Charge: +2 | Collision Energy: 32.5 | Percursor m/z: 398.203064 | Mass Diff. -0.003662 Da
Percolator Score: 1.00 | Hyperscore: 8.18 | Spectral Angle: 0.15 | Pearson's Correlation: 0.11
Intensity
m/z
y1
y2
y3
b1
b1
y1
b2
y2
y5+2
b3
y6+2
y3
b4
y4
b5
y5
b6
y6

**SFSHGTLKHLDLFAGCGGFTLAAEQTR | NC_005232.1 : 31572 : +. pep02**

Charge: +3 | Collision Energy: 25.0 | Percursor m/z: 975.152100 | Mass Diff. 2.005615 Da

Percolator Score: 0.89 | Hyperscore: 14.41 | Spectral Angle: 0.32 | Pearson's Correlation: 0.48

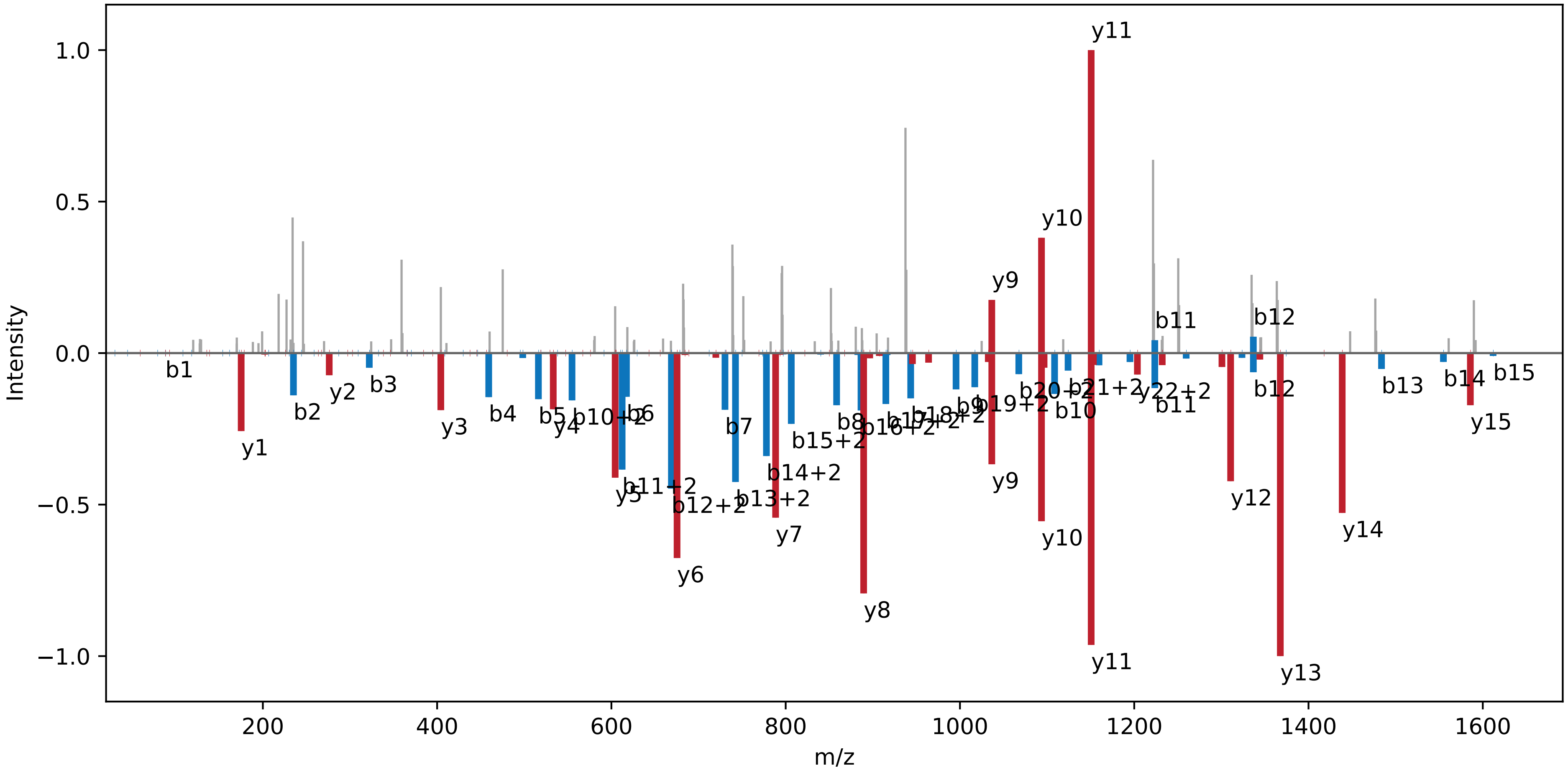

**CGQGPGDENDFYCLCTGK | NC_000911.1 : 1163779 : −. pep01**

Charge: +3 | Collision Energy: 28.0 | Percursor m/z: 694.273682 | Mass Diff. 2.994629 Da

Percolator Score: 1.00 | Hyperscore: 15.27 | Spectral Angle: 0.12 | Pearson's Correlation: 0.13

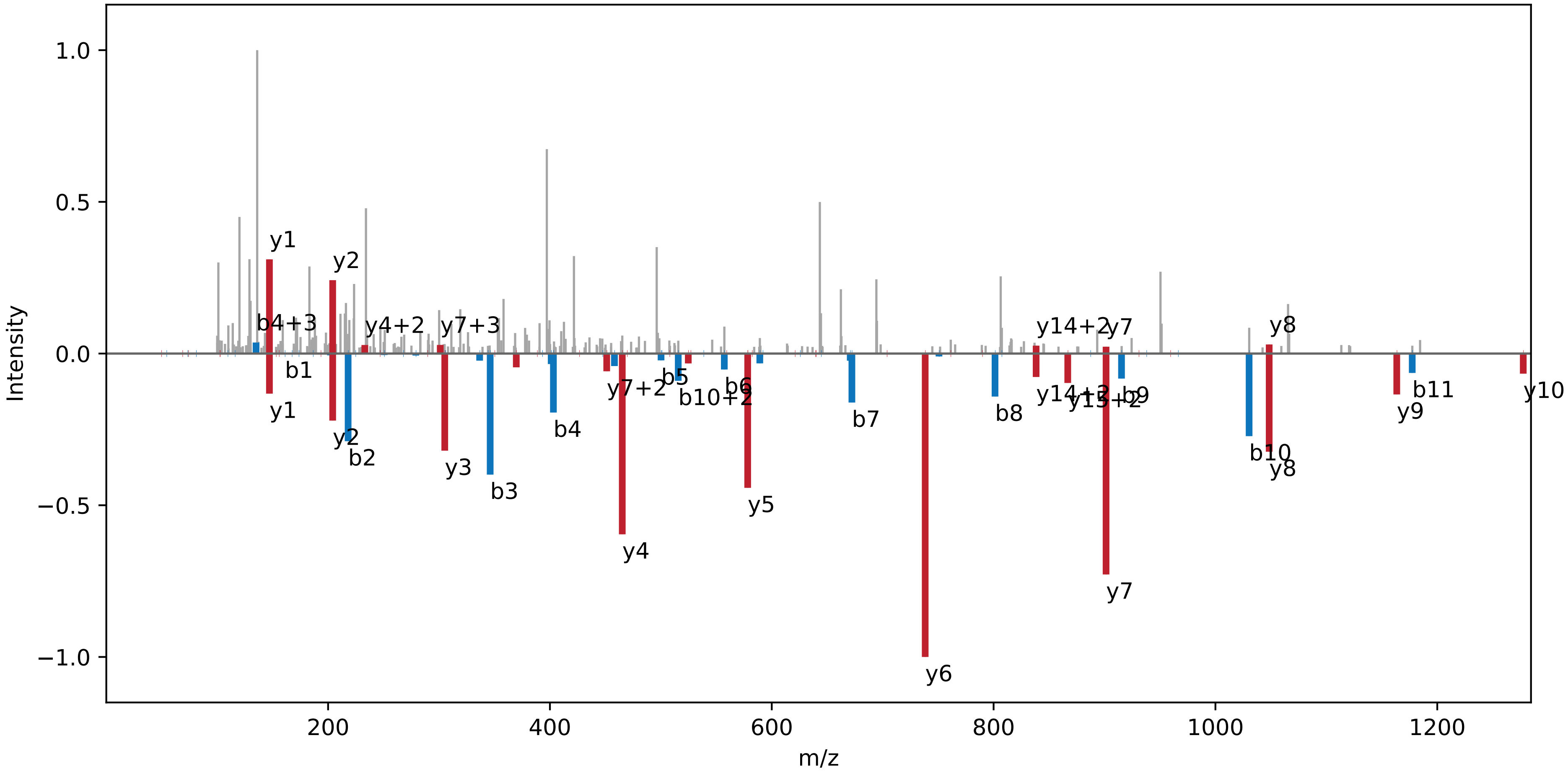

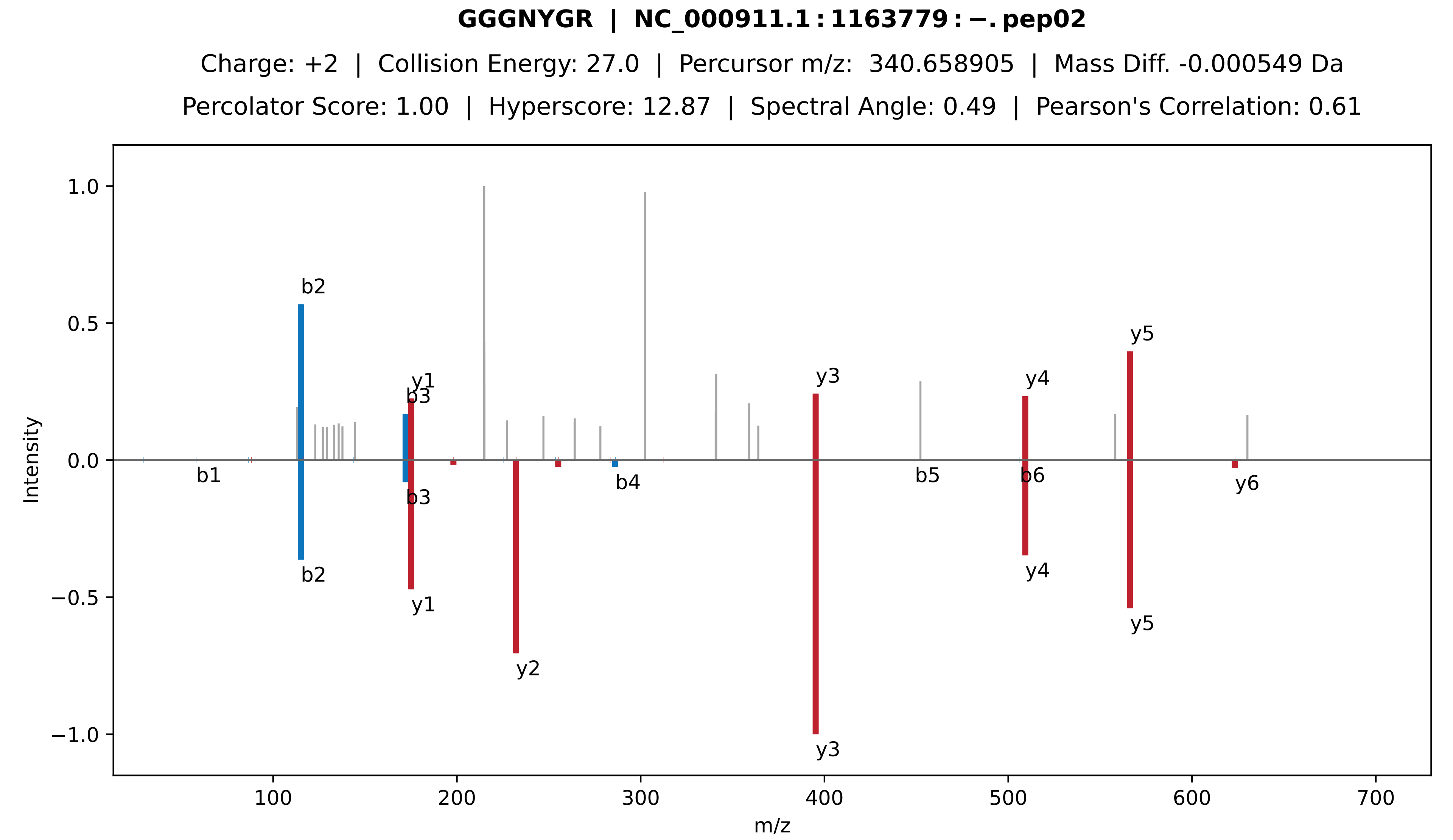
GGGNYGR | NC_000911.1:1163779:−.pep02
Charge: +2 | Collision Energy: 27.0 | Percursor m/z: 340.658905 | Mass Diff. -0.000549 Da
Percolator Score: 1.00 | Hyperscore: 12.87 | Spectral Angle: 0.49 | Pearson's Correlation: 0.61
Intensity
m/z
b1
b2
b3
b4
b5
b6
y1
y2
y3
y4
y5
y6
1.0
0.5
0.0
−0.5
−1.0
100
200
300
400
500
600
700

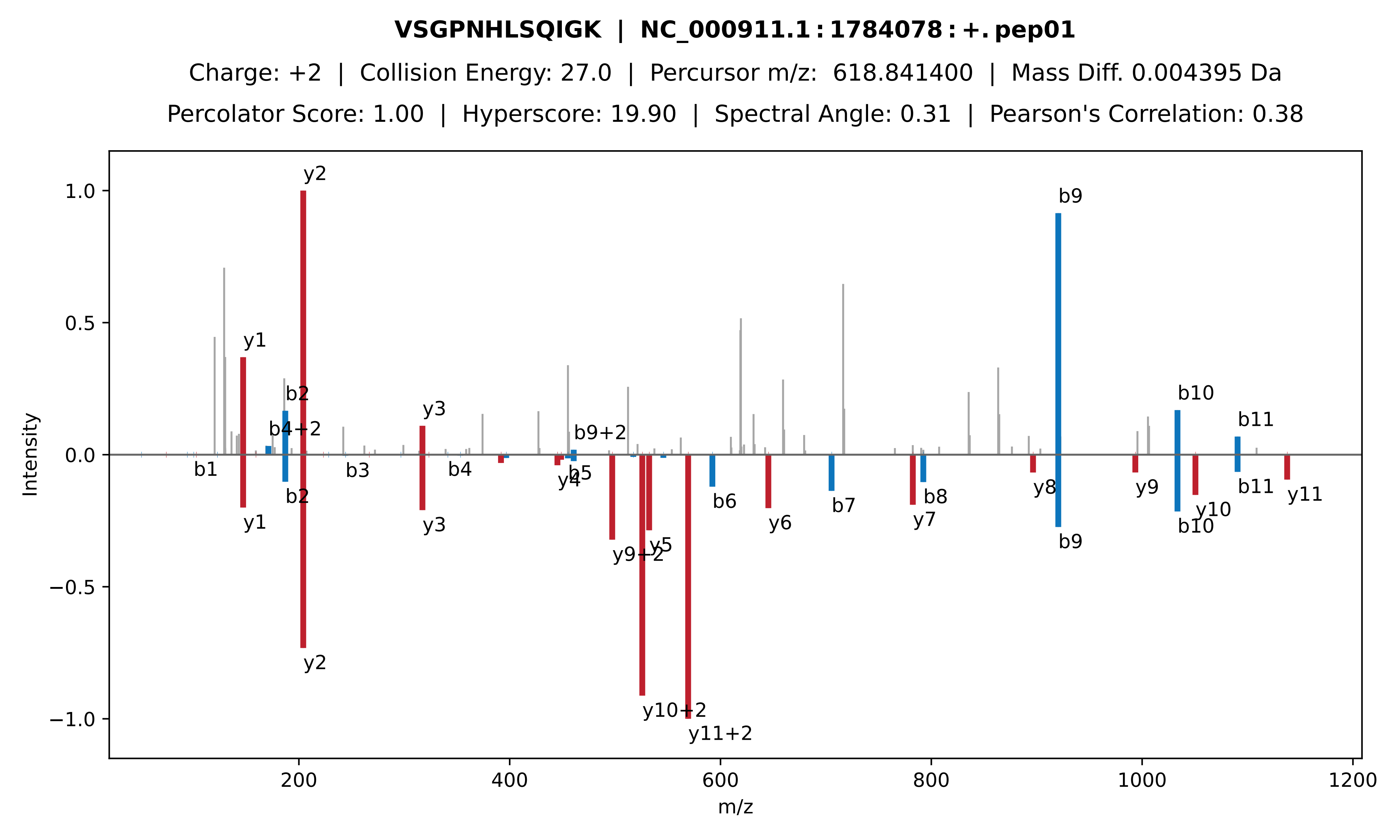
VSGPNHLSQIGK | NC_000911.1 : 1784078 : +. pep01
Charge: +2 | Collision Energy: 27.0 | Percursor m/z: 618.841400 | Mass Diff. 0.004395 Da
Percolator Score: 1.00 | Hyperscore: 19.90 | Spectral Angle: 0.31 | Pearson's Correlation: 0.38
Intensity
m/z
y2
b9
y1
b2
b4+2
y3
b9+2
b10
b11
b1
b3
b4
y4
b5
y9+2
y5
y10+2
y11+2
b6
y6
b7
y7
b8
y8
y9
y10
y11
1.0
0.5
0.0
−0.5
−1.0
200
400
600
800
1000
1200

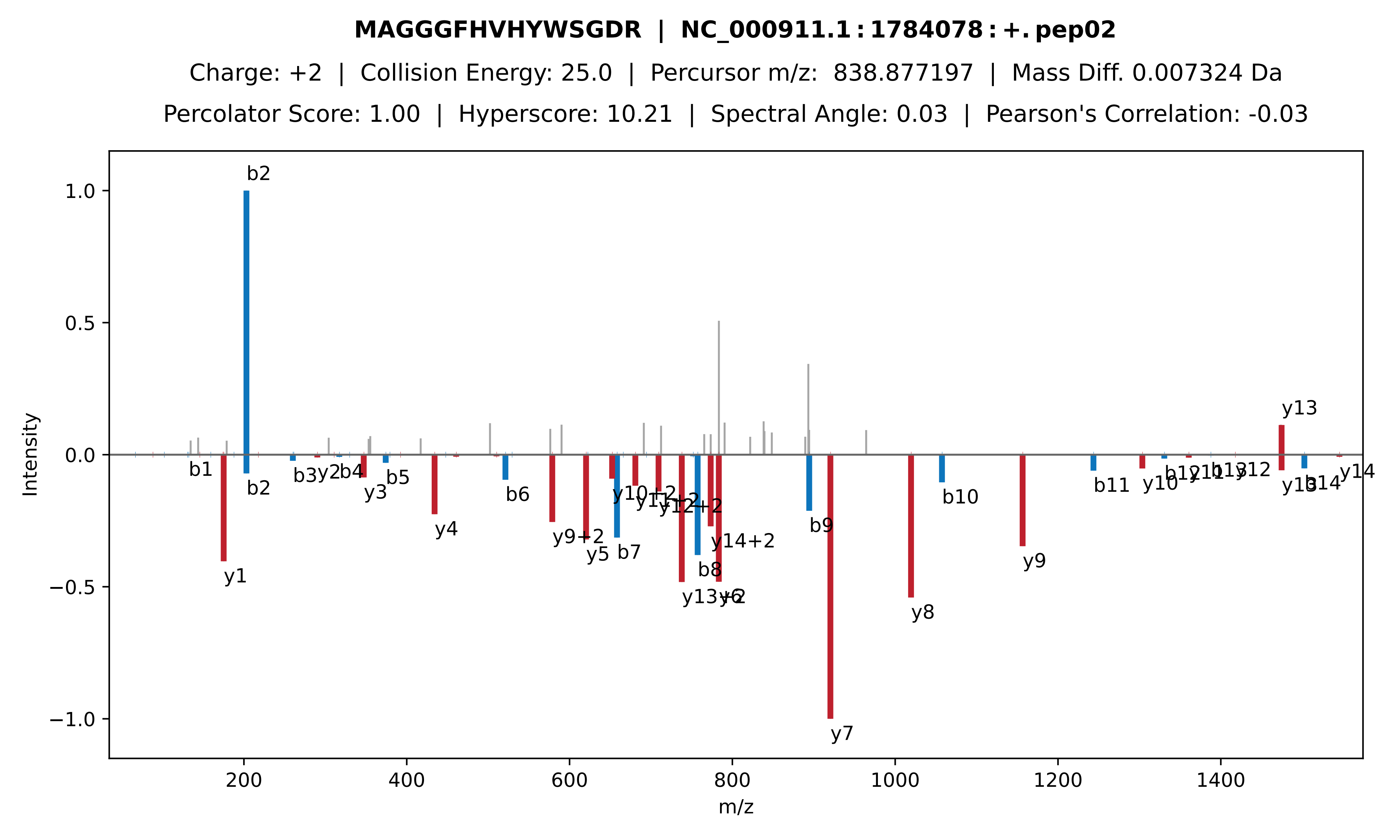

MAGGGFHVHYWSGDR | NC_000911.1 : 1784078 : +. pep02
Charge: +2 | Collision Energy: 25.0 | Percursor m/z: 838.877197 | Mass Diff. 0.007324 Da
Percolator Score: 1.00 | Hyperscore: 10.21 | Spectral Angle: 0.03 | Pearson's Correlation: -0.03
Intensity
m/z
b2
y13
b1
b3
y2
b4
y3
b5
y1
y4
b6
y9+2
y5
b7
y10+2
y11+2
y12+2
b8
y13+2
y6
y14+2
b9
y7
y8
b10
y9
b11
y10
b12
y11
b13
y12
y13
b14
y14

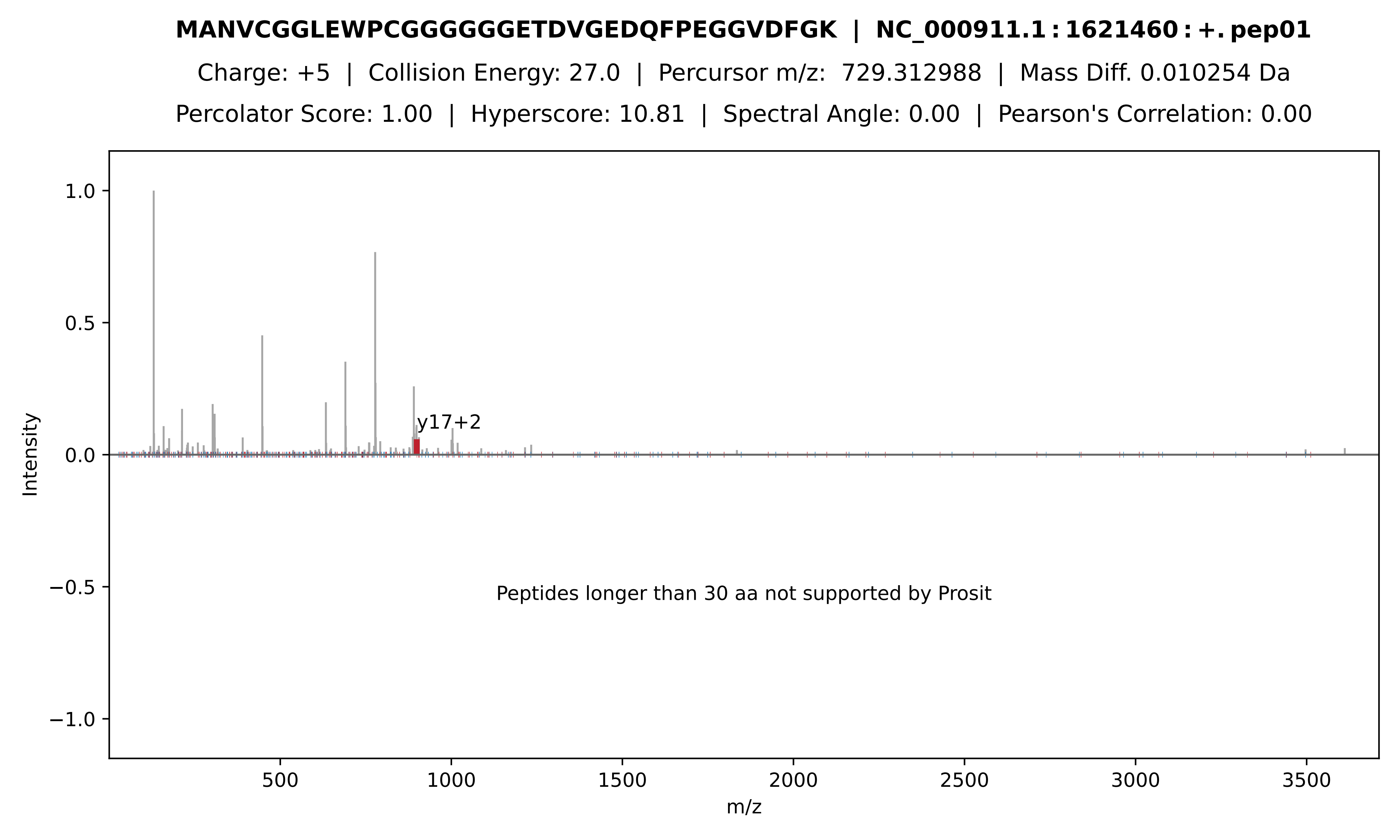

**MANVCGGLEWPCGGGGGGETDVGEDQFPEGGVDFGK | NC_000911.1 : 1621460 : +. pep01**
Charge: +5 | Collision Energy: 27.0 | Percursor m/z: 729.312988 | Mass Diff. 0.010254 Da
Percolator Score: 1.00 | Hyperscore: 10.81 | Spectral Angle: 0.00 | Pearson's Correlation: 0.00
y17+2
Peptides longer than 30 aa not supported by Prosit
Intensity
m/z
1.0
0.5
0.0
−0.5
−1.0
500
1000
1500
2000
2500
3000
3500

**c] − MANVCGGLEWPCGGGGGGETDVGEDQFPEGGVDFGK[TMT]VRGS[TMT]IFQFWD | NC_000911.1 : 1621460 :**

Charge: +5 | Collision Energy: 35.0 | Percursor m/z: 1076.299316 | Mass Diff. -1.011230 Da

Percolator Score: 1.00 | Hyperscore: 10.59 | Spectral Angle: 0.00 | Pearson's Correlation: 0.00

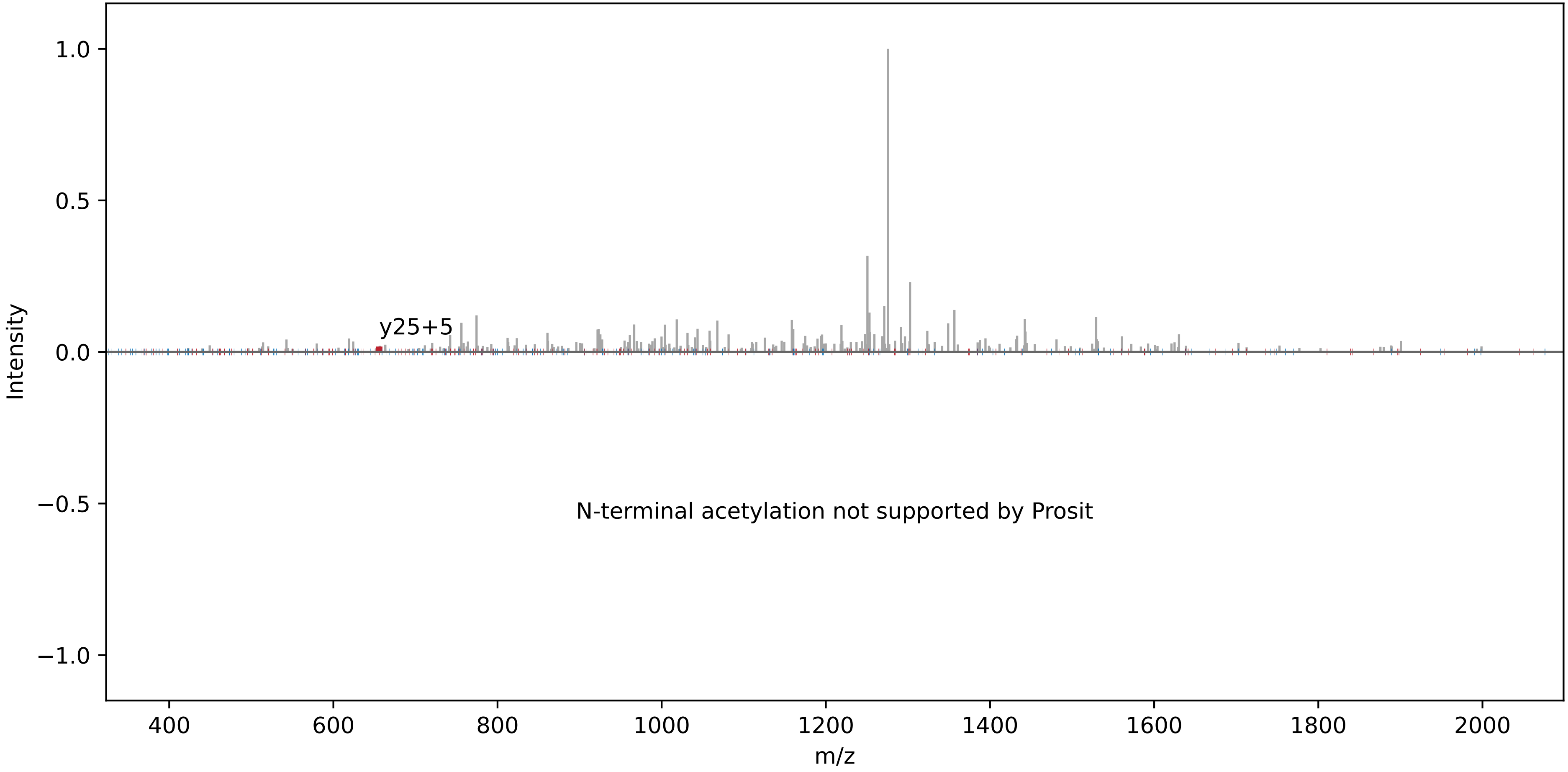

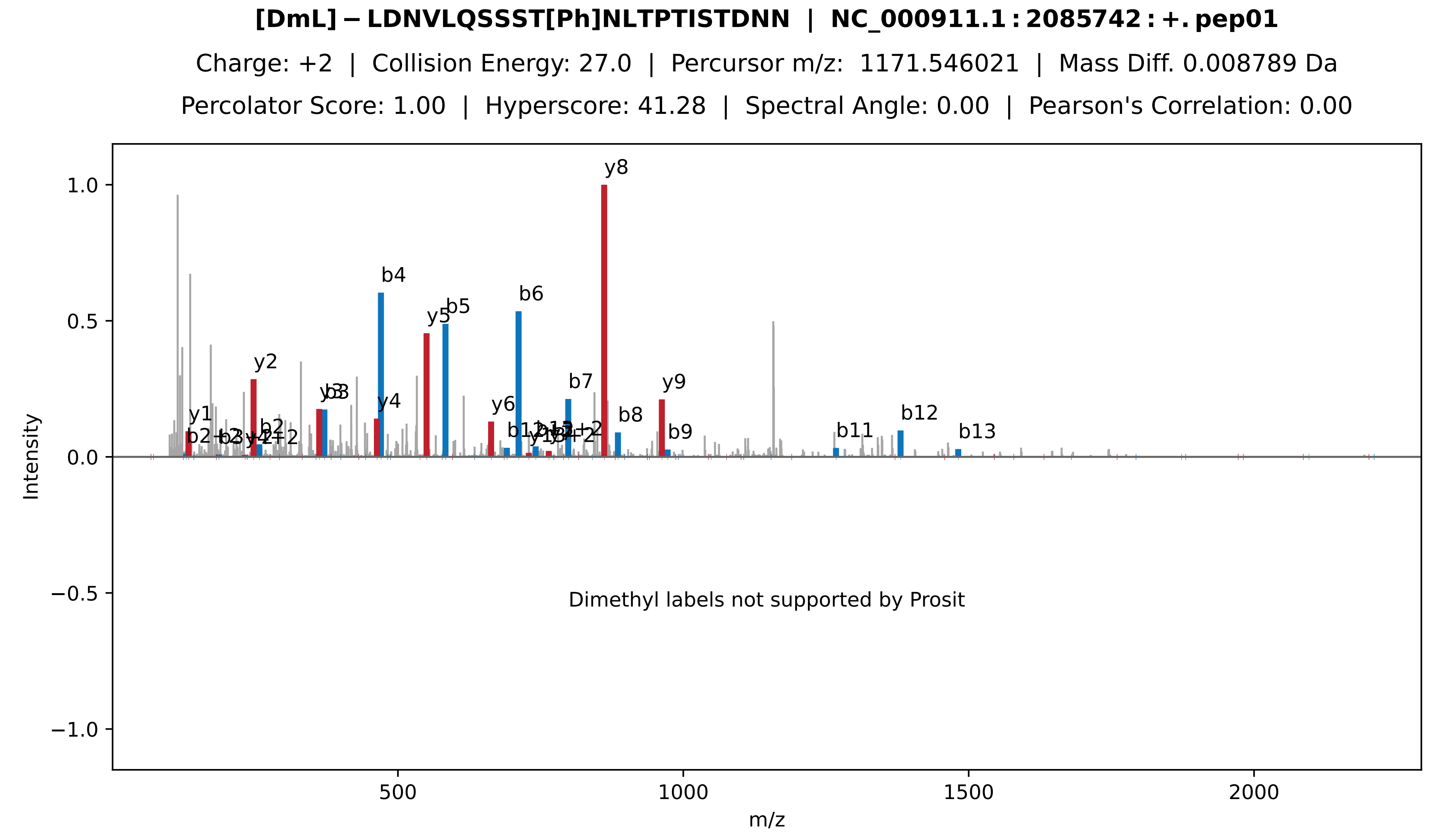
[DmL]−LDNVLQSSST[Ph]NLTPTISTDNN | NC_000911.1:2085742:+.pep01
Charge: +2 | Collision Energy: 27.0 | Percursor m/z: 1171.546021 | Mass Diff. 0.008789 Da
Percolator Score: 1.00 | Hyperscore: 41.28 | Spectral Angle: 0.00 | Pearson's Correlation: 0.00
Intensity
m/z
y8
b4
b6
b5
y5
y2
b3
y4
b7
y9
y6
b8
b9
b11
b12
b13
y1
b2
Dimethyl labels not supported by Prosit

**[DmH] − LWSWINSYGGLSDLK[DmL]K[DmH] | NC_000911.1 : 2085742 : +. pep02**

Charge: +3 | Collision Energy: 27.0 | Percursor m/z: 656.385000 | Mass Diff. -0.016968 Da

Percolator Score: 1.00 | Hyperscore: 38.65 | Spectral Angle: 0.00 | Pearson's Correlation: 0.00

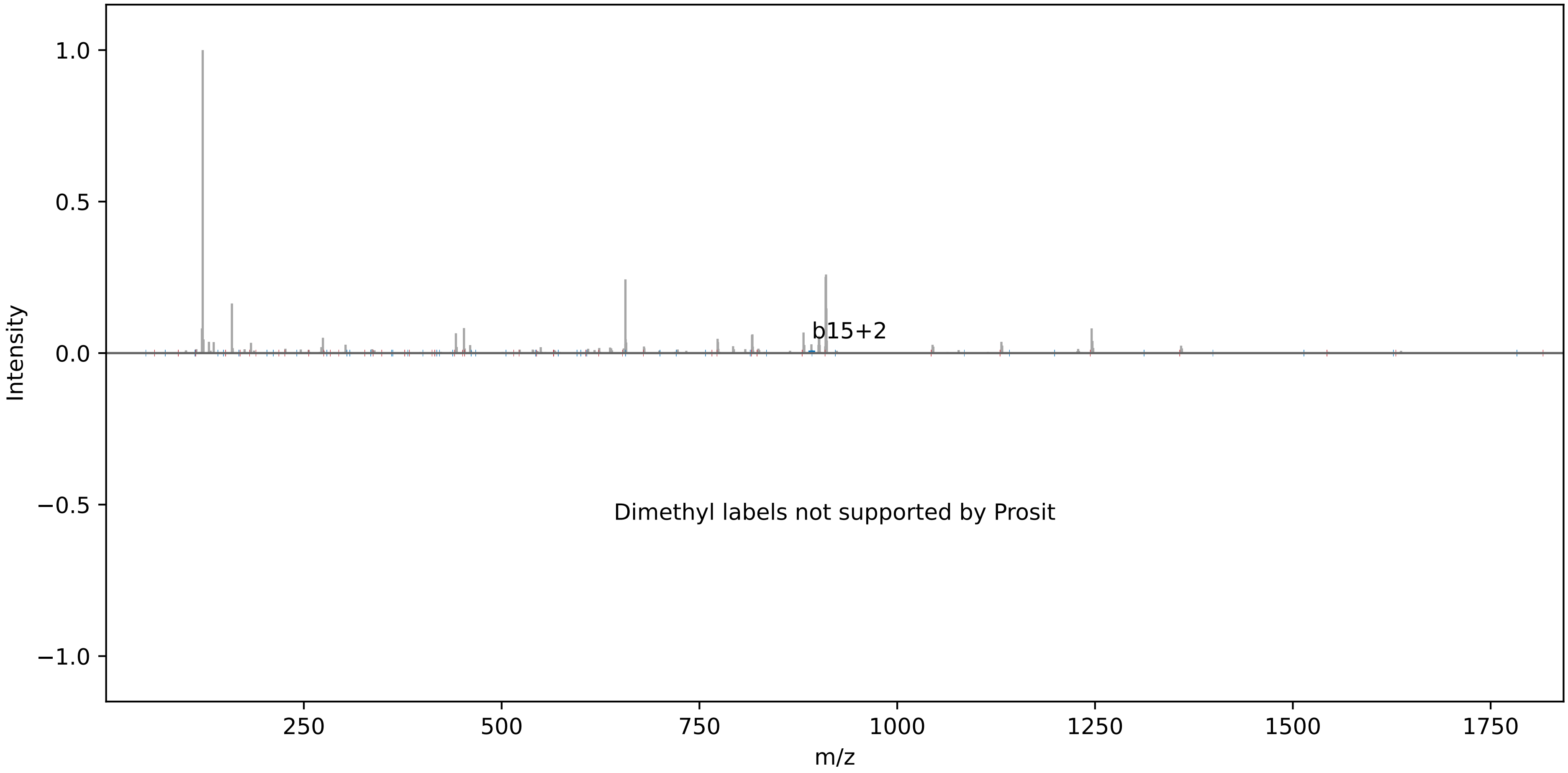

**MFTATIFQLLSCM[Ox]VHIFYSQQCRCTLDAVGIASCRCGVVAGLLCSK | NC_000911.1:1575411:−.pep01**

Charge: +5 | Collision Energy: 27.0 | Percursor m/z: 1070.312012 | Mass Diff. 1.993652 Da

Percolator Score: 1.00 | Hyperscore: 15.32 | Spectral Angle: 0.00 | Pearson's Correlation: 0.00

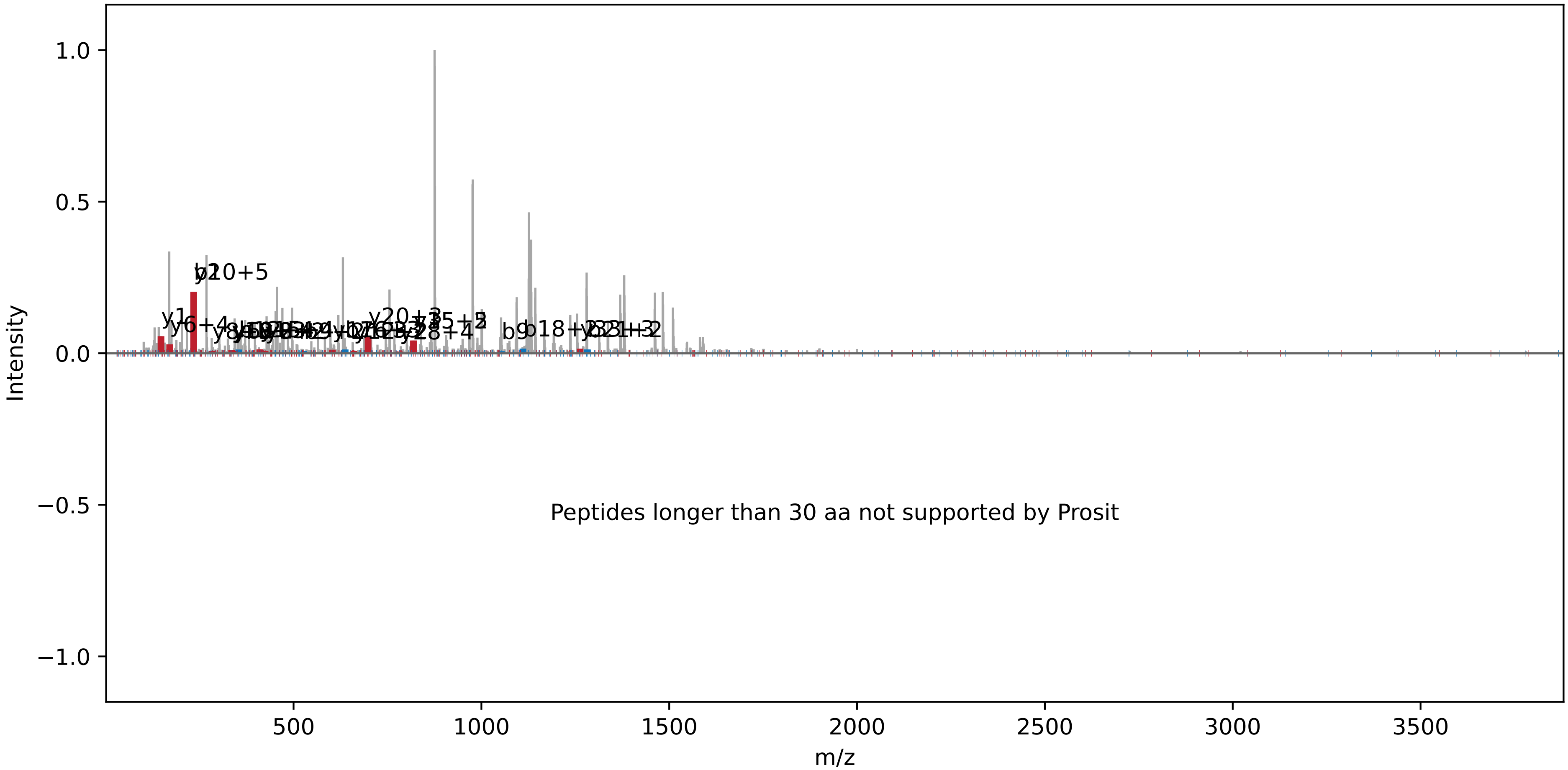

**[Ac] − FTATIFQLLSCMVHIFYSQQCRCTLDAVGIASCRCGVVAGLLCSK | NC_000911.1 : 1575411 : −. pep02**

Charge: +4 | Collision Energy: 27.0 | Percursor m/z: 1311.396362 | Mass Diff. 2.035645 Da

Percolator Score: 0.99 | Hyperscore: 14.70 | Spectral Angle: 0.00 | Pearson's Correlation: 0.00

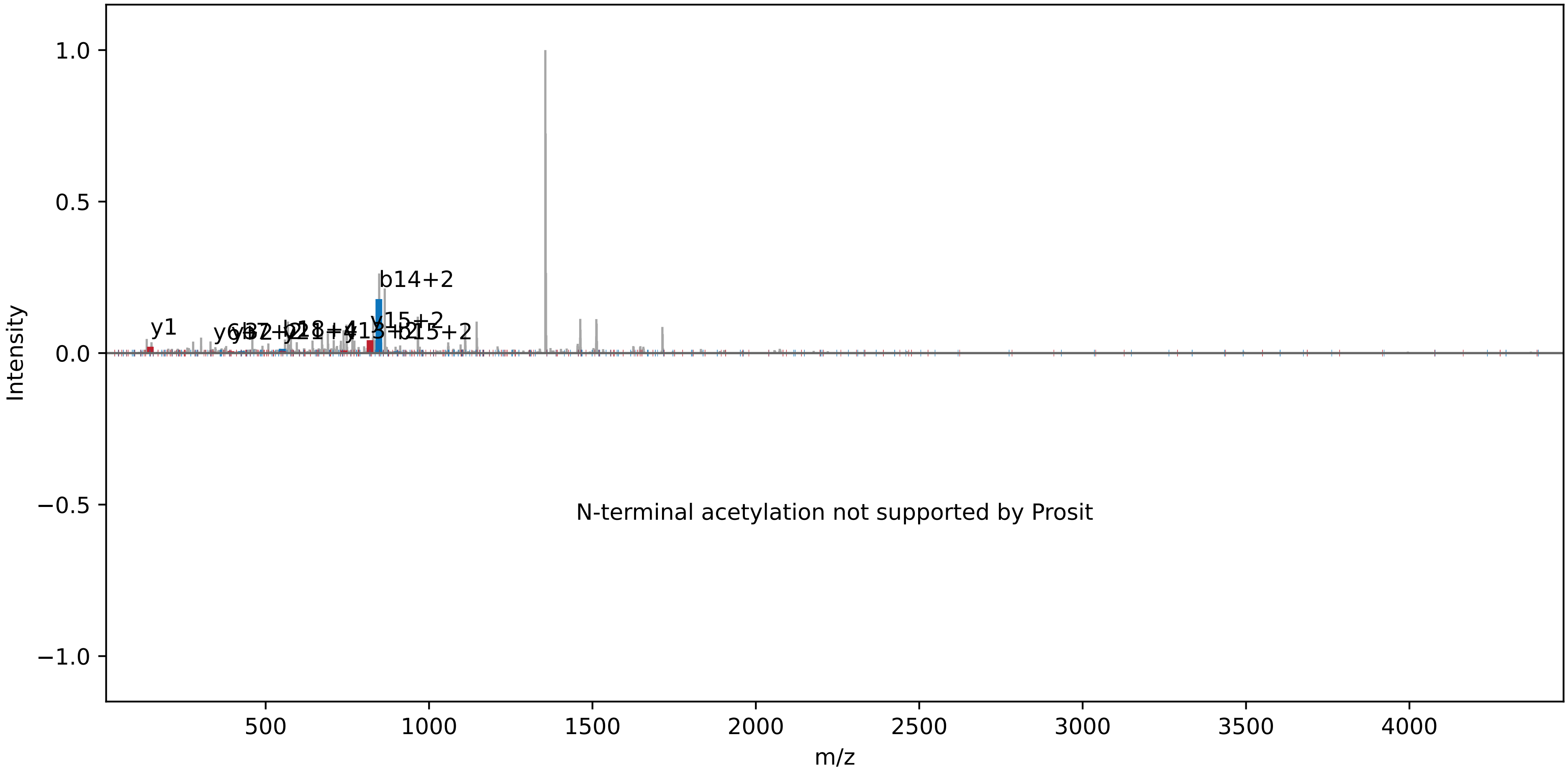

**M[Ox]VIPWPDAYPCCCKCPSCLPYLPPCGDLPFPILITPLTCKYYPR | NC_000911.1 : 1828167 : +. pep01**

Charge: +4 | Collision Energy: 27.0 | Percursor m/z: 1351.387207 | Mass Diff. 0.004883 Da

Percolator Score: 0.99 | Hyperscore: 10.23 | Spectral Angle: 0.00 | Pearson's Correlation: 0.00

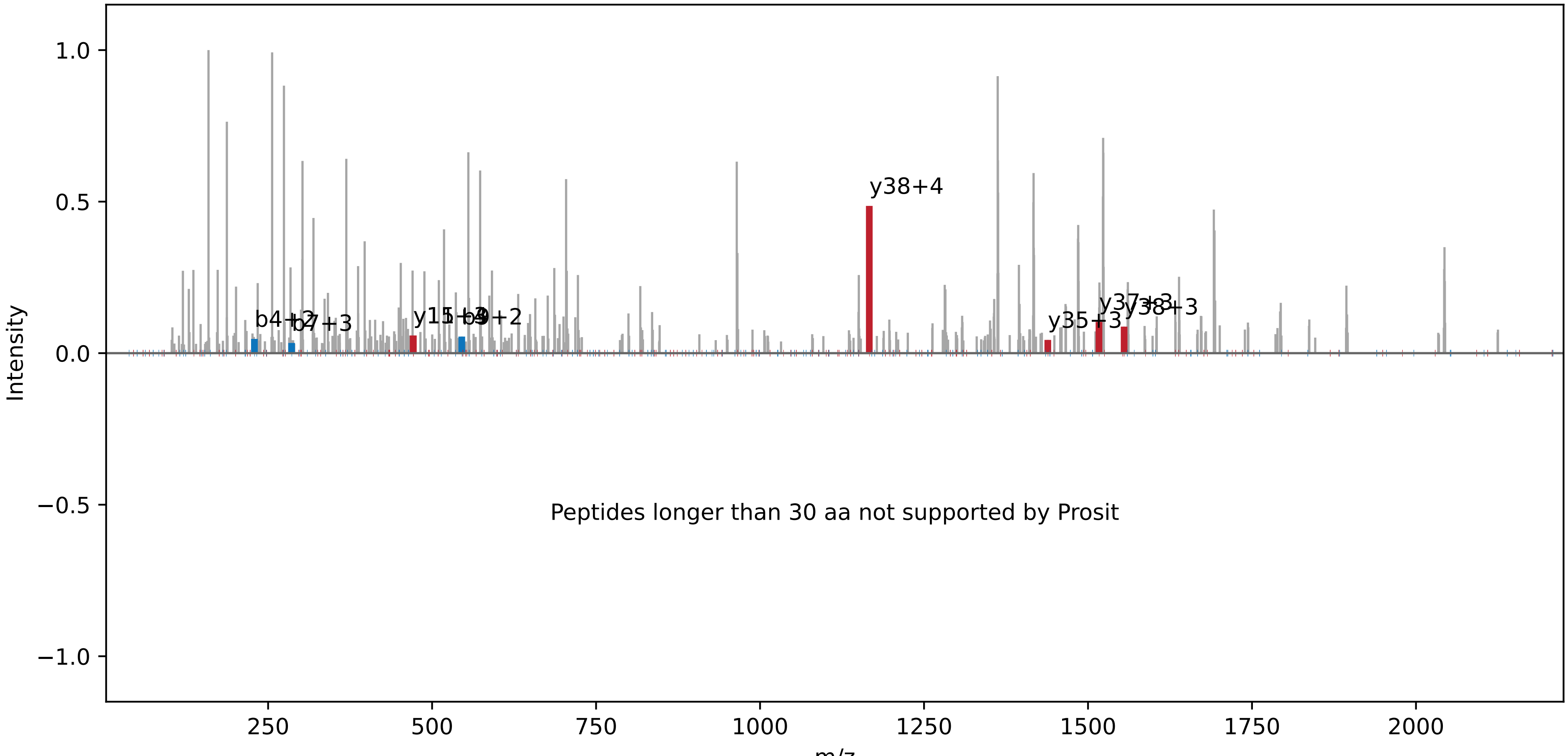

**VIPWPDAYPCCCKCPSCLPYLPPCGDLPFPILITPLTCKYYPR | NC_000911.1:1828167:+.pep02**

Charge: +5 | Collision Energy: 27.0 | Percursor m/z: 1051.913086 | Mass Diff. 0.040527 Da

Percolator Score: 1.00 | Hyperscore: 15.16 | Spectral Angle: 0.00 | Pearson's Correlation: 0.00

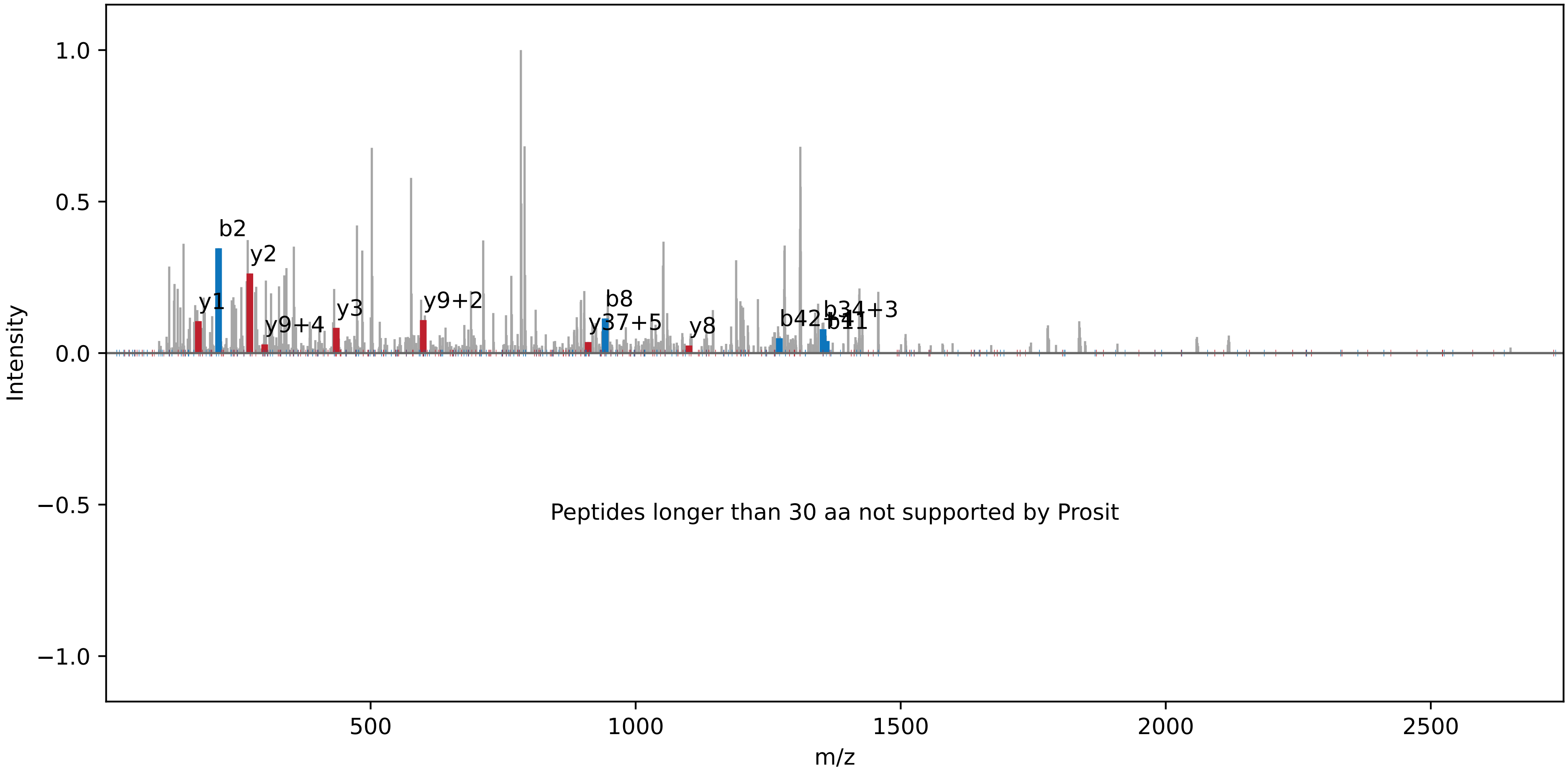

**SSLVTIFR | NC_005229.1 : 98992 : −. pep01**

Charge: +2 | Collision Energy: 27.0 | Percursor m/z: 461.772888 | Mass Diff. -0.000305 Da

Percolator Score: 1.00 | Hyperscore: 18.40 | Spectral Angle: 0.45 | Pearson's Correlation: 0.59

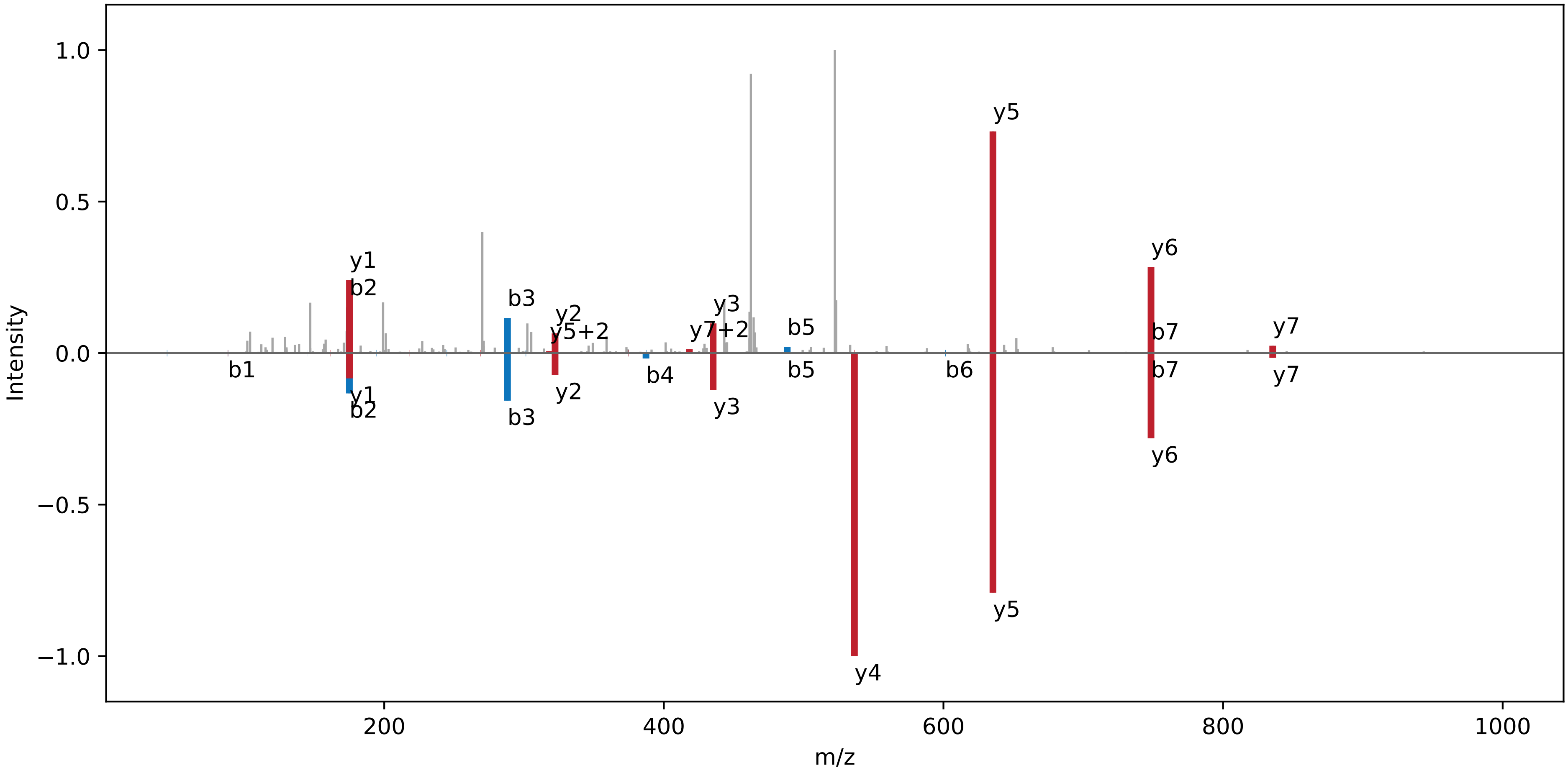

**PIQGAGELIR | NC_000911.1 : 2737711 : +. pep01**

Charge: +2 | Collision Energy: 27.0 | Percursor m/z: 527.305908 | Mass Diff. -0.000732 Da

Percolator Score: 1.00 | Hyperscore: 22.34 | Spectral Angle: 0.18 | Pearson's Correlation: 0.15

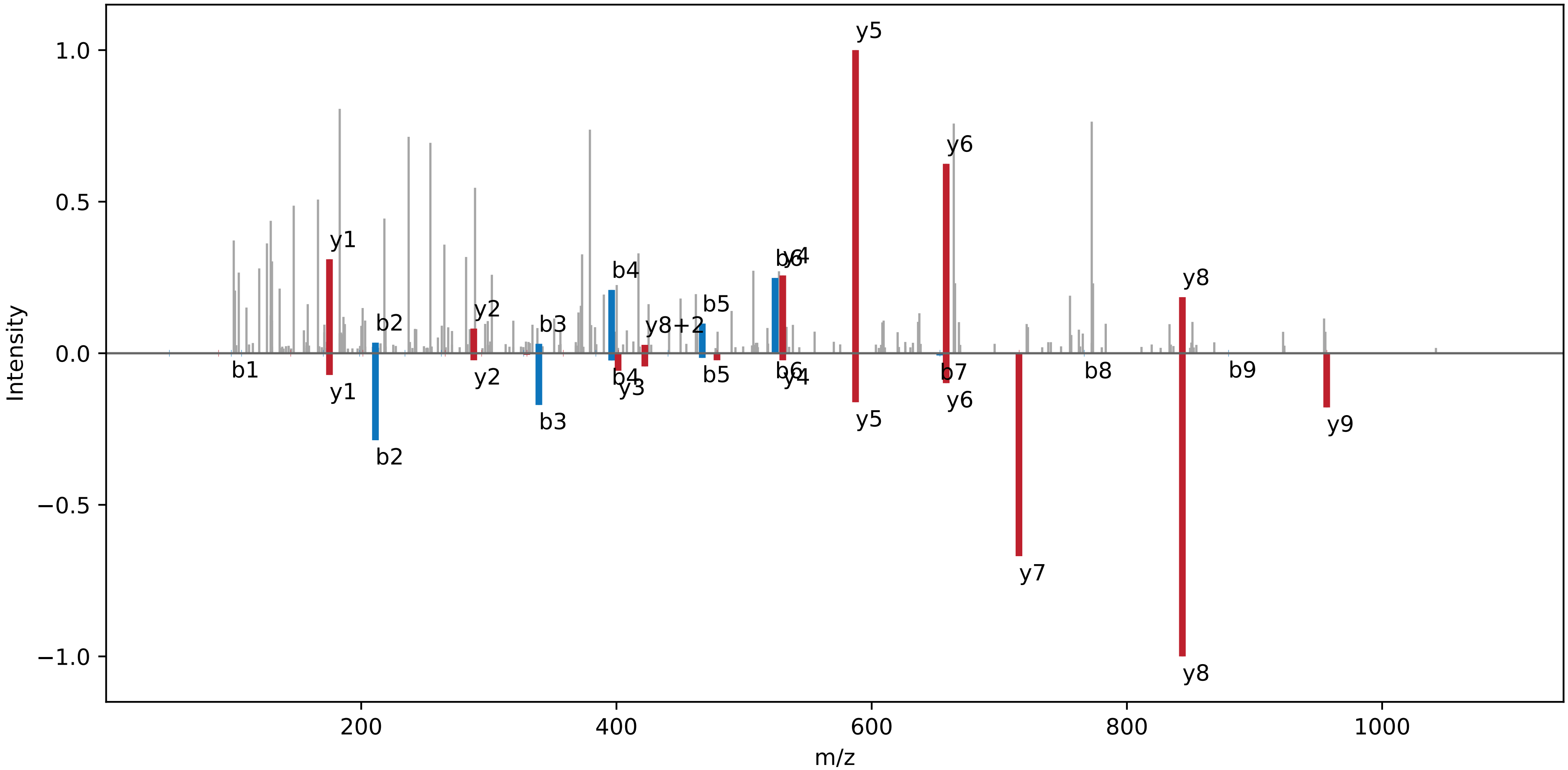

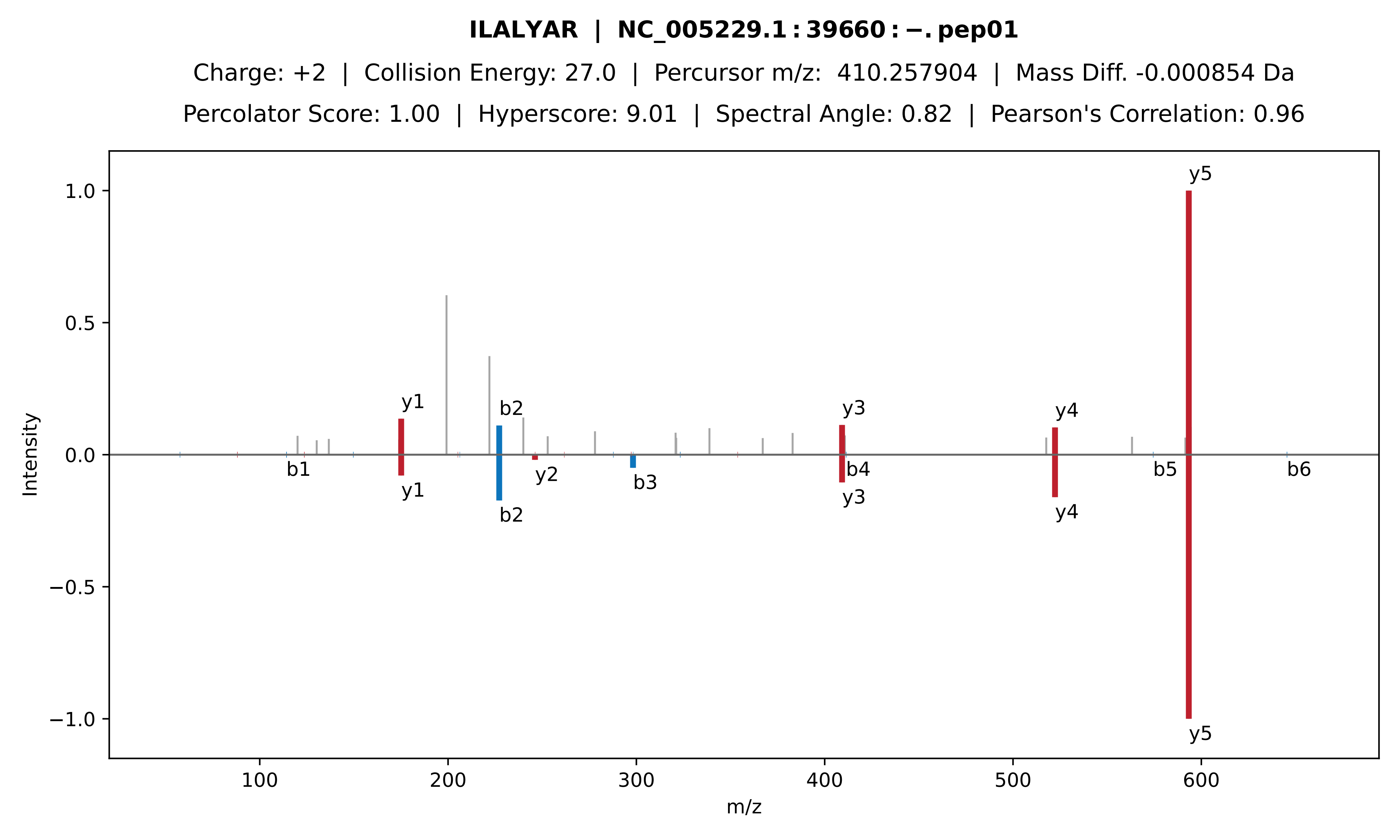

**ILALYAR | NC_005229.1 : 39660 : −. pep01**
Charge: +2 | Collision Energy: 27.0 | Percursor m/z: 410.257904 | Mass Diff. -0.000854 Da
Percolator Score: 1.00 | Hyperscore: 9.01 | Spectral Angle: 0.82 | Pearson's Correlation: 0.96
Intensity
m/z
y1
b2
y3
y4
y5
b1
y2
b3
b4
b5
b6

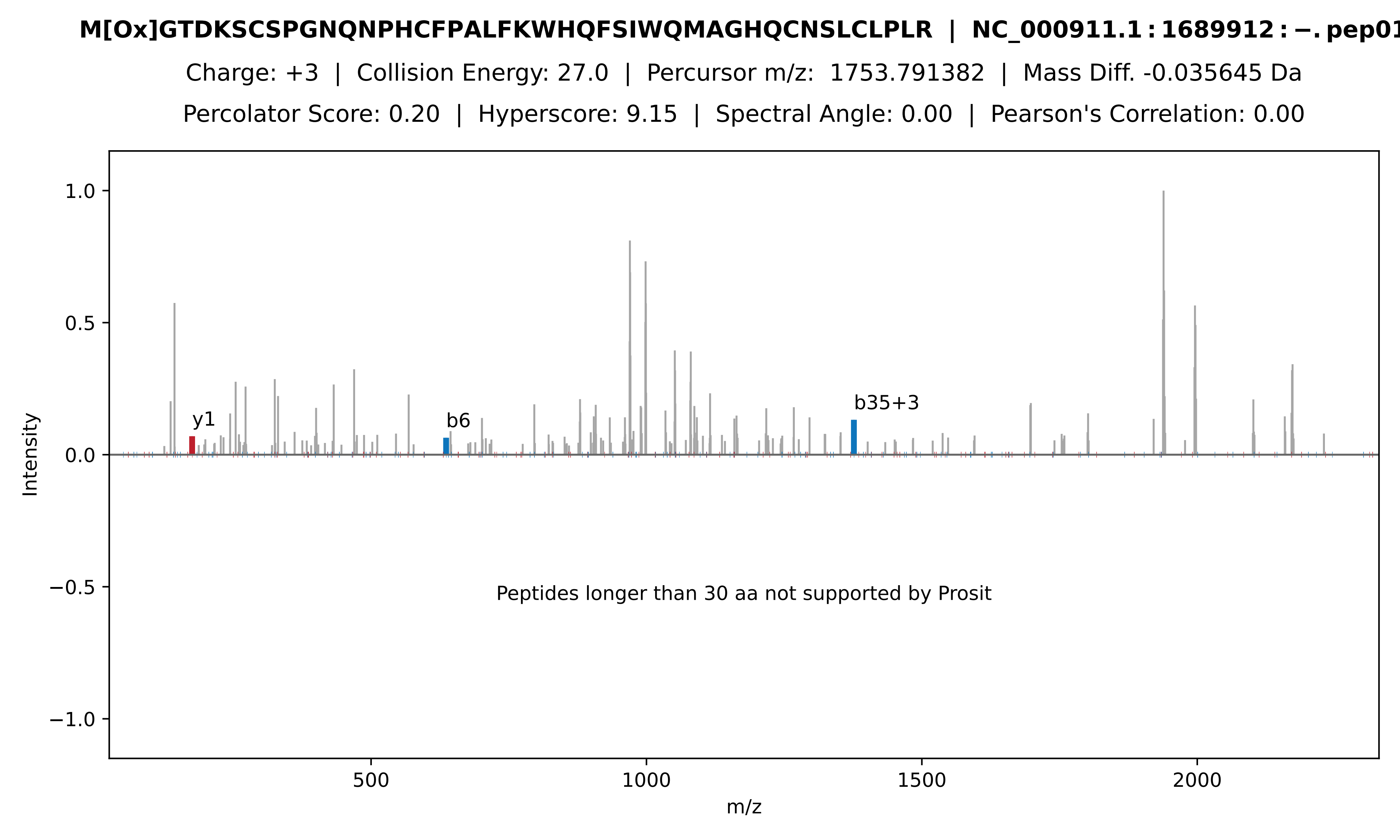
M[Ox]GTDKSCSPGNQNPHCFPALFKWHQFSIWQMAGHQCNSLCLPLR | NC_000911.1:1689912:−.pep01
Charge: +3 | Collision Energy: 27.0 | Percursor m/z: 1753.791382 | Mass Diff. -0.035645 Da
Percolator Score: 0.20 | Hyperscore: 9.15 | Spectral Angle: 0.00 | Pearson's Correlation: 0.00
Intensity
1.0
0.5
0.0
−0.5
−1.0
y1
b6
b35+3
Peptides longer than 30 aa not supported by Prosit
500
1000
1500
2000
m/z

**EAGLVLR | NC_005230.1 : 40013 : −. pep01**

Charge: +2 | Collision Energy: 27.0 | Percursor m/z: 379.231476 | Mass Diff. -0.000671 Da

Percolator Score: 0.99 | Hyperscore: 12.78 | Spectral Angle: 0.56 | Pearson's Correlation: 0.72

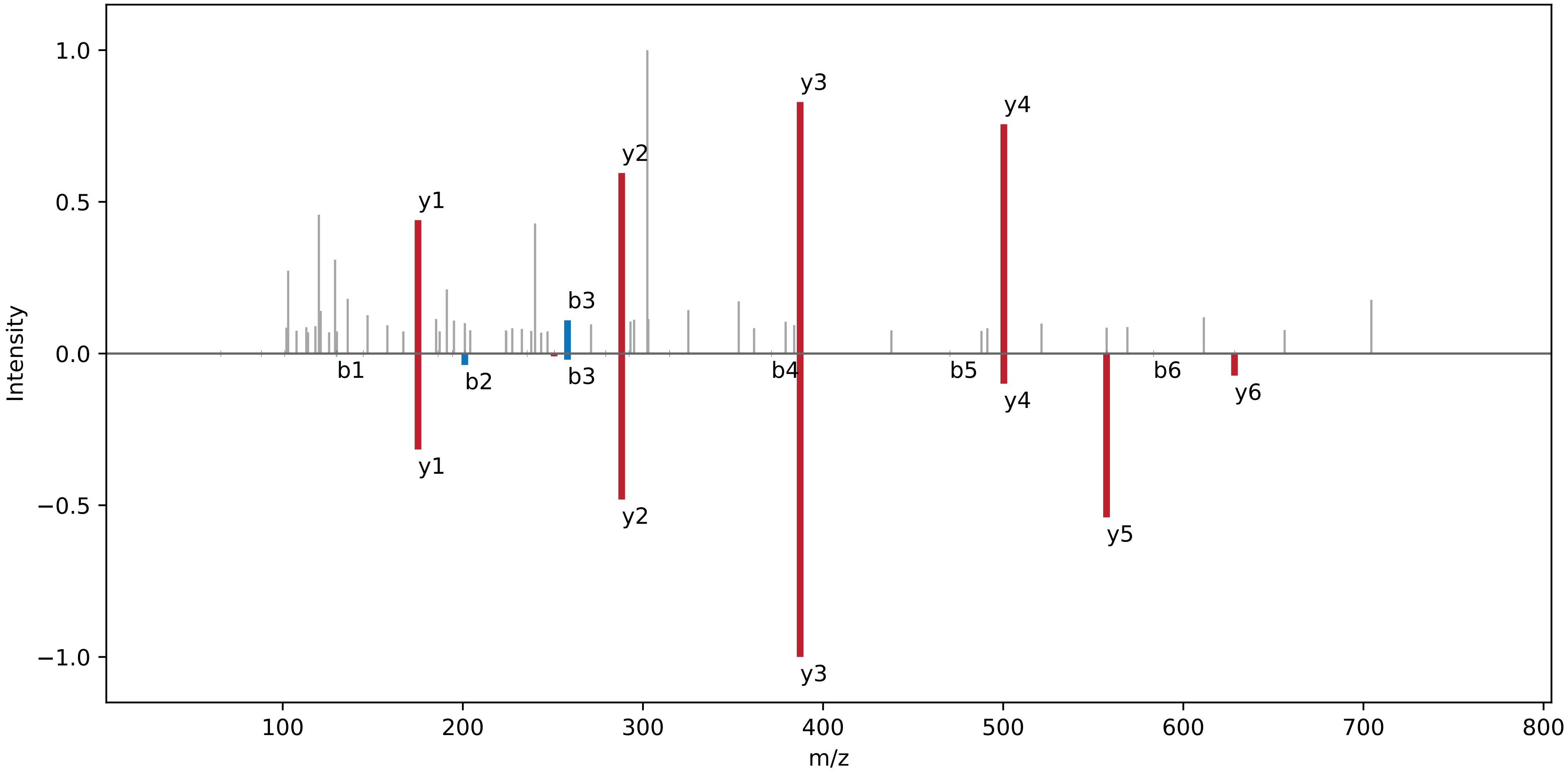

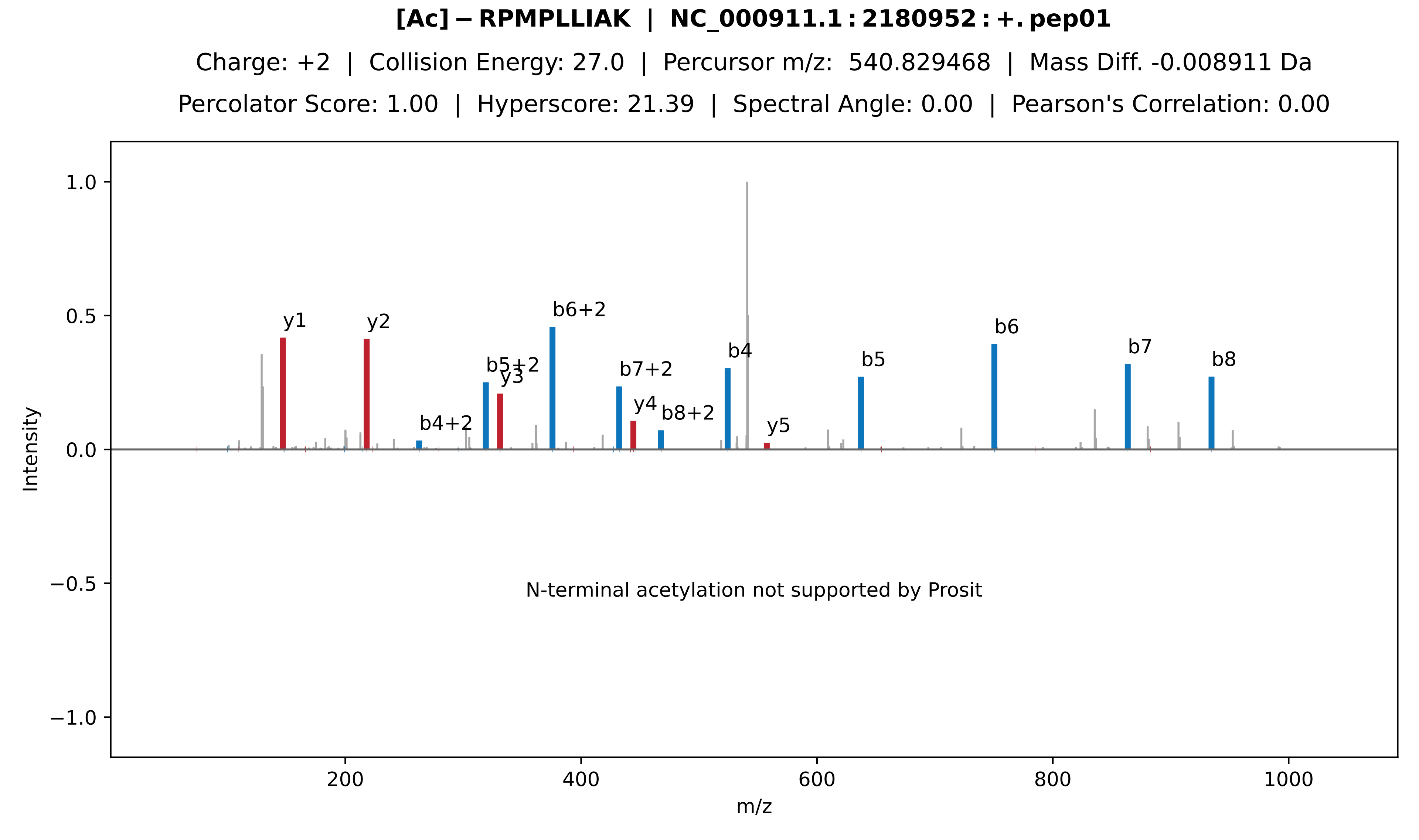
[Ac]−RPMPLLIAK | NC_000911.1:2180952:+.pep01
Charge: +2 | Collision Energy: 27.0 | Percursor m/z: 540.829468 | Mass Diff. -0.008911 Da
Percolator Score: 1.00 | Hyperscore: 21.39 | Spectral Angle: 0.00 | Pearson's Correlation: 0.00
y1
y2
b4+2
b5+2
y3
b6+2
b7+2
y4
b8+2
b4
y5
b5
b6
b7
b8
N-terminal acetylation not supported by Prosit
Intensity
m/z
1.0
0.5
0.0
−0.5
−1.0
200
400
600
800
1000

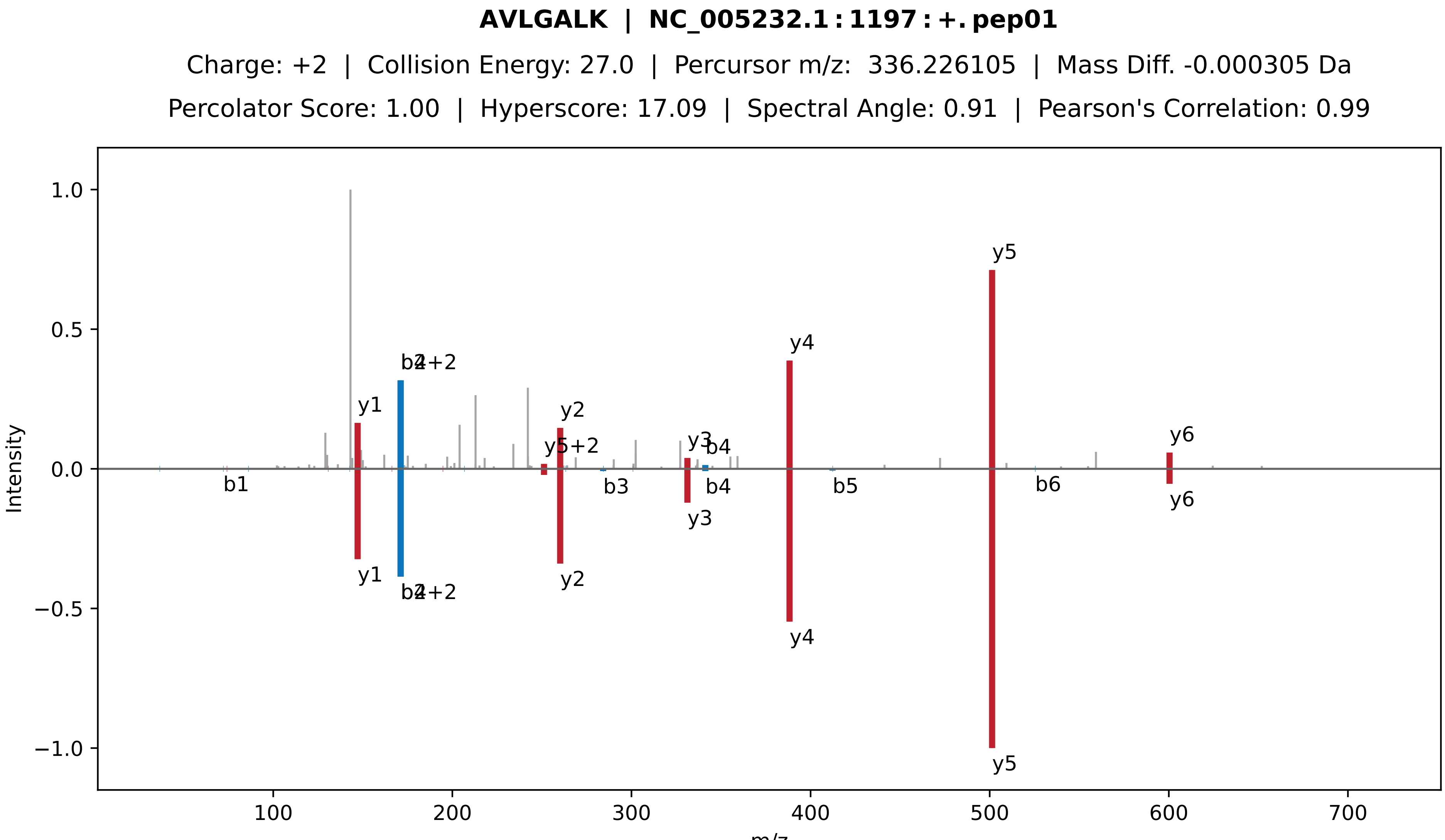
AVLGALK | NC_005232.1:1197:+.pep01
Charge: +2 | Collision Energy: 27.0 | Percursor m/z: 336.226105 | Mass Diff. -0.000305 Da
Percolator Score: 1.00 | Hyperscore: 17.09 | Spectral Angle: 0.91 | Pearson's Correlation: 0.99
1.0
0.5
0.0
−0.5
−1.0
Intensity
100
200
300
400
500
600
700
m/z
y1
b2
b4+2
y2
y5+2
y3
b4
y4
y5
y6
b1
b3
b5
b6

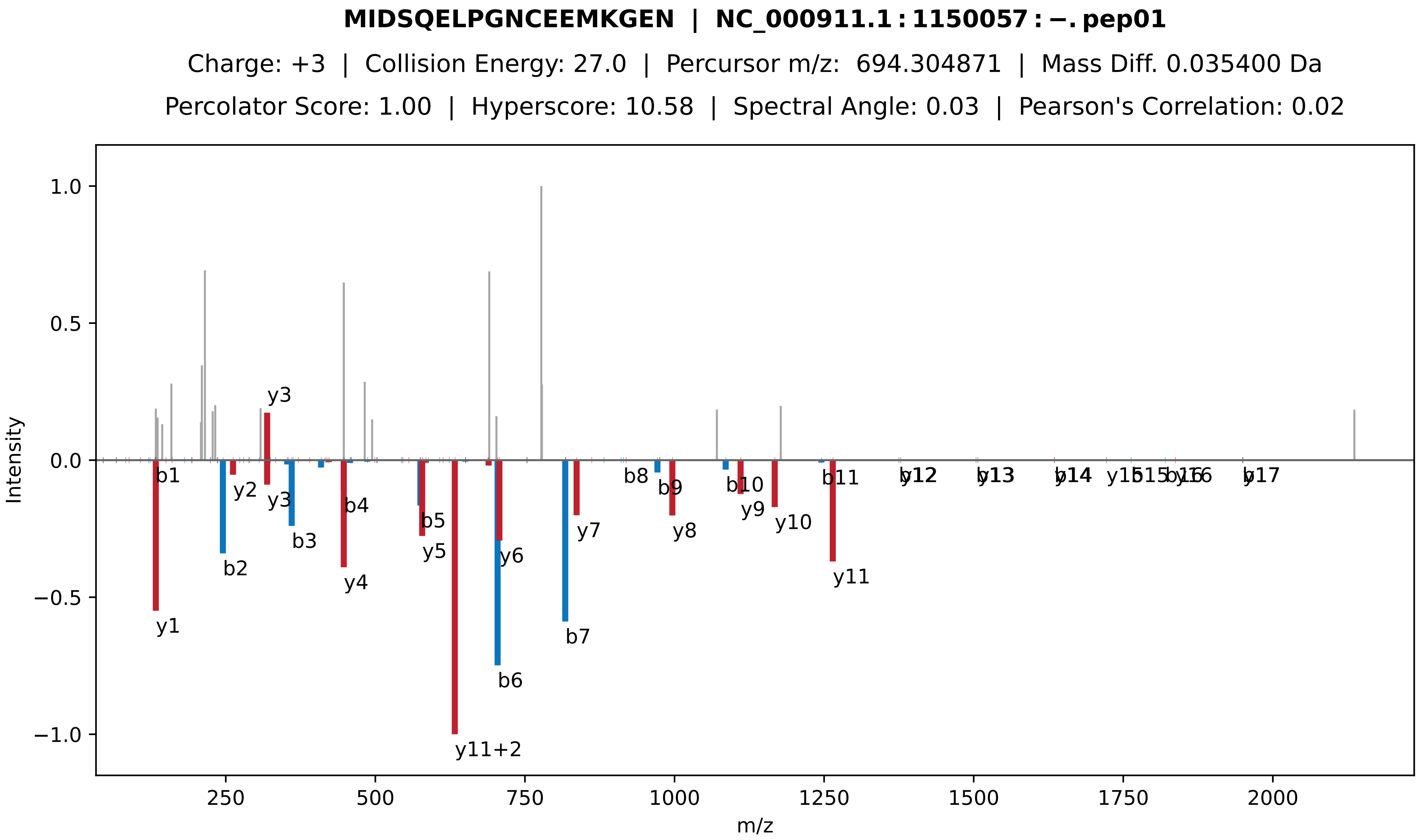

MIDSQELPGNCEEMKGEN | NC_000911.1 : 1150057 : −. pep01
Charge: +3 | Collision Energy: 27.0 | Percursor m/z: 694.304871 | Mass Diff. 0.035400 Da
Percolator Score: 1.00 | Hyperscore: 10.58 | Spectral Angle: 0.03 | Pearson's Correlation: 0.02
Intensity
m/z
1.0
0.5
0.0
−0.5
−1.0
250
500
750
1000
1250
1500
1750
2000
y3
b1
y2
y3
b4
b5
y5
y6
b2
b3
y4
y1
y11+2
b6
b7
y7
b8
b9
y8
b10
y9
y10
b11
y11
b12
y12
b13
y13
b14
y14
y15
b15
b16
y16
b17
y17

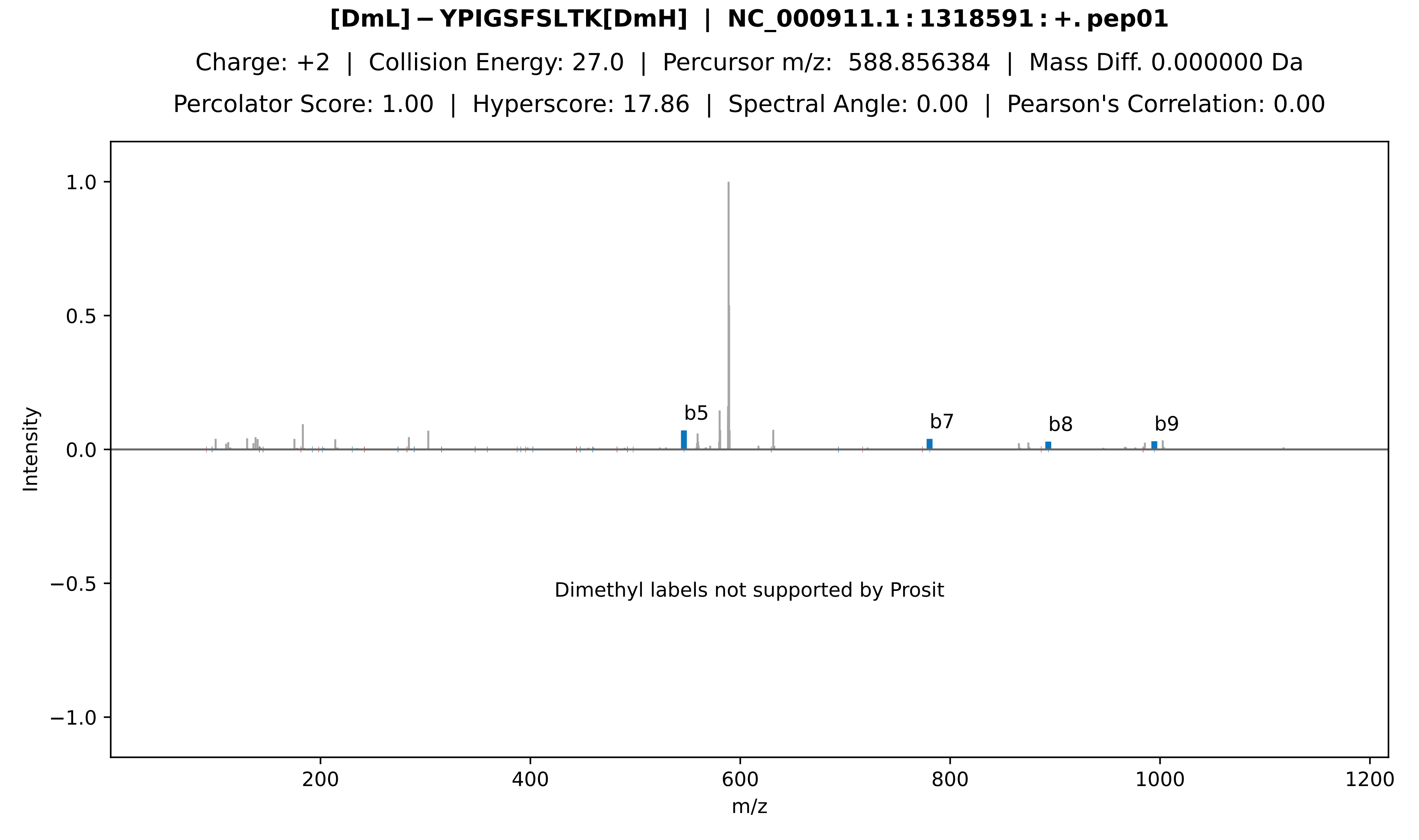
[DmL]−YPIGSFSLTK[DmH] | NC_000911.1:1318591:+.pep01
Charge: +2 | Collision Energy: 27.0 | Percursor m/z: 588.856384 | Mass Diff. 0.000000 Da
Percolator Score: 1.00 | Hyperscore: 17.86 | Spectral Angle: 0.00 | Pearson's Correlation: 0.00
b5
b7
b8
b9
Dimethyl labels not supported by Prosit
Intensity
m/z
1.0
0.5
0.0
−0.5
−1.0
200
400
600
800
1000
1200

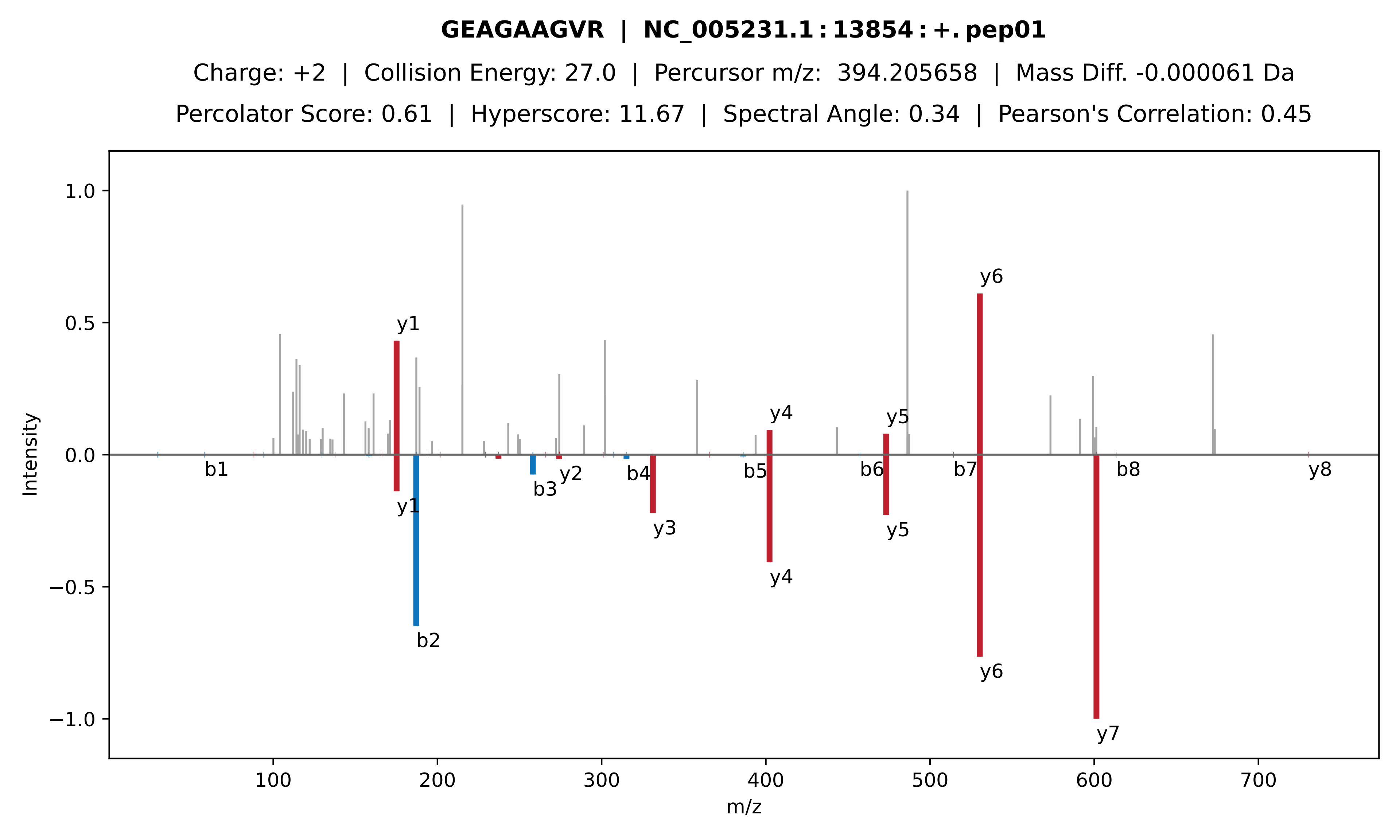
GEAGAAGVR | NC_005231.1 : 13854 : +. pep01
Charge: +2 | Collision Energy: 27.0 | Percursor m/z: 394.205658 | Mass Diff. -0.000061 Da
Percolator Score: 0.61 | Hyperscore: 11.67 | Spectral Angle: 0.34 | Pearson's Correlation: 0.45
Intensity
m/z
1.0
0.5
0.0
−0.5
−1.0
100
200
300
400
500
600
700
y1
y4
y5
y6
b1
b2
b3
b4
b5
b6
b7
b8
y1
y2
y3
y4
y5
y6
y7
y8

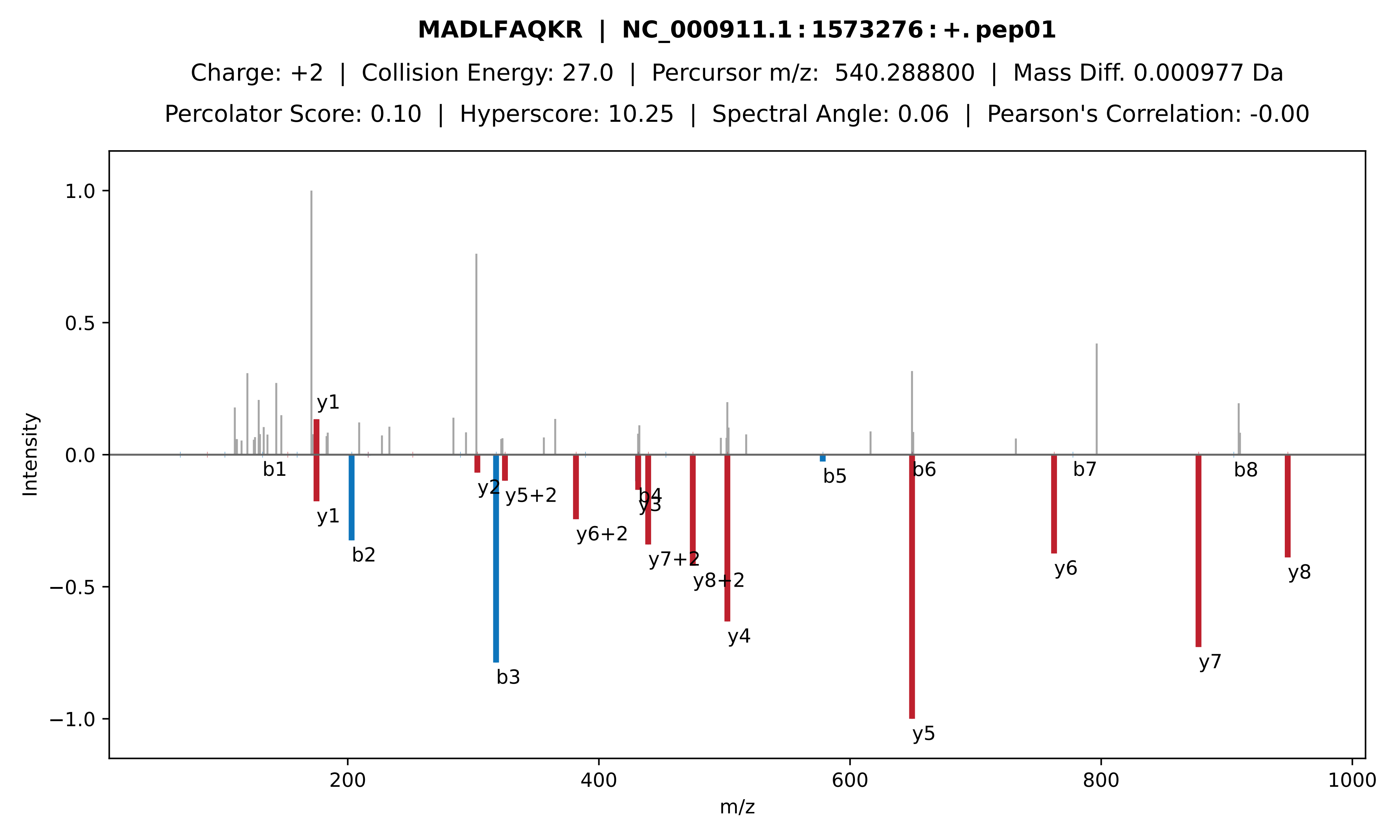
MADLFAQKR | NC_000911.1:1573276:+.pep01
Charge: +2 | Collision Energy: 27.0 | Percursor m/z: 540.288800 | Mass Diff. 0.000977 Da
Percolator Score: 0.10 | Hyperscore: 10.25 | Spectral Angle: 0.06 | Pearson's Correlation: -0.00
Intensity
m/z
1.0
0.5
0.0
−0.5
−1.0
200
400
600
800
1000
y1
b1
y1
b2
y2
b3
y5+2
y6+2
b4
y3
y7+2
y8+2
y4
b5
b6
y5
y6
b7
y7
b8
y8

**[DmM] − SAIGSGD | NC_000911.1 : 1785226 : +. pep01**

Charge: +2 | Collision Energy: 28.0 | Percursor m/z: 319.669006 | Mass Diff. -0.000488 Da

Percolator Score: 1.00 | Hyperscore: 10.12 | Spectral Angle: 0.00 | Pearson's Correlation: 0.00

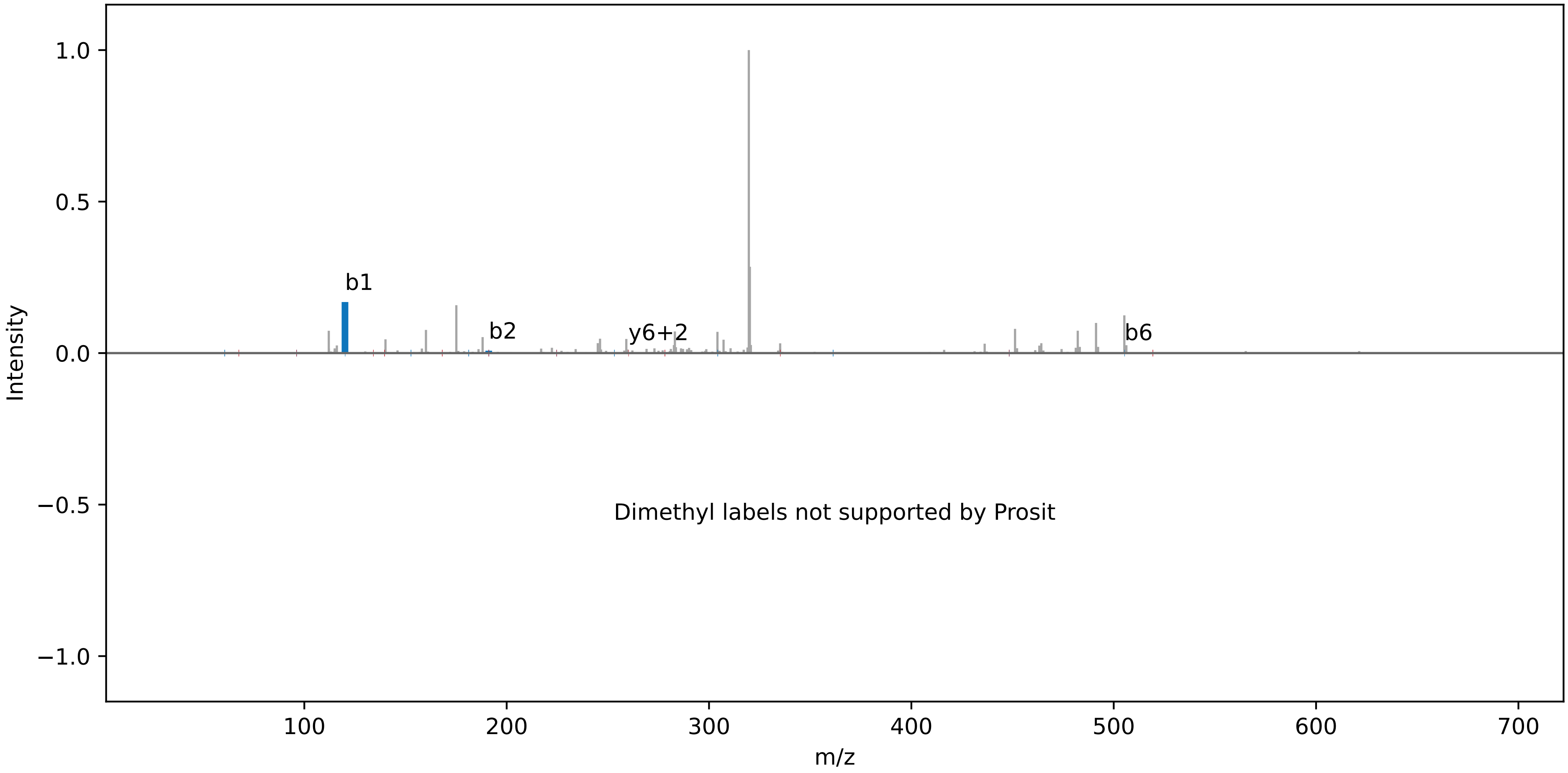

**TPQCAPPTYPK | NC_000911.1 : 1996578 : −. pep01**

Charge: +2 | Collision Energy: 27.0 | Percursor m/z: 630.309143 | Mass Diff. 0.000977 Da

Percolator Score: 1.00 | Hyperscore: 22.98 | Spectral Angle: 0.63 | Pearson's Correlation: 0.79

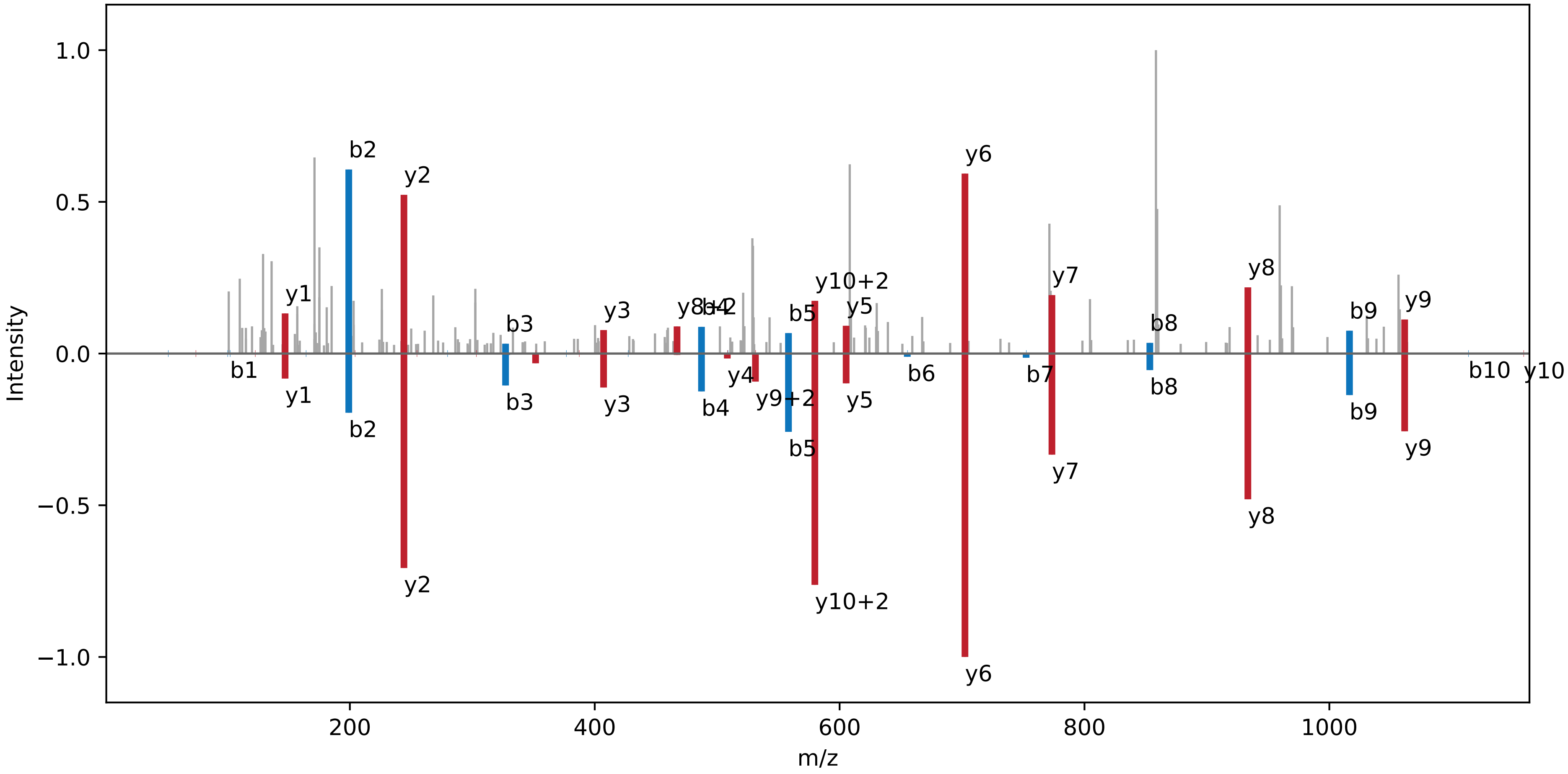

**[DmH] − RVTASVAPLLDTPSTTR | NC_000911.1 : 2081442 : −. pep01**

Charge: +3 | Collision Energy: 27.0 | Percursor m/z: 607.692200 | Mass Diff. 0.000000 Da

Percolator Score: 1.00 | Hyperscore: 25.42 | Spectral Angle: 0.00 | Pearson's Correlation: 0.00

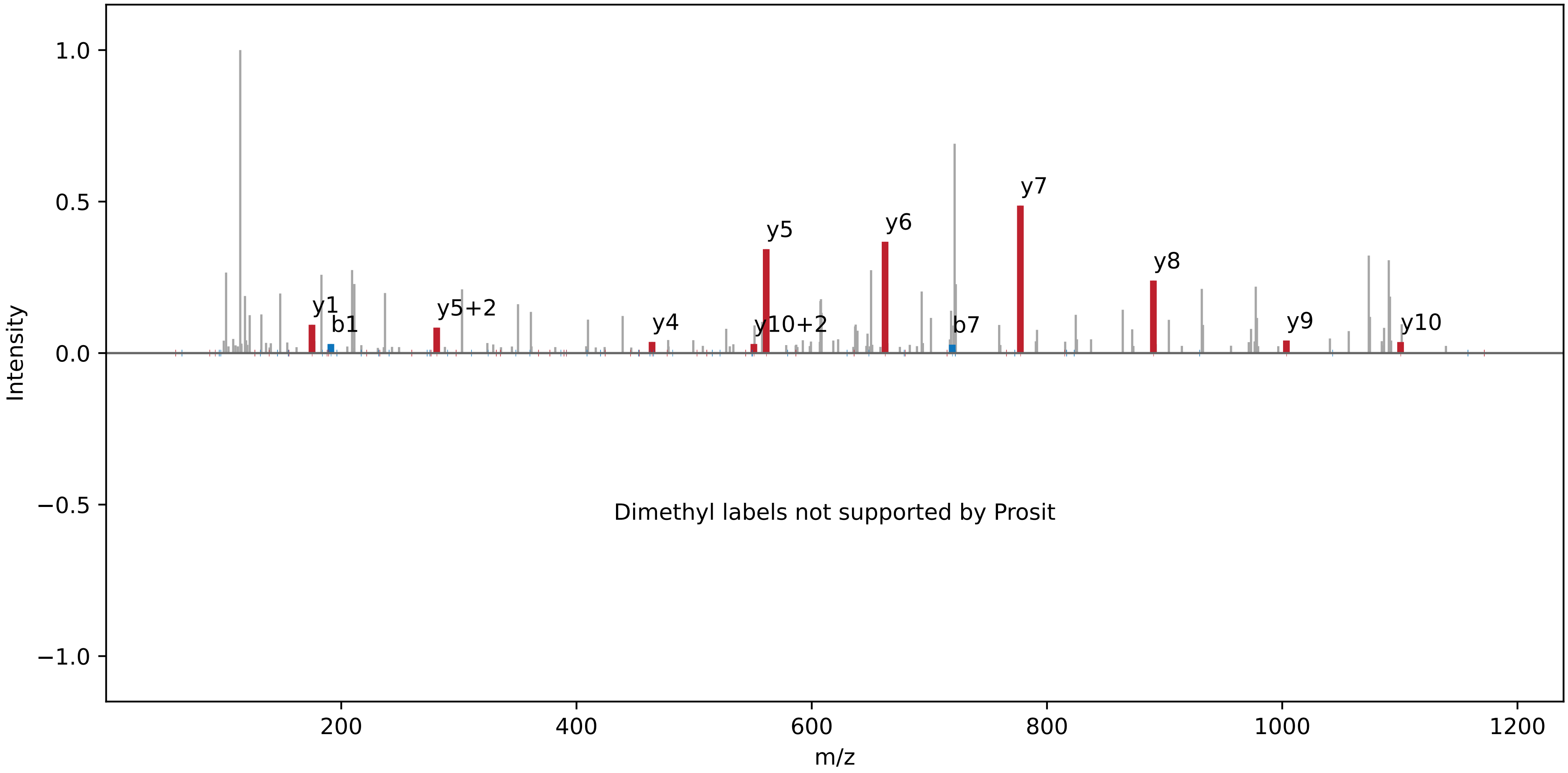

**[DmM] − AGAM[Ox]DT[Ph]IR | NC_000911.1 : 764875 : −. pep01**

Charge: +2 | Collision Energy: 27.0 | Percursor m/z: 481.720856 | Mass Diff. 0.002991 Da

Percolator Score: 1.00 | Hyperscore: 11.90 | Spectral Angle: 0.00 | Pearson's Correlation: 0.00

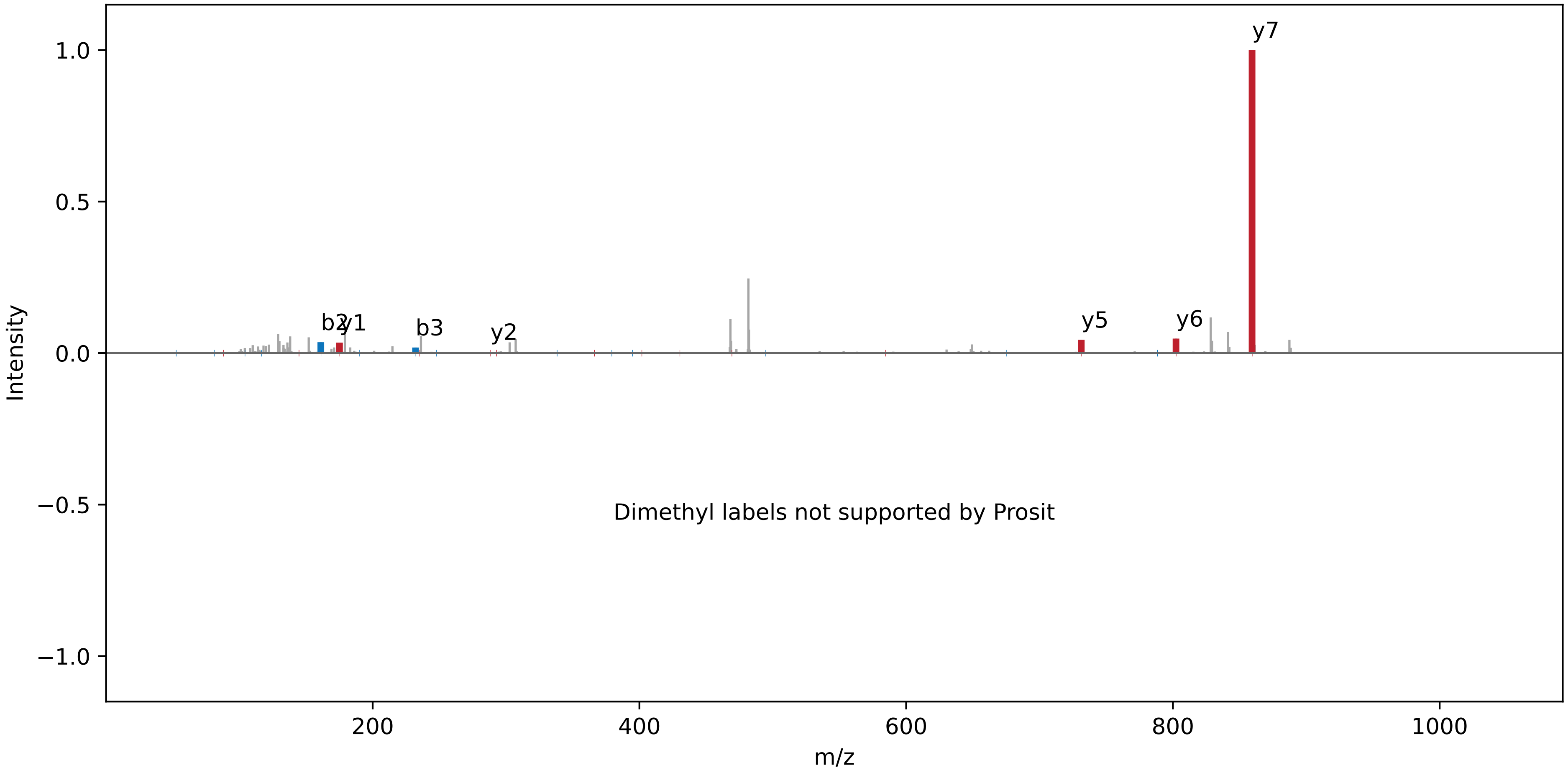

**[DmM] − PGLFVGLRL | NC_000911.1 : 2281858 : +. pep01**

Charge: +2 | Collision Energy: 27.0 | Percursor m/z: 502.336853 | Mass Diff. 0.004822 Da

Percolator Score: 1.00 | Hyperscore: 19.49 | Spectral Angle: 0.00 | Pearson's Correlation: 0.00

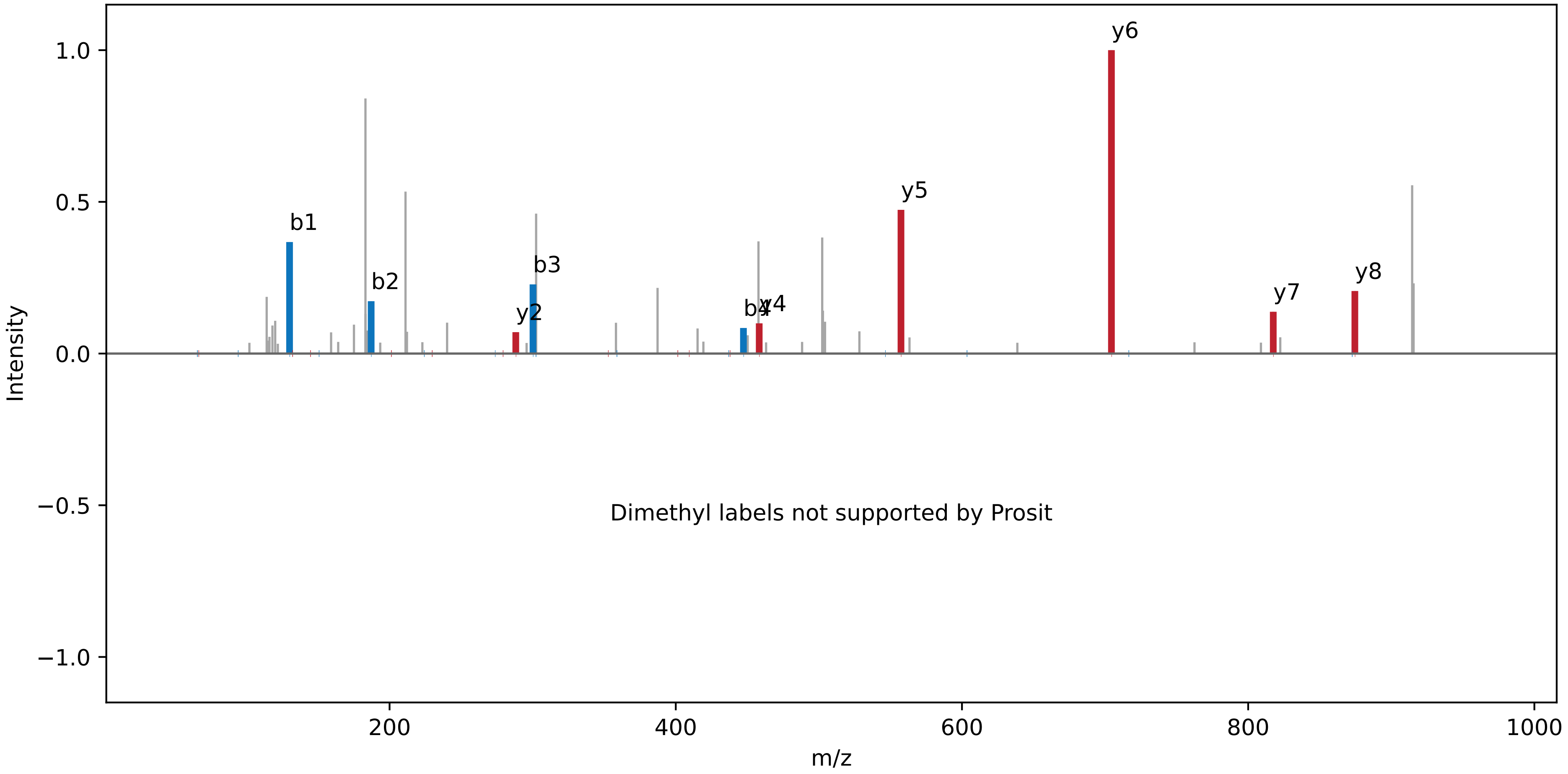

**SM[Ox]M[Ox]CSPHCWPR | NC_000911.1 : 2476692 : +. pep01**

Charge: +3 | Collision Energy: 27.0 | Percursor m/z: 494.191254 | Mass Diff. -0.001587 Da

Percolator Score: 1.00 | Hyperscore: 8.21 | Spectral Angle: 0.14 | Pearson's Correlation: 0.15

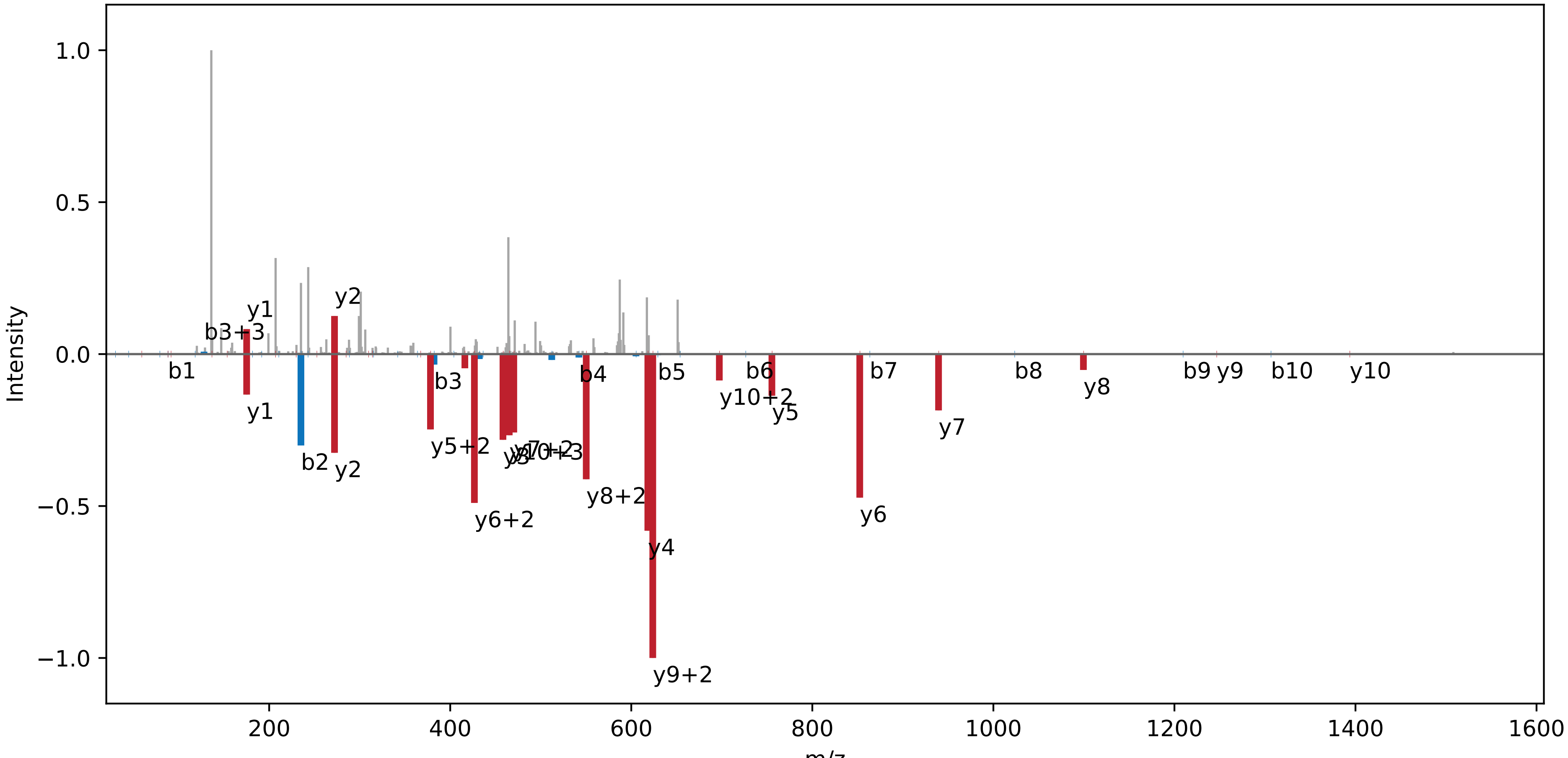

**NGNSSIGIGWGAPQSGGDLFC | NC_000911.1 : 2633382 : −. pep01**

Charge: +3 | Collision Energy: 27.0 | Percursor m/z: 698.647766 | Mass Diff. -0.005615 Da

Percolator Score: 1.00 | Hyperscore: 12.65 | Spectral Angle: 0.01 | Pearson's Correlation: -0.02

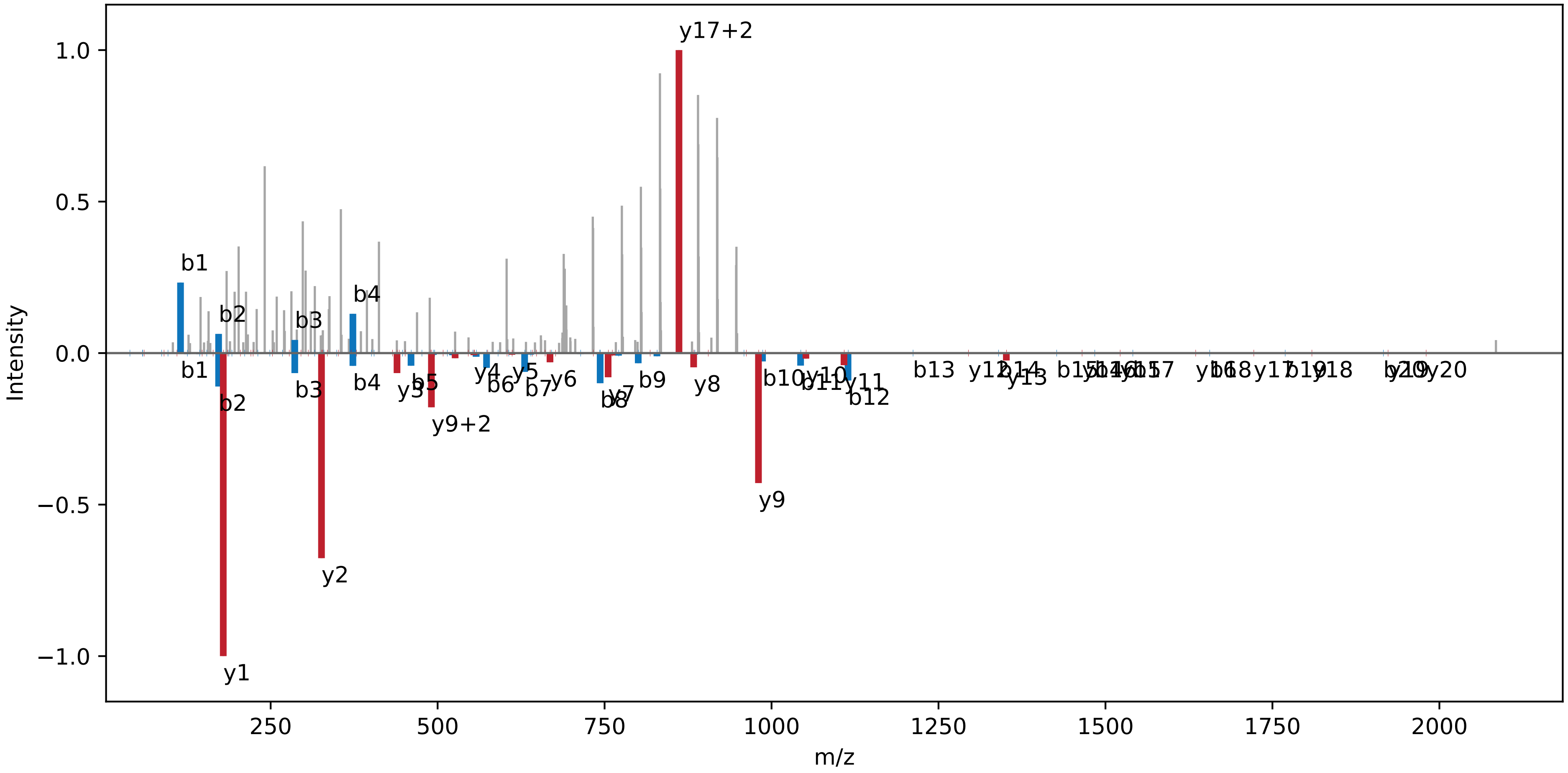

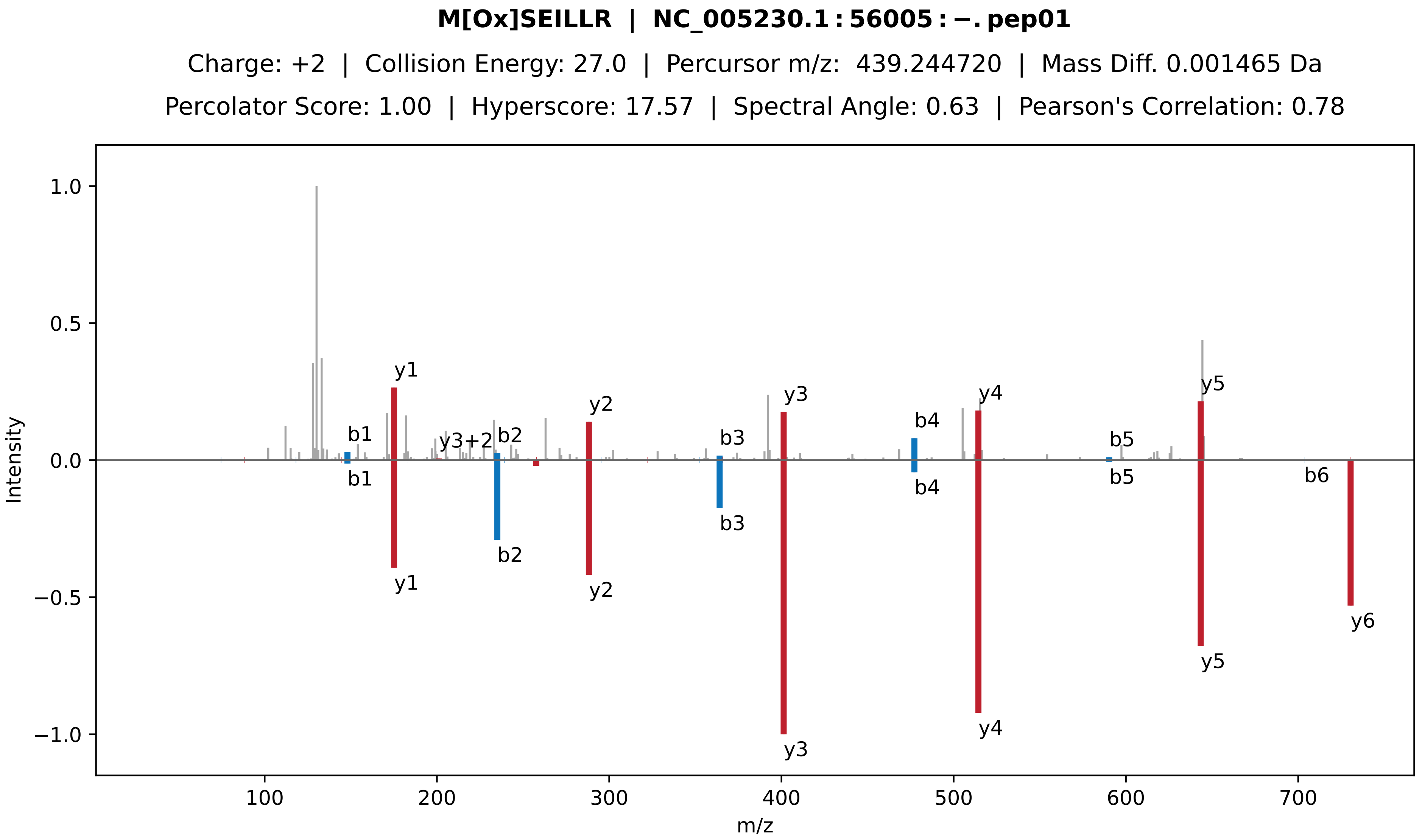

M[Ox]SEILLR | NC_005230.1 : 56005 : −. pep01
Charge: +2 | Collision Energy: 27.0 | Percursor m/z: 439.244720 | Mass Diff. 0.001465 Da
Percolator Score: 1.00 | Hyperscore: 17.57 | Spectral Angle: 0.63 | Pearson's Correlation: 0.78
Intensity
m/z
1.0
0.5
0.0
−0.5
−1.0
100
200
300
400
500
600
700
y1
b1
y3+2
b2
y2
b3
y3
b4
y4
b5
y5
b6
y6

**RLEPVLK | NC_000911.1 : 3260036 : −. pep01**

Charge: +2 | Collision Energy: 27.0 | Percursor m/z: 427.776031 | Mass Diff. 0.000244 Da

Percolator Score: 1.00 | Hyperscore: 14.29 | Spectral Angle: 0.68 | Pearson's Correlation: 0.85

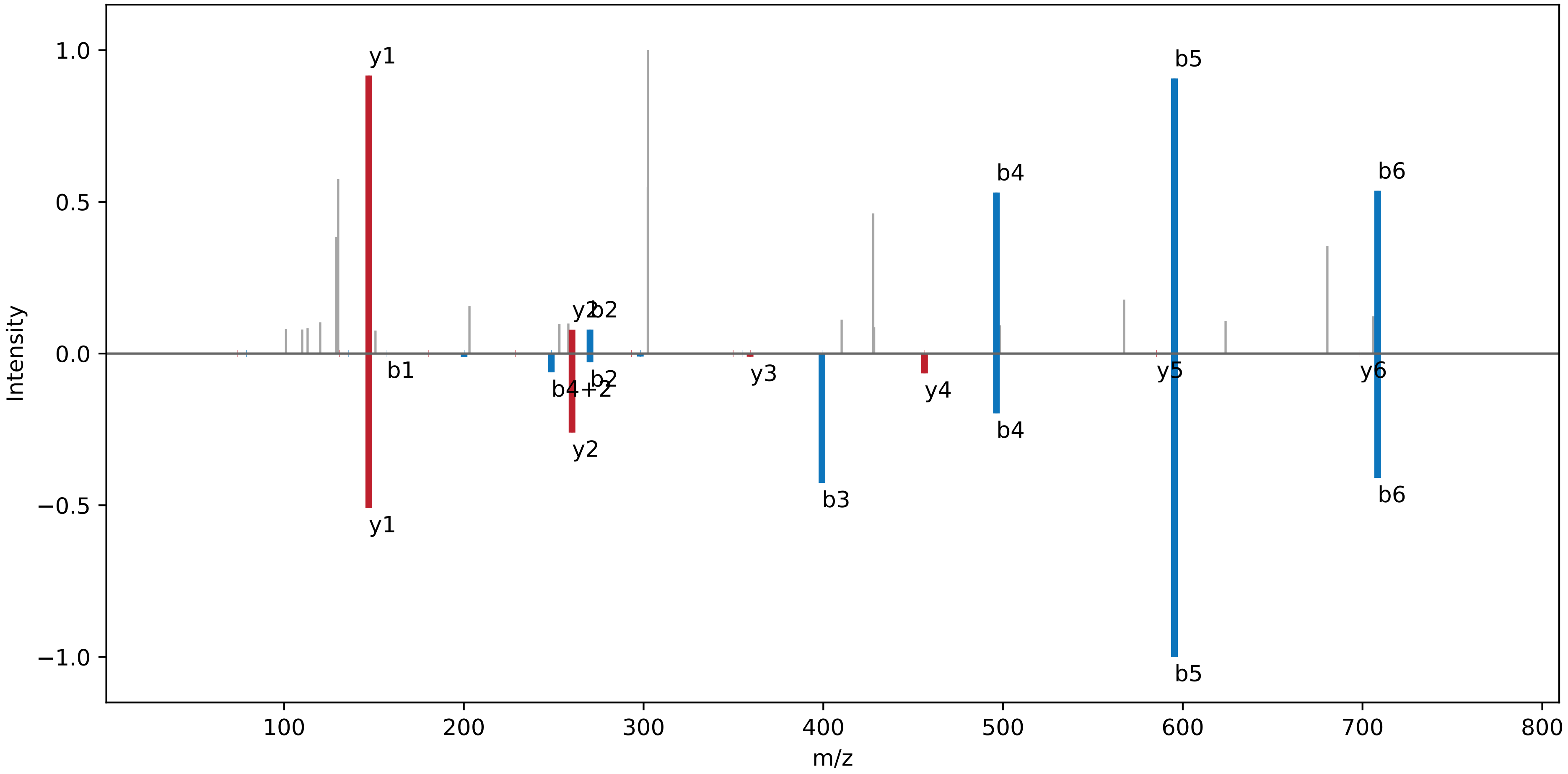

**PLAIEGR | NC_000911.1:3291170:+.pep01**

Charge: +2 | Collision Energy: 27.0 | Percursor m/z: 378.225555 | Mass Diff. 0.000061 Da

Percolator Score: 1.00 | Hyperscore: 10.42 | Spectral Angle: 0.82 | Pearson's Correlation: 0.96

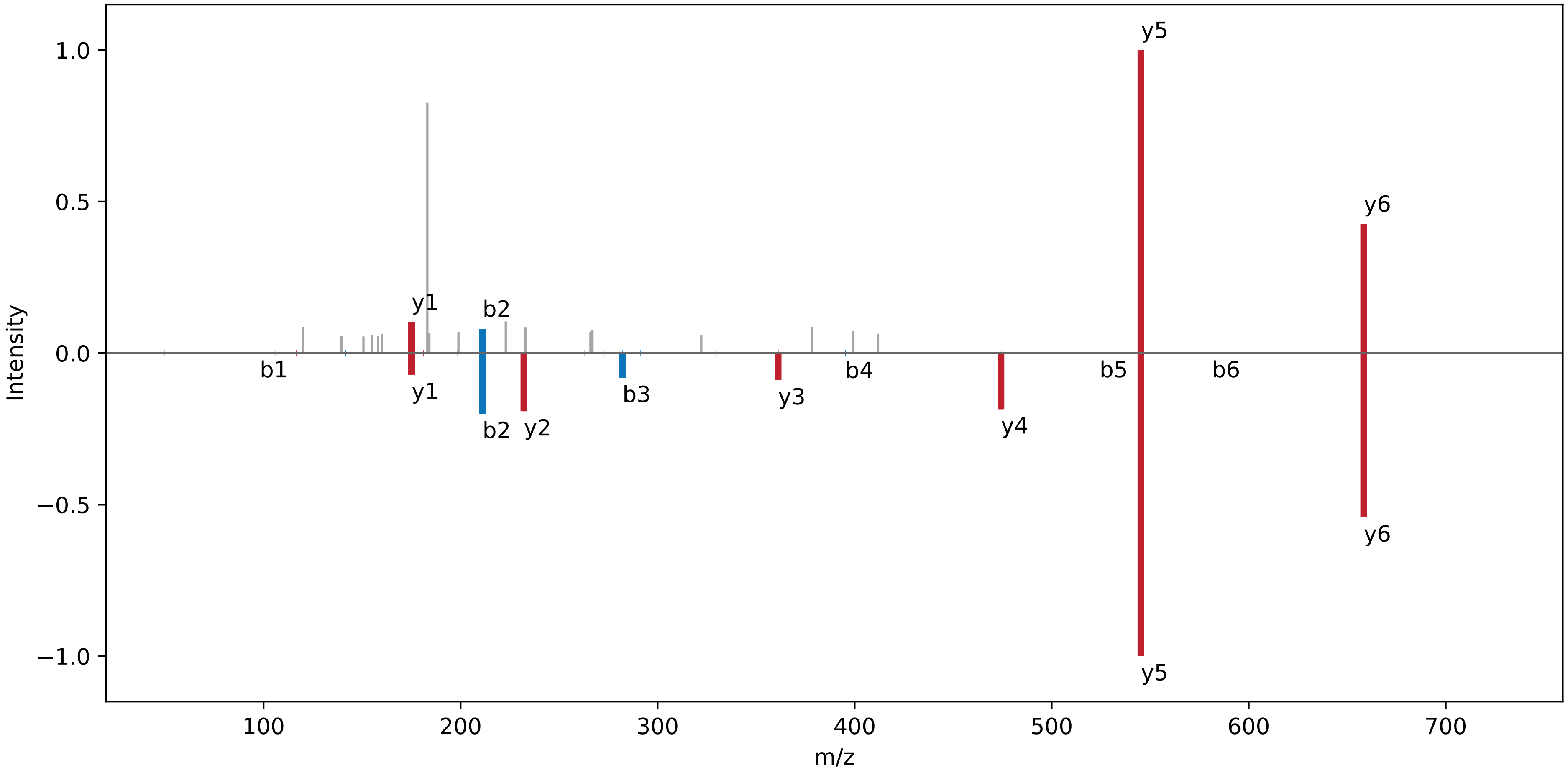

**GAGGLFGR | NC_000911.1 : 3306954 : −. pep01**

Charge: +2 | Collision Energy: 27.0 | Percursor m/z: 367.701263 | Mass Diff. -0.000549 Da

Percolator Score: 1.00 | Hyperscore: 12.97 | Spectral Angle: 0.51 | Pearson's Correlation: 0.67

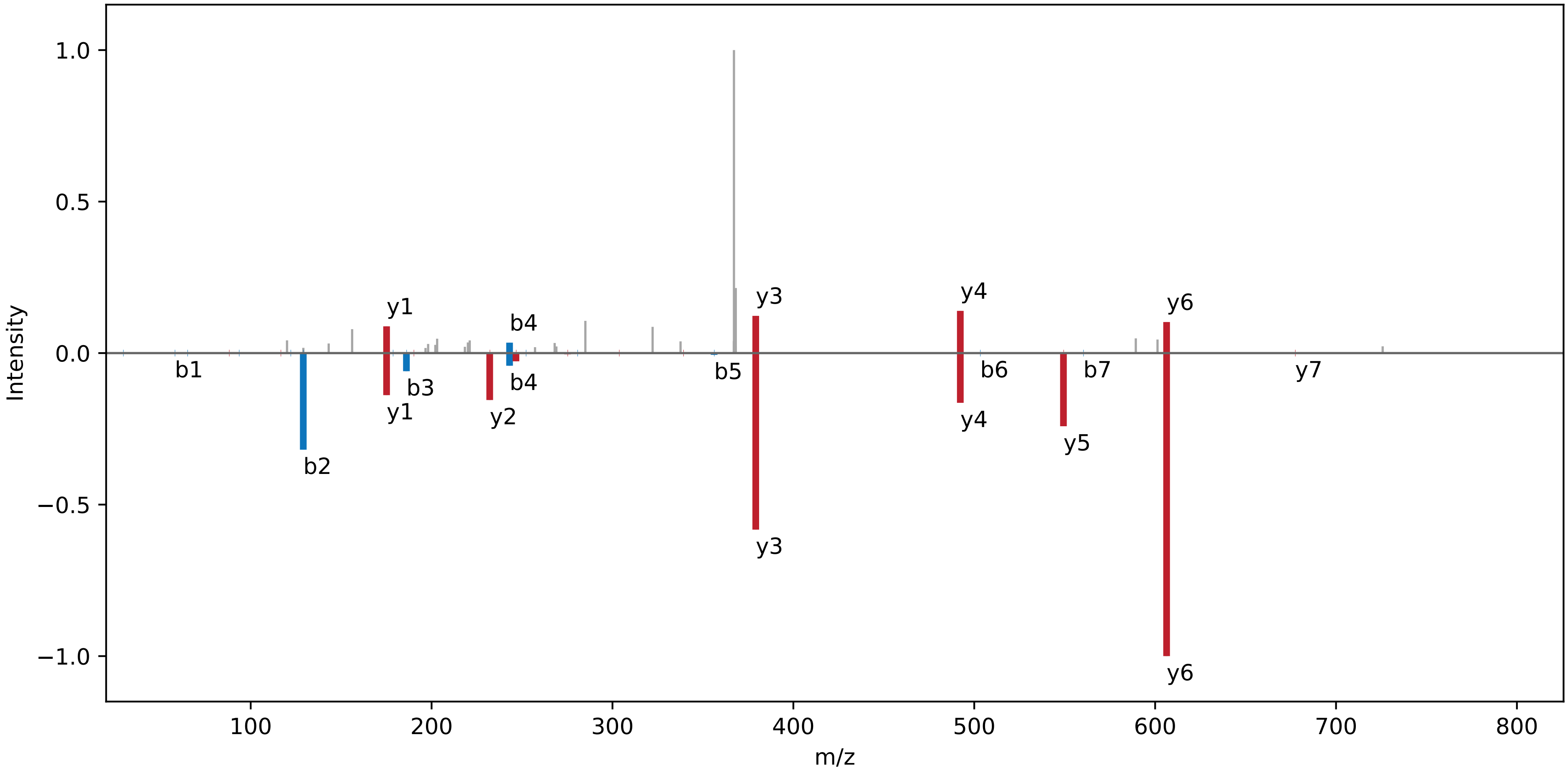

**WGAVWAK | NC_000911.1 : 495905 : +. pep01**

Charge: +2 | Collision Energy: 27.0 | Percursor m/z: 409.221802 | Mass Diff. -0.000061 Da

Percolator Score: 1.00 | Hyperscore: 16.15 | Spectral Angle: 0.51 | Pearson's Correlation: 0.64

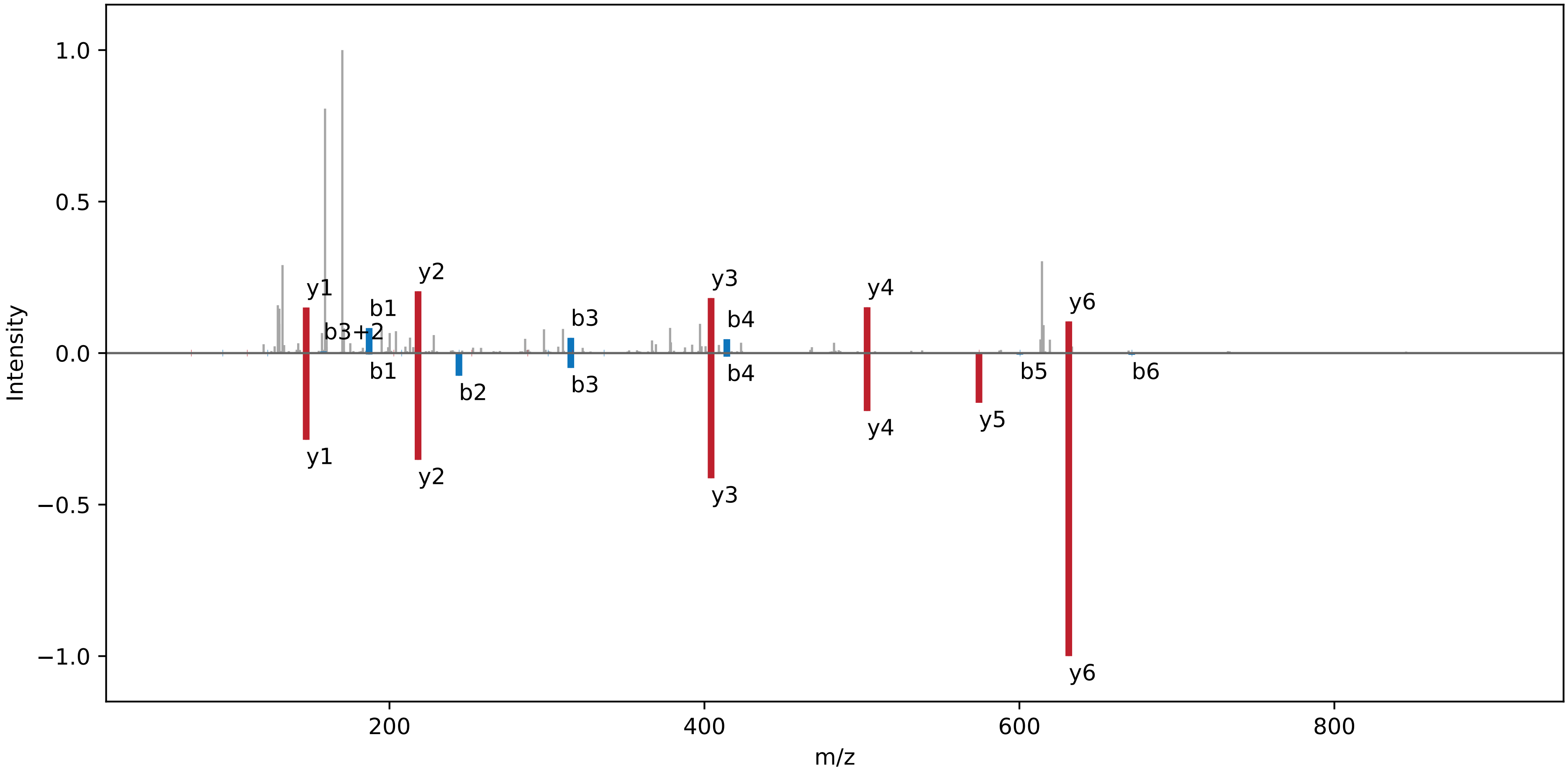

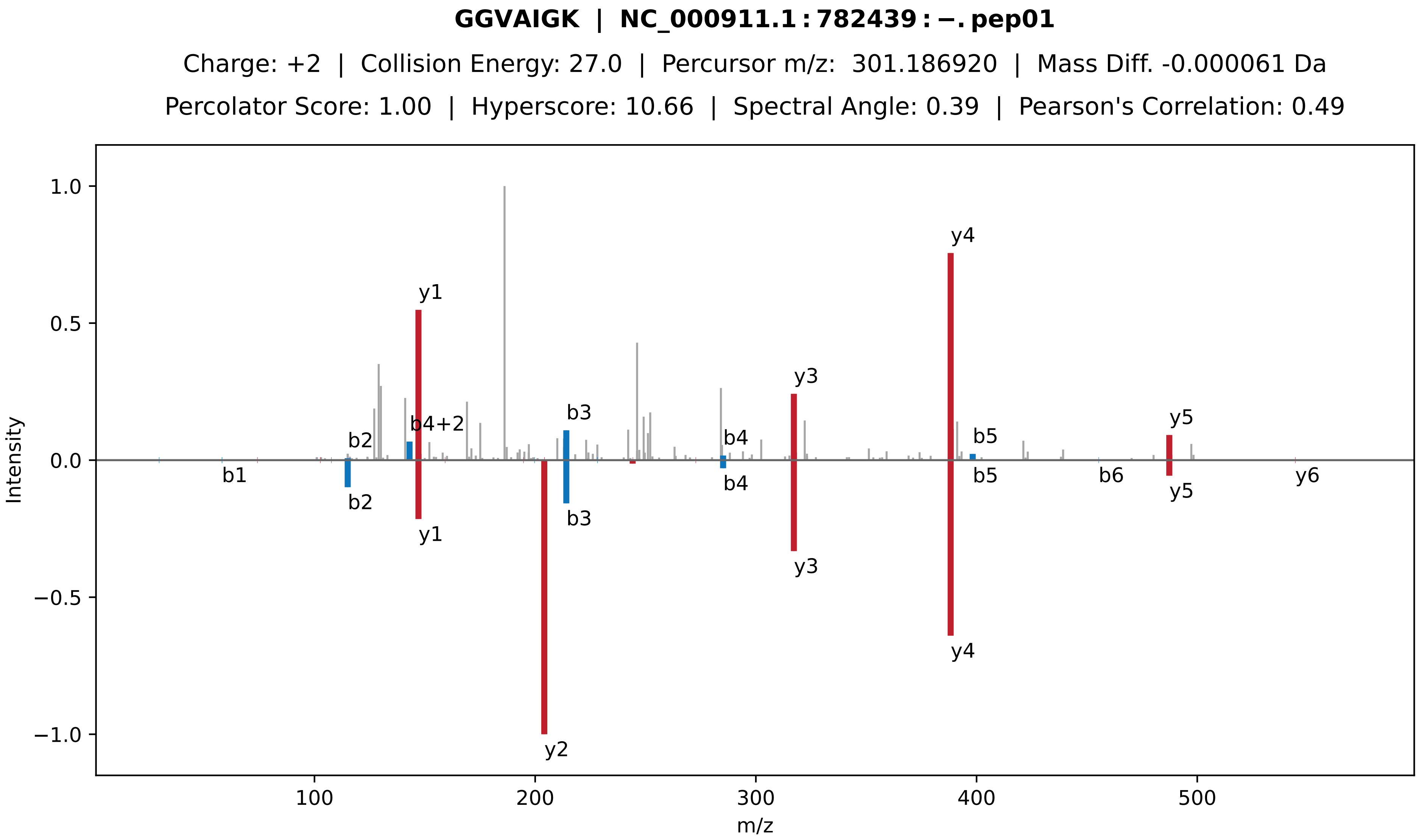
GGVAIGK | NC_000911.1:782439:−.pep01
Charge: +2 | Collision Energy: 27.0 | Percursor m/z: 301.186920 | Mass Diff. -0.000061 Da
Percolator Score: 1.00 | Hyperscore: 10.66 | Spectral Angle: 0.39 | Pearson's Correlation: 0.49
Intensity
m/z
y1
y2
y3
y4
y5
y6
b1
b2
b3
b4
b5
b6
b4+2

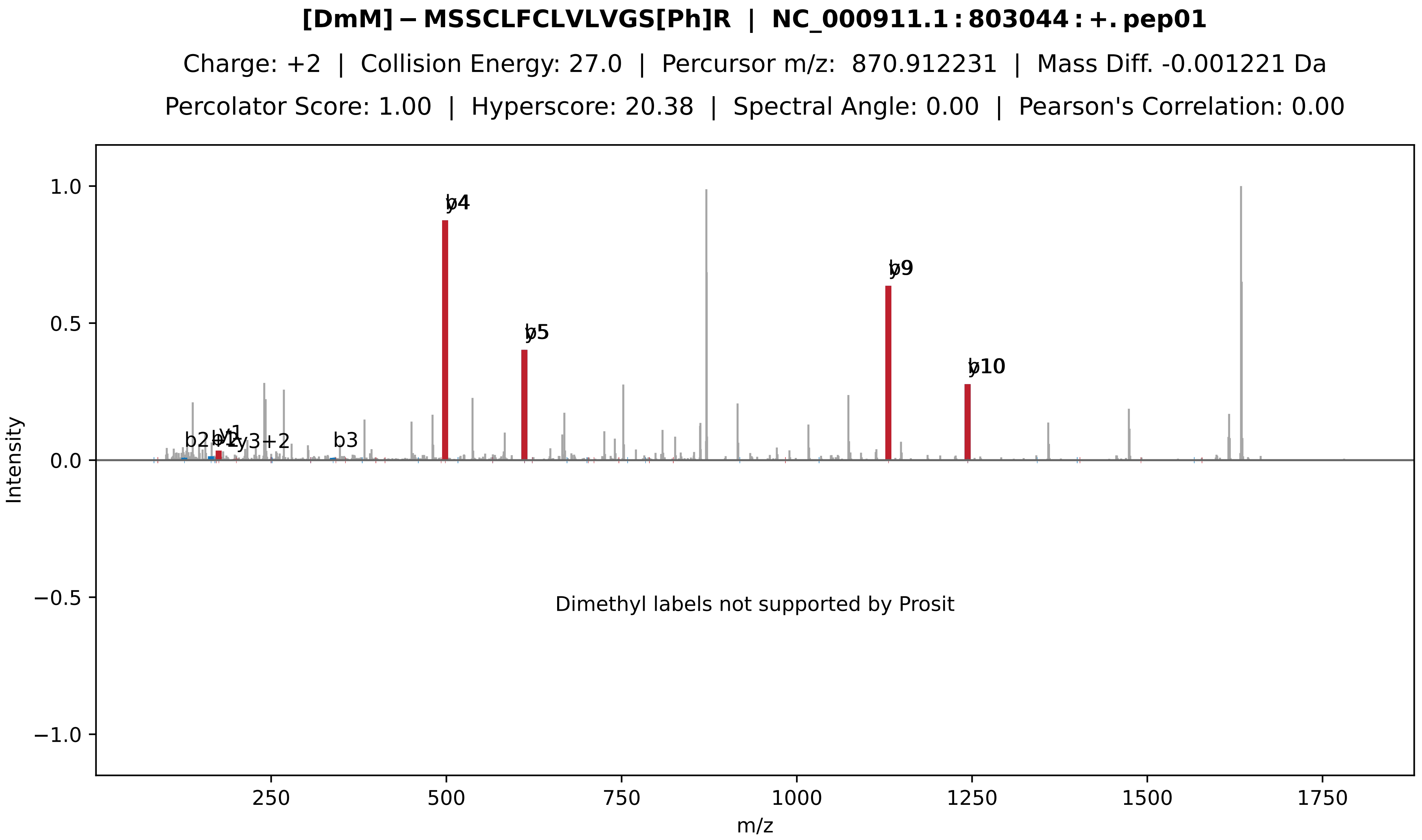
[DmM]−MSSCLFCLVLVGS[Ph]R | NC_000911.1:803044:+.pep01
Charge: +2 | Collision Energy: 27.0 | Percursor m/z: 870.912231 | Mass Diff. -0.001221 Da
Percolator Score: 1.00 | Hyperscore: 20.38 | Spectral Angle: 0.00 | Pearson's Correlation: 0.00
y4
y5
y9
y10
b3
Dimethyl labels not supported by Prosit
Intensity
m/z
1.0
0.5
0.0
−0.5
−1.0
250
500
750
1000
1250
1500
1750

**CLVLGAGRWVARPLCLGLLFACYHAIM[Ox]GLLVVPLGVALCCLFLPCR | NC_005232.1:23864:+.pep01**

Charge: +4 | Collision Energy: 27.0 | Percursor m/z: 1319.450439 | Mass Diff. 3.001953 Da

Percolator Score: 1.00 | Hyperscore: 19.23 | Spectral Angle: 0.00 | Pearson's Correlation: 0.00

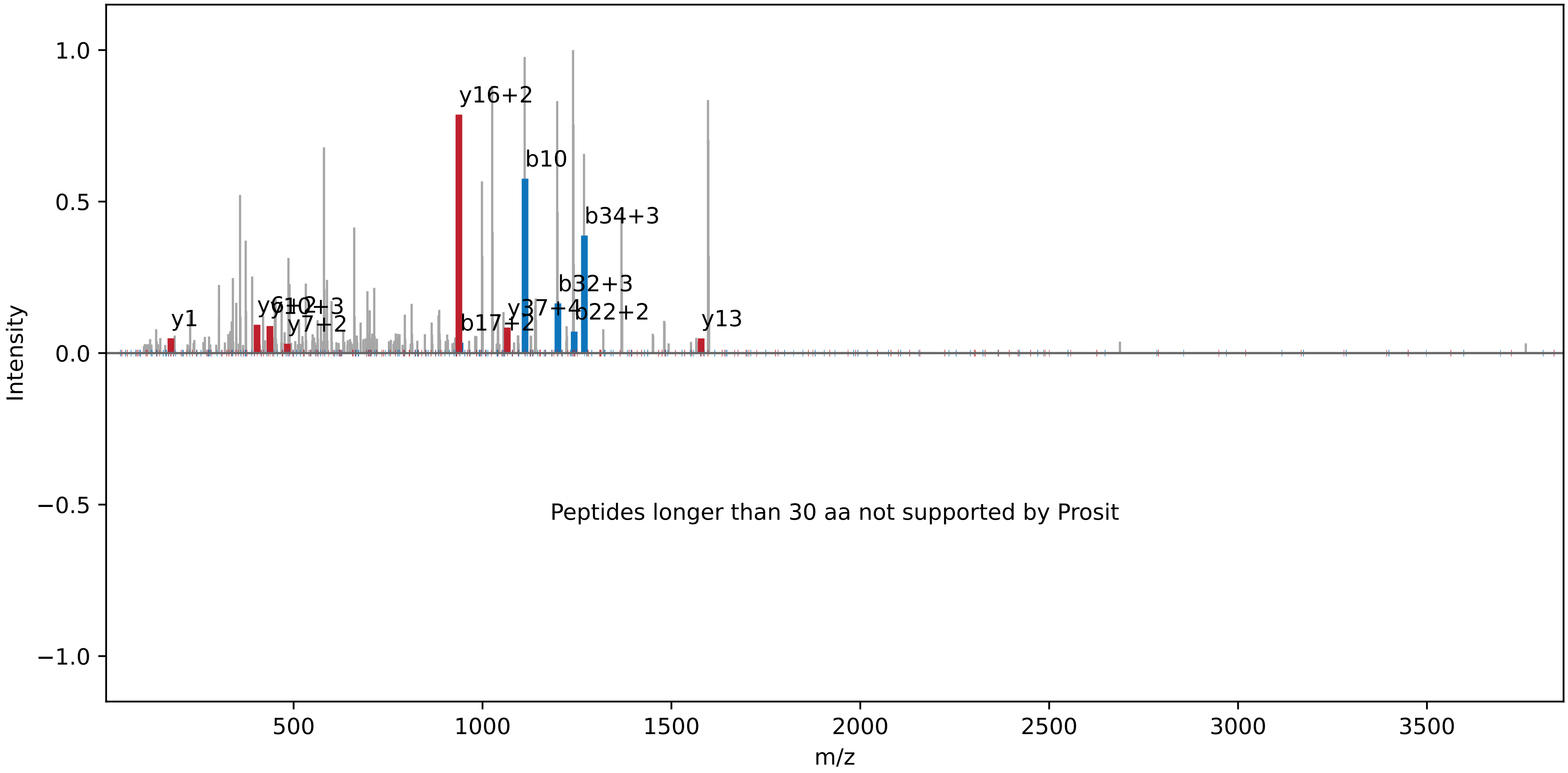

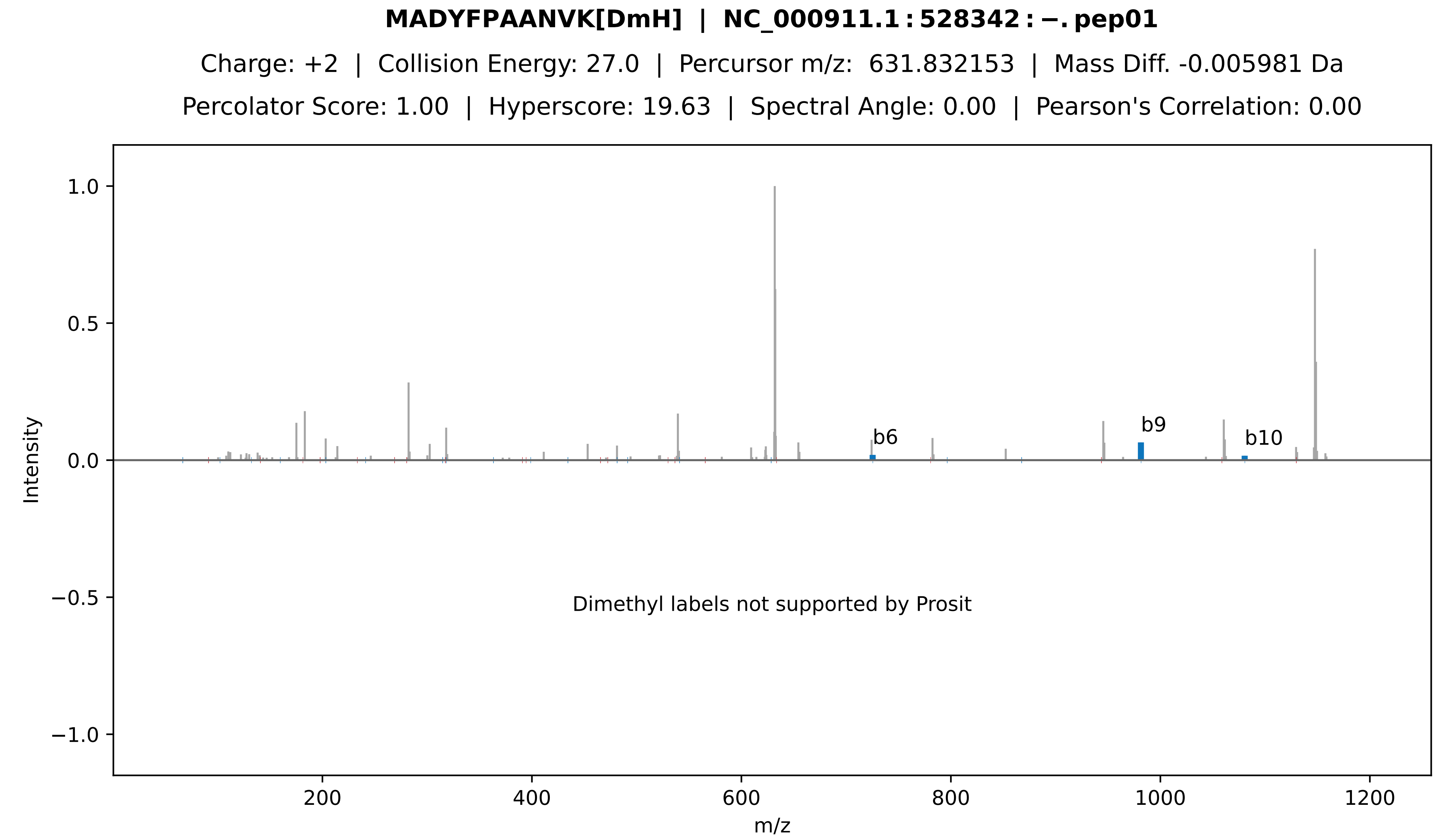

MADYFPAANVK[DmH] | NC_000911.1:528342:−.pep01
Charge: +2 | Collision Energy: 27.0 | Percursor m/z: 631.832153 | Mass Diff. -0.005981 Da
Percolator Score: 1.00 | Hyperscore: 19.63 | Spectral Angle: 0.00 | Pearson's Correlation: 0.00
Intensity
m/z
b6
b9
b10
Dimethyl labels not supported by Prosit

**[DmL]−VSLAVGFAIIHHR | NC_000911.1:298745:+.pep01**

Charge: +2 | Collision Energy: 27.0 | Percursor m/z: 724.431091 | Mass Diff. 0.001831 Da

Percolator Score: 1.00 | Hyperscore: 26.11 | Spectral Angle: 0.00 | Pearson's Correlation: 0.00

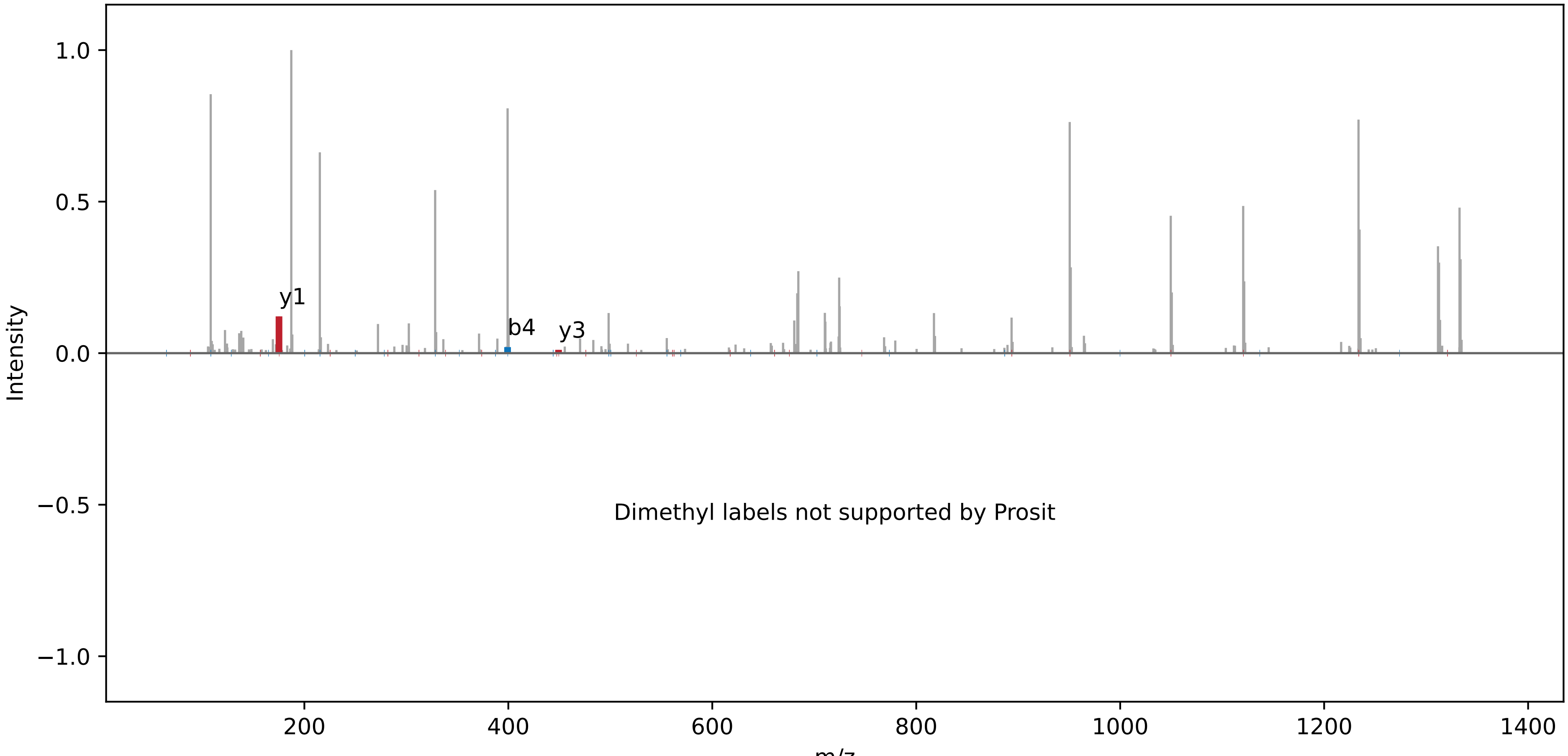

**[DmM] − PGLSGIIAP | NC_000911.1 : 323562 : +. pep01**

Charge: +2 | Collision Energy: 35.0 | Percursor m/z: 428.783752 | Mass Diff. 0.016296 Da

Percolator Score: 1.00 | Hyperscore: 24.68 | Spectral Angle: 0.00 | Pearson's Correlation: 0.00

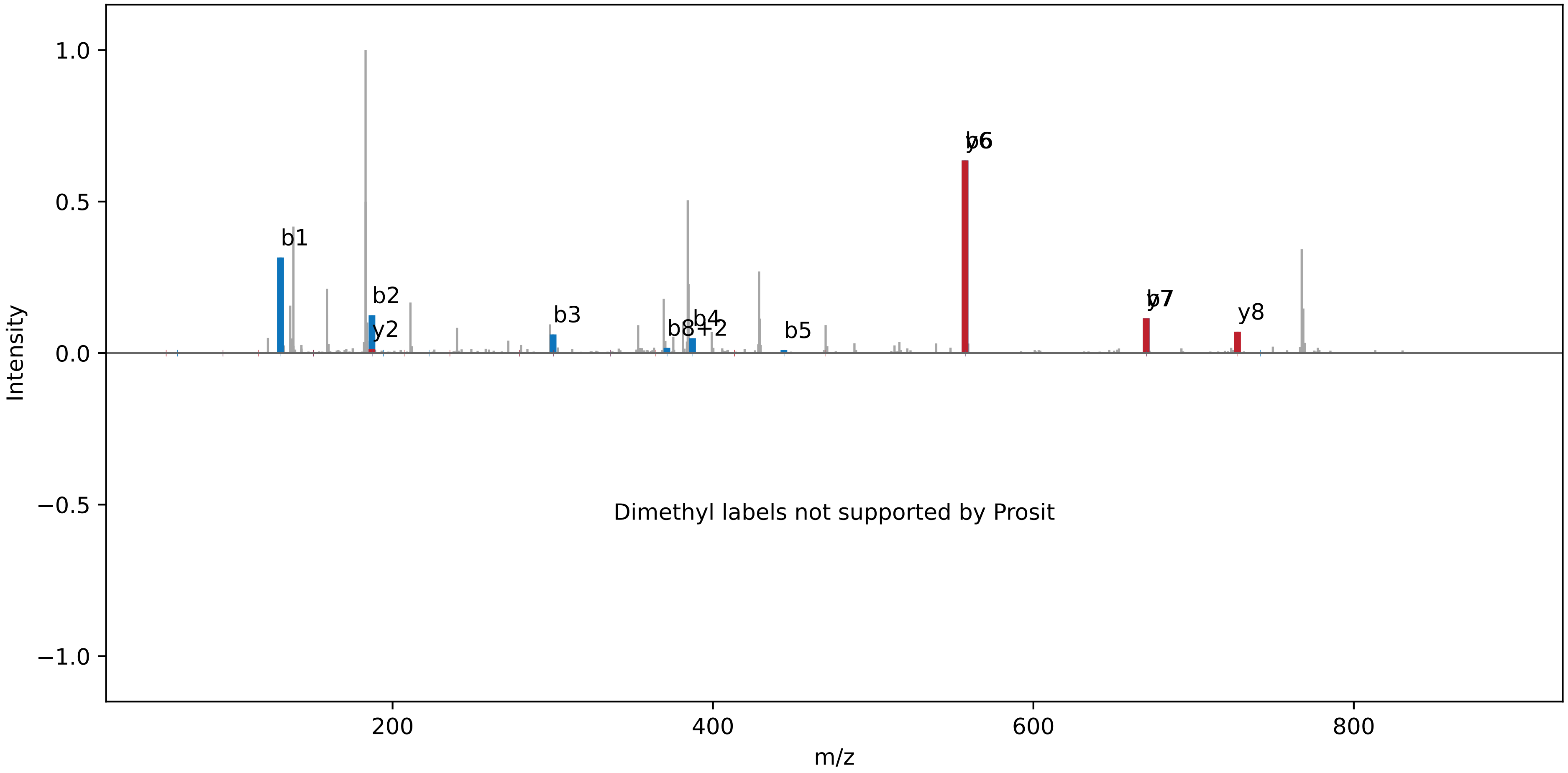

**[DmM] − MTVDGSSSWGQWPAGK[DmL] | NC_000911.1 : 1043772 : −. pep01**

Charge: +2 | Collision Energy: 27.0 | Percursor m/z: 877.433411 | Mass Diff. 0.008057 Da

Percolator Score: 1.00 | Hyperscore: 20.87 | Spectral Angle: 0.00 | Pearson's Correlation: 0.00

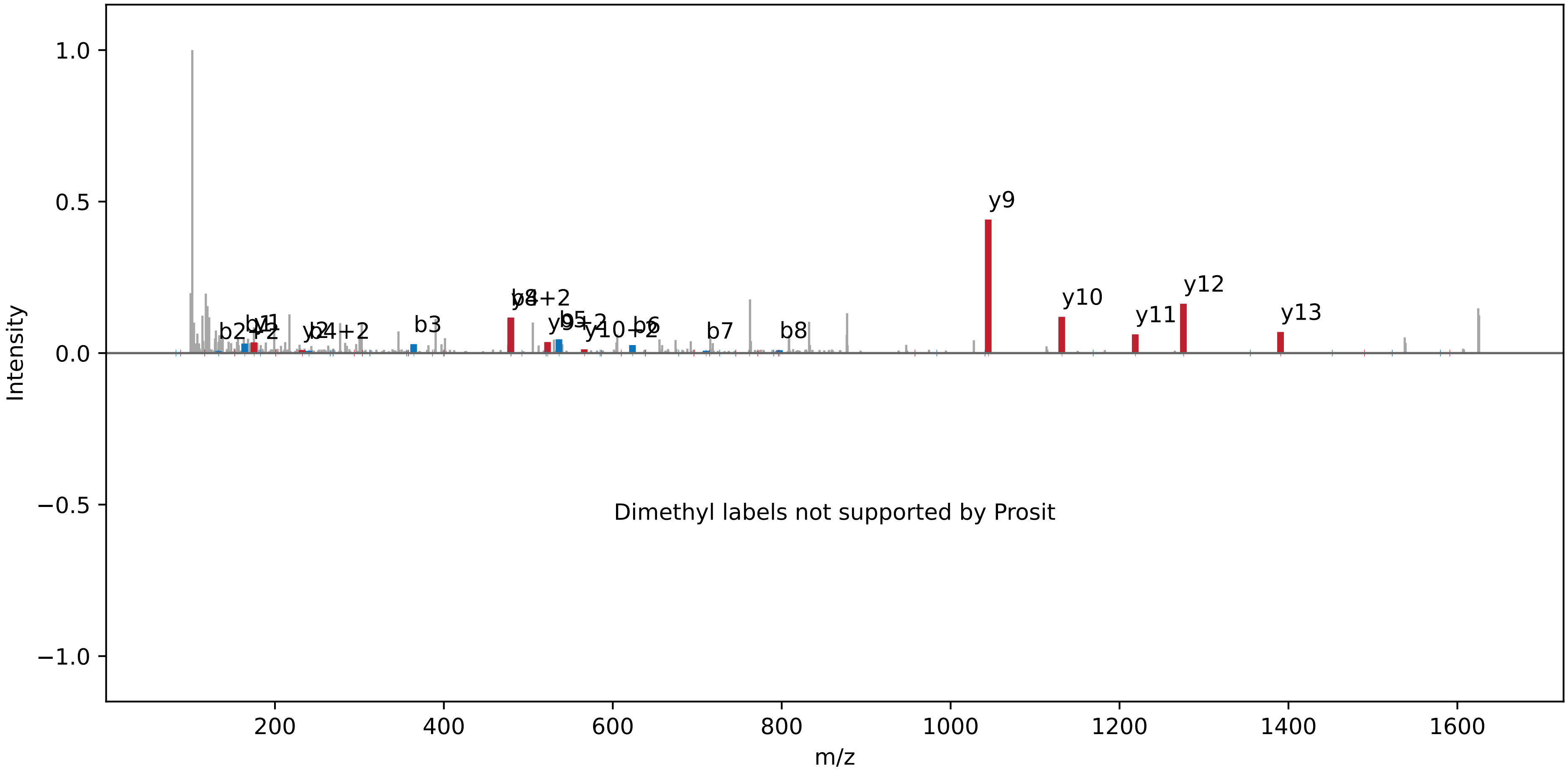

**[DmL] − PGVEGDLVK[DmM] | NC_000911.1 : 1125756 : +. pep01**

Charge: +2 | Collision Energy: 27.0 | Percursor m/z: 487.298004 | Mass Diff. 0.002136 Da

Percolator Score: 1.00 | Hyperscore: 14.39 | Spectral Angle: 0.00 | Pearson's Correlation: 0.00

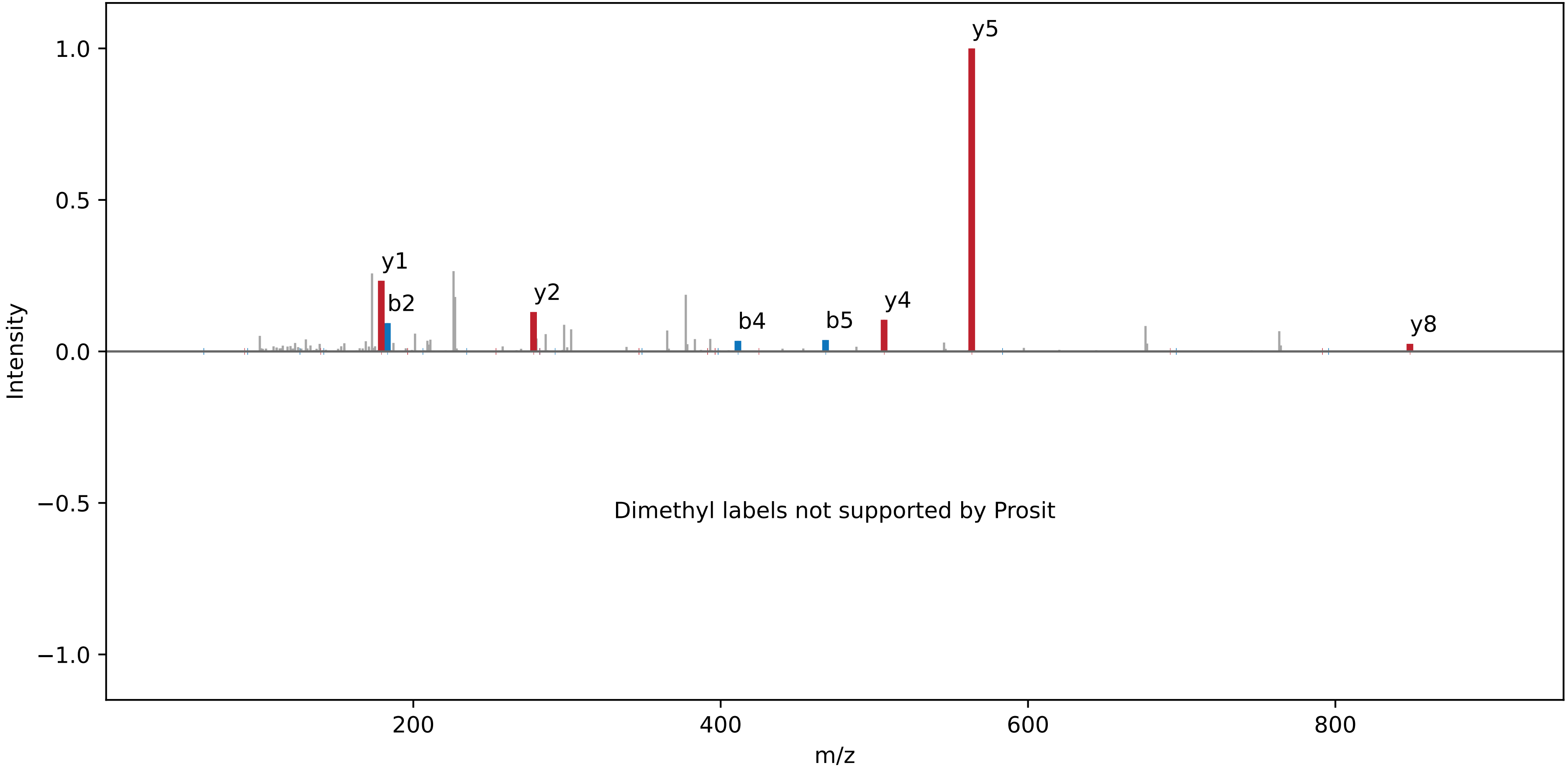

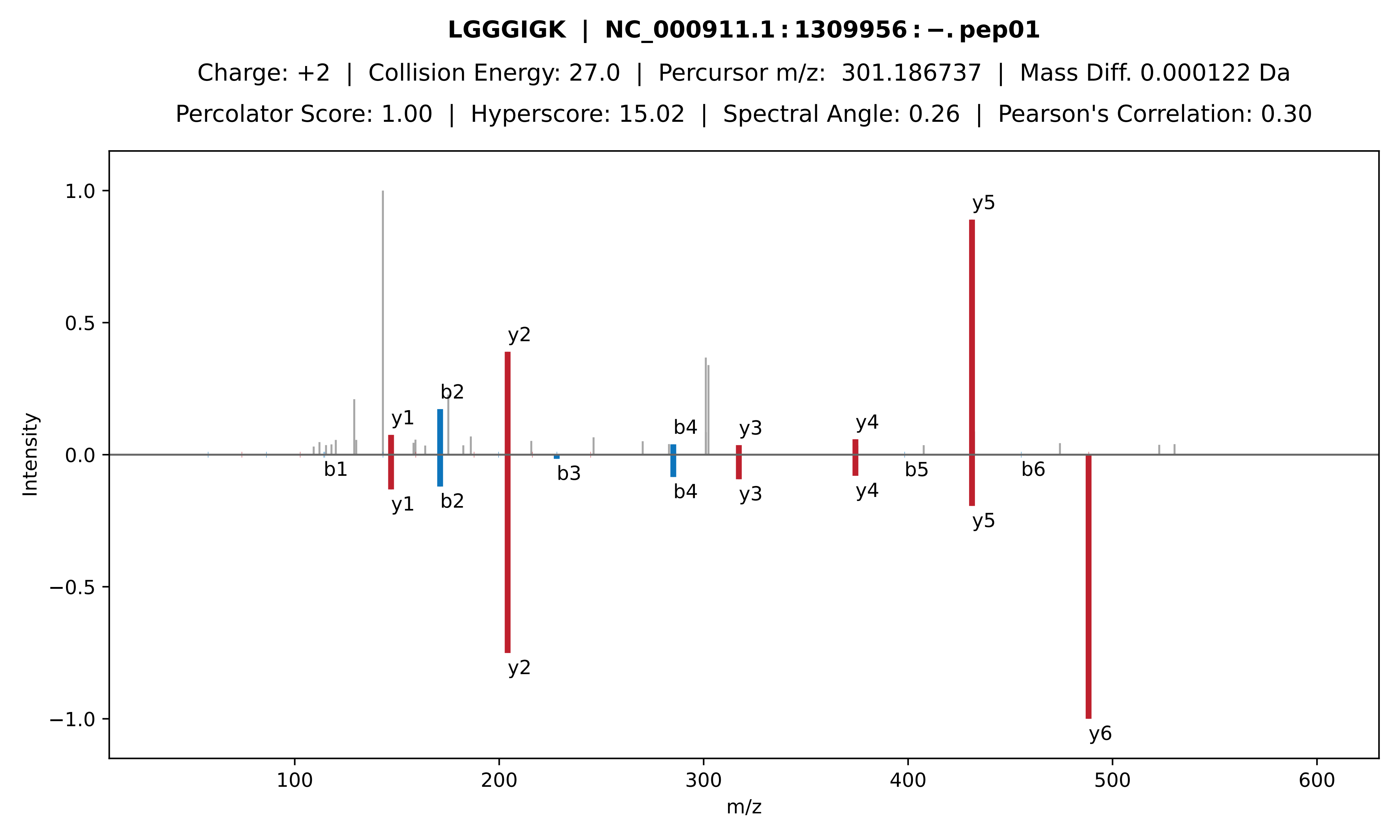

LGGGIGK | NC_000911.1 : 1309956 : −. pep01
Charge: +2 | Collision Energy: 27.0 | Percursor m/z: 301.186737 | Mass Diff. 0.000122 Da
Percolator Score: 1.00 | Hyperscore: 15.02 | Spectral Angle: 0.26 | Pearson's Correlation: 0.30
Intensity
m/z
1.0
0.5
0.0
−0.5
−1.0
100
200
300
400
500
600
y1
b2
y2
b4
y3
y4
y5
b1
b3
b5
b6
y6

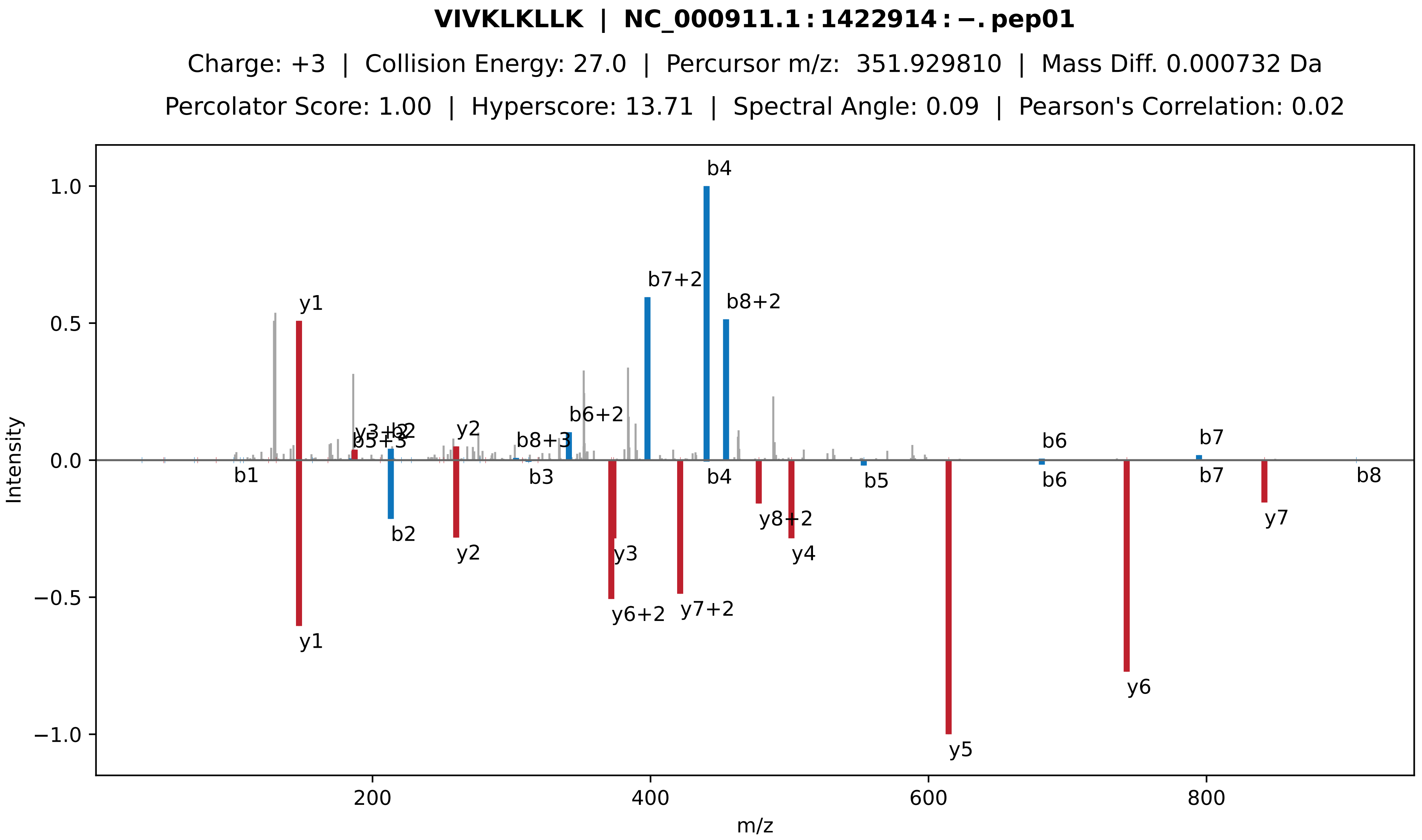
VIVKLKLLK | NC_000911.1:1422914:−.pep01
Charge: +3 | Collision Energy: 27.0 | Percursor m/z: 351.929810 | Mass Diff. 0.000732 Da
Percolator Score: 1.00 | Hyperscore: 13.71 | Spectral Angle: 0.09 | Pearson's Correlation: 0.02
Intensity
m/z
b4
b7+2
b8+2
y1
b6+2
y2
b8+3
b6
b7
b1
b3
b5
b8
b2
y3
y6+2
y7+2
y8+2
y4
y5
y6
y7

**PQNQGLSQEPK | NC_000911.1 : 502776 : −. pep01**

Charge: +2 | Collision Energy: 27.0 | Percursor m/z: 613.311523 | Mass Diff. -0.001099 Da

Percolator Score: 1.00 | Hyperscore: 19.37 | Spectral Angle: 0.20 | Pearson's Correlation: 0.20

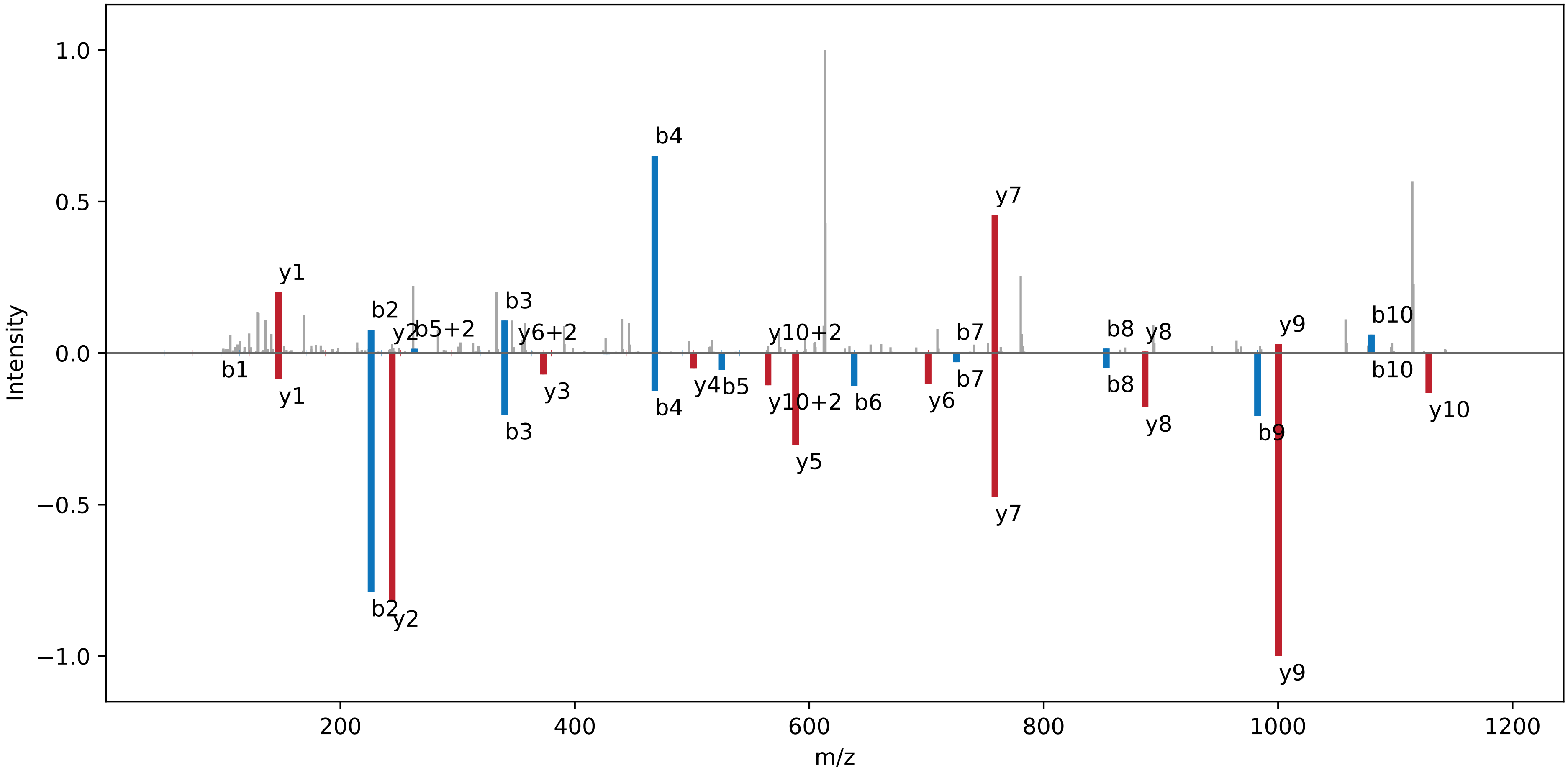

**[DmM] − FLM[Ox]IENK[DmM] | NC_000911.1 : 1527454 : +. pep01**

Charge: +2 | Collision Energy: 27.0 | Percursor m/z: 487.794586 | Mass Diff. -0.000427 Da

Percolator Score: 1.00 | Hyperscore: 16.24 | Spectral Angle: 0.00 | Pearson's Correlation: 0.00

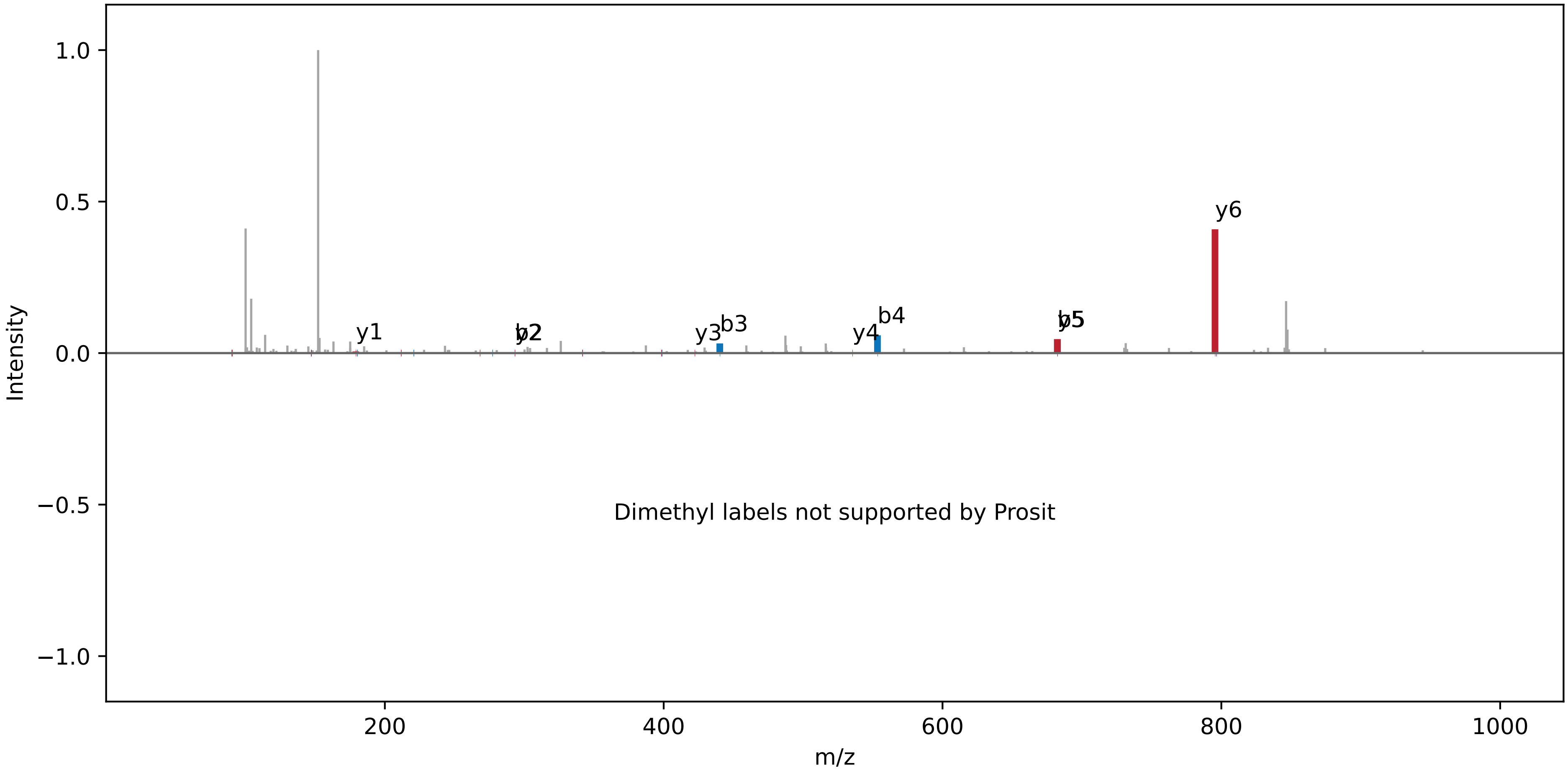

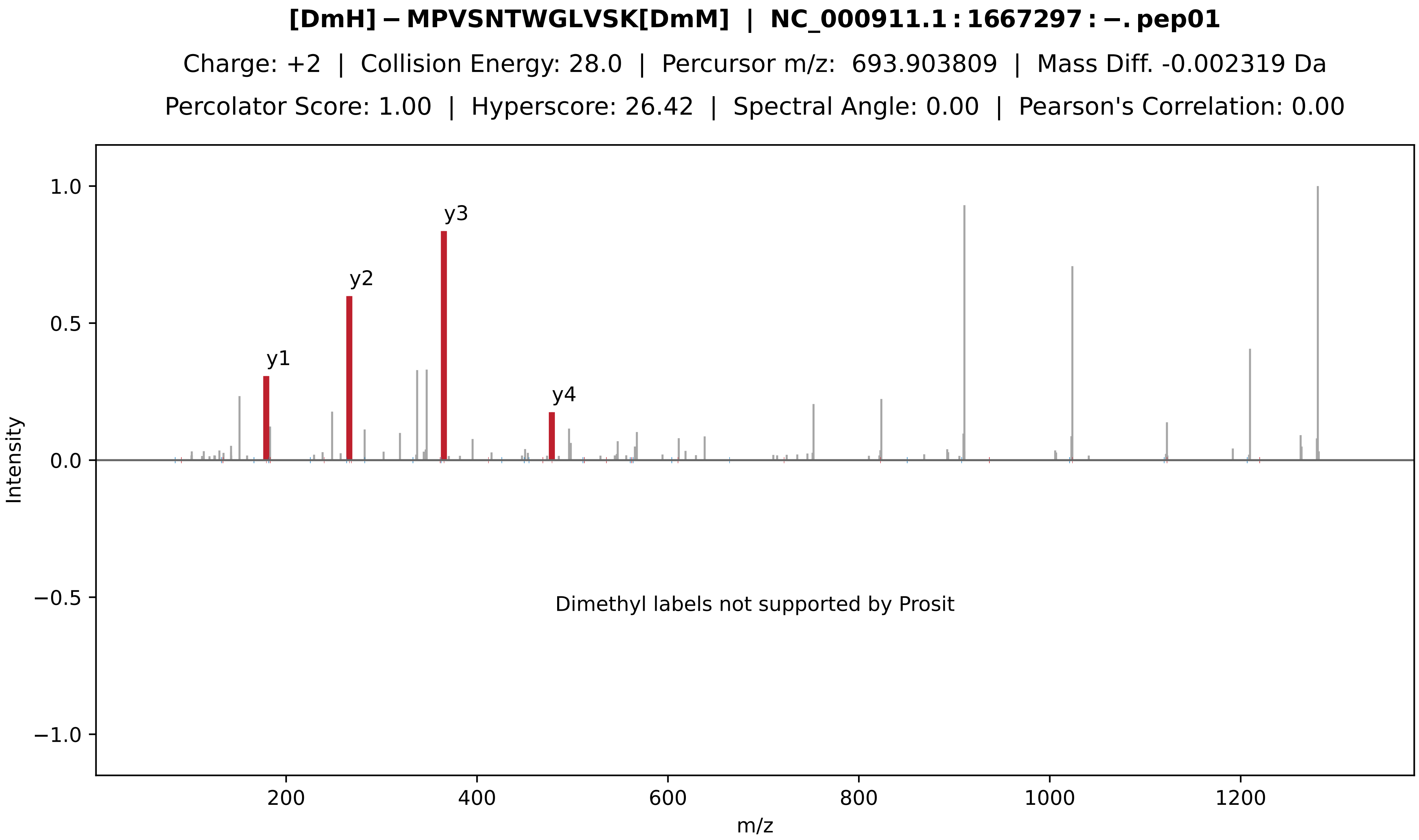

[DmH]−MPVSNTWGLVSK[DmM] | NC_000911.1:1667297:−.pep01
Charge: +2 | Collision Energy: 28.0 | Percursor m/z: 693.903809 | Mass Diff. -0.002319 Da
Percolator Score: 1.00 | Hyperscore: 26.42 | Spectral Angle: 0.00 | Pearson's Correlation: 0.00
y1
y2
y3
y4
Dimethyl labels not supported by Prosit
Intensity
m/z
1.0
0.5
0.0
−0.5
−1.0
200
400
600
800
1000
1200

**[Ac] − MPPPTGSESLSGLGYAGK | NC_000911.1 : 1779086 : −. pep01**

Charge: +2 | Collision Energy: 27.0 | Percursor m/z: 895.936800 | Mass Diff. 0.000732 Da

Percolator Score: 1.00 | Hyperscore: 24.09 | Spectral Angle: 0.00 | Pearson's Correlation: 0.00

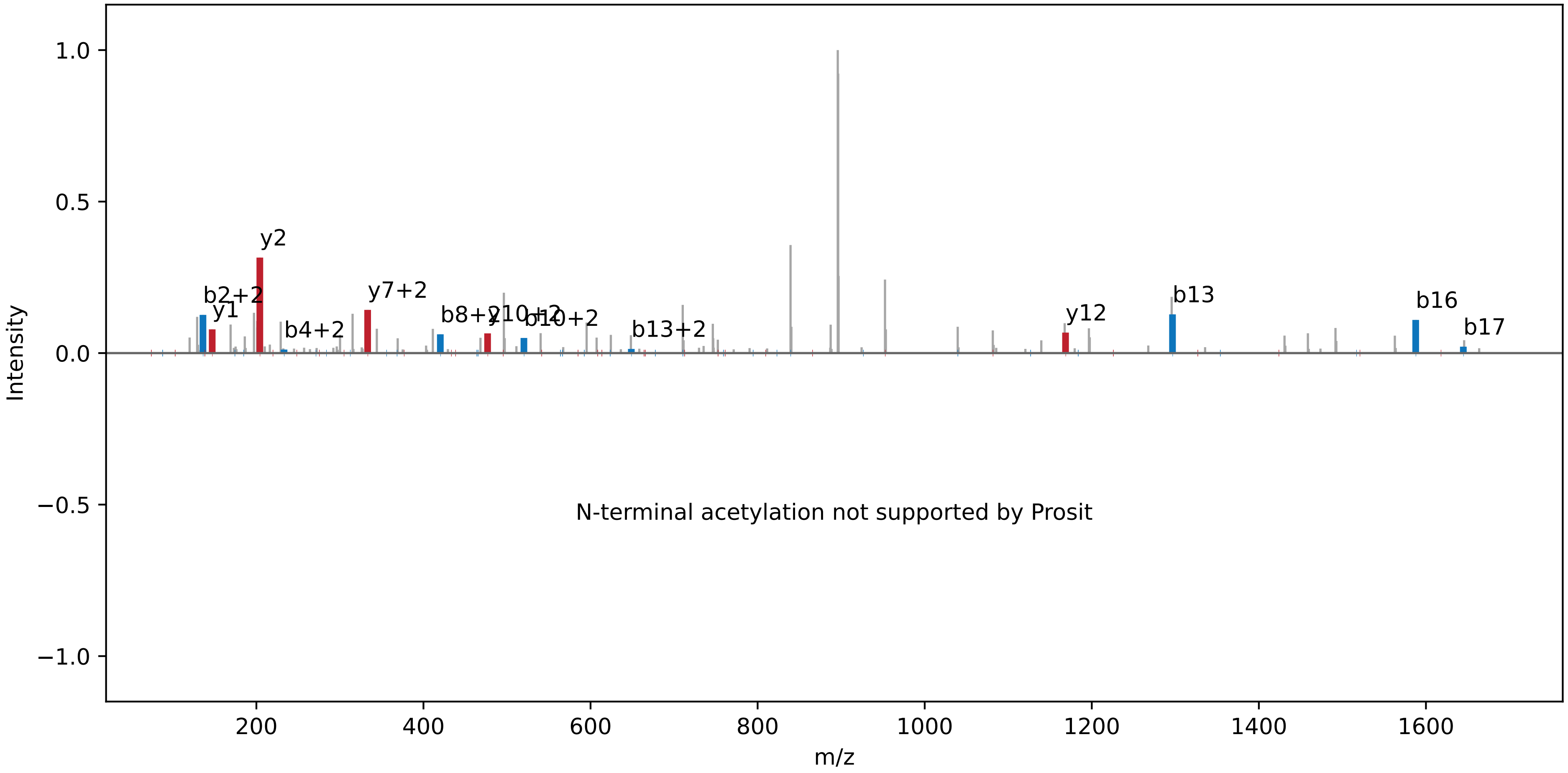

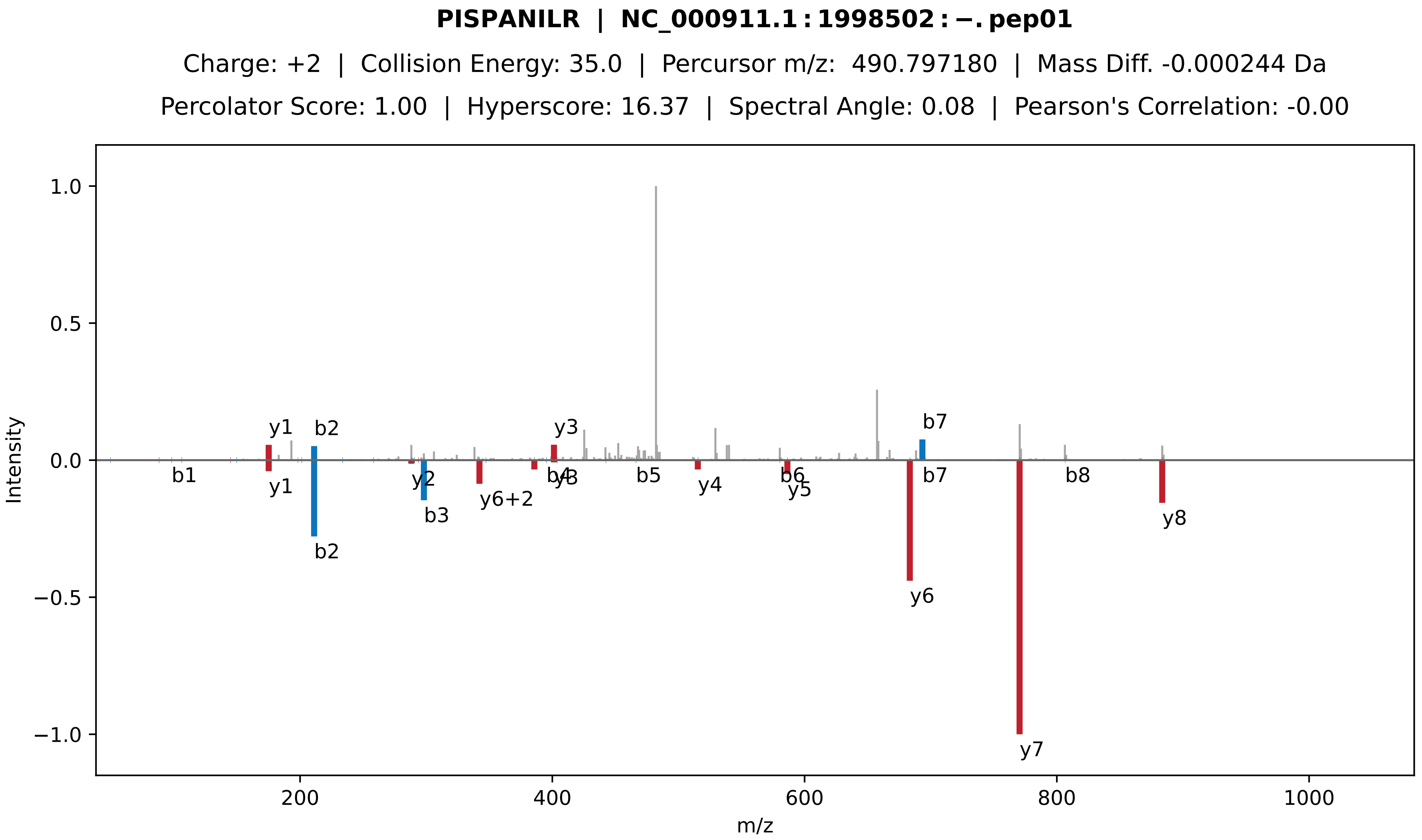

PISPANILR | NC_000911.1 : 1998502 : −. pep01
Charge: +2 | Collision Energy: 35.0 | Percursor m/z: 490.797180 | Mass Diff. -0.000244 Da
Percolator Score: 1.00 | Hyperscore: 16.37 | Spectral Angle: 0.08 | Pearson's Correlation: -0.00
Intensity
m/z
y1
b2
y3
b7
b1
y1
b2
y2
b3
y6+2
b4
y3
b5
y4
b6
y5
y6
b7
y7
b8
y8

**TGQGIGIHR | NC_000911.1 : 2396743 : −. pep01**

Charge: +2 | Collision Energy: 27.0 | Percursor m/z: 469.761993 | Mass Diff. -0.000366 Da

Percolator Score: 1.00 | Hyperscore: 20.05 | Spectral Angle: 0.34 | Pearson's Correlation: 0.36

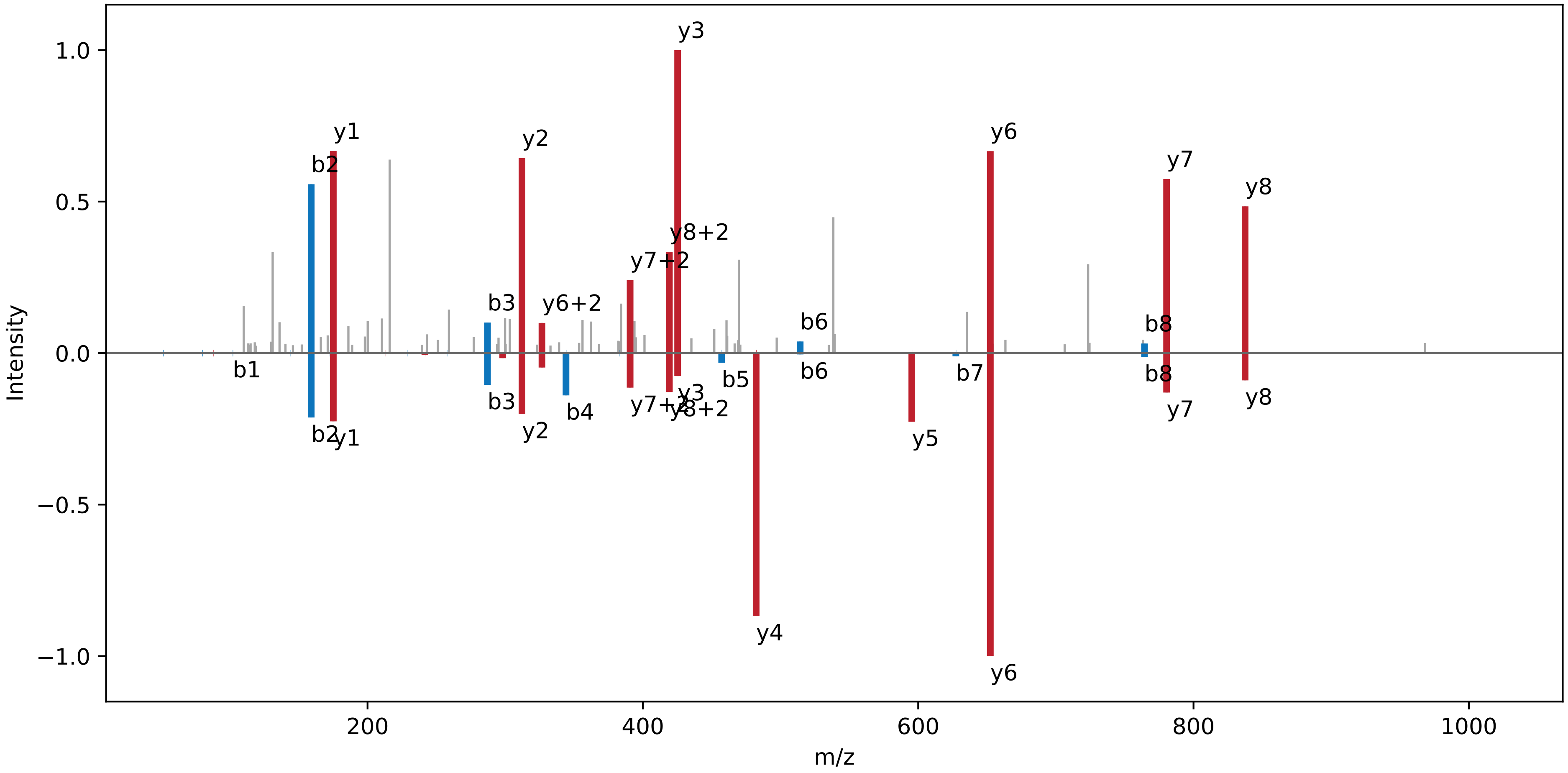

**AVAGFQNR | NC_000911.1 : 2601527 : −. pep01**

Charge: +2 | Collision Energy: 35.0 | Percursor m/z: 431.731720 | Mass Diff. 0.003235 Da

Percolator Score: 1.00 | Hyperscore: 14.17 | Spectral Angle: 0.09 | Pearson's Correlation: 0.02

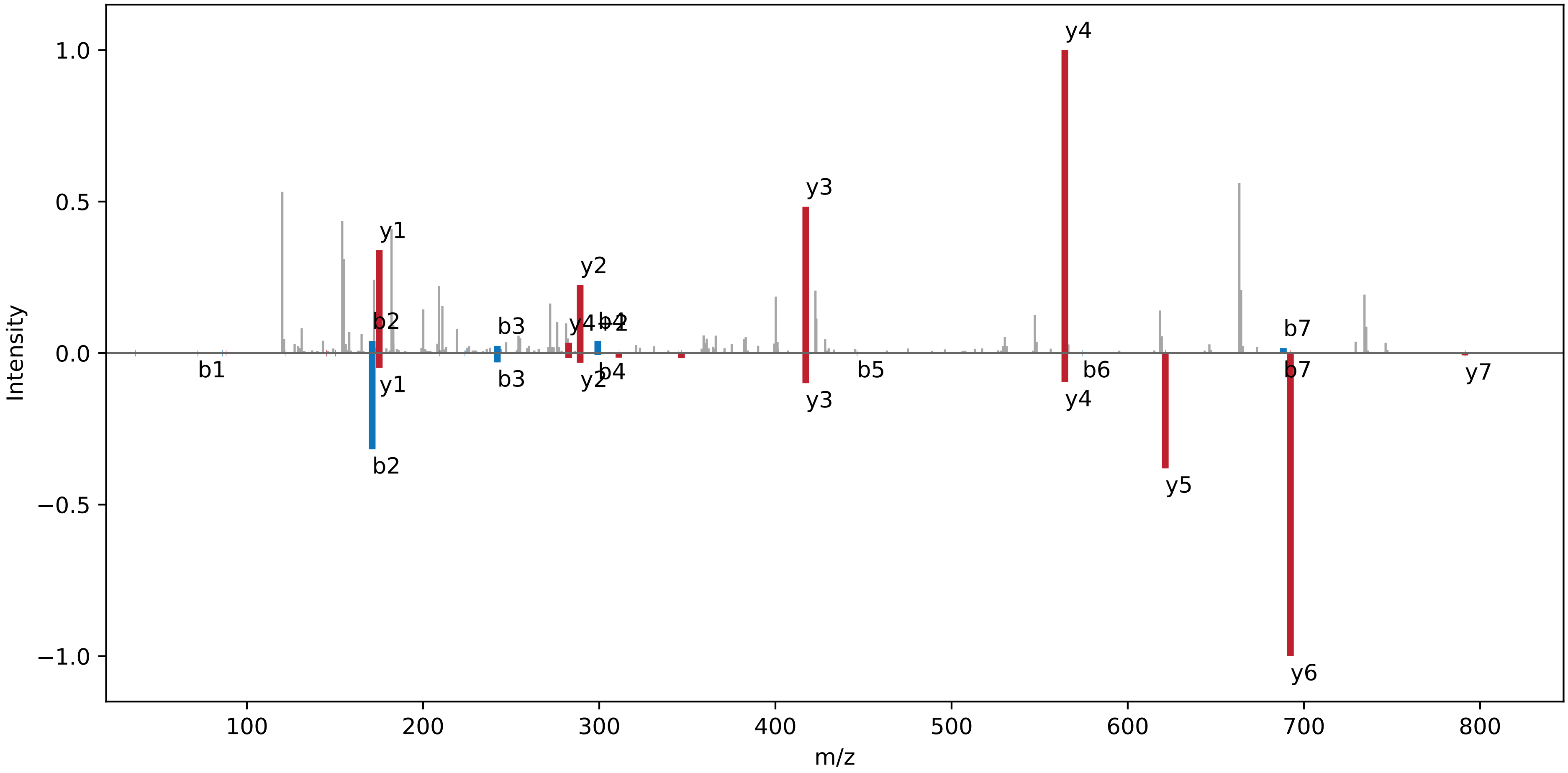

**LAPDPVG | NC_000911.1 : 2814522 : +. pep01**

Charge: +2 | Collision Energy: 27.0 | Percursor m/z: 334.688538 | Mass Diff. 0.009949 Da

Percolator Score: 1.00 | Hyperscore: 11.87 | Spectral Angle: 0.13 | Pearson's Correlation: 0.03

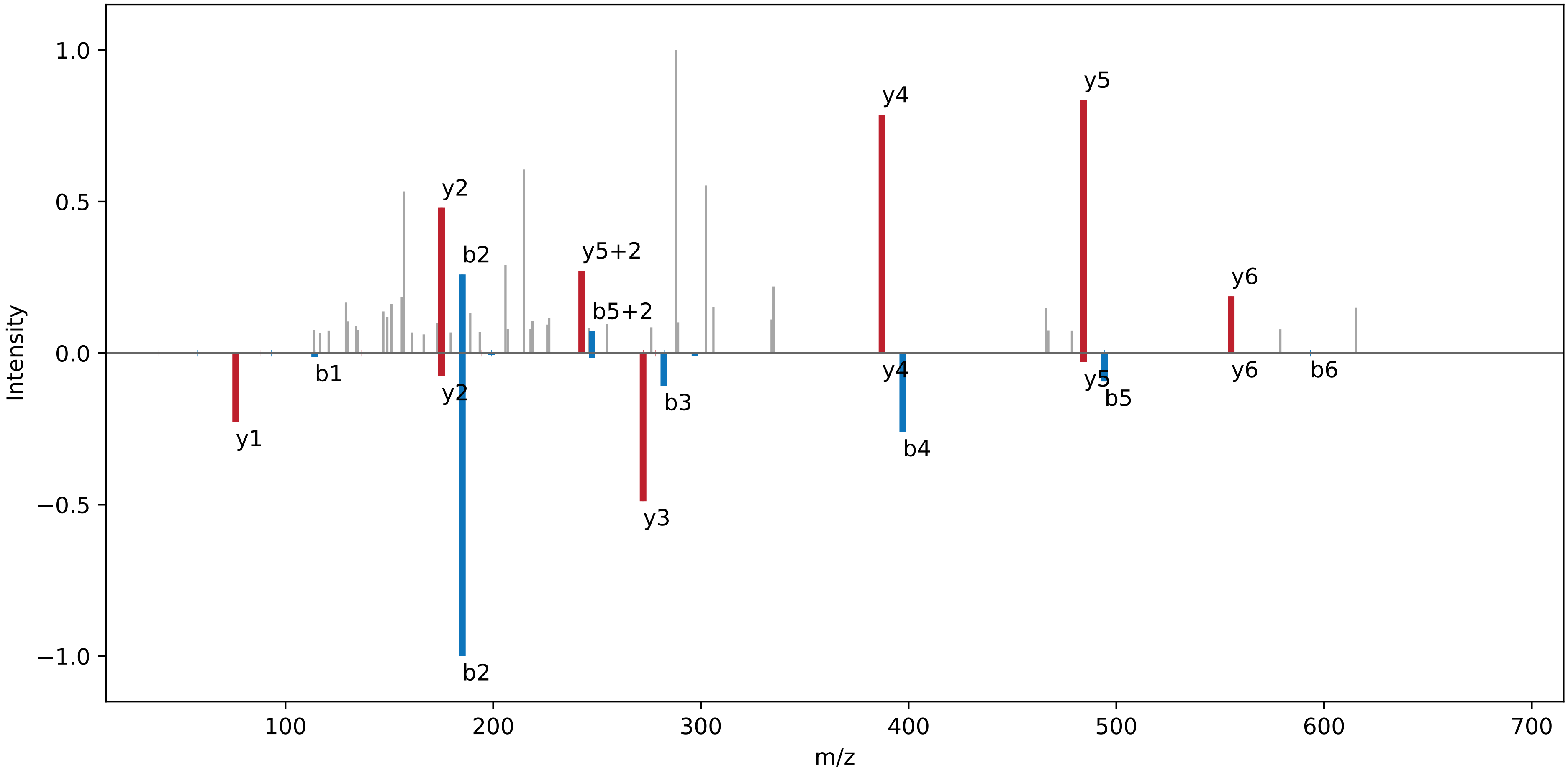

**[DmL] − AMGEILPPSKPLFPK[DmL] | NC_000911.1 : 2943281 : −. pep01**

Charge: +3 | Collision Energy: 28.0 | Percursor m/z: 560.997100 | Mass Diff. 0.001099 Da

Percolator Score: 0.99 | Hyperscore: 23.82 | Spectral Angle: 0.00 | Pearson's Correlation: 0.00

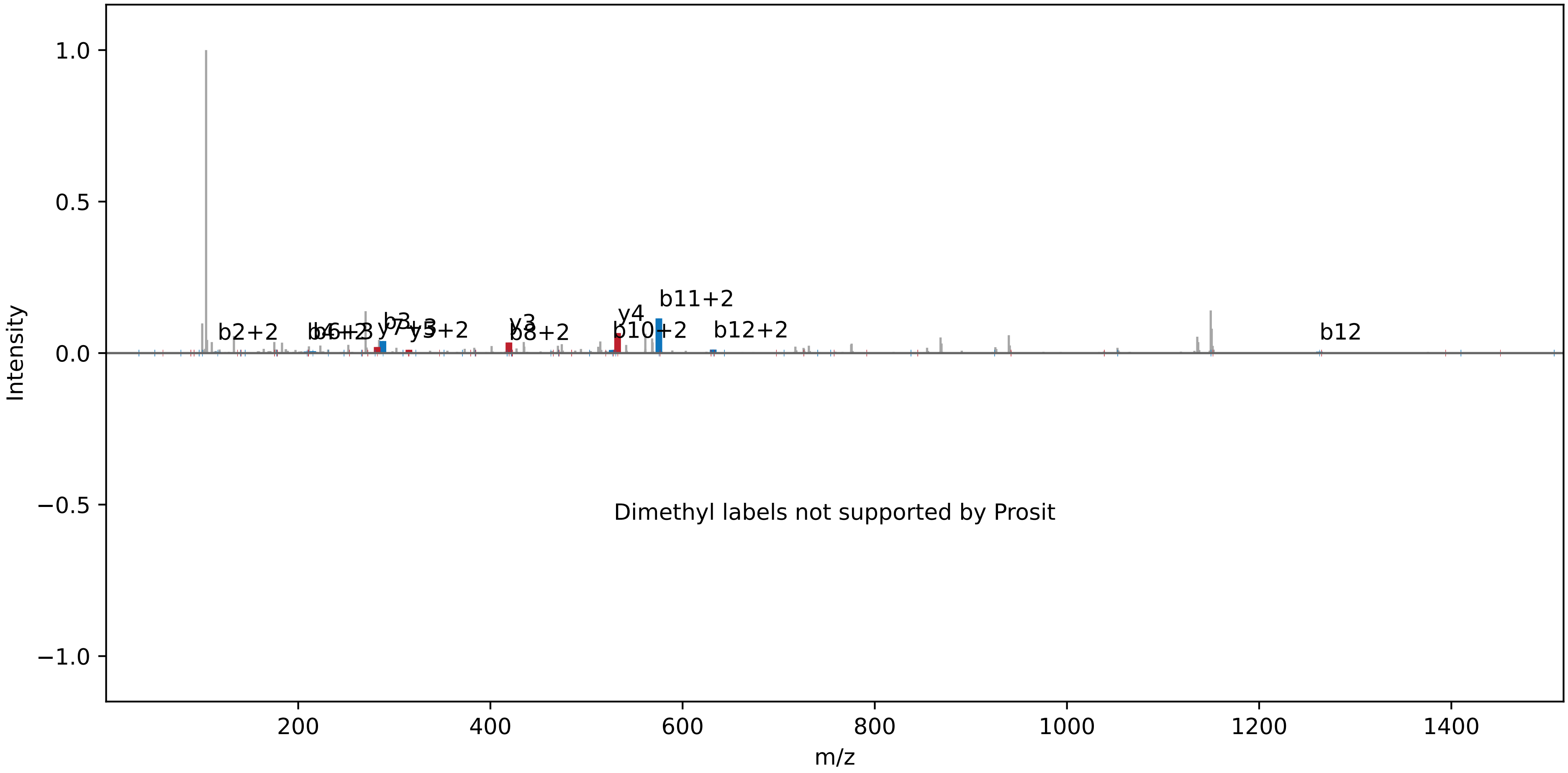

**RISAITAK | NC_000911.1 : 3038111 : −. pep01**

Charge: +2 | Collision Energy: 27.0 | Percursor m/z: 430.271729 | Mass Diff. 0.000000 Da

Percolator Score: 1.00 | Hyperscore: 16.65 | Spectral Angle: 0.81 | Pearson's Correlation: 0.95

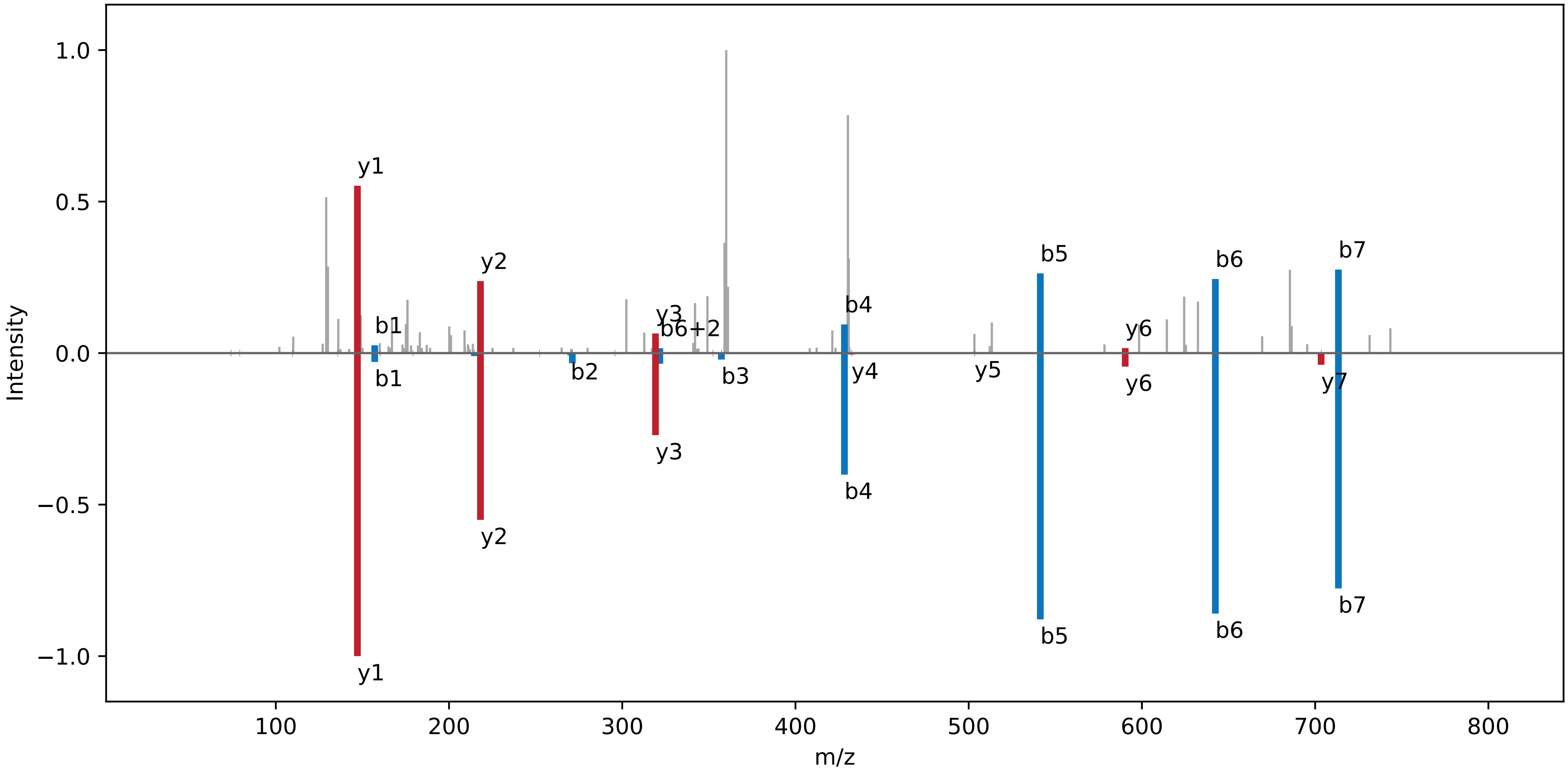

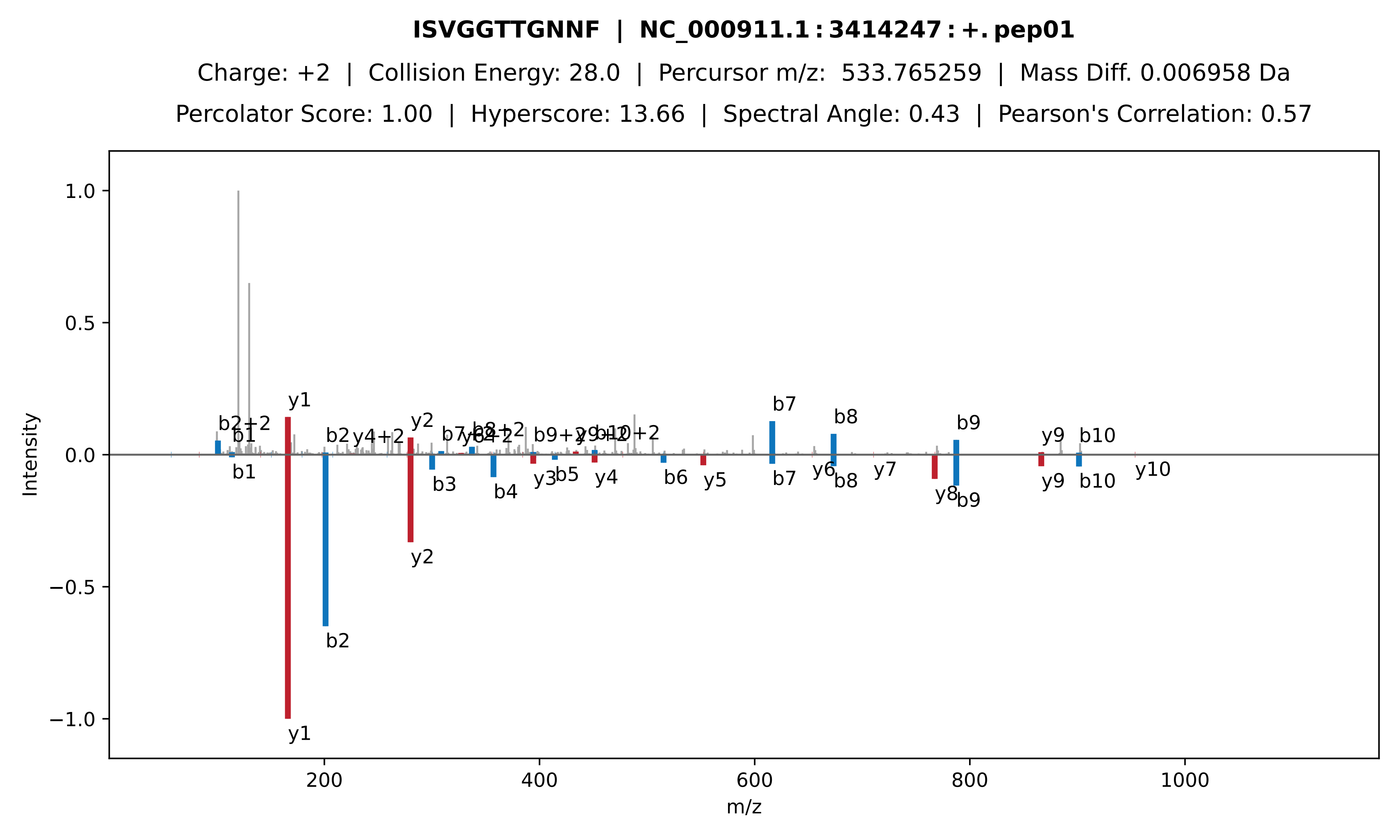

ISVGGTTGNNF | NC_000911.1:3414247:+.pep01
Charge: +2 | Collision Energy: 28.0 | Percursor m/z: 533.765259 | Mass Diff. 0.006958 Da
Percolator Score: 1.00 | Hyperscore: 13.66 | Spectral Angle: 0.43 | Pearson's Correlation: 0.57
Intensity
m/z

**LKLLLKVLK | NC_000911.1:358889:+.pep01**

Charge: +3 | Collision Energy: 27.0 | Percursor m/z: 356.601898 | Mass Diff. 0.000244 Da

Percolator Score: 1.00 | Hyperscore: 14.95 | Spectral Angle: 0.14 | Pearson's Correlation: 0.09

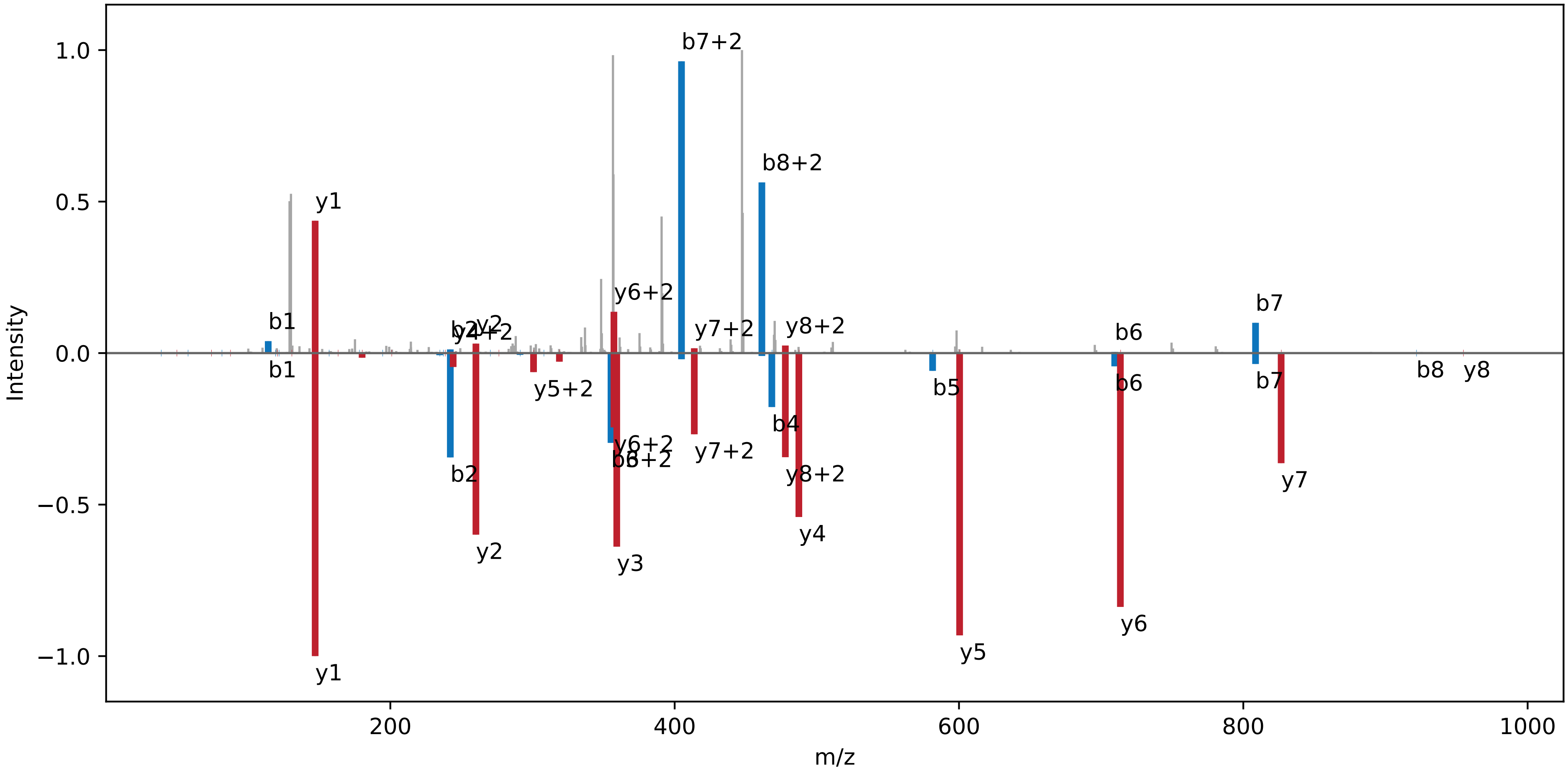

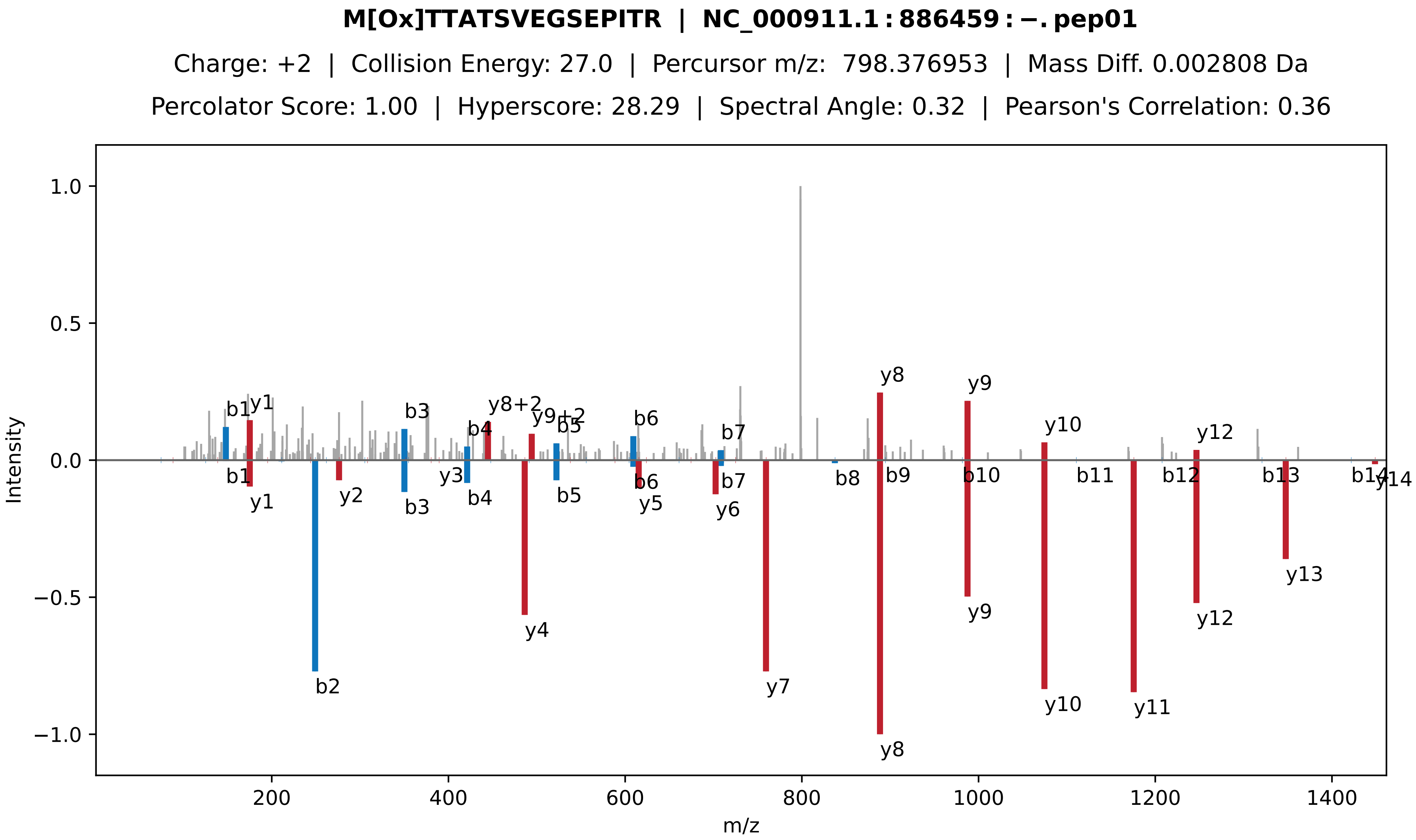

M[Ox]TTATSVEGSEPITR | NC_000911.1:886459:−.pep01
Charge: +2 | Collision Energy: 27.0 | Percursor m/z: 798.376953 | Mass Diff. 0.002808 Da
Percolator Score: 1.00 | Hyperscore: 28.29 | Spectral Angle: 0.32 | Pearson's Correlation: 0.36
Intensity
m/z
b1
y1
b2
y2
b3
y3
b4
y8+2
y4
y9+2
b5
b6
y5
b7
y6
y7
y8
b8
b9
y9
b10
y10
b11
y11
b12
y12
b13
y13
b14
y14

**ASRVGTCNGLAIDPSSTAILWAATM[Ox]AM[Ox]ASVICPLLGVPIDTCVSLIWPK | NC_005229.1 : 35502 : +. pep**

Charge: +6 | Collision Energy: 27.0 | Percursor m/z: 870.788330 | Mass Diff. 3.039551 Da

Percolator Score: 1.00 | Hyperscore: 12.89 | Spectral Angle: 0.00 | Pearson's Correlation: 0.00

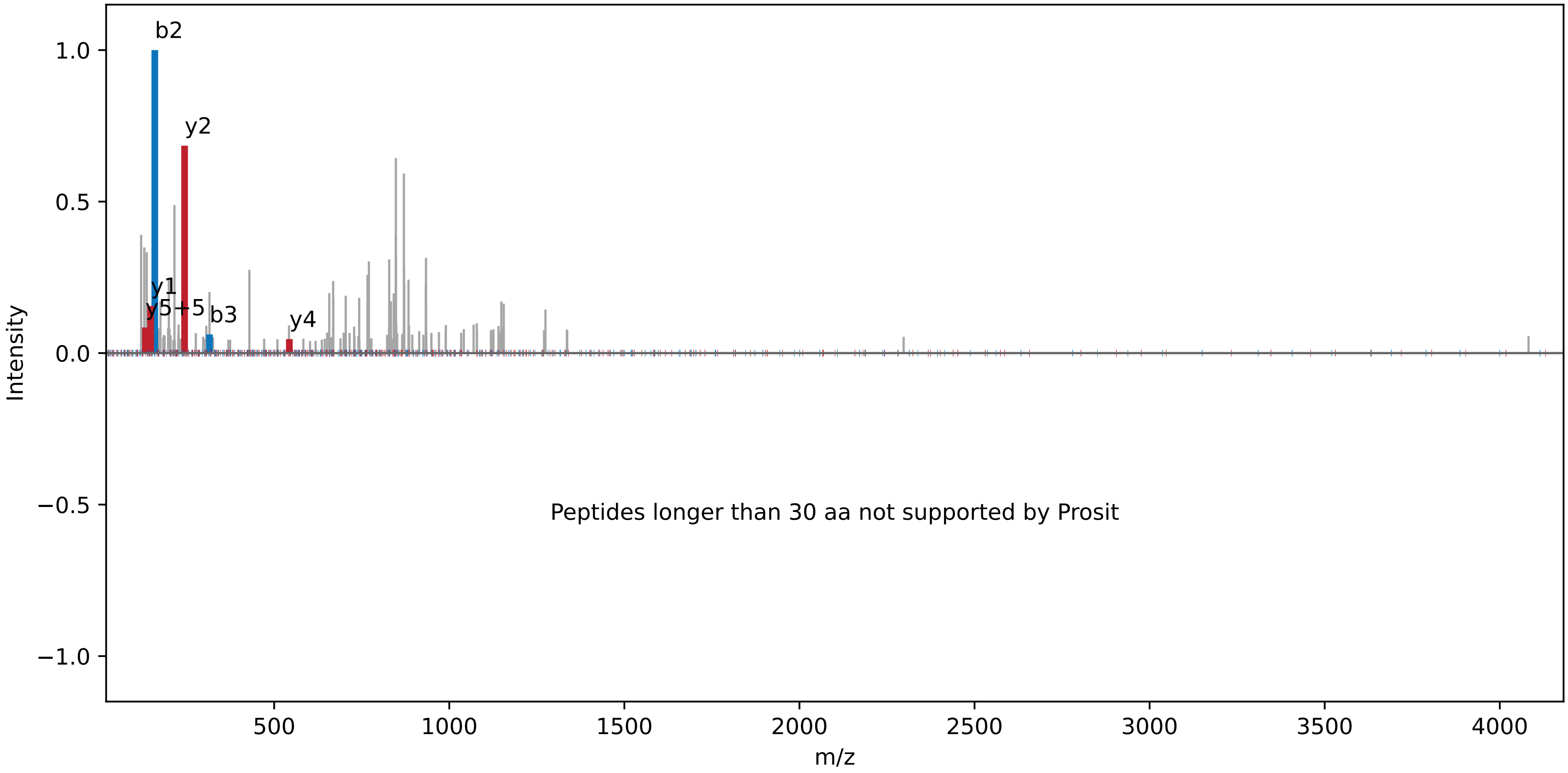

**STACHLLYLVVNIECEVSFCFFQRIIDNLICSIPSNNLLGRSLK | NC_005232.1 : 50265 : −. pep01**

Charge: +4 | Collision Energy: 27.0 | Percursor m/z: 1304.678589 | Mass Diff. 2.045898 Da

Percolator Score: 1.00 | Hyperscore: 27.80 | Spectral Angle: 0.00 | Pearson's Correlation: 0.00

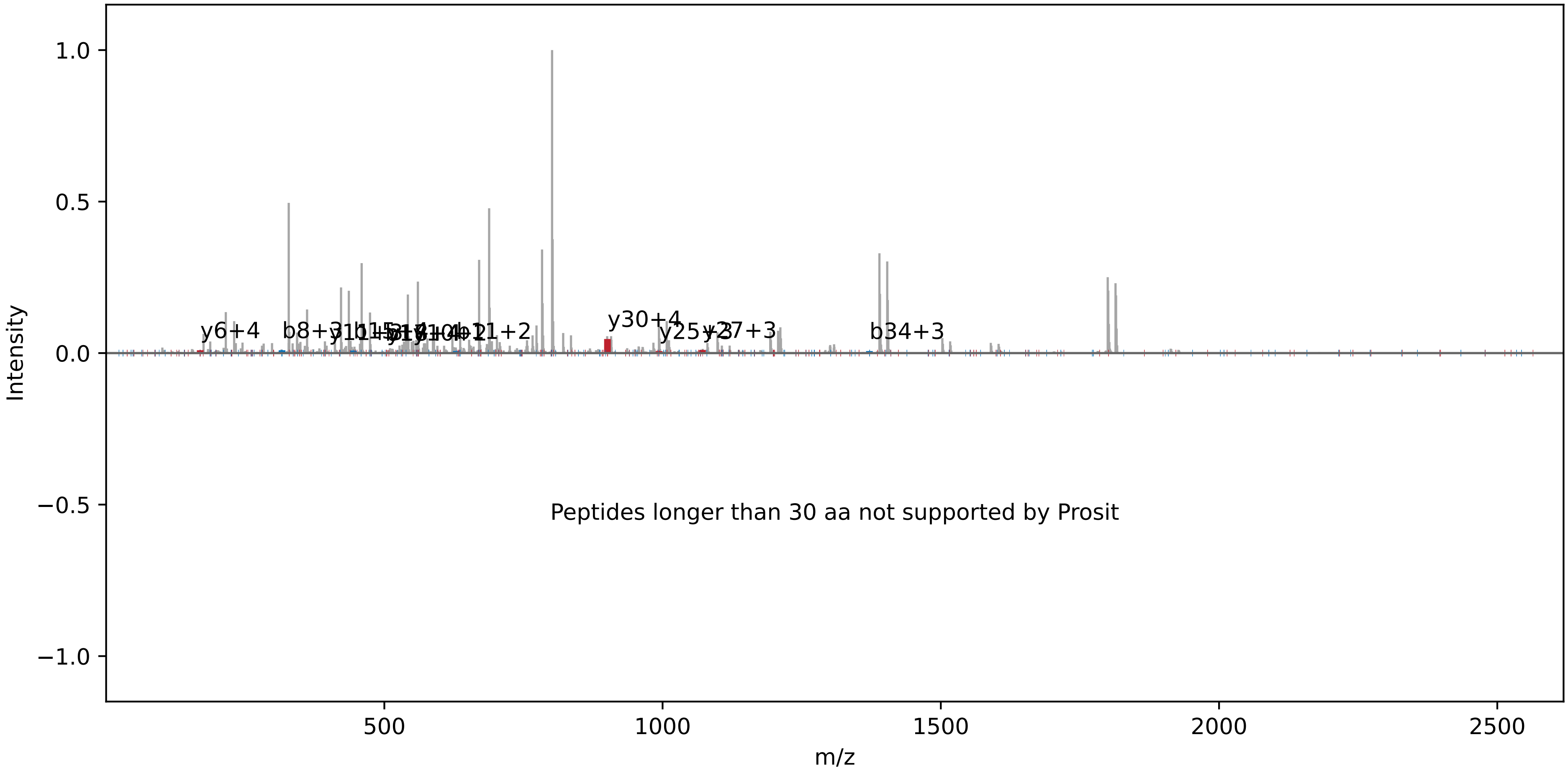
