## Supplementary Dataset 2 for "A revised genome annotation of the model cyanobacterium *Synechocystis* based on start and stop codon-enriched ribosome profiling and proteogenomics"

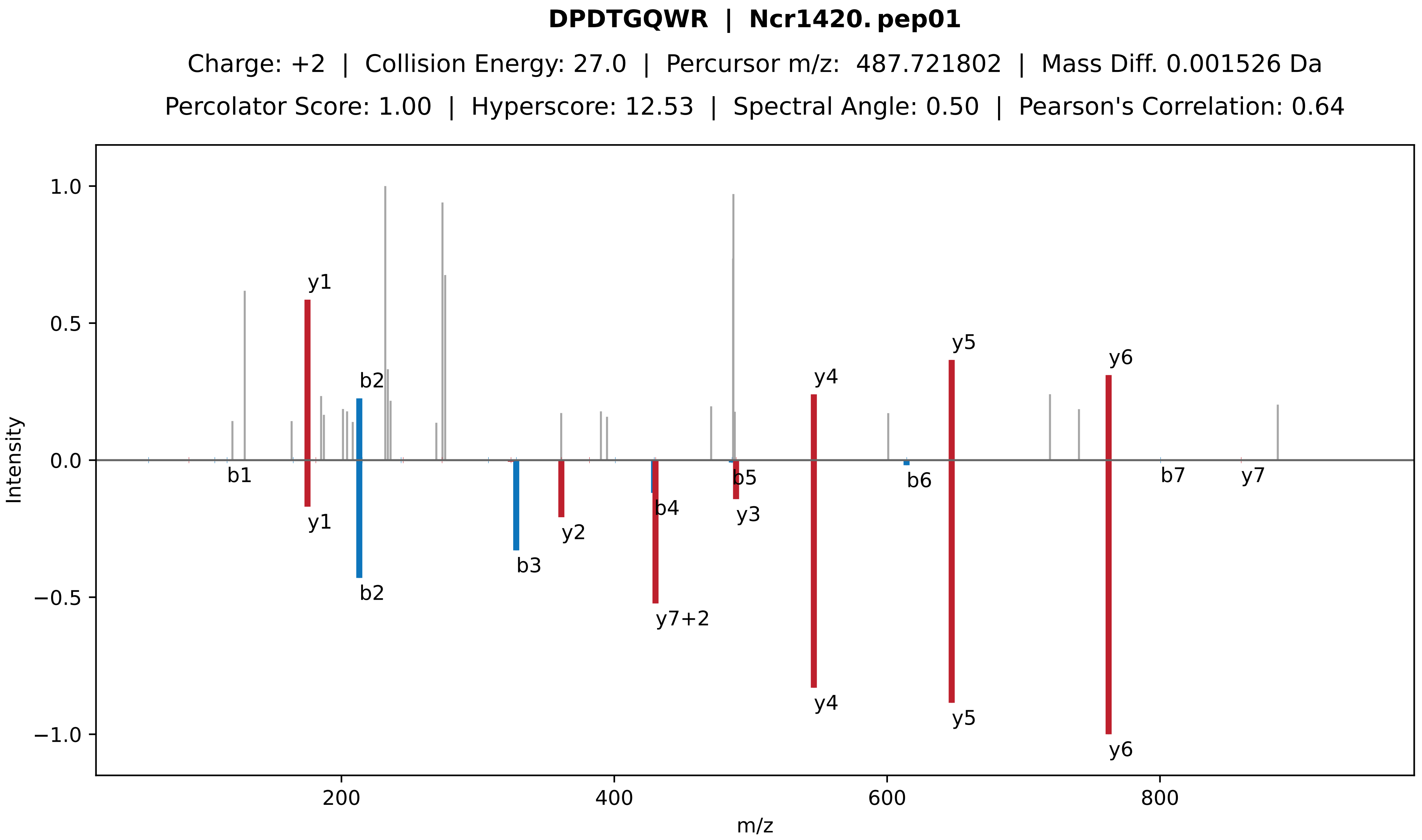
DPDTGQWR | Ncr1420. pep01
Charge: +2 | Collision Energy: 27.0 | Percursor m/z: 487.721802 | Mass Diff. 0.001526 Da
Percolator Score: 1.00 | Hyperscore: 12.53 | Spectral Angle: 0.50 | Pearson's Correlation: 0.64
Intensity
m/z
1.0
0.5
0.0
−0.5
−1.0
200
400
600
800
y1
b2
y4
y5
y6
b1
y1
b2
b3
y2
b4
y7+2
b5
y3
y4
b6
y5
y6
b7
y7

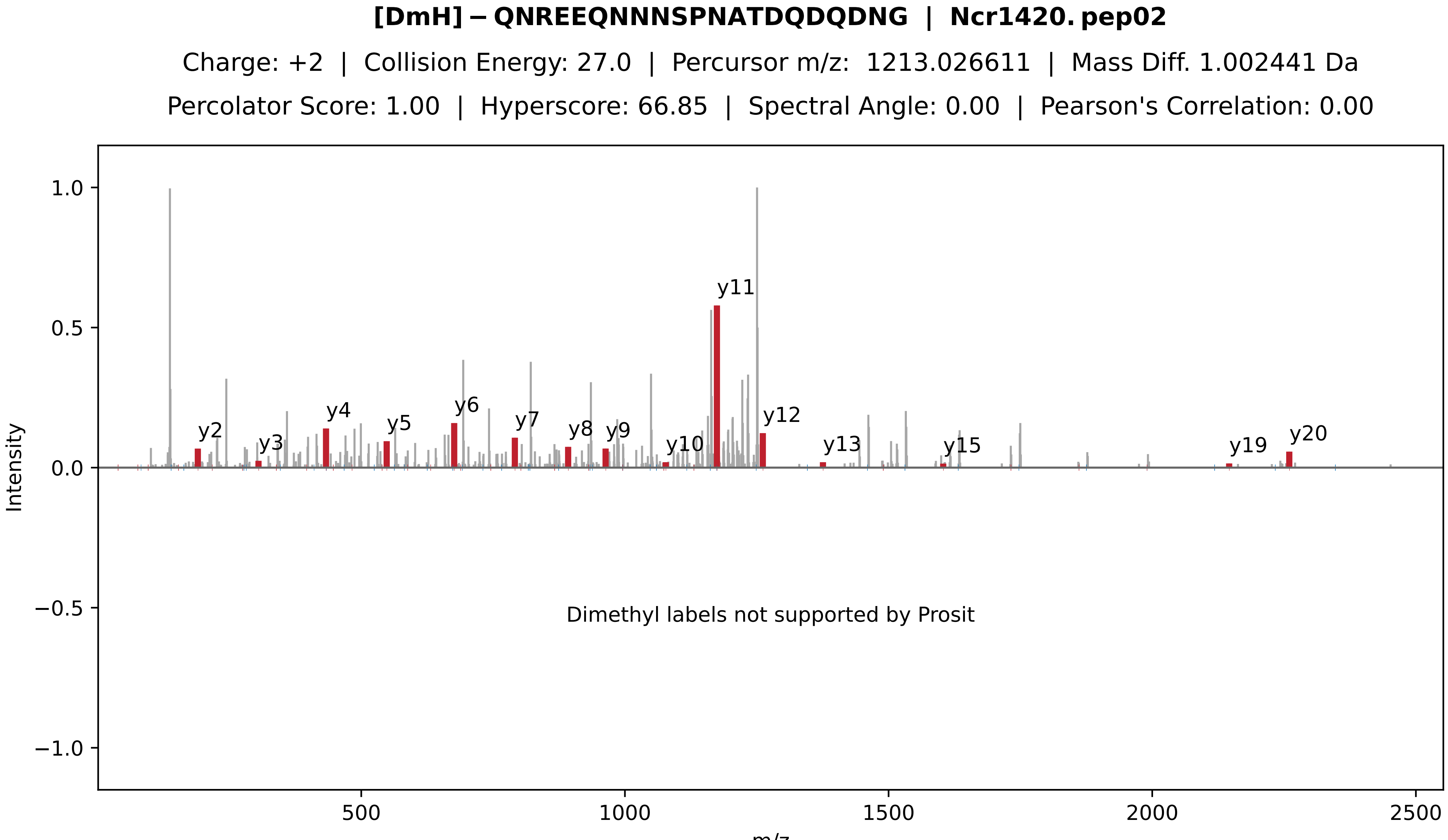

[DmH]−QNREEQNNNSPNATDQDQDNG | Ncr1420.pep02
Charge: +2 | Collision Energy: 27.0 | Percursor m/z: 1213.026611 | Mass Diff. 1.002441 Da
Percolator Score: 1.00 | Hyperscore: 66.85 | Spectral Angle: 0.00 | Pearson's Correlation: 0.00
Intensity
m/z
y2
y3
y4
y5
y6
y7
y8
y9
y10
y11
y12
y13
y15
y19
y20
Dimethyl labels not supported by Prosit

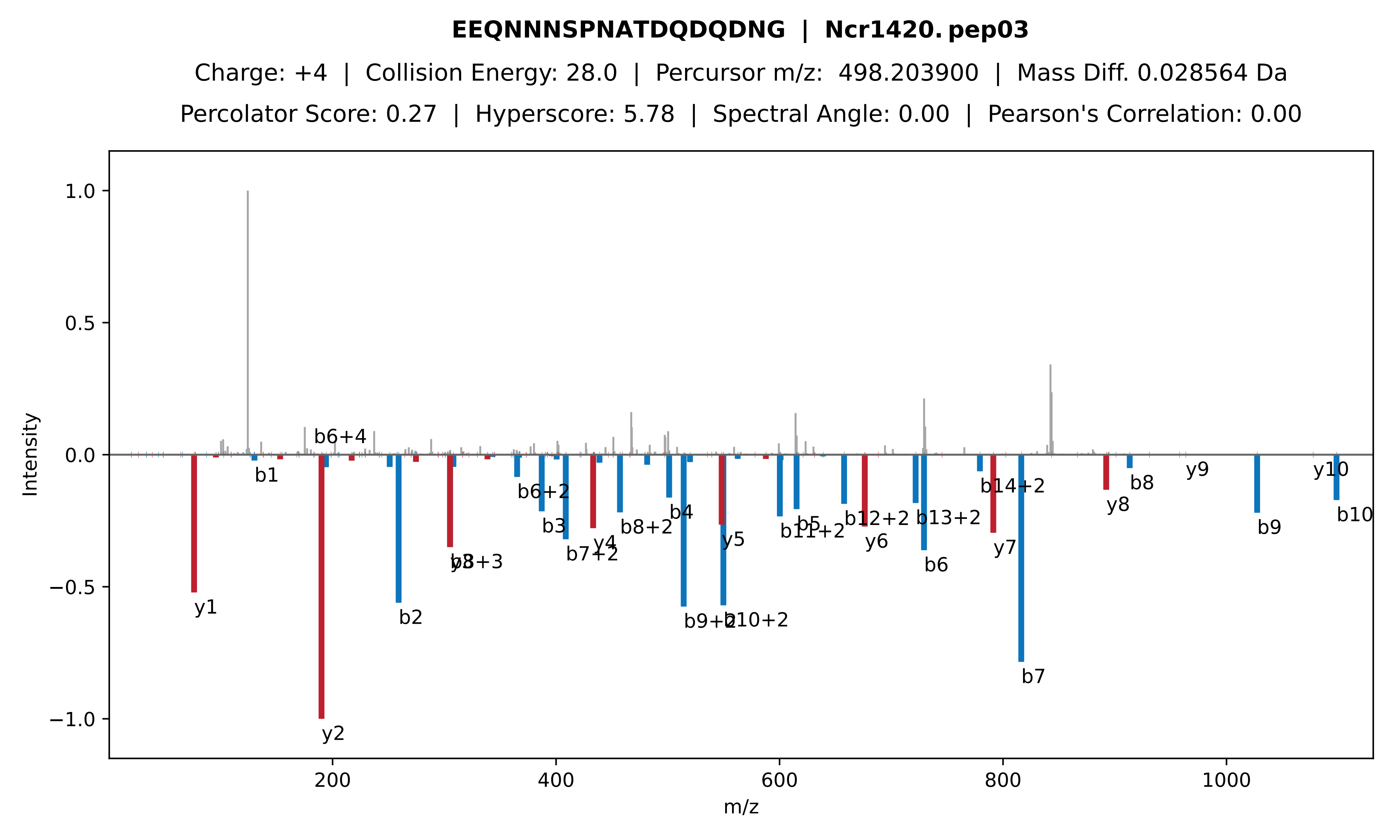

EEQNNNSPNATDQDQDNG | Ncr1420. pep03
Charge: +4 | Collision Energy: 28.0 | Percursor m/z: 498.203900 | Mass Diff. 0.028564 Da
Percolator Score: 0.27 | Hyperscore: 5.78 | Spectral Angle: 0.00 | Pearson's Correlation: 0.00
Intensity
m/z
1.0
0.5
0.0
−0.5
−1.0
200
400
600
800
1000
b6+4
b1
y1
y2
b2
b8+3
b6+2
b3
b7+2
y4
b8+2
b4
b9+2
b10+2
y5
b11+2
b5
b12+2
y6
b13+2
b6
b14+2
y7
b7
y8
b8
y9
b9
y10
b10

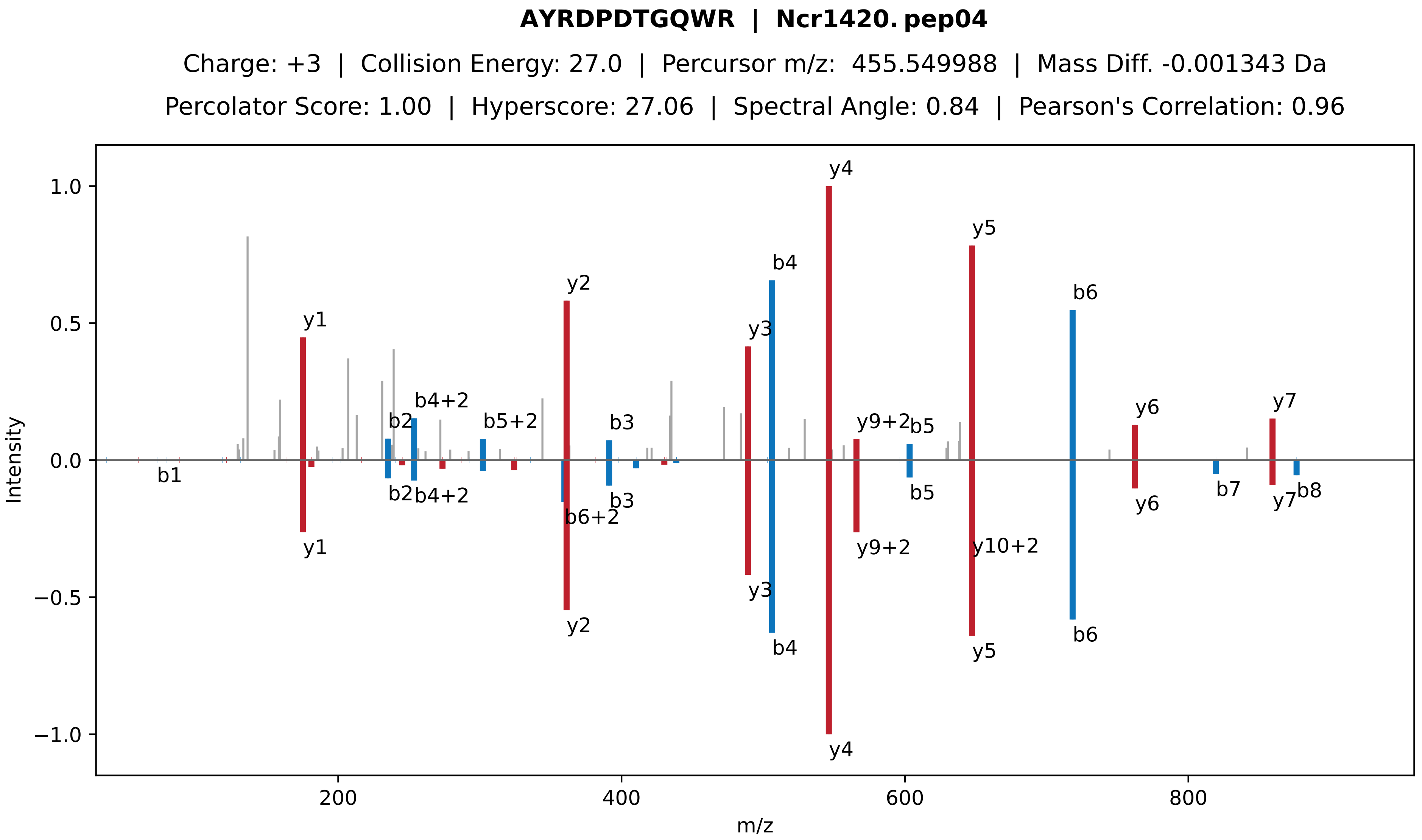

AYRDPDTGQWR | Ncr1420. pep04
Charge: +3 | Collision Energy: 27.0 | Percursor m/z: 455.549988 | Mass Diff. -0.001343 Da
Percolator Score: 1.00 | Hyperscore: 27.06 | Spectral Angle: 0.84 | Pearson's Correlation: 0.96
Intensity
m/z
y1
b1
b2
b4+2
b5+2
y2
b6+2
b3
y3
b4
y4
y9+2
b5
y5
y10+2
b6
y6
b7
y7
b8

**[DmH] − IAK[DmH]QNREEQNNNSPNATDQDQDNG | Ncr1420. pep05**

Charge: +3 | Collision Energy: 28.0 | Percursor m/z: 924.787200 | Mass Diff. 0.011719 Da

Percolator Score: 1.00 | Hyperscore: 69.71 | Spectral Angle: 0.00 | Pearson's Correlation: 0.00

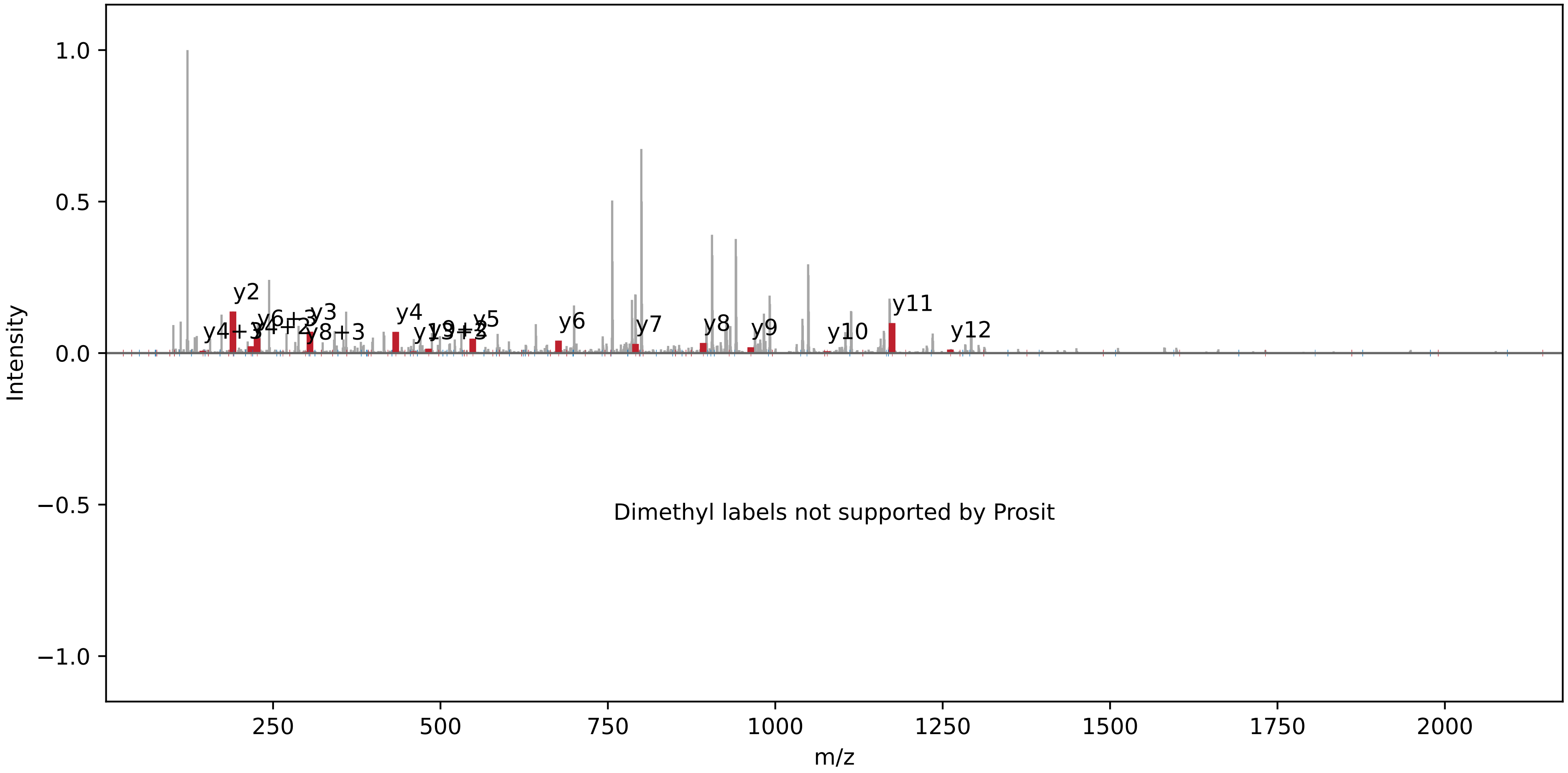

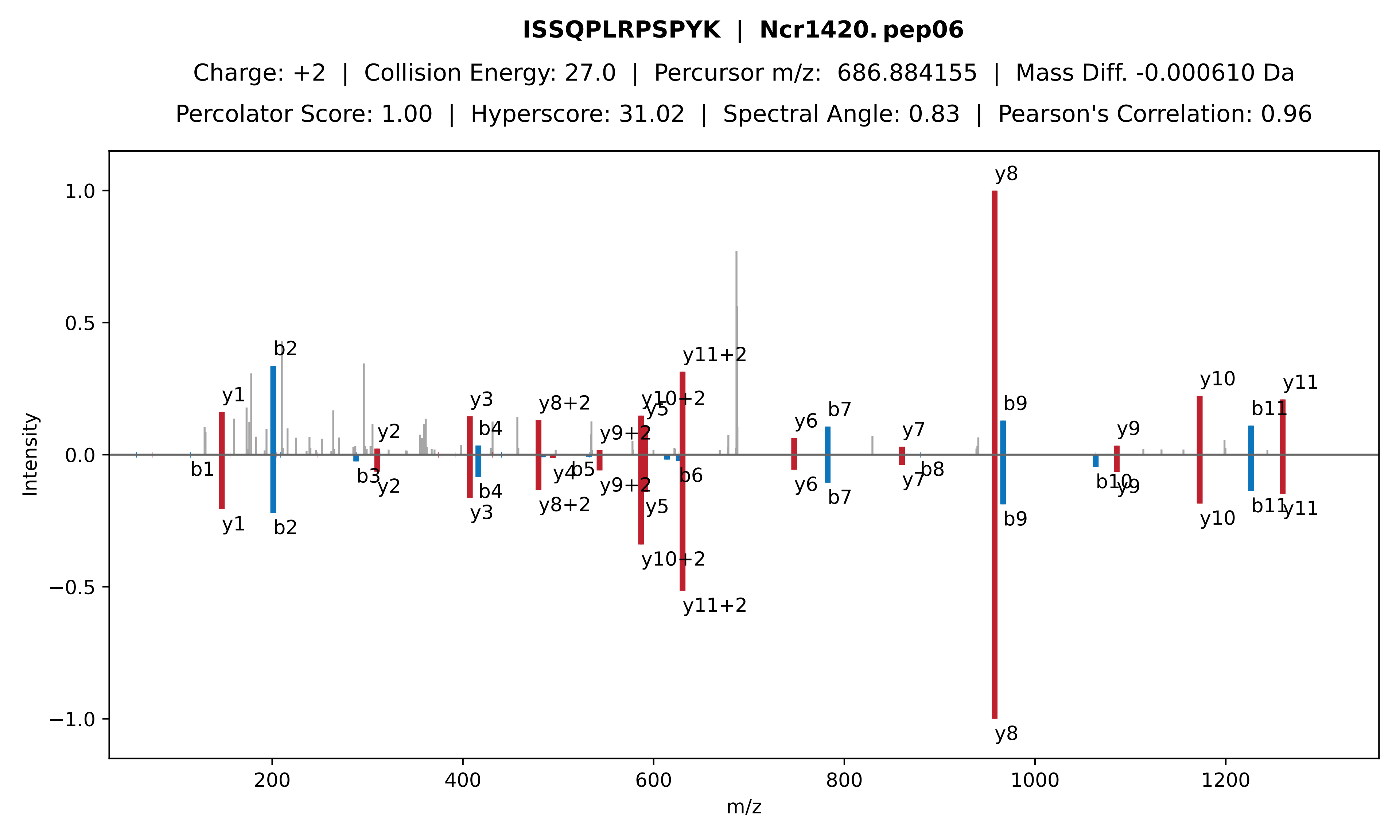
ISSQPLRPSPYK | Ncr1420. pep06
Charge: +2 | Collision Energy: 27.0 | Percursor m/z: 686.884155 | Mass Diff. -0.000610 Da
Percolator Score: 1.00 | Hyperscore: 31.02 | Spectral Angle: 0.83 | Pearson's Correlation: 0.96
Intensity
m/z
1.0
0.5
0.0
−0.5
−1.0
200
400
600
800
1000
1200
y1
b1
b2
b3
y2
y3
b4
y8+2
y4
b5
y9+2
y10+2
y5
y11+2
b6
y6
b7
y7
b8
y8
b9
b10
y9
y10
b11
y11

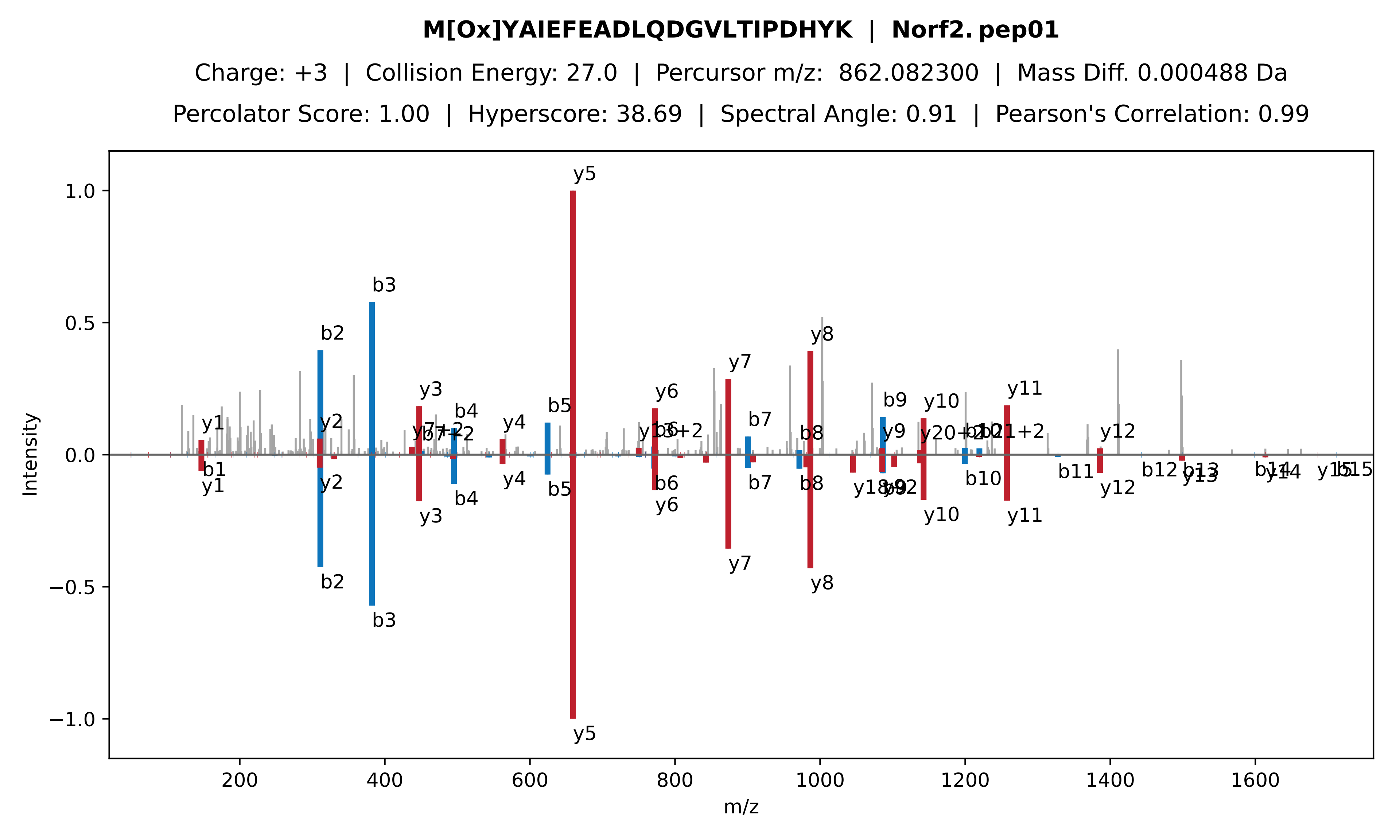

M[Ox]YAIEFEADLQDGVLTIPDHYK | Norf2. pep01
Charge: +3 | Collision Energy: 27.0 | Percursor m/z: 862.082300 | Mass Diff. 0.000488 Da
Percolator Score: 1.00 | Hyperscore: 38.69 | Spectral Angle: 0.91 | Pearson's Correlation: 0.99
Intensity
m/z
1.0
0.5
0.0
−0.5
−1.0
200
400
600
800
1000
1200
1400
1600
y1
b1
y2
b2
b3
y3
b4
y4
b5
y5
y6
b6
y7
b7
y8
b8
b9
y9
y10
b10
y11
b11
y12
b12
b13
y13
y14
b15

**ALSDHSATLIDEWHDPSEDDVWI | Norf2. pep02**

Charge: +3 | Collision Energy: 27.0 | Percursor m/z: 884.739502 | Mass Diff. 1.014404 Da

Percolator Score: 1.00 | Hyperscore: 32.23 | Spectral Angle: 0.73 | Pearson's Correlation: 0.91

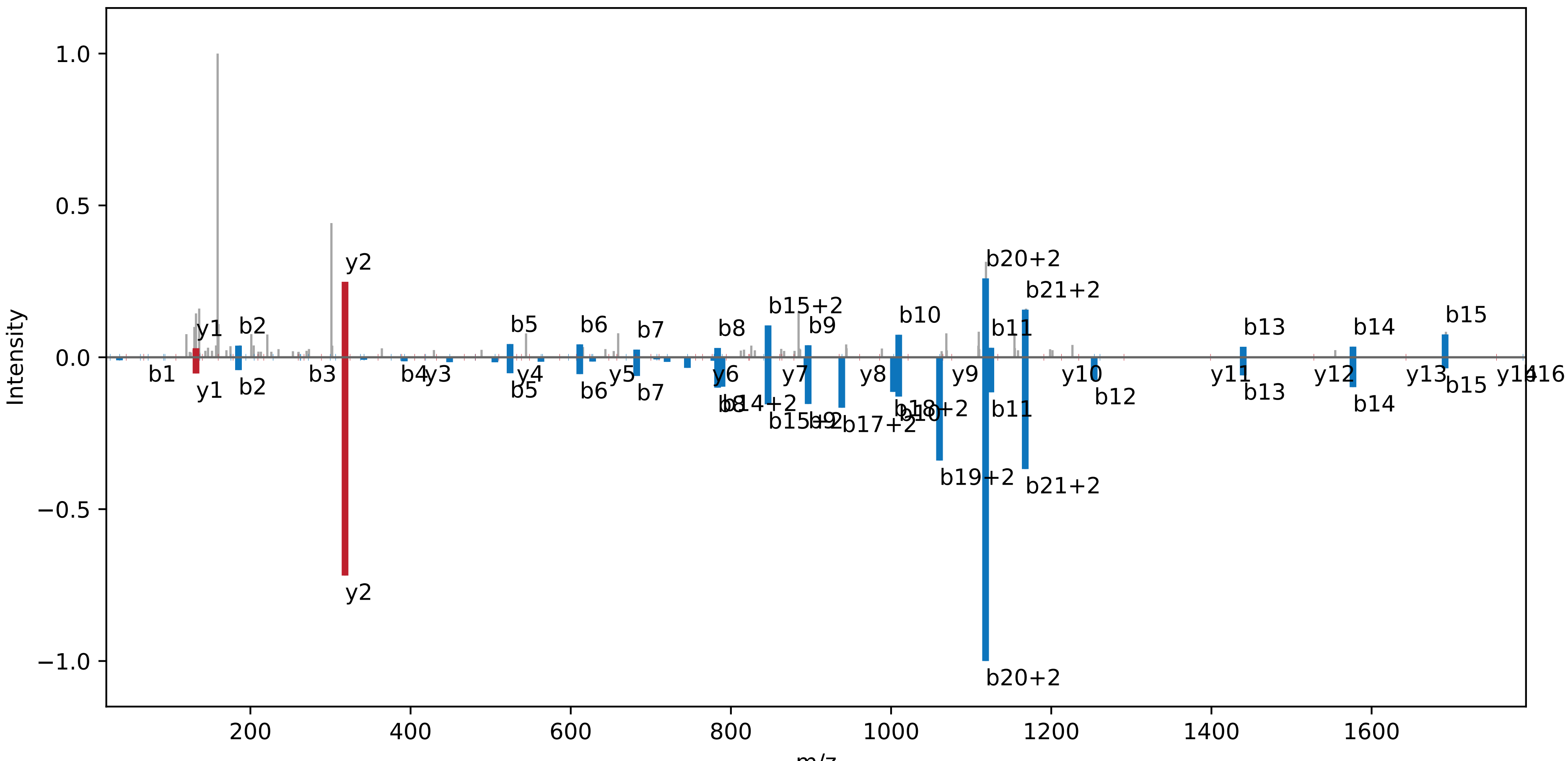

**VVIMLDGHGNELELR | Norf2. pep03**

Charge: +3 | Collision Energy: 27.0 | Percursor m/z: 565.634644 | Mass Diff. -0.000977 Da

Percolator Score: 1.00 | Hyperscore: 37.94 | Spectral Angle: 0.63 | Pearson's Correlation: 0.81

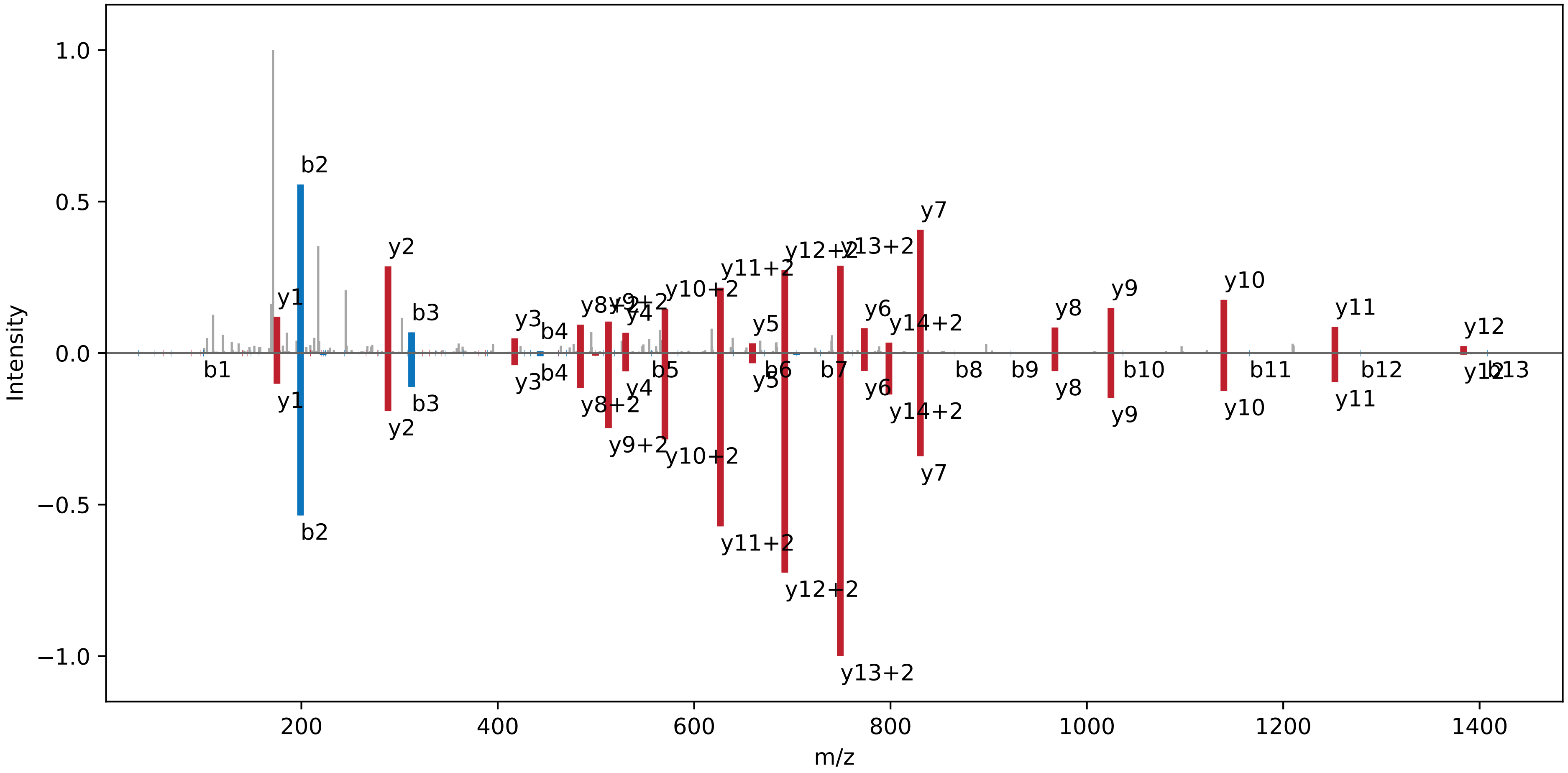

**YAIEFEADLQDGVLTIPDHYK | Norf2. pep04**

Charge: +3 | Collision Energy: 27.0 | Percursor m/z: 813.068420 | Mass Diff. 0.004883 Da

Percolator Score: 1.00 | Hyperscore: 20.83 | Spectral Angle: 0.81 | Pearson's Correlation: 0.96

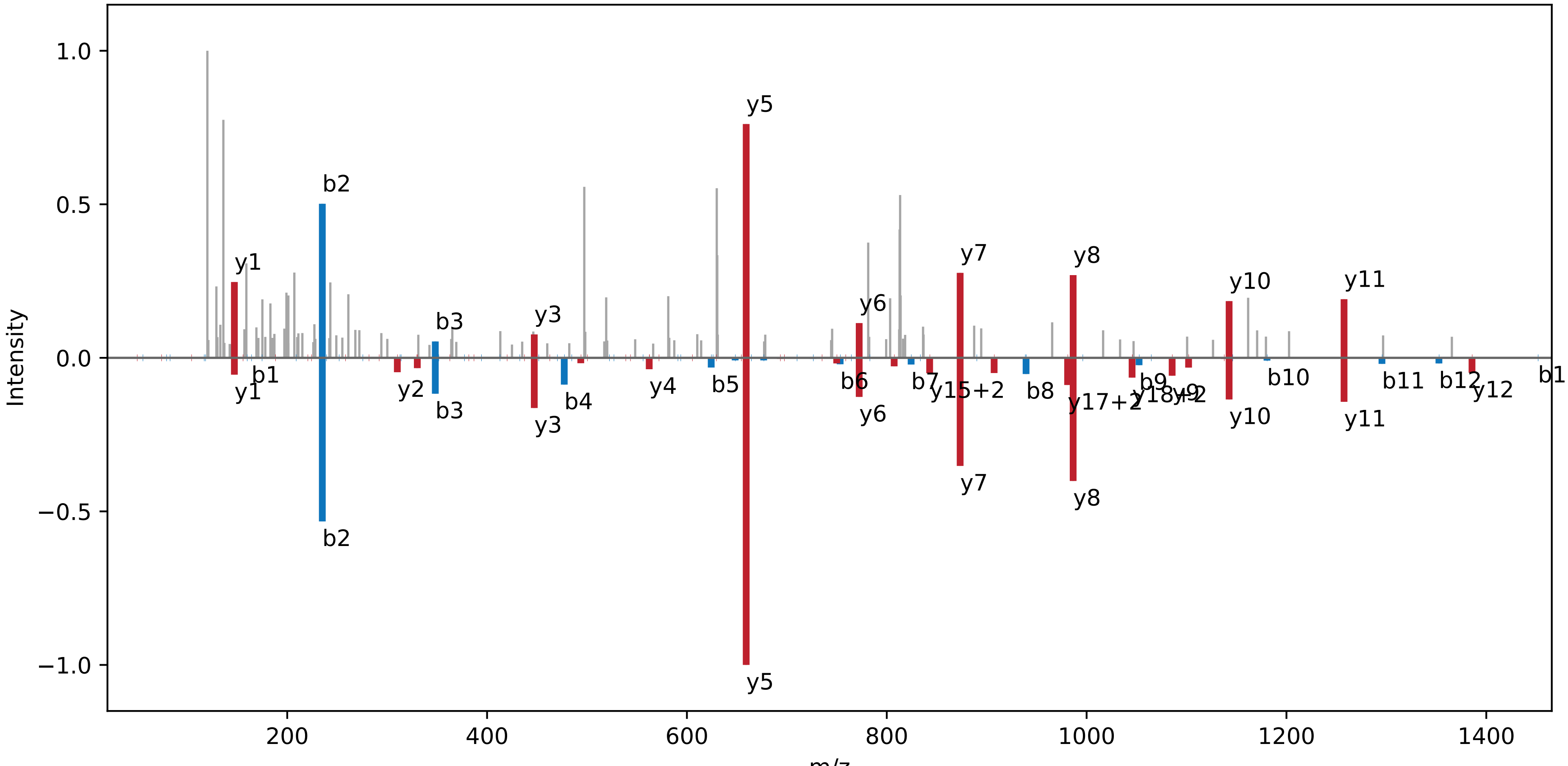

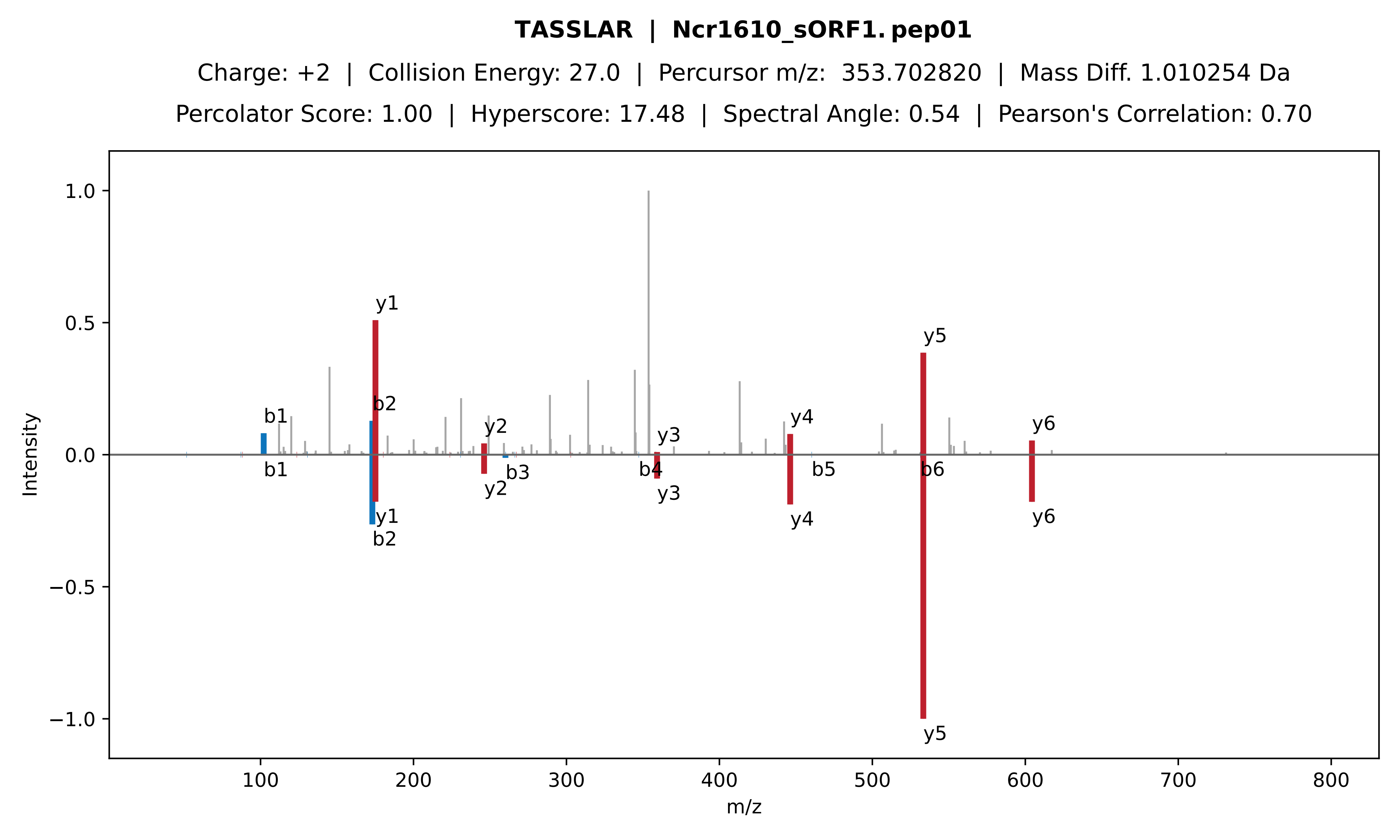

TASSLAR | Ncr1610_sORF1. pep01
Charge: +2 | Collision Energy: 27.0 | Percursor m/z: 353.702820 | Mass Diff. 1.010254 Da
Percolator Score: 1.00 | Hyperscore: 17.48 | Spectral Angle: 0.54 | Pearson's Correlation: 0.70
Intensity
m/z
1.0
0.5
0.0
−0.5
−1.0
100
200
300
400
500
600
700
800
b1
y1
b2
y2
y3
y4
y5
y6
b3
b4
b5
b6

**VYHLTPVVVA | Ncr1610_sORF1. pep02**

Charge: +2 | Collision Energy: 27.0 | Percursor m/z: 549.321411 | Mass Diff. -0.000244 Da

Percolator Score: 1.00 | Hyperscore: 17.24 | Spectral Angle: 0.64 | Pearson's Correlation: 0.82

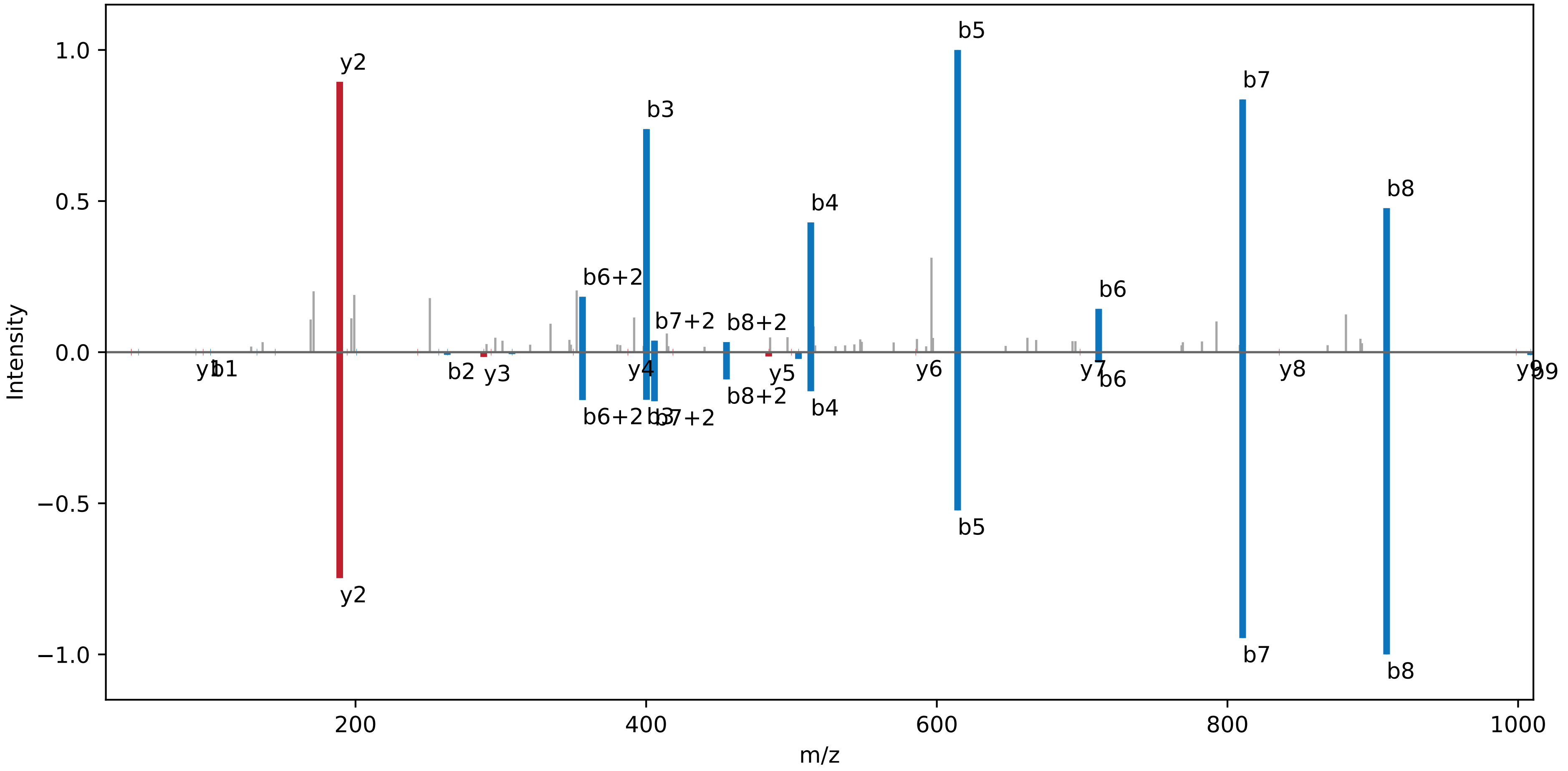

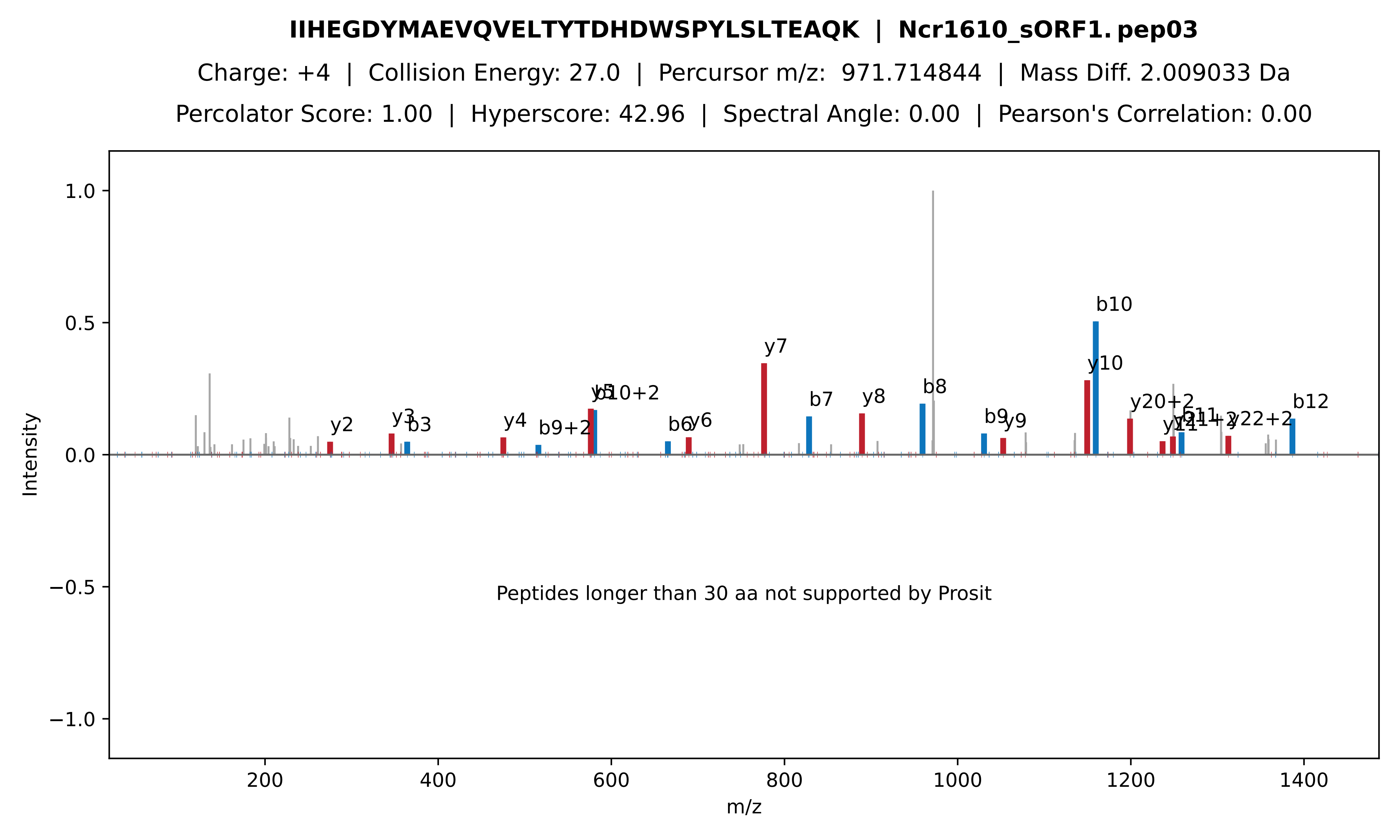
IIHEGDYMAEVQVELTYTDHDWSPYLSLTEAQK | Ncr1610_sORF1. pep03
Charge: +4 | Collision Energy: 27.0 | Percursor m/z: 971.714844 | Mass Diff. 2.009033 Da
Percolator Score: 1.00 | Hyperscore: 42.96 | Spectral Angle: 0.00 | Pearson's Correlation: 0.00
Intensity
m/z
1.0
0.5
0.0
−0.5
−1.0
200
400
600
800
1000
1200
1400
y2
y3
b3
y4
b9+2
b10+2
b6
y6
y7
b7
y8
b8
b9
y9
y10
b10
y20+2
y22+2
b12
Peptides longer than 30 aa not supported by Prosit

**LIEGLLGPQGEPEPELIPIPVNDRR | Sml0012. pep01**

Charge: +3 | Collision Energy: 35.0 | Percursor m/z: 918.172485 | Mass Diff. 1.000977 Da

Percolator Score: 1.00 | Hyperscore: 28.50 | Spectral Angle: 0.05 | Pearson's Correlation: 0.05

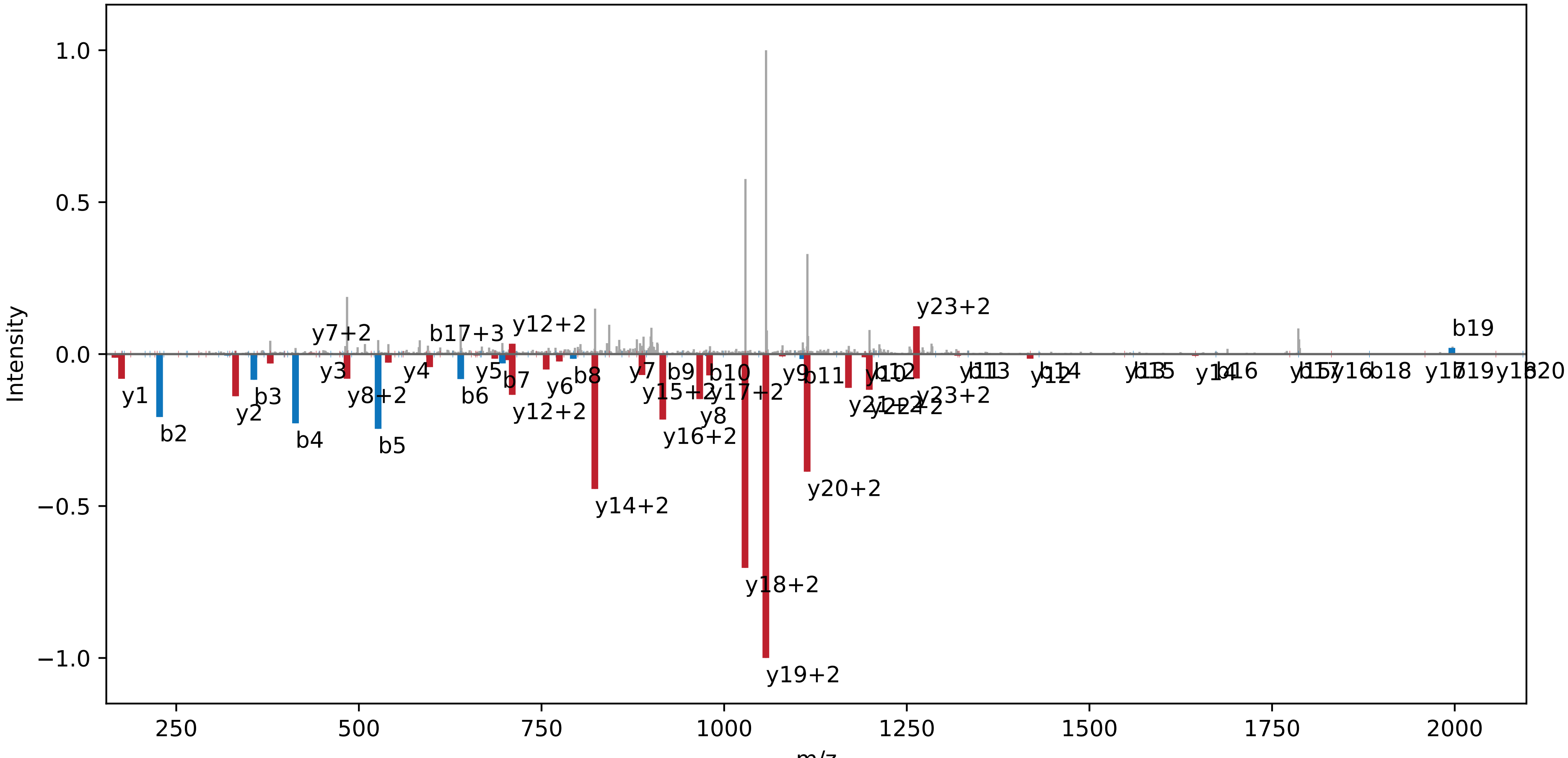

**GGVAIGK | rsbU − as. pep01**

Charge: +2 | Collision Energy: 27.0 | Percursor m/z: 301.186920 | Mass Diff. -0.000061 Da

Percolator Score: 1.00 | Hyperscore: 10.66 | Spectral Angle: 0.39 | Pearson's Correlation: 0.49

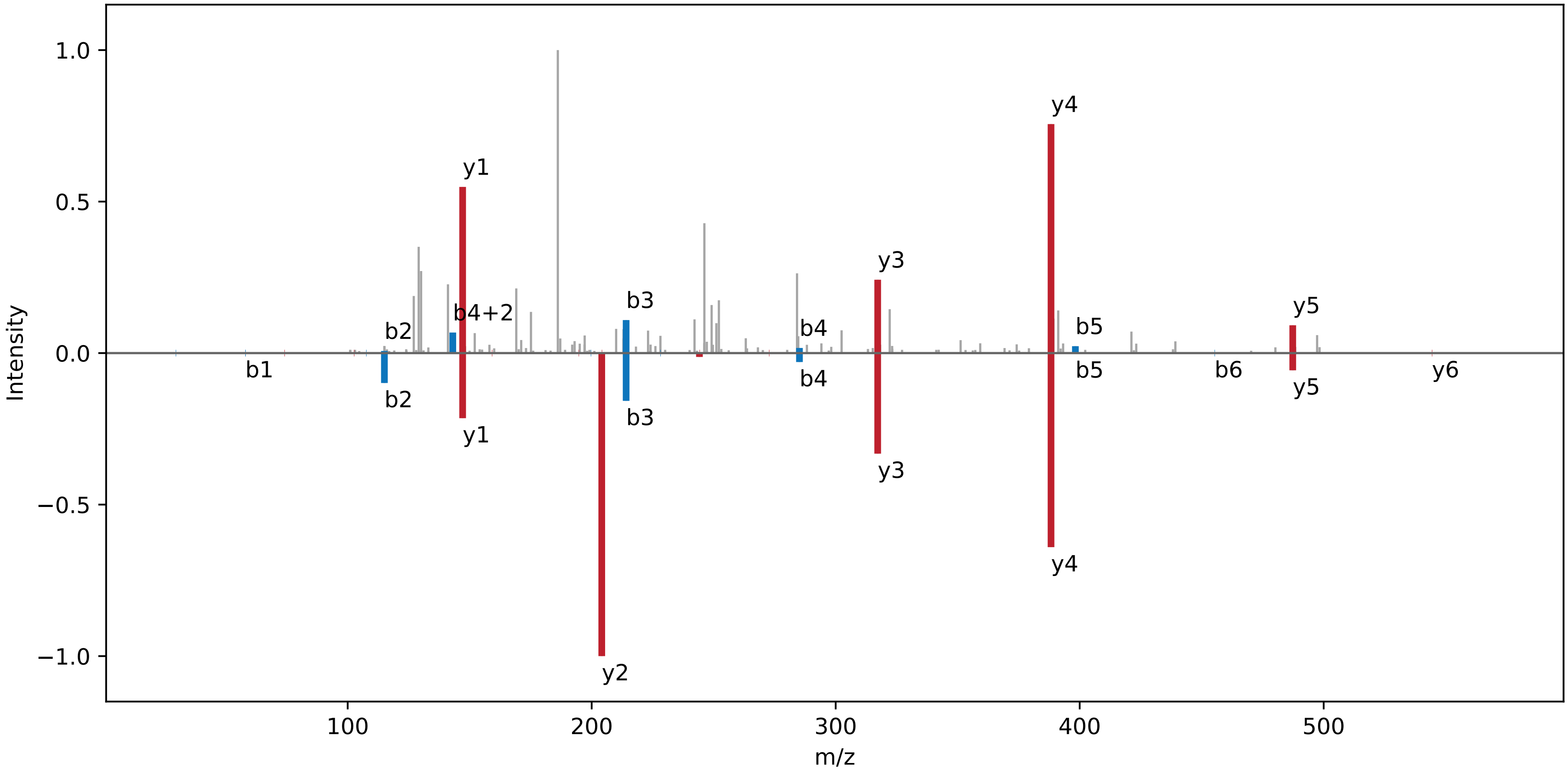
